# Designing of a mini-core that effectively represents 3004 diverse accessions of rice

**DOI:** 10.1101/762070

**Authors:** Angad Kumar, Shivendra Kumar, Manoj Prasad, Jitendra K. Thakur

## Abstract

Genetic diversity provides foundation for plant breeding and genetic research. As part of the 3K Rice Genome Project, over 3000 rice genomes were recently sequenced. We added four Indian rice accessions to it and made original panel of 3004 accessions. However, large set of germplasms are difficult to preserve and evaluate. Construction of core and mini-core collections is an efficient method for managing genetic resources. This study aims to designing of mini-core comprised of 520 accessions representing original panel. The designed mini-core captured most of the SNPs and represented all phenotypes and geographical regions. The mini-core was validated using different statistical analyses and had representation from all major groups including *japonica, indica, aus*/*boro* and aromatic/basmati. GWAS analyses with mini-core panel efficiently reproduced the marker-traits association identified among original 3004 panel. Expression analysis of trait-associated genes highlights the relevance of such mini-core panel. Haplotype analysis could also validate our mini-core panel. Apart from mini-core, we identified few regional and varietal specific marker-trait associations which were not evident in original panel. In this era of large-scale sequencing projects, such a strategy of designing mini-core will be very useful.

**One-sentence summary:** Designing of mini-core as manageable association panel that efficiently mirroring the large and diverse collection of 3004 rice accessions.

## INTRODUCTION

Rice (*Oryza sativa*) is among the primary staple crop fulfilling the nutritional requirement of more than half of the world’s population. An increase/improvement in its global production will have a direct impact on meeting the growing food demand of the world population. More than 90% of the world’s rice production is contributed by Asian countries majorly dominated by China and India (FAOSTAT, 2017). India is the second largest producer of rice (165.3 million tons) after China (208.4 million tons) contributing ∼ 22% of the total rice production of the world. Increase in the rice yield is achieved mainly through improved cropping methods, use of fertilizers and in many areas by intense irrigation. However, outcome of these strategies is now reaching saturation and is becoming limited. So, there is demand of finding alternative ways for yield improvement. Genetic improvement of rice cultivars and varieties can be an effective mean in this regard. The goal can be realized through improved breeding programs using marker-assisted selection and genetics methods which will help in identification of new sources of genetic variations that may help in increasing the productivity (McCouch et al., 2016).

Productivity/yield is a complex trait governed by multiple genes and depends upon the genetic composition and the environmental factors. The variability arises due to segregating alleles at multiple loci where the effect of each allele on the phenotypic trait is relatively small and the overall expression is also influenced by environmental conditions. Single nucleotide polymorphisms (SNPs) present throughout the genome are one of the major causes of allelic variation leading to genetic variability in a population. Quantitative genetic variations lead to a multitude of phenotypes which forms the basis for selection of improved cultivars for breeding and agricultural purpose. Identification of loci governing quantitative traits is critical for maintenance of variation within and among populations. Identification of quantitative trait loci (QTLs) by conventional method of linkage mapping or QTL mapping involves development of mapping population which is a time consuming process and captures limited number of recombination events based on the parental combinations. This methodology forms a part of markers assisted selection and biotechnological approach that has been utilized in large number of crops for identification of genes governing complex traits (Edgerton, 2009; Morrell et al., 2012).

However, with advancement in high-throughput genome sequencing and phenotyping methods, genome wide association studies (GWAS) have been initiated which have been proven to be more effective for crop improvement. It is a more efficient approach for identification of marker-trait association and has been utilized for recognition of genes or loci governing the complex trait (Singh and Singh, 2015; Huang *et al.*, 2012; Morrell *et al.*, 2012; Kump *et al.*, 2011; Famoso *et al.*, 2011; Ingvarsson and Street, 2011; Zhao *et al.*, 2011; Huang *et al.*, 2010; Breseghello and Sorrells, 2006; Gupta *et al.*, 2005). The advantage of GWAS is that it does not require development of mapping population. It explores the naturally available population for mining of available genomic diversity to assess marker-trait association. It also captures large number of historical recombination events prevalent in the natural population. The basic requirement for GWAS is a diverse panel beholding historical recombination events for greater genetic resolution (Morrell et al., 2012). This purpose is best served by a core collection which is designed to capture maximum available/possible diversity (genetic, phenotypic and geographical) of the entire population, with limited number of individuals having low or no kinship among them (Korte et al., 2012). Core collections have been used as an association panel for GWAS in different studies (El Bakkali et al., 2013; Zhang et al., 2014; Perseguini et al., 2015; Ambreen et al., 2018). In case of rice, limited attempts have been made to use a core collection as an association panel. Recently, with the availability of resequencing dataset for 3024 diverse rice accessions, a further deep and robust platform has been provided to elevate the marker-associated breeding efforts concerning various agronomic traits (Li et al., 2014; Alexandrov et al., 2015; Mansueto et al., 2016). Follow up studies has also explored the detailed structural variation and introgression pattern among the 3KRG dataset, further strengthening our understanding for rice diverse genome and traits domestication (Wang et al., 2018) (Fuentes et al., 2019). Even though this panel of 3000 accessions represents core collection of global rice accessions, but still its relatively large size would present difficulties for management and phenotype evaluation (Brown, 2011). So, the need to have a smaller subset mirroring such huge germplasm is warranted for convenient breeding efforts. Here, in this study, we have developed and utilized a mini-core collection (520 accessions) from the original collection of 3004 rice accession dataset, as an association panel for GWAS analysis using 2 million SNPs. While designing mini-core collection, we have considered genotypic (SNPs data), phenotypic data (18 yield-related traits), and representation from various regional genepools (geographical diversity) to accommodate maximum possible diversity. We demonstrate here that such analysis lead to identification of loci which can play an important role for increasing the yield of rice and will be helpful in identification of genes involved in regulation of these traits. The comparatively small size of the association panel designed in the study will be useful and convenient for various phenotype-genotype relationship studies which usually remain a major limitation in different plant breeding programs.

## RESULTS AND DISCUSSION

### Genome sequencing and identification of sequence polymorphism

We re-sequenced four rice accessions in our lab (LGR, PB 1121, Sonasal and Bindli) and identified 3564117 SNPs. We used the default parameters of BWA for alignment of reads sequences from all the four rice genotypes on Nipponbare reference genome. Only those SNPs with average base quality of 30, minimum read depth of 10 and minimum polymorphism call rate of > 90% were scored in order to minimize the detection of false positives. All the SNPs which were consecutive and adjacent to InDels were also eliminated. The read depth of these SNPs varied from 10 to more than 11000, with the overall sequencing depth for four rice accessions ranging from 42 to 48X. We clubbed these SNPs with the 3K Rice Genome (3KRG) project SNP dataset for this study (Alexandrov et al., 2015). Overall 18.9 million single nucleotide polymorphisms were identified among the 3,000 sequenced genomes with an average depth of ∼14X, ranging from ∼4X to 60X. For our analysis, in order to bring the 3KRG dataset to same level of quality we considered the filtered dataset (∼48 lakhs SNPs), corrected with excess of heterozygosity and linkage disequilibrium (LD). Finally, we merged both the datasets and deduced the common SNPs (∼20 lakhs) between the two studies (3KRG and four Indian genotypes). The distribution of common SNPs over different rice chromosomes showed non-uniform distribution. Maximum number of SNPs were located on the chromosome 1, 11 and 2 while least number of SNPs were found to be present on chromosome 9.

### Development of mini-core collection

In order to make a representative mini-core group, we utilized phenotypic data of 2266 rice accessions available for 18 yield-related traits and genotypic data (SNPs) of 3000 accessions of 3KRG dataset. The purpose of developing independent mini-cores using phenotypic data and genotypic data was to avoid trade off and to capture maximum possible phenotypic and genotypic variability prevalent in the original collection of rice accessions. Scanning of 2266 accessions resulted into a mini-core collection (CC1) of 227 accessions representing 10% of the initial collection. We added the four Indian accessions (LGR, PB 1121, Sonasal and Bindli) that were resequenced in our lab. Thus, the mini-core collection CC1 consisted of 231 accessions representing diversity in phenotypic traits. The 3000 accessions with their SNP data were analysed separately for development of another mini-core collection (CC2) consisting of 300 accessions representing 10% of the original collection and also included the four Indian accessions (LGR, PB 1121, Sonasal and Bindli) sequenced in our lab.

The mini-core collections, CC1 and CC2 were assessed for their coverage of phenotypic variations with reference to the original panel (Supplemental Table S1). It was found that none of these two mini-cores could capture the entire range for all the phenotypic traits prevalent in the original collection. The traits which could not be captured in the mini-cores included days to 80% heading, 100 grain weight, days to first flowering, grain width, panicle length and seedling height. These mini-core collections (CC1 and CC2) were further assessed for various evaluation criteria such as Shannon’s diversity index (*I*), Nei’s gene diversity (*H*), mean difference percentage (MD%), variance difference percentage (VD%), variable rate of coefficient of variance (VR%) and coincidence rate of range (CR%) to assess their efficiency to capture the maximum diversity prevalent in the original collection. The MD% of the developed mini-core collections was in the range of 2.8-4.08% which was well below the prescribed value of 20% (Supplemental Table S2). VD% representing the variance captured in the mini-core collections ranged from 19.78 to 39.77%. The VR% varied from 86 for CC1 to 107.68% for CC2. CC1 had the highest CR% value of 92% whereas CR% of CC2 was 91.1. The value of Shannon-Weaver index (*H*) ranged from 1.98 for CC1 to 2.25 for CC2. The value of Nei’s genetic diversity (*I*) was found to be higher for CC2 (0.79) in comparison to CC1 (0.77) (Supplemental Table S2).

The developed mini-core collections were also assessed for representation of all the varietal groups and regional genepools present in the original panel (Supplemental Table S3 & S4). The mini-core CC1 had highest representation of *indica* (129) followed by Temperate *japonica* (38), Intermediate (19), Tropical *japonica* (15), *japonica* (14), *aus/boro* (11) and Aromatic (5). The mini-core CC2 had highest representation of *indica* (171) followed by Intermediate (45), *aus/boro* (42), Aromatic and *japonica* (12 each), Tropical *japonica* (10) and Temperate *japonica* (8) (Supplemental Table S3). We also assessed the comparative distribution of accessions from different varietal groups into the mini-cores and found that CC2 had higher proportion of accessions from *aus/boro* (19.5% of the original representation), Intermediate (33.3% of the original representation) and Aromatic (16.9% of the original representation) groups. On the other hand, CC1 had only 5.1% representation of *aus/boro*, 14% of Intermediate and 7% of Aromatic group from the original collection (Supplemental Table S3). Accessions from Temperate *japonica* group were highly represented in CC1 (11.9% of the original representation) while only small proportion of Temperate *japonica* group (2.5% of original representation) were picked in CC2 (Supplemental Table S3). Comparable portions of accessions from *indica* (7.4% in CC1 and 9.8% in CC2), Tropical *japonica* (3.8% in CC1 and 2.5% in CC2) and *japonica* (10.6% in CC1 and 9% in CC2) groups were represented in both CC1 and CC2 mini-cores. Thus, none of the two mini-cores developed here consisted of 10% of representations from all the varietal groups (Supplemental Table S3). The developed mini-core collections were also assessed for distribution of accessions from different regional genepools. The mini-core CC1 consisted of 55 accessions from South Asia (6.9% of the original collection) followed by 52 accessions of South East Asia (5.1% of the original collection), 52 accessions of China (10.8% of original collection), 18 accessions of Europe (15.2% of original collection), 17 accessions of America (10.2% of original collection), 15 accessions each from East Asia and Africa (11.4% and 5.9% of the original collection, respectively), 4 accessions of Oceania (23.5% of original collection) and 3 accessions of Unknown origin (8.8% of original collection). CC2 comprised of 122 accessions from South Asia (15.5% of the original collection) followed by 70 accessions of South East Asia (6.9% of the original collection), 55 accessions of China (11.4% of the original collection), 23 accessions of Africa (9.1% of the original collection), 13 accessions of America (7.8% of the original collection), 8 accessions of East Asia (6% of the original collection), 6 accessions of Unknown origin (17.4% of the original collection), 2 accessions of Europe (1% of the original collection) and 1 accession from Oceania (5.9% of the original collection; Supplemental Table S4).

Only 28 accessions were found to be common between CC1 and CC2 showing that different accessions were selected on the basis of phenotypic and genotypic variation and justifying our concern of designing the mini-cores independently using just phenotypic or genotypic data. An ideal mini-core should represent maximum possible diversity present in the original collection. However, different evaluation criteria such as phenotypic range (Supplemental Table S1), MD%, VD%, VR%, CR%, Shannon and Nei’s diversity indices (Supplemental Table S2), varietal (Supplemental Table S3) and geographical coverage (Supplemental Table S4) revealed that none of the mini-cores (CC1 and CC2) were able to capture the complete diversity ranges of original collection to be considered as an ideal representative subset. Therefore, we merged CC1 and CC2 to develop mini-core collection (CC3) comprising of 520 non-redundant accessions (503 accessions by merging CC1 and CC2 + 17 accessions capturing the extreme values of phenotypic traits discussed below) in order to capture maximum possible allele/trait diversity and to prevent any trade-off between the two data sets (phenotypic and genotypic) when used in conjunction (Fig. 1). CC3 comprising of 520 accessions represented 17.3% of the initial collection (3004 accessions) fulfilling the initial requirement of size of an ideal core collection which should range between 5-20% of the original collection (Brown and Spilllane, 1999). CC3 was assessed for its representation of the original collection and various traits under consideration by different evaluation criteria (Supplemental Table S1-4). CC3 covered the entire range of traits from original collection including the traits which were not completely covered by CC1 and CC2 such as days to 80% heading, 100 grain weight, days to first flowering, grain width, panicle length and seedling height (Supplemental Table S1). CC3 was evaluated by different evaluation criteria. The MD% of CC3 was 2.9% within the range of 2.8 - 4.08% observed for CC1 and CC2 (Supplemental Table S2). The value of VD% representing the variance captured by CC3 accessions was 18.9% which is lower than the value of CC1 and CC2. The value of VR% captured in the mini-core CC3 was 109.3% (highest value among the three mini-cores). Also, CC3 was found to have the highest value of CR% (96.2) among all the three mini-cores. The value of Shannon-Weaver index (*H*) and Nei’s genetic diversity (*I*) for CC3 was found to be 2.17 and 0.79, respectively (Supplemental Table S2).

**Figure 1:**
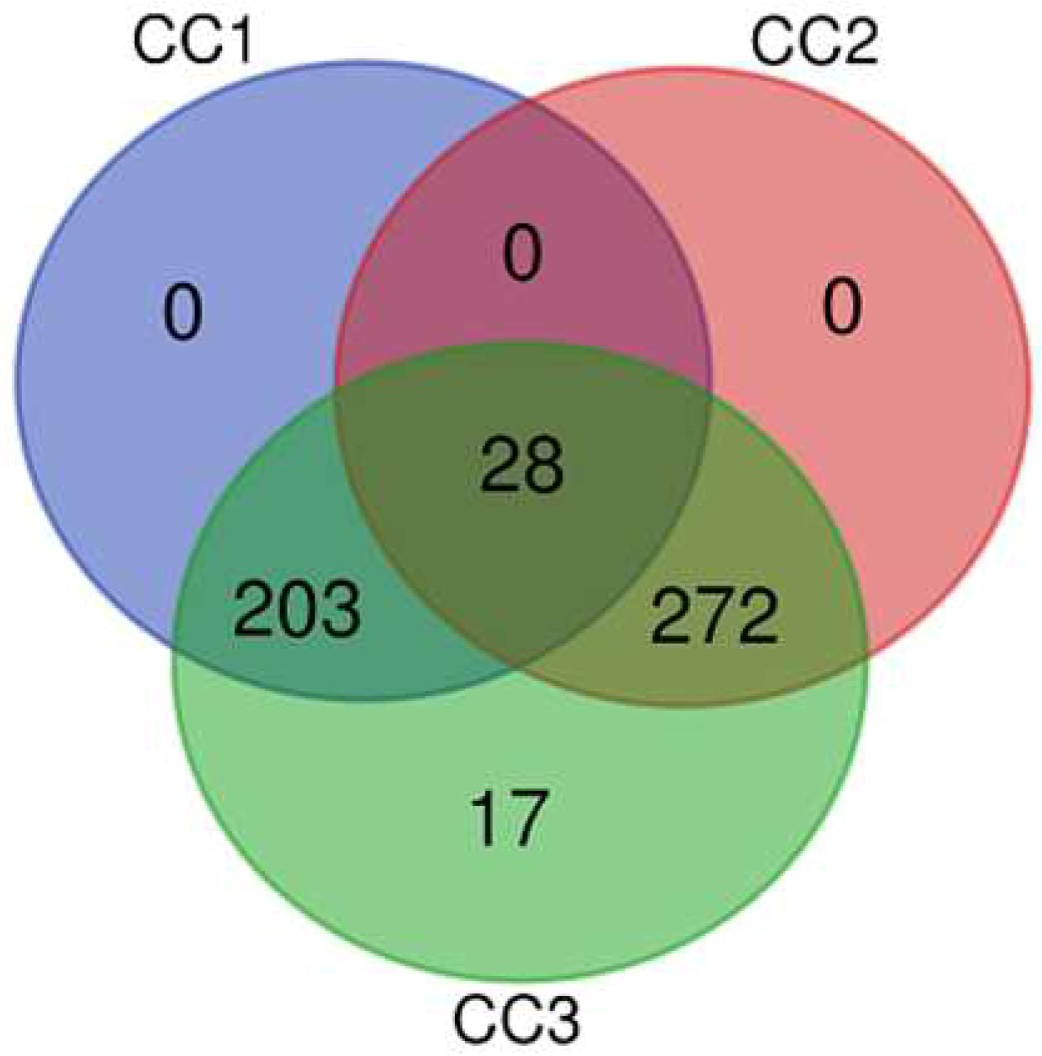
Venn diagram showing distribution of accessions in different mini-core collections developed in the present analysis. CC 1 represents mini-core designed using phenotypic data, CC2 represents mini-core designed using SNP data, CC3 represents merged mini-core collection.

The mini-core was also assessed for representation of all the varieties and regional gene pools of rice present in the initial collection (Supplemental Table S3 & S4). In the mini-core CC3, all the varieties had at-least 10% representation from original collection except Tropical *japonica* varietal group which had only 6.9% representation of original collection (27 accessions; Supplemental Table S3). Number of accessions and percentage representation of all the varietal groups of the original collection in the mini-core CC3 has been provided in Supplemental Table S3. Representation of accessions from different regional genepools of original collection varied from 12 to 29.4% in the mini-core CC3 (Supplemental Table S4). CC3 had highest contribution from South Asia (176 accessions; 22.4% of its original representation12% of its original representation), China (101 accessions; 20.95% of its original representation), Africa (38 accessions; 15% of its original representation), America (28 accessions; 16.9% of its original representation), East Asia (21 accessions; 15.9% of its original representation), Europe (19 accessions; 16.1% of its original representation), Unknown origin (9 accessions; 25.5% of its original representation) and Oceania (5 accessions; 29.4% of its original representation; Supplemental Table S4). Thus, mini-core collection CC3 with 520 accessions was found to fulfil all the criteria of capturing maximum possible diversity of the original panel (3004 accessions) in comparison to CC1 and CC2 developed using phenotypic and molecular data, respectively, and was considered further for its utility as an association panel.

### Correlation analysis between quantitative traits

Correlation analysis was performed between different quantitative traits under consideration. This exercise was done to identify the traits, if any, which were governed by same genes or which were developmentally or structurally related to each other. A significant positive correlation was observed between days to 80% heading (DEH) and days to first flower (DFF; 0.998). The correlation values for five trait combinations were found to be higher than 0.5; culm length and days to first flower (0.68), culm diameter and days to first flower (0.614), culm length and panicle length (0.557), culm length and days to 80% heading (0.555) and grain width and 100 grain weight (0.542). The correlation values for all the other trait combinations were found to be lower than 0.5. Though some of the traits studied here found to be negatively correlated but the correlation was not significant. The correlation coefficient of all the traits analysed is provided in Supplemental Table S5. The magnitude and direction of correlation between different traits helps in the selection decisions in breeding and crop improvement programs.

### Distance-based cluster analysis and principal component analysis (PCA)

Distance-based cluster analysis was performed to assess the grouping of accessions of original collection of rice genotypes. Analysis of the SNP data (2081521 SNPs) using Maximum-likelihood method grouped 3004 accessions into two major clusters (named as CL I and II) with internal subgroupings (Fig. 2A). The cluster CL I was found to contain maximum number of accessions (66%) of original collection while CL II showed presence of around 32% of accessions (Supplemental Table S6). Around 1.4% of the accessions did not belong to any of the groups (Fig. 2A). These un-clustered accessions (43) were mainly Intermediate (21) and *indica* (15) group genotypes. In addition, some of the *japonica* (3), tropical *japonica* (2), temperate *japonica* (1) and aromatic (1) genotypes also remained un-clustered. The cluster CL I comprise of 1987 accessions was further grouped into two sub-clusters CL Ia and CL Ib. The larger sub-cluster CL Ia consisting of 1771 accessions was majorly dominated by *indica* (1641) varieties whereas cluster CL Ib comprised of a total of 216 accessions with major contribution from *aus/boro* (172) followed by *indica* (25) varieties. CL II with 974 accessions was divided into three sub-clusters named as CL IIa, IIb and IIc. CL IIa being the largest sub-cluster of CCL II comprised of 519 accessions majorly dominated by Tropical *japonica* (329) and *japonica* (80). The sub-cluster CL IIb consisting of 358 accessions was dominated by Temperate *japonica* (250) genotypes. The Indian genotype included in the study (LGR) was part of sub-cluster IIb. CL IIc was the smallest sub-cluster of CL II comprising of 97 accessions with major representation from Aromatic (50) and Intermediate (23) varieties. Bindli, Sonasal and PB 1121 grouped together in sub-cluster CL IIc (Fig. 2A; Supplemental Table S6).

**Figure 2:**
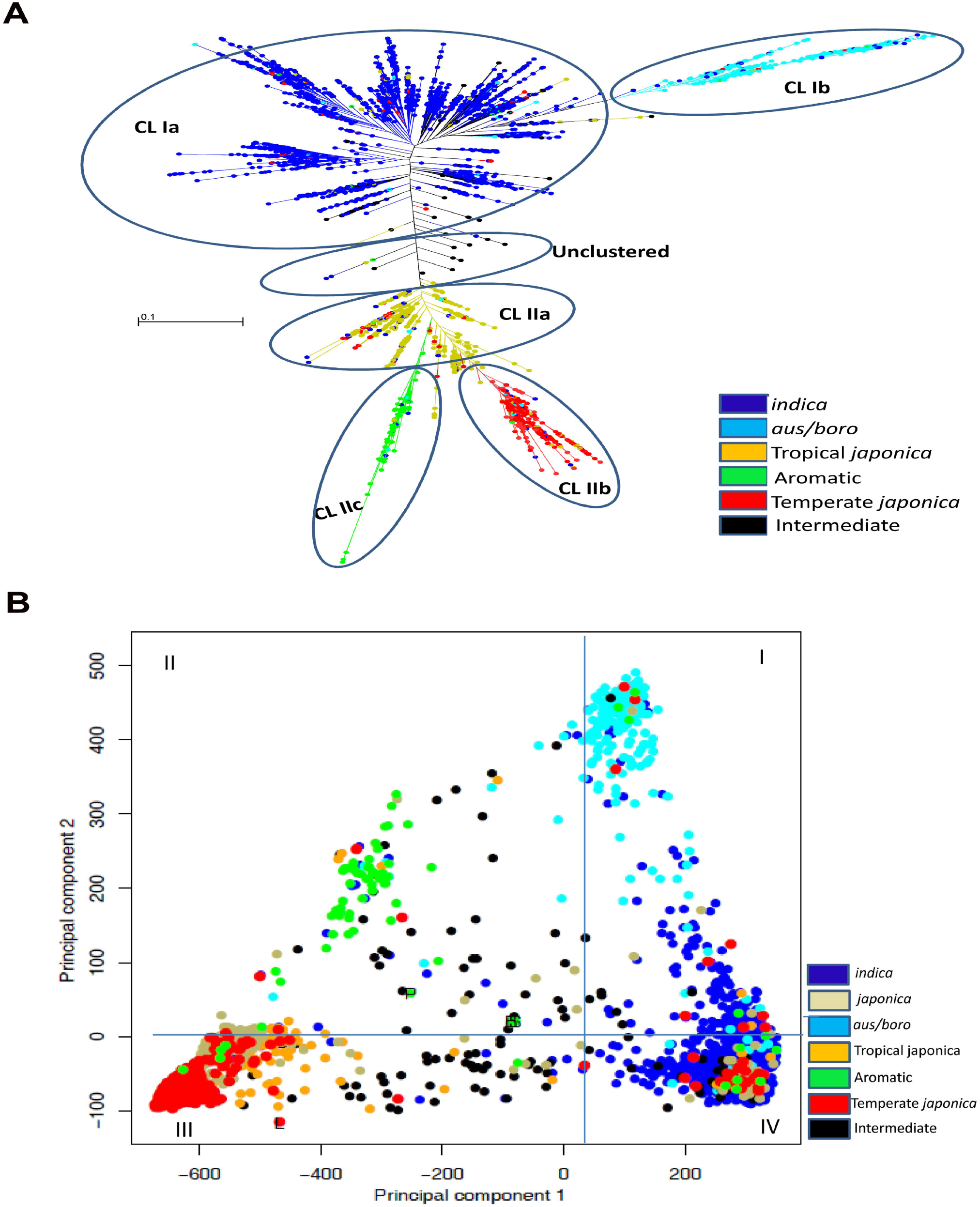
Phylogenetic representation among 3004 rice accessions based on polymorphic SNP markers. **(A)** Maximum-likelihood dendrogram illustrating the genetic relationship among. Two clusters designated as Cl **1-11** with further sub-clustering are shown. **(B)** Principal Component Analysis of 3004 accessions of original collection for principal component axes I and 2. Distribution of accessions in different quadrants (I-N) is shown. Varietal group color codes are provided in the figure.

In the Principal component analysis (PCA), 3004 accessions were evenly distributed on coordinate axes 1 and 2 covering 45.6% and 26% of total variance, respectively (Fig. 2B). Accessions from *indica* variety clustered together in PCA analysis showing congruence with distance-based analysis. Accessions from *indica* which forms the largest group of original collection and were majorly present in Cluster Ia of distance-based analysis were predominantly present in quadrant I and IV. Accessions from *japonica*, temperate *japonica* and tropical *japonica* which were part of Cluster II of distance-based analysis were found to be present in quadrant III and IV. Accessions from *aus/boro* were found to be limited in quadrant I while Aromatic accessions were found to be present in quadrant II. Accessions of Intermediate type were found to be spread in all the quadrants of PCA analysis and showed congruence with maximum likelihood dendrogram in which they were present among all the clusters in even proportion (Fig. 2B). Next, we checked the distribution of CC3 accessions on distance-based maximum likelihood dendrogram and PCA analysis of Initial collection of rice to assess their presence in all the clusters and quadrants. CC3 showed balanced representations from all the clusters in the range of 10.5%-25.7% of accessions of maximum likelihood dendrogram (Fig. 3A; Supplemental Table S7). A total of 18.1% of accessions from sub-cluster CL Ia, 19.4% of accessions from sub-cluster CL Ib, 25% of un-clustered accessions, 10.5% of accessions from sub-cluster IIa, 18% of accessions from sub-cluster IIb and 25.7% accessions from sub-cluster IIc were captured in CC3 from original collection (Fig. 3A; Supplemental Table S7). Similarly, accessions from all the quadrants of PCA analysis were found to be part of CC3 (Fig. 3B). Thus, CC3 had representation of accessions from all the clusters of maximum likelihood dendrogram and quadrants of PCA, capturing maximum possible genotypic diversity.

**Figure 3:**
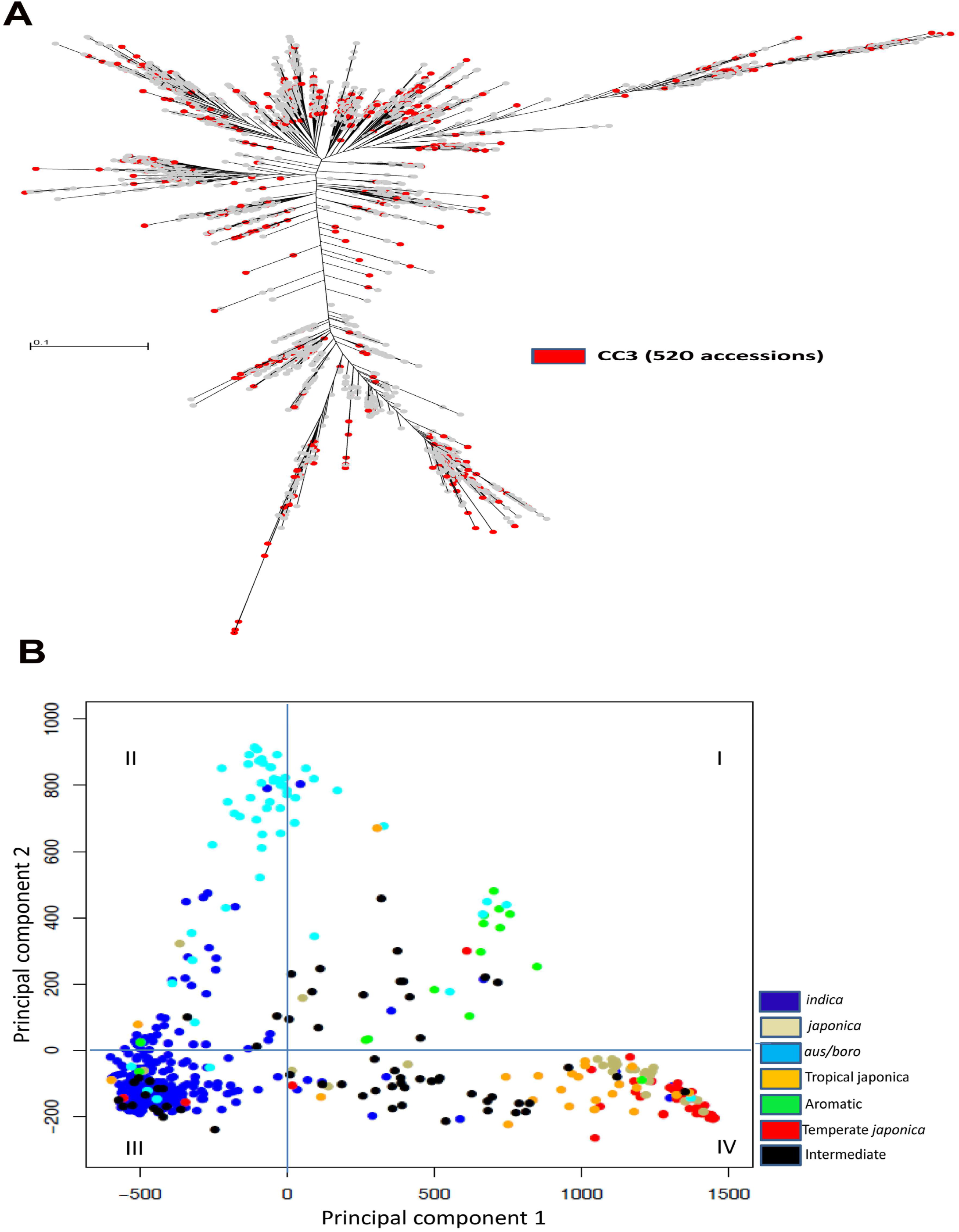
Phylogenetic representation showing distribution of CC3 accessions. **(A)** Distribution of CC3 accessions (520) in different clusters of maximum-likelihood dendrogram of original collection of rice (3004 accessions). CC3 accessions are shown with red dots. **(B)** Principal Component Analysis of 520 accessions of mini-core collection CC3 for principal component axes 1 and 2. Distribution of accessions in different quadrants (I-IV) is shown. Varietal group color codes are provided in the figure.

### Population structure analysis of original collection using FastSTRUCTURE

Population structure analysis for initial collection of rice (3004 accessions) was performed using the FastSTRUCTURE program. The best clustering was at K=7 and the clusters obtained were named as FSTR CL 1–7 (Fig. 4A). FSTR CL 1 consisted of 219 accessions majorly dominated by *aus/boro*(179) followed by *indica* (28; Supplemental Table S8) and showed congruence with CL Ib of maximum likelihood analysis. FSTR CL 2 comprised of 522 accessions having highest representation from tropical *japonica* (310) and *japonica* group (94) and was found to have similar accessions as in CL IIa of maximum likelihood analysis. Some of the *indica* (47 accessions), temperate *japonica* (35) and Intermediate group (27) accessions were also found in FSTR CL 2. The smallest cluster was FSTR CL 3 with 90 accessions and was dominated by Aromatic accessions (50) followed by Intermediate group accessions (19) and was showing congruence with CL IIc of maximum likelihood analysis. The largest cluster of population structure analysis was FSTR CL 4 comprising of 973 accessions with major representation from *indica* group (885) and minor contributions from Tropical *japonica* (26), Intermediate (21), Temperate *japonica* (17) and *aus/boro* group (14). FSTR CL 5 comprised of 372 accessions majorly representing Temperate *japonica* (248) and minor representations from Tropical *japonica* (35), Intermediate (29), *indica* (26) and *japonica* group (25) and was similar to CL IIb of maximum likelihood analysis. The FSTR CL 6 consisted of 323 accessions dominated by *indica* variety (297) while FSTR CL 7 comprised of a total of 505 accessions having contributions from *indica* (451) and Intermediate varieties (27). The FSTR CL 4, 6 and 7 corresponded with the CL Ia of maximum likelihood dendrogram, suggesting that the accessions of CLIa can be further divided into three subgroups. Number of accessions constituting different clusters as determined by FastSTRUCTURE analysis is provided in Supplemental Table S8. Next, we looked for admixed genotypes in all the groups and found that 41% (1242) of accessions were admixed in nature (Supplemental Table S9). Among all the clusters, FSTR CL 7 consisting of 505 accessions had more admixed individuals (351) than pure individuals (154 accessions) followed by FSTR CL 6 (145 admixed and 148 pure accessions; Supplemental Table S9). Assessment of admixtures within varietal groups revealed that Intermediate population had more admixtures (94) than pure (41) accessions while *indica* population had 821 admixed individuals (47%) out of 1743 accessions (Supplemental Table S9). Analysis of regional genepools revealed that only European region had more admixed (65) than pure individuals (53) (Supplemental Table S9). Distribution of CC3 accessions (520) in different clusters of FastSTRUCTURE analysis (FSTR CL 1 - 7) was assessed to check representation of individuals from all the clusters in the developed mini-core collection. CC3 captured 50 accessions (40 pure individuals with Q value > 80%) from a total of 219 accessions present in FSTR CL 1 (Supplemental Table S10). Forty two accessions (23 pure individual) were picked from 522 accessions of FSTR CL 2 in CC3 while 24 accessions (13 pure individuals) were part of CC3 from FSTR CL 3. CC3 captured 185 accessions (109 pure individuals) from the largest FastSTRUCTURE cluster FSTR CL 4 comprising of 973 accessions. Out of 372 accessions of FSTR CL 5, 74 accessions were present in CC3 (37 pure individuals). Out of the 323 accessions of FSTR CL 6, 61 accessions were part of CC3 (28 pure individuals) while 84 accessions (25 pure individuals) were present in CC3 representing FSTR CL 7 with 505 accessions (Supplemental Table S10). Thus, CC3 had representations of both pure and admixed accessions from all the 7 clusters of FastSTRUCTURE analysis derived for initial collection of rice. Therefore, we were able to verify the initial objective of development of a mini-core collection (CC3) representing maximum phenotypic, genotypic, varietal and geographical variability present in the original collection of rice consisting of 3004 rice accessions.

**Figure 4:**
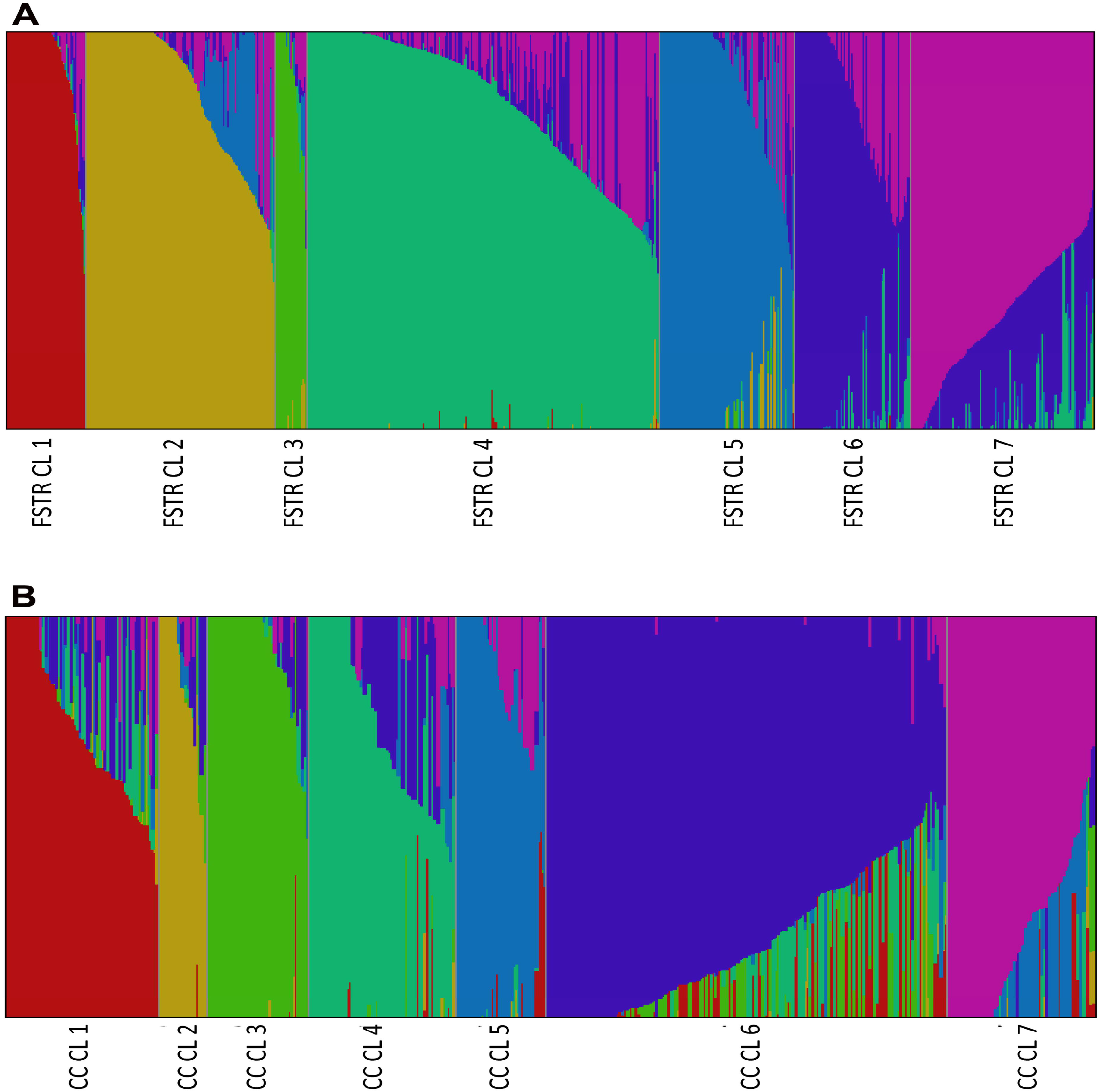
Population structure of original (3004) and mini-core (520) rice accessions at K=7, derived using the model based Bayesian algorithm. **(A)** Population structure of original (3004), each sub-population is represented by a different color code named as FSTR CLl-FSTR CL7.(B) Population structure of 520 rice accessions, each sub-population is represented by a different color code named as CC CLl - CC CL7. Each vertical bar represents a single rice accession. Admixed individuals have bars composed of multiple colors.

### Assessment of mini-core collection CC3 to assess its utility as an association panel

An association panel should have low population structure and low kinship among its members to avoid spurious marker-trait associations (Yu and Buckler, 2006; Zhu et al., 2008; Yang et al., 2010; Nachimuthu et al., 2015). Therefore, we performed population structure and kinship analysis among the accessions of the developed mini-core collection (CC3) to use it as an association panel.

### Population structure analysis of mini-core CC3

Estimation of the underlying population structure among 520 accessions of mini-core collection CC3 was done using FastSTRUCTURE program which delineated them into seven clusters (K = 7) named as CC CL1–CC CL7 (Fig. 4B; Supplemental Table S11). The cluster 1 (CC CL 1) comprised of 73 accessions and was dominated by *indica* (57) and Intermediate group accessions (12). CC CL2 was the smallest of all the clusters comprising of 23 accessions dominated by Aromatic (9) and Intermediate group (8) genotypes. The cluster CC CL3 consisting of 49 accessions was dominated by *aus/boro* (41). CC CL4 comprised of 70 accessions and had representations from *indica* (58 accessions) and Intermediate (8) populations. The cluster CC CL5 consisted of 43 accessions belonging to different varietal groups such as Tropical *japonica* (17), *japonica* (10) and Intermediate group (8). CC CL6 was the largest cluster consisting of 191 accessions majorly belonging to *indica* (165) and Intermediate (11) populations. CC CL7 comprised of 77 accessions majorly representing Temperate *japonica* (36), Intermediate (14) and *japonica* (9) population. The detailed distribution of accessions belonging to different varietal groups in 7 clusters of mini-core collection CC3 has been provided in Supplemental Table S11.

We found that 47% (245) of the accessions of mini-core collection CC3 were admixtures (Supplemental Table S12). CC CL1 comprised of a total of 73 accessions and had more admixed individuals (48) than pure individuals (25) followed by CC CL4 consisting of 70 accessions (42 admixed and 28 pure individuals (Supplemental Table S12). The clusters CC CL2, 5 and 7 had approximately equal number of pure and admixed accessions while CC CL3 and CC CL6 had higher number of pure individuals in comparison to admixed accessions. Admixture assessment of varietal group of CC3 showed that Intermediate group had more admixtures (47) than pure (14) individuals followed by *japonica* with 12 admixed and 11 pure accessions. The *indica* group had 144 admixed genotypes (49%) out of 295 accessions while Tropical *japonica* had 13 admixed accessions out of a total of 27 individuals. Analysis of regional gene pools of rice accessions revealed that South East Asia (63 admixed and 60 pure accessions), China (54 admixed and 47 pure accession), America (23 admixed and 15 pure accessions), Europe (13 admixed and 6 pure accessions) and Oceania (3 admixed and 2 pure accessions) pools had more admixed individuals than pure individuals present in CC3 (Supplemental Table S12). Detailed distribution of admixed and pure accessions from these different groups present in CC3 has been provided in Supplemental Table S12. Increased number of admixed individuals among populations derived using FastSTRUCTURE (CC CL1–CC CL7), varietal and geographical distribution/regional gene pool level confirms that CC3 comprised of more unrelated and admixed individuals in comparison to initial collection of rice and validates its suitability as an association panel.

### Kinship analysis of mini-core CC3 individuals

Kinship analysis between individuals of mini-core collection CC3 was performed to estimate co-ancestry among the accessions. Seventy percent of pairs of the CC3 accessions had a kinship value of less than zero while 25.4% accessions had kinship values ranging between 0-0.25 percent (Fig.5). Around 4.5% of CC3 accessions showed kinship value in the range of 0.25-0.50%, while only 0.1% of accessions had kinship value in the range of 0.5-0.75% (Fig. 5). Thus, the kinship value for most of the CC3 accessions exhibited absence or weak level of genetic relatedness among the individuals fulfilling the primary requirement of utilization of mini-core collection CC3 as an association panel.

**Figure 5:**
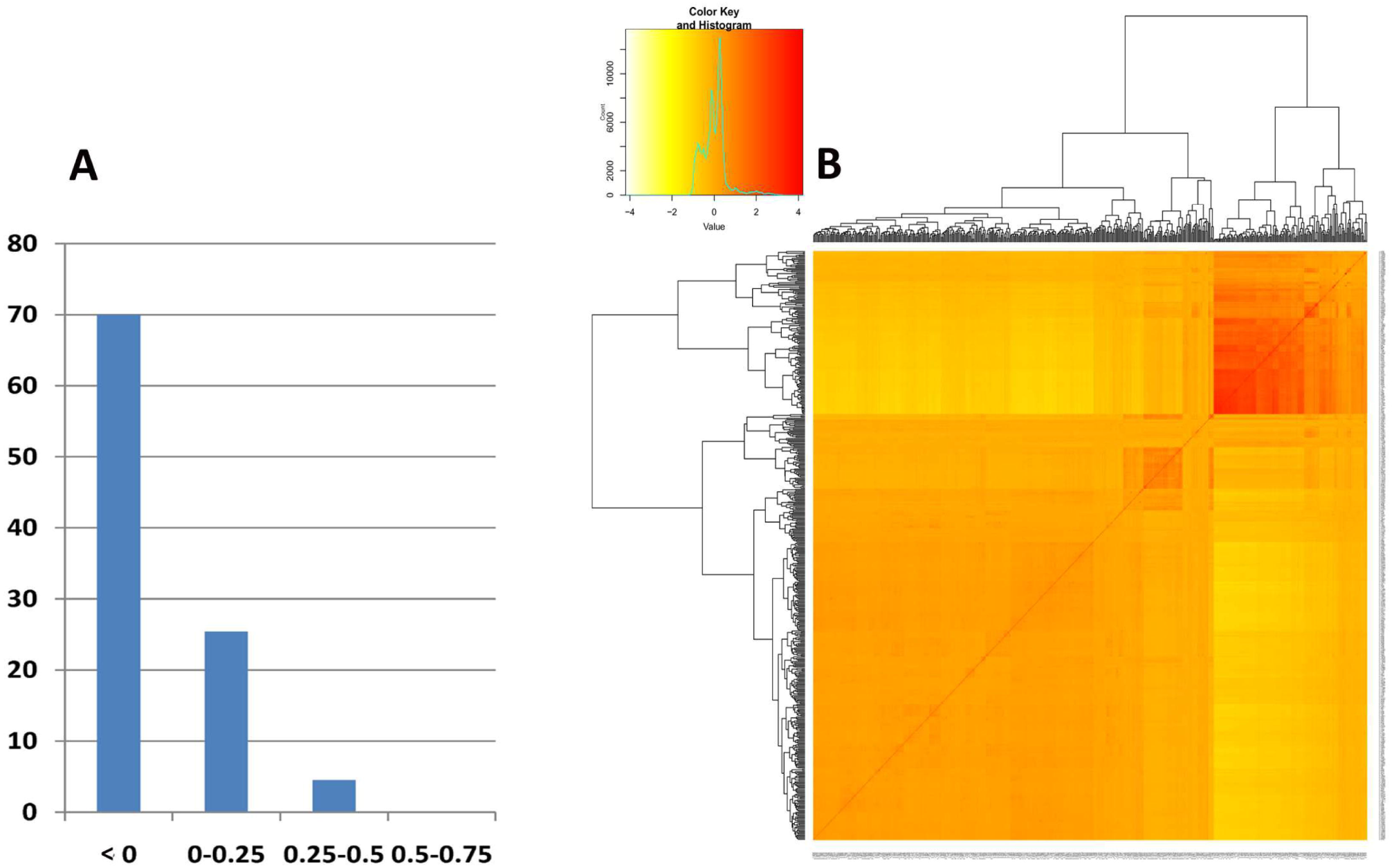
Kinship analysis between 520 accessions of mini-core CC3. **(A)** Histogram showing kinship status among mini-core accessions. **(B)** Kinship matrix for mini-core showing relatedness among accessions.

### GWAS with mini-core collection CC3

As the mini-core collection (CC3) showed low population structure and low kinship value, we proceeded to study its utilization for Genome Wide Association Studies (GWAS) in rice. GWAS analysis was performed on mini-core collection CC3 accessions (520) using 2081521 SNPs with MAF >0.02 and 18 yield-related traits of agronomic importance using the program GAPIT based on compressed mixed linear model (MLM). Association between markers and traits was considered to be significant at *P* < 1 × 10^−8^, FDR adjusted *P* value of < 0.05 and correlation value (R^2^) of >= 10%. Six out of the 18 traits analysed namely Endosperm Type (ET), Grain Length (GL), Grain Width (GW), Panicle axis (PA), Secondary branching (SB) and Seed coat color (SCC) showed significant marker-trait associations (Table 1). A total of 5924 SNPs were found to be significantly associated with the above mentioned six traits covering phenotypic variation from 10.4 to 61.6%. Three SNPs were found to be significantly associated with the trait Grain Length (GL) in our analysis which was present on chromosome 3. The most significant SNP (G/T) associated with the trait GL on chromosome 3 was at the position 16733441 with FDR adjusted *P* value of 1.4 × 10^−3^ and explaining phenotypic variation of 32.2% (Fig. 6A). This was previously reported as known loci for *GS3* (Fan et al., 2006). Another important grain trait, Grain Width (GW) showed significant association with 64 SNPs specifically present on chromosome 5. The most significant SNP (C/G) associated with the trait GW on chromosome 5 at position 5371949 had FDR adjusted *P* value of 2.8 × 10^−4^ and explained phenotypic variation of 34.2% (Fig. 6B). This SNP was found to be associated with previously characterized locus *qSW5* (Shomura et al., 2008). The trait Endosperm Type (ET) showed maximum number of significant associations with 3651 SNPs distributed on chromosomes 2, 4, 6, 8, 11 and 12. The most significant SNP (G/T) was found to be present on chromosome 6 at position 6294468 with FDR adjusted *P* value of 1.2 × 10^−8^ and explaining 29% of phenotypic variation (Fig. 6C). The other significant SNP (T/G) associated with ET on chromosome 6 was found at position 1765761 with FDR adjusted *P* value of 6.4 × 10^−8^ and detecting a phenotypic variation of 25%. This was also previously reported as known loci f*or Waxy* gene by various authors (GAO, 2003; Tian et al., 2009; Huang et al.,2010). The other significantly associated SNP were found to be present on chromosome number 2 at position 10277497 (C/T) with FDR adjusted *P* value of 1.7 × 10^−7^ and explaining phenotypic variation of 24% and on chromosome 8 at position 6048049 (G/A) with FDR adjusted *P* value of 4.3 × 10^−7^ and explaining phenotypic variation of 23.2% (Table 1). The trait Seed coat color (SCC) was found to be associated with 306 SNPs present on chromosome 7 in our analysis. The most significant SNP (T/C) associated with the trait SCC was present on chromosome 7 at position 6124457 with FDR adjusted *P* value of 4.5 × 10^−8^ and explaining a phenotypic variation of 61.6% (Fig. 6D). This SNP was found to be associated with *Rc* gene described in previous report as important locus for SCC (Sweeney et al., 2006). Another SNP significantly associated with the trait SCC (T/G) was present at position 6660825 on chromosome 7 with FDR adjusted *P* value of 1.6 × 10^−6^ and explaining phenotypic variation of 59.7%. The trait Secondary branching (SB) was found to be associated with 1779 SNPs distributed on chromosomes 2, 4, 6, 7, 9 and 11. The most significant SNP (C/T) associated with the trait SB was present on chromosome 2 at position 5032535 with FDR adjusted *P* value of 6.4 × 10^−7^ and explaining a phenotypic variation of 32% (Fig. 6E). Significantly associated SNP with the trait SB were also present on chromosome 4 at position 2521459 (A/G) and 12427420 (G/A) with FDR adjusted *P* value of 1.6 × 10^−4^ each and explaining phenotypic variation of 23.5 and 23.4%, respectively. The trait Panicle axis (PA) was significantly associated with 121 SNPs distributed on Chromosomes 2, 4, 6 and 10. The most significant SNP associated (A/C) with the trait PA was present on chromosome 4 at position 1075655 with FDR adjusted *P* value of 3.7 × 10^−4^ and explaining phenotypic variation of 24% (Fig. 6F). The SNPs that were significantly associated with the trait PA were also present on chromosomes 6 and 10 at positions 28676456 (G/A) and 14829875 (C/A) with FDR adjusted *P* value of 3.7 × 10^−4^ each and explaining phenotypic variation of 23.1 and 23%, respectively. Thus, identification of previously characterized QTLs for respective traits authenticates the utility and importance of mini-core CC3. The detailed distribution of all significantly associated 5924 SNPs, identified in the current analysis associated with the traits Endosperm Type (ET), Secondary branches (SB), Seed coat color (SCC), Panicle axis (PA), Grain Width (GW) and Grain Length (GL) and their chromosomal position has been provided in Supplemental Table S13.

**Table 1:** Association analysis of 520 accessions of mini-core collection CC3.

| Trait | Chr | Position | Major allele | Minor allele | Minor allele frequency | Nipp. allele | CC3 allele | P-value FDR adjusted | R<sup>2</sup> value (%) | Known loci |
| --- | --- | --- | --- | --- | --- | --- | --- | --- | --- | --- |
| Grain length | 3 | 16733441 | G | T | 0.36 | G | T | 1.4 X 10<sup>-3</sup> | 32.3 | *GS3* |
| Grain width | 5 | 5371949 | C | G | 0.46 | C | G | 2.8 X 10<sup>-4</sup> | 34.2 | *qSW5* |
| Endosperm type | 6 | 6294468 | G | T | 0.07 | G | T | 1.2 X 10<sup>-8</sup> | 29 |  |
| Endosperm type | 6 | 1765761 | T | G | 0.13 | T | G | 6.4 X 10<sup>-8</sup> | 25 | *Waxy* |
| Endosperm type | 2 | 10277497 | C | T | 0.026 | C | T | 1.7 X 10<sup>-7</sup> | 24 |  |
| Endosperm type | 8 | 6048049 | G | A | 0.035 | G | A | 4.3 X 10<sup>-7</sup> | 23.2 |  |
| Seed coat color | 7 | 6124457 | T | C | 0.456 | T | C | 4.5 X 10<sup>-8</sup> | 61.6 | *Rc* |
| Seed coat color | 7 | 6660825 | T | G | 0.454 | T | G | 1.6 X 10<sup>-6</sup> | 59.7 |  |
| Secondary branching | 2 | 5032535 | C | T | 0.013 | C | T | 6.4 X 10<sup>-7</sup> | 32 |  |
| Secondary branching | 4 | 2521459 | A | G | 0.052 | A | G | 1.6 X 10<sup>-4</sup> | 23.5 |  |
| Secondary branching | 4 | 12427420 | G | A | 0.208 | G | A | 1.6 X 10<sup>-4</sup> | 23.4 |  |
| Panicle axis | 4 | 1075655 | A | C | 0.013 | A | C | 3.7 X 10<sup>-4</sup> | 24 |  |
| Panicle axis | 6 | 28676456 | G | A | 0.0078 | G | A | 3.7 X 10<sup>-4</sup> | 23.1 |  |
| Panicle axis | 10 | 14829875 | C | A | 0.0078 | C | A | 3.7 X 10<sup>-4</sup> | 23 |  |

Nipp; Nipponbare, Chr; Chromosome

**Figure 6:**
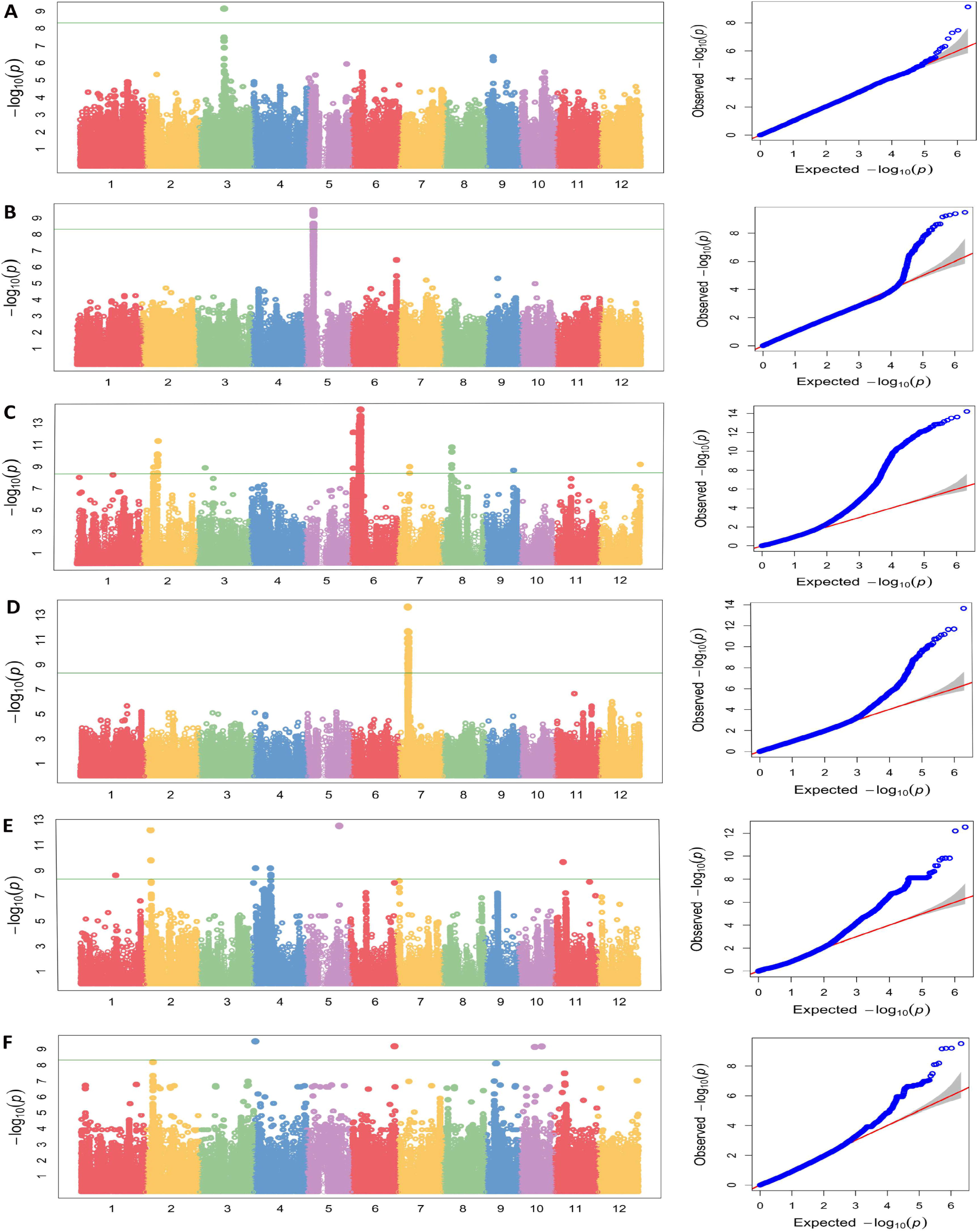
Genome-wide mapping for associations amon CC3 (520) accessions with various yield-related domesticated traits. **(A)** Manhattan and QQ plots of compressed MLM for grain length. Negative log_10_-transformed *P* values (y axis) values from the compressed mixed linear model are plotted against position (x axis) on different chromosomes. **(B)** Manhattan and QQ plots of compressed MLM for grain width, as plotted in A. (C) Manhattan and QQ plots of compressed MLM for endosperm type, as plotted in A. **(D)** Manhattan and QQ plots of compressed MLM for seed coat color, as plotted in A. **(E)** Manhattan and QQ plots of compressed MLM for secondary branching, as plotted in A. **(F)** Manhattan and QQ plots of compressed MLM for panicle axis, as plotted in A. Green line in each figure represents the genome-wide cut-off for significant association.

### GWAS using original (3004 accessions) panel

To further validate that the efficiency of designed mini-core collection CC3 (520 accessions) in terms of capturing the maximum possible marker-trait associations, we performed GWAS analyses with original collection of 3004 rice accessions for the above mentioned traits covering 500000 SNPs. The number of SNPs was reduced due to matrix limitations of the R program. In the case of original collection, only four [Endosperm Type (ET), Seed coat color (SCC), Grain Width (GW) and Grain Length (GL)] of the six traits showing significant marker-trait association in CC3 showed significant association. However, two traits namely 100 grain weight (HGW) and panicle threshability (PT) showed association in the case of original collection. These were missing in the analyses of CC3 mini-core (Supplemental Table S14). Overall, 1790 SNPs were found to be significantly associated with different traits covering phenotypic variation ranging from 5.6-51.4%. Grain Length (GL) was found to be associated with 325 SNPs present on chromosome 3 and 5. The most significant SNP (G/T) associated with the trait Grain Length (GL) present on chromosome 3 was at position 16733441 with FDR adjusted *P* value of 3.4 × 10^−43^ and explaining a phenotypic variation of 43.5% (Supplemental Figure S1A). Other SNP associated significantly with GL was found on chromosome 5 (G/A) at position 5361894 with FDR adjusted *P* value of 1.03 × 10^−9^ and detecting a phenotypic variation of 38.8%. Grain Width (GW) could be significantly associated with 737 SNPs specifically present on chromosome 5. Consistent with the earlier studies, the most significant SNP on chromosome 5 (C/T) was found at position 5371686 with FDR adjusted *P* value of 9.3 × 10^−34^ and explaining 51.4% of phenotypic variation (Supplemental Figure S1B). The other significant SNP at chromosome 5 (T/C) associated with GW was at 28019687 with FDR adjusted *P* value of 8.4 × 10^−6^ and detecting a phenotypic variation of 48%. The trait 100 grain weight (HGW) showed significant associations with 54 SNPs distributed on chromosomes 3 and 5. The most significant SNP was found to be present on chromosome 3 (G/T) at position 16733441 with FDR adjusted *P* value of 7.9 × 10^−5^ and explaining 35.2% of phenotypic variation. The other significant SNP at chromosome 5 (T/C) associated with HGW was at position 5375201 with FDR adjusted *P* value of 7.9 × 10^−5^ and detecting a phenotypic variation of 35.2% (Supplemental Figure S1C). The Endosperm Type (ET) trait was found to be associated with 503 SNPs present on chromosome 6. The most significant SNP (G/C) associated with the trait ET was present on chromosome 6 at position 1731808 with FDR adjusted *P* value of 1.03 × 10^−29^ and explaining a phenotypic variation of 20.2% (Supplemental Figure S1D). The other significant SNP at chromosome 6 (G/A) associated with ET was at position 6830286 with FDR adjusted *P* value of 3.4 × 10^−8^ and detecting a phenotypic variation of 15.6%. The trait Seed coat color (SCC) showed significant associations with 170 SNPs distributed on chromosome 2 and 7. The most significant SNP was found to be present on chromosome 7 (G/A) at position 6133394 with FDR adjusted *P* value of 6.6 × 10^−11^ and explaining 7.2% of phenotypic variation (Supplemental Figure S1E). The other significant SNP on chromosome 7 (G/T) associated with SCC was at position 6417000 with FDR adjusted *P* value of 1.7 × 10^−10^ and detecting a phenotypic variation of 7.1%. The other significantly associated SNP were found to be present on chromosome 7 (T/C) at position 6656052 with FDR adjusted *P* value of 1.8 × 10^−8^ and explaining phenotypic variation of 6.8% and on chromosome 2 (A/G) at position 32431463 with FDR adjusted *P* value of 3.7 × 10^−5^ and explaining phenotypic variation of 5.6%. The panicle threshability (PT) trait showed significant association with only 1 SNP present on chromosome 2 (C/T) at position 21739453 with FDR adjusted *P* value of 6.8 × 10^−3^ and explaining 16.4% of phenotypic variation (Supplemental Figure S1F). The detailed distribution of different SNPs associated with different traits such as Endosperm Type (ET), Seed coat color (SCC), Grain Width (GW), Grain Length (GL), 100 grain weight (HGW) and panicle threshability (PT) and their chromosomal positions for original collection of rice has been provided in Supplemental Table S14.

### GWAS based on regional and varietal grouping

Association analysis was also performed on different set of accessions based on varietal and regional genepool. The aim is to identify novel marker-trait association specific to a particular group which might have not been captured in case of association analysis of developed mini-core collection (CC3) and original collection of 3004 accessions of rice. Among the regional genepools, strong association were observed for various traits like Endosperm Type (ET), Grain Length (GL), Grain Width (GW), Panicle axis (PA), Secondary branching (SB), Seed coat color (SCC), 100 grain weight (HGW) and panicle threshability (PT) as evident in case of CC3 and original collection. Interestingly, panicle shattering (PS) was found to have strong association among African genotypes which was absent in case of CC3 and original panel of rice accessions. The most significant SNP was found to be present on chromosome 7 at position 19458367 with FDR adjusted *P* value of 2.9 × 10^−3^ and explaining 29.4% of phenotypic variation (Supplemental Figure S2A). The other significant SNP at chromosome 3 associated with PS was at position 15574478 with FDR adjusted *P* value of 3.3 × 10^−3^ and detecting a phenotypic variation of 28%.

In the case of varietal specific analysis, strong associations were again observed for numerous traits namely Endosperm Type (ET), Grain Length (GL), Grain Width (GW), Panicle axis (PA), Secondary branching (SB), Seed coat color (SCC), 100 grain weight (HGW) and panicle threshability (PT) as observed in CC3. Interestingly, here we identified one novel association for panicle shattering (PS) among the Temperate *japonica* group which was absent in the case of CC3 and the original panel of rice accessions. The most significant SNPs were found to be present on chromosomes 1 and chromosome 4 at positions 22646649 and 32213453 with FDR adjusted *P* value of 1.6 × 10^−7^ each, and explaining 61.5% of phenotypic variation each, respectively (Supplemental Figure S2B). The other significant SNP on chromosome 11 associated with PS was at position 4141451 with FDR adjusted *P* value of 1.6 × 10^−7^ and detecting a phenotypic variation of 54.7%. The other significantly associated SNP was found to be present on chromosome 2 at position 4724352 with FDR adjusted *P* value of 1.6 × 10^−7^ and explaining phenotypic variation of 54.7% and on chromosome 9 at position 169498 with FDR adjusted *P* value of 1.6 × 10^−7^ and explaining phenotypic variation of 54.7%.

### Linkage disequilibrium and haplotype analysis

To gain further insight into some of the less characterized marker-trait association such as panicle secondary branching found in the above analysis, the GWAS highlighted SNPs were studied for the LD pattern and haplotype block generation with their flanking nucleotides. In case of panicle secondary branching trait, 100 SNPs flanking the most associated SNP (SNP identity) were considered for the analysis and identification of the block containing the associated SNP. This block showed strong LD in a span of 2 kb, containing 7 neighboring SNPs including the associated one (Fig. 7A). Haplotype analysis of this block revealed that PSB_H1 (ATCAGGT) haplotype had the highest frequency (f=0.45) among all the haplotypes. Distribution of these haplotypes between light and dense panicle secondary branching revealed all but H4 haplotype having significant association with light level branching (Fig. 7A). Contrastingly, H4 haplotype showed inclination towards dense branching trait. One recent study has shown the association of gene *SHORT PANICLE* 1 (*SP1*) and elevated haplotype diversity among *japonica* than *indica* group (Jang et al., 2018).

**Figure 7:**
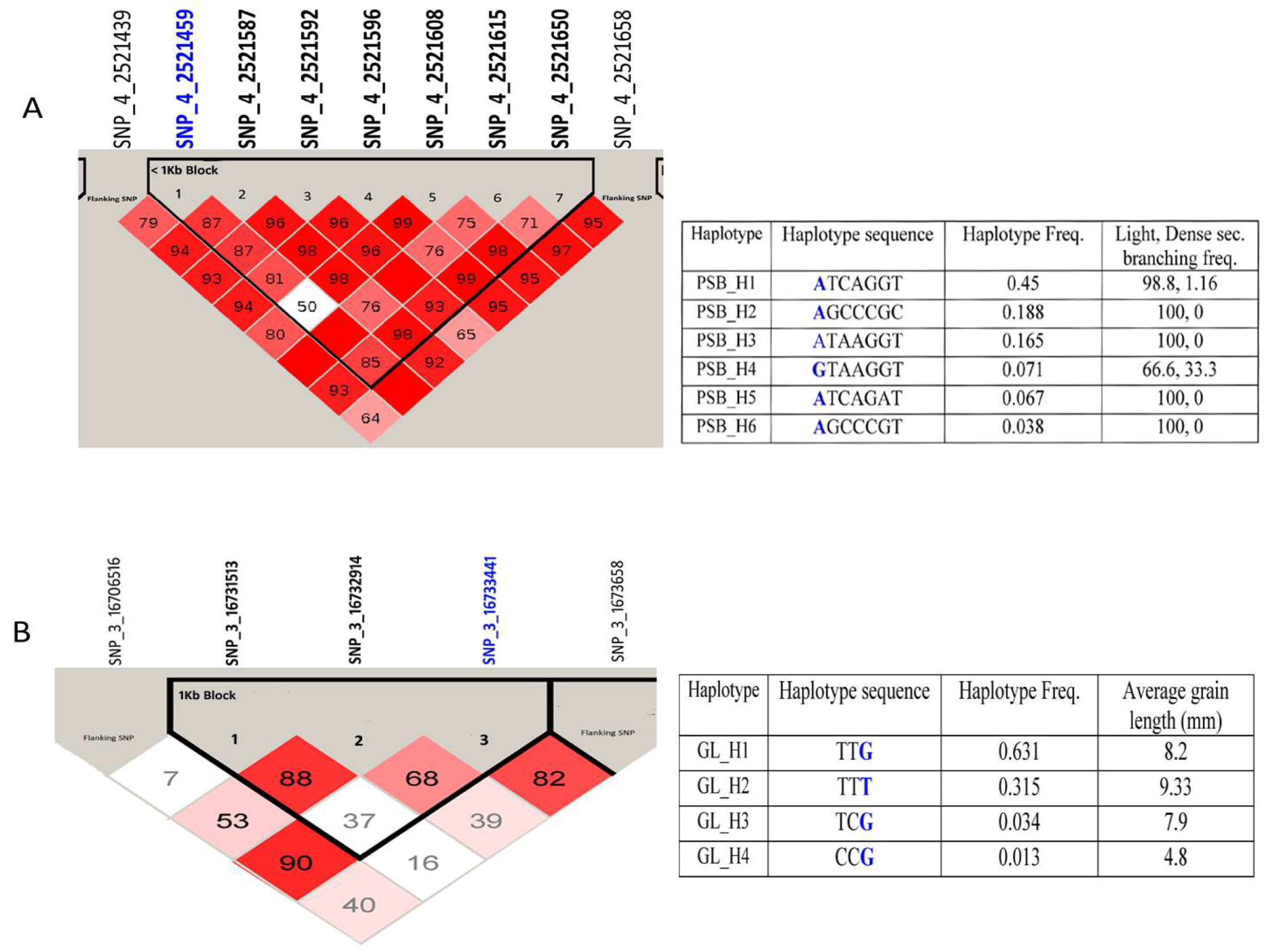
Linkage disequilibrium (LD) and haplotype analysis. **(A)** Depiction of strong LD and haplotype block containing the GWAS identified SNP for panicle secondary branching (PSB). The table shows the distribution of various haplotype for panicle secondary branching trait among core population. **(B)** Depiction of strong LD and haplotype block containing the GWAS identified SNP for grain length (GL). The table shows the distribution of various haplotype for panicle secondary branching trait among core population. GWAS identified SNP is highlight in blue. SNP id in bold format depicts Tag SNP. Red blocks, D’ (normalized linkage disequilibrium measure or D)≤ 1.0, with logarithm of odds (LOD) score ≥ 2.0; white blocks, D’ < 1.0 with LOD < 2.0; blue blocks, D’ = 1.0 with LOD < 2.0. Numbers in blocks denote D’ values. The genomic organization is described above the LD plot. LOD was defined as log10(Ll/L0), where L1 = likelihood of the data under linkage disequilibrium, and L0 = likelihood of the data under linkage equilibrium.

The grain length trait was also analyzed. LD analysis was performed for the identification of block containing the associated SNP. This block showed strong LD within a span of 1kb, containing 3 neighboring SNPs including the associated one (Fig. 7B). Haplotype analysis of this block revealed that GL_H1 (TTG) haplotype had the highest frequency (f=0.631) among all the haplotypes (GL_H2; TTT=0.315, GL_H3; TCG=0.034, GL_H4; CCG=0.013). Distribution of these haplotypes between long and short grain length revealed that GL_H2 in addition of being the second most frequent haplotype is also maximally associated with long rice grains, with average 9.33 mm grain length. Similarly, haplotype GL_H1 and GL_H3 identified with intermediate grain length i.e. showing mean value of 8.2 and 7.9 mm, respectively (Fig. 7B). On the other hand, H4 haplotype showed inclination towards short grain trait, having the least mean grain length of 4.8 mm. Above, haplotype observation was similar to one recent study dissecting the separate clustering of grain length haplotypes for varying size and different rice groups (Singh et al., 2017).

### GWAS correlation with expression data

Significantly associated SNPs were analysed for their position on the genome and association with gene(s) using the annotation data from RGAP (Kawahara et al., 2013). Rice has an estimated linkage disequilibrium value of around 200 Kb due to its self-pollinating nature (McNally et al., 2009). Keeping this in view, a genomic stretch of 100Kb up- and down-stream of the significantly associated SNPs was scanned to identify the genes. Expression level of genes/loci flanking the significantly associated SNPs was assessed in different tissues to be a probable candidate gene of function. The Grain Length (GL) trait which was found to be significantly associated with *GS3* in earlier studies (Fan et al., 2006) was highlighted in our analysis. We found one SNP (16733441;G/T) on chromosome 3 significantly associated with GL. This SNP belonged to the gene LOC_Os03g29260 showing high expression in seed of 5 DAP, embryo at 25 DAP and in pre- and post-emerging inflorescence (RGAP; Table 1). The Grain Width (GW) trait was found to be significantly associated with *qSW5* locus in earlier studies (Shomura et al., 2008). Here, we have identified identical SNP (C/G) at position 5371949 (chromosome 5; LOC_Os05g09500 and LOC_Os05g09550) showing strong association with Grain Width and higher expression in various tissues like seed of 5 DAP, embryo of 25 DAP and in pre- and post-emerging inflorescence (RGAP; Table 1). Significant marker-trait associations were observed for two traits namely 100 grain weight (HGW) and panicle threshability (PT) in the original collection (Supplemental Table S14). Interestingly, same two SNPs namely SNP (5371949;C/G) on chromosome 5 and SNP (16733441;G/T) on chromosome 3, were identified to influence the HGW trait. These SNPs were identified to be associated with Grain Length (GL) and Grain Width (GW) traits in CC3 accessions. A major QTL governing the traits of gelatinization temperature and amylose content, which in-turn affects Endosperm Type (ET), has been identified on chromosome 6 in various studies (GAO, 2003; Tian et al., 2009; Huang et al., 2010). In our analysis of mini-core collection CC3, the SNP (1765761;T/G) on chromosome 6 was found to be associated with Endosperm Type (ET). The gene encompassing this SNP could be *Waxy* (LOC_Os06g04200) showing higher expression in the seed of 5-10 DAP and in the endosperm of 25 DAP (RGAP; Table 1). Two other SNPs, one on chromosome 6 (6294468;G/T) and other on chromosome 2 (C/G;7413964) showed association with the ET trait, and were flanked by genes LOC_Os06g11812 and LOC_Os02g13840, respectively. These genes are predominately expressed in seed. On the similar note, the Secondary Branching (SB) trait significantly associated with SNP (C/T) at position 5032535 (chromosome 2; LOC_Os02g09780), SNP (A/G) at position 2521459 (chromosome 4; LOC_Os04g05050) and SNP (G/A) at position 1827355 (chromosome 7; LOC_Os07g04190) showed higher expression level in various tissues such as seed at 5 DAP, shoots, pre- and post-emergence inflorescence (RGAP; Table 1). The Seed Coat Color (SCC) trait was found to be associated with *Rc* gene in earlier studies (Sweeney et al., 2006). In our analysis, we found SNP (6124457;T/C) on chromosome 7, flanked by LOC_Os07g11020, which showed association with *Rc* gene. We also found SNP (T/G) at position 6660825 (chromosome 7; LOC_Os07g11900) associated with the trait SCC trait showing high expression level in various tissues such as seed at 5 and 10 DAP, and endosperm at 25 DAP (RGAP; Table 1). The Panicle Axis (PA) trait was found to be significantly associated with SNP (A/C) at position 1075655 (chromosome 4; LOC_Os04g02820 and LOC_Os04g02900) and SNP (G/A) at position 28676456 (chromosome 6; LOC_Os06g47320, LOC_Os06g47340 and LOC_Os06g47350) and showed higher expression level in different tissues such as pre- and post-emergence inflorescence, shoot, pistil and embryo at 25 DAP (RGAP; Table 1). The Panicle Threshability (PT) trait was found to be significantly associated with SNP (21739453;C/T) at chromosome 2. The expression database suggests the exclusive induction of flanking gene LOC_Os02g36150 in post-inflorescence (RGAP), indicating its plausible involvement in the trait. Thus, we were able to validate the known loci for various traits identified in earlier studies showing the effectiveness of CC3 in GWAS of different traits. Along with that we have also identified new loci showing strong marker trait associations which can be studied further to understand the underlying genetic make-up of these traits.

## CONCLUSIONS

Despite tremendous efforts, resolution of QTLs responsible for yield related traits and their causative genes have remained limited due to their complex nature. QTL mapping using a diverse panel and GWAS analysis has proven to be an effective tool to understand the genetic make-up. For GWAS analysis, estimation of underlying population structure of the panel under consideration is important which helps in avoiding spurious associations between phenotypes and genotypes (Pritchard and Rosenberg, 1999; Pritchard et al., 2000; Pritchard and Donnelly, 2001). Most of the earlier studies in rice have taken into consideration a particular population (Huang et al., 2010; Lu et al., 2015) which might have high level of structure and kinship for GWAS analysis resulting into spurious marker-trait associations. Here, this study for the first time utilized such large diversified (3004) rice germplasm, provided complete coverage of the global rice genepool. The mini-core was developed using program Core Hunter 3, based on 2 million genome-wide single nucleotide polymorphisms (SNPs), diverse 18 phenotypes and 89 country locations. The mini-core accounted for 17.3% of the original collection, and captured maximum of the SNP polymorphism, all phenotype and geographical regions. In addition, this panel showed low population structure and low or no kinship among the individuals of our panel, avoiding spurious marker-trait associations. Further, increase in number of admixed individuals in different clusters of structure analysis of CC3 showed that the panel was unstructured and diverse in nature, serving the purpose of using a mini-core for association analysis. Mini-core was validated using Shannon-Diversity Index.

On the utility front, the GWAS analyses with the designed mini-core panel found various novel marker-trait associations along with the validation of some of the earlier reported associations. More broadly, this analysis also provided a comparison between the mini-core CC3 and original collection and we were able to show that CC3 captured the associations prevalent in original collection and happens to be a representative subset. We found 5924 SNPs significantly associated with 6 different traits covering a phenotypic range of 10.4 – 61.6% in case of mini-core CC3, whereas identified 1790 SNPs significantly associated with 6 traits covering a phenotypic range of 4.9-51.4% for the original collection. Association analysis performed on different set of accessions based on varietal and regional genepool led to the identification of novel marker-trait associations which were not captured in CC3 and original collection of rice for the trait panicle shattering. This suggests that one marker can be very effective in one set of accessions but not in the other set. The accessions can be linked to each other by the geographical region or the other context.

Expression analysis highlighted the associated SNPs flanking gene(s), which possibly be involved in respective agronomic trait development. Overall, we identified at least one gene showing expression abundance at expected development stage for all the six associated agronomic traits. This study also performed haplotype analysis for two specific traits; panicle secondary branching and grain length. The secondary branching was dealt as it was not so well studied whereas grain length haplotype analysis was performed to reproduce the previous studies within mini-core. In conclusion, we were able to generate and validate one mini-core as robust, diversified, non-redundant and manageable association panel, efficiently mirroring the large and diverse collection of 3004 rice accessions. We suggest this relatively small subset can be effectively utilized for efficient agronomic traits evaluation, which in-turn be useful for marker assisted breeding program for rice crop improvement.

## MATERIALS AND METHODS

### Genotypic and phenotypic data of rice germplasm collection

The present study utilized SNP data of 3004 rice accession (hereafter named as original collection) and the phenotypic data for 18 yield related traits i.e. days to 80% heading (DEH), 100 grain weight (HGW), endosperm type (ET), days to first flower (DFF), grain length (GL), grain width (GW), leaf senescence (LS), panicle axis (PA), panicle length (PL), panicle shattering (PS), panicle threshability (PT), secondary branches (SB), seed coat color (SCC), seedling height (SH), spikelet fertility (SF), culm length (CL), culm number (CN) and culm diameter (CD) of agronomic value for the development of mini-core collections and association analysis. The data of 3000 rice genome project (3K RGP) for polymorphic SNPs (18.9 million) along with phenotypic data were retrieved from SNP-seek database (http://snp-seek.irri.org) (Alexandrov et al., 2015; Mansueto et al., 2017). In addition, we performed whole genome sequencing of 4 Indian accessions (LGR, PB-1121, Sonasal and Bindli) at a depth of 45X and collected phenotypic data for the above mentioned traits for the growing seasons 2016 and 2017. The accessions of initial collection of rice belonged to 89 countries representing all the regional pools and varieties of rice grown throughout the world.

### Isolation of genomic DNA, genome sequencing and SNP calling

All the four indian rice accessions [long grain: LGR (LG) and PB 1121 (PB); short grain: Sonasal (SN) and Bindli (BN)] were grown in the season of 2016 in NIPGR field. Rice seedling of 10 days old were used for the isolation of genomic DNA using Sigma GenEluteTM Plant gDNA kit. Integrity of genomic DNA was analyzed using Bioanalyzer 2100 (Agilent Technologies, Singapore). Samples for sequencing were prepared by Illumina TruSeq DNA sample preparation kit (Illumina, USA). Sequencing was performed with 90 bases pair-end chemistry on Illumina HiSeq 2000.

Subsequent standard bioinformatics analysis such as, raw reads quality check and removal of low quality bases (< Q30 Phred score) was done. The filtered reads were then mapped over the rice reference genome Nipponbare (IRGSP-1.0 pseudomolecule/MSU7) using BWA program with –q20 setting. Picard program was employed to get rid of duplicate reads.

Variant calling such single nucleotide polymorphisms (SNPs and InDels) were called via Genome Analysis TKLite-2.3-9 toolkit Unified Genotyper (GATK) (McKenna et al., 2010). The SNPs and InDels with a polymorphism call rate of < 90% were eliminated. After calling, in order to eliminate the low quality variants, a stringent of read depth ≥10 and quality score ≥ 30 was adopted to filter total variants and only good quality variants were considered for further analysis. Further, all the SNPs, those are consecutive and adjacent to InDels, were also eliminated. Plant Ensembl database was used to gene model for annotation. All the identified SNPs and InDels were annotated using customized VariMAT (SciGenome, India).

### Development of mini-core collections

The program Core Hunter 3 (De Beukelaer et al., 2018) was used for development of independent mini-core collections using phenotypic and genotypic data. A cut off value of 10% of the initial collection was used to design the mini-core collections with default parameters. The mini-cores were also assessed for coverage of entire range for all the quantitative traits with reference to the initial collection. The diversity captured within the mini-core collections with respect to the initial collection was assessed using different evaluation indices such as Shannon’s diversity index (*I*), Nei’s gene diversity (*H*), mean difference percentage (MD%), variance difference percentage (VD%), variable rate of coefficient of variance (VR%) and coincidence rate of range (CR%; Hu et al., 2000). Pearson correlation coefficient (r) was used to determine correlation between different quantitative traits using PAST version 3.10 (Hammer et al., 2001).

### Phylogenetic and population structure analysis

The SNP data was utilized to construct distance-based dendrogram through Maximum-likelihood method using SNP Phylo program (Lee et al., 2014). Principal Coordinate Analysis (PCoA) was performed to estimate the overall relationship between accessions. Bayesian analysis of the population structure was performed using fastSTRUCTURE program (Raj et al., 2014) which estimated the optimal K value for the dataset. Pair-wise kinship coefficient was estimated using the program SPAGeDi (Hardy and Vekemans, 2002). To estimate the proportion of ancestral contribution for each accession admixture model was followed. The analysis was performed independent of the geographical and varietal origin of accessions. Accessions with Q value (membership proportion) >= 80% were considered as pure and designated into a particular cluster while accessions with Q value < 80% were considered as admixtures.

### Genome-wide association analysis

All the Genome-wide association analysis were performed utilizing the program GAPIT (Lipka et al., 2012) based on compressed mixed linear model (MLM) for 18 rice agronomic traits. In case of original panel and developed mini-core, 2081521 and 500000 SNPs makers were utilized, respectively. Selected SNPs marker have minor allele frequency [MAF] > than 0.02.

### Analysis of expression data

Expression levels of all ORF’s flanking 100 Kb up- and down-stream of significantly associated SNPs were assessed in different tissues within RGAP MSUv7 expression database (Kawahara et al., 2013). Further, gene loci showing preferential expression within plant organ responsible for trait development was notified as the plausible causative gene(s) associated studied trait.

### LD and Haplotype analyses

Haplotypes were generated from the genotyped data. The linkage disequilibrium (LD) and haplotype analysis were performed using Haploview 4.2 (Barrett et al., 2005) with default parameters (MAF<0.001), HWE test (<0.001) and percent genotypes test (cut of value=75%). Four gamete rule method was employed to identify the more refined genomic block containing the associated SNP.

## ACKNOWLEDGMENTS

This work was financially supported by the grants BT/AB/NIPGR/SEED BIOLOGY/2012 and for Sub-DIC facility from Department of Biotechnology (DBT), Government of India, to JKT. A.K acknowledges University Grant Commission and NIPGR for the Fellowships. S.K acknowledges NPDF (DST) and Short-Term Research Fellowship from NIPGR. The authors are thankful to DBT-eLibrary Consortium (DeLCON) for providing access to literature.

