## Supplemental Tables S1-12,14 for "Designing of a mini-core that effectively represents 3004 diverse accessions of rice"

**Supplemental Table S1**

**Supplemental Table S2**

**Supplemental Table S3**

**Supplemental Table S4**

**Supplemental Table S5**

**Supplemental Table S6**

**Supplemental Table S7**

**Supplemental Table S8**

**Supplemental Table S9**

**Supplemental Table S10**

**Supplemental Table S11**

**Supplemental Table S12**

**Supplemental Table S13 (Excel sheet, uploaded separately)**

**Supplemental Table S14**

**Supplemental Table S1:** Range of quantitative traits in original collection and different core collections.

| **Traits** | **Original collection (3004 acc)** | | **CC1 (231acc)** | | **CC2 (300 acc)** | | **Merged CC1 &CC2 (503 acc)** | | **CC3 (503 + 17 = 520 acc)** | |
|---|---|---|---|---|---|---|---|---|---|---|
| | Max | Min | Max | Min | Max | Min | Max | Min | Max | Min |
| Days to 80% flowering | 184 | 50 | 175 | 50 | 175 | 52 | 175 | 50 | 184 | 50 |
| 100 GW (gm) | 5 | 0.98 | 4.6 | 0.98 | 4.6 | 0.98 | 4.6 | 0.98 | 5 | 0.98 |
| Days to 1st flowering | 182 | 45 | 171 | 46 | 171 | 45 | 171 | 45 | 182 | 45 |
| Grain length (mm) | 12.7 | 4.4 | 12.7 | 4.4 | 12.4 | 4.4 | 12.7 | 4.4 | 12.7 | 4.4 |
| Grain width (mm) | 4.4 | 1.5 | 4.1 | 1.7 | 4.3 | 2.1 | 4.4 | 1.7 | 4.4 | 1.5 |
| Panicle length (cm) | 37 | 13 | 36 | 13 | 36 | 13 | 36 | 13 | 37 | 13 |
| Seed coat color | 99 | 10 | 88 | 10 | 99 | 10 | 99 | 10 | 99 | 10 |
| Seedling height (cm) | 74 | 12 | 71 | 16 | 74 | 13 | 74 | 13 | 74 | 12 |
| Culm length (cm) | 204 | 27 | 204 | 35 | 181 | 27 | 204 | 27 | 204 | 27 |
| Culm number | 40 | 5 | 40 | 6 | 33 | 5 | 40 | 5 | 40 | 5 |
| Culm diameter (mm) | 9.1 | 2 | 9.1 | 2 | 7.7 | 3 | 9.1 | 2 | 9.1 | 2 |

Highlighted traits were not picked up for their complete range in any mini-core collection (CC1, CC2 and CC3). Seventeen accessions were included in CC3 (503+17 = 520 accessions) to cover the entire range for all the traits with respect to original collection.

**Supplemental Table S2:** Assessment of mini-core collections for various evaluation indices using phenotypic data.

| **Core collection** | **MD%** | **VD%** | **VR%** | **CR%** | **H** | **I** |
|---|---|---|---|---|---|---|
| CC1 (231 acc) | 4.08 | 39.77 | 86 | 92 | 2.25 | 0.79 |
| CC2 (300 acc) | 2.8 | 19.78 | 107.68 | 91.1 | 1.98 | 0.77 |
| CC3 (520 acc) | 2.9 | 18.9 | 109.3 | 96.2 | 2.17 | 0.79 |

MD% *(Mean difference percentage)*, VD% *variance difference percentage*, VR % *(Variable rate of coefficient of variance*), CR% (*coincidence rate of range*), H (Shannon diversity index), I (Nei's diversity index)

**Supplemental Table S3:** Distribution of accessions from different varietal groups in different mini-core collections to check proportionate representation from original collection.

| **Core/varietal group** | ***indica*** | **Tropical *japonica*** | **Temperate *japonica*** | ***japonica*** | ***aus/boro*** | **Intermediate** | **Aromatic (Basmati)** |
|---|---|---|---|---|---|---|---|
| Original collection (3004 acc) | **1743** | **388** | **320** | **132** | **215** | **135** | **71** |
| CC1 (231 acc) | **129** (7.4%) | **15** (3.8%) | **38** (11.9%) | **14** (10.6%) | **11** (5.1%) | **19** (14%) | **5** (7%) |
| CC2 (300 acc) | **171** (9.8%) | **10** (2.5%) | **8** (2.5%) | **12** (9.1%) | **42** (19.5%) | **45** (33.3%) | **12** (16.9%) |
| CC3 (520 acc) | **295** (16.9%) | **27** (6.9%) | **44** (13.4%) | **23** (17.4%) | **55** (25.6%) | **61** (44.4%) | **15** (21%) |

**Supplemental Table S4:** Distribution of accessions from different regional gene pools in different mini-core collections to check proportionate representation from original collection.

| **Core/ Regions** | **South-Asia** | **South East-Asia** | **China** | **Africa** | **America** | **Europe** | **East Asia** | **Oceania** | **Unknown** |
|---|---|---|---|---|---|---|---|---|---|
| Original collection (3004 acc) | **787** | **1016** | **482** | **252** | **166** | **118** | **132** | **17** | **34** |
| CC1 (231 acc) | **55** (6.9%) | **52** (5.1%) | **52** (10.8%) | **15** (5.9%) | **17** (10.2%) | **18** (15.2%) | **15** (11.4%) | **4** (23.5%) | **3** (8.8%) |
| CC2 (300 acc) | **122** (15.5%) | **70** (6.9%) | **55** (11.4%) | **23** (9.1%) | **13** (7.8%) | **2** (1%) | **8** (6%) | **1** (5.9%) | **6** (17.4%) |
| CC3 (520 acc) | **176** (22.4%) | **123** (12%) | **101** (20.95%) | **38** (15%) | **28** (16.9%) | **19** (16.1%) | **21** (15.9%) | **5** (29.4%) | **9** (25.5%) |

**Supplemental Table S5:** Correlation coefficient analysis for quantitative traits in CC3 accessions.

| Traits/traits | A | B | C | D | E | F | G | H | I | J | K |
|---|---|---|---|---|---|---|---|---|---|---|---|
| **A** | -- | | | | | | | | | | |
| **B** | -0.07 | -- | | | | | | | | | |
| **C** | **0.998** | -0.2 | -- | | | | | | | | |
| **D** | 0.07 | 0.44 | -0.04 | -- | | | | | | | |
| **E** | -0.227 | 0.542 | -0.27 | 0.021 | -- | | | | | | |
| **F** | 0.370 | 0.086 | 0.489 | 0.244 | -0.025 | -- | | | | | |
| **G** | -0.095 | 0.051 | -0.008 | -0.02 | 0.084 | -0.023 | -- | | | | |
| **H** | 0.139 | 0.037 | 0.156 | 0.153 | -0.021 | 0.08 | 0.158 | -- | | | |
| **I** | 0.555 | 0.065 | 0.68 | 0.165 | 0.005 | 0.557 | 0.069 | 0.292 | -- | | |
| **J** | -0.033 | -0.16 | -0.27 | -0.09 | -0.208 | -0.144 | 0.029 | -0.13 | -0.112 | -- | |
| **K** | 0.426 | 0.097 | 0.614 | 0.112 | -0.03 | 0.341 | -0.03 | 0.096 | 0.391 | -0.04 | -- |

*Values > 0.7 were significant, **values between 0.3-0.7 were weakly significant, P < 0.001
A - days to 80% heading (DEH), B - 100 grain weight (HGW), C - days to first flower (DFF), D - grain length (GL), E - grain width (GW), F - panicle length (PL), G - secondary branches (SB), H- seedling height (SH), I - culm length (CL), J - culm number (CN) and K - culm diameter (CD)

**Supplemental Table S6:** Distribution of 3004 accessions of original rice collection in different clusters of maximum likelihood dendrogram (based on different varietal group).

| Cluster/varietal group | *indica* | *japonica* | Tropical *japonica* | Temperate *japonica* | *aus/boro* | Intermediate | Aromatic (Basmati) |
|---|---|---|---|---|---|---|---|
| **Cluster Ia (1771 acc)** | 1641 (92.6%) | 7 | 27 | 30 | 30 | 26 | 10 |
| **Cluster Ib (216 acc)** | 25 | 1 | 2 | 7 | 172 (72.6%) | 5 | 4 |
| **Cluster IIa (519 acc)** | 35 | 80 (15.4%) | 329 (63.3%) | 31 | 4 | 35 | 5 |
| **Cluster IIb (358 acc)** | 18 | 36 | 22 | 250 (69.8%) | 6 | 25 | 1 |
| **Cluster IIc (97 acc)** | 9 | 5 | 6 | 1 | 3 | 23 (23.7%) | 50 (51.5%) |
| **Un-clustered accessions (43 acc)** | 15 (34.8%) | 3 | 2 | 1 | 0 | 21 (48.8%) | 1 |

**Supplemental Table S7:** Distribution of CC3 accessions (520) in different clusters of maximum-likelihood dendrogram of original collection of rice (3004 accessions).

| **Accession distribution in cluster of maximum-likelihood dendrogram of original collection (3004 accessions)** | **Distribution of accession from different clusters of maximum-likelihood dendrogram captured in CC3 (520 accessions)** |
|---|---|
| Cluster Ia - 1771 accessions | Cluster Ia - 322 accessions (18.1%) |
| Cluster Ib - 216 accessions | Cluster Ib - 42 accessions (19.4%) |
| Cluster IIa - 519 accessions | Cluster IIa - 55 accessions (10.5%) |
| Cluster IIb - 358 accessions | Cluster IIb - 65 accessions (18%) |
| Cluster IIc - 97 accessions | Cluster IIc - 25 accessions (25.7%) |
| Un-clustered group - 43 accessions | Un-clustered group - 11 accessions (25%) |

**Supplemental Table S8:** Distribution of different varietal group of original collection (3004 acc) in different clusters of FastSTRUCTURE analysis.

| **Varietal group/ Cluster** | **FSTR CL1 (219 acc)** | **FSTR CL2 (522 acc)** | **FSTR CL3 (90 acc)** | **FSTR CL4 (973 acc)** | **FSTR CL5 (372 acc)** | **FSTR CL6 (323 acc)** | **FSTR CL7 (505 acc)** |
|---|---|---|---|---|---|---|---|
| *indica* (1743 acc) | 28 | 47 | 9 | **885 (91%)** | 26 | **297 (92%)** | **451 (89.3%)** |
| *japonica* (132 acc) | 1 | **94 (18%)** | 4 | 5 | 25 | 0 | 3 |
| Temperate *japonica* (320 acc) | 4 | 35 | 1 | 17 | **248 (66.6%)** | 6 | 9 |
| Tropical *japonica* (388 acc) | 1 | **310 (59.3%)** | 2 | 26 | 35 | 5 | 9 |
| *aus/boro* (215 acc) | **179 (81.7%)** | 3 | 5 | 14 | 5 | 4 | 5 |
| Intermediate (135 acc) | 3 | **27** | **19 (21.1%)** | **21** | **29** | 9 | **27** |
| Aromatic (Basmati) (71 acc) | 3 | 6 | **50 (55.5%)** | 5 | 4 | 2 | 1 |

**Supplemental Table S9:** Analysis of original collection (3004 accessions) for admixtures through population structure using FastSTRUCTURE.

| Pure accessions (1762 acc) | | Admixtures (1242 acc) | |
|---|---|---|---|
| **Structure analysis (K=7)** | | **Structure analysis (K=7)** | |
| FSTR CL 1 | 189 | FSTR CL 1 | 30 |
| FSTR CL 2 | 330 | FSTR CL 2 | 192 |
| FSTR CL 3 | 64 | FSTR CL 3 | 26 |
| FSTR CL 4 | 591 | FSTR CL 4 | 382 |
| FSTR CL 5 | 256 | FSTR CL 5 | 116 |
| FSTR CL 6 | 148 | FSTR CL 6 | 145 |
| FSTR CL 7 | 154 | FSTR CL 7 | 351 |
| **Varietal group (K=7)** | | **Varietal group (K=7)** | |
| *indica* | 922 | *indica* | 821 |
| *japonica* | 90 | *japonica* | 42 |
| Temperate *japonica* | 234 | Temperate *japonica* | 86 |
| Tropical *japonica* | 236 | Tropical *japonica* | 152 |
| *aus/ boro* | 186 | *aus/ boro* | 29 |
| Aromatic (Basmati) | 53 | Aromatic (Basmati) | 18 |
| Intermediate | 41 | Intermediate | 94 |
| **Region wise (K=7)** | | **Region wise (K=7)** | |
| South Asia | 497 | South Asia | 290 |
| South East Asia | 567 | South East Asia | 449 |
| China | 275 | China | 207 |
| Africa | 166 | Africa | 86 |
| America | 87 | America | 79 |
| Europe | 53 | Europe | 65 |
| East Asia | 89 | East Asia | 43 |
| Oceania | 9 | Oceania | 8 |
| Unknown | 19 | Unknown | 15 |

Accessions with ≥ 80% genome similarity were considered as pure while accessions with < 80% shared genome were termed as admixtures. Accessions highlighted in red have around equal or more number of admixtures than pure accessions.

**Supplemental Table S10:** Distribution of CC3 accessions (520 accessions) in FastSTRUCTURE derived clusters of original collection of rice (3004 accessions).

| FastStructure Clusters | Accessions from original collection | Accessions of original collection with Q value > 80% (Pure) | Accessions of original collection with Q value < 80% (Admixtures) | Accessions picked from original collection in CC3 mini-core | Accessions of CC3 with Q value > 80% (Pure) | Accessions of CC3 with Q value < 80% (Admixture) |
|---|---|---|---|---|---|---|
| FSTR CL 1 | **219** | 189 | 30 | **50** | 40 | 10 |
| FSTR CL 2 | **522** | 330 | 192 | **42** | **23** | **19** |
| FSTR CL 3 | **90** | 64 | 26 | **24** | **13** | **11** |
| FSTR CL 4 | **973** | 591 | 382 | **185** | 109 | 76 |
| FSTR CL 5 | **372** | 256 | 116 | **74** | **37** | **37** |
| FSTR CL 6 | **323** | **148** | **145** | **61** | **28** | **33** |
| FSTR CL 7 | **505** | **154** | **351** | **84** | **25** | **59** |

Accessions with ≥ 80% genome similarity were considered as pure while accessions with < 80% shared genome were termed as admixtures. Accessions highlighted in red have around equal or more number of admixtures than pure accessions.

**Supplemental Table S11:** Distribution of CC3 accessions (520 acc) based on varietal groups in different clusters of FastSTRUCTURE analysis (K=7). Numbers in parentheses represents accessions.

| Cluster/ Varietal group | *indica* (295) | Tropical *japonica* (27) | Temperate *japonica* (44) | *Japonica* (23) | *aus/boro* (55) | Intermediate (61) | Aromatic (Basmati) (15) |
|---|---|---|---|---|---|---|---|
| CC CL1 (73 acc) | 57 | 0 | 1 | 1 | 1 | 12 | 1 |
| CC CL2 (23 acc) | 1 | 0 | 0 | 0 | 5 | 8 | 9 |
| CC CL3 (49 acc) | 6 | 0 | 1 | 1 | 41 | 0 | 0 |
| CC CL4 (70 acc) | 58 | 1 | 0 | 0 | 3 | 8 | 0 |
| CC CL5 (43 acc) | 3 | 17 | 4 | 10 | 0 | 8 | 1 |
| CC CL6 (191 acc) | 165 | 5 | 2 | 2 | 4 | 11 | 2 |
| CC CL7 (71 acc) | 5 | 4 | 36 | 9 | 1 | 14 | 2 |

**Supplemental Table S12:** Analysis of CC3 (520 accessions) for admixtures through population structure using FastSTRUCTURE.

| Pure accessions (275 accessions) | | Admixtures (245 accessions) | |
|---|---|---|---|
| **Structure analysis (K=7)** | | **Structure analysis (K=7)** | |
| CC CL1 | 25 | CC CL1 | 48 |
| CC CL2 | 13 | CC CL2 | 10 |
| CC CL3 | 40 | CC CL3 | 9 |
| CC CL4 | 28 | CC CL4 | 42 |
| CC CL5 | 23 | CC CL5 | 20 |
| CC CL6 | 109 | CC CL6 | 82 |
| CC CL7 | 37 | CC CL7 | 34 |
| **Varietal group (K=7)** | | **Varietal group (K=7)** | |
| *indica* | 151 | *indica* | 144 |
| *japonica* | 11 | *japonica* | 12 |
| Temperate *japonica* | 29 | Temperate *japonica* | 15 |
| Tropical *japonica* | 14 | Tropical *japonica* | 13 |
| *aus/ boro* | 45 | *aus/ boro* | 10 |
| Aromatic (Basmati) | 11 | Aromatic (Basmati) | 4 |
| Intermediate | 14 | Intermediate | 47 |
| **Region wise (K=7)** | | **Region wise (K=7)** | |
| South Asia | 106 | South Asia | 70 |
| South East Asia | 60 | South East Asia | 63 |
| China | 47 | China | 54 |
| Africa | 23 | Africa | 15 |
| America | 15 | America | 23 |
| Europe | 6 | Europe | 13 |
| East Asia | 11 | East Asia | 10 |
| Oceania | 2 | Oceania | 3 |
| Unknown | 5 | Unknown | 4 |

Accessions with ≥ 80% genome similarity were considered as pure while accessions with < 80% shared genome were termed as admixtures. Accessions highlighted in red have equal or more number of admixtures than pure accessions

**Supplemental Table S14:** Association analysis of 3004 accessions of original collection.

| Trait | Chr | Position | Major allele | Minor allele | Minor allele frequency | Nipp. allele | CC3 allele | P-value FDR adjusted | $R^2$ value (%) | Known loci |
|---|---|---|---|---|---|---|---|---|---|---|
| Grain length | 3 | 16733441 | G | T | 0.36 | G | T | $3.4 \times 10^{-43}$ | 43.5 | *GS3* |
| Grain length | 5 | 5361894 | G | A | 0.36 | G | A | $3.4 \times 10^{-9}$ | 38.8 | *qSW5* |
| Grain width | 5 | 5371686 | C | T | 0.49 | C | T | $9.3 \times 10^{-34}$ | 51.4 | *qSW5* |
| Grain width | 5 | 28019687 | T | C | 0.10 | T | C | $8.4 \times 10^{-6}$ | 48 | |
| Hundred Grain weight | 3 | 16733441 | G | T | 0.36 | G | T | $7.9 \times 10^{-5}$ | 35.2 | *GS3* |
| Hundred Grain weight | 5 | 5375201 | T | C | 0.48 | T | C | $7.9 \times 10^{-5}$ | 35.2 | *qSW5* |
| Endosperm type | 6 | 1731808 | G | C | 0.20 | G | C | $1.03 \times 10^{-29}$ | 20.2 | *waxy* |
| Endosperm type | 6 | 6830286 | G | A | 0.21 | G | A | $3.4 \times 10^{-8}$ | 15.6 | |
| Seed coat color | 7 | 6133394 | G | A | 0.26 | G | A | $6.6 \times 10^{-11}$ | 7.2 | *Rc* |
| Seed coat color | 7 | 6417000 | G | T | 0.32 | G | T | $1.7 \times 10^{-10}$ | 7.1 | |
| Seed coat color | 7 | 6656052 | T | C | 0.43 | T | C | $1.8 \times 10^{-8}$ | 6.8 | |
| Seed coat color | 2 | 32431463 | A | G | 0.27 | A | G | $3.7 \times 10^{-5}$ | 5.6 | |
| Panicle thresability | 2 | 21739453 | C | T | 0.23 | C | T | $6.8 \times 10^{-3}$ | 16.4 | |

Nipp; Nipponbare, Chr; Chromosome
