## Supplemental Tables S13 for "Designing of a mini-core that effectively represents 3004 diverse accessions of rice"

**Supplemental Table S13: Details of significantly associated SNPs (FDR adjusted p-value < 0.05) with different agronomic traits across the CC3 panel (520) rice accessions**

Endospem_type

| SNP | Chr. | Position | P.value | maf | nobs | $R^2$ with SNP | FDR_Adjusted_P-values |
|---|---|---|---|---|---|---|---|
| SNP_A.K_999520_6_6294468 | 6 | 6294468 | 6.05E-15 | 0.0786 | 299 | 0.291132498 | 1.26E-08 |
| SNP_A.K_999073_6_6254780 | 6 | 6254780 | 2.35E-14 | 0.08528 | 299 | 0.28089951 | 2.12E-08 |
| SNP_A.K_999754_6_6312567 | 6 | 6312567 | 3.06E-14 | 0.08863 | 299 | 0.278936803 | 2.12E-08 |
| SNP_A.K_999759_6_6312665 | 6 | 6312665 | 5.27E-14 | 0.08863 | 299 | 0.274878583 | 2.74E-08 |
| SNP_A.K_998917_6_6234398 | 6 | 6234398 | 6.99E-14 | 0.08528 | 299 | 0.272778142 | 2.91E-08 |
| SNP_A.K_1000416_6_6382958 | 6 | 6382958 | 1.17E-13 | 0.0903 | 299 | 0.268937006 | 3.11E-08 |
| SNP_A.K_993625_6_5780299 | 6 | 5780299 | 1.31E-13 | 0.0903 | 299 | 0.268123575 | 3.11E-08 |
| SNP_A.K_999753_6_6312564 | 6 | 6312564 | 1.36E-13 | 0.08863 | 299 | 0.267834759 | 3.11E-08 |
| SNP_A.K_1000516_6_6390607 | 6 | 6390607 | 1.43E-13 | 0.08696 | 299 | 0.267481017 | 3.11E-08 |
| SNP_A.K_998815_6_6226255 | 6 | 6226255 | 1.49E-13 | 0.08863 | 299 | 0.267140217 | 3.11E-08 |
| SNP_A.K_993650_6_5781607 | 6 | 5781607 | 2.35E-13 | 0.09197 | 299 | 0.263802539 | 4.32E-08 |
| SNP_A.K_994404_6_5856411 | 6 | 5856411 | 2.49E-13 | 0.08027 | 299 | 0.26337929 | 4.32E-08 |
| SNP_A.K_994355_6_5850932 | 6 | 5850932 | 2.71E-13 | 0.09532 | 299 | 0.262758569 | 4.33E-08 |
| SNP_A.K_995823_6_5951741 | 6 | 5951741 | 3.71E-13 | 0.0903 | 299 | 0.260434516 | 5.52E-08 |
| SNP_A.K_999049_6_6249991 | 6 | 6249991 | 4.52E-13 | 0.09197 | 299 | 0.258984871 | 6.12E-08 |
| SNP_A.K_995382_6_5931081 | 6 | 5931081 | 4.70E-13 | 0.12542 | 299 | 0.258700998 | 6.12E-08 |
| SNP_A.K_1000201_6_6354533 | 6 | 6354533 | 5.50E-13 | 0.08863 | 299 | 0.257551968 | 6.43E-08 |
| SNP_A.K_999736_6_6311774 | 6 | 6311774 | 5.83E-13 | 0.08361 | 299 | 0.257126879 | 6.43E-08 |
| SNP_A.K_999643_6_6302290 | 6 | 6302290 | 5.92E-13 | 0.08696 | 299 | 0.257008495 | 6.43E-08 |
| SNP_A.K_999737_6_6311779 | 6 | 6311779 | 7.16E-13 | 0.0903 | 299 | 0.255622588 | 6.43E-08 |
| SNP_A.K_993674_6_5783531 | 6 | 5783531 | 7.17E-13 | 0.08361 | 299 | 0.255609383 | 6.43E-08 |
| SNP_A.K_999522_6_6294475 | 6 | 6294475 | 7.50E-13 | 0.07525 | 299 | 0.255277734 | 6.43E-08 |
| SNP_A.K_995627_6_5943868 | 6 | 5943868 | 7.67E-13 | 0.12375 | 299 | 0.25511368 | 6.43E-08 |
| SNP_A.K_978696_6_1765761 | 6 | 1765761 | 7.79E-13 | 0.13043 | 299 | 0.255007736 | 6.43E-08 |
| SNP_A.K_994328_6_5845302 | 6 | 5845302 | 8.09E-13 | 0.09365 | 299 | 0.254729601 | 6.43E-08 |
| SNP_A.K_1000581_6_6397558 | 6 | 6397558 | 8.20E-13 | 0.08361 | 299 | 0.254631025 | 6.43E-08 |
| SNP_A.K_995269_6_5923617 | 6 | 5923617 | 8.34E-13 | 0.0903 | 299 | 0.254507649 | 6.43E-08 |
| SNP_A.K_1000563_6_6396376 | 6 | 6396376 | 1.03E-12 | 0.08696 | 299 | 0.252947395 | 7.68E-08 |
| SNP_A.K_999844_6_6327999 | 6 | 6327999 | 1.09E-12 | 0.09197 | 299 | 0.252538245 | 7.84E-08 |
| SNP_A.K_999914_6_6332248 | 6 | 6332248 | 1.20E-12 | 0.0903 | 299 | 0.251855476 | 8.18E-08 |
| SNP_A.K_999899_6_6331984 | 6 | 6331984 | 1.22E-12 | 0.13545 | 299 | 0.251738046 | 8.18E-08 |
| SNP_A.K_993639_6_5780757 | 6 | 5780757 | 1.40E-12 | 0.09532 | 299 | 0.25071359 | 9.13E-08 |
| SNP_A.K_998850_6_6228661 | 6 | 6228661 | 1.60E-12 | 0.08528 | 299 | 0.249751738 | 1.01E-07 |
| SNP_A.K_994487_6_5868508 | 6 | 5868508 | 1.79E-12 | 0.08361 | 299 | 0.248952856 | 1.07E-07 |
| SNP_A.K_994483_6_5868337 | 6 | 5868337 | 1.83E-12 | 0.08361 | 299 | 0.248795707 | 1.07E-07 |
| SNP_A.K_998813_6_6225970 | 6 | 6225970 | 1.86E-12 | 0.09365 | 299 | 0.248656659 | 1.07E-07 |
| SNP_A.K_994320_6_5843882 | 6 | 5843882 | 1.90E-12 | 0.08696 | 299 | 0.248524277 | 1.07E-07 |
| SNP_A.K_998795_6_6223701 | 6 | 6223701 | 2.02E-12 | 0.08696 | 299 | 0.248059107 | 1.11E-07 |
| SNP_A.K_999625_6_6301269 | 6 | 6301269 | 2.12E-12 | 0.08863 | 299 | 0.247711345 | 1.13E-07 |
| SNP_A.K_993671_6_5783170 | 6 | 5783170 | 2.29E-12 | 0.09365 | 299 | 0.247145893 | 1.19E-07 |
| SNP_A.K_995537_6_5940623 | 6 | 5940623 | 2.36E-12 | 0.08696 | 299 | 0.246924467 | 1.19E-07 |
| SNP_A.K_999271_6_6275695 | 6 | 6275695 | 2.44E-12 | 0.12709 | 299 | 0.246695873 | 1.19E-07 |
| SNP_A.K_999638_6_6301875 | 6 | 6301875 | 2.45E-12 | 0.09197 | 299 | 0.246652771 | 1.19E-07 |
| SNP_A.K_999167_6_6264479 | 6 | 6264479 | 2.65E-12 | 0.08528 | 299 | 0.246099147 | 1.25E-07 |
| SNP_A.K_996759_6_6024319 | 6 | 6024319 | 2.89E-12 | 0.09699 | 299 | 0.245475946 | 1.34E-07 |

| | | | | | | | |
|---|---|---|---|---|---|---|---|
| SNP_A.K_995533_6_5940415 | 6 | 5940415 | 3.27E-12 | 0.12542 | 299 | 0.24458706 | 1.48E-07 |
| SNP_A.K_999222_6_6268799 | 6 | 6268799 | 3.45E-12 | 0.08696 | 299 | 0.244196963 | 1.53E-07 |
| SNP_A.K_999333_6_6282344 | 6 | 6282344 | 3.97E-12 | 0.12375 | 299 | 0.243174944 | 1.72E-07 |
| SNP_A.K_999422_6_6287576 | 6 | 6287576 | 4.07E-12 | 0.08863 | 299 | 0.242989809 | 1.73E-07 |
| SNP_A.K_993654_6_5781741 | 6 | 5781741 | 4.28E-12 | 0.08528 | 299 | 0.242628753 | 1.76E-07 |
| SNP_A.K_328503_2_10277497 | 2 | 10277497 | 4.31E-12 | 0.02676 | 299 | 0.242580737 | 1.76E-07 |
| SNP_A.K_999338_6_6282725 | 6 | 6282725 | 4.50E-12 | 0.09532 | 299 | 0.242268681 | 1.80E-07 |
| SNP_A.K_994564_6_5873622 | 6 | 5873622 | 4.87E-12 | 0.08696 | 299 | 0.241700627 | 1.91E-07 |
| SNP_A.K_1000565_6_6396414 | 6 | 6396414 | 5.04E-12 | 0.08696 | 299 | 0.241459782 | 1.94E-07 |
| SNP_A.K_996298_6_5981812 | 6 | 5981812 | 5.38E-12 | 0.09197 | 299 | 0.240980002 | 2.04E-07 |
| SNP_A.K_995373_6_5929348 | 6 | 5929348 | 5.53E-12 | 0.0903 | 299 | 0.240788867 | 2.05E-07 |
| SNP_A.K_994236_6_5832007 | 6 | 5832007 | 5.67E-12 | 0.08863 | 299 | 0.240610008 | 2.07E-07 |
| SNP_A.K_994445_6_5860919 | 6 | 5860919 | 6.03E-12 | 0.08696 | 299 | 0.240168265 | 2.08E-07 |
| SNP_A.K_998775_6_6221000 | 6 | 6221000 | 6.07E-12 | 0.0903 | 299 | 0.240112443 | 2.08E-07 |
| SNP_A.K_998730_6_6218360 | 6 | 6218360 | 6.08E-12 | 0.09699 | 299 | 0.240100918 | 2.08E-07 |
| SNP_A.K_996570_6_6010335 | 6 | 6010335 | 6.09E-12 | 0.09532 | 299 | 0.240091998 | 2.08E-07 |
| SNP_A.K_995555_6_5941693 | 6 | 5941693 | 6.26E-12 | 0.11706 | 299 | 0.239891556 | 2.08E-07 |
| SNP_A.K_994403_6_5856394 | 6 | 5856394 | 6.29E-12 | 0.07525 | 299 | 0.239864161 | 2.08E-07 |
| SNP_A.K_995905_6_5954735 | 6 | 5954735 | 6.41E-12 | 0.08863 | 299 | 0.239726703 | 2.08E-07 |
| SNP_A.K_1000556_6_6395708 | 6 | 6395708 | 6.71E-12 | 0.08863 | 299 | 0.239396072 | 2.15E-07 |
| SNP_A.K_994224_6_5827383 | 6 | 5827383 | 7.12E-12 | 0.12375 | 299 | 0.238966799 | 2.25E-07 |
| SNP_A.K_991499_6_5493250 | 6 | 5493250 | 7.71E-12 | 0.07023 | 299 | 0.238395362 | 2.40E-07 |
| SNP_A.K_993655_6_5781748 | 6 | 5781748 | 8.77E-12 | 0.08863 | 299 | 0.237470308 | 2.62E-07 |
| SNP_A.K_996001_6_5957294 | 6 | 5957294 | 8.79E-12 | 0.09197 | 299 | 0.237452137 | 2.62E-07 |
| SNP_A.K_994316_6_5843510 | 6 | 5843510 | 9.07E-12 | 0.0903 | 299 | 0.237231643 | 2.62E-07 |
| SNP_A.K_999138_6_6263245 | 6 | 6263245 | 9.14E-12 | 0.08863 | 299 | 0.237175337 | 2.62E-07 |
| SNP_A.K_995822_6_5951726 | 6 | 5951726 | 9.15E-12 | 0.0903 | 299 | 0.237168794 | 2.62E-07 |
| SNP_A.K_997873_6_6155185 | 6 | 6155185 | 9.19E-12 | 0.08696 | 299 | 0.237131859 | 2.62E-07 |
| SNP_A.K_998814_6_6226124 | 6 | 6226124 | 9.72E-12 | 0.0903 | 299 | 0.236729792 | 2.74E-07 |
| SNP_A.K_998777_6_6221103 | 6 | 6221103 | 1.01E-11 | 0.14381 | 299 | 0.236434101 | 2.81E-07 |
| SNP_A.K_999213_6_6267310 | 6 | 6267310 | 1.14E-11 | 0.0903 | 299 | 0.235578142 | 3.13E-07 |
| SNP_A.K_996673_6_6016337 | 6 | 6016337 | 1.21E-11 | 0.09699 | 299 | 0.235135119 | 3.28E-07 |
| SNP_A.K_998846_6_6228322 | 6 | 6228322 | 1.28E-11 | 0.08863 | 299 | 0.234781757 | 3.41E-07 |
| SNP_A.K_993974_6_5804805 | 6 | 5804805 | 1.44E-11 | 0.1204 | 299 | 0.23392584 | 3.79E-07 |
| SNP_A.K_1000919_6_6423748 | 6 | 6423748 | 1.47E-11 | 0.08194 | 299 | 0.23376088 | 3.79E-07 |
| SNP_A.K_999099_6_6257056 | 6 | 6257056 | 1.48E-11 | 0.08863 | 299 | 0.233744064 | 3.79E-07 |
| SNP_A.K_999370_6_6284657 | 6 | 6284657 | 1.50E-11 | 0.08863 | 299 | 0.233637342 | 3.80E-07 |
| SNP_A.K_994715_6_5881999 | 6 | 5881999 | 1.53E-11 | 0.13378 | 299 | 0.23346448 | 3.84E-07 |
| SNP_A.K_999106_6_6257205 | 6 | 6257205 | 1.55E-11 | 0.09866 | 299 | 0.233393737 | 3.84E-07 |
| SNP_A.K_995348_6_5927290 | 6 | 5927290 | 1.57E-11 | 0.08696 | 299 | 0.233293012 | 3.85E-07 |
| SNP_A.K_1000444_6_6385368 | 6 | 6385368 | 1.72E-11 | 0.0903 | 299 | 0.232657628 | 4.16E-07 |
| SNP_A.K_1000505_6_6389577 | 6 | 6389577 | 1.74E-11 | 0.08361 | 299 | 0.232551898 | 4.17E-07 |
| SNP_A.K_999059_6_6250399 | 6 | 6250399 | 1.86E-11 | 0.09699 | 299 | 0.232103415 | 4.31E-07 |
| SNP_A.K_994387_6_5854645 | 6 | 5854645 | 1.86E-11 | 0.12542 | 299 | 0.232080526 | 4.31E-07 |
| SNP_A.K_1364837_8_6048049 | 8 | 6048049 | 1.87E-11 | 0.03512 | 299 | 0.232070063 | 4.31E-07 |
| SNP_A.K_999301_6_6277647 | 6 | 6277647 | 1.94E-11 | 0.08863 | 299 | 0.231776913 | 4.40E-07 |
| SNP_A.K_996745_6_6021469 | 6 | 6021469 | 1.94E-11 | 0.09365 | 299 | 0.23177393 | 4.40E-07 |
| SNP_A.K_995650_6_5946309 | 6 | 5946309 | 2.02E-11 | 0.12375 | 299 | 0.231497891 | 4.48E-07 |
| SNP_A.K_999622_6_6301063 | 6 | 6301063 | 2.03E-11 | 0.0903 | 299 | 0.231479213 | 4.48E-07 |
| SNP_A.K_993680_6_5784371 | 6 | 5784371 | 2.04E-11 | 0.09365 | 299 | 0.231415368 | 4.48E-07 |

| | | | | | | | |
|---|---|---|---|---|---|---|---|
| SNP_A.K_998940_6_6236471 | 6 | 6236471 | 2.17E-11 | 0.08194 | 299 | 0.231007352 | 4.69E-07 |
| SNP_A.K_997927_6_6159890 | 6 | 6159890 | 2.37E-11 | 0.09532 | 299 | 0.230365951 | 5.08E-07 |
| SNP_A.K_994378_6_5854002 | 6 | 5854002 | 2.40E-11 | 0.12375 | 299 | 0.23026806 | 5.10E-07 |
| SNP_A.K_999365_6_6284017 | 6 | 6284017 | 2.49E-11 | 0.09197 | 299 | 0.229998023 | 5.25E-07 |
| SNP_A.K_994475_6_5865633 | 6 | 5865633 | 2.61E-11 | 0.08361 | 299 | 0.229684479 | 5.43E-07 |
| SNP_A.K_995716_6_5948813 | 6 | 5948813 | 2.64E-11 | 0.09699 | 299 | 0.229605785 | 5.43E-07 |
| SNP_A.K_994749_6_5886035 | 6 | 5886035 | 2.66E-11 | 0.0903 | 299 | 0.229532749 | 5.43E-07 |
| SNP_A.K_994765_6_5888082 | 6 | 5888082 | 2.74E-11 | 0.10535 | 299 | 0.229321875 | 5.54E-07 |
| SNP_A.K_996782_6_6025389 | 6 | 6025389 | 2.80E-11 | 0.11037 | 299 | 0.229168538 | 5.61E-07 |
| SNP_A.K_995699_6_5948118 | 6 | 5948118 | 3.05E-11 | 0.08528 | 299 | 0.228578471 | 6.04E-07 |
| SNP_A.K_994401_6_5856335 | 6 | 5856335 | 3.12E-11 | 0.11037 | 299 | 0.228396763 | 6.08E-07 |
| SNP_A.K_999831_6_6327455 | 6 | 6327455 | 3.13E-11 | 0.08361 | 299 | 0.22837566 | 6.08E-07 |
| SNP_A.K_999103_6_6257135 | 6 | 6257135 | 3.16E-11 | 0.09866 | 299 | 0.22832674 | 6.08E-07 |
| SNP_A.K_993851_6_5796742 | 6 | 5796742 | 3.32E-11 | 0.12876 | 299 | 0.227962433 | 6.34E-07 |
| SNP_A.K_999348_6_6283162 | 6 | 6283162 | 3.35E-11 | 0.08361 | 299 | 0.227905489 | 6.34E-07 |
| SNP_A.K_995268_6_5923572 | 6 | 5923572 | 3.42E-11 | 0.09197 | 299 | 0.227752264 | 6.42E-07 |
| SNP_A.K_999191_6_6265220 | 6 | 6265220 | 3.53E-11 | 0.09365 | 299 | 0.227536371 | 6.54E-07 |
| SNP_A.K_996328_6_5986872 | 6 | 5986872 | 3.55E-11 | 0.09197 | 299 | 0.227491109 | 6.54E-07 |
| SNP_A.K_999730_6_6311365 | 6 | 6311365 | 3.73E-11 | 0.08528 | 299 | 0.227140086 | 6.73E-07 |
| SNP_A.K_998770_6_6220481 | 6 | 6220481 | 3.78E-11 | 0.08863 | 299 | 0.227047726 | 6.73E-07 |
| SNP_A.K_999683_6_6307536 | 6 | 6307536 | 3.78E-11 | 0.15385 | 299 | 0.227037986 | 6.73E-07 |
| SNP_A.K_996157_6_5970476 | 6 | 5970476 | 3.78E-11 | 0.09365 | 299 | 0.227036985 | 6.73E-07 |
| SNP_A.K_998794_6_6223638 | 6 | 6223638 | 3.93E-11 | 0.08863 | 299 | 0.226773695 | 6.81E-07 |
| SNP_A.K_999636_6_6301739 | 6 | 6301739 | 3.93E-11 | 0.0903 | 299 | 0.226767303 | 6.81E-07 |
| SNP_A.K_999110_6_6257284 | 6 | 6257284 | 3.97E-11 | 0.08696 | 299 | 0.226686742 | 6.81E-07 |
| SNP_A.K_999352_6_6283316 | 6 | 6283316 | 4.00E-11 | 0.09365 | 299 | 0.226647315 | 6.81E-07 |
| SNP_A.K_997171_6_6054645 | 6 | 6054645 | 4.01E-11 | 0.09197 | 299 | 0.226618588 | 6.81E-07 |
| SNP_A.K_998894_6_6231763 | 6 | 6231763 | 4.03E-11 | 0.0903 | 299 | 0.226595876 | 6.81E-07 |
| SNP_A.K_997686_6_6142451 | 6 | 6142451 | 4.12E-11 | 0.09866 | 299 | 0.226424241 | 6.90E-07 |
| SNP_A.K_994138_6_5817719 | 6 | 5817719 | 4.16E-11 | 0.11371 | 299 | 0.22636789 | 6.90E-07 |
| SNP_A.K_995379_6_5930135 | 6 | 5930135 | 4.18E-11 | 0.08528 | 299 | 0.226330728 | 6.90E-07 |
| SNP_A.K_998953_6_6240098 | 6 | 6240098 | 4.37E-11 | 0.08361 | 299 | 0.226022593 | 7.06E-07 |
| SNP_A.K_999349_6_6283176 | 6 | 6283176 | 4.37E-11 | 0.08361 | 299 | 0.226022593 | 7.06E-07 |
| SNP_A.K_1000295_6_6369047 | 6 | 6369047 | 4.38E-11 | 0.10368 | 299 | 0.226000895 | 7.06E-07 |
| SNP_A.K_994339_6_5847479 | 6 | 5847479 | 4.41E-11 | 0.08863 | 299 | 0.225948014 | 7.06E-07 |
| SNP_A.K_999672_6_6306686 | 6 | 6306686 | 4.53E-11 | 0.09699 | 299 | 0.225763641 | 7.13E-07 |
| SNP_A.K_995341_6_5926818 | 6 | 5926818 | 4.57E-11 | 0.09365 | 299 | 0.225703858 | 7.13E-07 |
| SNP_A.K_999241_6_6270064 | 6 | 6270064 | 4.57E-11 | 0.11873 | 299 | 0.225689748 | 7.13E-07 |
| SNP_A.K_999594_6_6299163 | 6 | 6299163 | 4.59E-11 | 0.08361 | 299 | 0.225664908 | 7.13E-07 |
| SNP_A.K_996367_6_5994895 | 6 | 5994895 | 4.72E-11 | 0.09365 | 299 | 0.225475195 | 7.27E-07 |
| SNP_A.K_996141_6_5964541 | 6 | 5964541 | 4.92E-11 | 0.09532 | 299 | 0.2251688 | 7.53E-07 |
| SNP_A.K_993961_6_5802562 | 6 | 5802562 | 5.02E-11 | 0.13211 | 299 | 0.225028614 | 7.53E-07 |
| SNP_A.K_993172_6_5712012 | 6 | 5712012 | 5.04E-11 | 0.11204 | 299 | 0.225006284 | 7.53E-07 |
| SNP_A.K_1002990_6_6640344 | 6 | 6640344 | 5.06E-11 | 0.08194 | 299 | 0.224974207 | 7.53E-07 |
| SNP_A.K_999276_6_6276073 | 6 | 6276073 | 5.10E-11 | 0.11538 | 299 | 0.224919978 | 7.53E-07 |
| SNP_A.K_999417_6_6287437 | 6 | 6287437 | 5.10E-11 | 0.08696 | 299 | 0.224919155 | 7.53E-07 |
| SNP_A.K_1364828_8_6046714 | 8 | 6046714 | 5.22E-11 | 0.04013 | 299 | 0.224755728 | 7.65E-07 |
| SNP_A.K_999468_6_6291568 | 6 | 6291568 | 5.26E-11 | 0.08528 | 299 | 0.22470558 | 7.65E-07 |
| SNP_A.K_998792_6_6222593 | 6 | 6222593 | 5.50E-11 | 0.10535 | 299 | 0.224386766 | 7.95E-07 |
| SNP_A.K_999407_6_6286564 | 6 | 6286564 | 5.55E-11 | 0.10201 | 299 | 0.224323477 | 7.96E-07 |

| | | | | | | | |
|---|---|---|---|---|---|---|---|
| SNP_A.K_996779_6_6025241 | 6 | 6025241 | 5.72E-11 | 0.10201 | 299 | 0.22410968 | 8.15E-07 |
| SNP_A.K_998957_6_6240837 | 6 | 6240837 | 5.76E-11 | 0.08027 | 299 | 0.224064111 | 8.15E-07 |
| SNP_A.K_994235_6_5831947 | 6 | 5831947 | 5.89E-11 | 0.08361 | 299 | 0.223900003 | 8.27E-07 |
| SNP_A.K_993991_6_5806864 | 6 | 5806864 | 5.92E-11 | 0.08863 | 299 | 0.223866352 | 8.27E-07 |
| SNP_A.K_995380_6_5930170 | 6 | 5930170 | 6.06E-11 | 0.08696 | 299 | 0.223698089 | 8.41E-07 |
| SNP_A.K_996285_6_5981490 | 6 | 5981490 | 6.20E-11 | 0.09532 | 299 | 0.223539852 | 8.54E-07 |
| SNP_A.K_995721_6_5948853 | 6 | 5948853 | 6.40E-11 | 0.09365 | 299 | 0.223318382 | 8.76E-07 |
| SNP_A.K_998913_6_6234067 | 6 | 6234067 | 6.66E-11 | 0.08194 | 299 | 0.22302949 | 9.06E-07 |
| SNP_A.K_1000338_6_6376429 | 6 | 6376429 | 7.09E-11 | 0.08528 | 299 | 0.222594845 | 9.58E-07 |
| SNP_A.K_325679_2_9921824 | 2 | 9921824 | 7.28E-11 | 0.0301 | 299 | 0.222405272 | 9.77E-07 |
| SNP_A.K_994998_6_5912028 | 6 | 5912028 | 7.42E-11 | 0.09365 | 299 | 0.222264471 | 9.91E-07 |
| SNP_A.K_1000523_6_6391183 | 6 | 6391183 | 7.86E-11 | 0.09866 | 299 | 0.221857799 | 1.04E-06 |
| SNP_A.K_328877_2_10321602 | 2 | 10321602 | 8.43E-11 | 0.01505 | 299 | 0.221371511 | 1.11E-06 |
| SNP_A.K_995384_6_5931170 | 6 | 5931170 | 8.68E-11 | 0.09866 | 299 | 0.221158898 | 1.13E-06 |
| SNP_A.K_1000506_6_6389604 | 6 | 6389604 | 8.71E-11 | 0.08696 | 299 | 0.221136996 | 1.13E-06 |
| SNP_A.K_998773_6_6220832 | 6 | 6220832 | 8.86E-11 | 0.09197 | 299 | 0.221013146 | 1.15E-06 |
| SNP_A.K_992488_6_5652615 | 6 | 5652615 | 8.94E-11 | 0.10535 | 299 | 0.22095153 | 1.15E-06 |
| SNP_A.K_1000225_6_6357159 | 6 | 6357159 | 9.64E-11 | 0.1087 | 299 | 0.220420623 | 1.22E-06 |
| SNP_A.K_325574_2_9907613 | 2 | 9907613 | 9.66E-11 | 0.02174 | 299 | 0.22040899 | 1.22E-06 |
| SNP_A.K_325662_2_9921207 | 2 | 9921207 | 9.66E-11 | 0.02174 | 299 | 0.22040899 | 1.22E-06 |
| SNP_A.K_996301_6_5981935 | 6 | 5981935 | 9.77E-11 | 0.09532 | 299 | 0.220325849 | 1.23E-06 |
| SNP_A.K_998232_6_6185754 | 6 | 6185754 | 9.93E-11 | 0.10033 | 299 | 0.22021643 | 1.24E-06 |
| SNP_A.K_998367_6_6191313 | 6 | 6191313 | 1.04E-10 | 0.09532 | 299 | 0.219886147 | 1.29E-06 |
| SNP_A.K_999668_6_6304733 | 6 | 6304733 | 1.05E-10 | 0.09866 | 299 | 0.219817076 | 1.29E-06 |
| SNP_A.K_999379_6_6285385 | 6 | 6285385 | 1.05E-10 | 0.08863 | 299 | 0.219800502 | 1.29E-06 |
| SNP_A.K_997784_6_6146505 | 6 | 6146505 | 1.06E-10 | 0.10033 | 299 | 0.219751378 | 1.29E-06 |
| SNP_A.K_1000246_6_6358278 | 6 | 6358278 | 1.07E-10 | 0.09699 | 299 | 0.219672564 | 1.30E-06 |
| SNP_A.K_999823_6_6327391 | 6 | 6327391 | 1.15E-10 | 0.08528 | 299 | 0.219209473 | 1.37E-06 |
| SNP_A.K_999457_6_6291033 | 6 | 6291033 | 1.15E-10 | 0.0903 | 299 | 0.219201928 | 1.37E-06 |
| SNP_A.K_997161_6_6051871 | 6 | 6051871 | 1.15E-10 | 0.10535 | 299 | 0.219174735 | 1.37E-06 |
| SNP_A.K_999735_6_6311754 | 6 | 6311754 | 1.16E-10 | 0.08863 | 299 | 0.219131293 | 1.37E-06 |
| SNP_A.K_995977_6_5956567 | 6 | 5956567 | 1.18E-10 | 0.08863 | 299 | 0.218991585 | 1.39E-06 |
| SNP_A.K_994226_6_5827705 | 6 | 5827705 | 1.22E-10 | 0.12207 | 299 | 0.218749157 | 1.43E-06 |
| SNP_A.K_998204_6_6184534 | 6 | 6184534 | 1.25E-10 | 0.09699 | 299 | 0.218612866 | 1.45E-06 |
| SNP_A.K_995479_6_5938571 | 6 | 5938571 | 1.25E-10 | 0.08863 | 299 | 0.218575758 | 1.45E-06 |
| SNP_A.K_994683_6_5880054 | 6 | 5880054 | 1.26E-10 | 0.08696 | 299 | 0.218519307 | 1.45E-06 |
| SNP_A.K_1000178_6_6351662 | 6 | 6351662 | 1.30E-10 | 0.08863 | 299 | 0.218297304 | 1.49E-06 |
| SNP_A.K_999547_6_6295268 | 6 | 6295268 | 1.31E-10 | 0.09699 | 299 | 0.21827985 | 1.49E-06 |
| SNP_A.K_997330_6_6088530 | 6 | 6088530 | 1.32E-10 | 0.09866 | 299 | 0.218212941 | 1.49E-06 |
| SNP_A.K_999921_6_6333270 | 6 | 6333270 | 1.32E-10 | 0.12876 | 299 | 0.218188464 | 1.49E-06 |
| SNP_A.K_999600_6_6299479 | 6 | 6299479 | 1.35E-10 | 0.08528 | 299 | 0.218050871 | 1.51E-06 |
| SNP_A.K_995786_6_5950706 | 6 | 5950706 | 1.38E-10 | 0.08696 | 299 | 0.217906507 | 1.53E-06 |
| SNP_A.K_999446_6_6290665 | 6 | 6290665 | 1.43E-10 | 0.08863 | 299 | 0.217645403 | 1.58E-06 |
| SNP_A.K_994160_6_5820762 | 6 | 5820762 | 1.45E-10 | 0.11706 | 299 | 0.217555703 | 1.58E-06 |
| SNP_A.K_998140_6_6177028 | 6 | 6177028 | 1.45E-10 | 0.09866 | 299 | 0.217547135 | 1.58E-06 |
| SNP_A.K_993967_6_5804373 | 6 | 5804373 | 1.46E-10 | 0.11371 | 299 | 0.217503734 | 1.58E-06 |
| SNP_A.K_999719_6_6308998 | 6 | 6308998 | 1.46E-10 | 0.08863 | 299 | 0.217500778 | 1.58E-06 |
| SNP_A.K_998346_6_6190557 | 6 | 6190557 | 1.47E-10 | 0.09699 | 299 | 0.217443201 | 1.59E-06 |
| SNP_A.K_997674_6_6142130 | 6 | 6142130 | 1.51E-10 | 0.09866 | 299 | 0.217284678 | 1.62E-06 |
| SNP_A.K_999200_6_6265562 | 6 | 6265562 | 1.53E-10 | 0.09197 | 299 | 0.217159067 | 1.64E-06 |

| | | | | | | | |
|---|---|---|---|---|---|---|---|
| SNP_A.K_999930_6_6333766 | 6 | 6333766 | 1.64E-10 | 0.12709 | 299 | 0.2167033 | 1.74E-06 |
| SNP_A.K_999506_6_6293835 | 6 | 6293835 | 1.69E-10 | 0.08696 | 299 | 0.216477565 | 1.78E-06 |
| SNP_A.K_1000567_6_6396433 | 6 | 6396433 | 1.71E-10 | 0.08194 | 299 | 0.216390543 | 1.80E-06 |
| SNP_A.K_1000288_6_6368327 | 6 | 6368327 | 1.82E-10 | 0.08863 | 299 | 0.215937673 | 1.91E-06 |
| SNP_A.K_1002807_6_6620235 | 6 | 6620235 | 2.02E-10 | 0.11037 | 299 | 0.215222312 | 2.10E-06 |
| SNP_A.K_994230_6_5830491 | 6 | 5830491 | 2.06E-10 | 0.12709 | 299 | 0.215087771 | 2.13E-06 |
| SNP_A.K_999763_6_6312704 | 6 | 6312704 | 2.20E-10 | 0.11204 | 299 | 0.214621477 | 2.27E-06 |
| SNP_A.K_1000276_6_6367416 | 6 | 6367416 | 2.32E-10 | 0.08863 | 299 | 0.214258931 | 2.38E-06 |
| SNP_A.K_325606_2_9916262 | 2 | 9916262 | 2.37E-10 | 0.02341 | 299 | 0.214110679 | 2.42E-06 |
| SNP_A.K_994168_6_5821410 | 6 | 5821410 | 2.40E-10 | 0.11371 | 299 | 0.214002085 | 2.44E-06 |
| SNP_A.K_996670_6_6016112 | 6 | 6016112 | 2.47E-10 | 0.09197 | 299 | 0.213824657 | 2.49E-06 |
| SNP_A.K_999424_6_6287793 | 6 | 6287793 | 2.81E-10 | 0.08696 | 299 | 0.212914965 | 2.82E-06 |
| SNP_A.K_996853_6_6030903 | 6 | 6030903 | 2.82E-10 | 0.08361 | 299 | 0.212899028 | 2.82E-06 |
| SNP_A.K_998664_6_6215959 | 6 | 6215959 | 2.88E-10 | 0.08863 | 299 | 0.212748261 | 2.87E-06 |
| SNP_A.K_997774_6_6146123 | 6 | 6146123 | 2.97E-10 | 0.10033 | 299 | 0.212535703 | 2.94E-06 |
| SNP_A.K_999039_6_6249543 | 6 | 6249543 | 3.06E-10 | 0.08863 | 299 | 0.212308196 | 3.02E-06 |
| SNP_A.K_994895_6_5900108 | 6 | 5900108 | 3.08E-10 | 0.08863 | 299 | 0.212269665 | 3.03E-06 |
| SNP_A.K_999697_6_6308164 | 6 | 6308164 | 3.11E-10 | 0.09866 | 299 | 0.212198968 | 3.04E-06 |
| SNP_A.K_325944_2_9963856 | 2 | 9963856 | 3.37E-10 | 0.02341 | 299 | 0.211637843 | 3.28E-06 |
| SNP_A.K_996162_6_5970569 | 6 | 5970569 | 3.66E-10 | 0.09866 | 299 | 0.211064866 | 3.55E-06 |
| SNP_A.K_995055_6_5914207 | 6 | 5914207 | 3.70E-10 | 0.0903 | 299 | 0.210990231 | 3.57E-06 |
| SNP_A.K_996665_6_6015908 | 6 | 6015908 | 3.74E-10 | 0.10368 | 299 | 0.210908237 | 3.59E-06 |
| SNP_A.K_998861_6_6229232 | 6 | 6229232 | 3.83E-10 | 0.09197 | 299 | 0.210749 | 3.66E-06 |
| SNP_A.K_1000595_6_6398937 | 6 | 6398937 | 3.94E-10 | 0.08696 | 299 | 0.210557665 | 3.74E-06 |
| SNP_A.K_996458_6_6002348 | 6 | 6002348 | 3.95E-10 | 0.10368 | 299 | 0.210527993 | 3.74E-06 |
| SNP_A.K_994340_6_5847625 | 6 | 5847625 | 3.99E-10 | 0.12207 | 299 | 0.210474658 | 3.75E-06 |
| SNP_A.K_999498_6_6293602 | 6 | 6293602 | 4.12E-10 | 0.08194 | 299 | 0.210239046 | 3.86E-06 |
| SNP_A.K_1001102_6_6442110 | 6 | 6442110 | 4.14E-10 | 0.12207 | 299 | 0.210214009 | 3.86E-06 |
| SNP_A.K_993757_6_5790986 | 6 | 5790986 | 4.16E-10 | 0.16555 | 299 | 0.210167446 | 3.87E-06 |
| SNP_A.K_994174_6_5821928 | 6 | 5821928 | 4.20E-10 | 0.11873 | 299 | 0.210116707 | 3.88E-06 |
| SNP_A.K_999214_6_6267616 | 6 | 6267616 | 4.21E-10 | 0.08027 | 299 | 0.210089434 | 3.88E-06 |
| SNP_A.K_998025_6_6166760 | 6 | 6166760 | 4.25E-10 | 0.10201 | 299 | 0.210021393 | 3.90E-06 |
| SNP_A.K_328398_2_10257252 | 2 | 10257252 | 4.42E-10 | 0.02174 | 299 | 0.209747133 | 4.04E-06 |
| SNP_A.K_993783_6_5793279 | 6 | 5793279 | 4.57E-10 | 0.08027 | 299 | 0.209524008 | 4.15E-06 |
| SNP_A.K_995342_6_5926823 | 6 | 5926823 | 4.59E-10 | 0.09197 | 299 | 0.209488605 | 4.15E-06 |
| SNP_A.K_999199_6_6265468 | 6 | 6265468 | 4.74E-10 | 0.08863 | 299 | 0.209266037 | 4.27E-06 |
| SNP_A.K_993618_6_5779707 | 6 | 5779707 | 5.13E-10 | 0.09532 | 299 | 0.208718551 | 4.60E-06 |
| SNP_A.K_999631_6_6301335 | 6 | 6301335 | 5.26E-10 | 0.09197 | 299 | 0.208543755 | 4.68E-06 |
| SNP_A.K_994858_6_5895786 | 6 | 5895786 | 5.26E-10 | 0.09699 | 299 | 0.208542749 | 4.68E-06 |
| SNP_A.K_999627_6_6301282 | 6 | 6301282 | 5.48E-10 | 0.08361 | 299 | 0.208259434 | 4.85E-06 |
| SNP_A.K_996134_6_5964453 | 6 | 5964453 | 5.93E-10 | 0.09197 | 299 | 0.207710447 | 5.23E-06 |
| SNP_A.K_997231_6_6058815 | 6 | 6058815 | 5.97E-10 | 0.10368 | 299 | 0.207664277 | 5.24E-06 |
| SNP_A.K_993865_6_5797885 | 6 | 5797885 | 6.10E-10 | 0.12709 | 299 | 0.207516825 | 5.32E-06 |
| SNP_A.K_998155_6_6179298 | 6 | 6179298 | 6.11E-10 | 0.09365 | 299 | 0.207499297 | 5.32E-06 |
| SNP_A.K_997217_6_6058367 | 6 | 6058367 | 6.41E-10 | 0.10033 | 299 | 0.207172494 | 5.56E-06 |
| SNP_A.K_994724_6_5883300 | 6 | 5883300 | 6.89E-10 | 0.1505 | 299 | 0.206669915 | 5.91E-06 |
| SNP_A.K_997319_6_6086995 | 6 | 6086995 | 6.89E-10 | 0.11204 | 299 | 0.206665577 | 5.91E-06 |
| SNP_A.K_992846_6_5681254 | 6 | 5681254 | 6.90E-10 | 0.11706 | 299 | 0.206655641 | 5.91E-06 |
| SNP_A.K_999855_6_6328477 | 6 | 6328477 | 6.93E-10 | 0.10368 | 299 | 0.206625551 | 5.91E-06 |
| SNP_A.K_998121_6_6174048 | 6 | 6174048 | 7.00E-10 | 0.10702 | 299 | 0.206559968 | 5.94E-06 |

| | | | | | | | |
|---|---|---|---|---|---|---|---|
| SNP_A.K_993063_6_5702831 | 6 | 5702831 | 7.21E-10 | 0.11538 | 299 | 0.206354677 | 6.10E-06 |
| SNP_A.K_1002263_6_6558482 | 6 | 6558482 | 7.25E-10 | 0.0903 | 299 | 0.206315658 | 6.11E-06 |
| SNP_A.K_996716_6_6018571 | 6 | 6018571 | 7.30E-10 | 0.09365 | 299 | 0.206264403 | 6.13E-06 |
| SNP_A.K_999432_6_6288177 | 6 | 6288177 | 7.33E-10 | 0.13712 | 299 | 0.206237196 | 6.13E-06 |
| SNP_A.K_2080292_12_27255011 | 12 | 27255011 | 7.41E-10 | 0.01003 | 299 | 0.206165269 | 6.17E-06 |
| SNP_A.K_1364824_8_6045628 | 8 | 6045628 | 7.45E-10 | 0.04348 | 299 | 0.206120314 | 6.18E-06 |
| SNP_A.K_999136_6_6263123 | 6 | 6263123 | 7.72E-10 | 0.09866 | 299 | 0.205880757 | 6.37E-06 |
| SNP_A.K_327043_2_10084832 | 2 | 10084832 | 8.22E-10 | 0.03512 | 299 | 0.205439283 | 6.77E-06 |
| SNP_A.K_994389_6_5854738 | 6 | 5854738 | 8.51E-10 | 0.12542 | 299 | 0.205205447 | 6.97E-06 |
| SNP_A.K_993111_6_5708648 | 6 | 5708648 | 8.55E-10 | 0.10702 | 299 | 0.205166015 | 6.98E-06 |
| SNP_A.K_996551_6_6008529 | 6 | 6008529 | 8.71E-10 | 0.09365 | 299 | 0.205037791 | 7.09E-06 |
| SNP_A.K_998056_6_6167657 | 6 | 6167657 | 8.80E-10 | 0.10033 | 299 | 0.204968785 | 7.13E-06 |
| SNP_A.K_998362_6_6190837 | 6 | 6190837 | 8.88E-10 | 0.09197 | 299 | 0.204908848 | 7.16E-06 |
| SNP_A.K_999270_6_6275636 | 6 | 6275636 | 9.16E-10 | 0.14381 | 299 | 0.204695269 | 7.36E-06 |
| SNP_A.K_994499_6_5869880 | 6 | 5869880 | 9.40E-10 | 0.1505 | 299 | 0.204513927 | 7.52E-06 |
| SNP_A.K_325590_2_9913723 | 2 | 9913723 | 9.52E-10 | 0.02508 | 299 | 0.204425813 | 7.57E-06 |
| SNP_A.K_998120_6_6173570 | 6 | 6173570 | 9.53E-10 | 0.09197 | 299 | 0.204415299 | 7.57E-06 |
| SNP_A.K_993062_6_5702799 | 6 | 5702799 | 9.73E-10 | 0.11873 | 299 | 0.204276579 | 7.70E-06 |
| SNP_A.K_326998_2_10078971 | 2 | 10078971 | 9.82E-10 | 0.02508 | 299 | 0.204210602 | 7.74E-06 |
| SNP_A.K_993364_6_5742301 | 6 | 5742301 | 9.89E-10 | 0.11706 | 299 | 0.204159144 | 7.77E-06 |
| SNP_A.K_999456_6_6291030 | 6 | 6291030 | 9.93E-10 | 0.08696 | 299 | 0.204132332 | 7.77E-06 |
| SNP_A.K_998159_6_6180389 | 6 | 6180389 | 1.09E-09 | 0.09532 | 299 | 0.203481086 | 8.50E-06 |
| SNP_A.K_997168_6_6053127 | 6 | 6053127 | 1.09E-09 | 0.09699 | 299 | 0.20346415 | 8.50E-06 |
| SNP_A.K_1198450_7_7870036 | 7 | 7870036 | 1.14E-09 | 0.01338 | 299 | 0.203176178 | 8.82E-06 |
| SNP_A.K_307510_2_7413964 | 2 | 7413964 | 1.15E-09 | 0.23913 | 299 | 0.203114333 | 8.87E-06 |
| SNP_A.K_993978_6_5805283 | 6 | 5805283 | 1.18E-09 | 0.12375 | 299 | 0.202963351 | 9.03E-06 |
| SNP_A.K_999628_6_6301297 | 6 | 6301297 | 1.19E-09 | 0.09365 | 299 | 0.202887943 | 9.10E-06 |
| SNP_A.K_998778_6_6221159 | 6 | 6221159 | 1.24E-09 | 0.13545 | 299 | 0.202599278 | 9.45E-06 |
| SNP_A.K_995371_6_5929105 | 6 | 5929105 | 1.30E-09 | 0.10033 | 299 | 0.202263331 | 9.89E-06 |
| SNP_A.K_996441_6_6001052 | 6 | 6001052 | 1.31E-09 | 0.09365 | 299 | 0.202239043 | 9.89E-06 |
| SNP_A.K_497857_3_5598569 | 3 | 5598569 | 1.34E-09 | 0.01338 | 299 | 0.202071352 | 1.01E-05 |
| SNP_A.K_995793_6_5950773 | 6 | 5950773 | 1.39E-09 | 0.12709 | 299 | 0.201825059 | 1.04E-05 |
| SNP_A.K_994761_6_5887861 | 6 | 5887861 | 1.39E-09 | 0.10033 | 299 | 0.201803684 | 1.04E-05 |
| SNP_A.K_992745_6_5678115 | 6 | 5678115 | 1.44E-09 | 0.11204 | 299 | 0.201541388 | 1.08E-05 |
| SNP_A.K_978593_6_1736459 | 6 | 1736459 | 1.50E-09 | 0.11873 | 299 | 0.201273405 | 1.12E-05 |
| SNP_A.K_1000603_6_6399151 | 6 | 6399151 | 1.53E-09 | 0.10033 | 299 | 0.201153436 | 1.13E-05 |
| SNP_A.K_993188_6_5716112 | 6 | 5716112 | 1.53E-09 | 0.11204 | 299 | 0.201141491 | 1.13E-05 |
| SNP_A.K_998831_6_6226827 | 6 | 6226827 | 1.57E-09 | 0.08361 | 299 | 0.200985173 | 1.15E-05 |
| SNP_A.K_1364816_8_6043498 | 8 | 6043498 | 1.59E-09 | 0.05017 | 299 | 0.200864649 | 1.16E-05 |
| SNP_A.K_999648_6_6302685 | 6 | 6302685 | 1.60E-09 | 0.09365 | 299 | 0.200855158 | 1.16E-05 |
| SNP_A.K_999113_6_6257528 | 6 | 6257528 | 1.60E-09 | 0.10368 | 299 | 0.200847193 | 1.16E-05 |
| SNP_A.K_1002710_6_6612229 | 6 | 6612229 | 1.65E-09 | 0.11706 | 299 | 0.200622778 | 1.20E-05 |
| SNP_A.K_999434_6_6288270 | 6 | 6288270 | 1.66E-09 | 0.15552 | 299 | 0.200573799 | 1.20E-05 |
| SNP_A.K_992450_6_5647061 | 6 | 5647061 | 1.67E-09 | 0.12207 | 299 | 0.200526506 | 1.21E-05 |
| SNP_A.K_995642_6_5945927 | 6 | 5945927 | 1.70E-09 | 0.14047 | 299 | 0.200438939 | 1.22E-05 |
| SNP_A.K_1000548_6_6393688 | 6 | 6393688 | 1.78E-09 | 0.10201 | 299 | 0.200120297 | 1.27E-05 |
| SNP_A.K_993238_6_5723513 | 6 | 5723513 | 1.88E-09 | 0.11873 | 299 | 0.199738343 | 1.34E-05 |
| SNP_A.K_993349_6_5740837 | 6 | 5740837 | 1.91E-09 | 0.11706 | 299 | 0.199604959 | 1.36E-05 |
| SNP_A.K_1000003_6_6340934 | 6 | 6340934 | 1.93E-09 | 0.13043 | 299 | 0.199533706 | 1.37E-05 |
| SNP_A.K_999188_6_6265147 | 6 | 6265147 | 1.95E-09 | 0.09532 | 299 | 0.199476845 | 1.37E-05 |

| | | | | | | | |
|---|---|---|---|---|---|---|---|
| SNP_A.K_997841_6_6151654 | 6 | 6151654 | 1.95E-09 | 0.09197 | 299 | 0.199473173 | 1.37E-05 |
| SNP_A.K_999398_6_6286418 | 6 | 6286418 | 2.07E-09 | 0.12207 | 299 | 0.199065592 | 1.45E-05 |
| SNP_A.K_992715_6_5675706 | 6 | 5675706 | 2.11E-09 | 0.12207 | 299 | 0.198946769 | 1.47E-05 |
| SNP_A.K_998348_6_6190678 | 6 | 6190678 | 2.15E-09 | 0.0903 | 299 | 0.198799538 | 1.50E-05 |
| SNP_A.K_998882_6_6230362 | 6 | 6230362 | 2.22E-09 | 0.1087 | 299 | 0.198568743 | 1.54E-05 |
| SNP_A.K_993059_6_5702626 | 6 | 5702626 | 2.24E-09 | 0.11873 | 299 | 0.198524494 | 1.55E-05 |
| SNP_A.K_998080_6_6170102 | 6 | 6170102 | 2.32E-09 | 0.10201 | 299 | 0.198291061 | 1.60E-05 |
| SNP_A.K_994910_6_5905982 | 6 | 5905982 | 2.33E-09 | 0.10033 | 299 | 0.198237357 | 1.60E-05 |
| SNP_A.K_994650_6_5877851 | 6 | 5877851 | 2.39E-09 | 0.08863 | 299 | 0.198076888 | 1.64E-05 |
| SNP_A.K_993130_6_5709698 | 6 | 5709698 | 2.41E-09 | 0.13211 | 299 | 0.198026611 | 1.64E-05 |
| SNP_A.K_997576_6_6109635 | 6 | 6109635 | 2.50E-09 | 0.09866 | 299 | 0.197777197 | 1.70E-05 |
| SNP_A.K_995844_6_5952431 | 6 | 5952431 | 2.52E-09 | 0.1689 | 299 | 0.197721212 | 1.71E-05 |
| SNP_A.K_998106_6_6171179 | 6 | 6171179 | 2.53E-09 | 0.09699 | 299 | 0.197671522 | 1.71E-05 |
| SNP_A.K_1599657_9_18583878 | 9 | 18583878 | 2.56E-09 | 0.03512 | 299 | 0.197599639 | 1.72E-05 |
| SNP_A.K_997667_6_6141730 | 6 | 6141730 | 2.57E-09 | 0.09699 | 299 | 0.197571555 | 1.72E-05 |
| SNP_A.K_996232_6_5976271 | 6 | 5976271 | 2.57E-09 | 0.09866 | 299 | 0.197569264 | 1.72E-05 |
| SNP_A.K_1004738_6_6926541 | 6 | 6926541 | 2.63E-09 | 0.11204 | 299 | 0.197420218 | 1.75E-05 |
| SNP_A.K_993296_6_5734229 | 6 | 5734229 | 2.65E-09 | 0.11204 | 299 | 0.197356248 | 1.76E-05 |
| SNP_A.K_996493_6_6003854 | 6 | 6003854 | 2.65E-09 | 0.09699 | 299 | 0.197355254 | 1.76E-05 |
| SNP_A.K_999637_6_6301795 | 6 | 6301795 | 2.73E-09 | 0.09699 | 299 | 0.197163784 | 1.80E-05 |
| SNP_A.K_994617_6_5876549 | 6 | 5876549 | 2.77E-09 | 0.16722 | 299 | 0.197066837 | 1.82E-05 |
| SNP_A.K_994205_6_5825330 | 6 | 5825330 | 2.78E-09 | 0.10201 | 299 | 0.197030353 | 1.83E-05 |
| SNP_A.K_998776_6_6221062 | 6 | 6221062 | 2.88E-09 | 0.10702 | 299 | 0.1967979 | 1.88E-05 |
| SNP_A.K_995381_6_5930184 | 6 | 5930184 | 2.90E-09 | 0.08696 | 299 | 0.196744383 | 1.89E-05 |
| SNP_A.K_993985_6_5806596 | 6 | 5806596 | 2.95E-09 | 0.12207 | 299 | 0.196634612 | 1.92E-05 |
| SNP_A.K_997932_6_6160193 | 6 | 6160193 | 3.01E-09 | 0.09866 | 299 | 0.196501591 | 1.95E-05 |
| SNP_A.K_994830_6_5893166 | 6 | 5893166 | 3.05E-09 | 0.12207 | 299 | 0.196409747 | 1.97E-05 |
| SNP_A.K_998444_6_6199199 | 6 | 6199199 | 3.14E-09 | 0.18729 | 299 | 0.196209278 | 2.02E-05 |
| SNP_A.K_999717_6_6308732 | 6 | 6308732 | 3.35E-09 | 0.10535 | 299 | 0.19576584 | 2.15E-05 |
| SNP_A.K_1000145_6_6348397 | 6 | 6348397 | 3.37E-09 | 0.14716 | 299 | 0.195724509 | 2.15E-05 |
| SNP_A.K_997943_6_6161115 | 6 | 6161115 | 3.37E-09 | 0.10033 | 299 | 0.195715328 | 2.15E-05 |
| SNP_A.K_994781_6_5888777 | 6 | 5888777 | 3.48E-09 | 0.09197 | 299 | 0.195506167 | 2.21E-05 |
| SNP_A.K_997484_6_6103259 | 6 | 6103259 | 3.55E-09 | 0.11538 | 299 | 0.195361241 | 2.25E-05 |
| SNP_A.K_1000308_6_6371694 | 6 | 6371694 | 3.57E-09 | 0.08696 | 299 | 0.195321496 | 2.26E-05 |
| SNP_A.K_994099_6_5814805 | 6 | 5814805 | 3.59E-09 | 0.16555 | 299 | 0.195276317 | 2.27E-05 |
| SNP_A.K_997693_6_6142570 | 6 | 6142570 | 3.66E-09 | 0.11538 | 299 | 0.195152155 | 2.30E-05 |
| SNP_A.K_1003050_6_6653023 | 6 | 6653023 | 3.70E-09 | 0.1505 | 299 | 0.195084029 | 2.32E-05 |
| SNP_A.K_996866_6_6031194 | 6 | 6031194 | 3.71E-09 | 0.0903 | 299 | 0.195051438 | 2.32E-05 |
| SNP_A.K_325949_2_9964036 | 2 | 9964036 | 3.74E-09 | 0.04682 | 299 | 0.195004907 | 2.32E-05 |
| SNP_A.K_325545_2_9901600 | 2 | 9901600 | 3.74E-09 | 0.03512 | 299 | 0.195004067 | 2.32E-05 |
| SNP_A.K_306863_2_7361415 | 2 | 7361415 | 4.03E-09 | 0.25585 | 299 | 0.19448783 | 2.50E-05 |
| SNP_A.K_994980_6_5909464 | 6 | 5909464 | 4.20E-09 | 0.08696 | 299 | 0.194206258 | 2.60E-05 |
| SNP_A.K_999266_6_6275506 | 6 | 6275506 | 4.27E-09 | 0.11037 | 299 | 0.194095652 | 2.63E-05 |
| SNP_A.K_1198617_7_7902877 | 7 | 7902877 | 4.49E-09 | 0.01338 | 299 | 0.193752478 | 2.76E-05 |
| SNP_A.K_993095_6_5707541 | 6 | 5707541 | 4.63E-09 | 0.11371 | 299 | 0.193537507 | 2.84E-05 |
| SNP_A.K_993046_6_5701032 | 6 | 5701032 | 4.76E-09 | 0.11371 | 299 | 0.193358821 | 2.90E-05 |
| SNP_A.K_325603_2_9915633 | 2 | 9915633 | 4.86E-09 | 0.02843 | 299 | 0.193213371 | 2.96E-05 |
| SNP_A.K_996676_6_6016570 | 6 | 6016570 | 4.91E-09 | 0.10033 | 299 | 0.193145828 | 2.98E-05 |
| SNP_A.K_996900_6_6033059 | 6 | 6033059 | 5.11E-09 | 0.12876 | 299 | 0.192867709 | 3.09E-05 |
| SNP_A.K_993138_6_5710380 | 6 | 5710380 | 5.47E-09 | 0.11371 | 299 | 0.192409335 | 3.30E-05 |

| | | | | | | | |
|---|---|---|---|---|---|---|---|
| SNP_A.K_1000268_6_6360016 | 6 | 6360016 | 5.53E-09 | 0.0903 | 299 | 0.192331104 | 3.33E-05 |
| SNP_A.K_996861_6_6031038 | 6 | 6031038 | 5.57E-09 | 0.10033 | 299 | 0.192275319 | 3.34E-05 |
| SNP_A.K_997575_6_6109614 | 6 | 6109614 | 5.64E-09 | 0.11037 | 299 | 0.192192965 | 3.37E-05 |
| SNP_A.K_153549_1_23835205 | 1 | 23835205 | 5.81E-09 | 0.01505 | 299 | 0.191993795 | 3.46E-05 |
| SNP_A.K_326609_2_10035213 | 2 | 10035213 | 6.04E-09 | 0.02843 | 299 | 0.191731305 | 3.59E-05 |
| SNP_A.K_994384_6_5854540 | 6 | 5854540 | 6.06E-09 | 0.12709 | 299 | 0.19170841 | 3.59E-05 |
| SNP_A.K_993016_6_5698260 | 6 | 5698260 | 6.20E-09 | 0.11371 | 299 | 0.191552293 | 3.66E-05 |
| SNP_A.K_997620_6_6139730 | 6 | 6139730 | 6.22E-09 | 0.10368 | 299 | 0.191532092 | 3.66E-05 |
| SNP_A.K_996772_6_6025063 | 6 | 6025063 | 6.26E-09 | 0.1087 | 299 | 0.191477797 | 3.68E-05 |
| SNP_A.K_992718_6_5675999 | 6 | 5675999 | 6.43E-09 | 0.11706 | 299 | 0.191303506 | 3.77E-05 |
| SNP_A.K_994382_6_5854415 | 6 | 5854415 | 6.45E-09 | 0.13043 | 299 | 0.191276028 | 3.77E-05 |
| SNP_A.K_992650_6_5668869 | 6 | 5668869 | 6.66E-09 | 0.13211 | 299 | 0.191061662 | 3.88E-05 |
| SNP_A.K_996430_6_6000692 | 6 | 6000692 | 6.70E-09 | 0.09699 | 299 | 0.191023156 | 3.89E-05 |
| SNP_A.K_999712_6_6308691 | 6 | 6308691 | 6.76E-09 | 0.1505 | 299 | 0.190963475 | 3.91E-05 |
| SNP_A.K_993023_6_5698597 | 6 | 5698597 | 6.76E-09 | 0.11204 | 299 | 0.190961745 | 3.91E-05 |
| SNP_A.K_992553_6_5656995 | 6 | 5656995 | 7.24E-09 | 0.13378 | 299 | 0.190494171 | 4.17E-05 |
| SNP_A.K_1004070_6_6825388 | 6 | 6825388 | 7.24E-09 | 0.1087 | 299 | 0.190488005 | 4.17E-05 |
| SNP_A.K_997858_6_6154262 | 6 | 6154262 | 7.40E-09 | 0.1505 | 299 | 0.190345751 | 4.24E-05 |
| SNP_A.K_993011_6_5698041 | 6 | 5698041 | 7.58E-09 | 0.11371 | 299 | 0.190178105 | 4.33E-05 |
| SNP_A.K_996454_6_6002242 | 6 | 6002242 | 7.60E-09 | 0.10033 | 299 | 0.190161912 | 4.33E-05 |
| SNP_A.K_993467_6_5754363 | 6 | 5754363 | 7.89E-09 | 0.13043 | 299 | 0.18990708 | 4.48E-05 |
| SNP_A.K_1002178_6_6551261 | 6 | 6551261 | 7.90E-09 | 0.15552 | 299 | 0.189898521 | 4.48E-05 |
| SNP_A.K_992496_6_5652972 | 6 | 5652972 | 8.04E-09 | 0.11873 | 299 | 0.189777034 | 4.55E-05 |
| SNP_A.K_996136_6_5964491 | 6 | 5964491 | 8.15E-09 | 0.09197 | 299 | 0.189687801 | 4.60E-05 |
| SNP_A.K_992460_6_5651689 | 6 | 5651689 | 8.24E-09 | 0.11873 | 299 | 0.189611262 | 4.64E-05 |
| SNP_A.K_1363253_8_5767274 | 8 | 5767274 | 8.33E-09 | 0.00836 | 299 | 0.189535749 | 4.67E-05 |
| SNP_A.K_994063_6_5812365 | 6 | 5812365 | 8.39E-09 | 0.12207 | 299 | 0.189492171 | 4.69E-05 |
| SNP_A.K_306698_2_7341926 | 2 | 7341926 | 8.52E-09 | 0.23244 | 299 | 0.189383676 | 4.75E-05 |
| SNP_A.K_1005628_6_7030955 | 6 | 7030955 | 8.56E-09 | 0.21572 | 299 | 0.189352197 | 4.76E-05 |
| SNP_A.K_992858_6_5681932 | 6 | 5681932 | 9.02E-09 | 0.11538 | 299 | 0.188993259 | 5.01E-05 |
| SNP_A.K_996395_6_5999382 | 6 | 5999382 | 9.18E-09 | 0.11037 | 299 | 0.188877564 | 5.08E-05 |
| SNP_A.K_998187_6_6183490 | 6 | 6183490 | 9.29E-09 | 0.09532 | 299 | 0.188798824 | 5.13E-05 |
| SNP_A.K_995998_6_5957240 | 6 | 5957240 | 9.52E-09 | 0.17726 | 299 | 0.188631278 | 5.24E-05 |
| SNP_A.K_993050_6_5701548 | 6 | 5701548 | 9.86E-09 | 0.11538 | 299 | 0.18839098 | 5.41E-05 |
| SNP_A.K_307042_2_7374688 | 2 | 7374688 | 9.98E-09 | 0.2408 | 299 | 0.188309537 | 5.45E-05 |
| SNP_A.K_8302_1_1541551 | 1 | 1541551 | 1.00E-08 | 0.00836 | 299 | 0.188292026 | 5.45E-05 |
| SNP_A.K_992636_6_5667426 | 6 | 5667426 | 1.00E-08 | 0.11538 | 299 | 0.188273973 | 5.45E-05 |
| SNP_A.K_996878_6_6031585 | 6 | 6031585 | 1.00E-08 | 0.09365 | 299 | 0.188273037 | 5.45E-05 |
| SNP_A.K_996483_6_6003523 | 6 | 6003523 | 1.04E-08 | 0.09197 | 299 | 0.188042564 | 5.63E-05 |
| SNP_A.K_993191_6_5716326 | 6 | 5716326 | 1.04E-08 | 0.11538 | 299 | 0.188010684 | 5.64E-05 |
| SNP_A.K_1364936_8_6060386 | 8 | 6060386 | 1.05E-08 | 0.08528 | 299 | 0.187983373 | 5.65E-05 |
| SNP_A.K_998936_6_6235674 | 6 | 6235674 | 1.07E-08 | 0.10368 | 299 | 0.187859581 | 5.73E-05 |
| SNP_A.K_992613_6_5665213 | 6 | 5665213 | 1.12E-08 | 0.1204 | 299 | 0.187510735 | 6.02E-05 |
| SNP_A.K_999789_6_6315470 | 6 | 6315470 | 1.16E-08 | 0.09532 | 299 | 0.187313425 | 6.18E-05 |
| SNP_A.K_995320_6_5926014 | 6 | 5926014 | 1.18E-08 | 0.1388 | 299 | 0.18715159 | 6.32E-05 |
| SNP_A.K_307278_2_7391064 | 2 | 7391064 | 1.23E-08 | 0.23746 | 299 | 0.186864705 | 6.56E-05 |
| SNP_A.K_993189_6_5716124 | 6 | 5716124 | 1.24E-08 | 0.11706 | 299 | 0.186857083 | 6.56E-05 |
| SNP_A.K_991466_6_5384404 | 6 | 5384404 | 1.26E-08 | 0.10201 | 299 | 0.186702277 | 6.70E-05 |
| SNP_A.K_997423_6_6098199 | 6 | 6098199 | 1.28E-08 | 0.12542 | 299 | 0.186601987 | 6.78E-05 |
| SNP_A.K_526927_3_11022317 | 3 | 11022317 | 1.32E-08 | 0.01171 | 299 | 0.186435061 | 6.93E-05 |

| | | | | | | | |
|---|---|---|---|---|---|---|---|
| SNP_A.K_1006018_6_7096695 | 6 | 7096695 | 1.32E-08 | 0.27926 | 299 | 0.186409256 | 6.94E-05 |
| SNP_A.K_999579_6_6297708 | 6 | 6297708 | 1.35E-08 | 0.09532 | 299 | 0.186264664 | 7.07E-05 |
| SNP_A.K_1002526_6_6581196 | 6 | 6581196 | 1.35E-08 | 0.18896 | 299 | 0.1862488 | 7.07E-05 |
| SNP_A.K_993096_6_5707553 | 6 | 5707553 | 1.36E-08 | 0.1087 | 299 | 0.186188045 | 7.12E-05 |
| SNP_A.K_994182_6_5822877 | 6 | 5822877 | 1.42E-08 | 0.08528 | 299 | 0.185937149 | 7.37E-05 |
| SNP_A.K_999652_6_6303233 | 6 | 6303233 | 1.47E-08 | 0.20234 | 299 | 0.185670457 | 7.64E-05 |
| SNP_A.K_998027_6_6166823 | 6 | 6166823 | 1.48E-08 | 0.11873 | 299 | 0.18565268 | 7.64E-05 |
| SNP_A.K_996412_6_5999758 | 6 | 5999758 | 1.48E-08 | 0.08194 | 299 | 0.185628911 | 7.65E-05 |
| SNP_A.K_993105_6_5708271 | 6 | 5708271 | 1.51E-08 | 0.11873 | 299 | 0.18551998 | 7.76E-05 |
| SNP_A.K_999654_6_6303294 | 6 | 6303294 | 1.52E-08 | 0.18729 | 299 | 0.185453178 | 7.81E-05 |
| SNP_A.K_999273_6_6275876 | 6 | 6275876 | 1.52E-08 | 0.11037 | 299 | 0.185439275 | 7.81E-05 |
| SNP_A.K_1830656_11_10572969 | 11 | 10572969 | 1.53E-08 | 0.16555 | 299 | 0.18541895 | 7.82E-05 |
| SNP_A.K_1000528_6_6391236 | 6 | 6391236 | 1.54E-08 | 0.10033 | 299 | 0.185369537 | 7.84E-05 |
| SNP_A.K_997297_6_6085744 | 6 | 6085744 | 1.54E-08 | 0.11371 | 299 | 0.185367603 | 7.84E-05 |
| SNP_A.K_326923_2_10070797 | 2 | 10070797 | 1.55E-08 | 0.06689 | 299 | 0.185330154 | 7.85E-05 |
| SNP_A.K_997199_6_6056708 | 6 | 6056708 | 1.55E-08 | 0.12709 | 299 | 0.185319017 | 7.85E-05 |
| SNP_A.K_978953_6_1826585 | 6 | 1826585 | 1.68E-08 | 0.29599 | 299 | 0.184787258 | 8.48E-05 |
| SNP_A.K_1364929_8_6059874 | 8 | 6059874 | 1.71E-08 | 0.07191 | 299 | 0.184657268 | 8.62E-05 |
| SNP_A.K_1005393_6_7011650 | 6 | 7011650 | 1.75E-08 | 0.27759 | 299 | 0.184520159 | 8.77E-05 |
| SNP_A.K_995322_6_5926036 | 6 | 5926036 | 1.76E-08 | 0.13545 | 299 | 0.184477492 | 8.81E-05 |
| SNP_A.K_1005601_6_7029459 | 6 | 7029459 | 1.79E-08 | 0.28428 | 299 | 0.184332113 | 8.98E-05 |
| SNP_A.K_998789_6_6222413 | 6 | 6222413 | 1.82E-08 | 0.12375 | 299 | 0.184235283 | 9.07E-05 |
| SNP_A.K_999701_6_6308189 | 6 | 6308189 | 1.82E-08 | 0.1689 | 299 | 0.184231349 | 9.07E-05 |
| SNP_A.K_992514_6_5654471 | 6 | 5654471 | 1.84E-08 | 0.11706 | 299 | 0.184177915 | 9.12E-05 |
| SNP_A.K_999896_6_6331874 | 6 | 6331874 | 1.85E-08 | 0.1806 | 299 | 0.184143599 | 9.13E-05 |
| SNP_A.K_1371260_8_7816922 | 8 | 7816922 | 1.85E-08 | 0.00669 | 299 | 0.18410837 | 9.13E-05 |
| SNP_A.K_1371321_8_7820142 | 8 | 7820142 | 1.85E-08 | 0.00669 | 299 | 0.18410837 | 9.13E-05 |
| SNP_A.K_1371325_8_7820415 | 8 | 7820415 | 1.85E-08 | 0.00669 | 299 | 0.18410837 | 9.13E-05 |
| SNP_A.K_993107_6_5708482 | 6 | 5708482 | 1.90E-08 | 0.11706 | 299 | 0.183930328 | 9.35E-05 |
| SNP_A.K_1002655_6_6601669 | 6 | 6601669 | 1.99E-08 | 0.10535 | 299 | 0.183644049 | 9.73E-05 |
| SNP_A.K_1003561_6_6739023 | 6 | 6739023 | 2.02E-08 | 0.00669 | 299 | 0.183523057 | 9.88E-05 |
| SNP_A.K_998103_6_6171160 | 6 | 6171160 | 2.03E-08 | 0.1087 | 299 | 0.183493967 | 9.90E-05 |
| SNP_A.K_991438_6_5269712 | 6 | 5269712 | 2.07E-08 | 0.04515 | 299 | 0.183356024 | 0.000100832 |
| SNP_A.K_996128_6_5964208 | 6 | 5964208 | 2.18E-08 | 0.09699 | 299 | 0.183015795 | 0.000105796 |
| SNP_A.K_995631_6_5944115 | 6 | 5944115 | 2.25E-08 | 0.11538 | 299 | 0.182806443 | 0.000108875 |
| SNP_A.K_994729_6_5884185 | 6 | 5884185 | 2.30E-08 | 0.14883 | 299 | 0.182666055 | 0.000110906 |
| SNP_A.K_992947_6_5690053 | 6 | 5690053 | 2.33E-08 | 0.11037 | 299 | 0.182561744 | 0.000112373 |
| SNP_A.K_994646_6_5877804 | 6 | 5877804 | 2.34E-08 | 0.18227 | 299 | 0.182541893 | 0.000112444 |
| SNP_A.K_1004693_6_6918935 | 6 | 6918935 | 2.42E-08 | 0.43144 | 299 | 0.182310319 | 0.000116103 |
| SNP_A.K_993211_6_5719217 | 6 | 5719217 | 2.59E-08 | 0.11037 | 299 | 0.181854967 | 0.00012393 |
| SNP_A.K_994849_6_5894471 | 6 | 5894471 | 2.70E-08 | 0.0903 | 299 | 0.181567786 | 0.000128899 |
| SNP_A.K_998342_6_6190281 | 6 | 6190281 | 2.71E-08 | 0.09365 | 299 | 0.181559118 | 0.000128899 |
| SNP_A.K_995798_6_5950839 | 6 | 5950839 | 2.73E-08 | 0.20401 | 299 | 0.181511267 | 0.000129521 |
| SNP_A.K_996409_6_5999704 | 6 | 5999704 | 2.77E-08 | 0.09866 | 299 | 0.181392724 | 0.00013152 |
| SNP_A.K_997460_6_6100243 | 6 | 6100243 | 2.78E-08 | 0.11538 | 299 | 0.18137351 | 0.000131596 |
| SNP_A.K_306726_2_7345691 | 2 | 7345691 | 2.81E-08 | 0.23746 | 299 | 0.181299525 | 0.000132748 |
| SNP_A.K_994212_6_5825721 | 6 | 5825721 | 2.89E-08 | 0.12375 | 299 | 0.181116982 | 0.000136086 |
| SNP_A.K_307161_2_7382785 | 2 | 7382785 | 2.94E-08 | 0.24415 | 299 | 0.180997347 | 0.000138213 |
| SNP_A.K_993156_6_5710885 | 6 | 5710885 | 2.95E-08 | 0.1204 | 299 | 0.180976752 | 0.000138324 |
| SNP_A.K_992511_6_5654191 | 6 | 5654191 | 3.10E-08 | 0.11037 | 299 | 0.180649671 | 0.000144884 |

| | | | | | | | |
|---|---|---|---|---|---|---|---|
| SNP_A.K_1005323_6_7006059 | 6 | 7006059 | 3.22E-08 | 0.12709 | 299 | 0.180379792 | 0.000150475 |
| SNP_A.K_993512_6_5763550 | 6 | 5763550 | 3.24E-08 | 0.11538 | 299 | 0.180339301 | 0.000151045 |
| SNP_A.K_1390296_8_10172119 | 8 | 10172119 | 3.27E-08 | 0.05518 | 299 | 0.180287689 | 0.000151868 |
| SNP_A.K_993321_6_5738277 | 6 | 5738277 | 3.32E-08 | 0.12709 | 299 | 0.18017528 | 0.000154083 |
| SNP_A.K_997951_6_6162089 | 6 | 6162089 | 3.35E-08 | 0.09532 | 299 | 0.180130625 | 0.000154765 |
| SNP_A.K_1005375_6_7010344 | 6 | 7010344 | 3.47E-08 | 0.2893 | 299 | 0.179883809 | 0.000160195 |
| SNP_A.K_993352_6_5740917 | 6 | 5740917 | 3.56E-08 | 0.11873 | 299 | 0.179706599 | 0.000163993 |
| SNP_A.K_1364977_8_6063489 | 8 | 6063489 | 3.57E-08 | 0.03177 | 299 | 0.179696531 | 0.000163993 |
| SNP_A.K_995327_6_5926112 | 6 | 5926112 | 3.61E-08 | 0.13043 | 299 | 0.179626284 | 0.00016535 |
| SNP_A.K_993280_6_5730059 | 6 | 5730059 | 3.78E-08 | 0.11706 | 299 | 0.179314926 | 0.000172811 |
| SNP_A.K_1364729_8_6038354 | 8 | 6038354 | 3.79E-08 | 0.03344 | 299 | 0.179284655 | 0.00017321 |
| SNP_A.K_1005319_6_7005641 | 6 | 7005641 | 3.89E-08 | 0.2893 | 299 | 0.179124832 | 0.000176992 |
| SNP_A.K_992730_6_5676591 | 6 | 5676591 | 3.96E-08 | 0.16054 | 299 | 0.179005494 | 0.000179771 |
| SNP_A.K_978697_6_1766020 | 6 | 1766020 | 4.00E-08 | 0.2893 | 299 | 0.178937576 | 0.000181203 |
| SNP_A.K_994381_6_5854407 | 6 | 5854407 | 4.08E-08 | 0.12709 | 299 | 0.17879303 | 0.000184743 |
| SNP_A.K_999265_6_6275495 | 6 | 6275495 | 4.11E-08 | 0.11538 | 299 | 0.178743394 | 0.00018571 |
| SNP_A.K_997414_6_6098080 | 6 | 6098080 | 4.14E-08 | 0.1204 | 299 | 0.178702085 | 0.000186452 |
| SNP_A.K_306805_2_7352465 | 2 | 7352465 | 4.22E-08 | 0.23579 | 299 | 0.178566344 | 0.000189849 |
| SNP_A.K_990608_6_4728236 | 6 | 4728236 | 4.51E-08 | 0.13043 | 299 | 0.178118049 | 0.000202527 |
| SNP_A.K_993344_6_5740379 | 6 | 5740379 | 4.56E-08 | 0.11037 | 299 | 0.178047778 | 0.000203781 |
| SNP_A.K_993215_6_5720734 | 6 | 5720734 | 4.56E-08 | 0.11037 | 299 | 0.178047778 | 0.000203781 |
| SNP_A.K_307280_2_7391076 | 2 | 7391076 | 4.59E-08 | 0.24749 | 299 | 0.178011149 | 0.000204295 |
| SNP_A.K_1005278_6_7002095 | 6 | 7002095 | 4.59E-08 | 0.23244 | 299 | 0.178002163 | 0.000204295 |
| SNP_A.K_993244_6_5723647 | 6 | 5723647 | 4.73E-08 | 0.11706 | 299 | 0.177812056 | 0.00020972 |
| SNP_A.K_1005225_6_6995275 | 6 | 6995275 | 4.82E-08 | 0.29766 | 299 | 0.177674323 | 0.000213617 |
| SNP_A.K_993527_6_5768225 | 6 | 5768225 | 4.86E-08 | 0.12709 | 299 | 0.177626505 | 0.000214689 |
| SNP_A.K_709527_4_8271604 | 4 | 8271604 | 4.97E-08 | 0.08863 | 299 | 0.177478561 | 0.000219014 |
| SNP_A.K_999709_6_6308568 | 6 | 6308568 | 5.00E-08 | 0.13545 | 299 | 0.177426978 | 0.000220239 |
| SNP_A.K_326867_2_10062054 | 2 | 10062054 | 5.18E-08 | 0.06187 | 299 | 0.177194607 | 0.000227528 |
| SNP_A.K_995752_6_5949795 | 6 | 5949795 | 5.31E-08 | 0.10535 | 299 | 0.177035522 | 0.000232504 |
| SNP_A.K_993626_6_5780312 | 6 | 5780312 | 5.58E-08 | 0.20736 | 299 | 0.176698865 | 0.000243975 |
| SNP_A.K_995615_6_5943431 | 6 | 5943431 | 5.83E-08 | 0.12207 | 299 | 0.176409103 | 0.000254231 |
| SNP_A.K_1005324_6_7006068 | 6 | 7006068 | 5.90E-08 | 0.29431 | 299 | 0.176319941 | 0.000257101 |
| SNP_A.K_992618_6_5665456 | 6 | 5665456 | 5.93E-08 | 0.11037 | 299 | 0.176289665 | 0.000257727 |
| SNP_A.K_1005960_6_7056095 | 6 | 7056095 | 6.07E-08 | 0.13712 | 299 | 0.176134696 | 0.000263215 |
| SNP_A.K_1004664_6_6914441 | 6 | 6914441 | 6.09E-08 | 0.28595 | 299 | 0.176113147 | 0.000263515 |
| SNP_A.K_1004361_6_6876125 | 6 | 6876125 | 6.10E-08 | 0.31271 | 299 | 0.176097928 | 0.000263567 |
| SNP_A.K_992518_6_5654573 | 6 | 5654573 | 6.14E-08 | 0.12709 | 299 | 0.176060872 | 0.000264482 |
| SNP_A.K_1004855_6_6936396 | 6 | 6936396 | 6.21E-08 | 0.28428 | 299 | 0.175981619 | 0.00026708 |
| SNP_A.K_978737_6_1772653 | 6 | 1772653 | 6.32E-08 | 0.3311 | 299 | 0.175864054 | 0.000271254 |
| SNP_A.K_992862_6_5682324 | 6 | 5682324 | 6.62E-08 | 0.18395 | 299 | 0.175557694 | 0.000283383 |
| SNP_A.K_1004786_6_6931701 | 6 | 6931701 | 6.71E-08 | 0.2893 | 299 | 0.175464633 | 0.000286765 |
| SNP_A.K_306866_2_7361495 | 2 | 7361495 | 6.77E-08 | 0.2709 | 299 | 0.17540827 | 0.000288483 |
| SNP_A.K_998780_6_6221178 | 6 | 6221178 | 6.78E-08 | 0.1505 | 299 | 0.175397283 | 0.000288483 |
| SNP_A.K_992619_6_5665847 | 6 | 5665847 | 7.02E-08 | 0.11204 | 299 | 0.175165347 | 0.000298059 |
| SNP_A.K_978802_6_1782689 | 6 | 1782689 | 7.03E-08 | 0.31104 | 299 | 0.175147792 | 0.000298187 |
| SNP_A.K_1005487_6_7016760 | 6 | 7016760 | 7.05E-08 | 0.29766 | 299 | 0.175134241 | 0.000298187 |
| SNP_A.K_1004360_6_6876114 | 6 | 6876114 | 7.06E-08 | 0.31438 | 299 | 0.175121683 | 0.000298187 |
| SNP_A.K_306734_2_7346476 | 2 | 7346476 | 7.09E-08 | 0.24749 | 299 | 0.175093512 | 0.00029884 |
| SNP_A.K_995321_6_5926034 | 6 | 5926034 | 7.23E-08 | 0.13378 | 299 | 0.174968728 | 0.000303218 |

| | | | | | | | |
|---|---|---|---|---|---|---|---|
| SNP_A.K_1004572_6_6907904 | 6 | 6907904 | 7.24E-08 | 0.29431 | 299 | 0.174959146 | 0.000303218 |
| SNP_A.K_1005314_6_7005301 | 6 | 7005301 | 7.24E-08 | 0.29264 | 299 | 0.174955859 | 0.000303218 |
| SNP_A.K_993576_6_5774842 | 6 | 5774842 | 7.34E-08 | 0.11371 | 299 | 0.174862718 | 0.000305821 |
| SNP_A.K_689897_4_4325774 | 4 | 4325774 | 7.35E-08 | 0.06355 | 299 | 0.174854845 | 0.000305821 |
| SNP_A.K_1364734_8_6038740 | 8 | 6038740 | 7.36E-08 | 0.03344 | 299 | 0.174847114 | 0.000305821 |
| SNP_A.K_306703_2_7342236 | 2 | 7342236 | 7.36E-08 | 0.23244 | 299 | 0.174845191 | 0.000305821 |
| SNP_A.K_1005454_6_7014355 | 6 | 7014355 | 7.42E-08 | 0.23579 | 299 | 0.174793855 | 0.000307566 |
| SNP_A.K_992580_6_5659658 | 6 | 5659658 | 7.49E-08 | 0.13545 | 299 | 0.174731433 | 0.000309836 |
| SNP_A.K_994371_6_5853160 | 6 | 5853160 | 7.55E-08 | 0.16722 | 299 | 0.174672812 | 0.000311698 |
| SNP_A.K_978579_6_1731808 | 6 | 1731808 | 7.57E-08 | 0.1388 | 299 | 0.174657347 | 0.000311698 |
| SNP_A.K_978695_6_1765448 | 6 | 1765448 | 7.58E-08 | 0.28094 | 299 | 0.174647045 | 0.000311698 |
| SNP_A.K_328725_2_10296603 | 2 | 10296603 | 7.59E-08 | 0.01505 | 299 | 0.174638512 | 0.000311698 |
| SNP_A.K_978710_6_1769240 | 6 | 1769240 | 7.75E-08 | 0.2893 | 299 | 0.17450448 | 0.00031739 |
| SNP_A.K_978734_6_1771928 | 6 | 1771928 | 7.80E-08 | 0.29933 | 299 | 0.174460367 | 0.000318866 |
| SNP_A.K_978793_6_1782083 | 6 | 1782083 | 7.85E-08 | 0.29933 | 299 | 0.17441119 | 0.000320593 |
| SNP_A.K_1830658_11_10573017 | 11 | 10573017 | 8.00E-08 | 0.17057 | 299 | 0.174288071 | 0.000325921 |
| SNP_A.K_307396_2_7400989 | 2 | 7400989 | 8.03E-08 | 0.24749 | 299 | 0.174264499 | 0.000326435 |
| SNP_A.K_326232_2_9986971 | 2 | 9986971 | 8.11E-08 | 0.03512 | 299 | 0.174199947 | 0.000328964 |
| SNP_A.K_306784_2_7350447 | 2 | 7350447 | 8.14E-08 | 0.23244 | 299 | 0.174174113 | 0.000329205 |
| SNP_A.K_993168_6_5711650 | 6 | 5711650 | 8.15E-08 | 0.12375 | 299 | 0.17416909 | 0.000329205 |
| SNP_A.K_993539_6_5769872 | 6 | 5769872 | 8.18E-08 | 0.11538 | 299 | 0.17414167 | 0.000329919 |
| SNP_A.K_1364944_8_6061267 | 8 | 6061267 | 8.42E-08 | 0.0903 | 299 | 0.173944625 | 0.000339147 |
| SNP_A.K_2065457_12_24399968 | 12 | 24399968 | 8.52E-08 | 0.02843 | 299 | 0.173871627 | 0.000342216 |
| SNP_A.K_978709_6_1769228 | 6 | 1769228 | 8.68E-08 | 0.28763 | 299 | 0.173743293 | 0.00034819 |
| SNP_A.K_992610_6_5665025 | 6 | 5665025 | 8.75E-08 | 0.11706 | 299 | 0.173694784 | 0.000348736 |
| SNP_A.K_1005888_6_7050492 | 6 | 7050492 | 8.75E-08 | 0.29264 | 299 | 0.173689182 | 0.000348736 |
| SNP_A.K_1433089_8_16617204 | 8 | 16617204 | 8.76E-08 | 0.08194 | 299 | 0.173686433 | 0.000348736 |
| SNP_A.K_996705_6_6017667 | 6 | 6017667 | 8.76E-08 | 0.10535 | 299 | 0.173681614 | 0.000348736 |
| SNP_A.K_994225_6_5827427 | 6 | 5827427 | 8.96E-08 | 0.12542 | 299 | 0.173532456 | 0.000355942 |
| SNP_A.K_992555_6_5657280 | 6 | 5657280 | 9.12E-08 | 0.1087 | 299 | 0.173411532 | 0.000361764 |
| SNP_A.K_1005223_6_6995207 | 6 | 6995207 | 9.25E-08 | 0.24247 | 299 | 0.173318839 | 0.00036613 |
| SNP_A.K_1599650_9_18583384 | 9 | 18583384 | 9.85E-08 | 0.03512 | 299 | 0.172903898 | 0.000388905 |
| SNP_A.K_995337_6_5926466 | 6 | 5926466 | 9.89E-08 | 0.13043 | 299 | 0.172871442 | 0.000390064 |
| SNP_A.K_527484_3_11137587 | 3 | 11137587 | 9.92E-08 | 0.01505 | 299 | 0.172853875 | 0.000390354 |
| SNP_A.K_998787_6_6222371 | 6 | 6222371 | 1.00E-07 | 0.18729 | 299 | 0.172781402 | 0.000393878 |
| SNP_A.K_992918_6_5684604 | 6 | 5684604 | 1.01E-07 | 0.1204 | 299 | 0.172735009 | 0.000395883 |
| SNP_A.K_1005321_6_7005868 | 6 | 7005868 | 1.01E-07 | 0.29431 | 299 | 0.172720675 | 0.00039599 |
| SNP_A.K_1371634_8_7860560 | 8 | 7860560 | 1.02E-07 | 0.00836 | 299 | 0.172655472 | 0.0003969 |
| SNP_A.K_1371474_8_7834834 | 8 | 7834834 | 1.02E-07 | 0.00836 | 299 | 0.172655472 | 0.0003969 |
| SNP_A.K_1371480_8_7835417 | 8 | 7835417 | 1.02E-07 | 0.00836 | 299 | 0.172655472 | 0.0003969 |
| SNP_A.K_1371699_8_7875943 | 8 | 7875943 | 1.02E-07 | 0.00836 | 299 | 0.172655472 | 0.0003969 |
| SNP_A.K_325759_2_9938123 | 2 | 9938123 | 1.05E-07 | 0.03177 | 299 | 0.172504387 | 0.000405249 |
| SNP_A.K_689896_4_4325625 | 4 | 4325625 | 1.05E-07 | 0.05686 | 299 | 0.172456224 | 0.000407431 |
| SNP_A.K_998160_6_6180544 | 6 | 6180544 | 1.07E-07 | 0.11371 | 299 | 0.172372211 | 0.00041114 |
| SNP_A.K_1005917_6_7052407 | 6 | 7052407 | 1.07E-07 | 0.27592 | 299 | 0.172371161 | 0.00041114 |
| SNP_A.K_1002499_6_6577922 | 6 | 6577922 | 1.08E-07 | 0.26923 | 299 | 0.172266245 | 0.000416324 |
| SNP_A.K_306697_2_7341888 | 2 | 7341888 | 1.08E-07 | 0.23746 | 299 | 0.172263103 | 0.000416324 |
| SNP_A.K_1005228_6_6995640 | 6 | 6995640 | 1.09E-07 | 0.29431 | 299 | 0.172246642 | 0.000416586 |
| SNP_A.K_992525_6_5654784 | 6 | 5654784 | 1.09E-07 | 0.11873 | 299 | 0.172219586 | 0.000417513 |
| SNP_A.K_1599597_9_18566375 | 9 | 18566375 | 1.10E-07 | 0.03846 | 299 | 0.172193794 | 0.000418364 |

| | | | | | | | |
|---|---|---|---|---|---|---|---|
| SNP_A.K_1364620_8_6029630 | 8 | 6029630 | 1.12E-07 | 0.09699 | 299 | 0.172066177 | 0.00042568 |
| SNP_A.K_993255_6_5724648 | 6 | 5724648 | 1.14E-07 | 0.11538 | 299 | 0.171914853 | 0.000434672 |
| SNP_A.K_942621_5_23062718 | 5 | 23062718 | 1.16E-07 | 0.21237 | 299 | 0.171835138 | 0.000439107 |
| SNP_A.K_328090_2_10197112 | 2 | 10197112 | 1.19E-07 | 0.01839 | 299 | 0.171645283 | 0.000450993 |
| SNP_A.K_673004_4_309896 | 4 | 309896 | 1.20E-07 | 0.04515 | 299 | 0.17156287 | 0.000455568 |
| SNP_A.K_997267_6_6079334 | 6 | 6079334 | 1.21E-07 | 0.15217 | 299 | 0.17155297 | 0.000455568 |
| SNP_A.K_1005633_6_7031828 | 6 | 7031828 | 1.21E-07 | 0.25251 | 299 | 0.171541871 | 0.000455568 |
| SNP_A.K_992886_6_5683022 | 6 | 5683022 | 1.22E-07 | 0.14047 | 299 | 0.171469752 | 0.00045949 |
| SNP_A.K_1005305_6_7003993 | 6 | 7003993 | 1.22E-07 | 0.27258 | 299 | 0.171460788 | 0.00045949 |
| SNP_A.K_978883_6_1794663 | 6 | 1794663 | 1.23E-07 | 0.30769 | 299 | 0.171417764 | 0.000461111 |
| SNP_A.K_977878_6_1608328 | 6 | 1608328 | 1.23E-07 | 0.15552 | 299 | 0.171413392 | 0.000461111 |
| SNP_A.K_993526_6_5768081 | 6 | 5768081 | 1.24E-07 | 0.11873 | 299 | 0.171391947 | 0.000461595 |
| SNP_A.K_993500_6_5759546 | 6 | 5759546 | 1.24E-07 | 0.11538 | 299 | 0.171382537 | 0.000461595 |
| SNP_A.K_992601_6_5664690 | 6 | 5664690 | 1.28E-07 | 0.11873 | 299 | 0.171166887 | 0.000475956 |
| SNP_A.K_997745_6_6145263 | 6 | 6145263 | 1.29E-07 | 0.14883 | 299 | 0.171119206 | 0.000478401 |
| SNP_A.K_2061896_12_23689543 | 12 | 23689543 | 1.29E-07 | 0.0301 | 299 | 0.171109073 | 0.000478401 |
| SNP_A.K_1005337_6_7007779 | 6 | 7007779 | 1.33E-07 | 0.301 | 299 | 0.170911273 | 0.000491972 |
| SNP_A.K_995316_6_5925953 | 6 | 5925953 | 1.36E-07 | 0.12876 | 299 | 0.170776814 | 0.000501134 |
| SNP_A.K_1004651_6_6913618 | 6 | 6913618 | 1.42E-07 | 0.31271 | 299 | 0.170477105 | 0.000523327 |
| SNP_A.K_1004658_6_6913983 | 6 | 6913983 | 1.43E-07 | 0.29264 | 299 | 0.170400037 | 0.000528496 |
| SNP_A.K_993885_6_5799259 | 6 | 5799259 | 1.44E-07 | 0.19732 | 299 | 0.170380358 | 0.000529128 |
| SNP_A.K_999650_6_6302785 | 6 | 6302785 | 1.50E-07 | 0.13043 | 299 | 0.170083734 | 0.000552321 |
| SNP_A.K_978775_6_1779985 | 6 | 1779985 | 1.51E-07 | 0.29766 | 299 | 0.170054341 | 0.000553314 |
| SNP_A.K_1005462_6_7015065 | 6 | 7015065 | 1.51E-07 | 0.27759 | 299 | 0.170048428 | 0.000553314 |
| SNP_A.K_1364942_8_6061118 | 8 | 6061118 | 1.52E-07 | 0.09532 | 299 | 0.170002599 | 0.00055523 |
| SNP_A.K_995211_6_5920302 | 6 | 5920302 | 1.52E-07 | 0.12375 | 299 | 0.170002181 | 0.00055523 |
| SNP_A.K_992494_6_5652924 | 6 | 5652924 | 1.60E-07 | 0.11706 | 299 | 0.169695498 | 0.000580469 |
| SNP_A.K_998161_6_6180565 | 6 | 6180565 | 1.60E-07 | 0.11037 | 299 | 0.169673965 | 0.000581339 |
| SNP_A.K_710548_4_8426976 | 4 | 8426976 | 1.63E-07 | 0.10033 | 299 | 0.169570014 | 0.000588892 |
| SNP_A.K_999662_6_6303577 | 6 | 6303577 | 1.63E-07 | 0.18395 | 299 | 0.169565194 | 0.000588892 |
| SNP_A.K_1364826_8_6046066 | 8 | 6046066 | 1.64E-07 | 0.09197 | 299 | 0.169523888 | 0.000590712 |
| SNP_A.K_997633_6_6139999 | 6 | 6139999 | 1.64E-07 | 0.11538 | 299 | 0.169521677 | 0.000590712 |
| SNP_A.K_1364970_8_6062391 | 8 | 6062391 | 1.64E-07 | 0.0903 | 299 | 0.169509918 | 0.000590736 |
| SNP_A.K_992874_6_5682720 | 6 | 5682720 | 1.65E-07 | 0.11371 | 299 | 0.169484316 | 0.000591996 |
| SNP_A.K_978751_6_1775037 | 6 | 1775037 | 1.65E-07 | 0.37793 | 299 | 0.169457393 | 0.000592012 |
| SNP_A.K_978718_6_1771148 | 6 | 1771148 | 1.66E-07 | 0.37458 | 299 | 0.169450414 | 0.000592012 |
| SNP_A.K_1390018_8_10145436 | 8 | 10145436 | 1.66E-07 | 0.04849 | 299 | 0.169449851 | 0.000592012 |
| SNP_A.K_1005908_6_7051226 | 6 | 7051226 | 1.67E-07 | 0.301 | 299 | 0.169406036 | 0.000594912 |
| SNP_A.K_978794_6_1782121 | 6 | 1782121 | 1.68E-07 | 0.30268 | 299 | 0.169352739 | 0.000598685 |
| SNP_A.K_1005446_6_7014242 | 6 | 7014242 | 1.68E-07 | 0.28094 | 299 | 0.169340185 | 0.000598793 |
| SNP_A.K_908233_5_17491226 | 5 | 17491226 | 1.69E-07 | 0.02341 | 299 | 0.169320745 | 0.000599526 |
| SNP_A.K_1005709_6_7034809 | 6 | 7034809 | 1.70E-07 | 0.24582 | 299 | 0.169279755 | 0.000602215 |
| SNP_A.K_993384_6_5744658 | 6 | 5744658 | 1.71E-07 | 0.12876 | 299 | 0.169238589 | 0.000603698 |
| SNP_A.K_978672_6_1755397 | 6 | 1755397 | 1.71E-07 | 0.27759 | 299 | 0.169236122 | 0.000603698 |
| SNP_A.K_998243_6_6186002 | 6 | 6186002 | 1.71E-07 | 0.13378 | 299 | 0.169227596 | 0.000603698 |
| SNP_A.K_1608644_9_20809208 | 9 | 20809208 | 1.71E-07 | 0.24916 | 299 | 0.169218399 | 0.000603698 |
| SNP_A.K_978719_6_1771154 | 6 | 1771154 | 1.75E-07 | 0.36455 | 299 | 0.169081588 | 0.000615241 |
| SNP_A.K_1004609_6_6910110 | 6 | 6910110 | 1.75E-07 | 0.29264 | 299 | 0.169062743 | 0.000615951 |
| SNP_A.K_1364831_8_6047477 | 8 | 6047477 | 1.81E-07 | 0.09699 | 299 | 0.16885224 | 0.000634751 |
| SNP_A.K_993141_6_5710490 | 6 | 5710490 | 1.83E-07 | 0.11706 | 299 | 0.168801329 | 0.00063857 |

| | | | | | | | |
|---|---|---|---|---|---|---|---|
| SNP_A.K_900798_5_16220448 | 5 | 16220448 | 1.83E-07 | 0.03679 | 299 | 0.168787267 | 0.000638853 |
| SNP_A.K_1004610_6_6910145 | 6 | 6910145 | 1.91E-07 | 0.29599 | 299 | 0.16849374 | 0.000666666 |
| SNP_A.K_1005916_6_7051934 | 6 | 7051934 | 1.95E-07 | 0.27592 | 299 | 0.168365648 | 0.000678544 |
| SNP_A.K_978671_6_1755368 | 6 | 1755368 | 1.97E-07 | 0.27759 | 299 | 0.16829238 | 0.000683242 |
| SNP_A.K_978704_6_1769141 | 6 | 1769141 | 1.97E-07 | 0.28595 | 299 | 0.168288717 | 0.000683242 |
| SNP_A.K_990584_6_4726929 | 6 | 4726929 | 1.97E-07 | 0.1204 | 299 | 0.168286794 | 0.000683242 |
| SNP_A.K_978670_6_1754821 | 6 | 1754821 | 1.99E-07 | 0.27425 | 299 | 0.168241744 | 0.000686762 |
| SNP_A.K_2080315_12_27256354 | 12 | 27256354 | 2.00E-07 | 0.00836 | 299 | 0.168202268 | 0.00068858 |
| SNP_A.K_2080229_12_27246573 | 12 | 27246573 | 2.00E-07 | 0.00836 | 299 | 0.168202268 | 0.00068858 |
| SNP_A.K_978783_6_1780336 | 6 | 1780336 | 2.01E-07 | 0.29933 | 299 | 0.168168354 | 0.000690971 |
| SNP_A.K_1363959_8_5860969 | 8 | 5860969 | 2.02E-07 | 0.01171 | 299 | 0.168146251 | 0.000692137 |
| SNP_A.K_998198_6_6184151 | 6 | 6184151 | 2.03E-07 | 0.13043 | 299 | 0.168108828 | 0.000694912 |
| SNP_A.K_1364838_8_6048202 | 8 | 6048202 | 2.04E-07 | 0.09197 | 299 | 0.168078108 | 0.000696995 |
| SNP_A.K_999539_6_6295100 | 6 | 6295100 | 2.09E-07 | 0.0786 | 299 | 0.167903841 | 0.000714408 |
| SNP_A.K_992521_6_5654624 | 6 | 5654624 | 2.09E-07 | 0.11706 | 299 | 0.167891463 | 0.000714571 |
| SNP_A.K_177897_1_28607327 | 1 | 28607327 | 2.10E-07 | 0.02341 | 299 | 0.167857368 | 0.000716945 |
| SNP_A.K_992471_6_5652099 | 6 | 5652099 | 2.11E-07 | 0.11706 | 299 | 0.167843818 | 0.000716945 |
| SNP_A.K_1005453_6_7014335 | 6 | 7014335 | 2.11E-07 | 0.27759 | 299 | 0.167837028 | 0.000716945 |
| SNP_A.K_306938_2_7369542 | 2 | 7369542 | 2.13E-07 | 0.21405 | 299 | 0.167789125 | 0.000720976 |
| SNP_A.K_307253_2_7388054 | 2 | 7388054 | 2.17E-07 | 0.24582 | 299 | 0.167654212 | 0.000734626 |
| SNP_A.K_710484_4_8424476 | 4 | 8424476 | 2.21E-07 | 0.07023 | 299 | 0.167540524 | 0.000745308 |
| SNP_A.K_992866_6_5682363 | 6 | 5682363 | 2.21E-07 | 0.18227 | 299 | 0.16753719 | 0.000745308 |
| SNP_A.K_1001774_6_6519962 | 6 | 6519962 | 2.24E-07 | 0.12709 | 299 | 0.167459681 | 0.000752869 |
| SNP_A.K_20623_1_3101817 | 1 | 3101817 | 2.28E-07 | 0.00669 | 299 | 0.16732109 | 0.000767565 |
| SNP_A.K_978784_6_1780935 | 6 | 1780935 | 2.31E-07 | 0.29097 | 299 | 0.167234651 | 0.000776406 |
| SNP_A.K_978701_6_1766460 | 6 | 1766460 | 2.32E-07 | 0.29431 | 299 | 0.167220169 | 0.000776855 |
| SNP_A.K_1608572_9_20787173 | 9 | 20787173 | 2.32E-07 | 0.23579 | 299 | 0.16720584 | 0.000777288 |
| SNP_A.K_171097_1_27544195 | 1 | 27544195 | 2.34E-07 | 0.02341 | 299 | 0.167163773 | 0.000780991 |
| SNP_A.K_307259_2_7388426 | 2 | 7388426 | 2.35E-07 | 0.2291 | 299 | 0.167126463 | 0.000784151 |
| SNP_A.K_992515_6_5654540 | 6 | 5654540 | 2.37E-07 | 0.12375 | 299 | 0.167058921 | 0.000790933 |
| SNP_A.K_978879_6_1794224 | 6 | 1794224 | 2.38E-07 | 0.30435 | 299 | 0.167034528 | 0.000792588 |
| SNP_A.K_1004393_6_6879131 | 6 | 6879131 | 2.39E-07 | 0.31271 | 299 | 0.167013556 | 0.000793837 |
| SNP_A.K_995255_6_5921567 | 6 | 5921567 | 2.40E-07 | 0.12542 | 299 | 0.166997673 | 0.000794479 |
| SNP_A.K_1005420_6_7013177 | 6 | 7013177 | 2.45E-07 | 0.30769 | 299 | 0.166861037 | 0.000809778 |
| SNP_A.K_978694_6_1765415 | 6 | 1765415 | 2.46E-07 | 0.27926 | 299 | 0.166836303 | 0.000811523 |
| SNP_A.K_1005912_6_7051414 | 6 | 7051414 | 2.60E-07 | 0.27926 | 299 | 0.166459675 | 0.000857748 |
| SNP_A.K_999314_6_6280780 | 6 | 6280780 | 2.62E-07 | 0.1388 | 299 | 0.166400297 | 0.000864122 |
| SNP_A.K_1005315_6_7005314 | 6 | 7005314 | 2.64E-07 | 0.25418 | 299 | 0.166357371 | 0.00086838 |
| SNP_A.K_1371330_8_7820516 | 8 | 7820516 | 2.65E-07 | 0.01672 | 299 | 0.16634379 | 0.000868795 |
| SNP_A.K_1004703_6_6921594 | 6 | 6921594 | 2.65E-07 | 0.27759 | 299 | 0.166326029 | 0.000869762 |
| SNP_A.K_1005500_6_7017178 | 6 | 7017178 | 2.69E-07 | 0.28595 | 299 | 0.166243626 | 0.000879294 |
| SNP_A.K_978950_6_1823332 | 6 | 1823332 | 2.71E-07 | 0.29264 | 299 | 0.166178652 | 0.000886592 |
| SNP_A.K_1005452_6_7014333 | 6 | 7014333 | 2.74E-07 | 0.23077 | 299 | 0.166109551 | 0.000894512 |
| SNP_A.K_307238_2_7387182 | 2 | 7387182 | 2.76E-07 | 0.23579 | 299 | 0.166063992 | 0.000899294 |
| SNP_A.K_993843_6_5796435 | 6 | 5796435 | 2.82E-07 | 0.1505 | 299 | 0.165929751 | 0.000916328 |
| SNP_A.K_306702_2_7342222 | 2 | 7342222 | 2.83E-07 | 0.23579 | 299 | 0.165908851 | 0.000917799 |
| SNP_A.K_1390048_8_10146895 | 8 | 10146895 | 2.84E-07 | 0.04849 | 299 | 0.165870405 | 0.00092154 |
| SNP_A.K_1371331_8_7820542 | 8 | 7820542 | 2.85E-07 | 0.01003 | 299 | 0.165858151 | 0.00092154 |
| SNP_A.K_1004359_6_6875866 | 6 | 6875866 | 2.85E-07 | 0.30435 | 299 | 0.165851162 | 0.00092154 |
| SNP_A.K_957685_5_26967577 | 5 | 26967577 | 2.87E-07 | 0.01839 | 299 | 0.165814958 | 0.000925171 |

| | | | | | | | |
|---|---|---|---|---|---|---|---|
| SNP_A.K_1371276_8_7817486 | 8 | 7817486 | 2.90E-07 | 0.01003 | 299 | 0.165742608 | 0.000933918 |
| SNP_A.K_1005232_6_6996522 | 6 | 6996522 | 2.91E-07 | 0.28261 | 299 | 0.165725444 | 0.000934902 |
| SNP_A.K_306916_2_7366560 | 2 | 7366560 | 2.92E-07 | 0.27759 | 299 | 0.165693046 | 0.000937515 |
| SNP_A.K_993883_6_5798993 | 6 | 5798993 | 2.92E-07 | 0.18395 | 299 | 0.165686649 | 0.000937515 |
| SNP_A.K_1004381_6_6878013 | 6 | 6878013 | 2.93E-07 | 0.13043 | 299 | 0.165669693 | 0.000938481 |
| SNP_A.K_1005441_6_7014033 | 6 | 7014033 | 2.95E-07 | 0.27759 | 299 | 0.165634434 | 0.000942058 |
| SNP_A.K_11421_1_1955684 | 1 | 1955684 | 3.00E-07 | 0.00669 | 299 | 0.165510143 | 0.000958495 |
| SNP_A.K_1004857_6_6936530 | 6 | 6936530 | 3.02E-07 | 0.2893 | 299 | 0.165469217 | 0.000962981 |
| SNP_A.K_992968_6_5691334 | 6 | 5691334 | 3.03E-07 | 0.14047 | 299 | 0.165445584 | 0.000963727 |
| SNP_A.K_306809_2_7352848 | 2 | 7352848 | 3.03E-07 | 0.23244 | 299 | 0.165443925 | 0.000963727 |
| SNP_A.K_995301_6_5924436 | 6 | 5924436 | 3.08E-07 | 0.12709 | 299 | 0.165348489 | 0.000976277 |
| SNP_A.K_1005385_6_7011329 | 6 | 7011329 | 3.08E-07 | 0.31773 | 299 | 0.165331111 | 0.000977362 |
| SNP_A.K_1004681_6_6916783 | 6 | 6916783 | 3.10E-07 | 0.301 | 299 | 0.165300713 | 0.000980172 |
| SNP_A.K_995291_6_5924019 | 6 | 5924019 | 3.10E-07 | 0.1204 | 299 | 0.165292114 | 0.000980172 |
| SNP_A.K_306727_2_7345818 | 2 | 7345818 | 3.11E-07 | 0.23411 | 299 | 0.165277709 | 0.000980827 |
| SNP_A.K_1371551_8_7852154 | 8 | 7852154 | 3.12E-07 | 0.01672 | 299 | 0.165261865 | 0.000981698 |
| SNP_A.K_990242_6_4702841 | 6 | 4702841 | 3.13E-07 | 0.2107 | 299 | 0.165239198 | 0.000983589 |
| SNP_A.K_999632_6_6301340 | 6 | 6301340 | 3.15E-07 | 0.19064 | 299 | 0.165184594 | 0.000990268 |
| SNP_A.K_1004177_6_6841690 | 6 | 6841690 | 3.16E-07 | 0.30602 | 299 | 0.165166474 | 0.000991497 |
| SNP_A.K_1005521_6_7018974 | 6 | 7018974 | 3.18E-07 | 0.23579 | 299 | 0.165133822 | 0.000993735 |
| SNP_A.K_992875_6_5682759 | 6 | 5682759 | 3.18E-07 | 0.1806 | 299 | 0.165131765 | 0.000993735 |
| SNP_A.K_1005332_6_7007554 | 6 | 7007554 | 3.23E-07 | 0.23244 | 299 | 0.165028861 | 0.001006804 |
| SNP_A.K_995293_6_5924075 | 6 | 5924075 | 3.23E-07 | 0.12876 | 299 | 0.165025814 | 0.001006804 |
| SNP_A.K_994373_6_5853168 | 6 | 5853168 | 3.25E-07 | 0.16388 | 299 | 0.164992829 | 0.001009575 |
| SNP_A.K_999030_6_6249316 | 6 | 6249316 | 3.25E-07 | 0.14047 | 299 | 0.16498797 | 0.001009575 |
| SNP_A.K_993265_6_5728362 | 6 | 5728362 | 3.27E-07 | 0.15719 | 299 | 0.164951437 | 0.001013671 |
| SNP_A.K_307374_2_7399685 | 2 | 7399685 | 3.28E-07 | 0.27425 | 299 | 0.164917112 | 0.001017445 |
| SNP_A.K_977872_6_1606079 | 6 | 1606079 | 3.29E-07 | 0.15719 | 299 | 0.164902864 | 0.001018131 |
| SNP_A.K_1004580_6_6908543 | 6 | 6908543 | 3.30E-07 | 0.31271 | 299 | 0.164883364 | 0.001019631 |
| SNP_A.K_1005382_6_7011231 | 6 | 7011231 | 3.36E-07 | 0.29933 | 299 | 0.164776897 | 0.001034695 |
| SNP_A.K_1005240_6_6997398 | 6 | 6997398 | 3.42E-07 | 0.33278 | 299 | 0.16465339 | 0.001052374 |
| SNP_A.K_307501_2_7413253 | 2 | 7413253 | 3.42E-07 | 0.23411 | 299 | 0.164645708 | 0.001052374 |
| SNP_A.K_1364820_8_6044920 | 8 | 6044920 | 3.43E-07 | 0.10033 | 299 | 0.164626183 | 0.00105394 |
| SNP_A.K_307160_2_7382743 | 2 | 7382743 | 3.46E-07 | 0.24916 | 299 | 0.164572531 | 0.001060989 |
| SNP_A.K_992454_6_5647197 | 6 | 5647197 | 3.48E-07 | 0.1204 | 299 | 0.164545578 | 0.001062503 |
| SNP_A.K_1004376_6_6877683 | 6 | 6877683 | 3.48E-07 | 0.33612 | 299 | 0.164543745 | 0.001062503 |
| SNP_A.K_993135_6_5709995 | 6 | 5709995 | 3.50E-07 | 0.12542 | 299 | 0.16450324 | 0.001067486 |
| SNP_A.K_1005456_6_7014542 | 6 | 7014542 | 3.54E-07 | 0.23244 | 299 | 0.164420851 | 0.001079334 |
| SNP_A.K_998069_6_6168624 | 6 | 6168624 | 3.56E-07 | 0.13545 | 299 | 0.164386956 | 0.001083314 |
| SNP_A.K_308590_2_7528214 | 2 | 7528214 | 3.58E-07 | 0.22408 | 299 | 0.164353549 | 0.001085988 |
| SNP_A.K_978752_6_1775050 | 6 | 1775050 | 3.58E-07 | 0.38963 | 299 | 0.164351473 | 0.001085988 |
| SNP_A.K_1364347_8_5991929 | 8 | 5991929 | 3.60E-07 | 0.01338 | 299 | 0.164312981 | 0.00108932 |
| SNP_A.K_1371504_8_7850184 | 8 | 7850184 | 3.60E-07 | 0.01672 | 299 | 0.164312105 | 0.00108932 |
| SNP_A.K_1005624_6_7030797 | 6 | 7030797 | 3.62E-07 | 0.23579 | 299 | 0.164269363 | 0.001094501 |
| SNP_A.K_690092_4_4404200 | 4 | 4404200 | 3.63E-07 | 0.04181 | 299 | 0.164261723 | 0.001094501 |
| SNP_A.K_993446_6_5751798 | 6 | 5751798 | 3.64E-07 | 0.14047 | 299 | 0.164239191 | 0.001096661 |
| SNP_A.K_1004346_6_6870440 | 6 | 6870440 | 3.68E-07 | 0.33278 | 299 | 0.164170816 | 0.001106502 |
| SNP_A.K_1364823_8_6045314 | 8 | 6045314 | 3.68E-07 | 0.05853 | 299 | 0.164161153 | 0.001106527 |
| SNP_A.K_977883_6_1612527 | 6 | 1612527 | 3.70E-07 | 0.15719 | 299 | 0.164139612 | 0.001108552 |
| SNP_A.K_1005425_6_7013618 | 6 | 7013618 | 3.73E-07 | 0.23411 | 299 | 0.164071859 | 0.001118403 |

| | | | | | | | |
|---|---|---|---|---|---|---|---|
| SNP_A.K_978673_6_1755645 | 6 | 1755645 | 3.81E-07 | 0.28428 | 299 | 0.163932842 | 0.001139593 |
| SNP_A.K_307532_2_7416038 | 2 | 7416038 | 3.82E-07 | 0.25753 | 299 | 0.163916012 | 0.001139593 |
| SNP_A.K_1198049_7_7832352 | 7 | 7832352 | 3.83E-07 | 0.01003 | 299 | 0.163910453 | 0.001139593 |
| SNP_A.K_1198080_7_7835380 | 7 | 7835380 | 3.83E-07 | 0.01003 | 299 | 0.163910453 | 0.001139593 |
| SNP_A.K_1004248_6_6853504 | 6 | 6853504 | 3.85E-07 | 0.32776 | 299 | 0.163870077 | 0.001143346 |
| SNP_A.K_326878_2_10064484 | 2 | 10064484 | 3.87E-07 | 0.07692 | 299 | 0.163840379 | 0.001143346 |
| SNP_A.K_1005508_6_7017838 | 6 | 7017838 | 3.87E-07 | 0.29264 | 299 | 0.163832646 | 0.001143346 |
| SNP_A.K_1005534_6_7019948 | 6 | 7019948 | 3.88E-07 | 0.24247 | 299 | 0.163823653 | 0.001143346 |
| SNP_A.K_307373_2_7399672 | 2 | 7399672 | 3.88E-07 | 0.21906 | 299 | 0.163822687 | 0.001143346 |
| SNP_A.K_328406_2_10261897 | 2 | 10261897 | 3.88E-07 | 0.04515 | 299 | 0.163817515 | 0.001143346 |
| SNP_A.K_992675_6_5671488 | 6 | 5671488 | 3.88E-07 | 0.14381 | 299 | 0.163813823 | 0.001143346 |
| SNP_A.K_326815_2_10057242 | 2 | 10057242 | 3.88E-07 | 0.06187 | 299 | 0.163812931 | 0.001143346 |
| SNP_A.K_1004155_6_6839214 | 6 | 6839214 | 3.89E-07 | 0.31605 | 299 | 0.163804572 | 0.001143346 |
| SNP_A.K_307405_2_7401533 | 2 | 7401533 | 3.90E-07 | 0.2291 | 299 | 0.163788139 | 0.001144586 |
| SNP_A.K_978764_6_1777321 | 6 | 1777321 | 3.91E-07 | 0.3495 | 299 | 0.163760958 | 0.001147703 |
| SNP_A.K_1371561_8_7852947 | 8 | 7852947 | 3.95E-07 | 0.01672 | 299 | 0.163700689 | 0.001156536 |
| SNP_A.K_1177595_7_2163967 | 7 | 2163967 | 3.96E-07 | 0.03177 | 299 | 0.163691965 | 0.001156536 |
| SNP_A.K_306945_2_7370258 | 2 | 7370258 | 4.01E-07 | 0.25418 | 299 | 0.163608478 | 0.001169656 |
| SNP_A.K_598481_3_24171758 | 3 | 24171758 | 4.02E-07 | 0.02007 | 299 | 0.16358251 | 0.001172635 |
| SNP_A.K_997783_6_6146427 | 6 | 6146427 | 4.03E-07 | 0.14214 | 299 | 0.163569566 | 0.0011733 |
| SNP_A.K_1005326_6_7006104 | 6 | 7006104 | 4.13E-07 | 0.28763 | 299 | 0.163405281 | 0.001198277 |
| SNP_A.K_307155_2_7382512 | 2 | 7382512 | 4.13E-07 | 0.24749 | 299 | 0.163402846 | 0.001198277 |
| SNP_A.K_1004399_6_6879583 | 6 | 6879583 | 4.14E-07 | 0.3194 | 299 | 0.163400011 | 0.001198277 |
| SNP_A.K_978951_6_1824082 | 6 | 1824082 | 4.14E-07 | 0.29431 | 299 | 0.163394225 | 0.001198277 |
| SNP_A.K_1004784_6_6931492 | 6 | 6931492 | 4.15E-07 | 0.28763 | 299 | 0.163380536 | 0.001199105 |
| SNP_A.K_389592_2_19885939 | 2 | 19885939 | 4.18E-07 | 0.02174 | 299 | 0.163322028 | 0.001208137 |
| SNP_A.K_306802_2_7351931 | 2 | 7351931 | 4.20E-07 | 0.23579 | 299 | 0.163294583 | 0.001211436 |
| SNP_A.K_1599602_9_18568232 | 9 | 18568232 | 4.21E-07 | 0.04348 | 299 | 0.163279034 | 0.001211436 |
| SNP_A.K_1364863_8_6049881 | 8 | 6049881 | 4.21E-07 | 0.08528 | 299 | 0.163276767 | 0.001211436 |
| SNP_A.K_307291_2_7391466 | 2 | 7391466 | 4.25E-07 | 0.23411 | 299 | 0.163225257 | 0.001219163 |
| SNP_A.K_1004180_6_6841855 | 6 | 6841855 | 4.25E-07 | 0.301 | 299 | 0.163216788 | 0.001219163 |
| SNP_A.K_1608612_9_20794226 | 9 | 20794226 | 4.27E-07 | 0.2408 | 299 | 0.163192935 | 0.00122183 |
| SNP_A.K_1390025_8_10145506 | 8 | 10145506 | 4.27E-07 | 0.04181 | 299 | 0.163184315 | 0.00122183 |
| SNP_A.K_457035_2_33231713 | 2 | 33231713 | 4.30E-07 | 0.02174 | 299 | 0.16314821 | 0.00122687 |
| SNP_A.K_1005422_6_7013265 | 6 | 7013265 | 4.31E-07 | 0.28595 | 299 | 0.163127805 | 0.001228996 |
| SNP_A.K_1550014_9_8775809 | 9 | 8775809 | 4.33E-07 | 0.02676 | 299 | 0.16309988 | 0.001232536 |
| SNP_A.K_1004152_6_6838670 | 6 | 6838670 | 4.35E-07 | 0.301 | 299 | 0.163072902 | 0.001235911 |
| SNP_A.K_998923_6_6234753 | 6 | 6234753 | 4.37E-07 | 0.13712 | 299 | 0.163038835 | 0.001240634 |
| SNP_A.K_1004895_6_6949684 | 6 | 6949684 | 4.39E-07 | 0.2893 | 299 | 0.163004486 | 0.001245432 |
| SNP_A.K_978772_6_1779545 | 6 | 1779545 | 4.40E-07 | 0.29097 | 299 | 0.162992238 | 0.001246056 |
| SNP_A.K_1005190_6_6990453 | 6 | 6990453 | 4.42E-07 | 0.25753 | 299 | 0.162967559 | 0.001249041 |
| SNP_A.K_990240_6_4702808 | 6 | 4702808 | 4.47E-07 | 0.21405 | 299 | 0.162887462 | 0.001262632 |
| SNP_A.K_1004385_6_6878700 | 6 | 6878700 | 4.48E-07 | 0.33946 | 299 | 0.162867287 | 0.001264796 |
| SNP_A.K_307377_2_7399770 | 2 | 7399770 | 4.53E-07 | 0.23579 | 299 | 0.162800441 | 0.001275991 |
| SNP_A.K_307461_2_7408743 | 2 | 7408743 | 4.56E-07 | 0.23077 | 299 | 0.162757855 | 0.001279782 |
| SNP_A.K_1005516_6_7018255 | 6 | 7018255 | 4.56E-07 | 0.29097 | 299 | 0.162752796 | 0.001279782 |
| SNP_A.K_1860753_11_15381015 | 11 | 15381015 | 4.57E-07 | 0.11037 | 299 | 0.162747242 | 0.001279782 |
| SNP_A.K_1371528_8_7851013 | 8 | 7851013 | 4.57E-07 | 0.01672 | 299 | 0.162745446 | 0.001279782 |
| SNP_A.K_306723_2_7345357 | 2 | 7345357 | 4.58E-07 | 0.23077 | 299 | 0.162724586 | 0.001282124 |
| SNP_A.K_995313_6_5925376 | 6 | 5925376 | 4.60E-07 | 0.13545 | 299 | 0.162705333 | 0.001284158 |

| | | | | | | | |
|---|---|---|---|---|---|---|---|
| SNP_A.K_306731_2_7346275 | 2 | 7346275 | 4.64E-07 | 0.23411 | 299 | 0.162641646 | 0.001294922 |
| SNP_A.K_1004079_6_6830286 | 6 | 6830286 | 4.75E-07 | 0.17057 | 299 | 0.162485675 | 0.001324243 |
| SNP_A.K_1364906_8_6054591 | 8 | 6054591 | 4.78E-07 | 0.08194 | 299 | 0.162446867 | 0.001330305 |
| SNP_A.K_307499_2_7413212 | 2 | 7413212 | 4.80E-07 | 0.2592 | 299 | 0.162426866 | 0.001332579 |
| SNP_A.K_1005694_6_7034225 | 6 | 7034225 | 4.83E-07 | 0.2709 | 299 | 0.162373592 | 0.001341635 |
| SNP_A.K_977907_6_1619620 | 6 | 1619620 | 4.84E-07 | 0.24582 | 299 | 0.162361395 | 0.001342338 |
| SNP_A.K_1004630_6_6912375 | 6 | 6912375 | 4.87E-07 | 0.2893 | 299 | 0.162321559 | 0.001348705 |
| SNP_A.K_1004861_6_6936730 | 6 | 6936730 | 4.90E-07 | 0.28595 | 299 | 0.162291159 | 0.00135316 |
| SNP_A.K_1364922_8_6057412 | 8 | 6057412 | 4.96E-07 | 0.0786 | 299 | 0.162205313 | 0.00136914 |
| SNP_A.K_978735_6_1772565 | 6 | 1772565 | 5.02E-07 | 0.31104 | 299 | 0.162130119 | 0.001383067 |
| SNP_A.K_978287_6_1669314 | 6 | 1669314 | 5.02E-07 | 0.1689 | 299 | 0.162119501 | 0.001383472 |
| SNP_A.K_1005208_6_6994515 | 6 | 6994515 | 5.05E-07 | 0.25251 | 299 | 0.162084781 | 0.001388967 |
| SNP_A.K_1390019_8_10145441 | 8 | 10145441 | 5.12E-07 | 0.05351 | 299 | 0.162002219 | 0.001404682 |
| SNP_A.K_997253_6_6061397 | 6 | 6061397 | 5.16E-07 | 0.06355 | 299 | 0.161939853 | 0.001416217 |
| SNP_A.K_1371604_8_7858316 | 8 | 7858316 | 5.18E-07 | 0.01003 | 299 | 0.161922168 | 0.001418168 |
| SNP_A.K_1003937_6_6807270 | 6 | 6807270 | 5.20E-07 | 0.301 | 299 | 0.161894009 | 0.001421663 |
| SNP_A.K_1608747_9_20829580 | 9 | 20829580 | 5.20E-07 | 0.24582 | 299 | 0.161888746 | 0.001421663 |
| SNP_A.K_992519_6_5654597 | 6 | 5654597 | 5.24E-07 | 0.11538 | 299 | 0.161842113 | 0.001429918 |
| SNP_A.K_995323_6_5926047 | 6 | 5926047 | 5.29E-07 | 0.21237 | 299 | 0.161785134 | 0.001440101 |
| SNP_A.K_978821_6_1784985 | 6 | 1784985 | 5.29E-07 | 0.301 | 299 | 0.161778334 | 0.001440101 |
| SNP_A.K_978803_6_1782840 | 6 | 1782840 | 5.37E-07 | 0.32107 | 299 | 0.161683297 | 0.001457286 |
| SNP_A.K_1000323_6_6374634 | 6 | 6374634 | 5.37E-07 | 0.10033 | 299 | 0.16168145 | 0.001457286 |
| SNP_A.K_1004036_6_6820916 | 6 | 6820916 | 5.38E-07 | 0.17224 | 299 | 0.161674762 | 0.001457286 |
| SNP_A.K_1005739_6_7036720 | 6 | 7036720 | 5.38E-07 | 0.23579 | 299 | 0.161664855 | 0.001457589 |
| SNP_A.K_1005004_6_6961307 | 6 | 6961307 | 5.41E-07 | 0.31605 | 299 | 0.161631715 | 0.001463064 |
| SNP_A.K_306994_2_7372249 | 2 | 7372249 | 5.45E-07 | 0.23411 | 299 | 0.161583618 | 0.001471912 |
| SNP_A.K_998978_6_6243940 | 6 | 6243940 | 5.47E-07 | 0.14214 | 299 | 0.161567123 | 0.001473705 |
| SNP_A.K_995328_6_5926145 | 6 | 5926145 | 5.50E-07 | 0.13043 | 299 | 0.161520386 | 0.001480714 |
| SNP_A.K_978799_6_1782431 | 6 | 1782431 | 5.51E-07 | 0.29933 | 299 | 0.161518997 | 0.001480714 |
| SNP_A.K_995248_6_5921315 | 6 | 5921315 | 5.53E-07 | 0.12207 | 299 | 0.16148686 | 0.001486063 |
| SNP_A.K_999629_6_6301310 | 6 | 6301310 | 5.56E-07 | 0.19398 | 299 | 0.161450882 | 0.00149014 |
| SNP_A.K_1004558_6_6901563 | 6 | 6901563 | 5.57E-07 | 0.2893 | 299 | 0.161448398 | 0.00149014 |
| SNP_A.K_978801_6_1782622 | 6 | 1782622 | 5.57E-07 | 0.29264 | 299 | 0.161441265 | 0.00149014 |
| SNP_A.K_1005038_6_6964239 | 6 | 6964239 | 5.58E-07 | 0.30268 | 299 | 0.161435094 | 0.00149014 |
| SNP_A.K_1004064_6_6823164 | 6 | 6823164 | 5.60E-07 | 0.30268 | 299 | 0.161412149 | 0.001493443 |
| SNP_A.K_1019370_6_9054292 | 6 | 9054292 | 5.62E-07 | 0.03344 | 299 | 0.161380418 | 0.00149876 |
| SNP_A.K_1009020_6_7457588 | 6 | 7457588 | 5.63E-07 | 0.07191 | 299 | 0.161370722 | 0.001499057 |
| SNP_A.K_993014_6_5698223 | 6 | 5698223 | 5.64E-07 | 0.19398 | 299 | 0.161355959 | 0.001500019 |
| SNP_A.K_1004398_6_6879577 | 6 | 6879577 | 5.65E-07 | 0.32107 | 299 | 0.161349754 | 0.001500019 |
| SNP_A.K_1005229_6_6995700 | 6 | 6995700 | 5.70E-07 | 0.29097 | 299 | 0.161288212 | 0.001512225 |
| SNP_A.K_978872_6_1793712 | 6 | 1793712 | 5.76E-07 | 0.32274 | 299 | 0.161220238 | 0.001523984 |
| SNP_A.K_978698_6_1766071 | 6 | 1766071 | 5.76E-07 | 0.30268 | 299 | 0.161219728 | 0.001523984 |
| SNP_A.K_1371525_8_7850968 | 8 | 7850968 | 5.77E-07 | 0.01672 | 299 | 0.16121237 | 0.001523984 |
| SNP_A.K_993841_6_5796393 | 6 | 5796393 | 5.82E-07 | 0.15217 | 299 | 0.16115914 | 0.001534453 |
| SNP_A.K_1004560_6_6904714 | 6 | 6904714 | 5.88E-07 | 0.30769 | 299 | 0.161085966 | 0.00154879 |
| SNP_A.K_1004751_6_6927221 | 6 | 6927221 | 5.89E-07 | 0.28261 | 299 | 0.161073741 | 0.00154879 |
| SNP_A.K_978768_6_1777747 | 6 | 1777747 | 5.89E-07 | 0.29933 | 299 | 0.161073247 | 0.00154879 |
| SNP_A.K_1504576_9_1146041 | 9 | 1146041 | 5.95E-07 | 0.04515 | 299 | 0.161012933 | 0.001561127 |
| SNP_A.K_1004888_6_6949456 | 6 | 6949456 | 6.02E-07 | 0.31104 | 299 | 0.160937846 | 0.00157639 |
| SNP_A.K_1005576_6_7027787 | 6 | 7027787 | 6.02E-07 | 0.28428 | 299 | 0.160932606 | 0.00157639 |

| | | | | | | | |
|---|---|---|---|---|---|---|---|
| SNP_A.K_1364969_8_6062335 | 8 | 6062335 | 6.04E-07 | 0.07358 | 299 | 0.16091682 | 0.001578204 |
| SNP_A.K_2081498_12_27527280 | 12 | 27527280 | 6.05E-07 | 0.02007 | 299 | 0.160905443 | 0.001578961 |
| SNP_A.K_1005335_6_7007691 | 6 | 7007691 | 6.06E-07 | 0.29599 | 299 | 0.160893611 | 0.00157983 |
| SNP_A.K_1006021_6_7096753 | 6 | 7096753 | 6.08E-07 | 0.27926 | 299 | 0.160871776 | 0.001583116 |
| SNP_A.K_1000536_6_6391746 | 6 | 6391746 | 6.09E-07 | 0.11204 | 299 | 0.160857773 | 0.001584517 |
| SNP_A.K_307066_2_7377265 | 2 | 7377265 | 6.11E-07 | 0.24916 | 299 | 0.160836869 | 0.001587592 |
| SNP_A.K_1004891_6_6949571 | 6 | 6949571 | 6.24E-07 | 0.30936 | 299 | 0.160699692 | 0.001619138 |
| SNP_A.K_1191357_7_6981949 | 7 | 6981949 | 6.25E-07 | 0.01171 | 299 | 0.160682593 | 0.001621345 |
| SNP_A.K_1364854_8_6049469 | 8 | 6049469 | 6.28E-07 | 0.07191 | 299 | 0.160658799 | 0.001625217 |
| SNP_A.K_1004213_6_6849830 | 6 | 6849830 | 6.31E-07 | 0.32441 | 299 | 0.160626069 | 0.001631323 |
| SNP_A.K_1608544_9_20775439 | 9 | 20775439 | 6.38E-07 | 0.23746 | 299 | 0.16055416 | 0.001645951 |
| SNP_A.K_1004577_6_6908410 | 6 | 6908410 | 6.38E-07 | 0.3311 | 299 | 0.160551295 | 0.001645951 |
| SNP_A.K_908231_5_17491203 | 5 | 17491203 | 6.43E-07 | 0.02508 | 299 | 0.160503084 | 0.00165605 |
| SNP_A.K_995318_6_5925965 | 6 | 5925965 | 6.47E-07 | 0.13545 | 299 | 0.160464928 | 0.001663661 |
| SNP_A.K_1000534_6_6391631 | 6 | 6391631 | 6.47E-07 | 0.1087 | 299 | 0.160456091 | 0.00166385 |
| SNP_A.K_993839_6_5796373 | 6 | 5796373 | 6.52E-07 | 0.15385 | 299 | 0.160405438 | 0.001674693 |
| SNP_A.K_1004725_6_6925686 | 6 | 6925686 | 6.55E-07 | 0.29097 | 299 | 0.16037787 | 0.001679682 |
| SNP_A.K_1599605_9_18569368 | 9 | 18569368 | 6.61E-07 | 0.04181 | 299 | 0.160323283 | 0.001691651 |
| SNP_A.K_991513_6_5536745 | 6 | 5536745 | 6.63E-07 | 0.15886 | 299 | 0.160302945 | 0.001694825 |
| SNP_A.K_993864_6_5797819 | 6 | 5797819 | 6.64E-07 | 0.16221 | 299 | 0.160294646 | 0.001694891 |
| SNP_A.K_978791_6_1782030 | 6 | 1782030 | 6.65E-07 | 0.29933 | 299 | 0.160277982 | 0.001697125 |
| SNP_A.K_1371573_8_7854648 | 8 | 7854648 | 6.66E-07 | 0.01003 | 299 | 0.160267092 | 0.001697868 |
| SNP_A.K_990266_6_4703810 | 6 | 4703810 | 6.86E-07 | 0.19064 | 299 | 0.160080159 | 0.00174488 |
| SNP_A.K_1004466_6_6884798 | 6 | 6884798 | 6.87E-07 | 0.30435 | 299 | 0.160066753 | 0.00174632 |
| SNP_A.K_1001130_6_6448378 | 6 | 6448378 | 6.91E-07 | 0.28763 | 299 | 0.160033402 | 0.001752116 |
| SNP_A.K_993406_6_5747985 | 6 | 5747985 | 6.91E-07 | 0.13545 | 299 | 0.160029076 | 0.001752116 |
| SNP_A.K_307351_2_7398399 | 2 | 7398399 | 6.96E-07 | 0.26589 | 299 | 0.159985483 | 0.001761672 |
| SNP_A.K_1004858_6_6936586 | 6 | 6936586 | 7.00E-07 | 0.2893 | 299 | 0.159946437 | 0.001770054 |
| SNP_A.K_998933_6_6235596 | 6 | 6235596 | 7.03E-07 | 0.14548 | 299 | 0.15991285 | 0.001774722 |
| SNP_A.K_1432894_8_16594506 | 8 | 16594506 | 7.04E-07 | 0.0786 | 299 | 0.159906685 | 0.001774722 |
| SNP_A.K_710043_4_8312041 | 4 | 8312041 | 7.04E-07 | 0.07358 | 299 | 0.15990536 | 0.001774722 |
| SNP_A.K_1005374_6_7010114 | 6 | 7010114 | 7.12E-07 | 0.24749 | 299 | 0.159838073 | 0.001790885 |
| SNP_A.K_993319_6_5738055 | 6 | 5738055 | 7.13E-07 | 0.12375 | 299 | 0.159821183 | 0.001793343 |
| SNP_A.K_1004183_6_6842183 | 6 | 6842183 | 7.14E-07 | 0.29933 | 299 | 0.159812507 | 0.001793554 |
| SNP_A.K_1004567_6_6906397 | 6 | 6906397 | 7.18E-07 | 0.29264 | 299 | 0.159782937 | 0.001798906 |
| SNP_A.K_990229_6_4701290 | 6 | 4701290 | 7.19E-07 | 0.199 | 299 | 0.159772328 | 0.001798906 |
| SNP_A.K_998934_6_6235624 | 6 | 6235624 | 7.19E-07 | 0.14381 | 299 | 0.159769351 | 0.001798906 |
| SNP_A.K_995312_6_5925280 | 6 | 5925280 | 7.21E-07 | 0.13712 | 299 | 0.159754669 | 0.00180078 |
| SNP_A.K_308772_2_7552377 | 2 | 7552377 | 7.23E-07 | 0.23244 | 299 | 0.159732774 | 0.001804646 |
| SNP_A.K_1608607_9_20794043 | 9 | 20794043 | 7.27E-07 | 0.24247 | 299 | 0.159700192 | 0.001810869 |
| SNP_A.K_2080242_12_27249340 | 12 | 27249340 | 7.27E-07 | 0.01003 | 299 | 0.159694557 | 0.001810869 |
| SNP_A.K_326781_2_10050198 | 2 | 10050198 | 7.28E-07 | 0.06522 | 299 | 0.159683999 | 0.001811626 |
| SNP_A.K_306774_2_7349593 | 2 | 7349593 | 7.37E-07 | 0.23244 | 299 | 0.159606947 | 0.001830889 |
| SNP_A.K_306985_2_7371938 | 2 | 7371938 | 7.40E-07 | 0.2408 | 299 | 0.159582539 | 0.001835539 |
| SNP_A.K_1003859_6_6797585 | 6 | 6797585 | 7.50E-07 | 0.01003 | 299 | 0.159494287 | 0.001858242 |
| SNP_A.K_978727_6_1771576 | 6 | 1771576 | 7.51E-07 | 0.36455 | 299 | 0.159480659 | 0.001859901 |
| SNP_A.K_1004859_6_6936639 | 6 | 6936639 | 7.57E-07 | 0.28763 | 299 | 0.159432358 | 0.001871455 |
| SNP_A.K_690102_4_4405254 | 4 | 4405254 | 7.58E-07 | 0.04682 | 299 | 0.159423249 | 0.001871838 |
| SNP_A.K_196591_1_31434679 | 1 | 31434679 | 7.65E-07 | 0.45987 | 299 | 0.159360207 | 0.00188772 |
| SNP_A.K_978143_6_1653092 | 6 | 1653092 | 7.71E-07 | 0.17559 | 299 | 0.159314976 | 0.001898565 |

| | | | | | | | |
|---|---|---|---|---|---|---|---|
| SNP_A.K_1006023_6_7097347 | 6 | 7097347 | 7.73E-07 | 0.28261 | 299 | 0.159294596 | 0.001902237 |
| SNP_A.K_1414837_8_13459023 | 8 | 13459023 | 7.77E-07 | 0.00669 | 299 | 0.15926482 | 0.001908658 |
| SNP_A.K_1004239_6_6853218 | 6 | 6853218 | 7.78E-07 | 0.30936 | 299 | 0.15925185 | 0.001910191 |
| SNP_A.K_1005950_6_7055238 | 6 | 7055238 | 7.83E-07 | 0.29766 | 299 | 0.159212507 | 0.001919449 |
| SNP_A.K_999038_6_6249481 | 6 | 6249481 | 7.86E-07 | 0.1388 | 299 | 0.159187752 | 0.001923096 |
| SNP_A.K_978741_6_1774311 | 6 | 1774311 | 7.86E-07 | 0.37458 | 299 | 0.159184699 | 0.001923096 |
| SNP_A.K_994710_6_5881365 | 6 | 5881365 | 7.88E-07 | 0.13043 | 299 | 0.159166436 | 0.001926209 |
| SNP_A.K_999620_6_6300986 | 6 | 6300986 | 7.95E-07 | 0.17893 | 299 | 0.159112652 | 0.001939835 |
| SNP_A.K_389624_2_19887421 | 2 | 19887421 | 8.06E-07 | 0.02341 | 299 | 0.159023082 | 0.001961545 |
| SNP_A.K_998927_6_6235106 | 6 | 6235106 | 8.06E-07 | 0.1388 | 299 | 0.159022922 | 0.001961545 |
| SNP_A.K_1004332_6_6868562 | 6 | 6868562 | 8.07E-07 | 0.30769 | 299 | 0.159012851 | 0.001961545 |
| SNP_A.K_994810_6_5891677 | 6 | 5891677 | 8.08E-07 | 0.27592 | 299 | 0.159006069 | 0.001961545 |
| SNP_A.K_1364551_8_6022345 | 8 | 6022345 | 8.09E-07 | 0.04348 | 299 | 0.159001635 | 0.001961545 |
| SNP_A.K_977942_6_1623289 | 6 | 1623289 | 8.10E-07 | 0.17726 | 299 | 0.158988005 | 0.001963349 |
| SNP_A.K_1433086_8_16617147 | 8 | 16617147 | 8.14E-07 | 0.08027 | 299 | 0.158958524 | 0.001969927 |
| SNP_A.K_1004790_6_6932649 | 6 | 6932649 | 8.15E-07 | 0.31773 | 299 | 0.158948036 | 0.001970797 |
| SNP_A.K_1005689_6_7034054 | 6 | 7034054 | 8.18E-07 | 0.14716 | 299 | 0.158923159 | 0.001976014 |
| SNP_A.K_709770_4_8285795 | 4 | 8285795 | 8.20E-07 | 0.11204 | 299 | 0.158911196 | 0.001977338 |
| SNP_A.K_307447_2_7408162 | 2 | 7408162 | 8.25E-07 | 0.26087 | 299 | 0.158872441 | 0.00198679 |
| SNP_A.K_673430_4_361606 | 4 | 361606 | 8.27E-07 | 0.04013 | 299 | 0.158853303 | 0.00199031 |
| SNP_A.K_992974_6_5693175 | 6 | 5693175 | 8.30E-07 | 0.2291 | 299 | 0.158829082 | 0.00199539 |
| SNP_A.K_307496_2_7413020 | 2 | 7413020 | 8.38E-07 | 0.23746 | 299 | 0.158770195 | 0.002011121 |
| SNP_A.K_1830578_11_10568064 | 11 | 10568064 | 8.40E-07 | 0.43144 | 299 | 0.158753935 | 0.002013807 |
| SNP_A.K_1004107_6_6834185 | 6 | 6834185 | 8.50E-07 | 0.30268 | 299 | 0.158678006 | 0.002034989 |
| SNP_A.K_978191_6_1659647 | 6 | 1659647 | 8.51E-07 | 0.17057 | 299 | 0.158663626 | 0.002037126 |
| SNP_A.K_1608576_9_20788401 | 9 | 20788401 | 8.65E-07 | 0.23411 | 299 | 0.158563861 | 0.002064796 |
| SNP_A.K_1830745_11_10577156 | 11 | 10577156 | 8.65E-07 | 0.13712 | 299 | 0.158560434 | 0.002064796 |
| SNP_A.K_307286_2_7391329 | 2 | 7391329 | 8.84E-07 | 0.23579 | 299 | 0.158417991 | 0.002107367 |
| SNP_A.K_1004143_6_6838009 | 6 | 6838009 | 8.85E-07 | 0.30936 | 299 | 0.158412094 | 0.002107367 |
| SNP_A.K_1005202_6_6993268 | 6 | 6993268 | 8.90E-07 | 0.24582 | 299 | 0.158371649 | 0.002118028 |
| SNP_A.K_1005304_6_7003659 | 6 | 7003659 | 8.99E-07 | 0.24247 | 299 | 0.158305737 | 0.002137061 |
| SNP_A.K_307002_2_7372595 | 2 | 7372595 | 9.01E-07 | 0.2408 | 299 | 0.158297318 | 0.002137376 |
| SNP_A.K_709177_4_8236729 | 4 | 8236729 | 9.08E-07 | 0.03177 | 299 | 0.158246815 | 0.002140745 |
| SNP_A.K_1003901_6_6802721 | 6 | 6802721 | 9.08E-07 | 0.31773 | 299 | 0.158244161 | 0.002140745 |
| SNP_A.K_1002949_6_6635225 | 6 | 6635225 | 9.08E-07 | 0.12207 | 299 | 0.15824237 | 0.002140745 |
| SNP_A.K_1005306_6_7004242 | 6 | 7004242 | 9.08E-07 | 0.28763 | 299 | 0.158241419 | 0.002140745 |
| SNP_A.K_710330_4_8336652 | 4 | 8336652 | 9.08E-07 | 0.07191 | 299 | 0.158240021 | 0.002140745 |
| SNP_A.K_999660_6_6303469 | 6 | 6303469 | 9.08E-07 | 0.19064 | 299 | 0.158239824 | 0.002140745 |
| SNP_A.K_306995_2_7372253 | 2 | 7372253 | 9.09E-07 | 0.23913 | 299 | 0.158235092 | 0.002140745 |
| SNP_A.K_1004579_6_6908451 | 6 | 6908451 | 9.14E-07 | 0.35452 | 299 | 0.158203442 | 0.002148711 |
| SNP_A.K_307239_2_7387199 | 2 | 7387199 | 9.20E-07 | 0.25753 | 299 | 0.158160872 | 0.002160319 |
| SNP_A.K_978782_6_1780293 | 6 | 1780293 | 9.26E-07 | 0.3194 | 299 | 0.158116959 | 0.002170814 |
| SNP_A.K_307397_2_7401005 | 2 | 7401005 | 9.27E-07 | 0.23746 | 299 | 0.158107041 | 0.002170814 |
| SNP_A.K_1005396_6_7011724 | 6 | 7011724 | 9.28E-07 | 0.26254 | 299 | 0.158100344 | 0.002170814 |
| SNP_A.K_307398_2_7401013 | 2 | 7401013 | 9.28E-07 | 0.23913 | 299 | 0.158099793 | 0.002170814 |
| SNP_A.K_997602_6_6111814 | 6 | 6111814 | 9.31E-07 | 0.1689 | 299 | 0.158081061 | 0.002174605 |
| SNP_A.K_1005399_6_7011794 | 6 | 7011794 | 9.42E-07 | 0.2709 | 299 | 0.158003661 | 0.002198064 |
| SNP_A.K_997445_6_6099901 | 6 | 6099901 | 9.58E-07 | 0.16388 | 299 | 0.157890531 | 0.002233971 |
| SNP_A.K_1003984_6_6812059 | 6 | 6812059 | 9.63E-07 | 0.30936 | 299 | 0.157856952 | 0.002240995 |
| SNP_A.K_307199_2_7384359 | 2 | 7384359 | 9.64E-07 | 0.24415 | 299 | 0.157855426 | 0.002240995 |

| | | | | | | | |
|---|---|---|---|---|---|---|---|
| SNP_A.K_1005890_6_7050562 | 6 | 7050562 | 9.68E-07 | 0.30435 | 299 | 0.157825808 | 0.002248672 |
| SNP_A.K_1005203_6_6993291 | 6 | 6993291 | 9.74E-07 | 0.29599 | 299 | 0.157783222 | 0.002258613 |
| SNP_A.K_994374_6_5853170 | 6 | 5853170 | 9.75E-07 | 0.17057 | 299 | 0.157777936 | 0.002258613 |
| SNP_A.K_307522_2_7415041 | 2 | 7415041 | 9.75E-07 | 0.24247 | 299 | 0.157775187 | 0.002258613 |
| SNP_A.K_710120_4_8317305 | 4 | 8317305 | 9.77E-07 | 0.09866 | 299 | 0.157762117 | 0.00226012 |
| SNP_A.K_978700_6_1766110 | 6 | 1766110 | 9.79E-07 | 0.30435 | 299 | 0.157752808 | 0.00226012 |
| SNP_A.K_1433091_8_16617234 | 8 | 16617234 | 9.79E-07 | 0.07692 | 299 | 0.157749083 | 0.00226012 |
| SNP_A.K_1005238_6_6997346 | 6 | 6997346 | 9.81E-07 | 0.33612 | 299 | 0.157739204 | 0.002261036 |
| SNP_A.K_994714_6_5881943 | 6 | 5881943 | 9.86E-07 | 0.1689 | 299 | 0.1577056 | 0.002270191 |
| SNP_A.K_1389908_8_10135213 | 8 | 10135213 | 9.97E-07 | 0.05351 | 299 | 0.157631766 | 0.002293479 |
| SNP_A.K_1364627_8_6030470 | 8 | 6030470 | 9.99E-07 | 0.01338 | 299 | 0.157618431 | 0.00229491 |
| SNP_A.K_1364938_8_6060610 | 8 | 6060610 | 1.00E-06 | 0.09197 | 299 | 0.157607954 | 0.00229491 |
| SNP_A.K_1004221_6_6851101 | 6 | 6851101 | 1.00E-06 | 0.30602 | 299 | 0.157606094 | 0.00229491 |
| SNP_A.K_1430891_8_16351157 | 8 | 16351157 | 1.01E-06 | 0.09197 | 299 | 0.157576405 | 0.002302836 |
| SNP_A.K_978800_6_1782566 | 6 | 1782566 | 1.01E-06 | 0.29766 | 299 | 0.157555375 | 0.00230773 |
| SNP_A.K_978943_6_1815277 | 6 | 1815277 | 1.02E-06 | 0.30769 | 299 | 0.157492976 | 0.002327343 |
| SNP_A.K_1433085_8_16617104 | 8 | 16617104 | 1.02E-06 | 0.0786 | 299 | 0.157472832 | 0.002331979 |
| SNP_A.K_710051_4_8312772 | 4 | 8312772 | 1.03E-06 | 0.07692 | 299 | 0.157451709 | 0.002336977 |
| SNP_A.K_978780_6_1780160 | 6 | 1780160 | 1.03E-06 | 0.35452 | 299 | 0.157435289 | 0.002339751 |
| SNP_A.K_999021_6_6248779 | 6 | 6248779 | 1.03E-06 | 0.13043 | 299 | 0.157429691 | 0.002339751 |
| SNP_A.K_998244_6_6186063 | 6 | 6186063 | 1.03E-06 | 0.09197 | 299 | 0.157411737 | 0.002343636 |
| SNP_A.K_389982_2_19960545 | 2 | 19960545 | 1.03E-06 | 0.01839 | 299 | 0.157394984 | 0.002347099 |
| SNP_A.K_1004075_6_6827225 | 6 | 6827225 | 1.04E-06 | 0.30769 | 299 | 0.157367635 | 0.002353726 |
| SNP_A.K_995290_6_5924002 | 6 | 5924002 | 1.04E-06 | 0.12709 | 299 | 0.157362373 | 0.002353726 |
| SNP_A.K_995641_6_5945460 | 6 | 5945460 | 1.05E-06 | 0.27926 | 299 | 0.15731682 | 0.002365378 |
| SNP_A.K_307025_2_7373332 | 2 | 7373332 | 1.05E-06 | 0.23579 | 299 | 0.15731597 | 0.002365378 |
| SNP_A.K_306724_2_7345368 | 2 | 7345368 | 1.05E-06 | 0.2408 | 299 | 0.157295962 | 0.00237007 |
| SNP_A.K_978723_6_1771530 | 6 | 1771530 | 1.06E-06 | 0.36789 | 299 | 0.157237872 | 0.002387328 |
| SNP_A.K_1005606_6_7029584 | 6 | 7029584 | 1.06E-06 | 0.13712 | 299 | 0.157231087 | 0.002387328 |
| SNP_A.K_1005573_6_7027557 | 6 | 7027557 | 1.06E-06 | 0.32776 | 299 | 0.157223235 | 0.002387328 |
| SNP_A.K_992723_6_5676274 | 6 | 5676274 | 1.06E-06 | 0.14214 | 299 | 0.157220391 | 0.002387328 |
| SNP_A.K_1371510_8_7850476 | 8 | 7850476 | 1.07E-06 | 0.01003 | 299 | 0.157201516 | 0.002391663 |
| SNP_A.K_992887_6_5683118 | 6 | 5683118 | 1.08E-06 | 0.12876 | 299 | 0.157110347 | 0.002422713 |
| SNP_A.K_994585_6_5874707 | 6 | 5874707 | 1.10E-06 | 0.25251 | 299 | 0.157001589 | 0.002457493 |
| SNP_A.K_1005402_6_7012375 | 6 | 7012375 | 1.10E-06 | 0.39298 | 299 | 0.156994755 | 0.002457493 |
| SNP_A.K_1005230_6_6996133 | 6 | 6996133 | 1.10E-06 | 0.23913 | 299 | 0.156994512 | 0.002457493 |
| SNP_A.K_1433147_8_16620252 | 8 | 16620252 | 1.10E-06 | 0.08194 | 299 | 0.156989339 | 0.002457493 |
| SNP_A.K_1004444_6_6882872 | 6 | 6882872 | 1.10E-06 | 0.30769 | 299 | 0.156981036 | 0.002457987 |
| SNP_A.K_710335_4_8336738 | 4 | 8336738 | 1.12E-06 | 0.08528 | 299 | 0.156900296 | 0.002483365 |
| SNP_A.K_306955_2_7370604 | 2 | 7370604 | 1.12E-06 | 0.24582 | 299 | 0.156900093 | 0.002483365 |
| SNP_A.K_1004336_6_6869601 | 6 | 6869601 | 1.12E-06 | 0.31773 | 299 | 0.156887525 | 0.002485498 |
| SNP_A.K_1005226_6_6995408 | 6 | 6995408 | 1.13E-06 | 0.24582 | 299 | 0.156844393 | 0.002499325 |
| SNP_A.K_978744_6_1774722 | 6 | 1774722 | 1.13E-06 | 0.30435 | 299 | 0.156833616 | 0.002500791 |
| SNP_A.K_1004629_6_6912352 | 6 | 6912352 | 1.13E-06 | 0.29766 | 299 | 0.1568234 | 0.002502046 |
| SNP_A.K_1371560_8_7852756 | 8 | 7852756 | 1.13E-06 | 0.02007 | 299 | 0.156790127 | 0.002512172 |
| SNP_A.K_1005320_6_7005799 | 6 | 7005799 | 1.14E-06 | 0.29933 | 299 | 0.156780981 | 0.002513026 |
| SNP_A.K_1005031_6_6963051 | 6 | 6963051 | 1.14E-06 | 0.30435 | 299 | 0.156751019 | 0.002521923 |
| SNP_A.K_1005490_6_7016833 | 6 | 7016833 | 1.15E-06 | 0.24749 | 299 | 0.156716321 | 0.002532694 |
| SNP_A.K_1005261_6_7000150 | 6 | 7000150 | 1.15E-06 | 0.2893 | 299 | 0.156679574 | 0.002544314 |
| SNP_A.K_990383_6_4712914 | 6 | 4712914 | 1.16E-06 | 0.20067 | 299 | 0.156649353 | 0.002553433 |

| | | | | | | | |
|---|---|---|---|---|---|---|---|
| SNP_A.K_978774_6_1779898 | 6 | 1779898 | 1.17E-06 | 0.29766 | 299 | 0.156615518 | 0.00256401 |
| SNP_A.K_1005197_6_6992009 | 6 | 6992009 | 1.17E-06 | 0.23746 | 299 | 0.156569686 | 0.002579378 |
| SNP_A.K_990353_6_4712175 | 6 | 4712175 | 1.18E-06 | 0.20234 | 299 | 0.156516642 | 0.002597715 |
| SNP_A.K_195535_1_31223944 | 1 | 31223944 | 1.19E-06 | 0.48829 | 299 | 0.156498206 | 0.00260233 |
| SNP_A.K_997541_6_6108212 | 6 | 6108212 | 1.19E-06 | 0.14381 | 299 | 0.156477776 | 0.002607753 |
| SNP_A.K_998495_6_6205629 | 6 | 6205629 | 1.20E-06 | 0.12709 | 299 | 0.156448859 | 0.002614558 |
| SNP_A.K_152999_1_23782002 | 1 | 23782002 | 1.20E-06 | 0.03679 | 299 | 0.156447092 | 0.002614558 |
| SNP_A.K_993888_6_5799435 | 6 | 5799435 | 1.20E-06 | 0.15552 | 299 | 0.156423508 | 0.002621285 |
| SNP_A.K_978128_6_1648960 | 6 | 1648960 | 1.21E-06 | 0.17559 | 299 | 0.156386306 | 0.002630839 |
| SNP_A.K_978882_6_1794586 | 6 | 1794586 | 1.21E-06 | 0.30435 | 299 | 0.156386142 | 0.002630839 |
| SNP_A.K_997881_6_6155417 | 6 | 6155417 | 1.22E-06 | 0.14716 | 299 | 0.156315141 | 0.002653378 |
| SNP_A.K_691242_4_4565343 | 4 | 4565343 | 1.22E-06 | 0.05017 | 299 | 0.156314047 | 0.002653378 |
| SNP_A.K_307422_2_7403442 | 2 | 7403442 | 1.22E-06 | 0.24247 | 299 | 0.156310131 | 0.002653378 |
| SNP_A.K_689899_4_4326361 | 4 | 4326361 | 1.22E-06 | 0.05853 | 299 | 0.156302345 | 0.002653781 |
| SNP_A.K_307424_2_7403784 | 2 | 7403784 | 1.22E-06 | 0.26254 | 299 | 0.156292707 | 0.002654942 |
| SNP_A.K_1433093_8_16617315 | 8 | 16617315 | 1.23E-06 | 0.08027 | 299 | 0.156284638 | 0.002655466 |
| SNP_A.K_62575_1_8994875 | 1 | 8994875 | 1.23E-06 | 0.01003 | 299 | 0.156250183 | 0.002666773 |
| SNP_A.K_997887_6_6155512 | 6 | 6155512 | 1.24E-06 | 0.14883 | 299 | 0.15621687 | 0.002677663 |
| SNP_A.K_1005378_6_7010978 | 6 | 7010978 | 1.24E-06 | 0.24582 | 299 | 0.156184597 | 0.002688172 |
| SNP_A.K_1364867_8_6050039 | 8 | 6050039 | 1.26E-06 | 0.08863 | 299 | 0.156123458 | 0.002705637 |
| SNP_A.K_1004882_6_6949017 | 6 | 6949017 | 1.26E-06 | 0.07023 | 299 | 0.156117114 | 0.002705637 |
| SNP_A.K_978693_6_1764700 | 6 | 1764700 | 1.26E-06 | 0.27258 | 299 | 0.15611366 | 0.002705637 |
| SNP_A.K_1001259_6_6459698 | 6 | 6459698 | 1.26E-06 | 0.29264 | 299 | 0.156113267 | 0.002705637 |
| SNP_A.K_307277_2_7390966 | 2 | 7390966 | 1.26E-06 | 0.23913 | 299 | 0.156108723 | 0.002705637 |
| SNP_A.K_307288_2_7391384 | 2 | 7391384 | 1.27E-06 | 0.23579 | 299 | 0.156062566 | 0.002716863 |
| SNP_A.K_978629_6_1747288 | 6 | 1747288 | 1.27E-06 | 0.28094 | 299 | 0.156062337 | 0.002716863 |
| SNP_A.K_307276_2_7390963 | 2 | 7390963 | 1.27E-06 | 0.23913 | 299 | 0.156057108 | 0.002716863 |
| SNP_A.K_1830665_11_10573230 | 11 | 10573230 | 1.27E-06 | 0.17224 | 299 | 0.156054931 | 0.002716863 |
| SNP_A.K_1000712_6_6413366 | 6 | 6413366 | 1.28E-06 | 0.35284 | 299 | 0.15602346 | 0.002727221 |
| SNP_A.K_1608555_9_20777132 | 9 | 20777132 | 1.28E-06 | 0.22575 | 299 | 0.156002169 | 0.00273275 |
| SNP_A.K_1004483_6_6886544 | 6 | 6886544 | 1.28E-06 | 0.30602 | 299 | 0.155996915 | 0.00273275 |
| SNP_A.K_978937_6_1814428 | 6 | 1814428 | 1.28E-06 | 0.32441 | 299 | 0.155989385 | 0.002733111 |
| SNP_A.K_978907_6_1800538 | 6 | 1800538 | 1.30E-06 | 0.30769 | 299 | 0.155891526 | 0.002771662 |
| SNP_A.K_998196_6_6184023 | 6 | 6184023 | 1.31E-06 | 0.12876 | 299 | 0.155877556 | 0.002774778 |
| SNP_A.K_1196546_7_7627910 | 7 | 7627910 | 1.31E-06 | 0.01171 | 299 | 0.155839172 | 0.002788338 |
| SNP_A.K_1433131_8_16618779 | 8 | 16618779 | 1.32E-06 | 0.08194 | 299 | 0.155827978 | 0.002790289 |
| SNP_A.K_689894_4_4325517 | 4 | 4325517 | 1.34E-06 | 0.05184 | 299 | 0.155721078 | 0.002833601 |
| SNP_A.K_1005227_6_6995623 | 6 | 6995623 | 1.34E-06 | 0.23579 | 299 | 0.155711919 | 0.002834704 |
| SNP_A.K_306837_2_7358494 | 2 | 7358494 | 1.34E-06 | 0.27258 | 299 | 0.155704921 | 0.002834869 |
| SNP_A.K_998179_6_6183266 | 6 | 6183266 | 1.35E-06 | 0.12709 | 299 | 0.155641664 | 0.00285965 |
| SNP_A.K_306859_2_7360914 | 2 | 7360914 | 1.36E-06 | 0.24415 | 299 | 0.155601204 | 0.002872489 |
| SNP_A.K_1004142_6_6837841 | 6 | 6837841 | 1.36E-06 | 0.31438 | 299 | 0.155599309 | 0.002872489 |
| SNP_A.K_1371552_8_7852191 | 8 | 7852191 | 1.37E-06 | 0.00669 | 299 | 0.155564099 | 0.002885149 |
| SNP_A.K_978885_6_1795798 | 6 | 1795798 | 1.37E-06 | 0.29431 | 299 | 0.155541147 | 0.002892415 |
| SNP_A.K_1004730_6_6925934 | 6 | 6925934 | 1.38E-06 | 0.29599 | 299 | 0.155522577 | 0.002897751 |
| SNP_A.K_1004391_6_6879086 | 6 | 6879086 | 1.38E-06 | 0.30936 | 299 | 0.155514268 | 0.002898526 |
| SNP_A.K_992731_6_5676620 | 6 | 5676620 | 1.39E-06 | 0.17391 | 299 | 0.155477542 | 0.002909272 |
| SNP_A.K_194836_1_30970379 | 1 | 30970379 | 1.39E-06 | 0.46823 | 299 | 0.155470302 | 0.002909272 |
| SNP_A.K_978862_6_1789336 | 6 | 1789336 | 1.39E-06 | 0.32274 | 299 | 0.15546116 | 0.002909272 |
| SNP_A.K_1000553_6_6394370 | 6 | 6394370 | 1.39E-06 | 0.11371 | 299 | 0.155459689 | 0.002909272 |

| | | | | | | | |
|---|---|---|---|---|---|---|---|
| SNP_A.K_978626_6_1746099 | 6 | 1746099 | 1.39E-06 | 0.27926 | 299 | 0.155457441 | 0.002909272 |
| SNP_A.K_307244_2_7387782 | 2 | 7387782 | 1.40E-06 | 0.23244 | 299 | 0.155426006 | 0.00292043 |
| SNP_A.K_978290_6_1669653 | 6 | 1669653 | 1.41E-06 | 0.24582 | 299 | 0.155361185 | 0.002946717 |
| SNP_A.K_1830646_11_10572639 | 11 | 10572639 | 1.42E-06 | 0.16054 | 299 | 0.155327777 | 0.002958923 |
| SNP_A.K_306730_2_7346016 | 2 | 7346016 | 1.43E-06 | 0.23746 | 299 | 0.155297983 | 0.002969534 |
| SNP_A.K_992651_6_5669933 | 6 | 5669933 | 1.43E-06 | 0.11873 | 299 | 0.155269247 | 0.002977897 |
| SNP_A.K_978881_6_1794548 | 6 | 1794548 | 1.43E-06 | 0.30769 | 299 | 0.155266691 | 0.002977897 |
| SNP_A.K_977864_6_1604765 | 6 | 1604765 | 1.44E-06 | 0.16388 | 299 | 0.155246585 | 0.002984138 |
| SNP_A.K_1364543_8_6019942 | 8 | 6019942 | 1.44E-06 | 0.08863 | 299 | 0.155232565 | 0.002987598 |
| SNP_A.K_1365025_8_6067016 | 8 | 6067016 | 1.45E-06 | 0.09197 | 299 | 0.155194053 | 0.003000169 |
| SNP_A.K_992743_6_5677380 | 6 | 5677380 | 1.45E-06 | 0.14381 | 299 | 0.155192311 | 0.003000169 |
| SNP_A.K_709961_4_8303580 | 4 | 8303580 | 1.46E-06 | 0.07525 | 299 | 0.15515565 | 0.003014132 |
| SNP_A.K_1001186_6_6453319 | 6 | 6453319 | 1.46E-06 | 0.2893 | 299 | 0.155139525 | 0.003018617 |
| SNP_A.K_1830651_11_10572843 | 11 | 10572843 | 1.47E-06 | 0.14047 | 299 | 0.155102465 | 0.003032859 |
| SNP_A.K_678443_4_986682 | 4 | 986682 | 1.48E-06 | 0.07358 | 299 | 0.155063385 | 0.003048119 |
| SNP_A.K_978946_6_1817571 | 6 | 1817571 | 1.49E-06 | 0.30268 | 299 | 0.155019387 | 0.003065778 |
| SNP_A.K_978875_6_1793947 | 6 | 1793947 | 1.50E-06 | 0.30268 | 299 | 0.154988466 | 0.003077347 |
| SNP_A.K_1004126_6_6835977 | 6 | 6835977 | 1.51E-06 | 0.32441 | 299 | 0.154947156 | 0.003093905 |
| SNP_A.K_1197718_7_7779740 | 7 | 7779740 | 1.51E-06 | 0.01171 | 299 | 0.154927736 | 0.0031001 |
| SNP_A.K_708903_4_8123035 | 4 | 8123035 | 1.51E-06 | 0.08696 | 299 | 0.154918742 | 0.003101333 |
| SNP_A.K_710336_4_8336745 | 4 | 8336745 | 1.52E-06 | 0.08696 | 299 | 0.154886278 | 0.00311379 |
| SNP_A.K_978823_6_1785260 | 6 | 1785260 | 1.52E-06 | 0.30602 | 299 | 0.154867338 | 0.003119155 |
| SNP_A.K_326118_2_9978595 | 2 | 9978595 | 1.53E-06 | 0.12709 | 299 | 0.154862302 | 0.003119155 |
| SNP_A.K_710482_4_8424260 | 4 | 8424260 | 1.54E-06 | 0.07692 | 299 | 0.15481411 | 0.00313928 |
| SNP_A.K_1383210_8_9134678 | 8 | 9134678 | 1.54E-06 | 0.11204 | 299 | 0.15479505 | 0.003145412 |
| SNP_A.K_998176_6_6183195 | 6 | 6183195 | 1.55E-06 | 0.13043 | 299 | 0.154756833 | 0.003160861 |
| SNP_A.K_1433129_8_16618538 | 8 | 16618538 | 1.57E-06 | 0.0786 | 299 | 0.154690044 | 0.003190384 |
| SNP_A.K_990338_6_4711401 | 6 | 4711401 | 1.57E-06 | 0.20569 | 299 | 0.154660058 | 0.003202005 |
| SNP_A.K_1364542_8_6019669 | 8 | 6019669 | 1.58E-06 | 0.08194 | 299 | 0.154638503 | 0.003209505 |
| SNP_A.K_307389_2_7400292 | 2 | 7400292 | 1.58E-06 | 0.24247 | 299 | 0.154627795 | 0.003210387 |
| SNP_A.K_992698_6_5672329 | 6 | 5672329 | 1.58E-06 | 0.19064 | 299 | 0.154624038 | 0.003210387 |
| SNP_A.K_978708_6_1769211 | 6 | 1769211 | 1.58E-06 | 0.32274 | 299 | 0.154616079 | 0.003211192 |
| SNP_A.K_1364538_8_6017574 | 8 | 6017574 | 1.59E-06 | 0.08696 | 299 | 0.15460744 | 0.003212335 |
| SNP_A.K_994756_6_5887341 | 6 | 5887341 | 1.59E-06 | 0.27759 | 299 | 0.15459842 | 0.003213671 |
| SNP_A.K_978699_6_1766108 | 6 | 1766108 | 1.60E-06 | 0.30769 | 299 | 0.154557295 | 0.003230934 |
| SNP_A.K_307032_2_7374120 | 2 | 7374120 | 1.61E-06 | 0.23579 | 299 | 0.154522294 | 0.003245234 |
| SNP_A.K_1608543_9_20775423 | 9 | 20775423 | 1.61E-06 | 0.2408 | 299 | 0.154493756 | 0.00325636 |
| SNP_A.K_254326_1_41018019 | 1 | 41018019 | 1.62E-06 | 0.03344 | 299 | 0.154476199 | 0.003262011 |
| SNP_A.K_1005267_6_7001492 | 6 | 7001492 | 1.62E-06 | 0.28261 | 299 | 0.154452524 | 0.003270753 |
| SNP_A.K_978886_6_1796372 | 6 | 1796372 | 1.65E-06 | 0.31104 | 299 | 0.154365261 | 0.003309197 |
| SNP_A.K_1005224_6_6995267 | 6 | 6995267 | 1.65E-06 | 0.24749 | 299 | 0.154364055 | 0.003309197 |
| SNP_A.K_978602_6_1738545 | 6 | 1738545 | 1.65E-06 | 0.25084 | 299 | 0.154355605 | 0.003310309 |
| SNP_A.K_995335_6_5926428 | 6 | 5926428 | 1.66E-06 | 0.13712 | 299 | 0.154312541 | 0.003329117 |
| SNP_A.K_998075_6_6169972 | 6 | 6169972 | 1.67E-06 | 0.26254 | 299 | 0.15427386 | 0.003345778 |
| SNP_A.K_993473_6_5755450 | 6 | 5755450 | 1.68E-06 | 0.13712 | 299 | 0.154217072 | 0.003371914 |
| SNP_A.K_978338_6_1679798 | 6 | 1679798 | 1.69E-06 | 0.25753 | 299 | 0.154208783 | 0.003372977 |
| SNP_A.K_1004443_6_6882848 | 6 | 6882848 | 1.70E-06 | 0.30936 | 299 | 0.154164554 | 0.003392767 |
| SNP_A.K_978636_6_1748174 | 6 | 1748174 | 1.70E-06 | 0.2709 | 299 | 0.154148986 | 0.003397649 |
| SNP_A.K_1830662_11_10573134 | 11 | 10573134 | 1.72E-06 | 0.16054 | 299 | 0.154099916 | 0.003420141 |
| SNP_A.K_1004559_6_6904539 | 6 | 6904539 | 1.72E-06 | 0.29933 | 299 | 0.154086433 | 0.003423151 |

| | | | | | | | |
|---|---|---|---|---|---|---|---|
| SNP_A.K_978011_6_1636239 | 6 | 1636239 | 1.72E-06 | 0.17726 | 299 | 0.154079808 | 0.003423151 |
| SNP_A.K_1004337_6_6869619 | 6 | 6869619 | 1.72E-06 | 0.32107 | 299 | 0.154075571 | 0.003423151 |
| SNP_A.K_978822_6_1785046 | 6 | 1785046 | 1.73E-06 | 0.30268 | 299 | 0.154051235 | 0.003431253 |
| SNP_A.K_993616_6_5779663 | 6 | 5779663 | 1.73E-06 | 0.12542 | 299 | 0.154046176 | 0.003431253 |
| SNP_A.K_306996_2_7372370 | 2 | 7372370 | 1.73E-06 | 0.23746 | 299 | 0.154032639 | 0.003431253 |
| SNP_A.K_978873_6_1793728 | 6 | 1793728 | 1.73E-06 | 0.33278 | 299 | 0.154031376 | 0.003431253 |
| SNP_A.K_1183403_7_4541227 | 7 | 4541227 | 1.73E-06 | 0.4097 | 299 | 0.154029287 | 0.003431253 |
| SNP_A.K_1910769_11_22488147 | 11 | 22488147 | 1.74E-06 | 0.04013 | 299 | 0.154016686 | 0.003434654 |
| SNP_A.K_307380_2_7399803 | 2 | 7399803 | 1.76E-06 | 0.2408 | 299 | 0.153939618 | 0.00347237 |
| SNP_A.K_998878_6_6230113 | 6 | 6230113 | 1.77E-06 | 0.26087 | 299 | 0.153907141 | 0.003486477 |
| SNP_A.K_997949_6_6162004 | 6 | 6162004 | 1.77E-06 | 0.17559 | 299 | 0.153895994 | 0.003487287 |
| SNP_A.K_1004543_6_6899633 | 6 | 6899633 | 1.77E-06 | 0.30435 | 299 | 0.153893337 | 0.003487287 |
| SNP_A.K_1364935_8_6060324 | 8 | 6060324 | 1.77E-06 | 0.09365 | 299 | 0.15388321 | 0.003489431 |
| SNP_A.K_1830640_11_10572252 | 11 | 10572252 | 1.78E-06 | 0.16555 | 299 | 0.153861277 | 0.003497934 |
| SNP_A.K_325494_2_9897261 | 2 | 9897261 | 1.79E-06 | 0.02676 | 299 | 0.153824126 | 0.003514693 |
| SNP_A.K_1004397_6_6879525 | 6 | 6879525 | 1.81E-06 | 0.31271 | 299 | 0.153765842 | 0.003543053 |
| SNP_A.K_307243_2_7387725 | 2 | 7387725 | 1.82E-06 | 0.24415 | 299 | 0.153714534 | 0.003567811 |
| SNP_A.K_998178_6_6183220 | 6 | 6183220 | 1.83E-06 | 0.13043 | 299 | 0.153691073 | 0.003577364 |
| SNP_A.K_978920_6_1804367 | 6 | 1804367 | 1.83E-06 | 0.30769 | 299 | 0.153680162 | 0.003579827 |
| SNP_A.K_993806_6_5794260 | 6 | 5794260 | 1.83E-06 | 0.06856 | 299 | 0.153674407 | 0.003579827 |
| SNP_A.K_1004611_6_6910377 | 6 | 6910377 | 1.84E-06 | 0.28261 | 299 | 0.153660459 | 0.003584164 |
| SNP_A.K_990243_6_4702847 | 6 | 4702847 | 1.85E-06 | 0.21237 | 299 | 0.153618122 | 0.003604244 |
| SNP_A.K_306808_2_7352837 | 2 | 7352837 | 1.86E-06 | 0.24247 | 299 | 0.153577043 | 0.003623737 |
| SNP_A.K_308786_2_7553821 | 2 | 7553821 | 1.86E-06 | 0.22241 | 299 | 0.153570656 | 0.003623913 |
| SNP_A.K_1003928_6_6806023 | 6 | 6806023 | 1.88E-06 | 0.31773 | 299 | 0.153518839 | 0.003647557 |
| SNP_A.K_1390329_8_10181900 | 8 | 10181900 | 1.88E-06 | 0.04348 | 299 | 0.153516334 | 0.003647557 |
| SNP_A.K_1004470_6_6884885 | 6 | 6884885 | 1.89E-06 | 0.31104 | 299 | 0.153482077 | 0.003663277 |
| SNP_A.K_1627148_10_1890862 | 10 | 1890862 | 1.89E-06 | 0.02676 | 299 | 0.153476329 | 0.003663277 |
| SNP_A.K_307381_2_7399817 | 2 | 7399817 | 1.90E-06 | 0.25585 | 299 | 0.153445284 | 0.003677422 |
| SNP_A.K_1004185_6_6842554 | 6 | 6842554 | 1.90E-06 | 0.32274 | 299 | 0.153426623 | 0.003683497 |
| SNP_A.K_1608611_9_20794190 | 9 | 20794190 | 1.90E-06 | 0.23077 | 299 | 0.153422506 | 0.003683497 |
| SNP_A.K_1004607_6_6910020 | 6 | 6910020 | 1.91E-06 | 0.31271 | 299 | 0.153401281 | 0.003691691 |
| SNP_A.K_1005284_6_7002464 | 6 | 7002464 | 1.91E-06 | 0.27592 | 299 | 0.153396045 | 0.003691691 |
| SNP_A.K_978931_6_1808335 | 6 | 1808335 | 1.92E-06 | 0.32943 | 299 | 0.153369449 | 0.003699428 |
| SNP_A.K_710337_4_8336757 | 4 | 8336757 | 1.92E-06 | 0.08528 | 299 | 0.153368878 | 0.003699428 |
| SNP_A.K_978771_6_1779134 | 6 | 1779134 | 1.92E-06 | 0.2893 | 299 | 0.153364439 | 0.003699428 |
| SNP_A.K_978749_6_1775031 | 6 | 1775031 | 1.93E-06 | 0.39298 | 299 | 0.153345624 | 0.003706745 |
| SNP_A.K_1004606_6_6909984 | 6 | 6909984 | 1.93E-06 | 0.31605 | 299 | 0.153337316 | 0.003708069 |
| SNP_A.K_978683_6_1760683 | 6 | 1760683 | 1.93E-06 | 0.27425 | 299 | 0.153319165 | 0.00371503 |
| SNP_A.K_978656_6_1750694 | 6 | 1750694 | 1.94E-06 | 0.30602 | 299 | 0.153296751 | 0.003724455 |
| SNP_A.K_996337_6_5988903 | 6 | 5988903 | 1.94E-06 | 0.27592 | 299 | 0.153289022 | 0.003725461 |
| SNP_A.K_1005290_6_7002660 | 6 | 7002660 | 1.95E-06 | 0.23077 | 299 | 0.153272694 | 0.003731417 |
| SNP_A.K_978456_6_1694918 | 6 | 1694918 | 1.95E-06 | 0.26421 | 299 | 0.153262804 | 0.003733676 |
| SNP_A.K_1003971_6_6809950 | 6 | 6809950 | 1.96E-06 | 0.31605 | 299 | 0.153246188 | 0.003738741 |
| SNP_A.K_995813_6_5951565 | 6 | 5951565 | 1.96E-06 | 0.30268 | 299 | 0.153242101 | 0.003738741 |
| SNP_A.K_389619_2_19887270 | 2 | 19887270 | 1.96E-06 | 0.04181 | 299 | 0.153231054 | 0.003741683 |
| SNP_A.K_996036_6_5958801 | 6 | 5958801 | 1.97E-06 | 0.2709 | 299 | 0.153193184 | 0.003759966 |
| SNP_A.K_978464_6_1696320 | 6 | 1696320 | 1.97E-06 | 0.24916 | 299 | 0.153187565 | 0.003759966 |
| SNP_A.K_978850_6_1787636 | 6 | 1787636 | 1.99E-06 | 0.35619 | 299 | 0.153133268 | 0.003788116 |
| SNP_A.K_990245_6_4702919 | 6 | 4702919 | 2.00E-06 | 0.21237 | 299 | 0.15309818 | 0.003803057 |

| | | | | | | | |
|---|---|---|---|---|---|---|---|
| SNP_A.K_1004608_6_6910068 | 6 | 6910068 | 2.00E-06 | 0.29431 | 299 | 0.1530959 | 0.003803057 |
| SNP_A.K_1004037_6_6820951 | 6 | 6820951 | 2.01E-06 | 0.29766 | 299 | 0.153073553 | 0.003810616 |
| SNP_A.K_1195882_7_7549486 | 7 | 7549486 | 2.01E-06 | 0.01171 | 299 | 0.153071204 | 0.003810616 |
| SNP_A.K_978748_6_1775022 | 6 | 1775022 | 2.02E-06 | 0.36957 | 299 | 0.153044637 | 0.003822781 |
| SNP_A.K_1006002_6_7095126 | 6 | 7095126 | 2.04E-06 | 0.27926 | 299 | 0.152981179 | 0.003856873 |
| SNP_A.K_1784823_11_3799601 | 11 | 3799601 | 2.04E-06 | 0.01003 | 299 | 0.152968963 | 0.003860638 |
| SNP_A.K_1004071_6_6825613 | 6 | 6825613 | 2.05E-06 | 0.29933 | 299 | 0.152945591 | 0.003871067 |
| SNP_A.K_1005408_6_7012542 | 6 | 7012542 | 2.05E-06 | 0.37458 | 299 | 0.152939536 | 0.003871171 |
| SNP_A.K_1858455_11_15010731 | 11 | 15010731 | 2.06E-06 | 0.0903 | 299 | 0.152925199 | 0.003876228 |
| SNP_A.K_307148_2_7381718 | 2 | 7381718 | 2.06E-06 | 0.24247 | 299 | 0.152915544 | 0.003878493 |
| SNP_A.K_307089_2_7378365 | 2 | 7378365 | 2.06E-06 | 0.25251 | 299 | 0.152906377 | 0.00388047 |
| SNP_A.K_709873_4_8300080 | 4 | 8300080 | 2.08E-06 | 0.06689 | 299 | 0.152855524 | 0.0039075 |
| SNP_A.K_306722_2_7345318 | 2 | 7345318 | 2.08E-06 | 0.24582 | 299 | 0.152849364 | 0.003907686 |
| SNP_A.K_1602001_9_19102756 | 9 | 19102756 | 2.08E-06 | 0.11037 | 299 | 0.152836979 | 0.003911629 |
| SNP_A.K_990307_6_4709831 | 6 | 4709831 | 2.09E-06 | 0.20569 | 299 | 0.152815239 | 0.003921236 |
| SNP_A.K_306908_2_7364967 | 2 | 7364967 | 2.12E-06 | 0.2408 | 299 | 0.15272388 | 0.003968569 |
| SNP_A.K_306811_2_7352987 | 2 | 7352987 | 2.12E-06 | 0.23913 | 299 | 0.152720496 | 0.003968569 |
| SNP_A.K_1830586_11_10568737 | 11 | 10568737 | 2.12E-06 | 0.1689 | 299 | 0.152719987 | 0.003968569 |
| SNP_A.K_2078892_12_27033066 | 12 | 27033066 | 2.13E-06 | 0.01171 | 299 | 0.152697111 | 0.003979028 |
| SNP_A.K_710052_4_8312782 | 4 | 8312782 | 2.14E-06 | 0.0786 | 299 | 0.152666518 | 0.003994273 |
| SNP_A.K_995957_6_5955892 | 6 | 5955892 | 2.14E-06 | 0.29933 | 299 | 0.152656886 | 0.003996631 |
| SNP_A.K_1004389_6_6878989 | 6 | 6878989 | 2.16E-06 | 0.31773 | 299 | 0.15260891 | 0.004022728 |
| SNP_A.K_1364553_8_6022926 | 8 | 6022926 | 2.16E-06 | 0.04682 | 299 | 0.152599433 | 0.004025013 |
| SNP_A.K_977891_6_1613741 | 6 | 1613741 | 2.18E-06 | 0.15886 | 299 | 0.152552266 | 0.004050799 |
| SNP_A.K_1390218_8_10163233 | 8 | 10163233 | 2.18E-06 | 0.04682 | 299 | 0.152537712 | 0.004052945 |
| SNP_A.K_977897_6_1616556 | 6 | 1616556 | 2.18E-06 | 0.23913 | 299 | 0.152537265 | 0.004052945 |
| SNP_A.K_1005520_6_7018966 | 6 | 7018966 | 2.19E-06 | 0.28261 | 299 | 0.152513731 | 0.004063725 |
| SNP_A.K_997271_6_6082627 | 6 | 6082627 | 2.19E-06 | 0.13712 | 299 | 0.15250851 | 0.004063725 |
| SNP_A.K_307442_2_7407713 | 2 | 7407713 | 2.20E-06 | 0.2592 | 299 | 0.152496114 | 0.004067886 |
| SNP_A.K_456553_2_33063997 | 2 | 33063997 | 2.21E-06 | 0.02676 | 299 | 0.152465079 | 0.004083788 |
| SNP_A.K_709845_4_8299480 | 4 | 8299480 | 2.21E-06 | 0.06856 | 299 | 0.152452578 | 0.004088043 |
| SNP_A.K_978860_6_1789007 | 6 | 1789007 | 2.22E-06 | 0.30936 | 299 | 0.152415635 | 0.004107776 |
| SNP_A.K_978851_6_1787641 | 6 | 1787641 | 2.23E-06 | 0.3612 | 299 | 0.152386913 | 0.004122374 |
| SNP_A.K_678702_4_994397 | 4 | 994397 | 2.24E-06 | 0.08863 | 299 | 0.15237503 | 0.004126286 |
| SNP_A.K_882922_5_7668003 | 5 | 7668003 | 2.24E-06 | 0.02676 | 299 | 0.152365281 | 0.004128844 |
| SNP_A.K_1005294_6_7002697 | 6 | 7002697 | 2.25E-06 | 0.28094 | 299 | 0.152351983 | 0.004133672 |
| SNP_A.K_307352_2_7398452 | 2 | 7398452 | 2.25E-06 | 0.26923 | 299 | 0.152345961 | 0.004133862 |
| SNP_A.K_1599653_9_18583529 | 9 | 18583529 | 2.25E-06 | 0.06355 | 299 | 0.152339363 | 0.004134423 |
| SNP_A.K_978855_6_1788384 | 6 | 1788384 | 2.27E-06 | 0.31271 | 299 | 0.152285534 | 0.004165254 |
| SNP_A.K_1003862_6_6798743 | 6 | 6798743 | 2.28E-06 | 0.17559 | 299 | 0.152261436 | 0.004177099 |
| SNP_A.K_1004890_6_6949544 | 6 | 6949544 | 2.29E-06 | 0.34114 | 299 | 0.152239092 | 0.004186795 |
| SNP_A.K_978646_6_1749357 | 6 | 1749357 | 2.29E-06 | 0.34615 | 299 | 0.152230287 | 0.004186795 |
| SNP_A.K_325546_2_9901614 | 2 | 9901614 | 2.29E-06 | 0.03846 | 299 | 0.152229048 | 0.004186795 |
| SNP_A.K_990585_6_4726935 | 6 | 4726935 | 2.29E-06 | 0.11706 | 299 | 0.15222364 | 0.004186795 |
| SNP_A.K_998065_6_6168312 | 6 | 6168312 | 2.29E-06 | 0.16555 | 299 | 0.152215683 | 0.004188266 |
| SNP_A.K_1364931_8_6060126 | 8 | 6060126 | 2.30E-06 | 0.08863 | 299 | 0.152187419 | 0.004202901 |
| SNP_A.K_1608676_9_20815198 | 9 | 20815198 | 2.31E-06 | 0.24247 | 299 | 0.152159872 | 0.004217124 |
| SNP_A.K_526677_3_10964889 | 3 | 10964889 | 2.32E-06 | 0.01003 | 299 | 0.152131984 | 0.00422423 |
| SNP_A.K_528922_3_11353842 | 3 | 11353842 | 2.32E-06 | 0.01003 | 299 | 0.152131984 | 0.00422423 |
| SNP_A.K_528340_3_11270505 | 3 | 11270505 | 2.32E-06 | 0.01003 | 299 | 0.152131984 | 0.00422423 |

| | | | | | | | |
|---|---|---|---|---|---|---|---|
| SNP_A.K_978277_6_1666751 | 6 | 1666751 | 2.34E-06 | 0.1806 | 299 | 0.15209696 | 0.004243437 |
| SNP_A.K_978601_6_1738375 | 6 | 1738375 | 2.35E-06 | 0.2592 | 299 | 0.152060704 | 0.004259464 |
| SNP_A.K_307150_2_7381999 | 2 | 7381999 | 2.35E-06 | 0.24582 | 299 | 0.152052219 | 0.004259464 |
| SNP_A.K_307451_2_7408539 | 2 | 7408539 | 2.35E-06 | 0.24247 | 299 | 0.152052173 | 0.004259464 |
| SNP_A.K_306732_2_7346355 | 2 | 7346355 | 2.35E-06 | 0.2408 | 299 | 0.152049997 | 0.004259464 |
| SNP_A.K_86565_1_11815553 | 1 | 11815553 | 2.38E-06 | 0.06522 | 299 | 0.151976496 | 0.004304359 |
| SNP_A.K_1005483_6_7016656 | 6 | 7016656 | 2.39E-06 | 0.29599 | 299 | 0.151952604 | 0.004315348 |
| SNP_A.K_307145_2_7381643 | 2 | 7381643 | 2.39E-06 | 0.24247 | 299 | 0.15194728 | 0.004315348 |
| SNP_A.K_978660_6_1751310 | 6 | 1751310 | 2.39E-06 | 0.27592 | 299 | 0.151942515 | 0.004315348 |
| SNP_A.K_978630_6_1747678 | 6 | 1747678 | 2.39E-06 | 0.28595 | 299 | 0.151937538 | 0.004315348 |
| SNP_A.K_1004204_6_6847599 | 6 | 6847599 | 2.40E-06 | 0.29599 | 299 | 0.15191918 | 0.004321094 |
| SNP_A.K_497656_3_5537385 | 3 | 5537385 | 2.40E-06 | 0.03846 | 299 | 0.151917727 | 0.004321094 |
| SNP_A.K_1004217_6_6849914 | 6 | 6849914 | 2.41E-06 | 0.31773 | 299 | 0.151908054 | 0.004323819 |
| SNP_A.K_978627_6_1746820 | 6 | 1746820 | 2.43E-06 | 0.26589 | 299 | 0.151853506 | 0.004356652 |
| SNP_A.K_1503487_9_1028218 | 9 | 1028218 | 2.44E-06 | 0.04515 | 299 | 0.151824603 | 0.00437238 |
| SNP_A.K_1005268_6_7001638 | 6 | 7001638 | 2.44E-06 | 0.27926 | 299 | 0.151815501 | 0.004373168 |
| SNP_A.K_978717_6_1771029 | 6 | 1771029 | 2.44E-06 | 0.38963 | 299 | 0.151812289 | 0.004373168 |
| SNP_A.K_1004791_6_6932852 | 6 | 6932852 | 2.45E-06 | 0.29933 | 299 | 0.151799913 | 0.004377772 |
| SNP_A.K_307273_2_7390634 | 2 | 7390634 | 2.45E-06 | 0.25084 | 299 | 0.151793119 | 0.004378605 |
| SNP_A.K_1830661_11_10573117 | 11 | 10573117 | 2.45E-06 | 0.16221 | 299 | 0.151786284 | 0.00437947 |
| SNP_A.K_995253_6_5921522 | 6 | 5921522 | 2.46E-06 | 0.12375 | 299 | 0.151773251 | 0.004384536 |
| SNP_A.K_306861_2_7361400 | 2 | 7361400 | 2.47E-06 | 0.26421 | 299 | 0.151743021 | 0.004401292 |
| SNP_A.K_978959_6_1831190 | 6 | 1831190 | 2.47E-06 | 0.3311 | 299 | 0.151736866 | 0.004401709 |
| SNP_A.K_990344_6_4711747 | 6 | 4711747 | 2.49E-06 | 0.20736 | 299 | 0.1516765 | 0.004435372 |
| SNP_A.K_990345_6_4711750 | 6 | 4711750 | 2.49E-06 | 0.20736 | 299 | 0.1516765 | 0.004435372 |
| SNP_A.K_690542_4_4448925 | 4 | 4448925 | 2.50E-06 | 0.0301 | 299 | 0.151669334 | 0.004436495 |
| SNP_A.K_1364999_8_6065775 | 8 | 6065775 | 2.51E-06 | 0.04515 | 299 | 0.15164175 | 0.004451649 |
| SNP_A.K_196625_1_31452357 | 1 | 31452357 | 2.52E-06 | 0.47324 | 299 | 0.151616014 | 0.004465582 |
| SNP_A.K_710042_4_8311970 | 4 | 8311970 | 2.52E-06 | 0.07525 | 299 | 0.151599341 | 0.004468043 |
| SNP_A.K_978637_6_1748297 | 6 | 1748297 | 2.52E-06 | 0.2893 | 299 | 0.151598537 | 0.004468043 |
| SNP_A.K_197054_1_31549174 | 1 | 31549174 | 2.52E-06 | 0.02676 | 299 | 0.151595925 | 0.004468043 |
| SNP_A.K_1005027_6_6962917 | 6 | 6962917 | 2.55E-06 | 0.31773 | 299 | 0.151532671 | 0.004508112 |
| SNP_A.K_1602090_9_19122093 | 9 | 19122093 | 2.56E-06 | 0.11706 | 299 | 0.151517647 | 0.004514759 |
| SNP_A.K_194653_1_30906130 | 1 | 30906130 | 2.56E-06 | 0.49498 | 299 | 0.151505635 | 0.004519314 |
| SNP_A.K_977945_6_1623530 | 6 | 1623530 | 2.57E-06 | 0.18227 | 299 | 0.151484021 | 0.004530598 |
| SNP_A.K_999571_6_6296890 | 6 | 6296890 | 2.58E-06 | 0.13545 | 299 | 0.151466142 | 0.004539292 |
| SNP_A.K_1005537_6_7020016 | 6 | 7020016 | 2.58E-06 | 0.30936 | 299 | 0.151459003 | 0.00454046 |
| SNP_A.K_1608677_9_20815201 | 9 | 20815201 | 2.59E-06 | 0.24247 | 299 | 0.151434542 | 0.004553811 |
| SNP_A.K_1004692_6_6918929 | 6 | 6918929 | 2.59E-06 | 0.38462 | 299 | 0.151426881 | 0.004554046 |
| SNP_A.K_306718_2_7345002 | 2 | 7345002 | 2.59E-06 | 0.26923 | 299 | 0.151423283 | 0.004554046 |
| SNP_A.K_307056_2_7376283 | 2 | 7376283 | 2.61E-06 | 0.2408 | 299 | 0.151377902 | 0.004582247 |
| SNP_A.K_326915_2_10070580 | 2 | 10070580 | 2.62E-06 | 0.05686 | 299 | 0.151365369 | 0.004587268 |
| SNP_A.K_307149_2_7381995 | 2 | 7381995 | 2.63E-06 | 0.24582 | 299 | 0.151333411 | 0.004606113 |
| SNP_A.K_978020_6_1638628 | 6 | 1638628 | 2.63E-06 | 0.16221 | 299 | 0.151319482 | 0.004610211 |
| SNP_A.K_307283_2_7391152 | 2 | 7391152 | 2.64E-06 | 0.24247 | 299 | 0.151316782 | 0.004610211 |
| SNP_A.K_691232_4_4563130 | 4 | 4563130 | 2.65E-06 | 0.07191 | 299 | 0.151293218 | 0.004618452 |
| SNP_A.K_308299_2_7489138 | 2 | 7489138 | 2.65E-06 | 0.21572 | 299 | 0.151290428 | 0.004618452 |
| SNP_A.K_1001335_6_6466173 | 6 | 6466173 | 2.65E-06 | 0.29097 | 299 | 0.151288952 | 0.004618452 |
| SNP_A.K_1364934_8_6060307 | 8 | 6060307 | 2.67E-06 | 0.07692 | 299 | 0.151239516 | 0.004649999 |
| SNP_A.K_195534_1_31223799 | 1 | 31223799 | 2.68E-06 | 0.48328 | 299 | 0.151198603 | 0.004674968 |

| | | | | | | | |
|---|---|---|---|---|---|---|---|
| SNP_A.K_977994_6_1633040 | 6 | 1633040 | 2.69E-06 | 0.16054 | 299 | 0.15119407 | 0.004674968 |
| SNP_A.K_1433081_8_16617048 | 8 | 16617048 | 2.69E-06 | 0.08361 | 299 | 0.151174379 | 0.00468531 |
| SNP_A.K_729659_4_12624469 | 4 | 12624469 | 2.71E-06 | 0.22408 | 299 | 0.151134312 | 0.004710502 |
| SNP_A.K_561038_3_17749930 | 3 | 17749930 | 2.72E-06 | 0.20903 | 299 | 0.151109365 | 0.004724771 |
| SNP_A.K_1371159_8_7804028 | 8 | 7804028 | 2.73E-06 | 0.01839 | 299 | 0.151100747 | 0.004727131 |
| SNP_A.K_1004009_6_6815691 | 6 | 6815691 | 2.73E-06 | 0.29431 | 299 | 0.15108966 | 0.004731303 |
| SNP_A.K_1004016_6_6818321 | 6 | 6818321 | 2.74E-06 | 0.301 | 299 | 0.151059846 | 0.004749221 |
| SNP_A.K_307393_2_7400764 | 2 | 7400764 | 2.75E-06 | 0.2408 | 299 | 0.151048531 | 0.004753586 |
| SNP_A.K_978457_6_1694922 | 6 | 1694922 | 2.75E-06 | 0.26756 | 299 | 0.151030736 | 0.004757008 |
| SNP_A.K_990236_6_4701817 | 6 | 4701817 | 2.76E-06 | 0.19398 | 299 | 0.151030235 | 0.004757008 |
| SNP_A.K_1600941_9_18863472 | 9 | 18863472 | 2.76E-06 | 0.13211 | 299 | 0.15102778 | 0.004757008 |
| SNP_A.K_990378_6_4712741 | 6 | 4712741 | 2.76E-06 | 0.19231 | 299 | 0.151009905 | 0.004766229 |
| SNP_A.K_978777_6_1780110 | 6 | 1780110 | 2.77E-06 | 0.30936 | 299 | 0.151004541 | 0.004766238 |
| SNP_A.K_534170_3_12412541 | 3 | 12412541 | 2.79E-06 | 0.02676 | 299 | 0.150939909 | 0.004807055 |
| SNP_A.K_1364835_8_6047734 | 8 | 6047734 | 2.80E-06 | 0.10368 | 299 | 0.15093417 | 0.004807055 |
| SNP_A.K_992969_6_5691412 | 6 | 5691412 | 2.80E-06 | 0.22742 | 299 | 0.150933396 | 0.004807055 |
| SNP_A.K_990244_6_4702916 | 6 | 4702916 | 2.82E-06 | 0.21405 | 299 | 0.150889613 | 0.004831929 |
| SNP_A.K_374685_2_17872020 | 2 | 17872020 | 2.82E-06 | 0.06355 | 299 | 0.150889376 | 0.004831929 |
| SNP_A.K_1004223_6_6851230 | 6 | 6851230 | 2.82E-06 | 0.32441 | 299 | 0.150877085 | 0.004837141 |
| SNP_A.K_307513_2_7414680 | 2 | 7414680 | 2.83E-06 | 0.24415 | 299 | 0.150855167 | 0.00484958 |
| SNP_A.K_1004695_6_6918963 | 6 | 6918963 | 2.84E-06 | 0.36455 | 299 | 0.150835329 | 0.004860491 |
| SNP_A.K_306804_2_7352438 | 2 | 7352438 | 2.86E-06 | 0.2408 | 299 | 0.150798449 | 0.004884294 |
| SNP_A.K_1005416_6_7013002 | 6 | 7013002 | 2.86E-06 | 0.35786 | 299 | 0.150788392 | 0.004887886 |
| SNP_A.K_1004060_6_6822590 | 6 | 6822590 | 2.86E-06 | 0.29599 | 299 | 0.150780036 | 0.004890196 |
| SNP_A.K_306842_2_7359415 | 2 | 7359415 | 2.87E-06 | 0.25418 | 299 | 0.150768268 | 0.004891681 |
| SNP_A.K_306904_2_7364811 | 2 | 7364811 | 2.87E-06 | 0.25585 | 299 | 0.150767482 | 0.004891681 |
| SNP_A.K_1005371_6_7009948 | 6 | 7009948 | 2.88E-06 | 0.23746 | 299 | 0.150753505 | 0.004898263 |
| SNP_A.K_1364851_8_6048892 | 8 | 6048892 | 2.89E-06 | 0.11873 | 299 | 0.150712025 | 0.00492578 |
| SNP_A.K_977979_6_1628422 | 6 | 1628422 | 2.90E-06 | 0.23746 | 299 | 0.15070483 | 0.004927241 |
| SNP_A.K_977966_6_1626063 | 6 | 1626063 | 2.92E-06 | 0.16722 | 299 | 0.150654012 | 0.004953042 |
| SNP_A.K_709696_4_8281788 | 4 | 8281788 | 2.92E-06 | 0.09197 | 299 | 0.150651848 | 0.004953042 |
| SNP_A.K_978728_6_1771698 | 6 | 1771698 | 2.92E-06 | 0.35452 | 299 | 0.150648006 | 0.004953042 |
| SNP_A.K_307047_2_7375132 | 2 | 7375132 | 2.92E-06 | 0.23746 | 299 | 0.150645026 | 0.004953042 |
| SNP_A.K_307448_2_7408166 | 2 | 7408166 | 2.93E-06 | 0.26589 | 299 | 0.150641439 | 0.004953042 |
| SNP_A.K_1005205_6_6994413 | 6 | 6994413 | 2.93E-06 | 0.30769 | 299 | 0.150637069 | 0.004953042 |
| SNP_A.K_523315_3_10296571 | 3 | 10296571 | 2.93E-06 | 0.22074 | 299 | 0.150634249 | 0.004953042 |
| SNP_A.K_978414_6_1688826 | 6 | 1688826 | 2.94E-06 | 0.26254 | 299 | 0.150613379 | 0.004965036 |
| SNP_A.K_307346_2_7398170 | 2 | 7398170 | 2.94E-06 | 0.2291 | 299 | 0.150604669 | 0.004967703 |
| SNP_A.K_709908_4_8301435 | 4 | 8301435 | 2.95E-06 | 0.06856 | 299 | 0.150590867 | 0.004974294 |
| SNP_A.K_1602099_9_19127183 | 9 | 19127183 | 2.95E-06 | 0.11204 | 299 | 0.150585531 | 0.004974374 |
| SNP_A.K_1001121_6_6447982 | 6 | 6447982 | 2.97E-06 | 0.3796 | 299 | 0.150537211 | 0.005007671 |
| SNP_A.K_1364832_8_6047589 | 8 | 6047589 | 2.98E-06 | 0.10702 | 299 | 0.150525526 | 0.005012684 |
| SNP_A.K_723291_4_11429838 | 4 | 11429838 | 2.99E-06 | 0.0602 | 299 | 0.150510785 | 0.005020079 |
| SNP_A.K_1608678_9_20815241 | 9 | 20815241 | 2.99E-06 | 0.25418 | 299 | 0.150497426 | 0.005026413 |
| SNP_A.K_991211_6_4870675 | 6 | 4870675 | 3.04E-06 | 0.05518 | 299 | 0.150394047 | 0.005103403 |
| SNP_A.K_996049_6_5959426 | 6 | 5959426 | 3.04E-06 | 0.3194 | 299 | 0.150388439 | 0.005103721 |
| SNP_A.K_306841_2_7359409 | 2 | 7359409 | 3.05E-06 | 0.24749 | 299 | 0.150375474 | 0.005109861 |
| SNP_A.K_306851_2_7360244 | 2 | 7360244 | 3.05E-06 | 0.26923 | 299 | 0.15036613 | 0.005113143 |
| SNP_A.K_307168_2_7383136 | 2 | 7383136 | 3.08E-06 | 0.23746 | 299 | 0.150313638 | 0.005142444 |
| SNP_A.K_978101_6_1647188 | 6 | 1647188 | 3.08E-06 | 0.25251 | 299 | 0.150311453 | 0.005142444 |

| | | | | | | | |
|---|---|---|---|---|---|---|---|
| SNP_A.K_709828_4_8298974 | 4 | 8298974 | 3.08E-06 | 0.06689 | 299 | 0.150310042 | 0.005142444 |
| SNP_A.K_1005637_6_7032073 | 6 | 7032073 | 3.08E-06 | 0.29097 | 299 | 0.150305233 | 0.005142444 |
| SNP_A.K_1390041_8_10146158 | 8 | 10146158 | 3.08E-06 | 0.04348 | 299 | 0.150303313 | 0.005142444 |
| SNP_A.K_999070_6_6254457 | 6 | 6254457 | 3.09E-06 | 0.13712 | 299 | 0.150288538 | 0.005150099 |
| SNP_A.K_1364993_8_6065627 | 8 | 6065627 | 3.10E-06 | 0.11204 | 299 | 0.150267317 | 0.005162921 |
| SNP_A.K_306779_2_7349769 | 2 | 7349769 | 3.12E-06 | 0.23579 | 299 | 0.150228207 | 0.005186737 |
| SNP_A.K_328328_2_10222383 | 2 | 10222383 | 3.12E-06 | 0.08027 | 299 | 0.150227282 | 0.005186737 |
| SNP_A.K_1608610_9_20794181 | 9 | 20794181 | 3.14E-06 | 0.23244 | 299 | 0.150188307 | 0.005213981 |
| SNP_A.K_990276_6_4704269 | 6 | 4704269 | 3.15E-06 | 0.20736 | 299 | 0.150155779 | 0.005235278 |
| SNP_A.K_1608503_9_20765723 | 9 | 20765723 | 3.16E-06 | 0.28763 | 299 | 0.150151171 | 0.005235278 |
| SNP_A.K_1364921_8_6057127 | 8 | 6057127 | 3.16E-06 | 0.12207 | 299 | 0.150146557 | 0.005235278 |
| SNP_A.K_1004568_6_6906786 | 6 | 6906786 | 3.17E-06 | 0.29431 | 299 | 0.150133198 | 0.00524195 |
| SNP_A.K_2080286_12_27254632 | 12 | 27254632 | 3.17E-06 | 0.01003 | 299 | 0.150127404 | 0.005242487 |
| SNP_A.K_978635_6_1748121 | 6 | 1748121 | 3.20E-06 | 0.29766 | 299 | 0.150056452 | 0.005296223 |
| SNP_A.K_1364547_8_6021345 | 8 | 6021345 | 3.21E-06 | 0.08528 | 299 | 0.150046496 | 0.005300189 |
| SNP_A.K_997563_6_6109438 | 6 | 6109438 | 3.23E-06 | 0.14548 | 299 | 0.15000867 | 0.005326849 |
| SNP_A.K_1004343_6_6870262 | 6 | 6870262 | 3.23E-06 | 0.32107 | 299 | 0.150003593 | 0.005326849 |
| SNP_A.K_978118_6_1647773 | 6 | 1647773 | 3.23E-06 | 0.19064 | 299 | 0.149998766 | 0.005326849 |
| SNP_A.K_1830694_11_10574633 | 11 | 10574633 | 3.25E-06 | 0.35953 | 299 | 0.149956772 | 0.005357383 |
| SNP_A.K_709794_4_8297841 | 4 | 8297841 | 3.26E-06 | 0.06689 | 299 | 0.149947517 | 0.005360831 |
| SNP_A.K_978721_6_1771190 | 6 | 1771190 | 3.27E-06 | 0.3495 | 299 | 0.149932489 | 0.005369085 |
| SNP_A.K_1953710_12_3620597 | 12 | 3620597 | 3.27E-06 | 0.04013 | 299 | 0.149919969 | 0.005375267 |
| SNP_A.K_325798_2_9943355 | 2 | 9943355 | 3.28E-06 | 0.07023 | 299 | 0.1499125 | 0.005377248 |
| SNP_A.K_1005563_6_7025295 | 6 | 7025295 | 3.28E-06 | 0.26421 | 299 | 0.149901454 | 0.005382216 |
| SNP_A.K_994589_6_5874944 | 6 | 5874944 | 3.28E-06 | 0.25753 | 299 | 0.149894935 | 0.005383414 |
| SNP_A.K_1005892_6_7050571 | 6 | 7050571 | 3.30E-06 | 0.31271 | 299 | 0.149873983 | 0.005396674 |
| SNP_A.K_255301_1_41158313 | 1 | 41158313 | 3.30E-06 | 0.17057 | 299 | 0.149857555 | 0.005406178 |
| SNP_A.K_1005568_6_7025357 | 6 | 7025357 | 3.31E-06 | 0.2592 | 299 | 0.149841816 | 0.005415125 |
| SNP_A.K_306925_2_7368747 | 2 | 7368747 | 3.33E-06 | 0.24415 | 299 | 0.149804558 | 0.005440549 |
| SNP_A.K_306881_2_7363278 | 2 | 7363278 | 3.33E-06 | 0.25418 | 299 | 0.149801467 | 0.005440549 |
| SNP_A.K_1602020_9_19105955 | 9 | 19105955 | 3.34E-06 | 0.11204 | 299 | 0.149792474 | 0.005442815 |
| SNP_A.K_1004008_6_6815570 | 6 | 6815570 | 3.34E-06 | 0.30268 | 299 | 0.149788667 | 0.005442815 |
| SNP_A.K_1004226_6_6851854 | 6 | 6851854 | 3.34E-06 | 0.33445 | 299 | 0.149778304 | 0.00544642 |
| SNP_A.K_992435_6_5646433 | 6 | 5646433 | 3.35E-06 | 0.32609 | 299 | 0.1497743 | 0.00544642 |
| SNP_A.K_1000644_6_6403466 | 6 | 6403466 | 3.36E-06 | 0.29431 | 299 | 0.149748725 | 0.005463782 |
| SNP_A.K_978115_6_1647659 | 6 | 1647659 | 3.37E-06 | 0.199 | 299 | 0.149733598 | 0.005470246 |
| SNP_A.K_1390009_8_10145157 | 8 | 10145157 | 3.37E-06 | 0.04682 | 299 | 0.149731026 | 0.005470246 |
| SNP_A.K_1830676_11_10573745 | 11 | 10573745 | 3.38E-06 | 0.16221 | 299 | 0.149717808 | 0.005477194 |
| SNP_A.K_194268_1_30836704 | 1 | 30836704 | 3.40E-06 | 0.49331 | 299 | 0.149676786 | 0.005507843 |
| SNP_A.K_1608551_9_20776534 | 9 | 20776534 | 3.44E-06 | 0.23579 | 299 | 0.14959114 | 0.005577123 |
| SNP_A.K_1390036_8_10146095 | 8 | 10146095 | 3.47E-06 | 0.04682 | 299 | 0.149539866 | 0.005617268 |
| SNP_A.K_999393_6_6286280 | 6 | 6286280 | 3.48E-06 | 0.11538 | 299 | 0.149530181 | 0.005620997 |
| SNP_A.K_307495_2_7413016 | 2 | 7413016 | 3.48E-06 | 0.24247 | 299 | 0.149525564 | 0.005620997 |
| SNP_A.K_1364915_8_6056729 | 8 | 6056729 | 3.49E-06 | 0.11371 | 299 | 0.149495303 | 0.005639717 |
| SNP_A.K_1005562_6_7024873 | 6 | 7024873 | 3.50E-06 | 0.26923 | 299 | 0.149494117 | 0.005639717 |
| SNP_A.K_1005258_6_6999987 | 6 | 6999987 | 3.50E-06 | 0.25753 | 299 | 0.149478653 | 0.005648879 |
| SNP_A.K_709988_4_8304496 | 4 | 8304496 | 3.52E-06 | 0.07692 | 299 | 0.149440811 | 0.005677729 |
| SNP_A.K_308298_2_7489125 | 2 | 7489125 | 3.55E-06 | 0.21739 | 299 | 0.149397303 | 0.005711748 |
| SNP_A.K_1005467_6_7015374 | 6 | 7015374 | 3.56E-06 | 0.32107 | 299 | 0.149367912 | 0.00573341 |
| SNP_A.K_978729_6_1771713 | 6 | 1771713 | 3.57E-06 | 0.35786 | 299 | 0.149361755 | 0.005734456 |

| | | | | | | | |
|---|---|---|---|---|---|---|---|
| SNP_A.K_997491_6_6104256 | 6 | 6104256 | 3.57E-06 | 0.14548 | 299 | 0.149353104 | 0.005737726 |
| SNP_A.K_978770_6_1779076 | 6 | 1779076 | 3.58E-06 | 0.29431 | 299 | 0.14934668 | 0.005738293 |
| SNP_A.K_709963_4_8303718 | 4 | 8303718 | 3.58E-06 | 0.07692 | 299 | 0.149342525 | 0.005738293 |
| SNP_A.K_1004218_6_6850083 | 6 | 6850083 | 3.60E-06 | 0.30435 | 299 | 0.149304437 | 0.005766396 |
| SNP_A.K_978858_6_1788706 | 6 | 1788706 | 3.60E-06 | 0.31438 | 299 | 0.149301103 | 0.005766396 |
| SNP_A.K_990232_6_4701592 | 6 | 4701592 | 3.62E-06 | 0.20234 | 299 | 0.149270623 | 0.00578866 |
| SNP_A.K_978623_6_1745339 | 6 | 1745339 | 3.62E-06 | 0.27258 | 299 | 0.149266351 | 0.00578866 |
| SNP_A.K_1608604_9_20793807 | 9 | 20793807 | 3.63E-06 | 0.23244 | 299 | 0.149256434 | 0.00579204 |
| SNP_A.K_1004796_6_6933103 | 6 | 6933103 | 3.63E-06 | 0.31271 | 299 | 0.149242625 | 0.00579204 |
| SNP_A.K_1371520_8_7850808 | 8 | 7850808 | 3.63E-06 | 0.01839 | 299 | 0.149241246 | 0.00579204 |
| SNP_A.K_1001296_6_6463011 | 6 | 6463011 | 3.64E-06 | 0.3194 | 299 | 0.149238219 | 0.00579204 |
| SNP_A.K_978903_6_1799767 | 6 | 1799767 | 3.64E-06 | 0.32107 | 299 | 0.149237881 | 0.00579204 |
| SNP_A.K_1005644_6_7032880 | 6 | 7032880 | 3.66E-06 | 0.23746 | 299 | 0.1492016 | 0.005820278 |
| SNP_A.K_978608_6_1740753 | 6 | 1740753 | 3.67E-06 | 0.26421 | 299 | 0.149173697 | 0.00583676 |
| SNP_A.K_1004926_6_6953368 | 6 | 6953368 | 3.67E-06 | 0.30268 | 299 | 0.149173526 | 0.00583676 |
| SNP_A.K_978618_6_1744863 | 6 | 1744863 | 3.68E-06 | 0.29264 | 299 | 0.149165493 | 0.005839581 |
| SNP_A.K_1005298_6_7002963 | 6 | 7002963 | 3.71E-06 | 0.28261 | 299 | 0.149110974 | 0.005884699 |
| SNP_A.K_1005356_6_7008916 | 6 | 7008916 | 3.71E-06 | 0.30936 | 299 | 0.149104636 | 0.005885113 |
| SNP_A.K_307531_2_7416006 | 2 | 7416006 | 3.72E-06 | 0.25418 | 299 | 0.149100703 | 0.005885113 |
| SNP_A.K_1001163_6_6449922 | 6 | 6449922 | 3.72E-06 | 0.37625 | 299 | 0.149089772 | 0.00589062 |
| SNP_A.K_977877_6_1607061 | 6 | 1607061 | 3.73E-06 | 0.15719 | 299 | 0.149071169 | 0.0058999 |
| SNP_A.K_978939_6_1814581 | 6 | 1814581 | 3.73E-06 | 0.32441 | 299 | 0.149069835 | 0.0058999 |
| SNP_A.K_977899_6_1617097 | 6 | 1617097 | 3.75E-06 | 0.23746 | 299 | 0.14903704 | 0.005922667 |
| SNP_A.K_1004345_6_6870342 | 6 | 6870342 | 3.75E-06 | 0.32943 | 299 | 0.149035236 | 0.005922667 |
| SNP_A.K_307480_2_7410467 | 2 | 7410467 | 3.76E-06 | 0.28595 | 299 | 0.14901703 | 0.005934924 |
| SNP_A.K_1830741_11_10577127 | 11 | 10577127 | 3.78E-06 | 0.1204 | 299 | 0.148994008 | 0.005947825 |
| SNP_A.K_978818_6_1783773 | 6 | 1783773 | 3.78E-06 | 0.31271 | 299 | 0.148981158 | 0.005947825 |
| SNP_A.K_514070_3_8391504 | 3 | 8391504 | 3.79E-06 | 0.02676 | 299 | 0.148978676 | 0.005947825 |
| SNP_A.K_514018_3_8386416 | 3 | 8386416 | 3.79E-06 | 0.02676 | 299 | 0.148978676 | 0.005947825 |
| SNP_A.K_514106_3_8401030 | 3 | 8401030 | 3.79E-06 | 0.02676 | 299 | 0.148978676 | 0.005947825 |
| SNP_A.K_1830648_11_10572698 | 11 | 10572698 | 3.80E-06 | 0.1689 | 299 | 0.14895562 | 0.005962327 |
| SNP_A.K_978798_6_1782236 | 6 | 1782236 | 3.80E-06 | 0.29766 | 299 | 0.148953266 | 0.005962327 |
| SNP_A.K_1001509_6_6485667 | 6 | 6485667 | 3.82E-06 | 0.29431 | 299 | 0.148919888 | 0.005988782 |
| SNP_A.K_1390023_8_10145472 | 8 | 10145472 | 3.83E-06 | 0.05518 | 299 | 0.148908134 | 0.005995204 |
| SNP_A.K_978839_6_1787223 | 6 | 1787223 | 3.84E-06 | 0.34281 | 299 | 0.148888642 | 0.006004333 |
| SNP_A.K_978838_6_1787221 | 6 | 1787221 | 3.84E-06 | 0.34281 | 299 | 0.148888642 | 0.006004333 |
| SNP_A.K_978767_6_1777598 | 6 | 1777598 | 3.84E-06 | 0.30769 | 299 | 0.148879574 | 0.006005084 |
| SNP_A.K_990331_6_4710968 | 6 | 4710968 | 3.85E-06 | 0.20067 | 299 | 0.148878163 | 0.006005084 |
| SNP_A.K_1001092_6_6441019 | 6 | 6441019 | 3.87E-06 | 0.39465 | 299 | 0.148833275 | 0.006038642 |
| SNP_A.K_389622_2_19887350 | 2 | 19887350 | 3.87E-06 | 0.02843 | 299 | 0.148832603 | 0.006038642 |
| SNP_A.K_1608554_9_20776623 | 9 | 20776623 | 3.90E-06 | 0.23579 | 299 | 0.148795508 | 0.006063606 |
| SNP_A.K_997482_6_6103027 | 6 | 6103027 | 3.90E-06 | 0.13378 | 299 | 0.148791914 | 0.006063606 |
| SNP_A.K_1004649_6_6913565 | 6 | 6913565 | 3.90E-06 | 0.32609 | 299 | 0.148791568 | 0.006063606 |
| SNP_A.K_978612_6_1741229 | 6 | 1741229 | 3.91E-06 | 0.25251 | 299 | 0.148766125 | 0.006083057 |
| SNP_A.K_306854_2_7360517 | 2 | 7360517 | 3.93E-06 | 0.24415 | 299 | 0.148746202 | 0.006097348 |
| SNP_A.K_978887_6_1796552 | 6 | 1796552 | 3.94E-06 | 0.31104 | 299 | 0.148713979 | 0.006123358 |
| SNP_A.K_1003932_6_6806269 | 6 | 6806269 | 3.95E-06 | 0.32107 | 299 | 0.148703627 | 0.006128638 |
| SNP_A.K_710050_4_8312755 | 4 | 8312755 | 3.97E-06 | 0.08027 | 299 | 0.148673667 | 0.006152629 |
| SNP_A.K_709698_4_8282162 | 4 | 8282162 | 4.01E-06 | 0.09197 | 299 | 0.148608227 | 0.006210841 |
| SNP_A.K_1005610_6_7029780 | 6 | 7029780 | 4.04E-06 | 0.23913 | 299 | 0.148561337 | 0.00625158 |

| | | | | | | | |
|---|---|---|---|---|---|---|---|
| SNP_A.K_993964_6_5804126 | 6 | 5804126 | 4.07E-06 | 0.20569 | 299 | 0.148518942 | 0.0062882 |
| SNP_A.K_308303_2_7489534 | 2 | 7489534 | 4.09E-06 | 0.22074 | 299 | 0.148487406 | 0.00631099 |
| SNP_A.K_709945_4_8302688 | 4 | 8302688 | 4.09E-06 | 0.07692 | 299 | 0.148484524 | 0.00631099 |
| SNP_A.K_992437_6_5646455 | 6 | 5646455 | 4.09E-06 | 0.11371 | 299 | 0.148481314 | 0.00631099 |
| SNP_A.K_977867_6_1605358 | 6 | 1605358 | 4.10E-06 | 0.16221 | 299 | 0.148464551 | 0.006322756 |
| SNP_A.K_998751_6_6219656 | 6 | 6219656 | 4.12E-06 | 0.12542 | 299 | 0.148439601 | 0.006342607 |
| SNP_A.K_978864_6_1791845 | 6 | 1791845 | 4.14E-06 | 0.29097 | 299 | 0.14840563 | 0.006367871 |
| SNP_A.K_1000032_6_6342793 | 6 | 6342793 | 4.14E-06 | 0.14214 | 299 | 0.148404485 | 0.006367871 |
| SNP_A.K_709962_4_8303650 | 4 | 8303650 | 4.18E-06 | 0.08027 | 299 | 0.148345698 | 0.006421543 |
| SNP_A.K_306849_2_7360112 | 2 | 7360112 | 4.19E-06 | 0.25585 | 299 | 0.148328231 | 0.00643424 |
| SNP_A.K_1004613_6_6910734 | 6 | 6910734 | 4.23E-06 | 0.30602 | 299 | 0.148266784 | 0.006487225 |
| SNP_A.K_678622_4_991811 | 4 | 991811 | 4.23E-06 | 0.06522 | 299 | 0.148265949 | 0.006487225 |
| SNP_A.K_978013_6_1636356 | 6 | 1636356 | 4.24E-06 | 0.16722 | 299 | 0.1482475 | 0.006501056 |
| SNP_A.K_1001222_6_6455553 | 6 | 6455553 | 4.25E-06 | 0.28261 | 299 | 0.148238301 | 0.006505564 |
| SNP_A.K_990231_6_4701562 | 6 | 4701562 | 4.26E-06 | 0.199 | 299 | 0.148215726 | 0.006523622 |
| SNP_A.K_1830419_11_10560646 | 11 | 10560646 | 4.27E-06 | 0.15886 | 299 | 0.148198391 | 0.006536412 |
| SNP_A.K_1005357_6_7008937 | 6 | 7008937 | 4.29E-06 | 0.2893 | 299 | 0.1481762 | 0.006554174 |
| SNP_A.K_1433083_8_16617071 | 8 | 16617071 | 4.30E-06 | 0.08528 | 299 | 0.148161475 | 0.006564369 |
| SNP_A.K_1004041_6_6821331 | 6 | 6821331 | 4.32E-06 | 0.30268 | 299 | 0.148122342 | 0.006599567 |
| SNP_A.K_709944_4_8302621 | 4 | 8302621 | 4.33E-06 | 0.07692 | 299 | 0.148108988 | 0.006608432 |
| SNP_A.K_307079_2_7378017 | 2 | 7378017 | 4.35E-06 | 0.25418 | 299 | 0.148089447 | 0.006621818 |
| SNP_A.K_978837_6_1787217 | 6 | 1787217 | 4.35E-06 | 0.34783 | 299 | 0.148086546 | 0.006621818 |
| SNP_A.K_456488_2_33045330 | 2 | 33045330 | 4.35E-06 | 0.11706 | 299 | 0.148079765 | 0.006623955 |
| SNP_A.K_1365002_8_6065905 | 8 | 6065905 | 4.40E-06 | 0.0903 | 299 | 0.148015654 | 0.0066854 |
| SNP_A.K_1004592_6_6908932 | 6 | 6908932 | 4.40E-06 | 0.29097 | 299 | 0.148009374 | 0.006687043 |
| SNP_A.K_978621_6_1745281 | 6 | 1745281 | 4.41E-06 | 0.27592 | 299 | 0.147993941 | 0.006693937 |
| SNP_A.K_306729_2_7345940 | 2 | 7345940 | 4.41E-06 | 0.22575 | 299 | 0.14799336 | 0.006693937 |
| SNP_A.K_306777_2_7349731 | 2 | 7349731 | 4.42E-06 | 0.23244 | 299 | 0.147986823 | 0.006695861 |
| SNP_A.K_1000868_6_6421629 | 6 | 6421629 | 4.42E-06 | 0.3913 | 299 | 0.147976179 | 0.006701343 |
| SNP_A.K_978835_6_1787160 | 6 | 1787160 | 4.43E-06 | 0.3612 | 299 | 0.147972194 | 0.006701343 |
| SNP_A.K_978866_6_1792588 | 6 | 1792588 | 4.43E-06 | 0.29097 | 299 | 0.147963765 | 0.006705252 |
| SNP_A.K_1608594_9_20792063 | 9 | 20792063 | 4.44E-06 | 0.23077 | 299 | 0.147959086 | 0.006705257 |
| SNP_A.K_990343_6_4711721 | 6 | 4711721 | 4.46E-06 | 0.20569 | 299 | 0.147928433 | 0.006730877 |
| SNP_A.K_1500203_9_688602 | 9 | 688602 | 4.46E-06 | 0.03512 | 299 | 0.147925217 | 0.006730877 |
| SNP_A.K_1003991_6_6813686 | 6 | 6813686 | 4.47E-06 | 0.34615 | 299 | 0.147912533 | 0.006739274 |
| SNP_A.K_307285_2_7391278 | 2 | 7391278 | 4.48E-06 | 0.24415 | 299 | 0.147891428 | 0.006751928 |
| SNP_A.K_978789_6_1781572 | 6 | 1781572 | 4.49E-06 | 0.29431 | 299 | 0.14788415 | 0.006751928 |
| SNP_A.K_977968_6_1626348 | 6 | 1626348 | 4.49E-06 | 0.16054 | 299 | 0.147882104 | 0.006751928 |
| SNP_A.K_138883_1_21888564 | 1 | 21888564 | 4.49E-06 | 0.01003 | 299 | 0.147881848 | 0.006751928 |
| SNP_A.K_1005727_6_7035477 | 6 | 7035477 | 4.50E-06 | 0.28763 | 299 | 0.147861589 | 0.006763992 |
| SNP_A.K_306889_2_7363995 | 2 | 7363995 | 4.50E-06 | 0.26254 | 299 | 0.147861076 | 0.006763992 |
| SNP_A.K_1390016_8_10145390 | 8 | 10145390 | 4.51E-06 | 0.04515 | 299 | 0.147851824 | 0.006765476 |
| SNP_A.K_1788806_11_4375003 | 11 | 4375003 | 4.51E-06 | 0.02174 | 299 | 0.147847761 | 0.006765476 |
| SNP_A.K_306865_2_7361459 | 2 | 7361459 | 4.51E-06 | 0.25585 | 299 | 0.147845757 | 0.006765476 |
| SNP_A.K_306793_2_7351367 | 2 | 7351367 | 4.54E-06 | 0.26589 | 299 | 0.147816173 | 0.0067906 |
| SNP_A.K_1004818_6_6933986 | 6 | 6933986 | 4.54E-06 | 0.301 | 299 | 0.147812662 | 0.0067906 |
| SNP_A.K_1518103_9_3562346 | 9 | 3562346 | 4.55E-06 | 0.04515 | 299 | 0.147799853 | 0.006799249 |
| SNP_A.K_999706_6_6308329 | 6 | 6308329 | 4.56E-06 | 0.14548 | 299 | 0.147786517 | 0.006808471 |
| SNP_A.K_1781570_11_3218786 | 11 | 3218786 | 4.57E-06 | 0.04013 | 299 | 0.147767687 | 0.00682196 |
| SNP_A.K_1500114_9_680219 | 9 | 680219 | 4.57E-06 | 0.04013 | 299 | 0.14776456 | 0.00682196 |

| | | | | | | | |
|---|---|---|---|---|---|---|---|
| SNP_A.K_978412_6_1688287 | 6 | 1688287 | 4.58E-06 | 0.2592 | 299 | 0.147750546 | 0.006831944 |
| SNP_A.K_306736_2_7346730 | 2 | 7346730 | 4.59E-06 | 0.25251 | 299 | 0.147736109 | 0.006842397 |
| SNP_A.K_978458_6_1695357 | 6 | 1695357 | 4.61E-06 | 0.26087 | 299 | 0.147715712 | 0.006855656 |
| SNP_A.K_1005724_6_7035415 | 6 | 7035415 | 4.61E-06 | 0.2893 | 299 | 0.147714459 | 0.006855656 |
| SNP_A.K_525716_3_10766180 | 3 | 10766180 | 4.61E-06 | 0.01171 | 299 | 0.147705458 | 0.006860355 |
| SNP_A.K_306728_2_7345840 | 2 | 7345840 | 4.62E-06 | 0.25084 | 299 | 0.147696102 | 0.006865439 |
| SNP_A.K_195521_1_31221088 | 1 | 31221088 | 4.64E-06 | 0.47826 | 299 | 0.147663546 | 0.006891804 |
| SNP_A.K_957708_5_26977103 | 5 | 26977103 | 4.65E-06 | 0.02843 | 299 | 0.147662282 | 0.006891804 |
| SNP_A.K_307379_2_7399794 | 2 | 7399794 | 4.66E-06 | 0.25418 | 299 | 0.147635296 | 0.006915859 |
| SNP_A.K_1001218_6_6455370 | 6 | 6455370 | 4.69E-06 | 0.2893 | 299 | 0.147603329 | 0.006945381 |
| SNP_A.K_307322_2_7394997 | 2 | 7394997 | 4.70E-06 | 0.24749 | 299 | 0.147593031 | 0.006951566 |
| SNP_A.K_978675_6_1756071 | 6 | 1756071 | 4.73E-06 | 0.28428 | 299 | 0.14754406 | 0.006999739 |
| SNP_A.K_456493_2_33047430 | 2 | 33047430 | 4.74E-06 | 0.1204 | 299 | 0.147538798 | 0.007000495 |
| SNP_A.K_752094_4_17831574 | 4 | 17831574 | 4.76E-06 | 0.02007 | 299 | 0.147504895 | 0.007032514 |
| SNP_A.K_1433038_8_16614032 | 8 | 16614032 | 4.78E-06 | 0.08194 | 299 | 0.147484519 | 0.007049837 |
| SNP_A.K_1390020_8_10145451 | 8 | 10145451 | 4.80E-06 | 0.06355 | 299 | 0.147453822 | 0.007070131 |
| SNP_A.K_978894_6_1797551 | 6 | 1797551 | 4.80E-06 | 0.29766 | 299 | 0.147450594 | 0.007070131 |
| SNP_A.K_1004253_6_6854257 | 6 | 6854257 | 4.80E-06 | 0.09866 | 299 | 0.147448995 | 0.007070131 |
| SNP_A.K_710324_4_8336412 | 4 | 8336412 | 4.80E-06 | 0.08528 | 299 | 0.147447452 | 0.007070131 |
| SNP_A.K_673444_4_362125 | 4 | 362125 | 4.81E-06 | 0.12709 | 299 | 0.147443286 | 0.007070131 |
| SNP_A.K_1005469_6_7015473 | 6 | 7015473 | 4.82E-06 | 0.27926 | 299 | 0.147430243 | 0.007079488 |
| SNP_A.K_1005029_6_6962960 | 6 | 6962960 | 4.83E-06 | 0.30936 | 299 | 0.147408445 | 0.007098524 |
| SNP_A.K_1001105_6_6443114 | 6 | 6443114 | 4.84E-06 | 0.38963 | 299 | 0.147399607 | 0.007099374 |
| SNP_A.K_307478_2_7410068 | 2 | 7410068 | 4.84E-06 | 0.26756 | 299 | 0.147398549 | 0.007099374 |
| SNP_A.K_978411_6_1688271 | 6 | 1688271 | 4.84E-06 | 0.26087 | 299 | 0.147394081 | 0.007099374 |
| SNP_A.K_1004679_6_6916461 | 6 | 6916461 | 4.85E-06 | 0.31605 | 299 | 0.147381951 | 0.007107779 |
| SNP_A.K_1608643_9_20809184 | 9 | 20809184 | 4.90E-06 | 0.28261 | 299 | 0.147323619 | 0.007167539 |
| SNP_A.K_307279_2_7391069 | 2 | 7391069 | 4.91E-06 | 0.4097 | 299 | 0.147302709 | 0.007185844 |
| SNP_A.K_994712_6_5881760 | 6 | 5881760 | 4.94E-06 | 0.16722 | 299 | 0.147264541 | 0.007223572 |
| SNP_A.K_307516_2_7414902 | 2 | 7414902 | 4.95E-06 | 0.2408 | 299 | 0.147257038 | 0.007226936 |
| SNP_A.K_1608749_9_20829793 | 9 | 20829793 | 4.96E-06 | 0.24415 | 299 | 0.147245469 | 0.007234881 |
| SNP_A.K_1002066_6_6539233 | 6 | 6539233 | 4.96E-06 | 0.31773 | 299 | 0.147237108 | 0.007239224 |
| SNP_A.K_1004714_6_6924931 | 6 | 6924931 | 4.98E-06 | 0.32609 | 299 | 0.147220566 | 0.007252801 |
| SNP_A.K_307376_2_7399758 | 2 | 7399758 | 4.98E-06 | 0.25251 | 299 | 0.147216031 | 0.007252843 |
| SNP_A.K_1004193_6_6844944 | 6 | 6844944 | 5.00E-06 | 0.301 | 299 | 0.147184703 | 0.00727822 |
| SNP_A.K_908260_5_17494874 | 5 | 17494874 | 5.00E-06 | 0.01839 | 299 | 0.147184601 | 0.00727822 |
| SNP_A.K_977906_6_1619467 | 6 | 1619467 | 5.01E-06 | 0.24247 | 299 | 0.147176511 | 0.0072823 |
| SNP_A.K_1001239_6_6457018 | 6 | 6457018 | 5.02E-06 | 0.3913 | 299 | 0.147164852 | 0.007289571 |
| SNP_A.K_726725_4_12038546 | 4 | 12038546 | 5.02E-06 | 0.07023 | 299 | 0.147161132 | 0.007289571 |
| SNP_A.K_1390012_8_10145235 | 8 | 10145235 | 5.05E-06 | 0.04849 | 299 | 0.14712883 | 0.007321206 |
| SNP_A.K_978055_6_1644446 | 6 | 1644446 | 5.06E-06 | 0.19398 | 299 | 0.147115028 | 0.007331841 |
| SNP_A.K_1608501_9_20765517 | 9 | 20765517 | 5.07E-06 | 0.30435 | 299 | 0.147099307 | 0.007339744 |
| SNP_A.K_1001165_6_6450074 | 6 | 6450074 | 5.07E-06 | 0.39632 | 299 | 0.147099165 | 0.007339744 |
| SNP_A.K_1004171_6_6840823 | 6 | 6840823 | 5.08E-06 | 0.30602 | 299 | 0.147086897 | 0.007348663 |
| SNP_A.K_1726582_10_16600979 | 10 | 16600979 | 5.11E-06 | 0.05853 | 299 | 0.147053665 | 0.007381644 |
| SNP_A.K_1001595_6_6496375 | 6 | 6496375 | 5.12E-06 | 0.29431 | 299 | 0.147031451 | 0.007402075 |
| SNP_A.K_997768_6_6145949 | 6 | 6145949 | 5.13E-06 | 0.15886 | 299 | 0.147021047 | 0.007408455 |
| SNP_A.K_1830670_11_10573397 | 11 | 10573397 | 5.14E-06 | 0.34615 | 299 | 0.147017007 | 0.007408455 |
| SNP_A.K_760943_4_19823560 | 4 | 19823560 | 5.17E-06 | 0.01003 | 299 | 0.146977118 | 0.007449439 |
| SNP_A.K_307402_2_7401363 | 2 | 7401363 | 5.19E-06 | 0.24916 | 299 | 0.146943653 | 0.007483171 |

| | | | | | | | |
|---|---|---|---|---|---|---|---|
| SNP_A.K_978731_6_1771843 | 6 | 1771843 | 5.21E-06 | 0.30268 | 299 | 0.146929978 | 0.007491416 |
| SNP_A.K_306715_2_7344401 | 2 | 7344401 | 5.21E-06 | 0.24916 | 299 | 0.146924342 | 0.007491416 |
| SNP_A.K_1003988_6_6812535 | 6 | 6812535 | 5.21E-06 | 0.37458 | 299 | 0.146923259 | 0.007491416 |
| SNP_A.K_1608627_9_20803563 | 9 | 20803563 | 5.23E-06 | 0.28261 | 299 | 0.146904365 | 0.007508301 |
| SNP_A.K_978941_6_1814778 | 6 | 1814778 | 5.26E-06 | 0.3194 | 299 | 0.146862348 | 0.007552372 |
| SNP_A.K_1261281_7_17331377 | 7 | 17331377 | 5.27E-06 | 0.01338 | 299 | 0.146856974 | 0.007553484 |
| SNP_A.K_1001226_6_6455942 | 6 | 6455942 | 5.27E-06 | 0.28595 | 299 | 0.146850729 | 0.007555626 |
| SNP_A.K_990355_6_4712248 | 6 | 4712248 | 5.28E-06 | 0.19565 | 299 | 0.146835385 | 0.007568488 |
| SNP_A.K_1005358_6_7008950 | 6 | 7008950 | 5.31E-06 | 0.24415 | 299 | 0.146797234 | 0.007608349 |
| SNP_A.K_306769_2_7349058 | 2 | 7349058 | 5.32E-06 | 0.25251 | 299 | 0.146787471 | 0.0076125 |
| SNP_A.K_307528_2_7415763 | 2 | 7415763 | 5.32E-06 | 0.2408 | 299 | 0.146784905 | 0.0076125 |
| SNP_A.K_996253_6_5977236 | 6 | 5977236 | 5.37E-06 | 0.31271 | 299 | 0.14673521 | 0.007666378 |
| SNP_A.K_726096_4_11913307 | 4 | 11913307 | 5.38E-06 | 0.09197 | 299 | 0.146721169 | 0.007677892 |
| SNP_A.K_308579_2_7527594 | 2 | 7527594 | 5.38E-06 | 0.23077 | 299 | 0.146712853 | 0.007682574 |
| SNP_A.K_1001094_6_6441068 | 6 | 6441068 | 5.39E-06 | 0.38796 | 299 | 0.14670278 | 0.007689366 |
| SNP_A.K_1830655_11_10572967 | 11 | 10572967 | 5.40E-06 | 0.3495 | 299 | 0.146695996 | 0.007692227 |
| SNP_A.K_1830732_11_10576194 | 11 | 10576194 | 5.43E-06 | 0.20569 | 299 | 0.146654972 | 0.00773624 |
| SNP_A.K_307127_2_7380892 | 2 | 7380892 | 5.46E-06 | 0.26589 | 299 | 0.146618741 | 0.007774705 |
| SNP_A.K_1830645_11_10572636 | 11 | 10572636 | 5.47E-06 | 0.35117 | 299 | 0.146613155 | 0.007776158 |
| SNP_A.K_1390366_8_10192958 | 8 | 10192958 | 5.48E-06 | 0.06522 | 299 | 0.146601935 | 0.007784444 |
| SNP_A.K_1004207_6_6848565 | 6 | 6848565 | 5.48E-06 | 0.30435 | 299 | 0.146594834 | 0.007787744 |
| SNP_A.K_1432028_8_16445400 | 8 | 16445400 | 5.52E-06 | 0.07191 | 299 | 0.146548814 | 0.007837392 |
| SNP_A.K_1004634_6_6912599 | 6 | 6912599 | 5.53E-06 | 0.27592 | 299 | 0.146545289 | 0.007837392 |
| SNP_A.K_1001127_6_6448295 | 6 | 6448295 | 5.54E-06 | 0.28261 | 299 | 0.146528844 | 0.007850572 |
| SNP_A.K_978902_6_1799753 | 6 | 1799753 | 5.54E-06 | 0.3194 | 299 | 0.146525764 | 0.007850572 |
| SNP_A.K_978116_6_1647676 | 6 | 1647676 | 5.59E-06 | 0.19398 | 299 | 0.146468439 | 0.007913052 |
| SNP_A.K_990377_6_4712728 | 6 | 4712728 | 5.60E-06 | 0.19732 | 299 | 0.146464584 | 0.007913052 |
| SNP_A.K_308126_2_7474982 | 2 | 7474982 | 5.60E-06 | 0.17057 | 299 | 0.146461799 | 0.007913052 |
| SNP_A.K_1005183_6_6989633 | 6 | 6989633 | 5.61E-06 | 0.25418 | 299 | 0.146447921 | 0.007924801 |
| SNP_A.K_978825_6_1785875 | 6 | 1785875 | 5.65E-06 | 0.29766 | 299 | 0.146407964 | 0.007964869 |
| SNP_A.K_990332_6_4711168 | 6 | 4711168 | 5.65E-06 | 0.20067 | 299 | 0.146406853 | 0.007964869 |
| SNP_A.K_978210_6_1661287 | 6 | 1661287 | 5.66E-06 | 0.17057 | 299 | 0.146392792 | 0.007976934 |
| SNP_A.K_307082_2_7378311 | 2 | 7378311 | 5.68E-06 | 0.24582 | 299 | 0.146374509 | 0.007994281 |
| SNP_A.K_307287_2_7391370 | 2 | 7391370 | 5.70E-06 | 0.24415 | 299 | 0.146349124 | 0.008016194 |
| SNP_A.K_1390256_8_10167379 | 8 | 10167379 | 5.70E-06 | 0.05017 | 299 | 0.146344593 | 0.008016194 |
| SNP_A.K_307403_2_7401413 | 2 | 7401413 | 5.70E-06 | 0.24749 | 299 | 0.146343932 | 0.008016194 |
| SNP_A.K_1005536_6_7019983 | 6 | 7019983 | 5.71E-06 | 0.24916 | 299 | 0.146338525 | 0.008017538 |
| SNP_A.K_306712_2_7343155 | 2 | 7343155 | 5.73E-06 | 0.27592 | 299 | 0.146316276 | 0.008039736 |
| SNP_A.K_1001171_6_6450938 | 6 | 6450938 | 5.73E-06 | 0.28094 | 299 | 0.14631213 | 0.008039736 |
| SNP_A.K_1025193_6_10030439 | 6 | 10030439 | 5.78E-06 | 0.0301 | 299 | 0.146255807 | 0.008105158 |
| SNP_A.K_678621_4_991792 | 4 | 991792 | 5.79E-06 | 0.06689 | 299 | 0.14624915 | 0.008108112 |
| SNP_A.K_990371_6_4712607 | 6 | 4712607 | 5.79E-06 | 0.19732 | 299 | 0.146242394 | 0.008111196 |
| SNP_A.K_992708_6_5674853 | 6 | 5674853 | 5.84E-06 | 0.13211 | 299 | 0.14619382 | 0.008164029 |
| SNP_A.K_996311_6_5983227 | 6 | 5983227 | 5.84E-06 | 0.27759 | 299 | 0.146192118 | 0.008164029 |
| SNP_A.K_306920_2_7367120 | 2 | 7367120 | 5.87E-06 | 0.27926 | 299 | 0.146162794 | 0.008195928 |
| SNP_A.K_978609_6_1740760 | 6 | 1740760 | 5.87E-06 | 0.25418 | 299 | 0.146157699 | 0.008196939 |
| SNP_A.K_978936_6_1814161 | 6 | 1814161 | 5.89E-06 | 0.32776 | 299 | 0.146137691 | 0.008217034 |
| SNP_A.K_999409_6_6286620 | 6 | 6286620 | 5.90E-06 | 0.15886 | 299 | 0.146123073 | 0.008230264 |
| SNP_A.K_1004222_6_6851166 | 6 | 6851166 | 5.92E-06 | 0.33278 | 299 | 0.146102637 | 0.008247182 |
| SNP_A.K_1003935_6_6806403 | 6 | 6806403 | 5.92E-06 | 0.33278 | 299 | 0.146100597 | 0.008247182 |

| | | | | | | | |
|---|---|---|---|---|---|---|---|
| SNP_A.K_710027_4_8309642 | 4 | 8309642 | 5.93E-06 | 0.07023 | 299 | 0.146095085 | 0.008247182 |
| SNP_A.K_528009_3_11220328 | 3 | 11220328 | 5.93E-06 | 0.01171 | 299 | 0.146092738 | 0.008247182 |
| SNP_A.K_1005898_6_7050648 | 6 | 7050648 | 5.96E-06 | 0.28595 | 299 | 0.146066538 | 0.008275411 |
| SNP_A.K_990270_6_4703934 | 6 | 4703934 | 5.97E-06 | 0.2291 | 299 | 0.146053287 | 0.008286993 |
| SNP_A.K_1005403_6_7012406 | 6 | 7012406 | 6.02E-06 | 0.29933 | 299 | 0.14600063 | 0.008349096 |
| SNP_A.K_978476_6_1699399 | 6 | 1699399 | 6.03E-06 | 0.26254 | 299 | 0.145989765 | 0.008349096 |
| SNP_A.K_1004224_6_6851623 | 6 | 6851623 | 6.03E-06 | 0.33278 | 299 | 0.145986983 | 0.008349096 |
| SNP_A.K_1830617_11_10570973 | 11 | 10570973 | 6.03E-06 | 0.17224 | 299 | 0.145984344 | 0.008349096 |
| SNP_A.K_1001238_6_6457002 | 6 | 6457002 | 6.03E-06 | 0.38127 | 299 | 0.145984018 | 0.008349096 |
| SNP_A.K_1004387_6_6878844 | 6 | 6878844 | 6.04E-06 | 0.3311 | 299 | 0.145979783 | 0.008349096 |
| SNP_A.K_978401_6_1687237 | 6 | 1687237 | 6.06E-06 | 0.25585 | 299 | 0.145955965 | 0.008374598 |
| SNP_A.K_1001516_6_6486229 | 6 | 6486229 | 6.07E-06 | 0.301 | 299 | 0.145947386 | 0.008380244 |
| SNP_A.K_995339_6_5926600 | 6 | 5926600 | 6.08E-06 | 0.13378 | 299 | 0.145938314 | 0.008386543 |
| SNP_A.K_978668_6_1752902 | 6 | 1752902 | 6.10E-06 | 0.28763 | 299 | 0.145916632 | 0.008404843 |
| SNP_A.K_1001704_6_6507129 | 6 | 6507129 | 6.10E-06 | 0.29431 | 299 | 0.145915837 | 0.008404843 |
| SNP_A.K_990370_6_4712590 | 6 | 4712590 | 6.14E-06 | 0.19732 | 299 | 0.145865836 | 0.008465032 |
| SNP_A.K_529113_3_11375408 | 3 | 11375408 | 6.17E-06 | 0.01171 | 299 | 0.145843577 | 0.008488852 |
| SNP_A.K_673216_4_336353 | 4 | 336353 | 6.19E-06 | 0.11204 | 299 | 0.145823236 | 0.008510197 |
| SNP_A.K_1005617_6_7030227 | 6 | 7030227 | 6.21E-06 | 0.24247 | 299 | 0.145798405 | 0.008537578 |
| SNP_A.K_389593_2_19885960 | 2 | 19885960 | 6.23E-06 | 0.02676 | 299 | 0.145779602 | 0.008557001 |
| SNP_A.K_1289457_7_22448833 | 7 | 22448833 | 6.27E-06 | 0.07358 | 299 | 0.145735103 | 0.00861092 |
| SNP_A.K_1004923_6_6952651 | 6 | 6952651 | 6.31E-06 | 0.33278 | 299 | 0.145690841 | 0.008664866 |
| SNP_A.K_306771_2_7349386 | 2 | 7349386 | 6.35E-06 | 0.23913 | 299 | 0.145652098 | 0.008711654 |
| SNP_A.K_977917_6_1621323 | 6 | 1621323 | 6.36E-06 | 0.17224 | 299 | 0.145641962 | 0.008719696 |
| SNP_A.K_978110_6_1647552 | 6 | 1647552 | 6.37E-06 | 0.17391 | 299 | 0.145631213 | 0.008728064 |
| SNP_A.K_1953760_12_3627401 | 12 | 3627401 | 6.38E-06 | 0.05518 | 299 | 0.14562738 | 0.008728064 |
| SNP_A.K_1602014_9_19103639 | 9 | 19103639 | 6.40E-06 | 0.12375 | 299 | 0.145609515 | 0.008746676 |
| SNP_A.K_1004329_6_6867306 | 6 | 6867306 | 6.40E-06 | 0.01171 | 299 | 0.145600266 | 0.008753555 |
| SNP_A.K_306720_2_7345201 | 2 | 7345201 | 6.42E-06 | 0.26421 | 299 | 0.145589914 | 0.008761953 |
| SNP_A.K_978437_6_1693594 | 6 | 1693594 | 6.43E-06 | 0.2709 | 299 | 0.145573981 | 0.008778002 |
| SNP_A.K_307392_2_7400563 | 2 | 7400563 | 6.47E-06 | 0.23077 | 299 | 0.145535574 | 0.008824978 |
| SNP_A.K_978747_6_1775018 | 6 | 1775018 | 6.50E-06 | 0.36455 | 299 | 0.145504948 | 0.008861447 |
| SNP_A.K_1001313_6_6464352 | 6 | 6464352 | 6.51E-06 | 0.28595 | 299 | 0.145494825 | 0.008868536 |
| SNP_A.K_978868_6_1793273 | 6 | 1793273 | 6.52E-06 | 0.32776 | 299 | 0.145488596 | 0.008868536 |
| SNP_A.K_1001533_6_6487629 | 6 | 6487629 | 6.52E-06 | 0.29766 | 299 | 0.145487246 | 0.008868536 |
| SNP_A.K_308316_2_7490145 | 2 | 7490145 | 6.52E-06 | 0.25585 | 299 | 0.145482568 | 0.008869216 |
| SNP_A.K_1601887_9_19087252 | 9 | 19087252 | 6.56E-06 | 0.1087 | 299 | 0.145450584 | 0.008896908 |
| SNP_A.K_1830739_11_10577097 | 11 | 10577097 | 6.56E-06 | 0.21906 | 299 | 0.145445531 | 0.008896908 |
| SNP_A.K_1005622_6_7030707 | 6 | 7030707 | 6.57E-06 | 0.24415 | 299 | 0.145439844 | 0.008896908 |
| SNP_A.K_1949539_12_2311934 | 12 | 2311934 | 6.57E-06 | 0.03512 | 299 | 0.145435135 | 0.008896908 |
| SNP_A.K_992476_6_5652202 | 6 | 5652202 | 6.58E-06 | 0.19565 | 299 | 0.145431457 | 0.008896908 |
| SNP_A.K_1004957_6_6957442 | 6 | 6957442 | 6.58E-06 | 0.34448 | 299 | 0.145431439 | 0.008896908 |
| SNP_A.K_1004935_6_6954996 | 6 | 6954996 | 6.58E-06 | 0.30435 | 299 | 0.145431017 | 0.008896908 |
| SNP_A.K_923814_5_19502484 | 5 | 19502484 | 6.58E-06 | 0.02007 | 299 | 0.145429196 | 0.008896908 |
| SNP_A.K_1004303_6_6864422 | 6 | 6864422 | 6.59E-06 | 0.39967 | 299 | 0.145415695 | 0.008904858 |
| SNP_A.K_456464_2_33040299 | 2 | 33040299 | 6.59E-06 | 0.12542 | 299 | 0.145413063 | 0.008904858 |
| SNP_A.K_999905_6_6332100 | 6 | 6332100 | 6.60E-06 | 0.19732 | 299 | 0.145408783 | 0.008904858 |
| SNP_A.K_2080297_12_27255245 | 12 | 27255245 | 6.60E-06 | 0.01171 | 299 | 0.145406842 | 0.008904858 |
| SNP_A.K_307144_2_7381532 | 2 | 7381532 | 6.66E-06 | 0.28261 | 299 | 0.145348518 | 0.008977098 |
| SNP_A.K_1390034_8_10146015 | 8 | 10146015 | 6.66E-06 | 0.04849 | 299 | 0.145346776 | 0.008977098 |

| | | | | | | | |
|---|---|---|---|---|---|---|---|
| SNP_A.K_1364900_8_6053953 | 8 | 6053953 | 6.67E-06 | 0.1087 | 299 | 0.145340417 | 0.008980201 |
| SNP_A.K_308610_2_7529549 | 2 | 7529549 | 6.68E-06 | 0.18896 | 299 | 0.145327669 | 0.008992271 |
| SNP_A.K_307215_2_7385328 | 2 | 7385328 | 6.70E-06 | 0.24582 | 299 | 0.145313046 | 0.009006998 |
| SNP_A.K_679299_4_1070800 | 4 | 1070800 | 6.72E-06 | 0.03177 | 299 | 0.145296604 | 0.009024315 |
| SNP_A.K_1001201_6_6454824 | 6 | 6454824 | 6.75E-06 | 0.29431 | 299 | 0.145268042 | 0.009058792 |
| SNP_A.K_690086_4_4403112 | 4 | 4403112 | 6.75E-06 | 0.05518 | 299 | 0.145260159 | 0.009062922 |
| SNP_A.K_307274_2_7390648 | 2 | 7390648 | 6.76E-06 | 0.27258 | 299 | 0.145256863 | 0.009062922 |
| SNP_A.K_1371150_8_7803778 | 8 | 7803778 | 6.79E-06 | 0.01003 | 299 | 0.145226104 | 0.009100682 |
| SNP_A.K_978899_6_1798773 | 6 | 1798773 | 6.80E-06 | 0.30268 | 299 | 0.145215281 | 0.009110206 |
| SNP_A.K_990407_6_4716115 | 6 | 4716115 | 6.83E-06 | 0.21405 | 299 | 0.145192036 | 0.009137448 |
| SNP_A.K_1123815_6_23600036 | 6 | 23600036 | 6.83E-06 | 0.01338 | 299 | 0.145184061 | 0.009142952 |
| SNP_A.K_990342_6_4711681 | 6 | 4711681 | 6.84E-06 | 0.20234 | 299 | 0.145175543 | 0.00914924 |
| SNP_A.K_1001182_6_6452751 | 6 | 6452751 | 6.85E-06 | 0.28428 | 299 | 0.14516527 | 0.009155586 |
| SNP_A.K_306750_2_7347644 | 2 | 7347644 | 6.86E-06 | 0.25418 | 299 | 0.145161327 | 0.009155586 |
| SNP_A.K_997638_6_6140248 | 6 | 6140248 | 6.86E-06 | 0.14883 | 299 | 0.145158773 | 0.009155586 |
| SNP_A.K_978940_6_1814671 | 6 | 1814671 | 6.88E-06 | 0.33612 | 299 | 0.145137824 | 0.009179697 |
| SNP_A.K_672676_4_269948 | 4 | 269948 | 6.89E-06 | 0.08027 | 299 | 0.145130114 | 0.009184872 |
| SNP_A.K_990379_6_4712788 | 6 | 4712788 | 6.90E-06 | 0.199 | 299 | 0.145121106 | 0.009191915 |
| SNP_A.K_1371622_8_7859409 | 8 | 7859409 | 6.92E-06 | 0.01338 | 299 | 0.145106962 | 0.009206348 |
| SNP_A.K_307530_2_7415968 | 2 | 7415968 | 6.94E-06 | 0.43478 | 299 | 0.14508854 | 0.009224231 |
| SNP_A.K_1001441_6_6481751 | 6 | 6481751 | 6.94E-06 | 0.30435 | 299 | 0.14508635 | 0.009224231 |
| SNP_A.K_997893_6_6155584 | 6 | 6155584 | 6.95E-06 | 0.14548 | 299 | 0.145079338 | 0.009228444 |
| SNP_A.K_1830667_11_10573250 | 11 | 10573250 | 6.99E-06 | 0.16221 | 299 | 0.145041616 | 0.009277046 |
| SNP_A.K_1004269_6_6858759 | 6 | 6858759 | 7.01E-06 | 0.3194 | 299 | 0.145023954 | 0.009296739 |
| SNP_A.K_710104_4_8315938 | 4 | 8315938 | 7.01E-06 | 0.08863 | 299 | 0.145019193 | 0.009297729 |
| SNP_A.K_306773_2_7349477 | 2 | 7349477 | 7.02E-06 | 0.25585 | 299 | 0.145010649 | 0.009304216 |
| SNP_A.K_1364925_8_6059013 | 8 | 6059013 | 7.03E-06 | 0.1204 | 299 | 0.145005168 | 0.009306261 |
| SNP_A.K_1005554_6_7024152 | 6 | 7024152 | 7.04E-06 | 0.27592 | 299 | 0.144991397 | 0.009319906 |
| SNP_A.K_1001213_6_6455233 | 6 | 6455233 | 7.05E-06 | 0.28763 | 299 | 0.144987646 | 0.009319906 |
| SNP_A.K_456443_2_33037266 | 2 | 33037266 | 7.07E-06 | 0.12542 | 299 | 0.144969297 | 0.009340718 |
| SNP_A.K_1004996_6_6960932 | 6 | 6960932 | 7.08E-06 | 0.32107 | 299 | 0.144960896 | 0.009347048 |
| SNP_A.K_1608608_9_20794082 | 9 | 20794082 | 7.11E-06 | 0.24916 | 299 | 0.144927901 | 0.00938468 |
| SNP_A.K_978779_6_1780142 | 6 | 1780142 | 7.11E-06 | 0.32441 | 299 | 0.14492705 | 0.00938468 |
| SNP_A.K_709946_4_8302690 | 4 | 8302690 | 7.13E-06 | 0.0786 | 299 | 0.144916787 | 0.009392312 |
| SNP_A.K_978012_6_1636252 | 6 | 1636252 | 7.13E-06 | 0.17391 | 299 | 0.144913736 | 0.009392312 |
| SNP_A.K_978099_6_1647177 | 6 | 1647177 | 7.13E-06 | 0.17224 | 299 | 0.144908887 | 0.009393484 |
| SNP_A.K_307348_2_7398308 | 2 | 7398308 | 7.14E-06 | 0.22742 | 299 | 0.144902896 | 0.009396335 |
| SNP_A.K_1005030_6_6963011 | 6 | 6963011 | 7.15E-06 | 0.32441 | 299 | 0.144895232 | 0.009401647 |
| SNP_A.K_1001535_6_6487891 | 6 | 6487891 | 7.16E-06 | 0.29933 | 299 | 0.144882289 | 0.009411635 |
| SNP_A.K_978831_6_1787037 | 6 | 1787037 | 7.17E-06 | 0.36789 | 299 | 0.144877436 | 0.009411635 |
| SNP_A.K_993148_6_5710593 | 6 | 5710593 | 7.17E-06 | 0.21405 | 299 | 0.144876314 | 0.009411635 |
| SNP_A.K_1433180_8_16622122 | 8 | 16622122 | 7.19E-06 | 0.08863 | 299 | 0.144862616 | 0.009425851 |
| SNP_A.K_978193_6_1659700 | 6 | 1659700 | 7.20E-06 | 0.25251 | 299 | 0.144853561 | 0.009429921 |
| SNP_A.K_995345_6_5927037 | 6 | 5927037 | 7.20E-06 | 0.22408 | 299 | 0.14485179 | 0.009429921 |
| SNP_A.K_710341_4_8336943 | 4 | 8336943 | 7.22E-06 | 0.07692 | 299 | 0.144828982 | 0.009455256 |
| SNP_A.K_673349_4_350462 | 4 | 350462 | 7.23E-06 | 0.14047 | 299 | 0.144826203 | 0.009455256 |
| SNP_A.K_709874_4_8300085 | 4 | 8300085 | 7.23E-06 | 0.06856 | 299 | 0.144822541 | 0.009455256 |
| SNP_A.K_994602_6_5875390 | 6 | 5875390 | 7.25E-06 | 0.27926 | 299 | 0.144809039 | 0.009469272 |
| SNP_A.K_307443_2_7407758 | 2 | 7407758 | 7.26E-06 | 0.25251 | 299 | 0.144797164 | 0.009480904 |
| SNP_A.K_978311_6_1672704 | 6 | 1672704 | 7.27E-06 | 0.1806 | 299 | 0.144784492 | 0.009493736 |

| | | | | | | | |
|---|---|---|---|---|---|---|---|
| SNP_A.K_1830674_11_10573648 | 11 | 10573648 | 7.29E-06 | 0.16221 | 299 | 0.144773088 | 0.009502701 |
| SNP_A.K_1433026_8_16611474 | 8 | 16611474 | 7.30E-06 | 0.08194 | 299 | 0.144763907 | 0.009502701 |
| SNP_A.K_999720_6_6309357 | 6 | 6309357 | 7.30E-06 | 0.11204 | 299 | 0.144761966 | 0.009502701 |
| SNP_A.K_709627_4_8278535 | 4 | 8278535 | 7.30E-06 | 0.08361 | 299 | 0.144761673 | 0.009502701 |
| SNP_A.K_978097_6_1647094 | 6 | 1647094 | 7.30E-06 | 0.17057 | 299 | 0.144758418 | 0.009502701 |
| SNP_A.K_1001140_6_6448787 | 6 | 6448787 | 7.32E-06 | 0.28763 | 299 | 0.144742347 | 0.009520381 |
| SNP_A.K_1788566_11_4353615 | 11 | 4353615 | 7.33E-06 | 0.01171 | 299 | 0.144738525 | 0.009520381 |
| SNP_A.K_306782_2_7350075 | 2 | 7350075 | 7.35E-06 | 0.2408 | 299 | 0.14471866 | 0.009544018 |
| SNP_A.K_997883_6_6155434 | 6 | 6155434 | 7.37E-06 | 0.14548 | 299 | 0.144698229 | 0.009568566 |
| SNP_A.K_1001291_6_6462734 | 6 | 6462734 | 7.38E-06 | 0.32274 | 299 | 0.144690886 | 0.009573581 |
| SNP_A.K_307087_2_7378353 | 2 | 7378353 | 7.41E-06 | 0.4097 | 299 | 0.144670341 | 0.009598085 |
| SNP_A.K_195475_1_31201552 | 1 | 31201552 | 7.41E-06 | 0.46656 | 299 | 0.144666557 | 0.009598085 |
| SNP_A.K_1000990_6_6434253 | 6 | 6434253 | 7.42E-06 | 0.33278 | 299 | 0.14466075 | 0.009600823 |
| SNP_A.K_977937_6_1622933 | 6 | 1622933 | 7.43E-06 | 0.16722 | 299 | 0.144652248 | 0.009602746 |
| SNP_A.K_307222_2_7386206 | 2 | 7386206 | 7.43E-06 | 0.2291 | 299 | 0.144651514 | 0.009602746 |
| SNP_A.K_978650_6_1750423 | 6 | 1750423 | 7.44E-06 | 0.29097 | 299 | 0.144641961 | 0.009604041 |
| SNP_A.K_978911_6_1803185 | 6 | 1803185 | 7.44E-06 | 0.32274 | 299 | 0.144639412 | 0.009604041 |
| SNP_A.K_977967_6_1626332 | 6 | 1626332 | 7.44E-06 | 0.24582 | 299 | 0.144638738 | 0.009604041 |
| SNP_A.K_1830769_11_10580048 | 11 | 10580048 | 7.45E-06 | 0.22241 | 299 | 0.144629706 | 0.009611647 |
| SNP_A.K_861107_5_3338604 | 5 | 3338604 | 7.48E-06 | 0.11037 | 299 | 0.144608984 | 0.009636852 |
| SNP_A.K_1005360_6_7009541 | 6 | 7009541 | 7.52E-06 | 0.25084 | 299 | 0.14457584 | 0.009680902 |
| SNP_A.K_977902_6_1618714 | 6 | 1618714 | 7.52E-06 | 0.24415 | 299 | 0.144568543 | 0.009685955 |
| SNP_A.K_997890_6_6155559 | 6 | 6155559 | 7.55E-06 | 0.14548 | 299 | 0.144544092 | 0.00971703 |
| SNP_A.K_306884_2_7363474 | 2 | 7363474 | 7.58E-06 | 0.26421 | 299 | 0.144519132 | 0.009748985 |
| SNP_A.K_457094_2_33254450 | 2 | 33254450 | 7.59E-06 | 0.02676 | 299 | 0.144510883 | 0.009755538 |
| SNP_A.K_306857_2_7360809 | 2 | 7360809 | 7.61E-06 | 0.23746 | 299 | 0.144497023 | 0.009770664 |
| SNP_A.K_1005047_6_6965191 | 6 | 6965191 | 7.62E-06 | 0.31104 | 299 | 0.144491239 | 0.009773472 |
| SNP_A.K_978786_6_1781213 | 6 | 1781213 | 7.62E-06 | 0.29599 | 299 | 0.144485172 | 0.009776716 |
| SNP_A.K_990368_6_4712581 | 6 | 4712581 | 7.64E-06 | 0.19732 | 299 | 0.144475143 | 0.009786025 |
| SNP_A.K_1004542_6_6898884 | 6 | 6898884 | 7.66E-06 | 0.30435 | 299 | 0.144453598 | 0.009811568 |
| SNP_A.K_306800_2_7351743 | 2 | 7351743 | 7.66E-06 | 0.2709 | 299 | 0.144450434 | 0.009811568 |
| SNP_A.K_1001680_6_6503208 | 6 | 6503208 | 7.67E-06 | 0.30435 | 299 | 0.144446659 | 0.009811568 |
| SNP_A.K_1004931_6_6954602 | 6 | 6954602 | 7.69E-06 | 0.30769 | 299 | 0.144431721 | 0.009821516 |
| SNP_A.K_978820_6_1784882 | 6 | 1784882 | 7.69E-06 | 0.35786 | 299 | 0.144427073 | 0.009821516 |
| SNP_A.K_401067_2_21817285 | 2 | 21817285 | 7.69E-06 | 0.05184 | 299 | 0.144426052 | 0.009821516 |
| SNP_A.K_1004622_6_6911174 | 6 | 6911174 | 7.70E-06 | 0.29264 | 299 | 0.144422823 | 0.009821516 |
| SNP_A.K_1001077_6_6439881 | 6 | 6439881 | 7.70E-06 | 0.29097 | 299 | 0.144417132 | 0.009821516 |
| SNP_A.K_1004597_6_6909152 | 6 | 6909152 | 7.71E-06 | 0.33445 | 299 | 0.144416628 | 0.009821516 |
| SNP_A.K_195007_1_31034367 | 1 | 31034367 | 7.71E-06 | 0.41973 | 299 | 0.144411884 | 0.009822788 |
| SNP_A.K_456490_2_33046782 | 2 | 33046782 | 7.82E-06 | 0.11538 | 299 | 0.144319535 | 0.009947679 |
| SNP_A.K_307523_2_7415150 | 2 | 7415150 | 7.83E-06 | 0.25251 | 299 | 0.144315733 | 0.009947679 |
| SNP_A.K_1001106_6_6443861 | 6 | 6443861 | 7.83E-06 | 0.28763 | 299 | 0.14431419 | 0.009947679 |
| SNP_A.K_1364901_8_6054078 | 8 | 6054078 | 7.83E-06 | 0.12375 | 299 | 0.144313183 | 0.009947679 |
| SNP_A.K_990381_6_4712821 | 6 | 4712821 | 7.83E-06 | 0.19565 | 299 | 0.144311526 | 0.009947679 |
| SNP_A.K_1608605_9_20793997 | 9 | 20793997 | 7.86E-06 | 0.24247 | 299 | 0.144290107 | 0.009974963 |
| SNP_A.K_306918_2_7366655 | 2 | 7366655 | 7.87E-06 | 0.26421 | 299 | 0.144284039 | 0.009978347 |
| SNP_A.K_326864_2_10061941 | 2 | 10061941 | 7.90E-06 | 0.06187 | 299 | 0.144257925 | 0.010007601 |
| SNP_A.K_978923_6_1804747 | 6 | 1804747 | 7.90E-06 | 0.31773 | 299 | 0.144256976 | 0.010007601 |
| SNP_A.K_1608591_9_20791532 | 9 | 20791532 | 7.90E-06 | 0.24247 | 299 | 0.144253635 | 0.010007601 |
| SNP_A.K_1608618_9_20800390 | 9 | 20800390 | 7.93E-06 | 0.23579 | 299 | 0.144234419 | 0.010024831 |

| | | | | | | | |
|---|---|---|---|---|---|---|---|
| SNP_A.K_1001113_6_6446295 | 6 | 6446295 | 7.93E-06 | 0.2893 | 299 | 0.144231039 | 0.010024831 |
| SNP_A.K_1001083_6_6440024 | 6 | 6440024 | 7.94E-06 | 0.29431 | 299 | 0.144228311 | 0.010024831 |
| SNP_A.K_1005214_6_6995083 | 6 | 6995083 | 7.94E-06 | 0.34114 | 299 | 0.144227093 | 0.010024831 |
| SNP_A.K_1001080_6_6439976 | 6 | 6439976 | 7.95E-06 | 0.28261 | 299 | 0.144214843 | 0.010034256 |
| SNP_A.K_978005_6_1634991 | 6 | 1634991 | 7.95E-06 | 0.17057 | 299 | 0.144213327 | 0.010034256 |
| SNP_A.K_978284_6_1668603 | 6 | 1668603 | 7.98E-06 | 0.24582 | 299 | 0.144195137 | 0.010056744 |
| SNP_A.K_307028_2_7373779 | 2 | 7373779 | 8.01E-06 | 0.25418 | 299 | 0.14416756 | 0.010094094 |
| SNP_A.K_709676_4_8280696 | 4 | 8280696 | 8.03E-06 | 0.10033 | 299 | 0.144153285 | 0.010110533 |
| SNP_A.K_978932_6_1808359 | 6 | 1808359 | 8.09E-06 | 0.32274 | 299 | 0.144105924 | 0.010178776 |
| SNP_A.K_977939_6_1623005 | 6 | 1623005 | 8.09E-06 | 0.1689 | 299 | 0.144102541 | 0.010178776 |
| SNP_A.K_1000853_6_6421185 | 6 | 6421185 | 8.12E-06 | 0.28595 | 299 | 0.14407824 | 0.010211219 |
| SNP_A.K_1782998_11_3531933 | 11 | 3531933 | 8.13E-06 | 0.01171 | 299 | 0.144074474 | 0.010211219 |
| SNP_A.K_994292_6_5839856 | 6 | 5839856 | 8.17E-06 | 0.24247 | 299 | 0.144044482 | 0.01024859 |
| SNP_A.K_194681_1_30914490 | 1 | 30914490 | 8.17E-06 | 0.49666 | 299 | 0.144043406 | 0.01024859 |
| SNP_A.K_752455_4_17906629 | 4 | 17906629 | 8.18E-06 | 0.04348 | 299 | 0.144030837 | 0.010262572 |
| SNP_A.K_978732_6_1771856 | 6 | 1771856 | 8.20E-06 | 0.30769 | 299 | 0.144018843 | 0.01026399 |
| SNP_A.K_307344_2_7397484 | 2 | 7397484 | 8.20E-06 | 0.23579 | 299 | 0.144018704 | 0.01026399 |
| SNP_A.K_307349_2_7398319 | 2 | 7398319 | 8.20E-06 | 0.22742 | 299 | 0.144018411 | 0.01026399 |
| SNP_A.K_977893_6_1616223 | 6 | 1616223 | 8.21E-06 | 0.24582 | 299 | 0.144011869 | 0.010268324 |
| SNP_A.K_201445_1_32182693 | 1 | 32182693 | 8.22E-06 | 0.16555 | 299 | 0.144005927 | 0.010269129 |
| SNP_A.K_1371280_8_7817601 | 8 | 7817601 | 8.22E-06 | 0.01338 | 299 | 0.144003689 | 0.010269129 |
| SNP_A.K_526307_3_10871057 | 3 | 10871057 | 8.23E-06 | 0.01171 | 299 | 0.14399429 | 0.010278069 |
| SNP_A.K_978207_6_1660474 | 6 | 1660474 | 8.25E-06 | 0.17726 | 299 | 0.143981078 | 0.010287446 |
| SNP_A.K_990369_6_4712588 | 6 | 4712588 | 8.25E-06 | 0.19732 | 299 | 0.143980796 | 0.010287446 |
| SNP_A.K_978238_6_1664035 | 6 | 1664035 | 8.26E-06 | 0.17391 | 299 | 0.143973003 | 0.010293827 |
| SNP_A.K_1001310_6_6463980 | 6 | 6463980 | 8.27E-06 | 0.28763 | 299 | 0.143964492 | 0.010301373 |
| SNP_A.K_990248_6_4703038 | 6 | 4703038 | 8.31E-06 | 0.20569 | 299 | 0.14393583 | 0.010341478 |
| SNP_A.K_978826_6_1786490 | 6 | 1786490 | 8.31E-06 | 0.35619 | 299 | 0.143931887 | 0.010341675 |
| SNP_A.K_978804_6_1782855 | 6 | 1782855 | 8.32E-06 | 0.33612 | 299 | 0.143922445 | 0.010350158 |
| SNP_A.K_1000923_6_6424119 | 6 | 6424119 | 8.33E-06 | 0.32943 | 299 | 0.143916573 | 0.010350158 |
| SNP_A.K_978934_6_1813106 | 6 | 1813106 | 8.34E-06 | 0.33612 | 299 | 0.143913286 | 0.010350158 |
| SNP_A.K_1830420_11_10560666 | 11 | 10560666 | 8.34E-06 | 0.15552 | 299 | 0.143911382 | 0.010350158 |
| SNP_A.K_978610_6_1740777 | 6 | 1740777 | 8.35E-06 | 0.26087 | 299 | 0.14390176 | 0.010359573 |
| SNP_A.K_990360_6_4712355 | 6 | 4712355 | 8.37E-06 | 0.20067 | 299 | 0.143887217 | 0.010374151 |
| SNP_A.K_306788_2_7350832 | 2 | 7350832 | 8.37E-06 | 0.23244 | 299 | 0.143885158 | 0.010374151 |
| SNP_A.K_990361_6_4712375 | 6 | 4712375 | 8.39E-06 | 0.19732 | 299 | 0.143871189 | 0.010390665 |
| SNP_A.K_1804142_11_7035804 | 11 | 7035804 | 8.41E-06 | 0.01839 | 299 | 0.143860142 | 0.010402453 |
| SNP_A.K_1830629_11_10571417 | 11 | 10571417 | 8.43E-06 | 0.36957 | 299 | 0.143843877 | 0.010417929 |
| SNP_A.K_1390322_8_10181340 | 8 | 10181340 | 8.43E-06 | 0.04682 | 299 | 0.143843045 | 0.010417929 |
| SNP_A.K_1364871_8_6050143 | 8 | 6050143 | 8.44E-06 | 0.10033 | 299 | 0.143832618 | 0.010422993 |
| SNP_A.K_1600930_9_18862335 | 9 | 18862335 | 8.44E-06 | 0.14047 | 299 | 0.143832354 | 0.010422993 |
| SNP_A.K_1004501_6_6893948 | 6 | 6893948 | 8.46E-06 | 0.30769 | 299 | 0.143817522 | 0.010433629 |
| SNP_A.K_990365_6_4712489 | 6 | 4712489 | 8.47E-06 | 0.20234 | 299 | 0.143814745 | 0.010433629 |
| SNP_A.K_1004062_6_6822849 | 6 | 6822849 | 8.47E-06 | 0.34114 | 299 | 0.143814475 | 0.010433629 |
| SNP_A.K_308323_2_7490402 | 2 | 7490402 | 8.48E-06 | 0.19732 | 299 | 0.143804175 | 0.010444274 |
| SNP_A.K_677721_4_906195 | 4 | 906195 | 8.49E-06 | 0.15385 | 299 | 0.143796714 | 0.01045029 |
| SNP_A.K_978628_6_1747015 | 6 | 1747015 | 8.51E-06 | 0.27592 | 299 | 0.143784652 | 0.010463846 |
| SNP_A.K_1390259_8_10167487 | 8 | 10167487 | 8.53E-06 | 0.04849 | 299 | 0.143763716 | 0.010491984 |
| SNP_A.K_990363_6_4712411 | 6 | 4712411 | 8.55E-06 | 0.19732 | 299 | 0.143749093 | 0.010503612 |
| SNP_A.K_990364_6_4712416 | 6 | 4712416 | 8.55E-06 | 0.19732 | 299 | 0.143749093 | 0.010503612 |

| | | | | | | | |
|---|---|---|---|---|---|---|---|
| SNP_A.K_990357_6_4712317 | 6 | 4712317 | 8.56E-06 | 0.20401 | 299 | 0.143740957 | 0.010510793 |
| SNP_A.K_1004375_6_6877639 | 6 | 6877639 | 8.57E-06 | 0.34281 | 299 | 0.143734416 | 0.010515358 |
| SNP_A.K_710234_4_8323620 | 4 | 8323620 | 8.63E-06 | 0.11706 | 299 | 0.143689835 | 0.010582745 |
| SNP_A.K_308770_2_7552253 | 2 | 7552253 | 8.64E-06 | 0.4097 | 299 | 0.143681256 | 0.010588006 |
| SNP_A.K_990322_6_4710600 | 6 | 4710600 | 8.65E-06 | 0.19732 | 299 | 0.143679138 | 0.010588006 |
| SNP_A.K_1608635_9_20804039 | 9 | 20804039 | 8.66E-06 | 0.23579 | 299 | 0.14367221 | 0.010589412 |
| SNP_A.K_990392_6_4714508 | 6 | 4714508 | 8.66E-06 | 0.20401 | 299 | 0.143668944 | 0.010589412 |
| SNP_A.K_1004396_6_6879402 | 6 | 6879402 | 8.66E-06 | 0.30936 | 299 | 0.143667025 | 0.010589412 |
| SNP_A.K_326590_2_10034363 | 2 | 10034363 | 8.67E-06 | 0.0602 | 299 | 0.143661641 | 0.010592119 |
| SNP_A.K_995278_6_5923820 | 6 | 5923820 | 8.69E-06 | 0.1204 | 299 | 0.143648423 | 0.010603815 |
| SNP_A.K_978405_6_1687961 | 6 | 1687961 | 8.69E-06 | 0.25585 | 299 | 0.143647095 | 0.010603815 |
| SNP_A.K_996292_6_5981704 | 6 | 5981704 | 8.72E-06 | 0.28261 | 299 | 0.143625649 | 0.01062731 |
| SNP_A.K_678411_4_985889 | 4 | 985889 | 8.72E-06 | 0.06689 | 299 | 0.143625469 | 0.01062731 |
| SNP_A.K_995214_6_5920403 | 6 | 5920403 | 8.74E-06 | 0.13712 | 299 | 0.143608605 | 0.010649165 |
| SNP_A.K_600991_3_24583196 | 3 | 24583196 | 8.76E-06 | 0.01839 | 299 | 0.143593923 | 0.010667426 |
| SNP_A.K_709808_4_8298240 | 4 | 8298240 | 8.78E-06 | 0.06856 | 299 | 0.14357967 | 0.010684556 |
| SNP_A.K_710325_4_8336426 | 4 | 8336426 | 8.79E-06 | 0.08528 | 299 | 0.143571397 | 0.010684556 |
| SNP_A.K_990327_6_4710755 | 6 | 4710755 | 8.79E-06 | 0.19231 | 299 | 0.143571363 | 0.010684556 |
| SNP_A.K_1600854_9_18853309 | 9 | 18853309 | 8.80E-06 | 0.12709 | 299 | 0.143565761 | 0.010684556 |
| SNP_A.K_1004419_6_6881380 | 6 | 6881380 | 8.81E-06 | 0.3311 | 299 | 0.143562349 | 0.010684556 |
| SNP_A.K_990325_6_4710703 | 6 | 4710703 | 8.81E-06 | 0.199 | 299 | 0.143560462 | 0.010684556 |
| SNP_A.K_195455_1_31196590 | 1 | 31196590 | 8.81E-06 | 0.47492 | 299 | 0.143557575 | 0.010684556 |
| SNP_A.K_977879_6_1608540 | 6 | 1608540 | 8.82E-06 | 0.1689 | 299 | 0.143551907 | 0.010687815 |
| SNP_A.K_1004914_6_6952274 | 6 | 6952274 | 8.84E-06 | 0.35619 | 299 | 0.14354187 | 0.010694032 |
| SNP_A.K_1004709_6_6922808 | 6 | 6922808 | 8.84E-06 | 0.20401 | 299 | 0.143540761 | 0.010694032 |
| SNP_A.K_308776_2_7552444 | 2 | 7552444 | 8.84E-06 | 0.22742 | 299 | 0.143534905 | 0.010697621 |
| SNP_A.K_1432213_8_16469057 | 8 | 16469057 | 8.88E-06 | 0.07525 | 299 | 0.14350954 | 0.010733947 |
| SNP_A.K_1390011_8_10145219 | 8 | 10145219 | 8.89E-06 | 0.04849 | 299 | 0.14350189 | 0.010740572 |
| SNP_A.K_306977_2_7371459 | 2 | 7371459 | 8.91E-06 | 0.26087 | 299 | 0.143489626 | 0.010754973 |
| SNP_A.K_1608645_9_20809573 | 9 | 20809573 | 8.95E-06 | 0.22742 | 299 | 0.143458982 | 0.010800432 |
| SNP_A.K_978175_6_1657731 | 6 | 1657731 | 8.97E-06 | 0.1689 | 299 | 0.143446968 | 0.010809116 |
| SNP_A.K_1005359_6_7009533 | 6 | 7009533 | 8.98E-06 | 0.25753 | 299 | 0.143440539 | 0.010809116 |
| SNP_A.K_1001442_6_6481777 | 6 | 6481777 | 8.98E-06 | 0.30769 | 299 | 0.143440288 | 0.010809116 |
| SNP_A.K_307323_2_7395007 | 2 | 7395007 | 8.98E-06 | 0.26756 | 299 | 0.143439056 | 0.010809116 |
| SNP_A.K_990348_6_4711834 | 6 | 4711834 | 8.99E-06 | 0.20569 | 299 | 0.143434239 | 0.010811017 |
| SNP_A.K_996198_6_5974181 | 6 | 5974181 | 9.00E-06 | 0.28094 | 299 | 0.143421898 | 0.01082567 |
| SNP_A.K_1830677_11_10573765 | 11 | 10573765 | 9.02E-06 | 0.16555 | 299 | 0.143408661 | 0.010841868 |
| SNP_A.K_1001287_6_6461845 | 6 | 6461845 | 9.10E-06 | 0.28261 | 299 | 0.143356358 | 0.010924719 |
| SNP_A.K_978788_6_1781475 | 6 | 1781475 | 9.11E-06 | 0.31605 | 299 | 0.143348588 | 0.010931712 |
| SNP_A.K_978929_6_1808243 | 6 | 1808243 | 9.14E-06 | 0.33445 | 299 | 0.143326686 | 0.010962949 |
| SNP_A.K_2080240_12_27248622 | 12 | 27248622 | 9.15E-06 | 0.01171 | 299 | 0.143318082 | 0.010971407 |
| SNP_A.K_1433082_8_16617064 | 8 | 16617064 | 9.16E-06 | 0.08696 | 299 | 0.143312071 | 0.010975417 |
| SNP_A.K_978736_6_1772612 | 6 | 1772612 | 9.17E-06 | 0.33612 | 299 | 0.143305861 | 0.010979775 |
| SNP_A.K_999260_6_6271990 | 6 | 6271990 | 9.19E-06 | 0.16722 | 299 | 0.143290405 | 0.011000055 |
| SNP_A.K_1804486_11_7079280 | 11 | 7079280 | 9.20E-06 | 0.01839 | 299 | 0.143280305 | 0.011011137 |
| SNP_A.K_709792_4_8297830 | 4 | 8297830 | 9.23E-06 | 0.07023 | 299 | 0.143264431 | 0.011025871 |
| SNP_A.K_709793_4_8297839 | 4 | 8297839 | 9.23E-06 | 0.07023 | 299 | 0.143264431 | 0.011025871 |
| SNP_A.K_1001842_6_6524271 | 6 | 6524271 | 9.24E-06 | 0.30435 | 299 | 0.143255585 | 0.011031226 |
| SNP_A.K_1196545_7_7627744 | 7 | 7627744 | 9.24E-06 | 0.01505 | 299 | 0.143253807 | 0.011031226 |
| SNP_A.K_978865_6_1792217 | 6 | 1792217 | 9.25E-06 | 0.29599 | 299 | 0.143250345 | 0.011031226 |

| | | | | | | | |
|---|---|---|---|---|---|---|---|
| SNP_A.K_709848_4_8299533 | 4 | 8299533 | 9.26E-06 | 0.07023 | 299 | 0.143240813 | 0.011041381 |
| SNP_A.K_1370949_8_7793086 | 8 | 7793086 | 9.30E-06 | 0.01338 | 299 | 0.143213613 | 0.011082172 |
| SNP_A.K_1005544_6_7022810 | 6 | 7022810 | 9.33E-06 | 0.29766 | 299 | 0.143195337 | 0.011107584 |
| SNP_A.K_401149_2_21824321 | 2 | 21824321 | 9.33E-06 | 0.05017 | 299 | 0.143190605 | 0.011109465 |
| SNP_A.K_2080313_12_27256266 | 12 | 27256266 | 9.35E-06 | 0.01338 | 299 | 0.143179097 | 0.011123148 |
| SNP_A.K_990507_6_4722308 | 6 | 4722308 | 9.36E-06 | 0.20903 | 299 | 0.143170488 | 0.011128015 |
| SNP_A.K_1608606_9_20794000 | 9 | 20794000 | 9.37E-06 | 0.24415 | 299 | 0.143168676 | 0.011128015 |
| SNP_A.K_1600942_9_18863483 | 9 | 18863483 | 9.37E-06 | 0.12876 | 299 | 0.14316537 | 0.011128015 |
| SNP_A.K_978205_6_1660310 | 6 | 1660310 | 9.38E-06 | 0.18227 | 299 | 0.143158367 | 0.011132802 |
| SNP_A.K_978182_6_1658336 | 6 | 1658336 | 9.39E-06 | 0.17391 | 299 | 0.143155143 | 0.011132802 |
| SNP_A.K_306905_2_7364891 | 2 | 7364891 | 9.39E-06 | 0.2408 | 299 | 0.143151709 | 0.011132802 |
| SNP_A.K_1001221_6_6455512 | 6 | 6455512 | 9.42E-06 | 0.35953 | 299 | 0.143134044 | 0.011157296 |
| SNP_A.K_719171_4_10646119 | 4 | 10646119 | 9.45E-06 | 0.06856 | 299 | 0.143111461 | 0.011190469 |
| SNP_A.K_307472_2_7409894 | 2 | 7409894 | 9.47E-06 | 0.25251 | 299 | 0.143096032 | 0.011211172 |
| SNP_A.K_1365034_8_6067527 | 8 | 6067527 | 9.48E-06 | 0.10201 | 299 | 0.143088784 | 0.011217534 |
| SNP_A.K_690714_4_4490974 | 4 | 4490974 | 9.49E-06 | 0.06187 | 299 | 0.143083148 | 0.011221067 |
| SNP_A.K_709525_4_8271593 | 4 | 8271593 | 9.50E-06 | 0.08361 | 299 | 0.143075949 | 0.011227354 |
| SNP_A.K_978190_6_1659353 | 6 | 1659353 | 9.61E-06 | 0.17559 | 299 | 0.143002797 | 0.011350331 |
| SNP_A.K_1804471_11_7078714 | 11 | 7078714 | 9.65E-06 | 0.01505 | 299 | 0.142976422 | 0.011387898 |
| SNP_A.K_978402_6_1687864 | 6 | 1687864 | 9.66E-06 | 0.25585 | 299 | 0.142974473 | 0.011387898 |
| SNP_A.K_1349475_8_3699993 | 8 | 3699993 | 9.69E-06 | 0.01672 | 299 | 0.142953728 | 0.011418507 |
| SNP_A.K_1001010_6_6436341 | 6 | 6436341 | 9.70E-06 | 0.2893 | 299 | 0.14294822 | 0.011421898 |
| SNP_A.K_990336_6_4711317 | 6 | 4711317 | 9.70E-06 | 0.20736 | 299 | 0.14294253 | 0.011425621 |
| SNP_A.K_1004338_6_6869730 | 6 | 6869730 | 9.73E-06 | 0.30435 | 299 | 0.142928763 | 0.011443822 |
| SNP_A.K_1000874_6_6421756 | 6 | 6421756 | 9.73E-06 | 0.28763 | 299 | 0.142924085 | 0.011445745 |
| SNP_A.K_978171_6_1657666 | 6 | 1657666 | 9.75E-06 | 0.2709 | 299 | 0.142913419 | 0.011456045 |
| SNP_A.K_1005480_6_7016563 | 6 | 7016563 | 9.75E-06 | 0.30602 | 299 | 0.142911139 | 0.011456045 |
| SNP_A.K_978692_6_1764553 | 6 | 1764553 | 9.80E-06 | 0.2709 | 299 | 0.142878714 | 0.011507911 |
| SNP_A.K_1005347_6_7008789 | 6 | 7008789 | 9.81E-06 | 0.37625 | 299 | 0.142873465 | 0.01151089 |
| SNP_A.K_1004779_6_6930481 | 6 | 6930481 | 9.82E-06 | 0.29599 | 299 | 0.142868476 | 0.011513403 |
| SNP_A.K_1382752_8_9115654 | 8 | 9115654 | 9.84E-06 | 0.27759 | 299 | 0.142852759 | 0.011535298 |
| SNP_A.K_1001155_6_6449633 | 6 | 6449633 | 9.85E-06 | 0.28595 | 299 | 0.142845889 | 0.011541227 |
| SNP_A.K_978404_6_1687944 | 6 | 1687944 | 9.86E-06 | 0.25251 | 299 | 0.142838379 | 0.01154832 |
| SNP_A.K_994847_6_5894454 | 6 | 5894454 | 9.87E-06 | 0.29431 | 299 | 0.142832762 | 0.011551995 |
| SNP_A.K_1830697_11_10574825 | 11 | 10574825 | 9.90E-06 | 0.17224 | 299 | 0.142817374 | 0.011572554 |
| SNP_A.K_996389_6_5998902 | 6 | 5998902 | 9.90E-06 | 0.11204 | 299 | 0.142814248 | 0.011572554 |
| SNP_A.K_978186_6_1658488 | 6 | 1658488 | 9.91E-06 | 0.25418 | 299 | 0.142808252 | 0.011576934 |
| SNP_A.K_129805_1_18618568 | 1 | 18618568 | 9.94E-06 | 0.35953 | 299 | 0.142792825 | 0.011598452 |
| SNP_A.K_995628_6_5943898 | 6 | 5943898 | 9.98E-06 | 0.36789 | 299 | 0.142763456 | 0.011645437 |
| SNP_A.K_1608491_9_20763576 | 9 | 20763576 | 9.99E-06 | 0.27425 | 299 | 0.142758844 | 0.011647329 |
| SNP_A.K_1000873_6_6421751 | 6 | 6421751 | 1.00E-05 | 0.28595 | 299 | 0.142750618 | 0.011655685 |
| SNP_A.K_1005199_6_6993058 | 6 | 6993058 | 1.00E-05 | 0.24582 | 299 | 0.142747125 | 0.011655685 |
| SNP_A.K_978716_6_1770131 | 6 | 1770131 | 1.00E-05 | 0.32441 | 299 | 0.142738361 | 0.011665181 |
| SNP_A.K_709787_4_8297724 | 4 | 8297724 | 1.00E-05 | 0.07023 | 299 | 0.142731199 | 0.011671712 |
| SNP_A.K_978762_6_1776093 | 6 | 1776093 | 1.00E-05 | 0.36789 | 299 | 0.142727659 | 0.011671712 |
| SNP_A.K_1608642_9_20809180 | 9 | 20809180 | 1.00E-05 | 0.29599 | 299 | 0.142723154 | 0.011673001 |
| SNP_A.K_15401_1_2381814 | 1 | 2381814 | 1.01E-05 | 0.02007 | 299 | 0.142718594 | 0.011673001 |
| SNP_A.K_995836_6_5952246 | 6 | 5952246 | 1.01E-05 | 0.29766 | 299 | 0.142716271 | 0.011673001 |
| SNP_A.K_196309_1_31386612 | 1 | 31386612 | 1.01E-05 | 0.06689 | 299 | 0.142700885 | 0.011685889 |
| SNP_A.K_456473_2_33042482 | 2 | 33042482 | 1.01E-05 | 0.12876 | 299 | 0.142699791 | 0.011685889 |

| | | | | | | | |
|---|---|---|---|---|---|---|---|
| SNP_A.K_990362_6_4712388 | 6 | 4712388 | 1.01E-05 | 0.19398 | 299 | 0.142698568 | 0.011685889 |
| SNP_A.K_1005008_6_6961751 | 6 | 6961751 | 1.01E-05 | 0.32943 | 299 | 0.142693501 | 0.011688666 |
| SNP_A.K_709846_4_8299492 | 4 | 8299492 | 1.02E-05 | 0.07358 | 299 | 0.142654071 | 0.011754602 |
| SNP_A.K_710086_4_8315557 | 4 | 8315557 | 1.02E-05 | 0.0786 | 299 | 0.142632424 | 0.011785062 |
| SNP_A.K_1433032_8_16612288 | 8 | 16612288 | 1.02E-05 | 0.08194 | 299 | 0.142630473 | 0.011785062 |
| SNP_A.K_990380_6_4712804 | 6 | 4712804 | 1.02E-05 | 0.20067 | 299 | 0.142623407 | 0.011791574 |
| SNP_A.K_673427_4_361172 | 4 | 361172 | 1.02E-05 | 0.12876 | 299 | 0.142606253 | 0.011816768 |
| SNP_A.K_978288_6_1669353 | 6 | 1669353 | 1.02E-05 | 0.24916 | 299 | 0.142596893 | 0.011827561 |
| SNP_A.K_1183725_7_4599480 | 7 | 4599480 | 1.03E-05 | 0.0602 | 299 | 0.142589224 | 0.011835226 |
| SNP_A.K_308318_2_7490175 | 2 | 7490175 | 1.03E-05 | 0.2592 | 299 | 0.142584831 | 0.01183682 |
| SNP_A.K_1364919_8_6056951 | 8 | 6056951 | 1.03E-05 | 0.12876 | 299 | 0.142574891 | 0.011838976 |
| SNP_A.K_978874_6_1793756 | 6 | 1793756 | 1.03E-05 | 0.32776 | 299 | 0.142572397 | 0.011838976 |
| SNP_A.K_977930_6_1622032 | 6 | 1622032 | 1.03E-05 | 0.20569 | 299 | 0.142571047 | 0.011838976 |
| SNP_A.K_978197_6_1659789 | 6 | 1659789 | 1.03E-05 | 0.17224 | 299 | 0.142569551 | 0.011838976 |
| SNP_A.K_978180_6_1658262 | 6 | 1658262 | 1.03E-05 | 0.17893 | 299 | 0.142564672 | 0.011841492 |
| SNP_A.K_990282_6_4704596 | 6 | 4704596 | 1.03E-05 | 0.20067 | 299 | 0.142559644 | 0.011844288 |
| SNP_A.K_990230_6_4701471 | 6 | 4701471 | 1.03E-05 | 0.20569 | 299 | 0.142536277 | 0.011881204 |
| SNP_A.K_277614_2_2954938 | 2 | 2954938 | 1.04E-05 | 0.06689 | 299 | 0.142527921 | 0.01188327 |
| SNP_A.K_978074_6_1645345 | 6 | 1645345 | 1.04E-05 | 0.17559 | 299 | 0.14252489 | 0.01188327 |
| SNP_A.K_1001472_6_6482907 | 6 | 6482907 | 1.04E-05 | 0.2893 | 299 | 0.142521458 | 0.01188327 |
| SNP_A.K_678728_4_995364 | 4 | 995364 | 1.04E-05 | 0.07023 | 299 | 0.142515498 | 0.01188327 |
| SNP_A.K_978765_6_1777372 | 6 | 1777372 | 1.04E-05 | 0.33445 | 299 | 0.142514267 | 0.01188327 |
| SNP_A.K_307394_2_7400887 | 2 | 7400887 | 1.04E-05 | 0.2592 | 299 | 0.142513661 | 0.01188327 |
| SNP_A.K_978178_6_1657844 | 6 | 1657844 | 1.04E-05 | 0.19732 | 299 | 0.14251058 | 0.01188327 |
| SNP_A.K_1005584_6_7028359 | 6 | 7028359 | 1.04E-05 | 0.33278 | 299 | 0.142503531 | 0.011889875 |
| SNP_A.K_978095_6_1647085 | 6 | 1647085 | 1.04E-05 | 0.17391 | 299 | 0.142484869 | 0.011918174 |
| SNP_A.K_990444_6_4718339 | 6 | 4718339 | 1.05E-05 | 0.47826 | 299 | 0.142467019 | 0.011945024 |
| SNP_A.K_306875_2_7362996 | 2 | 7362996 | 1.05E-05 | 0.27926 | 299 | 0.142460841 | 0.011946378 |
| SNP_A.K_1608557_9_20777260 | 9 | 20777260 | 1.05E-05 | 0.23746 | 299 | 0.142459301 | 0.011946378 |
| SNP_A.K_306770_2_7349370 | 2 | 7349370 | 1.05E-05 | 0.23746 | 299 | 0.142440238 | 0.011972646 |
| SNP_A.K_307100_2_7379180 | 2 | 7379180 | 1.05E-05 | 0.24749 | 299 | 0.142437838 | 0.011972646 |
| SNP_A.K_709872_4_8300067 | 4 | 8300067 | 1.05E-05 | 0.07191 | 299 | 0.142434817 | 0.011972646 |
| SNP_A.K_978880_6_1794363 | 6 | 1794363 | 1.05E-05 | 0.301 | 299 | 0.14242705 | 0.011979773 |
| SNP_A.K_977903_6_1618777 | 6 | 1618777 | 1.05E-05 | 0.24582 | 299 | 0.142418002 | 0.011979773 |
| SNP_A.K_978300_6_1671803 | 6 | 1671803 | 1.05E-05 | 0.25251 | 299 | 0.142417704 | 0.011979773 |
| SNP_A.K_1433066_8_16616204 | 8 | 16616204 | 1.05E-05 | 0.08194 | 299 | 0.142412924 | 0.011979773 |
| SNP_A.K_1830766_11_10579998 | 11 | 10579998 | 1.05E-05 | 0.28261 | 299 | 0.142410812 | 0.011979773 |
| SNP_A.K_1855176_11_14565502 | 11 | 14565502 | 1.05E-05 | 0.03512 | 299 | 0.142410117 | 0.011979773 |
| SNP_A.K_1608538_9_20773152 | 9 | 20773152 | 1.06E-05 | 0.32609 | 299 | 0.14239613 | 0.011993486 |
| SNP_A.K_678203_4_968678 | 4 | 968678 | 1.06E-05 | 0.14214 | 299 | 0.142392363 | 0.011993486 |
| SNP_A.K_280510_2_3619907 | 2 | 3619907 | 1.06E-05 | 0.01338 | 299 | 0.142389506 | 0.011993486 |
| SNP_A.K_1001534_6_6487642 | 6 | 6487642 | 1.06E-05 | 0.301 | 299 | 0.142388925 | 0.011993486 |
| SNP_A.K_672551_4_259476 | 4 | 259476 | 1.06E-05 | 0.10201 | 299 | 0.142380361 | 0.012003072 |
| SNP_A.K_1005216_6_6995106 | 6 | 6995106 | 1.06E-05 | 0.30435 | 299 | 0.142373616 | 0.012007637 |
| SNP_A.K_977978_6_1628167 | 6 | 1628167 | 1.06E-05 | 0.24582 | 299 | 0.142370741 | 0.012007637 |
| SNP_A.K_978707_6_1769205 | 6 | 1769205 | 1.06E-05 | 0.3495 | 299 | 0.142367539 | 0.012007637 |
| SNP_A.K_306974_2_7371290 | 2 | 7371290 | 1.07E-05 | 0.24582 | 299 | 0.14233782 | 0.012057193 |
| SNP_A.K_307508_2_7413925 | 2 | 7413925 | 1.07E-05 | 0.23913 | 299 | 0.142323904 | 0.012073691 |
| SNP_A.K_1600877_9_18857982 | 9 | 18857982 | 1.07E-05 | 0.12542 | 299 | 0.142322184 | 0.012073691 |
| SNP_A.K_1000627_6_6401387 | 6 | 6401387 | 1.08E-05 | 0.28261 | 299 | 0.14228685 | 0.012134219 |

| | | | | | | | |
|---|---|---|---|---|---|---|---|
| SNP_A.K_306817_2_7355363 | 2 | 7355363 | 1.08E-05 | 0.27926 | 299 | 0.142272417 | 0.012155135 |
| SNP_A.K_993636_6_5780657 | 6 | 5780657 | 1.08E-05 | 0.29264 | 299 | 0.142265593 | 0.012158185 |
| SNP_A.K_977875_6_1606487 | 6 | 1606487 | 1.08E-05 | 0.15552 | 299 | 0.142263916 | 0.012158185 |
| SNP_A.K_1004825_6_6934278 | 6 | 6934278 | 1.09E-05 | 0.29766 | 299 | 0.142225694 | 0.012224692 |
| SNP_A.K_990247_6_4703017 | 6 | 4703017 | 1.09E-05 | 0.20736 | 299 | 0.142203836 | 0.012259202 |
| SNP_A.K_1004867_6_6937822 | 6 | 6937822 | 1.09E-05 | 0.31271 | 299 | 0.142200836 | 0.012259202 |
| SNP_A.K_1000743_6_6414136 | 6 | 6414136 | 1.09E-05 | 0.29599 | 299 | 0.14217849 | 0.012295617 |
| SNP_A.K_1001532_6_6487625 | 6 | 6487625 | 1.10E-05 | 0.30435 | 299 | 0.142153228 | 0.012337786 |
| SNP_A.K_1006016_6_7096567 | 6 | 7096567 | 1.10E-05 | 0.31104 | 299 | 0.142146868 | 0.012343444 |
| SNP_A.K_456491_2_33046845 | 2 | 33046845 | 1.10E-05 | 0.06522 | 299 | 0.142139677 | 0.012350718 |
| SNP_A.K_306839_2_7358520 | 2 | 7358520 | 1.10E-05 | 0.26589 | 299 | 0.142129592 | 0.012363612 |
| SNP_A.K_752052_4_17818945 | 4 | 17818945 | 1.10E-05 | 0.02174 | 299 | 0.142122773 | 0.012370183 |
| SNP_A.K_1719483_10_15806214 | 10 | 15806214 | 1.10E-05 | 0.03512 | 299 | 0.142115844 | 0.012376975 |
| SNP_A.K_978681_6_1760250 | 6 | 1760250 | 1.11E-05 | 0.27258 | 299 | 0.142107886 | 0.012378048 |
| SNP_A.K_1001316_6_6464817 | 6 | 6464817 | 1.11E-05 | 0.29264 | 299 | 0.14210678 | 0.012378048 |
| SNP_A.K_1004869_6_6938024 | 6 | 6938024 | 1.11E-05 | 0.33278 | 299 | 0.142102746 | 0.012378048 |
| SNP_A.K_1000989_6_6434190 | 6 | 6434190 | 1.11E-05 | 0.33445 | 299 | 0.142098845 | 0.012378048 |
| SNP_A.K_194494_1_30878531 | 1 | 30878531 | 1.11E-05 | 0.3796 | 299 | 0.142098165 | 0.012378048 |
| SNP_A.K_978315_6_1674540 | 6 | 1674540 | 1.11E-05 | 0.29264 | 299 | 0.14209324 | 0.012380972 |
| SNP_A.K_1390017_8_10145392 | 8 | 10145392 | 1.11E-05 | 0.04849 | 299 | 0.142087518 | 0.012385451 |
| SNP_A.K_326817_2_10057358 | 2 | 10057358 | 1.11E-05 | 0.06355 | 299 | 0.142082253 | 0.012389044 |
| SNP_A.K_1001750_6_6514639 | 6 | 6514639 | 1.11E-05 | 0.30435 | 299 | 0.142069324 | 0.012407558 |
| SNP_A.K_307375_2_7399723 | 2 | 7399723 | 1.11E-05 | 0.25251 | 299 | 0.142057289 | 0.012422248 |
| SNP_A.K_1365035_8_6067627 | 8 | 6067627 | 1.12E-05 | 0.12876 | 299 | 0.142053911 | 0.012422248 |
| SNP_A.K_1001132_6_6448523 | 6 | 6448523 | 1.12E-05 | 0.28763 | 299 | 0.142050581 | 0.012422248 |
| SNP_A.K_710488_4_8424729 | 4 | 8424729 | 1.12E-05 | 0.07023 | 299 | 0.142048146 | 0.012422248 |
| SNP_A.K_1608579_9_20789844 | 9 | 20789844 | 1.12E-05 | 0.22408 | 299 | 0.142035694 | 0.012439898 |
| SNP_A.K_977874_6_1606181 | 6 | 1606181 | 1.12E-05 | 0.15217 | 299 | 0.142029915 | 0.012444537 |
| SNP_A.K_709804_4_8298098 | 4 | 8298098 | 1.12E-05 | 0.07191 | 299 | 0.142019118 | 0.012452346 |
| SNP_A.K_709805_4_8298101 | 4 | 8298101 | 1.12E-05 | 0.07191 | 299 | 0.142019118 | 0.012452346 |
| SNP_A.K_709200_4_8238535 | 4 | 8238535 | 1.12E-05 | 0.08528 | 299 | 0.142011438 | 0.012453921 |
| SNP_A.K_1004348_6_6870694 | 6 | 6870694 | 1.12E-05 | 0.36288 | 299 | 0.142010562 | 0.012453921 |
| SNP_A.K_990354_6_4712180 | 6 | 4712180 | 1.12E-05 | 0.14047 | 299 | 0.142007753 | 0.012453921 |
| SNP_A.K_1004999_6_6961072 | 6 | 6961072 | 1.12E-05 | 0.32274 | 299 | 0.142004734 | 0.012453921 |
| SNP_A.K_194918_1_30999355 | 1 | 30999355 | 1.13E-05 | 0.40635 | 299 | 0.141994931 | 0.012466461 |
| SNP_A.K_999746_6_6312241 | 6 | 6312241 | 1.13E-05 | 0.16221 | 299 | 0.141967264 | 0.012510939 |
| SNP_A.K_1000814_6_6417622 | 6 | 6417622 | 1.13E-05 | 0.33278 | 299 | 0.141965465 | 0.012510939 |
| SNP_A.K_709881_4_8300221 | 4 | 8300221 | 1.14E-05 | 0.07191 | 299 | 0.141933116 | 0.012561326 |
| SNP_A.K_194269_1_30836940 | 1 | 30836940 | 1.14E-05 | 0.48997 | 299 | 0.141933091 | 0.012561326 |
| SNP_A.K_1364872_8_6050156 | 8 | 6050156 | 1.14E-05 | 0.10201 | 299 | 0.141899733 | 0.012616152 |
| SNP_A.K_1003989_6_6812561 | 6 | 6812561 | 1.14E-05 | 0.38629 | 299 | 0.141898586 | 0.012616152 |
| SNP_A.K_978161_6_1656538 | 6 | 1656538 | 1.14E-05 | 0.17559 | 299 | 0.141894683 | 0.012617194 |
| SNP_A.K_1389938_8_10136380 | 8 | 10136380 | 1.15E-05 | 0.05017 | 299 | 0.141862127 | 0.012675123 |
| SNP_A.K_1005688_6_7034049 | 6 | 7034049 | 1.15E-05 | 0.27592 | 299 | 0.141844219 | 0.012698684 |
| SNP_A.K_308458_2_7517440 | 2 | 7517440 | 1.15E-05 | 0.19565 | 299 | 0.141843553 | 0.012698684 |
| SNP_A.K_307154_2_7382405 | 2 | 7382405 | 1.16E-05 | 0.39967 | 299 | 0.141821044 | 0.012736897 |
| SNP_A.K_679467_4_1085937 | 4 | 1085937 | 1.16E-05 | 0.02508 | 299 | 0.141814205 | 0.012743842 |
| SNP_A.K_1349619_8_3724870 | 8 | 3724870 | 1.16E-05 | 0.01839 | 299 | 0.141806628 | 0.012752125 |
| SNP_A.K_978632_6_1747812 | 6 | 1747812 | 1.16E-05 | 0.26756 | 299 | 0.141803336 | 0.012752125 |
| SNP_A.K_308430_2_7513181 | 2 | 7513181 | 1.16E-05 | 0.199 | 299 | 0.141799647 | 0.01275278 |

| | | | | | | | |
|---|---|---|---|---|---|---|---|
| SNP_A.K_1000762_6_6415502 | 6 | 6415502 | 1.16E-05 | 0.34448 | 299 | 0.141793156 | 0.012759051 |
| SNP_A.K_1432449_8_16508250 | 8 | 16508250 | 1.16E-05 | 0.07692 | 299 | 0.141789464 | 0.012759719 |
| SNP_A.K_2062771_12_23886161 | 12 | 23886161 | 1.16E-05 | 0.02676 | 299 | 0.141780668 | 0.012770621 |
| SNP_A.K_978196_6_1659757 | 6 | 1659757 | 1.17E-05 | 0.17391 | 299 | 0.141771047 | 0.012781603 |
| SNP_A.K_456445_2_33037326 | 2 | 33037326 | 1.17E-05 | 0.11873 | 299 | 0.141768485 | 0.012781603 |
| SNP_A.K_978599_6_1737975 | 6 | 1737975 | 1.17E-05 | 0.25585 | 299 | 0.141764554 | 0.012782766 |
| SNP_A.K_995648_6_5946264 | 6 | 5946264 | 1.17E-05 | 0.38796 | 299 | 0.141756754 | 0.012791701 |
| SNP_A.K_325940_2_9962957 | 2 | 9962957 | 1.17E-05 | 0.04849 | 299 | 0.14174879 | 0.012800192 |
| SNP_A.K_990339_6_4711556 | 6 | 4711556 | 1.17E-05 | 0.20067 | 299 | 0.141744031 | 0.012800192 |
| SNP_A.K_994786_6_5888958 | 6 | 5888958 | 1.17E-05 | 0.28094 | 299 | 0.14174249 | 0.012800192 |
| SNP_A.K_978146_6_1654064 | 6 | 1654064 | 1.17E-05 | 0.17391 | 299 | 0.141737769 | 0.012802964 |
| SNP_A.K_1001713_6_6508525 | 6 | 6508525 | 1.17E-05 | 0.29599 | 299 | 0.141733595 | 0.012804299 |
| SNP_A.K_978139_6_1651588 | 6 | 1651588 | 1.17E-05 | 0.25585 | 299 | 0.141730425 | 0.012804299 |
| SNP_A.K_312090_2_7906329 | 2 | 7906329 | 1.18E-05 | 0.28094 | 299 | 0.141720432 | 0.012817688 |
| SNP_A.K_1000623_6_6401217 | 6 | 6401217 | 1.18E-05 | 0.28428 | 299 | 0.141691166 | 0.012869985 |
| SNP_A.K_990372_6_4712627 | 6 | 4712627 | 1.19E-05 | 0.20569 | 299 | 0.141667383 | 0.012905464 |
| SNP_A.K_978298_6_1671382 | 6 | 1671382 | 1.19E-05 | 0.24916 | 299 | 0.141666971 | 0.012905464 |
| SNP_A.K_194450_1_30873906 | 1 | 30873906 | 1.19E-05 | 0.37291 | 299 | 0.141647864 | 0.012937478 |
| SNP_A.K_1001224_6_6455819 | 6 | 6455819 | 1.19E-05 | 0.28595 | 299 | 0.141640922 | 0.012944823 |
| SNP_A.K_1001598_6_6497124 | 6 | 6497124 | 1.19E-05 | 0.28595 | 299 | 0.141635465 | 0.012949155 |
| SNP_A.K_29305_1_4222839 | 1 | 4222839 | 1.19E-05 | 0.10368 | 299 | 0.141616019 | 0.012981981 |
| SNP_A.K_995647_6_5946239 | 6 | 5946239 | 1.20E-05 | 0.45485 | 299 | 0.141604926 | 0.012989015 |
| SNP_A.K_456685_2_33105070 | 2 | 33105070 | 1.20E-05 | 0.01839 | 299 | 0.141602609 | 0.012989015 |
| SNP_A.K_456579_2_33074293 | 2 | 33074293 | 1.20E-05 | 0.01839 | 299 | 0.141602609 | 0.012989015 |
| SNP_A.K_1364924_8_6057881 | 8 | 6057881 | 1.20E-05 | 0.07525 | 299 | 0.141571473 | 0.013045888 |
| SNP_A.K_1000812_6_6417528 | 6 | 6417528 | 1.21E-05 | 0.3311 | 299 | 0.14154773 | 0.013087812 |
| SNP_A.K_1608485_9_20762901 | 9 | 20762901 | 1.21E-05 | 0.28261 | 299 | 0.14153687 | 0.013103335 |
| SNP_A.K_990337_6_4711363 | 6 | 4711363 | 1.21E-05 | 0.20401 | 299 | 0.141531685 | 0.013105101 |
| SNP_A.K_709528_4_8271656 | 4 | 8271656 | 1.21E-05 | 0.08194 | 299 | 0.14152939 | 0.013105101 |
| SNP_A.K_710480_4_8424227 | 4 | 8424227 | 1.21E-05 | 0.07023 | 299 | 0.141522854 | 0.013111747 |
| SNP_A.K_990358_6_4712319 | 6 | 4712319 | 1.21E-05 | 0.20903 | 299 | 0.141517264 | 0.013116452 |
| SNP_A.K_307076_2_7377974 | 2 | 7377974 | 1.22E-05 | 0.2592 | 299 | 0.141502426 | 0.013135127 |
| SNP_A.K_994898_6_5900200 | 6 | 5900200 | 1.22E-05 | 0.07023 | 299 | 0.141501599 | 0.013135127 |
| SNP_A.K_1781564_11_3218614 | 11 | 3218614 | 1.22E-05 | 0.0301 | 299 | 0.141469503 | 0.013194673 |
| SNP_A.K_978592_6_1736235 | 6 | 1736235 | 1.23E-05 | 0.26254 | 299 | 0.141446401 | 0.013234883 |
| SNP_A.K_990457_6_4718859 | 6 | 4718859 | 1.23E-05 | 0.19064 | 299 | 0.141443537 | 0.013234883 |
| SNP_A.K_1001315_6_6464725 | 6 | 6464725 | 1.23E-05 | 0.2893 | 299 | 0.141425583 | 0.013265394 |
| SNP_A.K_1000696_6_6412405 | 6 | 6412405 | 1.23E-05 | 0.32274 | 299 | 0.141416412 | 0.013277649 |
| SNP_A.K_1389980_8_10141596 | 8 | 10141596 | 1.24E-05 | 0.05184 | 299 | 0.141404433 | 0.01329578 |
| SNP_A.K_978633_6_1747877 | 6 | 1747877 | 1.24E-05 | 0.28261 | 299 | 0.1413948 | 0.013304916 |
| SNP_A.K_1000981_6_6432167 | 6 | 6432167 | 1.24E-05 | 0.33278 | 299 | 0.141393482 | 0.013304916 |
| SNP_A.K_1389968_8_10139482 | 8 | 10139482 | 1.24E-05 | 0.04682 | 299 | 0.141386695 | 0.013312234 |
| SNP_A.K_1001305_6_6463410 | 6 | 6463410 | 1.25E-05 | 0.29599 | 299 | 0.141354333 | 0.013373188 |
| SNP_A.K_678572_4_990276 | 4 | 990276 | 1.25E-05 | 0.06856 | 299 | 0.141341585 | 0.013391608 |
| SNP_A.K_978665_6_1751903 | 6 | 1751903 | 1.25E-05 | 0.30936 | 299 | 0.1413372 | 0.013391608 |
| SNP_A.K_978104_6_1647239 | 6 | 1647239 | 1.25E-05 | 0.25418 | 299 | 0.141335728 | 0.013391608 |
| SNP_A.K_1004642_6_6913326 | 6 | 6913326 | 1.26E-05 | 0.29933 | 299 | 0.141294837 | 0.013470987 |
| SNP_A.K_307054_2_7376254 | 2 | 7376254 | 1.26E-05 | 0.23244 | 299 | 0.141291178 | 0.013471798 |
| SNP_A.K_993837_6_5796336 | 6 | 5796336 | 1.26E-05 | 0.15886 | 299 | 0.14127519 | 0.013498738 |
| SNP_A.K_678146_4_964582 | 4 | 964582 | 1.26E-05 | 0.16054 | 299 | 0.141266065 | 0.013511157 |

| | | | | | | | |
|---|---|---|---|---|---|---|---|
| SNP_A.K_709791_4_8297819 | 4 | 8297819 | 1.26E-05 | 0.07191 | 299 | 0.14125945 | 0.013518259 |
| SNP_A.K_978603_6_1738798 | 6 | 1738798 | 1.26E-05 | 0.2592 | 299 | 0.141254163 | 0.013522547 |
| SNP_A.K_1005024_6_6962876 | 6 | 6962876 | 1.27E-05 | 0.32274 | 299 | 0.141249051 | 0.013526467 |
| SNP_A.K_709871_4_8300058 | 4 | 8300058 | 1.27E-05 | 0.07358 | 299 | 0.141232403 | 0.013554941 |
| SNP_A.K_1382750_8_9115622 | 8 | 9115622 | 1.27E-05 | 0.28261 | 299 | 0.141219674 | 0.013575118 |
| SNP_A.K_710116_4_8316706 | 4 | 8316706 | 1.28E-05 | 0.08194 | 299 | 0.141201031 | 0.013607971 |
| SNP_A.K_994345_6_5849451 | 6 | 5849451 | 1.28E-05 | 0.2291 | 299 | 0.141194072 | 0.013615884 |
| SNP_A.K_710119_4_8317289 | 4 | 8317289 | 1.28E-05 | 0.09866 | 299 | 0.141180813 | 0.0136373 |
| SNP_A.K_1608628_9_20803584 | 9 | 20803584 | 1.28E-05 | 0.28763 | 299 | 0.141169055 | 0.013655532 |
| SNP_A.K_1600932_9_18862404 | 9 | 18862404 | 1.28E-05 | 0.1204 | 299 | 0.141161142 | 0.013660293 |
| SNP_A.K_990335_6_4711227 | 6 | 4711227 | 1.28E-05 | 0.21237 | 299 | 0.141159261 | 0.013660293 |
| SNP_A.K_990350_6_4711965 | 6 | 4711965 | 1.29E-05 | 0.20569 | 299 | 0.141153298 | 0.013660293 |
| SNP_A.K_993073_6_5704964 | 6 | 5704964 | 1.29E-05 | 0.15385 | 299 | 0.141149284 | 0.013660293 |
| SNP_A.K_978896_6_1798493 | 6 | 1798493 | 1.29E-05 | 0.33779 | 299 | 0.141147967 | 0.013660293 |
| SNP_A.K_1001766_6_6519580 | 6 | 6519580 | 1.29E-05 | 0.28595 | 299 | 0.141146266 | 0.013660293 |
| SNP_A.K_1608596_9_20792124 | 9 | 20792124 | 1.29E-05 | 0.24247 | 299 | 0.141144084 | 0.013660293 |
| SNP_A.K_1001237_6_6456904 | 6 | 6456904 | 1.29E-05 | 0.29431 | 299 | 0.141134045 | 0.013674892 |
| SNP_A.K_306775_2_7349677 | 2 | 7349677 | 1.29E-05 | 0.2408 | 299 | 0.141107255 | 0.013721569 |
| SNP_A.K_978634_6_1748034 | 6 | 1748034 | 1.29E-05 | 0.28763 | 299 | 0.141105883 | 0.013721569 |
| SNP_A.K_1608493_9_20763734 | 9 | 20763734 | 1.30E-05 | 0.29431 | 299 | 0.141095995 | 0.013735918 |
| SNP_A.K_307492_2_7412882 | 2 | 7412882 | 1.30E-05 | 0.23579 | 299 | 0.14105705 | 0.013813236 |
| SNP_A.K_990340_6_4711574 | 6 | 4711574 | 1.31E-05 | 0.20067 | 299 | 0.141041255 | 0.013836334 |
| SNP_A.K_1390240_8_10166668 | 8 | 10166668 | 1.31E-05 | 0.04849 | 299 | 0.141039774 | 0.013836334 |
| SNP_A.K_990329_6_4710809 | 6 | 4710809 | 1.31E-05 | 0.20903 | 299 | 0.141036722 | 0.013836334 |
| SNP_A.K_456494_2_33047934 | 2 | 33047934 | 1.31E-05 | 0.06689 | 299 | 0.141023689 | 0.013857672 |
| SNP_A.K_978026_6_1640830 | 6 | 1640830 | 1.32E-05 | 0.28428 | 299 | 0.141000145 | 0.013902003 |
| SNP_A.K_306971_2_7371153 | 2 | 7371153 | 1.32E-05 | 0.25585 | 299 | 0.140992618 | 0.013911405 |
| SNP_A.K_996231_6_5976246 | 6 | 5976246 | 1.32E-05 | 0.37291 | 299 | 0.140972382 | 0.013948661 |
| SNP_A.K_1005058_6_6966888 | 6 | 6966888 | 1.33E-05 | 0.31271 | 299 | 0.140950914 | 0.01398873 |
| SNP_A.K_978679_6_1758924 | 6 | 1758924 | 1.33E-05 | 0.27592 | 299 | 0.140945351 | 0.01399388 |
| SNP_A.K_1794676_11_5474936 | 11 | 5474936 | 1.33E-05 | 0.44147 | 299 | 0.140932248 | 0.014015644 |
| SNP_A.K_1608598_9_20792178 | 9 | 20792178 | 1.33E-05 | 0.23411 | 299 | 0.140925555 | 0.014023304 |
| SNP_A.K_1600805_9_18842812 | 9 | 18842812 | 1.33E-05 | 0.12542 | 299 | 0.140913923 | 0.014041872 |
| SNP_A.K_257550_1_41599614 | 1 | 41599614 | 1.34E-05 | 0.02843 | 299 | 0.140898696 | 0.014061401 |
| SNP_A.K_710114_4_8316691 | 4 | 8316691 | 1.34E-05 | 0.0786 | 299 | 0.140898657 | 0.014061401 |
| SNP_A.K_1005473_6_7015713 | 6 | 7015713 | 1.34E-05 | 0.35953 | 299 | 0.140880704 | 0.014094034 |
| SNP_A.K_1004146_6_6838409 | 6 | 6838409 | 1.34E-05 | 0.31271 | 299 | 0.140875735 | 0.014097932 |
| SNP_A.K_709754_4_8284876 | 4 | 8284876 | 1.35E-05 | 0.08361 | 299 | 0.140854373 | 0.014138236 |
| SNP_A.K_977963_6_1625935 | 6 | 1625935 | 1.35E-05 | 0.17057 | 299 | 0.140849848 | 0.014141168 |
| SNP_A.K_1004357_6_6875266 | 6 | 6875266 | 1.35E-05 | 0.3194 | 299 | 0.140833672 | 0.014170042 |
| SNP_A.K_977912_6_1621176 | 6 | 1621176 | 1.36E-05 | 0.16388 | 299 | 0.140804211 | 0.014224532 |
| SNP_A.K_1433063_8_16615989 | 8 | 16615989 | 1.36E-05 | 0.07525 | 299 | 0.14079982 | 0.014224532 |
| SNP_A.K_308426_2_7512251 | 2 | 7512251 | 1.36E-05 | 0.22241 | 299 | 0.140795107 | 0.014224532 |
| SNP_A.K_990239_6_4702703 | 6 | 4702703 | 1.36E-05 | 0.20903 | 299 | 0.140790689 | 0.014224532 |
| SNP_A.K_709825_4_8298559 | 4 | 8298559 | 1.36E-05 | 0.07023 | 299 | 0.140790483 | 0.014224532 |
| SNP_A.K_990328_6_4710774 | 6 | 4710774 | 1.36E-05 | 0.20569 | 299 | 0.140790075 | 0.014224532 |
| SNP_A.K_2068303_12_24973861 | 12 | 24973861 | 1.36E-05 | 0.02174 | 299 | 0.140785143 | 0.014228424 |
| SNP_A.K_1608514_9_20770019 | 9 | 20770019 | 1.36E-05 | 0.29431 | 299 | 0.14077144 | 0.01424949 |
| SNP_A.K_709880_4_8300209 | 4 | 8300209 | 1.37E-05 | 0.07191 | 299 | 0.140769355 | 0.01424949 |
| SNP_A.K_907940_5_17467198 | 5 | 17467198 | 1.37E-05 | 0.02174 | 299 | 0.140760245 | 0.014262768 |

| | | | | | | | |
|---|---|---|---|---|---|---|---|
| SNP_A.K_1004356_6_6875233 | 6 | 6875233 | 1.37E-05 | 0.31438 | 299 | 0.140747966 | 0.014283179 |
| SNP_A.K_306890_2_7364030 | 2 | 7364030 | 1.37E-05 | 0.2709 | 299 | 0.140733949 | 0.014302191 |
| SNP_A.K_978138_6_1651318 | 6 | 1651318 | 1.37E-05 | 0.17224 | 299 | 0.140732697 | 0.014302191 |
| SNP_A.K_992970_6_5692884 | 6 | 5692884 | 1.37E-05 | 0.23077 | 299 | 0.140728012 | 0.014302191 |
| SNP_A.K_1830772_11_10580303 | 11 | 10580303 | 1.37E-05 | 0.21906 | 299 | 0.140725299 | 0.014302191 |
| SNP_A.K_307533_2_7416065 | 2 | 7416065 | 1.37E-05 | 0.25753 | 299 | 0.140723472 | 0.014302191 |
| SNP_A.K_1000986_6_6433922 | 6 | 6433922 | 1.38E-05 | 0.32441 | 299 | 0.140720425 | 0.014302191 |
| SNP_A.K_1000975_6_6430830 | 6 | 6430830 | 1.39E-05 | 0.33445 | 299 | 0.140671939 | 0.014404488 |
| SNP_A.K_307336_2_7396717 | 2 | 7396717 | 1.39E-05 | 0.23244 | 299 | 0.140667264 | 0.014407368 |
| SNP_A.K_996278_6_5981197 | 6 | 5981197 | 1.39E-05 | 0.28094 | 299 | 0.140664324 | 0.014407368 |
| SNP_A.K_2079158_12_27061420 | 12 | 27061420 | 1.39E-05 | 0.01505 | 299 | 0.140651251 | 0.014429828 |
| SNP_A.K_1608614_9_20798975 | 9 | 20798975 | 1.40E-05 | 0.23579 | 299 | 0.140600152 | 0.014539041 |
| SNP_A.K_978385_6_1684894 | 6 | 1684894 | 1.40E-05 | 0.25585 | 299 | 0.140586901 | 0.014562126 |
| SNP_A.K_709795_4_8297870 | 4 | 8297870 | 1.41E-05 | 0.07023 | 299 | 0.14056676 | 0.014601069 |
| SNP_A.K_1608630_9_20803874 | 9 | 20803874 | 1.41E-05 | 0.24247 | 299 | 0.14055807 | 0.014609775 |
| SNP_A.K_709572_4_8274312 | 4 | 8274312 | 1.41E-05 | 0.09866 | 299 | 0.140556647 | 0.014609775 |
| SNP_A.K_1005049_6_6966049 | 6 | 6966049 | 1.41E-05 | 0.35117 | 299 | 0.140547813 | 0.014622822 |
| SNP_A.K_1005037_6_6964073 | 6 | 6964073 | 1.42E-05 | 0.29097 | 299 | 0.140539112 | 0.014631304 |
| SNP_A.K_978863_6_1789622 | 6 | 1789622 | 1.42E-05 | 0.31438 | 299 | 0.140537813 | 0.014631304 |
| SNP_A.K_307027_2_7373443 | 2 | 7373443 | 1.42E-05 | 0.25753 | 299 | 0.140520845 | 0.014654701 |
| SNP_A.K_1001443_6_6481781 | 6 | 6481781 | 1.42E-05 | 0.30936 | 299 | 0.1405196 | 0.014654701 |
| SNP_A.K_306713_2_7343894 | 2 | 7343894 | 1.42E-05 | 0.23077 | 299 | 0.140515353 | 0.014654701 |
| SNP_A.K_1004415_6_6880450 | 6 | 6880450 | 1.42E-05 | 0.38294 | 299 | 0.140515047 | 0.014654701 |
| SNP_A.K_990359_6_4712337 | 6 | 4712337 | 1.43E-05 | 0.20569 | 299 | 0.140462045 | 0.014770108 |
| SNP_A.K_1001834_6_6523614 | 6 | 6523614 | 1.44E-05 | 0.31438 | 299 | 0.140450198 | 0.014790342 |
| SNP_A.K_995725_6_5948957 | 6 | 5948957 | 1.44E-05 | 0.49666 | 299 | 0.140442704 | 0.014800468 |
| SNP_A.K_978212_6_1661829 | 6 | 1661829 | 1.44E-05 | 0.17726 | 299 | 0.140438321 | 0.014803354 |
| SNP_A.K_1365031_8_6067247 | 8 | 6067247 | 1.44E-05 | 0.11037 | 299 | 0.140433663 | 0.014806886 |
| SNP_A.K_978898_6_1798756 | 6 | 1798756 | 1.44E-05 | 0.29599 | 299 | 0.14042219 | 0.014826314 |
| SNP_A.K_710033_4_8310218 | 4 | 8310218 | 1.44E-05 | 0.07692 | 299 | 0.140407668 | 0.014852898 |
| SNP_A.K_990384_6_4712968 | 6 | 4712968 | 1.45E-05 | 0.19565 | 299 | 0.140390681 | 0.014885305 |
| SNP_A.K_1577941_9_13755181 | 9 | 13755181 | 1.45E-05 | 0.05017 | 299 | 0.140375891 | 0.014908738 |
| SNP_A.K_1000982_6_6432282 | 6 | 6432282 | 1.45E-05 | 0.33612 | 299 | 0.140374416 | 0.014908738 |
| SNP_A.K_1001480_6_6483261 | 6 | 6483261 | 1.45E-05 | 0.29431 | 299 | 0.140365239 | 0.014922929 |
| SNP_A.K_978399_6_1686955 | 6 | 1686955 | 1.46E-05 | 0.25418 | 299 | 0.140345302 | 0.014958488 |
| SNP_A.K_1001283_6_6461389 | 6 | 6461389 | 1.46E-05 | 0.37291 | 299 | 0.140343857 | 0.014958488 |
| SNP_A.K_1004367_6_6876933 | 6 | 6876933 | 1.46E-05 | 0.36288 | 299 | 0.140320908 | 0.015003818 |
| SNP_A.K_1004806_6_6933539 | 6 | 6933539 | 1.47E-05 | 0.30769 | 299 | 0.140318378 | 0.015003818 |
| SNP_A.K_995303_6_5924486 | 6 | 5924486 | 1.47E-05 | 0.1388 | 299 | 0.140309831 | 0.01501663 |
| SNP_A.K_28907_1_4179493 | 1 | 4179493 | 1.47E-05 | 0.12876 | 299 | 0.140304777 | 0.015021194 |
| SNP_A.K_993622_6_5780207 | 6 | 5780207 | 1.47E-05 | 0.29264 | 299 | 0.140299041 | 0.015022214 |
| SNP_A.K_1001243_6_6457259 | 6 | 6457259 | 1.47E-05 | 0.37124 | 299 | 0.140298105 | 0.015022214 |
| SNP_A.K_678658_4_993423 | 4 | 993423 | 1.47E-05 | 0.06856 | 299 | 0.140292066 | 0.015029122 |
| SNP_A.K_978123_6_1647999 | 6 | 1647999 | 1.47E-05 | 0.27759 | 299 | 0.14028198 | 0.015045617 |
| SNP_A.K_996334_6_5988201 | 6 | 5988201 | 1.48E-05 | 0.21906 | 299 | 0.140269054 | 0.01506887 |
| SNP_A.K_307514_2_7414867 | 2 | 7414867 | 1.48E-05 | 0.43144 | 299 | 0.140261157 | 0.015072848 |
| SNP_A.K_709753_4_8284869 | 4 | 8284869 | 1.48E-05 | 0.08361 | 299 | 0.140261152 | 0.015072848 |
| SNP_A.K_990346_6_4711807 | 6 | 4711807 | 1.48E-05 | 0.14381 | 299 | 0.140254619 | 0.01508097 |
| SNP_A.K_194660_1_30907310 | 1 | 30907310 | 1.48E-05 | 0.45485 | 299 | 0.140244828 | 0.015096841 |
| SNP_A.K_721670_4_11034116 | 4 | 11034116 | 1.49E-05 | 0.06187 | 299 | 0.140231942 | 0.015120099 |

| | | | | | | | |
|---|---|---|---|---|---|---|---|
| SNP_A.K_1600928_9_18862234 | 9 | 18862234 | 1.49E-05 | 0.14548 | 299 | 0.140220866 | 0.015139081 |
| SNP_A.K_1001270_6_6459969 | 6 | 6459969 | 1.49E-05 | 0.3913 | 299 | 0.140209209 | 0.01515948 |
| SNP_A.K_1004503_6_6894925 | 6 | 6894925 | 1.49E-05 | 0.32274 | 299 | 0.140197739 | 0.015168038 |
| SNP_A.K_1001247_6_6458325 | 6 | 6458325 | 1.49E-05 | 0.37458 | 299 | 0.140196984 | 0.015168038 |
| SNP_A.K_993112_6_5708683 | 6 | 5708683 | 1.49E-05 | 0.15217 | 299 | 0.140196321 | 0.015168038 |
| SNP_A.K_977995_6_1633260 | 6 | 1633260 | 1.49E-05 | 0.25084 | 299 | 0.140192474 | 0.015169827 |
| SNP_A.K_1001241_6_6457118 | 6 | 6457118 | 1.50E-05 | 0.29933 | 299 | 0.140182285 | 0.015179429 |
| SNP_A.K_1432429_8_16500875 | 8 | 16500875 | 1.50E-05 | 0.08361 | 299 | 0.140179756 | 0.015179429 |
| SNP_A.K_1600799_9_18841702 | 9 | 18841702 | 1.50E-05 | 0.13712 | 299 | 0.14017917 | 0.015179429 |
| SNP_A.K_709887_4_8300318 | 4 | 8300318 | 1.50E-05 | 0.07358 | 299 | 0.140171067 | 0.015186588 |
| SNP_A.K_709763_4_8285277 | 4 | 8285277 | 1.50E-05 | 0.08696 | 299 | 0.140169993 | 0.015186588 |
| SNP_A.K_978038_6_1642041 | 6 | 1642041 | 1.50E-05 | 0.20067 | 299 | 0.140156913 | 0.015210495 |
| SNP_A.K_978216_6_1662090 | 6 | 1662090 | 1.51E-05 | 0.1806 | 299 | 0.140113886 | 0.015306444 |
| SNP_A.K_998481_6_6203672 | 6 | 6203672 | 1.51E-05 | 0.15552 | 299 | 0.140109851 | 0.015307177 |
| SNP_A.K_1000932_6_6425020 | 6 | 6425020 | 1.51E-05 | 0.29097 | 299 | 0.140107412 | 0.015307177 |
| SNP_A.K_1004382_6_6878100 | 6 | 6878100 | 1.52E-05 | 0.4097 | 299 | 0.140102678 | 0.01531116 |
| SNP_A.K_1005655_6_7033271 | 6 | 7033271 | 1.52E-05 | 0.27258 | 299 | 0.140091415 | 0.015330897 |
| SNP_A.K_978346_6_1679943 | 6 | 1679943 | 1.52E-05 | 0.25753 | 299 | 0.140078576 | 0.015354475 |
| SNP_A.K_710115_4_8316699 | 4 | 8316699 | 1.52E-05 | 0.08027 | 299 | 0.140073422 | 0.015359496 |
| SNP_A.K_978389_6_1685030 | 6 | 1685030 | 1.53E-05 | 0.25753 | 299 | 0.140063014 | 0.015377238 |
| SNP_A.K_307167_2_7383081 | 2 | 7383081 | 1.53E-05 | 0.25084 | 299 | 0.140054858 | 0.015387044 |
| SNP_A.K_978669_6_1753579 | 6 | 1753579 | 1.53E-05 | 0.27258 | 299 | 0.140052818 | 0.015387044 |
| SNP_A.K_978015_6_1636847 | 6 | 1636847 | 1.53E-05 | 0.25251 | 299 | 0.140047358 | 0.015392831 |
| SNP_A.K_995226_6_5920765 | 6 | 5920765 | 1.54E-05 | 0.12375 | 299 | 0.140012293 | 0.015460553 |
| SNP_A.K_995683_6_5947774 | 6 | 5947774 | 1.54E-05 | 0.301 | 299 | 0.140012243 | 0.015460553 |
| SNP_A.K_978395_6_1686505 | 6 | 1686505 | 1.54E-05 | 0.25585 | 299 | 0.14000873 | 0.015460553 |
| SNP_A.K_978206_6_1660357 | 6 | 1660357 | 1.54E-05 | 0.19064 | 299 | 0.140006602 | 0.015460553 |
| SNP_A.K_709837_4_8299264 | 4 | 8299264 | 1.54E-05 | 0.07191 | 299 | 0.140004145 | 0.015460553 |
| SNP_A.K_978706_6_1769202 | 6 | 1769202 | 1.55E-05 | 0.35117 | 299 | 0.139970065 | 0.015536258 |
| SNP_A.K_1004956_6_6957433 | 6 | 6957433 | 1.55E-05 | 0.35786 | 299 | 0.139946342 | 0.015575207 |
| SNP_A.K_307385_2_7400113 | 2 | 7400113 | 1.55E-05 | 0.25753 | 299 | 0.139945071 | 0.015575207 |
| SNP_A.K_8635_1_1627181 | 1 | 1627181 | 1.55E-05 | 0.01505 | 299 | 0.139944989 | 0.015575207 |
| SNP_A.K_307153_2_7382225 | 2 | 7382225 | 1.56E-05 | 0.23913 | 299 | 0.139931414 | 0.015594239 |
| SNP_A.K_1925869_11_24770827 | 11 | 24770827 | 1.56E-05 | 0.02174 | 299 | 0.139926373 | 0.015594239 |
| SNP_A.K_307048_2_7375415 | 2 | 7375415 | 1.56E-05 | 0.23579 | 299 | 0.139925755 | 0.015594239 |
| SNP_A.K_456446_2_33037529 | 2 | 33037529 | 1.56E-05 | 0.06689 | 299 | 0.139925019 | 0.015594239 |
| SNP_A.K_1001337_6_6466195 | 6 | 6466195 | 1.56E-05 | 0.2893 | 299 | 0.1399202 | 0.015598585 |
| SNP_A.K_1600819_9_18843940 | 9 | 18843940 | 1.56E-05 | 0.12876 | 299 | 0.139914106 | 0.015602184 |
| SNP_A.K_307509_2_7413958 | 2 | 7413958 | 1.56E-05 | 0.43144 | 299 | 0.139912639 | 0.015602184 |
| SNP_A.K_993378_6_5743531 | 6 | 5743531 | 1.56E-05 | 0.09866 | 299 | 0.139901376 | 0.015615375 |
| SNP_A.K_1004085_6_6832717 | 6 | 6832717 | 1.56E-05 | 0.16722 | 299 | 0.139901183 | 0.015615375 |
| SNP_A.K_710385_4_8339488 | 4 | 8339488 | 1.57E-05 | 0.08361 | 299 | 0.139895904 | 0.015620876 |
| SNP_A.K_1003645_6_6752756 | 6 | 6752756 | 1.57E-05 | 0.01338 | 299 | 0.139891061 | 0.015625309 |
| SNP_A.K_978691_6_1764516 | 6 | 1764516 | 1.57E-05 | 0.27592 | 299 | 0.1398874 | 0.015626837 |
| SNP_A.K_1001242_6_6457170 | 6 | 6457170 | 1.57E-05 | 0.29766 | 299 | 0.13988077 | 0.01563568 |
| SNP_A.K_1005322_6_7005978 | 6 | 7005978 | 1.57E-05 | 0.35117 | 299 | 0.139863431 | 0.015670943 |
| SNP_A.K_1001471_6_6482862 | 6 | 6482862 | 1.58E-05 | 0.29264 | 299 | 0.139849505 | 0.015693366 |
| SNP_A.K_1600929_9_18862283 | 9 | 18862283 | 1.58E-05 | 0.14214 | 299 | 0.139848285 | 0.015693366 |
| SNP_A.K_1004225_6_6851835 | 6 | 6851835 | 1.58E-05 | 0.33278 | 299 | 0.139841291 | 0.015703163 |
| SNP_A.K_710475_4_8423968 | 4 | 8423968 | 1.58E-05 | 0.07191 | 299 | 0.139831347 | 0.01571277 |

| | | | | | | | |
|---|---|---|---|---|---|---|---|
| SNP_A.K_710025_4_8309576 | 4 | 8309576 | 1.58E-05 | 0.07191 | 299 | 0.139831347 | 0.01571277 |
| SNP_A.K_978945_6_1816874 | 6 | 1816874 | 1.58E-05 | 0.30435 | 299 | 0.139824084 | 0.015723258 |
| SNP_A.K_992480_6_5652278 | 6 | 5652278 | 1.59E-05 | 0.10201 | 299 | 0.139807449 | 0.015756999 |
| SNP_A.K_1382670_8_9112847 | 8 | 9112847 | 1.59E-05 | 0.28763 | 299 | 0.139800042 | 0.015767881 |
| SNP_A.K_995818_6_5951679 | 6 | 5951679 | 1.60E-05 | 0.301 | 299 | 0.139749097 | 0.015887371 |
| SNP_A.K_978147_6_1654365 | 6 | 1654365 | 1.60E-05 | 0.17559 | 299 | 0.139745511 | 0.015888783 |
| SNP_A.K_1382762_8_9115992 | 8 | 9115992 | 1.61E-05 | 0.27926 | 299 | 0.13973824 | 0.015899428 |
| SNP_A.K_1005546_6_7022823 | 6 | 7022823 | 1.61E-05 | 0.26087 | 299 | 0.139735013 | 0.01589995 |
| SNP_A.K_978877_6_1793978 | 6 | 1793978 | 1.61E-05 | 0.30936 | 299 | 0.139728365 | 0.015909047 |
| SNP_A.K_1001236_6_6456861 | 6 | 6456861 | 1.61E-05 | 0.29431 | 299 | 0.139721225 | 0.015919386 |
| SNP_A.K_306883_2_7363429 | 2 | 7363429 | 1.61E-05 | 0.27258 | 299 | 0.139707907 | 0.015938945 |
| SNP_A.K_374744_2_17878213 | 2 | 17878213 | 1.61E-05 | 0.05017 | 299 | 0.139707405 | 0.015938945 |
| SNP_A.K_672910_4_298014 | 4 | 298014 | 1.62E-05 | 0.10201 | 299 | 0.139698404 | 0.015953992 |
| SNP_A.K_1005007_6_6961747 | 6 | 6961747 | 1.62E-05 | 0.32609 | 299 | 0.139693292 | 0.015959277 |
| SNP_A.K_990275_6_4704180 | 6 | 4704180 | 1.62E-05 | 0.2107 | 299 | 0.139681921 | 0.015973743 |
| SNP_A.K_978107_6_1647427 | 6 | 1647427 | 1.62E-05 | 0.2592 | 299 | 0.139681526 | 0.015973743 |
| SNP_A.K_1608521_9_20772075 | 9 | 20772075 | 1.62E-05 | 0.28261 | 299 | 0.13966442 | 0.016009274 |
| SNP_A.K_1000582_6_6397565 | 6 | 6397565 | 1.63E-05 | 0.30435 | 299 | 0.139649928 | 0.01603319 |
| SNP_A.K_1004807_6_6933542 | 6 | 6933542 | 1.63E-05 | 0.30602 | 299 | 0.139648939 | 0.01603319 |
| SNP_A.K_1005302_6_7003399 | 6 | 7003399 | 1.63E-05 | 0.25418 | 299 | 0.13964541 | 0.016034524 |
| SNP_A.K_1001359_6_6466728 | 6 | 6466728 | 1.63E-05 | 0.28763 | 299 | 0.139632986 | 0.016058352 |
| SNP_A.K_580861_3_21143949 | 3 | 21143949 | 1.64E-05 | 0.40468 | 299 | 0.139600571 | 0.016132966 |
| SNP_A.K_978217_6_1662107 | 6 | 1662107 | 1.64E-05 | 0.1806 | 299 | 0.139584612 | 0.016161285 |
| SNP_A.K_1830624_11_10571212 | 11 | 10571212 | 1.65E-05 | 0.35117 | 299 | 0.13958345 | 0.016161285 |
| SNP_A.K_307527_2_7415701 | 2 | 7415701 | 1.65E-05 | 0.23579 | 299 | 0.139575045 | 0.016175072 |
| SNP_A.K_1001208_6_6455014 | 6 | 6455014 | 1.65E-05 | 0.37458 | 299 | 0.139559141 | 0.016204429 |
| SNP_A.K_1019445_6_9094776 | 6 | 9094776 | 1.65E-05 | 0.03512 | 299 | 0.139557556 | 0.016204429 |
| SNP_A.K_195113_1_31046295 | 1 | 31046295 | 1.65E-05 | 0.40803 | 299 | 0.139550743 | 0.016214196 |
| SNP_A.K_1607052_9_20264813 | 9 | 20264813 | 1.66E-05 | 0.24916 | 299 | 0.139543637 | 0.016224723 |
| SNP_A.K_978246_6_1664631 | 6 | 1664631 | 1.66E-05 | 0.1806 | 299 | 0.139538406 | 0.016226824 |
| SNP_A.K_737118_4_13970219 | 4 | 13970219 | 1.66E-05 | 0.02508 | 299 | 0.139535418 | 0.016226824 |
| SNP_A.K_1433094_8_16617450 | 8 | 16617450 | 1.66E-05 | 0.08361 | 299 | 0.139533859 | 0.016226824 |
| SNP_A.K_1000926_6_6424491 | 6 | 6424491 | 1.66E-05 | 0.32776 | 299 | 0.139530336 | 0.016228208 |
| SNP_A.K_1005486_6_7016696 | 6 | 7016696 | 1.66E-05 | 0.12375 | 299 | 0.139517382 | 0.016253736 |
| SNP_A.K_1183795_7_4603176 | 7 | 4603176 | 1.66E-05 | 0.06689 | 299 | 0.13951247 | 0.016258688 |
| SNP_A.K_1608475_9_20762236 | 9 | 20762236 | 1.67E-05 | 0.2893 | 299 | 0.139498955 | 0.016285712 |
| SNP_A.K_678442_4_986649 | 4 | 986649 | 1.67E-05 | 0.06187 | 299 | 0.139473297 | 0.016337067 |
| SNP_A.K_1001734_6_6511771 | 6 | 6511771 | 1.67E-05 | 0.30769 | 299 | 0.139473028 | 0.016337067 |
| SNP_A.K_990333_6_4711187 | 6 | 4711187 | 1.68E-05 | 0.20903 | 299 | 0.139461553 | 0.016358974 |
| SNP_A.K_1001338_6_6466197 | 6 | 6466197 | 1.68E-05 | 0.2893 | 299 | 0.139446204 | 0.01639092 |
| SNP_A.K_1830680_11_10574148 | 11 | 10574148 | 1.68E-05 | 0.39632 | 299 | 0.139438861 | 0.016402221 |
| SNP_A.K_307328_2_7395346 | 2 | 7395346 | 1.69E-05 | 0.25585 | 299 | 0.13941029 | 0.016446186 |
| SNP_A.K_483254_3_3232920 | 3 | 3232920 | 1.69E-05 | 0.00334 | 299 | 0.139407048 | 0.016446186 |
| SNP_A.K_758816_4_19381497 | 4 | 19381497 | 1.69E-05 | 0.00334 | 299 | 0.139407048 | 0.016446186 |
| SNP_A.K_1393679_8_10503898 | 8 | 10503898 | 1.69E-05 | 0.00334 | 299 | 0.139407048 | 0.016446186 |
| SNP_A.K_2070141_12_25468009 | 12 | 25468009 | 1.69E-05 | 0.00334 | 299 | 0.139407048 | 0.016446186 |
| SNP_A.K_1004407_6_6879927 | 6 | 6879927 | 1.69E-05 | 0.41137 | 299 | 0.139401241 | 0.016453562 |
| SNP_A.K_1004970_6_6959150 | 6 | 6959150 | 1.70E-05 | 0.31605 | 299 | 0.139389046 | 0.01647753 |
| SNP_A.K_978849_6_1787372 | 6 | 1787372 | 1.70E-05 | 0.42809 | 299 | 0.13937753 | 0.016499772 |
| SNP_A.K_1000924_6_6424130 | 6 | 6424130 | 1.70E-05 | 0.33779 | 299 | 0.139363329 | 0.016529045 |

| | | | | | | | |
|---|---|---|---|---|---|---|---|
| SNP_A.K_978596_6_1736937 | 6 | 1736937 | 1.71E-05 | 0.27759 | 299 | 0.139350882 | 0.016553796 |
| SNP_A.K_306968_2_7371107 | 2 | 7371107 | 1.71E-05 | 0.25753 | 299 | 0.139344565 | 0.016556101 |
| SNP_A.K_1600845_9_18849267 | 9 | 18849267 | 1.71E-05 | 0.14047 | 299 | 0.13934409 | 0.016556101 |
| SNP_A.K_28912_1_4181491 | 1 | 4181491 | 1.71E-05 | 0.10201 | 299 | 0.139339123 | 0.016561359 |
| SNP_A.K_1001703_6_6506973 | 6 | 6506973 | 1.71E-05 | 0.31271 | 299 | 0.139324503 | 0.016591857 |
| SNP_A.K_307009_2_7372928 | 2 | 7372928 | 1.72E-05 | 0.25753 | 299 | 0.139316132 | 0.016606046 |
| SNP_A.K_1001426_6_6481468 | 6 | 6481468 | 1.72E-05 | 0.29933 | 299 | 0.139306282 | 0.016624127 |
| SNP_A.K_1001613_6_6498392 | 6 | 6498392 | 1.72E-05 | 0.40635 | 299 | 0.139300553 | 0.01663142 |
| SNP_A.K_1263508_7_18652113 | 7 | 18652113 | 1.72E-05 | 0.49164 | 299 | 0.139291909 | 0.016646371 |
| SNP_A.K_725201_4_11762294 | 4 | 11762294 | 1.73E-05 | 0.09197 | 299 | 0.139282052 | 0.016664527 |
| SNP_A.K_1794678_11_5475941 | 11 | 5475941 | 1.73E-05 | 0.43813 | 299 | 0.139276112 | 0.016672406 |
| SNP_A.K_194802_1_30962904 | 1 | 30962904 | 1.73E-05 | 0.49498 | 299 | 0.13926621 | 0.016690715 |
| SNP_A.K_308459_2_7517454 | 2 | 7517454 | 1.73E-05 | 0.26087 | 299 | 0.139261373 | 0.016695708 |
| SNP_A.K_994759_6_5887555 | 6 | 5887555 | 1.74E-05 | 0.29097 | 299 | 0.139239152 | 0.016746551 |
| SNP_A.K_1001301_6_6463227 | 6 | 6463227 | 1.74E-05 | 0.29097 | 299 | 0.139220981 | 0.016773329 |
| SNP_A.K_773092_4_22078266 | 4 | 22078266 | 1.74E-05 | 0.03177 | 299 | 0.139220214 | 0.016773329 |
| SNP_A.K_914842_5_18422620 | 5 | 18422620 | 1.74E-05 | 0.03177 | 299 | 0.139220214 | 0.016773329 |
| SNP_A.K_978807_6_1783373 | 6 | 1783373 | 1.75E-05 | 0.3913 | 299 | 0.139198555 | 0.016817751 |
| SNP_A.K_2078610_12_27000452 | 12 | 27000452 | 1.75E-05 | 0.01672 | 299 | 0.139197578 | 0.016817751 |
| SNP_A.K_1001252_6_6458927 | 6 | 6458927 | 1.75E-05 | 0.29097 | 299 | 0.13919138 | 0.016826421 |
| SNP_A.K_1001084_6_6440036 | 6 | 6440036 | 1.75E-05 | 0.33445 | 299 | 0.139183284 | 0.01684014 |
| SNP_A.K_1004916_6_6952374 | 6 | 6952374 | 1.75E-05 | 0.31605 | 299 | 0.139174191 | 0.016856525 |
| SNP_A.K_1004362_6_6876460 | 6 | 6876460 | 1.76E-05 | 0.32107 | 299 | 0.13915259 | 0.016904203 |
| SNP_A.K_1001358_6_6466717 | 6 | 6466717 | 1.76E-05 | 0.29097 | 299 | 0.139150432 | 0.016904203 |
| SNP_A.K_1004618_6_6910879 | 6 | 6910879 | 1.77E-05 | 0.3194 | 299 | 0.139134094 | 0.016940005 |
| SNP_A.K_679337_4_1073273 | 4 | 1073273 | 1.77E-05 | 0.04348 | 299 | 0.139127974 | 0.016948554 |
| SNP_A.K_1608534_9_20772805 | 9 | 20772805 | 1.77E-05 | 0.30936 | 299 | 0.139109384 | 0.016990493 |
| SNP_A.K_996661_6_6015678 | 6 | 6015678 | 1.78E-05 | 0.48161 | 299 | 0.139078692 | 0.017065085 |
| SNP_A.K_326667_2_10037678 | 2 | 10037678 | 1.78E-05 | 0.06689 | 299 | 0.139073088 | 0.017072318 |
| SNP_A.K_1608728_9_20820729 | 9 | 20820729 | 1.79E-05 | 0.31438 | 299 | 0.139050745 | 0.017124716 |
| SNP_A.K_307170_2_7383164 | 2 | 7383164 | 1.79E-05 | 0.23913 | 299 | 0.139043096 | 0.017137509 |
| SNP_A.K_990334_6_4711194 | 6 | 4711194 | 1.80E-05 | 0.20903 | 299 | 0.139028149 | 0.017169842 |
| SNP_A.K_326860_2_10061629 | 2 | 10061629 | 1.80E-05 | 0.06187 | 299 | 0.139025323 | 0.017169842 |
| SNP_A.K_306820_2_7355642 | 2 | 7355642 | 1.81E-05 | 0.23913 | 299 | 0.138994046 | 0.017246848 |
| SNP_A.K_978827_6_1786554 | 6 | 1786554 | 1.81E-05 | 0.38796 | 299 | 0.138984672 | 0.017264447 |
| SNP_A.K_977952_6_1624213 | 6 | 1624213 | 1.81E-05 | 0.20903 | 299 | 0.138978311 | 0.017273856 |
| SNP_A.K_1831308_11_10618138 | 11 | 10618138 | 1.82E-05 | 0.13043 | 299 | 0.138933487 | 0.017383489 |
| SNP_A.K_995838_6_5952331 | 6 | 5952331 | 1.82E-05 | 0.29933 | 299 | 0.138932399 | 0.017383489 |
| SNP_A.K_978888_6_1796720 | 6 | 1796720 | 1.83E-05 | 0.3194 | 299 | 0.138902069 | 0.017458874 |
| SNP_A.K_978654_6_1750520 | 6 | 1750520 | 1.83E-05 | 0.29097 | 299 | 0.138891967 | 0.01747304 |
| SNP_A.K_2068321_12_24995670 | 12 | 24995670 | 1.84E-05 | 0.02174 | 299 | 0.138891128 | 0.01747304 |
| SNP_A.K_277981_2_2983844 | 2 | 2983844 | 1.85E-05 | 0.06856 | 299 | 0.138852401 | 0.017572099 |
| SNP_A.K_307458_2_7408644 | 2 | 7408644 | 1.85E-05 | 0.25585 | 299 | 0.138849332 | 0.017572575 |
| SNP_A.K_978463_6_1696139 | 6 | 1696139 | 1.85E-05 | 0.25585 | 299 | 0.138833667 | 0.017606352 |
| SNP_A.K_1608561_9_20777961 | 9 | 20777961 | 1.85E-05 | 0.25084 | 299 | 0.138831372 | 0.017606352 |
| SNP_A.K_1953761_12_3627549 | 12 | 3627549 | 1.86E-05 | 0.05351 | 299 | 0.138809929 | 0.017657963 |
| SNP_A.K_978371_6_1682730 | 6 | 1682730 | 1.86E-05 | 0.2592 | 299 | 0.138803465 | 0.017667921 |
| SNP_A.K_1600851_9_18851906 | 9 | 18851906 | 1.86E-05 | 0.12542 | 299 | 0.138796245 | 0.017674589 |
| SNP_A.K_978938_6_1814552 | 6 | 1814552 | 1.86E-05 | 0.31605 | 299 | 0.138795295 | 0.017674589 |
| SNP_A.K_1857590_11_14815205 | 11 | 14815205 | 1.86E-05 | 0.02676 | 299 | 0.138792049 | 0.017675588 |

| | | | | | | | |
|---|---|---|---|---|---|---|---|
| SNP_A.K_993237_6_5723448 | 6 | 5723448 | 1.87E-05 | 0.47826 | 299 | 0.138785565 | 0.017681605 |
| SNP_A.K_995851_6_5952494 | 6 | 5952494 | 1.87E-05 | 0.48495 | 299 | 0.138783937 | 0.017681605 |
| SNP_A.K_1001477_6_6483102 | 6 | 6483102 | 1.87E-05 | 0.29097 | 299 | 0.138773647 | 0.017681605 |
| SNP_A.K_1001476_6_6483093 | 6 | 6483093 | 1.87E-05 | 0.29097 | 299 | 0.138773647 | 0.017681605 |
| SNP_A.K_992585_6_5659711 | 6 | 5659711 | 1.87E-05 | 0.49331 | 299 | 0.138772595 | 0.017681605 |
| SNP_A.K_992584_6_5659709 | 6 | 5659709 | 1.87E-05 | 0.49331 | 299 | 0.138772595 | 0.017681605 |
| SNP_A.K_977993_6_1632138 | 6 | 1632138 | 1.88E-05 | 0.25251 | 299 | 0.138727582 | 0.017799573 |
| SNP_A.K_1001250_6_6458797 | 6 | 6458797 | 1.88E-05 | 0.37793 | 299 | 0.138721923 | 0.017806034 |
| SNP_A.K_1000978_6_6431297 | 6 | 6431297 | 1.89E-05 | 0.33278 | 299 | 0.138719531 | 0.017806034 |
| SNP_A.K_993887_6_5799400 | 6 | 5799400 | 1.89E-05 | 0.17893 | 299 | 0.138711485 | 0.017811549 |
| SNP_A.K_923810_5_19502424 | 5 | 19502424 | 1.89E-05 | 0.02843 | 299 | 0.138709157 | 0.017811549 |
| SNP_A.K_1004942_6_6956596 | 6 | 6956596 | 1.89E-05 | 0.29766 | 299 | 0.13870895 | 0.017811549 |
| SNP_A.K_308573_2_7527333 | 2 | 7527333 | 1.89E-05 | 0.2107 | 299 | 0.138696134 | 0.017839532 |
| SNP_A.K_978652_6_1750477 | 6 | 1750477 | 1.90E-05 | 0.27592 | 299 | 0.138685315 | 0.01786193 |
| SNP_A.K_1000748_6_6414480 | 6 | 6414480 | 1.90E-05 | 0.29097 | 299 | 0.138667935 | 0.017902894 |
| SNP_A.K_710549_4_8426985 | 4 | 8426985 | 1.90E-05 | 0.10535 | 299 | 0.138664009 | 0.017905889 |
| SNP_A.K_1925432_11_24737276 | 11 | 24737276 | 1.91E-05 | 0.02174 | 299 | 0.138642218 | 0.017958864 |
| SNP_A.K_1600886_9_18859623 | 9 | 18859623 | 1.91E-05 | 0.1388 | 299 | 0.138639566 | 0.017958864 |
| SNP_A.K_725250_4_11767812 | 4 | 11767812 | 1.91E-05 | 0.09866 | 299 | 0.138624337 | 0.017993956 |
| SNP_A.K_2067441_12_24765123 | 12 | 24765123 | 1.92E-05 | 0.04181 | 299 | 0.138619819 | 0.017998665 |
| SNP_A.K_306846_2_7359890 | 2 | 7359890 | 1.92E-05 | 0.27425 | 299 | 0.138605099 | 0.018026192 |
| SNP_A.K_1183398_7_4540222 | 7 | 4540222 | 1.92E-05 | 0.3796 | 299 | 0.13860442 | 0.018026192 |
| SNP_A.K_678978_4_1020672 | 4 | 1020672 | 1.92E-05 | 0.0301 | 299 | 0.138600603 | 0.018028926 |
| SNP_A.K_1608651_9_20810430 | 9 | 20810430 | 1.93E-05 | 0.30769 | 299 | 0.138586484 | 0.018058992 |
| SNP_A.K_1389904_8_10134749 | 8 | 10134749 | 1.93E-05 | 0.04849 | 299 | 0.138584338 | 0.018058992 |
| SNP_A.K_1001475_6_6483066 | 6 | 6483066 | 1.93E-05 | 0.29264 | 299 | 0.13857876 | 0.018066765 |
| SNP_A.K_978218_6_1662157 | 6 | 1662157 | 1.93E-05 | 0.19064 | 299 | 0.138562887 | 0.018103941 |
| SNP_A.K_1600723_9_18826389 | 9 | 18826389 | 1.93E-05 | 0.15385 | 299 | 0.138558299 | 0.01810891 |
| SNP_A.K_1389983_8_10141955 | 8 | 10141955 | 1.94E-05 | 0.05351 | 299 | 0.138543708 | 0.018136658 |
| SNP_A.K_196605_1_31436461 | 1 | 31436461 | 1.94E-05 | 0.47324 | 299 | 0.138541565 | 0.018136658 |
| SNP_A.K_1600786_9_18837702 | 9 | 18837702 | 1.94E-05 | 0.16054 | 299 | 0.138540058 | 0.018136658 |
| SNP_A.K_725829_4_11867677 | 4 | 11867677 | 1.95E-05 | 0.0903 | 299 | 0.138521572 | 0.018181503 |
| SNP_A.K_709970_4_8303846 | 4 | 8303846 | 1.96E-05 | 0.07023 | 299 | 0.13848457 | 0.018236058 |
| SNP_A.K_709972_4_8303874 | 4 | 8303874 | 1.96E-05 | 0.07023 | 299 | 0.13848457 | 0.018236058 |
| SNP_A.K_709789_4_8297798 | 4 | 8297798 | 1.96E-05 | 0.07023 | 299 | 0.13848457 | 0.018236058 |
| SNP_A.K_709815_4_8298327 | 4 | 8298327 | 1.96E-05 | 0.07023 | 299 | 0.13848457 | 0.018236058 |
| SNP_A.K_710479_4_8424176 | 4 | 8424176 | 1.96E-05 | 0.07023 | 299 | 0.13848457 | 0.018236058 |
| SNP_A.K_978381_6_1684625 | 6 | 1684625 | 1.96E-05 | 0.26756 | 299 | 0.138483965 | 0.018236058 |
| SNP_A.K_1600954_9_18866383 | 9 | 18866383 | 1.96E-05 | 0.13211 | 299 | 0.138482719 | 0.018236058 |
| SNP_A.K_307808_2_7448965 | 2 | 7448965 | 1.96E-05 | 0.47157 | 299 | 0.138470189 | 0.018259048 |
| SNP_A.K_307246_2_7387872 | 2 | 7387872 | 1.96E-05 | 0.24916 | 299 | 0.138469072 | 0.018259048 |
| SNP_A.K_977896_6_1616444 | 6 | 1616444 | 1.97E-05 | 0.15886 | 299 | 0.138456112 | 0.018288271 |
| SNP_A.K_1608537_9_20773129 | 9 | 20773129 | 1.97E-05 | 0.3194 | 299 | 0.138450403 | 0.018296587 |
| SNP_A.K_1608517_9_20770921 | 9 | 20770921 | 1.97E-05 | 0.29766 | 299 | 0.138446862 | 0.018298643 |
| SNP_A.K_672649_4_268874 | 4 | 268874 | 1.97E-05 | 0.10368 | 299 | 0.138443022 | 0.018301569 |
| SNP_A.K_978715_6_1770024 | 6 | 1770024 | 1.97E-05 | 0.30936 | 299 | 0.13843342 | 0.018321158 |
| SNP_A.K_1001445_6_6481844 | 6 | 6481844 | 1.97E-05 | 0.30602 | 299 | 0.138427661 | 0.018328901 |
| SNP_A.K_978185_6_1658471 | 6 | 1658471 | 1.98E-05 | 0.2592 | 299 | 0.138425095 | 0.018328901 |
| SNP_A.K_307086_2_7378350 | 2 | 7378350 | 1.98E-05 | 0.4097 | 299 | 0.138418638 | 0.018339424 |
| SNP_A.K_992003_6_5607940 | 6 | 5607940 | 1.98E-05 | 0.18896 | 299 | 0.138403726 | 0.018374471 |

| | | | | | | | |
|---|---|---|---|---|---|---|---|
| SNP_A.K_307061_2_7376491 | 2 | 7376491 | 1.98E-05 | 0.23077 | 299 | 0.13839961 | 0.018378233 |
| SNP_A.K_1001746_6_6513819 | 6 | 6513819 | 1.99E-05 | 0.30936 | 299 | 0.138373621 | 0.018445603 |
| SNP_A.K_978170_6_1657659 | 6 | 1657659 | 1.99E-05 | 0.19732 | 299 | 0.138363987 | 0.018459544 |
| SNP_A.K_978620_6_1745032 | 6 | 1745032 | 2.00E-05 | 0.27759 | 299 | 0.138361244 | 0.018459544 |
| SNP_A.K_978232_6_1663253 | 6 | 1663253 | 2.00E-05 | 0.16388 | 299 | 0.138359006 | 0.018459544 |
| SNP_A.K_306845_2_7359597 | 2 | 7359597 | 2.00E-05 | 0.26087 | 299 | 0.13835499 | 0.018459544 |
| SNP_A.K_977882_6_1611020 | 6 | 1611020 | 2.00E-05 | 0.23077 | 299 | 0.138353194 | 0.018459544 |
| SNP_A.K_673336_4_349555 | 4 | 349555 | 2.00E-05 | 0.08027 | 299 | 0.138351951 | 0.018459544 |
| SNP_A.K_173304_1_27886048 | 1 | 27886048 | 2.00E-05 | 0.02007 | 299 | 0.138339284 | 0.018475066 |
| SNP_A.K_978901_6_1799707 | 6 | 1799707 | 2.00E-05 | 0.31271 | 299 | 0.138339272 | 0.018475066 |
| SNP_A.K_306742_2_7347142 | 2 | 7347142 | 2.00E-05 | 0.24415 | 299 | 0.138338204 | 0.018475066 |
| SNP_A.K_678915_4_1007842 | 4 | 1007842 | 2.00E-05 | 0.02843 | 299 | 0.13833249 | 0.018482703 |
| SNP_A.K_726139_4_11919898 | 4 | 11919898 | 2.00E-05 | 0.09365 | 299 | 0.138329976 | 0.018482703 |
| SNP_A.K_709573_4_8274315 | 4 | 8274315 | 2.01E-05 | 0.10033 | 299 | 0.138311166 | 0.018526017 |
| SNP_A.K_996216_6_5975664 | 6 | 5975664 | 2.01E-05 | 0.15886 | 299 | 0.13830955 | 0.018526017 |
| SNP_A.K_247987_1_40109178 | 1 | 40109178 | 2.01E-05 | 0.02341 | 299 | 0.138305238 | 0.018528647 |
| SNP_A.K_306874_2_7362893 | 2 | 7362893 | 2.01E-05 | 0.23913 | 299 | 0.13830305 | 0.018528647 |
| SNP_A.K_321179_2_8956660 | 2 | 8956660 | 2.02E-05 | 0.03679 | 299 | 0.138292303 | 0.018551925 |
| SNP_A.K_413974_2_23757310 | 2 | 23757310 | 2.02E-05 | 0.05351 | 299 | 0.138278893 | 0.018576411 |
| SNP_A.K_978033_6_1641241 | 6 | 1641241 | 2.02E-05 | 0.20234 | 299 | 0.1382767 | 0.018576411 |
| SNP_A.K_307254_2_7388069 | 2 | 7388069 | 2.02E-05 | 0.26756 | 299 | 0.138275565 | 0.018576411 |
| SNP_A.K_456482_2_33043765 | 2 | 33043765 | 2.03E-05 | 0.06689 | 299 | 0.138248186 | 0.018648691 |
| SNP_A.K_1371601_8_7858209 | 8 | 7858209 | 2.03E-05 | 0.01338 | 299 | 0.138242041 | 0.018653432 |
| SNP_A.K_1431482_8_16377860 | 8 | 16377860 | 2.03E-05 | 0.13712 | 299 | 0.138239155 | 0.018653432 |
| SNP_A.K_526222_3_10849783 | 3 | 10849783 | 2.03E-05 | 0.01338 | 299 | 0.138236426 | 0.018653432 |
| SNP_A.K_726193_4_11927469 | 4 | 11927469 | 2.04E-05 | 0.09866 | 299 | 0.138235416 | 0.018653432 |
| SNP_A.K_978061_6_1644579 | 6 | 1644579 | 2.04E-05 | 0.18562 | 299 | 0.138229308 | 0.01866322 |
| SNP_A.K_326354_2_9993240 | 2 | 9993240 | 2.05E-05 | 0.0786 | 299 | 0.138197113 | 0.018748472 |
| SNP_A.K_710121_4_8317314 | 4 | 8317314 | 2.05E-05 | 0.11371 | 299 | 0.138194625 | 0.018748472 |
| SNP_A.K_978678_6_1758589 | 6 | 1758589 | 2.05E-05 | 0.30602 | 299 | 0.138192104 | 0.018748472 |
| SNP_A.K_1005547_6_7022827 | 6 | 7022827 | 2.06E-05 | 0.26254 | 299 | 0.138158388 | 0.01884032 |
| SNP_A.K_710485_4_8424659 | 4 | 8424659 | 2.06E-05 | 0.07023 | 299 | 0.138149211 | 0.018851088 |
| SNP_A.K_709822_4_8298510 | 4 | 8298510 | 2.06E-05 | 0.07023 | 299 | 0.138149211 | 0.018851088 |
| SNP_A.K_1608577_9_20788929 | 9 | 20788929 | 2.07E-05 | 0.24582 | 299 | 0.138142924 | 0.018861544 |
| SNP_A.K_1001261_6_6459730 | 6 | 6459730 | 2.07E-05 | 0.3796 | 299 | 0.138137781 | 0.018867751 |
| SNP_A.K_1004288_6_6862878 | 6 | 6862878 | 2.07E-05 | 0.33612 | 299 | 0.138135289 | 0.018867751 |
| SNP_A.K_456492_2_33047027 | 2 | 33047027 | 2.07E-05 | 0.07023 | 299 | 0.138128276 | 0.01888039 |
| SNP_A.K_978682_6_1760469 | 6 | 1760469 | 2.07E-05 | 0.3495 | 299 | 0.13812048 | 0.018895378 |
| SNP_A.K_1001303_6_6463326 | 6 | 6463326 | 2.08E-05 | 0.30268 | 299 | 0.138110992 | 0.01891544 |
| SNP_A.K_726266_4_11932060 | 4 | 11932060 | 2.08E-05 | 0.09699 | 299 | 0.138089721 | 0.018970811 |
| SNP_A.K_1004491_6_6891800 | 6 | 6891800 | 2.09E-05 | 0.3194 | 299 | 0.138080739 | 0.018989444 |
| SNP_A.K_1600935_9_18862680 | 9 | 18862680 | 2.09E-05 | 0.13378 | 299 | 0.13806202 | 0.019037363 |
| SNP_A.K_1001552_6_6490436 | 6 | 6490436 | 2.10E-05 | 0.29264 | 299 | 0.138051782 | 0.019059854 |
| SNP_A.K_1004814_6_6933844 | 6 | 6933844 | 2.10E-05 | 0.31104 | 299 | 0.138035747 | 0.01909986 |
| SNP_A.K_1600755_9_18833875 | 9 | 18833875 | 2.11E-05 | 0.13545 | 299 | 0.138010894 | 0.019166644 |
| SNP_A.K_709824_4_8298546 | 4 | 8298546 | 2.11E-05 | 0.07191 | 299 | 0.138007703 | 0.019167941 |
| SNP_A.K_1001350_6_6466490 | 6 | 6466490 | 2.12E-05 | 0.29431 | 299 | 0.137990775 | 0.019202507 |
| SNP_A.K_685936_4_2520135 | 4 | 2520135 | 2.12E-05 | 0.06522 | 299 | 0.137989952 | 0.019202507 |
| SNP_A.K_278114_2_3027804 | 2 | 3027804 | 2.12E-05 | 0.11538 | 299 | 0.137988021 | 0.019202507 |
| SNP_A.K_990278_6_4704331 | 6 | 4704331 | 2.12E-05 | 0.19064 | 299 | 0.137977654 | 0.019225534 |

| | | | | | | | |
|---|---|---|---|---|---|---|---|
| SNP_A.K_1389975_8_10140363 | 8 | 10140363 | 2.12E-05 | 0.05184 | 299 | 0.137972532 | 0.019225534 |
| SNP_A.K_513667_3_8307268 | 3 | 8307268 | 2.12E-05 | 0.02508 | 299 | 0.137972168 | 0.019225534 |
| SNP_A.K_1726549_10_16598195 | 10 | 16598195 | 2.14E-05 | 0.05686 | 299 | 0.137917181 | 0.019384898 |
| SNP_A.K_456435_2_33036164 | 2 | 33036164 | 2.15E-05 | 0.07023 | 299 | 0.137899963 | 0.019429267 |
| SNP_A.K_307517_2_7414933 | 2 | 7414933 | 2.16E-05 | 0.43645 | 299 | 0.137858234 | 0.019549338 |
| SNP_A.K_1608523_9_20772268 | 9 | 20772268 | 2.16E-05 | 0.28595 | 299 | 0.137849235 | 0.019568657 |
| SNP_A.K_710386_4_8339494 | 4 | 8339494 | 2.17E-05 | 0.08361 | 299 | 0.137840931 | 0.019585849 |
| SNP_A.K_977894_6_1616300 | 6 | 1616300 | 2.17E-05 | 0.24415 | 299 | 0.137818401 | 0.01964719 |
| SNP_A.K_1001202_6_6454837 | 6 | 6454837 | 2.18E-05 | 0.38796 | 299 | 0.137804733 | 0.019681141 |
| SNP_A.K_1000966_6_6428623 | 6 | 6428623 | 2.19E-05 | 0.35953 | 299 | 0.137781069 | 0.019746335 |
| SNP_A.K_710462_4_8422304 | 4 | 8422304 | 2.19E-05 | 0.09365 | 299 | 0.13776692 | 0.019781971 |
| SNP_A.K_990269_6_4703881 | 6 | 4703881 | 2.21E-05 | 0.19565 | 299 | 0.137719904 | 0.019920923 |
| SNP_A.K_995031_6_5912817 | 6 | 5912817 | 2.21E-05 | 0.16722 | 299 | 0.137714811 | 0.01992833 |
| SNP_A.K_1004788_6_6932381 | 6 | 6932381 | 2.21E-05 | 0.33445 | 299 | 0.137710131 | 0.019934447 |
| SNP_A.K_1001196_6_6454626 | 6 | 6454626 | 2.23E-05 | 0.3194 | 299 | 0.137670765 | 0.020050226 |
| SNP_A.K_978060_6_1644571 | 6 | 1644571 | 2.23E-05 | 0.18729 | 299 | 0.137656622 | 0.020086419 |
| SNP_A.K_978471_6_1697015 | 6 | 1697015 | 2.23E-05 | 0.26756 | 299 | 0.137651297 | 0.020094645 |
| SNP_A.K_312076_2_7903907 | 2 | 7903907 | 2.24E-05 | 0.27592 | 299 | 0.137632158 | 0.020146838 |
| SNP_A.K_1001307_6_6463602 | 6 | 6463602 | 2.24E-05 | 0.29933 | 299 | 0.137628025 | 0.020151296 |
| SNP_A.K_525231_3_10652553 | 3 | 10652553 | 2.24E-05 | 0.01171 | 299 | 0.137618181 | 0.020162771 |
| SNP_A.K_997787_6_6147326 | 6 | 6147326 | 2.25E-05 | 0.14214 | 299 | 0.137611557 | 0.020162771 |
| SNP_A.K_1371659_8_7863147 | 8 | 7863147 | 2.25E-05 | 0.02676 | 299 | 0.13760999 | 0.020162771 |
| SNP_A.K_978426_6_1689652 | 6 | 1689652 | 2.25E-05 | 0.28428 | 299 | 0.137598207 | 0.020162771 |
| SNP_A.K_995138_6_5916987 | 6 | 5916987 | 2.25E-05 | 0.44816 | 299 | 0.137591515 | 0.020162771 |
| SNP_A.K_307081_2_7378298 | 2 | 7378298 | 2.25E-05 | 0.24749 | 299 | 0.137588904 | 0.020162771 |
| SNP_A.K_477471_3_1827167 | 3 | 1827167 | 2.26E-05 | 0.01003 | 299 | 0.137578099 | 0.020162771 |
| SNP_A.K_1804384_11_7066211 | 11 | 7066211 | 2.26E-05 | 0.01003 | 299 | 0.137578099 | 0.020162771 |
| SNP_A.K_1804390_11_7066400 | 11 | 7066400 | 2.26E-05 | 0.01003 | 299 | 0.137578099 | 0.020162771 |
| SNP_A.K_1805586_11_7213751 | 11 | 7213751 | 2.26E-05 | 0.01003 | 299 | 0.137578099 | 0.020162771 |
| SNP_A.K_1803419_11_6912490 | 11 | 6912490 | 2.26E-05 | 0.01003 | 299 | 0.137578099 | 0.020162771 |
| SNP_A.K_1804371_11_7064881 | 11 | 7064881 | 2.26E-05 | 0.01003 | 299 | 0.137578099 | 0.020162771 |
| SNP_A.K_1804392_11_7066673 | 11 | 7066673 | 2.26E-05 | 0.01003 | 299 | 0.137578099 | 0.020162771 |
| SNP_A.K_1804556_11_7086080 | 11 | 7086080 | 2.26E-05 | 0.01003 | 299 | 0.137578099 | 0.020162771 |
| SNP_A.K_1805532_11_7199542 | 11 | 7199542 | 2.26E-05 | 0.01003 | 299 | 0.137578099 | 0.020162771 |
| SNP_A.K_1805590_11_7215506 | 11 | 7215506 | 2.26E-05 | 0.01003 | 299 | 0.137578099 | 0.020162771 |
| SNP_A.K_907830_5_17459266 | 5 | 17459266 | 2.26E-05 | 0.04682 | 299 | 0.137577455 | 0.020162771 |
| SNP_A.K_1001185_6_6453015 | 6 | 6453015 | 2.26E-05 | 0.28094 | 299 | 0.137574274 | 0.020162771 |
| SNP_A.K_307391_2_7400428 | 2 | 7400428 | 2.26E-05 | 0.24916 | 299 | 0.137571924 | 0.020162771 |
| SNP_A.K_709910_4_8301630 | 4 | 8301630 | 2.26E-05 | 0.07191 | 299 | 0.137570005 | 0.020162771 |
| SNP_A.K_306924_2_7368454 | 2 | 7368454 | 2.27E-05 | 0.28428 | 299 | 0.137550182 | 0.020192058 |
| SNP_A.K_456485_2_33044400 | 2 | 33044400 | 2.27E-05 | 0.07023 | 299 | 0.137550091 | 0.020192058 |
| SNP_A.K_307088_2_7378360 | 2 | 7378360 | 2.27E-05 | 0.4097 | 299 | 0.137548183 | 0.020192058 |
| SNP_A.K_709765_4_8285352 | 4 | 8285352 | 2.27E-05 | 0.0903 | 299 | 0.137546027 | 0.020192058 |
| SNP_A.K_1389979_8_10141545 | 8 | 10141545 | 2.27E-05 | 0.05017 | 299 | 0.137545636 | 0.020192058 |
| SNP_A.K_527065_3_11050383 | 3 | 11050383 | 2.27E-05 | 0.01338 | 299 | 0.137543443 | 0.020192058 |
| SNP_A.K_307289_2_7391398 | 2 | 7391398 | 2.27E-05 | 0.2592 | 299 | 0.137541892 | 0.020192058 |
| SNP_A.K_978806_6_1783005 | 6 | 1783005 | 2.28E-05 | 0.40301 | 299 | 0.13750833 | 0.020290859 |
| SNP_A.K_995205_6_5919314 | 6 | 5919314 | 2.29E-05 | 0.33278 | 299 | 0.13750247 | 0.020292759 |
| SNP_A.K_709691_4_8281401 | 4 | 8281401 | 2.29E-05 | 0.08696 | 299 | 0.137502341 | 0.020292759 |
| SNP_A.K_725251_4_11767851 | 4 | 11767851 | 2.29E-05 | 0.09866 | 299 | 0.137496979 | 0.020292881 |

| | | | | | | | |
|---|---|---|---|---|---|---|---|
| SNP_A.K_1001101_6_6441961 | 6 | 6441961 | 2.29E-05 | 0.38294 | 299 | 0.13749691 | 0.020292881 |
| SNP_A.K_978357_6_1681604 | 6 | 1681604 | 2.29E-05 | 0.29097 | 299 | 0.137493816 | 0.020294164 |
| SNP_A.K_996769_6_6025039 | 6 | 6025039 | 2.29E-05 | 0.17057 | 299 | 0.137481521 | 0.02032501 |
| SNP_A.K_263557_1_42826008 | 1 | 42826008 | 2.29E-05 | 0.00167 | 299 | 0.137478554 | 0.020325894 |
| SNP_A.K_978096_6_1647087 | 6 | 1647087 | 2.30E-05 | 0.18227 | 299 | 0.13746739 | 0.020353154 |
| SNP_A.K_1432892_8_16594308 | 8 | 16594308 | 2.31E-05 | 0.08528 | 299 | 0.137437204 | 0.020441869 |
| SNP_A.K_710463_4_8422319 | 4 | 8422319 | 2.31E-05 | 0.09197 | 299 | 0.137424169 | 0.020475353 |
| SNP_A.K_725491_4_11817067 | 4 | 11817067 | 2.31E-05 | 0.09866 | 299 | 0.137420653 | 0.020478039 |
| SNP_A.K_2073409_12_26025908 | 12 | 26025908 | 2.32E-05 | 0.02676 | 299 | 0.13741393 | 0.02049112 |
| SNP_A.K_1001467_6_6482727 | 6 | 6482727 | 2.32E-05 | 0.29264 | 299 | 0.13740637 | 0.020506928 |
| SNP_A.K_1608498_9_20765459 | 9 | 20765459 | 2.33E-05 | 0.30435 | 299 | 0.137391774 | 0.02054561 |
| SNP_A.K_908259_5_17494747 | 5 | 17494747 | 2.33E-05 | 0.02174 | 299 | 0.1373877 | 0.020550134 |
| SNP_A.K_2071311_12_25697089 | 12 | 25697089 | 2.33E-05 | 0.07023 | 299 | 0.137384063 | 0.020551379 |
| SNP_A.K_725359_4_11798174 | 4 | 11798174 | 2.33E-05 | 0.11037 | 299 | 0.137381956 | 0.020551379 |
| SNP_A.K_1830597_11_10569405 | 11 | 10569405 | 2.33E-05 | 0.40468 | 299 | 0.137369376 | 0.020577835 |
| SNP_A.K_1608746_9_20829513 | 9 | 20829513 | 2.33E-05 | 0.26087 | 299 | 0.137368467 | 0.020577835 |
| SNP_A.K_978611_6_1741111 | 6 | 1741111 | 2.35E-05 | 0.28763 | 299 | 0.137336856 | 0.020663875 |
| SNP_A.K_1608657_9_20811030 | 9 | 20811030 | 2.35E-05 | 0.2592 | 299 | 0.137336742 | 0.020663875 |
| SNP_A.K_86123_1_11757862 | 1 | 11757862 | 2.35E-05 | 0.23913 | 299 | 0.137323988 | 0.020696853 |
| SNP_A.K_978016_6_1637153 | 6 | 1637153 | 2.35E-05 | 0.16054 | 299 | 0.137316963 | 0.020711108 |
| SNP_A.K_978380_6_1684543 | 6 | 1684543 | 2.36E-05 | 0.26421 | 299 | 0.137311334 | 0.020720803 |
| SNP_A.K_994834_6_5893355 | 6 | 5893355 | 2.36E-05 | 0.40635 | 299 | 0.137287122 | 0.020791546 |
| SNP_A.K_977991_6_1631305 | 6 | 1631305 | 2.37E-05 | 0.16054 | 299 | 0.137267979 | 0.020845811 |
| SNP_A.K_307986_2_7461417 | 2 | 7461417 | 2.38E-05 | 0.46823 | 299 | 0.137253546 | 0.020884653 |
| SNP_A.K_1000961_6_6427421 | 6 | 6427421 | 2.39E-05 | 0.33445 | 299 | 0.137218541 | 0.020988751 |
| SNP_A.K_1600748_9_18831447 | 9 | 18831447 | 2.39E-05 | 0.14047 | 299 | 0.137216793 | 0.020988751 |
| SNP_A.K_1001503_6_6484980 | 6 | 6484980 | 2.39E-05 | 0.32107 | 299 | 0.137212016 | 0.020995768 |
| SNP_A.K_1001370_6_6471406 | 6 | 6471406 | 2.40E-05 | 0.29264 | 299 | 0.137204644 | 0.021011418 |
| SNP_A.K_1004448_6_6883190 | 6 | 6883190 | 2.41E-05 | 0.48328 | 299 | 0.137174964 | 0.02109289 |
| SNP_A.K_752428_4_17906030 | 4 | 17906030 | 2.41E-05 | 0.06355 | 299 | 0.137172206 | 0.02109289 |
| SNP_A.K_752429_4_17906042 | 4 | 17906042 | 2.41E-05 | 0.06355 | 299 | 0.137172206 | 0.02109289 |
| SNP_A.K_990324_6_4710649 | 6 | 4710649 | 2.41E-05 | 0.19732 | 299 | 0.137165186 | 0.021107452 |
| SNP_A.K_679324_4_1072950 | 4 | 1072950 | 2.41E-05 | 0.03512 | 299 | 0.137158933 | 0.021119467 |
| SNP_A.K_993958_6_5802505 | 6 | 5802505 | 2.42E-05 | 0.23411 | 299 | 0.137145003 | 0.02115718 |
| SNP_A.K_1004866_6_6937792 | 6 | 6937792 | 2.42E-05 | 0.32943 | 299 | 0.137129057 | 0.021201731 |
| SNP_A.K_673117_4_318549 | 4 | 318549 | 2.43E-05 | 0.04013 | 299 | 0.137124027 | 0.021202411 |
| SNP_A.K_306781_2_7350003 | 2 | 7350003 | 2.43E-05 | 0.26087 | 299 | 0.137121191 | 0.021202411 |
| SNP_A.K_1831275_11_10615343 | 11 | 10615343 | 2.43E-05 | 0.13211 | 299 | 0.137120895 | 0.021202411 |
| SNP_A.K_1004026_6_6820085 | 6 | 6820085 | 2.43E-05 | 0.3495 | 299 | 0.137108784 | 0.021226543 |
| SNP_A.K_978795_6_1782192 | 6 | 1782192 | 2.43E-05 | 0.2893 | 299 | 0.137108408 | 0.021226543 |
| SNP_A.K_307107_2_7379755 | 2 | 7379755 | 2.44E-05 | 0.31104 | 299 | 0.137098198 | 0.021251964 |
| SNP_A.K_201151_1_32138464 | 1 | 32138464 | 2.44E-05 | 0.41304 | 299 | 0.137094004 | 0.021252365 |
| SNP_A.K_1000984_6_6433493 | 6 | 6433493 | 2.44E-05 | 0.34114 | 299 | 0.137092785 | 0.021252365 |
| SNP_A.K_1004962_6_6958745 | 6 | 6958745 | 2.44E-05 | 0.301 | 299 | 0.137081255 | 0.021282275 |
| SNP_A.K_1608584_9_20790624 | 9 | 20790624 | 2.45E-05 | 0.25251 | 299 | 0.137066965 | 0.021321541 |
| SNP_A.K_1600927_9_18862219 | 9 | 18862219 | 2.45E-05 | 0.14716 | 299 | 0.137063809 | 0.021323272 |
| SNP_A.K_977983_6_1629391 | 6 | 1629391 | 2.48E-05 | 0.34783 | 299 | 0.136997516 | 0.021533465 |
| SNP_A.K_977920_6_1621853 | 6 | 1621853 | 2.48E-05 | 0.1806 | 299 | 0.136995925 | 0.021533465 |
| SNP_A.K_1295469_7_23350404 | 7 | 23350404 | 2.48E-05 | 0.02174 | 299 | 0.1369922 | 0.021533465 |
| SNP_A.K_2078813_12_27025327 | 12 | 27025327 | 2.48E-05 | 0.01338 | 299 | 0.136991284 | 0.021533465 |

| | | | | | | | |
|---|---|---|---|---|---|---|---|
| SNP_A.K_1002100_6_6540944 | 6 | 6540944 | 2.48E-05 | 0.31605 | 299 | 0.136977164 | 0.021572647 |
| SNP_A.K_708869_4_8120145 | 4 | 8120145 | 2.49E-05 | 0.08361 | 299 | 0.13695015 | 0.021656069 |
| SNP_A.K_709864_4_8299780 | 4 | 8299780 | 2.50E-05 | 0.07191 | 299 | 0.136946371 | 0.021659993 |
| SNP_A.K_673451_4_362839 | 4 | 362839 | 2.50E-05 | 0.13378 | 299 | 0.136943494 | 0.021660828 |
| SNP_A.K_1600872_9_18856877 | 9 | 18856877 | 2.51E-05 | 0.13712 | 299 | 0.136921581 | 0.021727053 |
| SNP_A.K_1600800_9_18841897 | 9 | 18841897 | 2.51E-05 | 0.13378 | 299 | 0.13691348 | 0.021736572 |
| SNP_A.K_992674_6_5671450 | 6 | 5671450 | 2.51E-05 | 0.1806 | 299 | 0.136913086 | 0.021736572 |
| SNP_A.K_1371565_8_7853041 | 8 | 7853041 | 2.51E-05 | 0.01839 | 299 | 0.136908845 | 0.021736572 |
| SNP_A.K_978025_6_1640482 | 6 | 1640482 | 2.51E-05 | 0.26254 | 299 | 0.136907847 | 0.021736572 |
| SNP_A.K_992464_6_5651782 | 6 | 5651782 | 2.51E-05 | 0.20234 | 299 | 0.13690567 | 0.021736572 |
| SNP_A.K_1371571_8_7854031 | 8 | 7854031 | 2.51E-05 | 0.02508 | 299 | 0.13690102 | 0.02174354 |
| SNP_A.K_709677_4_8280706 | 4 | 8280706 | 2.52E-05 | 0.09866 | 299 | 0.136881121 | 0.021803097 |
| SNP_A.K_996297_6_5981792 | 6 | 5981792 | 2.53E-05 | 0.2709 | 299 | 0.136865485 | 0.02184071 |
| SNP_A.K_307459_2_7408671 | 2 | 7408671 | 2.53E-05 | 0.25585 | 299 | 0.13686499 | 0.02184071 |
| SNP_A.K_709755_4_8284891 | 4 | 8284891 | 2.53E-05 | 0.08696 | 299 | 0.136858322 | 0.02185471 |
| SNP_A.K_1001331_6_6466073 | 6 | 6466073 | 2.53E-05 | 0.29097 | 299 | 0.136855459 | 0.021855551 |
| SNP_A.K_708363_4_8040750 | 4 | 8040750 | 2.53E-05 | 0.08696 | 299 | 0.13684883 | 0.021869434 |
| SNP_A.K_996203_6_5974871 | 6 | 5974871 | 2.54E-05 | 0.31271 | 299 | 0.136825269 | 0.021942085 |
| SNP_A.K_709503_4_8270885 | 4 | 8270885 | 2.55E-05 | 0.12207 | 299 | 0.136813366 | 0.021974377 |
| SNP_A.K_1001705_6_6507492 | 6 | 6507492 | 2.55E-05 | 0.29097 | 299 | 0.136798735 | 0.022016228 |
| SNP_A.K_1001082_6_6439998 | 6 | 6439998 | 2.56E-05 | 0.38294 | 299 | 0.136782931 | 0.022062267 |
| SNP_A.K_978184_6_1658453 | 6 | 1658453 | 2.56E-05 | 0.1689 | 299 | 0.136772817 | 0.022088488 |
| SNP_A.K_978663_6_1751722 | 6 | 1751722 | 2.57E-05 | 0.29264 | 299 | 0.136757313 | 0.022133637 |
| SNP_A.K_990274_6_4704175 | 6 | 4704175 | 2.57E-05 | 0.21237 | 299 | 0.136751479 | 0.022144938 |
| SNP_A.K_307228_2_7386440 | 2 | 7386440 | 2.58E-05 | 0.45987 | 299 | 0.136744386 | 0.022160666 |
| SNP_A.K_977900_6_1617418 | 6 | 1617418 | 2.58E-05 | 0.24247 | 299 | 0.136738411 | 0.022169637 |
| SNP_A.K_978312_6_1673229 | 6 | 1673229 | 2.58E-05 | 0.18729 | 299 | 0.136736615 | 0.022169637 |
| SNP_A.K_306877_2_7363185 | 2 | 7363185 | 2.58E-05 | 0.26756 | 299 | 0.136728505 | 0.02218897 |
| SNP_A.K_1383139_8_9131826 | 8 | 9131826 | 2.59E-05 | 0.27759 | 299 | 0.136706668 | 0.022256665 |
| SNP_A.K_1001097_6_6441285 | 6 | 6441285 | 2.59E-05 | 0.31104 | 299 | 0.136703935 | 0.022257117 |
| SNP_A.K_978435_6_1693320 | 6 | 1693320 | 2.60E-05 | 0.26421 | 299 | 0.136684356 | 0.022317046 |
| SNP_A.K_1389976_8_10140629 | 8 | 10140629 | 2.61E-05 | 0.05017 | 299 | 0.136670199 | 0.022357932 |
| SNP_A.K_1001306_6_6463514 | 6 | 6463514 | 2.61E-05 | 0.29933 | 299 | 0.136663169 | 0.022373624 |
| SNP_A.K_708896_4_8122472 | 4 | 8122472 | 2.61E-05 | 0.08696 | 299 | 0.136654773 | 0.022394176 |
| SNP_A.K_726170_4_11925305 | 4 | 11925305 | 2.62E-05 | 0.09699 | 299 | 0.136646823 | 0.022413169 |
| SNP_A.K_77686_1_10887934 | 1 | 10887934 | 2.62E-05 | 0.29599 | 299 | 0.136643649 | 0.022415214 |
| SNP_A.K_2063301_12_23966670 | 12 | 23966670 | 2.62E-05 | 0.02676 | 299 | 0.136634136 | 0.02243979 |
| SNP_A.K_1001520_6_6486754 | 6 | 6486754 | 2.63E-05 | 0.29431 | 299 | 0.13662272 | 0.022471166 |
| SNP_A.K_998067_6_6168370 | 6 | 6168370 | 2.63E-05 | 0.16054 | 299 | 0.136613806 | 0.022493675 |
| SNP_A.K_1432395_8_16491078 | 8 | 16491078 | 2.64E-05 | 0.07692 | 299 | 0.136601498 | 0.022505403 |
| SNP_A.K_1432342_8_16485849 | 8 | 16485849 | 2.64E-05 | 0.06522 | 299 | 0.136601316 | 0.022505403 |
| SNP_A.K_1001300_6_6463217 | 6 | 6463217 | 2.64E-05 | 0.29766 | 299 | 0.13660048 | 0.022505403 |
| SNP_A.K_1000994_6_6434753 | 6 | 6434753 | 2.64E-05 | 0.34281 | 299 | 0.136600151 | 0.022505403 |
| SNP_A.K_326938_2_10071519 | 2 | 10071519 | 2.64E-05 | 0.06522 | 299 | 0.136587224 | 0.022542292 |
| SNP_A.K_978674_6_1755963 | 6 | 1755963 | 2.65E-05 | 0.28261 | 299 | 0.136578045 | 0.022565844 |
| SNP_A.K_990311_6_4710030 | 6 | 4710030 | 2.65E-05 | 0.44147 | 299 | 0.136572618 | 0.022576001 |
| SNP_A.K_1390369_8_10193189 | 8 | 10193189 | 2.66E-05 | 0.05518 | 299 | 0.136548118 | 0.022651371 |
| SNP_A.K_990265_6_4703786 | 6 | 4703786 | 2.66E-05 | 0.17726 | 299 | 0.136546412 | 0.022651371 |
| SNP_A.K_218589_1_35319613 | 1 | 35319613 | 2.66E-05 | 0.19231 | 299 | 0.136541049 | 0.022661349 |
| SNP_A.K_325359_2_9874961 | 2 | 9874961 | 2.67E-05 | 0.02508 | 299 | 0.136529795 | 0.02268382 |

| | | | | | | | |
|---|---|---|---|---|---|---|---|
| SNP_A.K_990410_6_4716244 | 6 | 4716244 | 2.67E-05 | 0.1505 | 299 | 0.136527919 | 0.02268382 |
| SNP_A.K_307410_2_7401717 | 2 | 7401717 | 2.67E-05 | 0.25251 | 299 | 0.136527051 | 0.02268382 |
| SNP_A.K_254769_1_41088166 | 1 | 41088166 | 2.67E-05 | 0.1505 | 299 | 0.136522043 | 0.022692553 |
| SNP_A.K_1608738_9_20829099 | 9 | 20829099 | 2.67E-05 | 0.3913 | 299 | 0.136512106 | 0.022709896 |
| SNP_A.K_1004282_6_6862485 | 6 | 6862485 | 2.67E-05 | 0.34114 | 299 | 0.136512067 | 0.022709896 |
| SNP_A.K_998062_6_6168082 | 6 | 6168082 | 2.68E-05 | 0.18562 | 299 | 0.136502183 | 0.022736208 |
| SNP_A.K_1001991_6_6534115 | 6 | 6534115 | 2.68E-05 | 0.33612 | 299 | 0.136497229 | 0.022744783 |
| SNP_A.K_312201_2_7922869 | 2 | 7922869 | 2.68E-05 | 0.27759 | 299 | 0.136491656 | 0.022755599 |
| SNP_A.K_709752_4_8284859 | 4 | 8284859 | 2.69E-05 | 0.08696 | 299 | 0.136477763 | 0.022796454 |
| SNP_A.K_1608574_9_20787870 | 9 | 20787870 | 2.70E-05 | 0.24916 | 299 | 0.136451083 | 0.022883709 |
| SNP_A.K_1001695_6_6504275 | 6 | 6504275 | 2.70E-05 | 0.29264 | 299 | 0.136443502 | 0.022901889 |
| SNP_A.K_978705_6_1769155 | 6 | 1769155 | 2.71E-05 | 0.36455 | 299 | 0.136432706 | 0.022931764 |
| SNP_A.K_996276_6_5981139 | 6 | 5981139 | 2.71E-05 | 0.27425 | 299 | 0.136420812 | 0.022964827 |
| SNP_A.K_1001309_6_6463852 | 6 | 6463852 | 2.71E-05 | 0.301 | 299 | 0.13641848 | 0.022964827 |
| SNP_A.K_312097_2_7907094 | 2 | 7907094 | 2.72E-05 | 0.26756 | 299 | 0.136410587 | 0.022984224 |
| SNP_A.K_709971_4_8303867 | 4 | 8303867 | 2.72E-05 | 0.07191 | 299 | 0.136405374 | 0.022993871 |
| SNP_A.K_196596_1_31435797 | 1 | 31435797 | 2.72E-05 | 0.47826 | 299 | 0.136397584 | 0.022999825 |
| SNP_A.K_710384_4_8339426 | 4 | 8339426 | 2.72E-05 | 0.08528 | 299 | 0.136396278 | 0.022999825 |
| SNP_A.K_1001760_6_6515087 | 6 | 6515087 | 2.72E-05 | 0.31271 | 299 | 0.136396053 | 0.022999825 |
| SNP_A.K_306997_2_7372458 | 2 | 7372458 | 2.73E-05 | 0.25585 | 299 | 0.136386361 | 0.023025834 |
| SNP_A.K_992030_6_5613461 | 6 | 5613461 | 2.73E-05 | 0.20569 | 299 | 0.136379696 | 0.023038469 |
| SNP_A.K_709817_4_8298364 | 4 | 8298364 | 2.73E-05 | 0.07191 | 299 | 0.136375225 | 0.023038469 |
| SNP_A.K_709816_4_8298357 | 4 | 8298357 | 2.73E-05 | 0.07191 | 299 | 0.136375225 | 0.023038469 |
| SNP_A.K_1001355_6_6466652 | 6 | 6466652 | 2.73E-05 | 0.29599 | 299 | 0.136367789 | 0.023056296 |
| SNP_A.K_456442_2_33037031 | 2 | 33037031 | 2.74E-05 | 0.07358 | 299 | 0.136360167 | 0.023069456 |
| SNP_A.K_326712_2_10041616 | 2 | 10041616 | 2.74E-05 | 0.06689 | 299 | 0.136357362 | 0.023069456 |
| SNP_A.K_1001685_6_6503725 | 6 | 6503725 | 2.74E-05 | 0.29264 | 299 | 0.136355139 | 0.023069456 |
| SNP_A.K_990376_6_4712704 | 6 | 4712704 | 2.74E-05 | 0.19565 | 299 | 0.136353974 | 0.023069456 |
| SNP_A.K_2078834_12_27026248 | 12 | 27026248 | 2.74E-05 | 0.01505 | 299 | 0.136351059 | 0.023070789 |
| SNP_A.K_307017_2_7373145 | 2 | 7373145 | 2.76E-05 | 0.23913 | 299 | 0.136320895 | 0.023168671 |
| SNP_A.K_1001257_6_6459463 | 6 | 6459463 | 2.76E-05 | 0.28428 | 299 | 0.136319249 | 0.023168671 |
| SNP_A.K_710493_4_8424912 | 4 | 8424912 | 2.76E-05 | 0.07191 | 299 | 0.136310152 | 0.023181543 |
| SNP_A.K_978878_6_1793998 | 6 | 1793998 | 2.76E-05 | 0.31271 | 299 | 0.136309827 | 0.023181543 |
| SNP_A.K_995531_6_5940386 | 6 | 5940386 | 2.76E-05 | 0.43813 | 299 | 0.136308105 | 0.023181543 |
| SNP_A.K_978145_6_1653942 | 6 | 1653942 | 2.77E-05 | 0.25753 | 299 | 0.136280852 | 0.023267417 |
| SNP_A.K_978613_6_1742656 | 6 | 1742656 | 2.77E-05 | 0.27926 | 299 | 0.13627969 | 0.023267417 |
| SNP_A.K_1526094_9_5007108 | 9 | 5007108 | 2.78E-05 | 0.01839 | 299 | 0.136271559 | 0.023275043 |
| SNP_A.K_1005042_6_6964425 | 6 | 6964425 | 2.78E-05 | 0.301 | 299 | 0.136271109 | 0.023275043 |
| SNP_A.K_307452_2_7408551 | 2 | 7408551 | 2.78E-05 | 0.2408 | 299 | 0.136269999 | 0.023275043 |
| SNP_A.K_1830608_11_10569818 | 11 | 10569818 | 2.78E-05 | 0.43478 | 299 | 0.136262986 | 0.023291557 |
| SNP_A.K_978289_6_1669545 | 6 | 1669545 | 2.80E-05 | 0.24582 | 299 | 0.136222174 | 0.023433301 |
| SNP_A.K_978836_6_1787195 | 6 | 1787195 | 2.80E-05 | 0.41806 | 299 | 0.136218715 | 0.023436727 |
| SNP_A.K_1830860_11_10585376 | 11 | 10585376 | 2.80E-05 | 0.15719 | 299 | 0.136210482 | 0.023457904 |
| SNP_A.K_999389_6_6286152 | 6 | 6286152 | 2.81E-05 | 0.16221 | 299 | 0.136207245 | 0.023460515 |
| SNP_A.K_1001304_6_6463372 | 6 | 6463372 | 2.81E-05 | 0.30435 | 299 | 0.136192367 | 0.023506472 |
| SNP_A.K_2079159_12_27061427 | 12 | 27061427 | 2.81E-05 | 0.01672 | 299 | 0.136189041 | 0.023509428 |
| SNP_A.K_993662_6_5782161 | 6 | 5782161 | 2.82E-05 | 0.27926 | 299 | 0.136175377 | 0.023550956 |
| SNP_A.K_1001369_6_6470488 | 6 | 6470488 | 2.82E-05 | 0.2893 | 299 | 0.136169382 | 0.023563896 |
| SNP_A.K_325804_2_9944617 | 2 | 9944617 | 2.82E-05 | 0.06689 | 299 | 0.136166573 | 0.023564942 |
| SNP_A.K_990367_6_4712578 | 6 | 4712578 | 2.83E-05 | 0.19565 | 299 | 0.136161871 | 0.023573063 |

| | | | | | | | |
|---|---|---|---|---|---|---|---|
| SNP_A.K_1608490_9_20763555 | 9 | 20763555 | 2.83E-05 | 0.30769 | 299 | 0.136156387 | 0.023584114 |
| SNP_A.K_1608307_9_20667615 | 9 | 20667615 | 2.83E-05 | 0.14214 | 299 | 0.136152014 | 0.023589404 |
| SNP_A.K_308496_2_7521137 | 2 | 7521137 | 2.83E-05 | 0.24582 | 299 | 0.136149921 | 0.023589404 |
| SNP_A.K_709585_4_8274829 | 4 | 8274829 | 2.85E-05 | 0.08361 | 299 | 0.136114819 | 0.023711565 |
| SNP_A.K_1001116_6_6447008 | 6 | 6447008 | 2.85E-05 | 0.38629 | 299 | 0.136104418 | 0.023736254 |
| SNP_A.K_678575_4_990318 | 4 | 990318 | 2.85E-05 | 0.07358 | 299 | 0.136103211 | 0.023736254 |
| SNP_A.K_725739_4_11838620 | 4 | 11838620 | 2.85E-05 | 0.10368 | 299 | 0.136099535 | 0.0237406 |
| SNP_A.K_306790_2_7350886 | 2 | 7350886 | 2.86E-05 | 0.26087 | 299 | 0.136095186 | 0.02374376 |
| SNP_A.K_999886_6_6331605 | 6 | 6331605 | 2.86E-05 | 0.3913 | 299 | 0.136093656 | 0.02374376 |
| SNP_A.K_709830_4_8299133 | 4 | 8299133 | 2.86E-05 | 0.07358 | 299 | 0.136083267 | 0.023765492 |
| SNP_A.K_725661_4_11829199 | 4 | 11829199 | 2.86E-05 | 0.09699 | 299 | 0.136082852 | 0.023765492 |
| SNP_A.K_1608650_9_20810420 | 9 | 20810420 | 2.86E-05 | 0.30936 | 299 | 0.136075263 | 0.023784616 |
| SNP_A.K_710339_4_8336880 | 4 | 8336880 | 2.87E-05 | 0.07358 | 299 | 0.136072214 | 0.023786628 |
| SNP_A.K_1371599_8_7858082 | 8 | 7858082 | 2.87E-05 | 0.01839 | 299 | 0.136061928 | 0.023815965 |
| SNP_A.K_1004047_6_6821588 | 6 | 6821588 | 2.87E-05 | 0.33278 | 299 | 0.136058072 | 0.023820831 |
| SNP_A.K_1004351_6_6874789 | 6 | 6874789 | 2.87E-05 | 0.38629 | 299 | 0.136055615 | 0.023820831 |
| SNP_A.K_1001342_6_6466271 | 6 | 6466271 | 2.88E-05 | 0.29933 | 299 | 0.136047693 | 0.023841279 |
| SNP_A.K_1600694_9_18822060 | 9 | 18822060 | 2.88E-05 | 0.14214 | 299 | 0.136039605 | 0.023861704 |
| SNP_A.K_307052_2_7376096 | 2 | 7376096 | 2.88E-05 | 0.22742 | 299 | 0.136037274 | 0.023861704 |
| SNP_A.K_191198_1_30224468 | 1 | 30224468 | 2.89E-05 | 0.07023 | 299 | 0.136023781 | 0.02390331 |
| SNP_A.K_253109_1_40858909 | 1 | 40858909 | 2.89E-05 | 0.15886 | 299 | 0.136016202 | 0.023922544 |
| SNP_A.K_990234_6_4701775 | 6 | 4701775 | 2.90E-05 | 0.19565 | 299 | 0.136000112 | 0.023964512 |
| SNP_A.K_992530_6_5655041 | 6 | 5655041 | 2.90E-05 | 0.21405 | 299 | 0.136000021 | 0.023964512 |
| SNP_A.K_1123686_6_23551707 | 6 | 23551707 | 2.90E-05 | 0.01338 | 299 | 0.135997636 | 0.023964512 |
| SNP_A.K_673421_4_359064 | 4 | 359064 | 2.90E-05 | 0.1505 | 299 | 0.135993846 | 0.023969403 |
| SNP_A.K_710489_4_8424747 | 4 | 8424747 | 2.91E-05 | 0.07191 | 299 | 0.135978879 | 0.024016836 |
| SNP_A.K_307441_2_7407663 | 2 | 7407663 | 2.92E-05 | 0.301 | 299 | 0.135961861 | 0.024072202 |
| SNP_A.K_990382_6_4712830 | 6 | 4712830 | 2.92E-05 | 0.19398 | 299 | 0.135951707 | 0.024101442 |
| SNP_A.K_978944_6_1816748 | 6 | 1816748 | 2.92E-05 | 0.3194 | 299 | 0.135947596 | 0.024106506 |
| SNP_A.K_1600921_9_18862120 | 9 | 18862120 | 2.92E-05 | 0.15552 | 299 | 0.135945387 | 0.024106506 |
| SNP_A.K_995309_6_5925001 | 6 | 5925001 | 2.93E-05 | 0.14381 | 299 | 0.135938106 | 0.024120661 |
| SNP_A.K_1004522_6_6896526 | 6 | 6896526 | 2.93E-05 | 0.40803 | 299 | 0.135936694 | 0.024120661 |
| SNP_A.K_1001314_6_6464707 | 6 | 6464707 | 2.94E-05 | 0.28261 | 299 | 0.135919739 | 0.024176046 |
| SNP_A.K_1000980_6_6432165 | 6 | 6432165 | 2.94E-05 | 0.34281 | 299 | 0.135910775 | 0.024200874 |
| SNP_A.K_307988_2_7461440 | 2 | 7461440 | 2.94E-05 | 0.47659 | 299 | 0.135904919 | 0.024208718 |
| SNP_A.K_1001512_6_6485964 | 6 | 6485964 | 2.94E-05 | 0.29097 | 299 | 0.135903748 | 0.024208718 |
| SNP_A.K_990349_6_4711884 | 6 | 4711884 | 2.95E-05 | 0.199 | 299 | 0.135896728 | 0.024226118 |
| SNP_A.K_726190_4_11927244 | 4 | 11927244 | 2.96E-05 | 0.09699 | 299 | 0.135878723 | 0.024285816 |
| SNP_A.K_978600_6_1738285 | 6 | 1738285 | 2.96E-05 | 0.24582 | 299 | 0.135872185 | 0.024301421 |
| SNP_A.K_978597_6_1737366 | 6 | 1737366 | 2.96E-05 | 0.24916 | 299 | 0.135864932 | 0.0243198 |
| SNP_A.K_1196448_7_7613481 | 7 | 7613481 | 2.97E-05 | 0.01003 | 299 | 0.135853202 | 0.024347668 |
| SNP_A.K_978754_6_1775148 | 6 | 1775148 | 2.97E-05 | 0.36455 | 299 | 0.135852741 | 0.024347668 |
| SNP_A.K_308483_2_7520041 | 2 | 7520041 | 2.97E-05 | 0.23913 | 299 | 0.135846043 | 0.02436395 |
| SNP_A.K_1000993_6_6434624 | 6 | 6434624 | 2.98E-05 | 0.34615 | 299 | 0.135819461 | 0.024450678 |
| SNP_A.K_725228_4_11763666 | 4 | 11763666 | 2.98E-05 | 0.09866 | 299 | 0.135818678 | 0.024450678 |
| SNP_A.K_709123_4_8230845 | 4 | 8230845 | 2.99E-05 | 0.08528 | 299 | 0.135808741 | 0.024465186 |
| SNP_A.K_1001805_6_6522044 | 6 | 6522044 | 2.99E-05 | 0.29264 | 299 | 0.135807678 | 0.024465186 |
| SNP_A.K_1001234_6_6456767 | 6 | 6456767 | 2.99E-05 | 0.38963 | 299 | 0.135807499 | 0.024465186 |
| SNP_A.K_710363_4_8338696 | 4 | 8338696 | 2.99E-05 | 0.09197 | 299 | 0.135798891 | 0.024488991 |
| SNP_A.K_1004409_6_6880031 | 6 | 6880031 | 3.00E-05 | 0.43144 | 299 | 0.135785705 | 0.024530632 |

| | | | | | | | |
|---|---|---|---|---|---|---|---|
| SNP_A.K_1001809_6_6522189 | 6 | 6522189 | 3.00E-05 | 0.32274 | 299 | 0.135782808 | 0.02453227 |
| SNP_A.K_978686_6_1761272 | 6 | 1761272 | 3.00E-05 | 0.38629 | 299 | 0.13577535 | 0.024551672 |
| SNP_A.K_201239_1_32154607 | 1 | 32154607 | 3.01E-05 | 0.42308 | 299 | 0.135762809 | 0.024590914 |
| SNP_A.K_195003_1_31033495 | 1 | 31033495 | 3.01E-05 | 0.41304 | 299 | 0.135757329 | 0.02460265 |
| SNP_A.K_901127_5_16261519 | 5 | 16261519 | 3.01E-05 | 0.04348 | 299 | 0.1357528 | 0.024610682 |
| SNP_A.K_709840_4_8299280 | 4 | 8299280 | 3.02E-05 | 0.07525 | 299 | 0.135740847 | 0.024647735 |
| SNP_A.K_710409_4_8348757 | 4 | 8348757 | 3.02E-05 | 0.12709 | 299 | 0.135735789 | 0.024657858 |
| SNP_A.K_1001514_6_6486150 | 6 | 6486150 | 3.02E-05 | 0.29766 | 299 | 0.135732586 | 0.02466073 |
| SNP_A.K_963849_5_28325523 | 5 | 28325523 | 3.03E-05 | 0.03177 | 299 | 0.135729498 | 0.024663153 |
| SNP_A.K_312078_2_7904374 | 2 | 7904374 | 3.03E-05 | 0.2709 | 299 | 0.135720923 | 0.02468707 |
| SNP_A.K_1001470_6_6482800 | 6 | 6482800 | 3.03E-05 | 0.29766 | 299 | 0.135715281 | 0.024699511 |
| SNP_A.K_708510_4_8059196 | 4 | 8059196 | 3.04E-05 | 0.10201 | 299 | 0.135707202 | 0.024721522 |
| SNP_A.K_990273_6_4704167 | 6 | 4704167 | 3.04E-05 | 0.21237 | 299 | 0.135697622 | 0.024746885 |
| SNP_A.K_307252_2_7388042 | 2 | 7388042 | 3.04E-05 | 0.25753 | 299 | 0.135695813 | 0.024746885 |
| SNP_A.K_1000947_6_6425850 | 6 | 6425850 | 3.04E-05 | 0.29766 | 299 | 0.135692992 | 0.024748294 |
| SNP_A.K_709884_4_8300266 | 4 | 8300266 | 3.05E-05 | 0.07358 | 299 | 0.135687647 | 0.024759618 |
| SNP_A.K_996325_6_5986021 | 6 | 5986021 | 3.06E-05 | 0.27759 | 299 | 0.135661474 | 0.024852965 |
| SNP_A.K_1431609_8_16396015 | 8 | 16396015 | 3.07E-05 | 0.07692 | 299 | 0.135646897 | 0.024900802 |
| SNP_A.K_1004378_6_6877859 | 6 | 6877859 | 3.07E-05 | 0.37458 | 299 | 0.135640557 | 0.024916148 |
| SNP_A.K_1830634_11_10571660 | 11 | 10571660 | 3.07E-05 | 0.1689 | 299 | 0.135636631 | 0.024921953 |
| SNP_A.K_1004430_6_6881755 | 6 | 6881755 | 3.07E-05 | 0.31773 | 299 | 0.135632821 | 0.024927312 |
| SNP_A.K_1500264_9_693467 | 9 | 693467 | 3.08E-05 | 0.04515 | 299 | 0.135622891 | 0.024956896 |
| SNP_A.K_996596_6_6011384 | 6 | 6011384 | 3.08E-05 | 0.47659 | 299 | 0.135619864 | 0.024959164 |
| SNP_A.K_978024_6_1640478 | 6 | 1640478 | 3.09E-05 | 0.26087 | 299 | 0.135601716 | 0.025021412 |
| SNP_A.K_306765_2_7348681 | 2 | 7348681 | 3.09E-05 | 0.2709 | 299 | 0.135596071 | 0.025033533 |
| SNP_A.K_709849_4_8299546 | 4 | 8299546 | 3.09E-05 | 0.07358 | 299 | 0.135591319 | 0.025033533 |
| SNP_A.K_752442_4_17906421 | 4 | 17906421 | 3.09E-05 | 0.06355 | 299 | 0.135591311 | 0.025033533 |
| SNP_A.K_990233_6_4701766 | 6 | 4701766 | 3.11E-05 | 0.19732 | 299 | 0.135565737 | 0.025125579 |
| SNP_A.K_1005354_6_7008884 | 6 | 7008884 | 3.12E-05 | 0.34783 | 299 | 0.135545876 | 0.025188754 |
| SNP_A.K_709819_4_8298430 | 4 | 8298430 | 3.12E-05 | 0.07358 | 299 | 0.13554502 | 0.025188754 |
| SNP_A.K_1432909_8_16600919 | 8 | 16600919 | 3.12E-05 | 0.1388 | 299 | 0.13553416 | 0.025222417 |
| SNP_A.K_710127_4_8317606 | 4 | 8317606 | 3.12E-05 | 0.07191 | 299 | 0.135527987 | 0.025234912 |
| SNP_A.K_307053_2_7376127 | 2 | 7376127 | 3.13E-05 | 0.2408 | 299 | 0.13552615 | 0.025234912 |
| SNP_A.K_709783_4_8297645 | 4 | 8297645 | 3.13E-05 | 0.07191 | 299 | 0.135520031 | 0.02524649 |
| SNP_A.K_978598_6_1737830 | 6 | 1737830 | 3.13E-05 | 0.24415 | 299 | 0.135518375 | 0.02524649 |
| SNP_A.K_1830625_11_10571305 | 11 | 10571305 | 3.13E-05 | 0.36455 | 299 | 0.135507395 | 0.025280729 |
| SNP_A.K_312488_2_7965690 | 2 | 7965690 | 3.14E-05 | 0.18729 | 299 | 0.13550033 | 0.025299295 |
| SNP_A.K_1001325_6_6465280 | 6 | 6465280 | 3.14E-05 | 0.301 | 299 | 0.135495772 | 0.025307803 |
| SNP_A.K_997999_6_6165543 | 6 | 6165543 | 3.15E-05 | 0.39632 | 299 | 0.135479544 | 0.025363263 |
| SNP_A.K_1001166_6_6450433 | 6 | 6450433 | 3.15E-05 | 0.28261 | 299 | 0.135467512 | 0.025401922 |
| SNP_A.K_978841_6_1787283 | 6 | 1787283 | 3.16E-05 | 0.35786 | 299 | 0.135464023 | 0.025406167 |
| SNP_A.K_1600813_9_18843542 | 9 | 18843542 | 3.16E-05 | 0.13211 | 299 | 0.135456627 | 0.025426184 |
| SNP_A.K_990272_6_4704113 | 6 | 4704113 | 3.16E-05 | 0.22241 | 299 | 0.135449913 | 0.025443464 |
| SNP_A.K_307345_2_7398163 | 2 | 7398163 | 3.17E-05 | 0.42475 | 299 | 0.135444013 | 0.025457467 |
| SNP_A.K_978162_6_1656541 | 6 | 1656541 | 3.17E-05 | 0.26087 | 299 | 0.135438021 | 0.025466775 |
| SNP_A.K_1000747_6_6414441 | 6 | 6414441 | 3.17E-05 | 0.39298 | 299 | 0.135435817 | 0.025466775 |
| SNP_A.K_1002094_6_6540688 | 6 | 6540688 | 3.17E-05 | 0.31271 | 299 | 0.135434416 | 0.025466775 |
| SNP_A.K_307439_2_7407407 | 2 | 7407407 | 3.18E-05 | 0.23913 | 299 | 0.135422591 | 0.025494862 |
| SNP_A.K_1383174_8_9133384 | 8 | 9133384 | 3.18E-05 | 0.11037 | 299 | 0.135421765 | 0.025494862 |
| SNP_A.K_978763_6_1776123 | 6 | 1776123 | 3.18E-05 | 0.37793 | 299 | 0.135420186 | 0.025494862 |

| | | | | | | | |
|---|---|---|---|---|---|---|---|
| SNP_A.K_978018_6_1637636 | 6 | 1637636 | 3.18E-05 | 0.16221 | 299 | 0.135415406 | 0.02550439 |
| SNP_A.K_1925645_11_24752398 | 11 | 24752398 | 3.18E-05 | 0.04348 | 299 | 0.135409606 | 0.025518054 |
| SNP_A.K_978606_6_1740472 | 6 | 1740472 | 3.19E-05 | 0.26756 | 299 | 0.135401956 | 0.025539235 |
| SNP_A.K_923563_5_19487977 | 5 | 19487977 | 3.19E-05 | 0.03344 | 299 | 0.135399337 | 0.025540026 |
| SNP_A.K_1004835_6_6935110 | 6 | 6935110 | 3.19E-05 | 0.31773 | 299 | 0.135388356 | 0.025574748 |
| SNP_A.K_1831010_11_10595865 | 11 | 10595865 | 3.20E-05 | 0.14548 | 299 | 0.135383848 | 0.025578743 |
| SNP_A.K_978605_6_1740167 | 6 | 1740167 | 3.20E-05 | 0.25753 | 299 | 0.13538253 | 0.025578743 |
| SNP_A.K_978133_6_1649405 | 6 | 1649405 | 3.20E-05 | 0.29264 | 299 | 0.135370387 | 0.025618257 |
| SNP_A.K_1432938_8_16606416 | 8 | 16606416 | 3.21E-05 | 0.11538 | 299 | 0.135362341 | 0.025629555 |
| SNP_A.K_307350_2_7398349 | 2 | 7398349 | 3.21E-05 | 0.24247 | 299 | 0.13536185 | 0.025629555 |
| SNP_A.K_993998_6_5807106 | 6 | 5807106 | 3.21E-05 | 0.19732 | 299 | 0.135360356 | 0.025629555 |
| SNP_A.K_1608533_9_20772772 | 9 | 20772772 | 3.21E-05 | 0.30769 | 299 | 0.13535718 | 0.025632644 |
| SNP_A.K_1550753_9_9008975 | 9 | 9008975 | 3.21E-05 | 0.11037 | 299 | 0.135351477 | 0.02563729 |
| SNP_A.K_306794_2_7351554 | 2 | 7351554 | 3.21E-05 | 0.26087 | 299 | 0.135351209 | 0.02563729 |
| SNP_A.K_978661_6_1751603 | 6 | 1751603 | 3.23E-05 | 0.28428 | 299 | 0.135318878 | 0.025759367 |
| SNP_A.K_709501_4_8270845 | 4 | 8270845 | 3.23E-05 | 0.10033 | 299 | 0.135315946 | 0.025761493 |
| SNP_A.K_923866_5_19510547 | 5 | 19510547 | 3.24E-05 | 0.02341 | 299 | 0.135304886 | 0.025796896 |
| SNP_A.K_195081_1_31042288 | 1 | 31042288 | 3.25E-05 | 0.4097 | 299 | 0.135288094 | 0.025855877 |
| SNP_A.K_1608556_9_20777207 | 9 | 20777207 | 3.25E-05 | 0.2709 | 299 | 0.135281054 | 0.025874895 |
| SNP_A.K_709943_4_8302612 | 4 | 8302612 | 3.25E-05 | 0.07358 | 299 | 0.135275841 | 0.025886422 |
| SNP_A.K_307023_2_7373310 | 2 | 7373310 | 3.26E-05 | 0.25084 | 299 | 0.135268203 | 0.025907932 |
| SNP_A.K_978046_6_1642528 | 6 | 1642528 | 3.26E-05 | 0.16555 | 299 | 0.135254999 | 0.025952394 |
| SNP_A.K_1000979_6_6431705 | 6 | 6431705 | 3.27E-05 | 0.33946 | 299 | 0.135249106 | 0.025966769 |
| SNP_A.K_1600915_9_18861734 | 9 | 18861734 | 3.27E-05 | 0.12876 | 299 | 0.135243365 | 0.025980532 |
| SNP_A.K_326858_2_10061504 | 2 | 10061504 | 3.29E-05 | 0.06856 | 299 | 0.135210995 | 0.026104471 |
| SNP_A.K_1004897_6_6949731 | 6 | 6949731 | 3.29E-05 | 0.15385 | 299 | 0.135206502 | 0.026113138 |
| SNP_A.K_709814_4_8298320 | 4 | 8298320 | 3.30E-05 | 0.07358 | 299 | 0.135192863 | 0.026159784 |
| SNP_A.K_978889_6_1796800 | 6 | 1796800 | 3.30E-05 | 0.33612 | 299 | 0.135190327 | 0.026160346 |
| SNP_A.K_709372_4_8259282 | 4 | 8259282 | 3.30E-05 | 0.0786 | 299 | 0.135182471 | 0.026183025 |
| SNP_A.K_306717_2_7344672 | 2 | 7344672 | 3.30E-05 | 0.301 | 299 | 0.135178883 | 0.02618797 |
| SNP_A.K_673423_4_360463 | 4 | 360463 | 3.30E-05 | 0.13211 | 299 | 0.135175725 | 0.026191134 |
| SNP_A.K_218762_1_35347467 | 1 | 35347467 | 3.31E-05 | 0.16722 | 299 | 0.135166942 | 0.026211585 |
| SNP_A.K_307425_2_7403811 | 2 | 7403811 | 3.31E-05 | 0.24916 | 299 | 0.135166017 | 0.026211585 |
| SNP_A.K_1582285_9_15055495 | 9 | 15055495 | 3.32E-05 | 0.07358 | 299 | 0.135149812 | 0.026269143 |
| SNP_A.K_978466_6_1696748 | 6 | 1696748 | 3.32E-05 | 0.32107 | 299 | 0.135142716 | 0.026279262 |
| SNP_A.K_2078873_12_27029165 | 12 | 27029165 | 3.32E-05 | 0.01672 | 299 | 0.1351426 | 0.026279262 |
| SNP_A.K_1389969_8_10139556 | 8 | 10139556 | 3.33E-05 | 0.05184 | 299 | 0.135133473 | 0.02630739 |
| SNP_A.K_994952_6_5907602 | 6 | 5907602 | 3.33E-05 | 0.14716 | 299 | 0.135127689 | 0.026318753 |
| SNP_A.K_978019_6_1637921 | 6 | 1637921 | 3.33E-05 | 0.15886 | 299 | 0.135125972 | 0.026318753 |
| SNP_A.K_1001985_6_6533545 | 6 | 6533545 | 3.34E-05 | 0.32107 | 299 | 0.135109625 | 0.026377169 |
| SNP_A.K_726244_4_11930570 | 4 | 11930570 | 3.35E-05 | 0.09699 | 299 | 0.135091283 | 0.026444099 |
| SNP_A.K_709826_4_8298777 | 4 | 8298777 | 3.35E-05 | 0.07191 | 299 | 0.135080404 | 0.026479795 |
| SNP_A.K_977953_6_1624236 | 6 | 1624236 | 3.36E-05 | 0.25251 | 299 | 0.135076215 | 0.026484103 |
| SNP_A.K_1005353_6_7008881 | 6 | 7008881 | 3.36E-05 | 0.35117 | 299 | 0.135074608 | 0.026484103 |
| SNP_A.K_1001802_6_6521640 | 6 | 6521640 | 3.36E-05 | 0.34615 | 299 | 0.13507005 | 0.026493246 |
| SNP_A.K_1001288_6_6461986 | 6 | 6461986 | 3.36E-05 | 0.28094 | 299 | 0.135067096 | 0.026495645 |
| SNP_A.K_307184_2_7383647 | 2 | 7383647 | 3.38E-05 | 0.42977 | 299 | 0.135037616 | 0.026609951 |
| SNP_A.K_923993_5_19533988 | 5 | 19533988 | 3.38E-05 | 0.0301 | 299 | 0.135027533 | 0.026642267 |
| SNP_A.K_990387_6_4713985 | 6 | 4713985 | 3.38E-05 | 0.19565 | 299 | 0.135024315 | 0.026642267 |
| SNP_A.K_1000672_6_6409420 | 6 | 6409420 | 3.39E-05 | 0.28763 | 299 | 0.135021133 | 0.026642267 |

| | | | | | | | |
|---|---|---|---|---|---|---|---|
| SNP_A.K_307460_2_7408685 | 2 | 7408685 | 3.39E-05 | 0.28094 | 299 | 0.135020454 | 0.026642267 |
| SNP_A.K_1004136_6_6836413 | 6 | 6836413 | 3.40E-05 | 0.36789 | 299 | 0.134995356 | 0.02673326 |
| SNP_A.K_1002053_6_6538464 | 6 | 6538464 | 3.40E-05 | 0.31773 | 299 | 0.134994238 | 0.02673326 |
| SNP_A.K_990385_6_4713382 | 6 | 4713382 | 3.41E-05 | 0.46154 | 299 | 0.134982995 | 0.026770948 |
| SNP_A.K_710477_4_8424078 | 4 | 8424078 | 3.41E-05 | 0.07692 | 299 | 0.134971136 | 0.026801344 |
| SNP_A.K_990373_6_4712654 | 6 | 4712654 | 3.41E-05 | 0.19565 | 299 | 0.134971103 | 0.026801344 |
| SNP_A.K_86876_1_11849418 | 1 | 11849418 | 3.43E-05 | 0.09532 | 299 | 0.134935842 | 0.026935396 |
| SNP_A.K_1001776_6_6520039 | 6 | 6520039 | 3.43E-05 | 0.29431 | 299 | 0.134934958 | 0.026935396 |
| SNP_A.K_194947_1_31004275 | 1 | 31004275 | 3.44E-05 | 0.41137 | 299 | 0.134924094 | 0.026971769 |
| SNP_A.K_1004836_6_6935117 | 6 | 6935117 | 3.44E-05 | 0.32107 | 299 | 0.134916904 | 0.026992432 |
| SNP_A.K_230050_1_37526891 | 1 | 37526891 | 3.45E-05 | 0.01003 | 299 | 0.134907378 | 0.027023145 |
| SNP_A.K_978906_6_1800232 | 6 | 1800232 | 3.46E-05 | 0.32776 | 299 | 0.134888593 | 0.02709373 |
| SNP_A.K_709866_4_8299906 | 4 | 8299906 | 3.47E-05 | 0.08194 | 299 | 0.134871846 | 0.027150142 |
| SNP_A.K_678979_4_1020675 | 4 | 1020675 | 3.47E-05 | 0.03177 | 299 | 0.13487077 | 0.027150142 |
| SNP_A.K_990284_6_4704748 | 6 | 4704748 | 3.48E-05 | 0.17559 | 299 | 0.134845384 | 0.027222169 |
| SNP_A.K_1001098_6_6441292 | 6 | 6441292 | 3.48E-05 | 0.38796 | 299 | 0.134844879 | 0.027222169 |
| SNP_A.K_1005295_6_7002710 | 6 | 7002710 | 3.48E-05 | 0.32943 | 299 | 0.134843188 | 0.027222169 |
| SNP_A.K_995372_6_5929160 | 6 | 5929160 | 3.48E-05 | 0.36789 | 299 | 0.134841281 | 0.027222169 |
| SNP_A.K_1002155_6_6548233 | 6 | 6548233 | 3.48E-05 | 0.41304 | 299 | 0.134840633 | 0.027222169 |
| SNP_A.K_978049_6_1643938 | 6 | 1643938 | 3.49E-05 | 0.19398 | 299 | 0.134839917 | 0.027222169 |
| SNP_A.K_28875_1_4175841 | 1 | 4175841 | 3.49E-05 | 0.13712 | 299 | 0.134823844 | 0.027281557 |
| SNP_A.K_1830666_11_10573233 | 11 | 10573233 | 3.50E-05 | 0.35619 | 299 | 0.134818356 | 0.02728202 |
| SNP_A.K_307067_2_7377321 | 2 | 7377321 | 3.50E-05 | 0.39799 | 299 | 0.134815893 | 0.02728202 |
| SNP_A.K_129753_1_18609914 | 1 | 18609914 | 3.50E-05 | 0.34281 | 299 | 0.134814708 | 0.02728202 |
| SNP_A.K_1001111_6_6445293 | 6 | 6445293 | 3.50E-05 | 0.38796 | 299 | 0.134810112 | 0.02728202 |
| SNP_A.K_306872_2_7362171 | 2 | 7362171 | 3.50E-05 | 0.24415 | 299 | 0.134808218 | 0.02728202 |
| SNP_A.K_1001567_6_6492126 | 6 | 6492126 | 3.50E-05 | 0.29599 | 299 | 0.134807882 | 0.02728202 |
| SNP_A.K_978449_6_1694640 | 6 | 1694640 | 3.50E-05 | 0.26756 | 299 | 0.134807239 | 0.02728202 |
| SNP_A.K_1831277_11_10615374 | 11 | 10615374 | 3.51E-05 | 0.13378 | 299 | 0.134798734 | 0.027306711 |
| SNP_A.K_1831340_11_10620605 | 11 | 10620605 | 3.51E-05 | 0.13211 | 299 | 0.134793762 | 0.027306711 |
| SNP_A.K_1288082_7_22238441 | 7 | 22238441 | 3.51E-05 | 0.02007 | 299 | 0.134792565 | 0.027306711 |
| SNP_A.K_710476_4_8423994 | 4 | 8423994 | 3.51E-05 | 0.07525 | 299 | 0.134791442 | 0.027306711 |
| SNP_A.K_994771_6_5888257 | 6 | 5888257 | 3.51E-05 | 0.27759 | 299 | 0.134789789 | 0.027306711 |
| SNP_A.K_709823_4_8298539 | 4 | 8298539 | 3.52E-05 | 0.07191 | 299 | 0.134782458 | 0.027328346 |
| SNP_A.K_1000840_6_6420727 | 6 | 6420727 | 3.53E-05 | 0.40134 | 299 | 0.134767986 | 0.027377926 |
| SNP_A.K_1001330_6_6466063 | 6 | 6466063 | 3.53E-05 | 0.29264 | 299 | 0.134766359 | 0.027377926 |
| SNP_A.K_995310_6_5925041 | 6 | 5925041 | 3.53E-05 | 0.14883 | 299 | 0.134755744 | 0.027413934 |
| SNP_A.K_1001517_6_6486511 | 6 | 6486511 | 3.54E-05 | 0.29599 | 299 | 0.134750576 | 0.027426238 |
| SNP_A.K_996048_6_5959414 | 6 | 5959414 | 3.54E-05 | 0.45485 | 299 | 0.134744244 | 0.027443629 |
| SNP_A.K_709888_4_8300350 | 4 | 8300350 | 3.54E-05 | 0.07525 | 299 | 0.134737002 | 0.027465003 |
| SNP_A.K_2078507_12_26993854 | 12 | 26993854 | 3.55E-05 | 0.01672 | 299 | 0.134731326 | 0.027479564 |
| SNP_A.K_1000943_6_6425717 | 6 | 6425717 | 3.55E-05 | 0.301 | 299 | 0.13472561 | 0.027494306 |
| SNP_A.K_996189_6_5973818 | 6 | 5973818 | 3.55E-05 | 0.32943 | 299 | 0.134717749 | 0.027503961 |
| SNP_A.K_201282_1_32159629 | 1 | 32159629 | 3.55E-05 | 0.42475 | 299 | 0.134717038 | 0.027503961 |
| SNP_A.K_709415_4_8265494 | 4 | 8265494 | 3.55E-05 | 0.11204 | 299 | 0.134715514 | 0.027503961 |
| SNP_A.K_1001875_6_6528518 | 6 | 6528518 | 3.56E-05 | 0.3796 | 299 | 0.134714044 | 0.027503961 |
| SNP_A.K_1830660_11_10573074 | 11 | 10573074 | 3.56E-05 | 0.36957 | 299 | 0.134702328 | 0.027544994 |
| SNP_A.K_1600925_9_18862205 | 9 | 18862205 | 3.59E-05 | 0.15886 | 299 | 0.134657316 | 0.027707853 |
| SNP_A.K_1002141_6_6547812 | 6 | 6547812 | 3.59E-05 | 0.41973 | 299 | 0.134657266 | 0.027707853 |
| SNP_A.K_218780_1_35351216 | 1 | 35351216 | 3.59E-05 | 0.16555 | 299 | 0.134653358 | 0.027707853 |

| | | | | | | | |
|---|---|---|---|---|---|---|---|
| SNP_A.K_306972_2_7371274 | 2 | 7371274 | 3.59E-05 | 0.25418 | 299 | 0.134652471 | 0.027707853 |
| SNP_A.K_1002045_6_6537942 | 6 | 6537942 | 3.59E-05 | 0.31773 | 299 | 0.134652398 | 0.027707853 |
| SNP_A.K_307518_2_7414942 | 2 | 7414942 | 3.59E-05 | 0.43645 | 299 | 0.134651243 | 0.027707853 |
| SNP_A.K_995250_6_5921457 | 6 | 5921457 | 3.60E-05 | 0.1806 | 299 | 0.134642437 | 0.027736388 |
| SNP_A.K_1600741_9_18830687 | 9 | 18830687 | 3.60E-05 | 0.23077 | 299 | 0.134640107 | 0.02773639 |
| SNP_A.K_709853_4_8299610 | 4 | 8299610 | 3.60E-05 | 0.07358 | 299 | 0.134635678 | 0.02774565 |
| SNP_A.K_1000461_6_6386487 | 6 | 6386487 | 3.60E-05 | 0.27926 | 299 | 0.134632062 | 0.027751331 |
| SNP_A.K_978198_6_1659942 | 6 | 1659942 | 3.60E-05 | 0.18227 | 299 | 0.134627513 | 0.027758603 |
| SNP_A.K_1001749_6_6514610 | 6 | 6514610 | 3.61E-05 | 0.3194 | 299 | 0.134624158 | 0.027758603 |
| SNP_A.K_710481_4_8424250 | 4 | 8424250 | 3.61E-05 | 0.07358 | 299 | 0.134623435 | 0.027758603 |
| SNP_A.K_306812_2_7354862 | 2 | 7354862 | 3.62E-05 | 0.26589 | 299 | 0.134605655 | 0.027826897 |
| SNP_A.K_28979_1_4192836 | 1 | 4192836 | 3.62E-05 | 0.09365 | 299 | 0.134601827 | 0.027833556 |
| SNP_A.K_1001169_6_6450577 | 6 | 6450577 | 3.63E-05 | 0.28094 | 299 | 0.134586916 | 0.027889321 |
| SNP_A.K_307731_2_7443498 | 2 | 7443498 | 3.63E-05 | 0.45151 | 299 | 0.134579849 | 0.027910369 |
| SNP_A.K_1004211_6_6849785 | 6 | 6849785 | 3.64E-05 | 0.32609 | 299 | 0.134567649 | 0.027954247 |
| SNP_A.K_996108_6_5962408 | 6 | 5962408 | 3.64E-05 | 0.47826 | 299 | 0.134562527 | 0.02796289 |
| SNP_A.K_997627_6_6139934 | 6 | 6139934 | 3.64E-05 | 0.14548 | 299 | 0.134560926 | 0.02796289 |
| SNP_A.K_1004880_6_6942548 | 6 | 6942548 | 3.64E-05 | 0.4097 | 299 | 0.134558746 | 0.02796289 |
| SNP_A.K_374682_2_17871647 | 2 | 17871647 | 3.67E-05 | 0.05351 | 299 | 0.134518298 | 0.028130081 |
| SNP_A.K_709416_4_8265516 | 4 | 8265516 | 3.67E-05 | 0.08863 | 299 | 0.134516624 | 0.028130081 |
| SNP_A.K_996020_6_5958060 | 6 | 5958060 | 3.68E-05 | 0.29599 | 299 | 0.13450236 | 0.028163467 |
| SNP_A.K_709788_4_8297782 | 4 | 8297782 | 3.68E-05 | 0.07191 | 299 | 0.134499449 | 0.028163467 |
| SNP_A.K_995626_6_5943865 | 6 | 5943865 | 3.68E-05 | 0.46488 | 299 | 0.134497736 | 0.028163467 |
| SNP_A.K_307359_2_7398637 | 2 | 7398637 | 3.68E-05 | 0.44649 | 299 | 0.134495729 | 0.028163467 |
| SNP_A.K_709869_4_8299971 | 4 | 8299971 | 3.68E-05 | 0.07525 | 299 | 0.134495284 | 0.028163467 |
| SNP_A.K_709870_4_8299994 | 4 | 8299994 | 3.68E-05 | 0.07525 | 299 | 0.134495284 | 0.028163467 |
| SNP_A.K_1001518_6_6486525 | 6 | 6486525 | 3.69E-05 | 0.29264 | 299 | 0.134485577 | 0.028172043 |
| SNP_A.K_710037_4_8310626 | 4 | 8310626 | 3.69E-05 | 0.07692 | 299 | 0.134482065 | 0.028172043 |
| SNP_A.K_1005581_6_7028127 | 6 | 7028127 | 3.69E-05 | 0.13712 | 299 | 0.134481637 | 0.028172043 |
| SNP_A.K_978432_6_1690843 | 6 | 1690843 | 3.69E-05 | 0.24916 | 299 | 0.134480711 | 0.028172043 |
| SNP_A.K_990366_6_4712521 | 6 | 4712521 | 3.69E-05 | 0.20234 | 299 | 0.134479758 | 0.028172043 |
| SNP_A.K_1001482_6_6483366 | 6 | 6483366 | 3.69E-05 | 0.29264 | 299 | 0.134479519 | 0.028172043 |
| SNP_A.K_198084_1_31695572 | 1 | 31695572 | 3.69E-05 | 0.46154 | 299 | 0.134475331 | 0.028180479 |
| SNP_A.K_995623_6_5943761 | 6 | 5943761 | 3.70E-05 | 0.45652 | 299 | 0.134464015 | 0.028220896 |
| SNP_A.K_1001187_6_6453566 | 6 | 6453566 | 3.72E-05 | 0.27926 | 299 | 0.134429113 | 0.028359205 |
| SNP_A.K_1004989_6_6960170 | 6 | 6960170 | 3.72E-05 | 0.30268 | 299 | 0.134426988 | 0.028359205 |
| SNP_A.K_1500598_9_725474 | 9 | 725474 | 3.72E-05 | 0.03679 | 299 | 0.134425393 | 0.028359205 |
| SNP_A.K_998754_6_6219912 | 6 | 6219912 | 3.72E-05 | 0.26589 | 299 | 0.134424067 | 0.028359205 |
| SNP_A.K_442795_2_28971305 | 2 | 28971305 | 3.73E-05 | 0.03679 | 299 | 0.134420145 | 0.028366518 |
| SNP_A.K_1001344_6_6466344 | 6 | 6466344 | 3.73E-05 | 0.29264 | 299 | 0.134406218 | 0.028418275 |
| SNP_A.K_1001473_6_6483025 | 6 | 6483025 | 3.74E-05 | 0.29264 | 299 | 0.134401789 | 0.028418275 |
| SNP_A.K_1001474_6_6483046 | 6 | 6483046 | 3.74E-05 | 0.29264 | 299 | 0.134401789 | 0.028418275 |
| SNP_A.K_306766_2_7348832 | 2 | 7348832 | 3.75E-05 | 0.25084 | 299 | 0.134374389 | 0.028531975 |
| SNP_A.K_456449_2_33038925 | 2 | 33038925 | 3.76E-05 | 0.13211 | 299 | 0.134368154 | 0.028549863 |
| SNP_A.K_800197_4_28269322 | 4 | 28269322 | 3.77E-05 | 0.15886 | 299 | 0.134347394 | 0.028633845 |
| SNP_A.K_990280_6_4704420 | 6 | 4704420 | 3.78E-05 | 0.43311 | 299 | 0.134335447 | 0.028677848 |
| SNP_A.K_709835_4_8299235 | 4 | 8299235 | 3.79E-05 | 0.07358 | 299 | 0.134310919 | 0.028779461 |
| SNP_A.K_710421_4_8361024 | 4 | 8361024 | 3.80E-05 | 0.17224 | 299 | 0.134294152 | 0.028845814 |
| SNP_A.K_978425_6_1689632 | 6 | 1689632 | 3.80E-05 | 0.27926 | 299 | 0.134289698 | 0.02885574 |
| SNP_A.K_1001508_6_6485463 | 6 | 6485463 | 3.81E-05 | 0.29766 | 299 | 0.134282493 | 0.028871231 |

| | | | | | | | |
|---|---|---|---|---|---|---|---|
| SNP_A.K_311714_2_7854503 | 2 | 7854503 | 3.81E-05 | 0.20903 | 299 | 0.134281745 | 0.028871231 |
| SNP_A.K_760915_4_19816982 | 4 | 19816982 | 3.82E-05 | 0.01338 | 299 | 0.134271269 | 0.028908862 |
| SNP_A.K_1925314_11_24727277 | 11 | 24727277 | 3.83E-05 | 0.02843 | 299 | 0.134254577 | 0.028975189 |
| SNP_A.K_999127_6_6257882 | 6 | 6257882 | 3.83E-05 | 0.15886 | 299 | 0.134249585 | 0.02898766 |
| SNP_A.K_306957_2_7370659 | 2 | 7370659 | 3.83E-05 | 0.2408 | 299 | 0.134241129 | 0.02901613 |
| SNP_A.K_726191_4_11927318 | 4 | 11927318 | 3.84E-05 | 0.10201 | 299 | 0.134236964 | 0.02902481 |
| SNP_A.K_307309_2_7394056 | 2 | 7394056 | 3.85E-05 | 0.23579 | 299 | 0.134216569 | 0.029102358 |
| SNP_A.K_2080302_12_27255343 | 12 | 27255343 | 3.85E-05 | 0.02508 | 299 | 0.134215627 | 0.029102358 |
| SNP_A.K_996245_6_5976867 | 6 | 5976867 | 3.85E-05 | 0.47659 | 299 | 0.134208149 | 0.029126428 |
| SNP_A.K_307115_2_7380610 | 2 | 7380610 | 3.86E-05 | 0.40134 | 299 | 0.134195886 | 0.029172719 |
| SNP_A.K_992002_6_5607592 | 6 | 5607592 | 3.87E-05 | 0.19398 | 299 | 0.134174491 | 0.029256024 |
| SNP_A.K_977990_6_1631244 | 6 | 1631244 | 3.88E-05 | 0.16555 | 299 | 0.134172762 | 0.029256024 |
| SNP_A.K_1500115_9_680282 | 9 | 680282 | 3.88E-05 | 0.04515 | 299 | 0.134170019 | 0.029256024 |
| SNP_A.K_725704_4_11832175 | 4 | 11832175 | 3.88E-05 | 0.09365 | 299 | 0.134168848 | 0.029256024 |
| SNP_A.K_1001258_6_6459473 | 6 | 6459473 | 3.88E-05 | 0.37793 | 299 | 0.134160634 | 0.029283676 |
| SNP_A.K_923781_5_19501635 | 5 | 19501635 | 3.89E-05 | 0.03344 | 299 | 0.134152439 | 0.029311271 |
| SNP_A.K_1608488_9_20763316 | 9 | 20763316 | 3.89E-05 | 0.28261 | 299 | 0.134143552 | 0.029342125 |
| SNP_A.K_2064710_12_24187992 | 12 | 24187992 | 3.90E-05 | 0.0301 | 299 | 0.134138534 | 0.029354935 |
| SNP_A.K_307942_2_7457763 | 2 | 7457763 | 3.90E-05 | 0.45485 | 299 | 0.134133806 | 0.029366404 |
| SNP_A.K_325288_2_9863611 | 2 | 9863611 | 3.90E-05 | 0.1087 | 299 | 0.13412711 | 0.029387078 |
| SNP_A.K_977885_6_1612551 | 6 | 1612551 | 3.91E-05 | 0.17893 | 299 | 0.134115831 | 0.029429231 |
| SNP_A.K_374879_2_17910705 | 2 | 17910705 | 3.92E-05 | 0.04013 | 299 | 0.134104376 | 0.029469832 |
| SNP_A.K_710118_4_8317277 | 4 | 8317277 | 3.92E-05 | 0.09365 | 299 | 0.134102626 | 0.029469832 |
| SNP_A.K_1830696_11_10574736 | 11 | 10574736 | 3.92E-05 | 0.1388 | 299 | 0.134100144 | 0.029470828 |
| SNP_A.K_1432058_8_16448292 | 8 | 16448292 | 3.92E-05 | 0.0786 | 299 | 0.134096412 | 0.029477692 |
| SNP_A.K_990308_6_4709872 | 6 | 4709872 | 3.94E-05 | 0.20067 | 299 | 0.134066298 | 0.029602655 |
| SNP_A.K_990252_6_4703350 | 6 | 4703350 | 3.94E-05 | 0.4398 | 299 | 0.134064588 | 0.029602655 |
| SNP_A.K_1004912_6_6952238 | 6 | 6952238 | 3.94E-05 | 0.37124 | 299 | 0.134063031 | 0.029602655 |
| SNP_A.K_993957_6_5802362 | 6 | 5802362 | 3.95E-05 | 0.20234 | 299 | 0.134057314 | 0.029618921 |
| SNP_A.K_1004229_6_6851932 | 6 | 6851932 | 3.96E-05 | 0.32441 | 299 | 0.134033287 | 0.029704717 |
| SNP_A.K_710287_4_8334822 | 4 | 8334822 | 3.96E-05 | 0.08027 | 299 | 0.134032586 | 0.029704717 |
| SNP_A.K_978291_6_1669858 | 6 | 1669858 | 3.96E-05 | 0.2592 | 299 | 0.13403235 | 0.029704717 |
| SNP_A.K_1004974_6_6959305 | 6 | 6959305 | 3.96E-05 | 0.31605 | 299 | 0.134030012 | 0.029705076 |
| SNP_A.K_1001526_6_6487294 | 6 | 6487294 | 3.97E-05 | 0.29766 | 299 | 0.134019796 | 0.029742708 |
| SNP_A.K_990277_6_4704301 | 6 | 4704301 | 3.97E-05 | 0.20569 | 299 | 0.134017457 | 0.029743083 |
| SNP_A.K_710398_4_8340344 | 4 | 8340344 | 3.98E-05 | 0.07525 | 299 | 0.13400927 | 0.029771157 |
| SNP_A.K_978809_6_1783411 | 6 | 1783411 | 3.99E-05 | 0.35452 | 299 | 0.133990631 | 0.029848885 |
| SNP_A.K_1001420_6_6481121 | 6 | 6481121 | 3.99E-05 | 0.32274 | 299 | 0.133985901 | 0.029860634 |
| SNP_A.K_978374_6_1683886 | 6 | 1683886 | 4.00E-05 | 0.26756 | 299 | 0.133968465 | 0.029932872 |
| SNP_A.K_1001324_6_6465263 | 6 | 6465263 | 4.01E-05 | 0.30769 | 299 | 0.133959428 | 0.029946099 |
| SNP_A.K_307245_2_7387789 | 2 | 7387789 | 4.01E-05 | 0.23913 | 299 | 0.133958834 | 0.029946099 |
| SNP_A.K_307536_2_7416263 | 2 | 7416263 | 4.01E-05 | 0.2592 | 299 | 0.133957057 | 0.029946099 |
| SNP_A.K_307258_2_7388404 | 2 | 7388404 | 4.01E-05 | 0.44816 | 299 | 0.13395667 | 0.029946099 |
| SNP_A.K_1001500_6_6484753 | 6 | 6484753 | 4.01E-05 | 0.32943 | 299 | 0.1339539 | 0.029948567 |
| SNP_A.K_218548_1_35311188 | 1 | 35311188 | 4.01E-05 | 0.15719 | 299 | 0.133950875 | 0.029952252 |
| SNP_A.K_978845_6_1787322 | 6 | 1787322 | 4.02E-05 | 0.39298 | 299 | 0.133938214 | 0.030001931 |
| SNP_A.K_978472_6_1698496 | 6 | 1698496 | 4.03E-05 | 0.25753 | 299 | 0.133933747 | 0.030008877 |
| SNP_A.K_978388_6_1684979 | 6 | 1684979 | 4.03E-05 | 0.27425 | 299 | 0.13393226 | 0.030008877 |
| SNP_A.K_1001357_6_6466671 | 6 | 6466671 | 4.03E-05 | 0.29599 | 299 | 0.13392362 | 0.030039424 |
| SNP_A.K_1001479_6_6483202 | 6 | 6483202 | 4.04E-05 | 0.29431 | 299 | 0.133918192 | 0.030054636 |

| | | | | | | | |
|---|---|---|---|---|---|---|---|
| SNP_A.K_1000825_6_6418834 | 6 | 6418834 | 4.04E-05 | 0.39465 | 299 | 0.133907165 | 0.030089063 |
| SNP_A.K_1001756_6_6514964 | 6 | 6514964 | 4.04E-05 | 0.29766 | 299 | 0.133905515 | 0.030089063 |
| SNP_A.K_709762_4_8285211 | 4 | 8285211 | 4.04E-05 | 0.09365 | 299 | 0.133904263 | 0.030089063 |
| SNP_A.K_725358_4_11797977 | 4 | 11797977 | 4.05E-05 | 0.10033 | 299 | 0.133900958 | 0.030094143 |
| SNP_A.K_710327_4_8336476 | 4 | 8336476 | 4.05E-05 | 0.07358 | 299 | 0.133894997 | 0.030101387 |
| SNP_A.K_196663_1_31474094 | 1 | 31474094 | 4.05E-05 | 0.30936 | 299 | 0.13389496 | 0.030101387 |
| SNP_A.K_1000864_6_6421527 | 6 | 6421527 | 4.05E-05 | 0.28261 | 299 | 0.133892048 | 0.030104598 |
| SNP_A.K_38247_1_5512773 | 1 | 5512773 | 4.06E-05 | 0.0301 | 299 | 0.133885572 | 0.030124902 |
| SNP_A.K_1000912_6_6423386 | 6 | 6423386 | 4.06E-05 | 0.28595 | 299 | 0.133880973 | 0.030136216 |
| SNP_A.K_1001718_6_6508930 | 6 | 6508930 | 4.06E-05 | 0.30936 | 299 | 0.133876037 | 0.030149155 |
| SNP_A.K_1600842_9_18848742 | 9 | 18848742 | 4.07E-05 | 0.13712 | 299 | 0.133866225 | 0.030185532 |
| SNP_A.K_1001219_6_6455402 | 6 | 6455402 | 4.07E-05 | 0.35953 | 299 | 0.133863023 | 0.030190163 |
| SNP_A.K_1466264_8_21215845 | 8 | 21215845 | 4.07E-05 | 0.04013 | 299 | 0.133859345 | 0.030197088 |
| SNP_A.K_129814_1_18619202 | 1 | 18619202 | 4.08E-05 | 0.21405 | 299 | 0.133846963 | 0.03024591 |
| SNP_A.K_978607_6_1740733 | 6 | 1740733 | 4.08E-05 | 0.26756 | 299 | 0.133843266 | 0.030252951 |
| SNP_A.K_995388_6_5931368 | 6 | 5931368 | 4.09E-05 | 0.47324 | 299 | 0.133835629 | 0.030255259 |
| SNP_A.K_306814_2_7355244 | 2 | 7355244 | 4.09E-05 | 0.301 | 299 | 0.133834318 | 0.030255259 |
| SNP_A.K_978454_6_1694808 | 6 | 1694808 | 4.09E-05 | 0.27425 | 299 | 0.133833198 | 0.030255259 |
| SNP_A.K_673337_4_349572 | 4 | 349572 | 4.09E-05 | 0.08361 | 299 | 0.13383165 | 0.030255259 |
| SNP_A.K_978869_6_1793572 | 6 | 1793572 | 4.09E-05 | 0.3194 | 299 | 0.133831621 | 0.030255259 |
| SNP_A.K_1608492_9_20763701 | 9 | 20763701 | 4.10E-05 | 0.29766 | 299 | 0.133824051 | 0.030280996 |
| SNP_A.K_1390306_8_10178028 | 8 | 10178028 | 4.11E-05 | 0.06689 | 299 | 0.133802127 | 0.030376102 |
| SNP_A.K_1001530_6_6487508 | 6 | 6487508 | 4.12E-05 | 0.29599 | 299 | 0.13379317 | 0.030394816 |
| SNP_A.K_677697_4_904276 | 4 | 904276 | 4.12E-05 | 0.20067 | 299 | 0.133791869 | 0.030394816 |
| SNP_A.K_92744_1_12757601 | 1 | 12757601 | 4.12E-05 | 0.14548 | 299 | 0.133791575 | 0.030394816 |
| SNP_A.K_978251_6_1665566 | 6 | 1665566 | 4.12E-05 | 0.20067 | 299 | 0.133789291 | 0.030395094 |
| SNP_A.K_87017_1_11862775 | 1 | 11862775 | 4.13E-05 | 0.07692 | 299 | 0.133775874 | 0.030446992 |
| SNP_A.K_1534450_9_5986609 | 9 | 5986609 | 4.13E-05 | 0.12542 | 299 | 0.133774127 | 0.030446992 |
| SNP_A.K_1001167_6_6450534 | 6 | 6450534 | 4.13E-05 | 0.29933 | 299 | 0.133766271 | 0.030470335 |
| SNP_A.K_218488_1_35296897 | 1 | 35296897 | 4.14E-05 | 0.1689 | 299 | 0.133764867 | 0.030470335 |
| SNP_A.K_709367_4_8259002 | 4 | 8259002 | 4.14E-05 | 0.08696 | 299 | 0.133752369 | 0.030512502 |
| SNP_A.K_1001754_6_6514921 | 6 | 6514921 | 4.14E-05 | 0.30268 | 299 | 0.133751739 | 0.030512502 |
| SNP_A.K_1608518_9_20771552 | 9 | 20771552 | 4.15E-05 | 0.27258 | 299 | 0.133747809 | 0.030520811 |
| SNP_A.K_1364928_8_6059477 | 8 | 6059477 | 4.15E-05 | 0.12207 | 299 | 0.133745475 | 0.030521362 |
| SNP_A.K_28934_1_4183585 | 1 | 4183585 | 4.15E-05 | 0.08863 | 299 | 0.13373789 | 0.030547455 |
| SNP_A.K_1001561_6_6491839 | 6 | 6491839 | 4.16E-05 | 0.29766 | 299 | 0.133734523 | 0.030553045 |
| SNP_A.K_710499_4_8425039 | 4 | 8425039 | 4.16E-05 | 0.07525 | 299 | 0.133729372 | 0.030567318 |
| SNP_A.K_978148_6_1654369 | 6 | 1654369 | 4.17E-05 | 0.26087 | 299 | 0.13371953 | 0.030604464 |
| SNP_A.K_993233_6_5723185 | 6 | 5723185 | 4.17E-05 | 0.47324 | 299 | 0.133713001 | 0.030625494 |
| SNP_A.K_1000974_6_6430386 | 6 | 6430386 | 4.17E-05 | 0.32943 | 299 | 0.133710755 | 0.030625639 |
| SNP_A.K_752461_4_17906704 | 4 | 17906704 | 4.17E-05 | 0.06522 | 299 | 0.133707976 | 0.030628395 |
| SNP_A.K_996121_6_5963687 | 6 | 5963687 | 4.18E-05 | 0.49331 | 299 | 0.133703394 | 0.030636212 |
| SNP_A.K_908144_5_17480756 | 5 | 17480756 | 4.18E-05 | 0.03344 | 299 | 0.133700425 | 0.030636212 |
| SNP_A.K_1001495_6_6484023 | 6 | 6484023 | 4.18E-05 | 0.30268 | 299 | 0.133698203 | 0.030636212 |
| SNP_A.K_993451_6_5752386 | 6 | 5752386 | 4.18E-05 | 0.13712 | 299 | 0.133694353 | 0.030636212 |
| SNP_A.K_1488082_8_26998369 | 8 | 26998369 | 4.18E-05 | 0.49833 | 299 | 0.133690622 | 0.030636212 |
| SNP_A.K_1183415_7_4548555 | 7 | 4548555 | 4.19E-05 | 0.44482 | 299 | 0.133689815 | 0.030636212 |
| SNP_A.K_456486_2_33045111 | 2 | 33045111 | 4.19E-05 | 0.07191 | 299 | 0.13368887 | 0.030636212 |
| SNP_A.K_709855_4_8299648 | 4 | 8299648 | 4.19E-05 | 0.0786 | 299 | 0.133688687 | 0.030636212 |
| SNP_A.K_978150_6_1654538 | 6 | 1654538 | 4.19E-05 | 0.27759 | 299 | 0.13368615 | 0.030637822 |

| | | | | | | | |
|---|---|---|---|---|---|---|---|
| SNP_A.K_1830751_11_10577260 | 11 | 10577260 | 4.20E-05 | 0.13211 | 299 | 0.133673732 | 0.03068769 |
| SNP_A.K_1001499_6_6484709 | 6 | 6484709 | 4.20E-05 | 0.30602 | 299 | 0.133671525 | 0.030687696 |
| SNP_A.K_1004347_6_6870573 | 6 | 6870573 | 4.20E-05 | 0.37793 | 299 | 0.133661456 | 0.030726158 |
| SNP_A.K_306894_2_7364239 | 2 | 7364239 | 4.21E-05 | 0.26421 | 299 | 0.133655598 | 0.030744045 |
| SNP_A.K_990326_6_4710709 | 6 | 4710709 | 4.21E-05 | 0.19398 | 299 | 0.133652778 | 0.030747067 |
| SNP_A.K_709851_4_8299584 | 4 | 8299584 | 4.21E-05 | 0.07525 | 299 | 0.13365049 | 0.030747484 |
| SNP_A.K_1600700_9_18822430 | 9 | 18822430 | 4.22E-05 | 0.16221 | 299 | 0.133643236 | 0.03077224 |
| SNP_A.K_1001525_6_6487286 | 6 | 6487286 | 4.22E-05 | 0.29933 | 299 | 0.133638578 | 0.030784284 |
| SNP_A.K_981235_6_2810540 | 6 | 2810540 | 4.23E-05 | 0.02174 | 299 | 0.133629602 | 0.03081753 |
| SNP_A.K_978503_6_1707890 | 6 | 1707890 | 4.23E-05 | 0.26923 | 299 | 0.133625499 | 0.030823856 |
| SNP_A.K_306767_2_7348975 | 2 | 7348975 | 4.23E-05 | 0.2408 | 299 | 0.133623183 | 0.030823856 |
| SNP_A.K_708703_4_8087285 | 4 | 8087285 | 4.23E-05 | 0.08696 | 299 | 0.133621717 | 0.030823856 |
| SNP_A.K_1600950_9_18864583 | 9 | 18864583 | 4.25E-05 | 0.12375 | 299 | 0.133593665 | 0.030951069 |
| SNP_A.K_978421_6_1689373 | 6 | 1689373 | 4.26E-05 | 0.26589 | 299 | 0.133577935 | 0.031017869 |
| SNP_A.K_1001354_6_6466629 | 6 | 6466629 | 4.29E-05 | 0.301 | 299 | 0.133529173 | 0.031243151 |
| SNP_A.K_1004128_6_6836042 | 6 | 6836042 | 4.29E-05 | 0.42475 | 299 | 0.13352812 | 0.031243151 |
| SNP_A.K_709366_4_8258985 | 4 | 8258985 | 4.30E-05 | 0.08863 | 299 | 0.133513814 | 0.031288404 |
| SNP_A.K_710317_4_8336224 | 4 | 8336224 | 4.30E-05 | 0.07525 | 299 | 0.133513341 | 0.031288404 |
| SNP_A.K_1608658_9_20811050 | 9 | 20811050 | 4.31E-05 | 0.27258 | 299 | 0.133512457 | 0.031288404 |
| SNP_A.K_1804325_11_7061474 | 11 | 7061474 | 4.32E-05 | 0.01171 | 299 | 0.133496292 | 0.031358142 |
| SNP_A.K_1004354_6_6874923 | 6 | 6874923 | 4.32E-05 | 0.37291 | 299 | 0.133491307 | 0.031372107 |
| SNP_A.K_1488852_8_27109890 | 8 | 27109890 | 4.32E-05 | 0.301 | 299 | 0.133484931 | 0.031393042 |
| SNP_A.K_1001356_6_6466660 | 6 | 6466660 | 4.33E-05 | 0.29933 | 299 | 0.133481546 | 0.031399023 |
| SNP_A.K_513362_3_8240961 | 3 | 8240961 | 4.33E-05 | 0.03177 | 299 | 0.133477403 | 0.031408809 |
| SNP_A.K_690484_4_4430184 | 4 | 4430184 | 4.33E-05 | 0.03512 | 299 | 0.133474042 | 0.031414682 |
| SNP_A.K_709974_4_8304052 | 4 | 8304052 | 4.34E-05 | 0.07525 | 299 | 0.133468708 | 0.031430438 |
| SNP_A.K_999761_6_6312684 | 6 | 6312684 | 4.34E-05 | 0.30268 | 299 | 0.133463742 | 0.031434487 |
| SNP_A.K_977948_6_1623891 | 6 | 1623891 | 4.34E-05 | 0.20401 | 299 | 0.13346353 | 0.031434487 |
| SNP_A.K_997452_6_6100121 | 6 | 6100121 | 4.34E-05 | 0.37291 | 299 | 0.133460272 | 0.031439864 |
| SNP_A.K_1000970_6_6429793 | 6 | 6429793 | 4.35E-05 | 0.08528 | 299 | 0.133447889 | 0.031491003 |
| SNP_A.K_194493_1_30878501 | 1 | 30878501 | 4.36E-05 | 0.37291 | 299 | 0.133440003 | 0.031519636 |
| SNP_A.K_312180_2_7920505 | 2 | 7920505 | 4.36E-05 | 0.26589 | 299 | 0.133428526 | 0.031566354 |
| SNP_A.K_306741_2_7347106 | 2 | 7347106 | 4.37E-05 | 0.24916 | 299 | 0.133420673 | 0.031592324 |
| SNP_A.K_307500_2_7413232 | 2 | 7413232 | 4.37E-05 | 0.42475 | 299 | 0.133419004 | 0.031592324 |
| SNP_A.K_308616_2_7529715 | 2 | 7529715 | 4.37E-05 | 0.18729 | 299 | 0.133414599 | 0.031603532 |
| SNP_A.K_1924815_11_24669026 | 11 | 24669026 | 4.38E-05 | 0.01839 | 299 | 0.133402987 | 0.031612561 |
| SNP_A.K_1924832_11_24669843 | 11 | 24669843 | 4.38E-05 | 0.01839 | 299 | 0.133402987 | 0.031612561 |
| SNP_A.K_1924678_11_24662948 | 11 | 24662948 | 4.38E-05 | 0.01839 | 299 | 0.133402987 | 0.031612561 |
| SNP_A.K_1924680_11_24662984 | 11 | 24662984 | 4.38E-05 | 0.01839 | 299 | 0.133402987 | 0.031612561 |
| SNP_A.K_1001321_6_6465099 | 6 | 6465099 | 4.38E-05 | 0.29599 | 299 | 0.133401921 | 0.031612561 |
| SNP_A.K_995649_6_5946272 | 6 | 5946272 | 4.39E-05 | 0.4699 | 299 | 0.133390077 | 0.031661309 |
| SNP_A.K_709706_4_8282569 | 4 | 8282569 | 4.40E-05 | 0.0903 | 299 | 0.133373827 | 0.03173241 |
| SNP_A.K_1000915_6_6423464 | 6 | 6423464 | 4.40E-05 | 0.28428 | 299 | 0.133370828 | 0.031736584 |
| SNP_A.K_1001521_6_6487041 | 6 | 6487041 | 4.41E-05 | 0.30602 | 299 | 0.133366394 | 0.031748024 |
| SNP_A.K_723448_4_11457373 | 4 | 11457373 | 4.41E-05 | 0.0786 | 299 | 0.133364086 | 0.031748718 |
| SNP_A.K_1001466_6_6482649 | 6 | 6482649 | 4.41E-05 | 0.29431 | 299 | 0.133361612 | 0.031750249 |
| SNP_A.K_310601_2_7750247 | 2 | 7750247 | 4.41E-05 | 0.18896 | 299 | 0.133356573 | 0.031758055 |
| SNP_A.K_1001974_6_6533183 | 6 | 6533183 | 4.42E-05 | 0.4097 | 299 | 0.133353767 | 0.031758055 |
| SNP_A.K_678555_4_989658 | 4 | 989658 | 4.42E-05 | 0.10368 | 299 | 0.133353001 | 0.031758055 |
| SNP_A.K_996146_6_5964975 | 6 | 5964975 | 4.42E-05 | 0.33779 | 299 | 0.133351394 | 0.031758055 |

| | | | | | | | |
|---|---|---|---|---|---|---|---|
| SNP_A.K_996138_6_5964506 | 6 | 5964506 | 4.43E-05 | 0.33612 | 299 | 0.133333256 | 0.031838995 |
| SNP_A.K_10661_1_1870871 | 1 | 1870871 | 4.43E-05 | 0.02508 | 299 | 0.133326564 | 0.031861969 |
| SNP_A.K_978460_6_1695530 | 6 | 1695530 | 4.45E-05 | 0.25084 | 299 | 0.133301527 | 0.031977008 |
| SNP_A.K_307446_2_7408100 | 2 | 7408100 | 4.45E-05 | 0.29599 | 299 | 0.133298817 | 0.031977008 |
| SNP_A.K_1595429_9_17659880 | 9 | 17659880 | 4.46E-05 | 0.03679 | 299 | 0.133294137 | 0.031977008 |
| SNP_A.K_1382962_8_9125343 | 8 | 9125343 | 4.46E-05 | 0.28261 | 299 | 0.133293609 | 0.031977008 |
| SNP_A.K_673426_4_360850 | 4 | 360850 | 4.46E-05 | 0.13211 | 299 | 0.133293132 | 0.031977008 |
| SNP_A.K_1001574_6_6492463 | 6 | 6492463 | 4.48E-05 | 0.29766 | 299 | 0.133267475 | 0.032096983 |
| SNP_A.K_978081_6_1646331 | 6 | 1646331 | 4.48E-05 | 0.27926 | 299 | 0.13326178 | 0.032115069 |
| SNP_A.K_678721_4_995235 | 4 | 995235 | 4.50E-05 | 0.07525 | 299 | 0.133240836 | 0.032208285 |
| SNP_A.K_1830643_11_10572466 | 11 | 10572466 | 4.50E-05 | 0.16054 | 299 | 0.133239278 | 0.032208285 |
| SNP_A.K_1183407_7_4542597 | 7 | 4542597 | 4.50E-05 | 0.45652 | 299 | 0.133230936 | 0.032240049 |
| SNP_A.K_978956_6_1830253 | 6 | 1830253 | 4.51E-05 | 0.43144 | 299 | 0.13322647 | 0.032251913 |
| SNP_A.K_1578128_9_14012595 | 9 | 14012595 | 4.52E-05 | 0.05518 | 299 | 0.13320607 | 0.032345844 |
| SNP_A.K_710415_4_8349159 | 4 | 8349159 | 4.53E-05 | 0.15719 | 299 | 0.133194872 | 0.0323827 |
| SNP_A.K_1001527_6_6487331 | 6 | 6487331 | 4.53E-05 | 0.29766 | 299 | 0.133193065 | 0.0323827 |
| SNP_A.K_1608578_9_20789266 | 9 | 20789266 | 4.53E-05 | 0.22241 | 299 | 0.133192461 | 0.0323827 |
| SNP_A.K_1001506_6_6485316 | 6 | 6485316 | 4.54E-05 | 0.29766 | 299 | 0.133185084 | 0.032404398 |
| SNP_A.K_1184050_7_4647314 | 7 | 4647314 | 4.54E-05 | 0.05184 | 299 | 0.133183953 | 0.032404398 |
| SNP_A.K_710334_4_8336704 | 4 | 8336704 | 4.55E-05 | 0.07358 | 299 | 0.133167714 | 0.03247725 |
| SNP_A.K_993267_6_5728710 | 6 | 5728710 | 4.55E-05 | 0.25084 | 299 | 0.133164026 | 0.032477669 |
| SNP_A.K_1608508_9_20767377 | 9 | 20767377 | 4.55E-05 | 0.2709 | 299 | 0.133163331 | 0.032477669 |
| SNP_A.K_709938_4_8302518 | 4 | 8302518 | 4.55E-05 | 0.08863 | 299 | 0.133159397 | 0.032486909 |
| SNP_A.K_1001211_6_6455190 | 6 | 6455190 | 4.56E-05 | 0.36288 | 299 | 0.133148555 | 0.032531879 |
| SNP_A.K_709947_4_8302717 | 4 | 8302717 | 4.56E-05 | 0.07358 | 299 | 0.133146422 | 0.032531879 |
| SNP_A.K_996280_6_5981227 | 6 | 5981227 | 4.57E-05 | 0.27926 | 299 | 0.133139683 | 0.032546887 |
| SNP_A.K_978926_6_1807778 | 6 | 1807778 | 4.57E-05 | 0.32107 | 299 | 0.133139235 | 0.032546887 |
| SNP_A.K_1001413_6_6480208 | 6 | 6480208 | 4.57E-05 | 0.301 | 299 | 0.1331369 | 0.032547866 |
| SNP_A.K_310757_2_7765221 | 2 | 7765221 | 4.58E-05 | 0.19231 | 299 | 0.133128904 | 0.032578239 |
| SNP_A.K_709973_4_8303935 | 4 | 8303935 | 4.58E-05 | 0.07525 | 299 | 0.133117958 | 0.032623988 |
| SNP_A.K_1600732_9_18826824 | 9 | 18826824 | 4.59E-05 | 0.13545 | 299 | 0.133110375 | 0.032643118 |
| SNP_A.K_1924699_11_24663452 | 11 | 24663452 | 4.59E-05 | 0.02676 | 299 | 0.133108417 | 0.032643118 |
| SNP_A.K_375437_2_17971845 | 2 | 17971845 | 4.59E-05 | 0.05184 | 299 | 0.133107852 | 0.032643118 |
| SNP_A.K_1389600_8_10109469 | 8 | 10109469 | 4.60E-05 | 0.04682 | 299 | 0.133103059 | 0.032656923 |
| SNP_A.K_991248_6_4883785 | 6 | 4883785 | 4.60E-05 | 0.05017 | 299 | 0.133096708 | 0.032678853 |
| SNP_A.K_1004901_6_6951392 | 6 | 6951392 | 4.60E-05 | 0.35284 | 299 | 0.133090203 | 0.032690793 |
| SNP_A.K_1608757_9_20835803 | 9 | 20835803 | 4.60E-05 | 0.25418 | 299 | 0.133090137 | 0.032690793 |
| SNP_A.K_1001757_6_6514987 | 6 | 6514987 | 4.62E-05 | 0.29599 | 299 | 0.133070071 | 0.032784376 |
| SNP_A.K_978502_6_1707825 | 6 | 1707825 | 4.62E-05 | 0.26421 | 299 | 0.133067672 | 0.03278574 |
| SNP_A.K_990580_6_4726737 | 6 | 4726737 | 4.63E-05 | 0.48997 | 299 | 0.133053011 | 0.03285128 |
| SNP_A.K_1001464_6_6482465 | 6 | 6482465 | 4.64E-05 | 0.30268 | 299 | 0.133037145 | 0.032919356 |
| SNP_A.K_978439_6_1694076 | 6 | 1694076 | 4.65E-05 | 0.2709 | 299 | 0.133034082 | 0.032919356 |
| SNP_A.K_1001332_6_6466081 | 6 | 6466081 | 4.65E-05 | 0.29431 | 299 | 0.133033623 | 0.032919356 |
| SNP_A.K_1001112_6_6445573 | 6 | 6445573 | 4.67E-05 | 0.28763 | 299 | 0.13299732 | 0.033099221 |
| SNP_A.K_977871_6_1606052 | 6 | 1606052 | 4.68E-05 | 0.17893 | 299 | 0.132993894 | 0.033106044 |
| SNP_A.K_194669_1_30907718 | 1 | 30907718 | 4.68E-05 | 0.49833 | 299 | 0.132989068 | 0.033120269 |
| SNP_A.K_1608393_9_20744625 | 9 | 20744625 | 4.69E-05 | 0.26756 | 299 | 0.132976989 | 0.033162055 |
| SNP_A.K_1001199_6_6454785 | 6 | 6454785 | 4.69E-05 | 0.28763 | 299 | 0.132976779 | 0.033162055 |
| SNP_A.K_1183417_7_4550844 | 7 | 4550844 | 4.69E-05 | 0.44147 | 299 | 0.132974769 | 0.033162055 |
| SNP_A.K_1382668_8_9112821 | 8 | 9112821 | 4.69E-05 | 0.27926 | 299 | 0.132968772 | 0.033182515 |

| | | | | | | | |
|---|---|---|---|---|---|---|---|
| SNP_A.K_707412_4_7274495 | 4 | 7274495 | 4.71E-05 | 0.12876 | 299 | 0.132953185 | 0.033253816 |
| SNP_A.K_1001769_6_6519659 | 6 | 6519659 | 4.72E-05 | 0.29431 | 299 | 0.132930204 | 0.0333646 |
| SNP_A.K_1000891_6_6422360 | 6 | 6422360 | 4.73E-05 | 0.3913 | 299 | 0.13292302 | 0.033391523 |
| SNP_A.K_1001191_6_6454480 | 6 | 6454480 | 4.73E-05 | 0.4097 | 299 | 0.132920519 | 0.033393077 |
| SNP_A.K_1001810_6_6522193 | 6 | 6522193 | 4.73E-05 | 0.33612 | 299 | 0.132918476 | 0.033393077 |
| SNP_A.K_478931_3_2119566 | 3 | 2119566 | 4.74E-05 | 0.20569 | 299 | 0.132911542 | 0.033406709 |
| SNP_A.K_1001515_6_6486195 | 6 | 6486195 | 4.74E-05 | 0.29766 | 299 | 0.132907997 | 0.033406709 |
| SNP_A.K_1518023_9_3556650 | 9 | 3556650 | 4.74E-05 | 0.04849 | 299 | 0.132906385 | 0.033406709 |
| SNP_A.K_1000857_6_6421314 | 6 | 6421314 | 4.74E-05 | 0.28595 | 299 | 0.132906301 | 0.033406709 |
| SNP_A.K_218521_1_35301396 | 1 | 35301396 | 4.74E-05 | 0.15719 | 299 | 0.1329053 | 0.033406709 |
| SNP_A.K_1000991_6_6434353 | 6 | 6434353 | 4.75E-05 | 0.32776 | 299 | 0.132896024 | 0.033444858 |
| SNP_A.K_307487_2_7412193 | 2 | 7412193 | 4.75E-05 | 0.28428 | 299 | 0.132893179 | 0.033448723 |
| SNP_A.K_708508_4_8059153 | 4 | 8059153 | 4.75E-05 | 0.0903 | 299 | 0.13288907 | 0.033459338 |
| SNP_A.K_201238_1_32154591 | 1 | 32154591 | 4.76E-05 | 0.43813 | 299 | 0.132881073 | 0.033484295 |
| SNP_A.K_1002036_6_6537606 | 6 | 6537606 | 4.76E-05 | 0.31773 | 299 | 0.132879944 | 0.033484295 |
| SNP_A.K_456484_2_33043808 | 2 | 33043808 | 4.76E-05 | 0.12542 | 299 | 0.132876184 | 0.033484295 |
| SNP_A.K_29312_1_4223119 | 1 | 4223119 | 4.76E-05 | 0.08528 | 299 | 0.132875923 | 0.033484295 |
| SNP_A.K_1804145_11_7036188 | 11 | 7036188 | 4.78E-05 | 0.01171 | 299 | 0.132860556 | 0.033555148 |
| SNP_A.K_710400_4_8340448 | 4 | 8340448 | 4.79E-05 | 0.08863 | 299 | 0.132847832 | 0.033604282 |
| SNP_A.K_1364758_8_6039539 | 8 | 6039539 | 4.79E-05 | 0.11706 | 299 | 0.132847154 | 0.033604282 |
| SNP_A.K_145431_1_22973504 | 1 | 22973504 | 4.79E-05 | 0.04181 | 299 | 0.132844416 | 0.033607625 |
| SNP_A.K_978168_6_1657445 | 6 | 1657445 | 4.80E-05 | 0.16722 | 299 | 0.13283531 | 0.033645137 |
| SNP_A.K_1002032_6_6537549 | 6 | 6537549 | 4.80E-05 | 0.3194 | 299 | 0.132827945 | 0.033673343 |
| SNP_A.K_218927_1_35383812 | 1 | 35383812 | 4.81E-05 | 0.17224 | 299 | 0.132815827 | 0.033716446 |
| SNP_A.K_709839_4_8299274 | 4 | 8299274 | 4.81E-05 | 0.07692 | 299 | 0.132815704 | 0.033716446 |
| SNP_A.K_306970_2_7371140 | 2 | 7371140 | 4.82E-05 | 0.28595 | 299 | 0.132807746 | 0.033747915 |
| SNP_A.K_978353_6_1681467 | 6 | 1681467 | 4.83E-05 | 0.25084 | 299 | 0.132789859 | 0.033832965 |
| SNP_A.K_990416_6_4717114 | 6 | 4717114 | 4.83E-05 | 0.14047 | 299 | 0.13278736 | 0.033835075 |
| SNP_A.K_252893_1_40833008 | 1 | 40833008 | 4.84E-05 | 0.13545 | 299 | 0.132778558 | 0.033871227 |
| SNP_A.K_723964_4_11555307 | 4 | 11555307 | 4.84E-05 | 0.09699 | 299 | 0.132772613 | 0.033882322 |
| SNP_A.K_1004334_6_6869144 | 6 | 6869144 | 4.84E-05 | 0.47826 | 299 | 0.132772292 | 0.033882322 |
| SNP_A.K_306844_2_7359558 | 2 | 7359558 | 4.85E-05 | 0.25084 | 299 | 0.132762994 | 0.033906233 |
| SNP_A.K_999888_6_6331631 | 6 | 6331631 | 4.85E-05 | 0.17726 | 299 | 0.132759603 | 0.033906233 |
| SNP_A.K_307007_2_7372806 | 2 | 7372806 | 4.85E-05 | 0.24247 | 299 | 0.132758954 | 0.033906233 |
| SNP_A.K_306966_2_7371078 | 2 | 7371078 | 4.86E-05 | 0.2408 | 299 | 0.132757626 | 0.033906233 |
| SNP_A.K_891860_5_14667564 | 5 | 14667564 | 4.86E-05 | 0.04348 | 299 | 0.132754524 | 0.033906233 |
| SNP_A.K_690053_4_4370646 | 4 | 4370646 | 4.86E-05 | 0.03512 | 299 | 0.132754363 | 0.033906233 |
| SNP_A.K_1831294_11_10617044 | 11 | 10617044 | 4.86E-05 | 0.13211 | 299 | 0.132753145 | 0.033906233 |
| SNP_A.K_2078681_12_27008327 | 12 | 27008327 | 4.86E-05 | 0.01672 | 299 | 0.132749178 | 0.03391633 |
| SNP_A.K_1600749_9_18832562 | 9 | 18832562 | 4.87E-05 | 0.14716 | 299 | 0.132740862 | 0.033949981 |
| SNP_A.K_710055_4_8312932 | 4 | 8312932 | 4.88E-05 | 0.08027 | 299 | 0.132729917 | 0.033990887 |
| SNP_A.K_1004933_6_6954923 | 6 | 6954923 | 4.88E-05 | 0.30268 | 299 | 0.132727027 | 0.033990887 |
| SNP_A.K_306868_2_7361652 | 2 | 7361652 | 4.88E-05 | 0.23913 | 299 | 0.13272702 | 0.033990887 |
| SNP_A.K_1925491_11_24740300 | 11 | 24740300 | 4.88E-05 | 0.03512 | 299 | 0.132720201 | 0.034016507 |
| SNP_A.K_513950_3_8371268 | 3 | 8371268 | 4.90E-05 | 0.03177 | 299 | 0.13270239 | 0.034101909 |
| SNP_A.K_999084_6_6255049 | 6 | 6255049 | 4.90E-05 | 0.18395 | 299 | 0.132696103 | 0.034124723 |
| SNP_A.K_1002033_6_6537564 | 6 | 6537564 | 4.91E-05 | 0.32107 | 299 | 0.132689376 | 0.034146324 |
| SNP_A.K_710129_4_8317686 | 4 | 8317686 | 4.91E-05 | 0.07191 | 299 | 0.132687948 | 0.034146324 |
| SNP_A.K_1364940_8_6060842 | 8 | 6060842 | 4.92E-05 | 0.14214 | 299 | 0.132674913 | 0.03419686 |
| SNP_A.K_307372_2_7399623 | 2 | 7399623 | 4.92E-05 | 0.26756 | 299 | 0.132674494 | 0.03419686 |

| | | | | | | | |
|---|---|---|---|---|---|---|---|
| SNP_A.K_999572_6_6296929 | 6 | 6296929 | 4.92E-05 | 0.14047 | 299 | 0.132670845 | 0.034202139 |
| SNP_A.K_1383213_8_9134920 | 8 | 9134920 | 4.92E-05 | 0.27926 | 299 | 0.132669343 | 0.034202139 |
| SNP_A.K_307284_2_7391180 | 2 | 7391180 | 4.93E-05 | 0.24582 | 299 | 0.132662428 | 0.034228484 |
| SNP_A.K_170234_1_27321565 | 1 | 27321565 | 4.93E-05 | 0.01672 | 299 | 0.132658369 | 0.034239241 |
| SNP_A.K_709999_4_8305024 | 4 | 8305024 | 4.94E-05 | 0.09365 | 299 | 0.132651359 | 0.034260109 |
| SNP_A.K_1001334_6_6466119 | 6 | 6466119 | 4.94E-05 | 0.28763 | 299 | 0.132650373 | 0.034260109 |
| SNP_A.K_1262477_7_18237155 | 7 | 18237155 | 4.94E-05 | 0.01003 | 299 | 0.132645862 | 0.034264378 |
| SNP_A.K_902298_5_16467041 | 5 | 16467041 | 4.94E-05 | 0.05351 | 299 | 0.132643365 | 0.034264378 |
| SNP_A.K_978440_6_1694270 | 6 | 1694270 | 4.95E-05 | 0.26923 | 299 | 0.132643215 | 0.034264378 |
| SNP_A.K_307303_2_7393503 | 2 | 7393503 | 4.95E-05 | 0.28428 | 299 | 0.132641245 | 0.034264378 |
| SNP_A.K_1608529_9_20772659 | 9 | 20772659 | 4.95E-05 | 0.30602 | 299 | 0.13263625 | 0.034269291 |
| SNP_A.K_1001524_6_6487264 | 6 | 6487264 | 4.95E-05 | 0.29933 | 299 | 0.132636178 | 0.034269291 |
| SNP_A.K_328368_2_10227929 | 2 | 10227929 | 4.96E-05 | 0.0301 | 299 | 0.132626743 | 0.03430952 |
| SNP_A.K_194477_1_30877018 | 1 | 30877018 | 4.96E-05 | 0.4214 | 299 | 0.132619513 | 0.034337717 |
| SNP_A.K_1364927_8_6059438 | 8 | 6059438 | 4.97E-05 | 0.11706 | 299 | 0.132613479 | 0.034348383 |
| SNP_A.K_978356_6_1681580 | 6 | 1681580 | 4.97E-05 | 0.27592 | 299 | 0.132613403 | 0.034348383 |
| SNP_A.K_978249_6_1665555 | 6 | 1665555 | 4.97E-05 | 0.19732 | 299 | 0.132607271 | 0.034370601 |
| SNP_A.K_994746_6_5885600 | 6 | 5885600 | 4.98E-05 | 0.32274 | 299 | 0.132601156 | 0.03439274 |
| SNP_A.K_1001511_6_6485937 | 6 | 6485937 | 4.98E-05 | 0.301 | 299 | 0.13259597 | 0.034409796 |
| SNP_A.K_307018_2_7373163 | 2 | 7373163 | 4.99E-05 | 0.24582 | 299 | 0.132586393 | 0.034438139 |
| SNP_A.K_678668_4_993589 | 4 | 993589 | 4.99E-05 | 0.14381 | 299 | 0.132586366 | 0.034438139 |
| SNP_A.K_1831276_11_10615359 | 11 | 10615359 | 4.99E-05 | 0.13545 | 299 | 0.132584578 | 0.034438139 |
| SNP_A.K_991318_6_4924063 | 6 | 4924063 | 5.00E-05 | 0.05184 | 299 | 0.132576987 | 0.034467198 |
| SNP_A.K_277567_2_2951792 | 2 | 2951792 | 5.00E-05 | 0.07525 | 299 | 0.132575141 | 0.034467198 |
| SNP_A.K_709245_4_8242597 | 4 | 8242597 | 5.02E-05 | 0.10535 | 299 | 0.132554774 | 0.03456021 |
| SNP_A.K_1600692_9_18821938 | 9 | 18821938 | 5.03E-05 | 0.13378 | 299 | 0.132542616 | 0.03456021 |
| SNP_A.K_993232_6_5723162 | 6 | 5723162 | 5.03E-05 | 0.23244 | 299 | 0.132539595 | 0.03456021 |
| SNP_A.K_709781_4_8297600 | 4 | 8297600 | 5.03E-05 | 0.07191 | 299 | 0.132538111 | 0.03456021 |
| SNP_A.K_710478_4_8424155 | 4 | 8424155 | 5.03E-05 | 0.07191 | 299 | 0.132538111 | 0.03456021 |
| SNP_A.K_710491_4_8424867 | 4 | 8424867 | 5.03E-05 | 0.07191 | 299 | 0.132538111 | 0.03456021 |
| SNP_A.K_709811_4_8298263 | 4 | 8298263 | 5.03E-05 | 0.07191 | 299 | 0.132538111 | 0.03456021 |
| SNP_A.K_709889_4_8300393 | 4 | 8300393 | 5.03E-05 | 0.07191 | 299 | 0.132538111 | 0.03456021 |
| SNP_A.K_709890_4_8300404 | 4 | 8300404 | 5.03E-05 | 0.07191 | 299 | 0.132538111 | 0.03456021 |
| SNP_A.K_990271_6_4703970 | 6 | 4703970 | 5.03E-05 | 0.28428 | 299 | 0.132537538 | 0.03456021 |
| SNP_A.K_307673_2_7439718 | 2 | 7439718 | 5.03E-05 | 0.4699 | 299 | 0.132532434 | 0.034576964 |
| SNP_A.K_1005471_6_7015520 | 6 | 7015520 | 5.04E-05 | 0.36789 | 299 | 0.13252223 | 0.034621902 |
| SNP_A.K_725377_4_11799603 | 4 | 11799603 | 5.05E-05 | 0.10368 | 299 | 0.132515782 | 0.034641062 |
| SNP_A.K_736328_4_13863692 | 4 | 13863692 | 5.05E-05 | 0.02174 | 299 | 0.132514631 | 0.034641062 |
| SNP_A.K_326682_2_10038715 | 2 | 10038715 | 5.05E-05 | 0.06856 | 299 | 0.132508401 | 0.034664102 |
| SNP_A.K_991320_6_4925180 | 6 | 4925180 | 5.06E-05 | 0.05518 | 299 | 0.132502359 | 0.034686118 |
| SNP_A.K_1001108_6_6444719 | 6 | 6444719 | 5.07E-05 | 0.39465 | 299 | 0.132490285 | 0.034741587 |
| SNP_A.K_978415_6_1688913 | 6 | 1688913 | 5.08E-05 | 0.25084 | 299 | 0.132476971 | 0.03480404 |
| SNP_A.K_1001216_6_6455322 | 6 | 6455322 | 5.08E-05 | 0.29431 | 299 | 0.132474083 | 0.034808624 |
| SNP_A.K_709960_4_8303544 | 4 | 8303544 | 5.08E-05 | 0.0786 | 299 | 0.132470003 | 0.034819846 |
| SNP_A.K_1005511_6_7017893 | 6 | 7017893 | 5.09E-05 | 0.45318 | 299 | 0.132464102 | 0.034841193 |
| SNP_A.K_678676_4_993666 | 4 | 993666 | 5.10E-05 | 0.08027 | 299 | 0.132445797 | 0.034921704 |
| SNP_A.K_1001743_6_6512565 | 6 | 6512565 | 5.10E-05 | 0.29933 | 299 | 0.132444228 | 0.034921704 |
| SNP_A.K_990398_6_4715599 | 6 | 4715599 | 5.11E-05 | 0.20736 | 299 | 0.132442547 | 0.034921704 |
| SNP_A.K_978187_6_1658641 | 6 | 1658641 | 5.11E-05 | 0.24749 | 299 | 0.132441408 | 0.034921704 |
| SNP_A.K_218885_1_35377097 | 1 | 35377097 | 5.11E-05 | 0.15552 | 299 | 0.132437492 | 0.034930623 |

| | | | | | | | |
|---|---|---|---|---|---|---|---|
| SNP_A.K_129747_1_18608511 | 1 | 18608511 | 5.11E-05 | 0.34615 | 299 | 0.132435694 | 0.034930623 |
| SNP_A.K_195105_1_31045650 | 1 | 31045650 | 5.12E-05 | 0.4214 | 299 | 0.132431004 | 0.034945312 |
| SNP_A.K_709662_4_8279620 | 4 | 8279620 | 5.12E-05 | 0.08863 | 299 | 0.132428437 | 0.034948171 |
| SNP_A.K_793819_4_26293740 | 4 | 26293740 | 5.12E-05 | 0.03512 | 299 | 0.132423478 | 0.034955603 |
| SNP_A.K_994787_6_5889002 | 6 | 5889002 | 5.12E-05 | 0.27926 | 299 | 0.132422996 | 0.034955603 |
| SNP_A.K_673168_4_329777 | 4 | 329777 | 5.13E-05 | 0.10535 | 299 | 0.132415159 | 0.034987892 |
| SNP_A.K_1600724_9_18826447 | 9 | 18826447 | 5.13E-05 | 0.15217 | 299 | 0.132408093 | 0.035015908 |
| SNP_A.K_1002030_6_6537520 | 6 | 6537520 | 5.14E-05 | 0.31773 | 299 | 0.132401867 | 0.03502038 |
| SNP_A.K_992572_6_5659479 | 6 | 5659479 | 5.14E-05 | 0.49666 | 299 | 0.13240142 | 0.03502038 |
| SNP_A.K_1854276_11_14365813 | 11 | 14365813 | 5.14E-05 | 0.07023 | 299 | 0.13240114 | 0.03502038 |
| SNP_A.K_1001657_6_6501047 | 6 | 6501047 | 5.14E-05 | 0.44147 | 299 | 0.132398406 | 0.035024203 |
| SNP_A.K_28910_1_4181265 | 1 | 4181265 | 5.14E-05 | 0.11037 | 299 | 0.132395821 | 0.035027204 |
| SNP_A.K_710328_4_8336593 | 4 | 8336593 | 5.15E-05 | 0.07525 | 299 | 0.132389395 | 0.035051688 |
| SNP_A.K_1608653_9_20810683 | 9 | 20810683 | 5.15E-05 | 0.35619 | 299 | 0.132383848 | 0.03505513 |
| SNP_A.K_908188_5_17488720 | 5 | 17488720 | 5.15E-05 | 0.02341 | 299 | 0.132383483 | 0.03505513 |
| SNP_A.K_710399_4_8340356 | 4 | 8340356 | 5.16E-05 | 0.07692 | 299 | 0.132382639 | 0.03505513 |
| SNP_A.K_312079_2_7904690 | 2 | 7904690 | 5.16E-05 | 0.26923 | 299 | 0.132376809 | 0.035076317 |
| SNP_A.K_1004902_6_6951396 | 6 | 6951396 | 5.16E-05 | 0.35284 | 299 | 0.13237152 | 0.035094489 |
| SNP_A.K_478910_3_2116439 | 3 | 2116439 | 5.17E-05 | 0.20401 | 299 | 0.13236141 | 0.035139714 |
| SNP_A.K_1608479_9_20762642 | 9 | 20762642 | 5.18E-05 | 0.29599 | 299 | 0.132358107 | 0.035146781 |
| SNP_A.K_194271_1_30837003 | 1 | 30837003 | 5.18E-05 | 0.38462 | 299 | 0.132353608 | 0.035160568 |
| SNP_A.K_990460_6_4718899 | 6 | 4718899 | 5.19E-05 | 0.1505 | 299 | 0.132336476 | 0.035245383 |
| SNP_A.K_978166_6_1656871 | 6 | 1656871 | 5.19E-05 | 0.26421 | 299 | 0.132334394 | 0.035245611 |
| SNP_A.K_526646_3_10954259 | 3 | 10954259 | 5.20E-05 | 0.01672 | 299 | 0.132324075 | 0.035292235 |
| SNP_A.K_1925566_11_24745599 | 11 | 24745599 | 5.21E-05 | 0.0301 | 299 | 0.132321591 | 0.035294737 |
| SNP_A.K_1123826_6_23600406 | 6 | 23600406 | 5.21E-05 | 0.01672 | 299 | 0.132315233 | 0.035319079 |
| SNP_A.K_710061_4_8313271 | 4 | 8313271 | 5.22E-05 | 0.09197 | 299 | 0.132308683 | 0.035344536 |
| SNP_A.K_307502_2_7413268 | 2 | 7413268 | 5.22E-05 | 0.42809 | 299 | 0.132299536 | 0.03538468 |
| SNP_A.K_978844_6_1787313 | 6 | 1787313 | 5.24E-05 | 0.39465 | 299 | 0.132283782 | 0.035462269 |
| SNP_A.K_996193_6_5973917 | 6 | 5973917 | 5.25E-05 | 0.3194 | 299 | 0.132271163 | 0.035522249 |
| SNP_A.K_1364879_8_6050366 | 8 | 6050366 | 5.25E-05 | 0.1204 | 299 | 0.132263734 | 0.035552855 |
| SNP_A.K_990310_6_4709952 | 6 | 4709952 | 5.26E-05 | 0.20234 | 299 | 0.132251832 | 0.035603489 |
| SNP_A.K_995693_6_5947977 | 6 | 5947977 | 5.26E-05 | 0.34281 | 299 | 0.132250753 | 0.035603489 |
| SNP_A.K_1001522_6_6487092 | 6 | 6487092 | 5.27E-05 | 0.33779 | 299 | 0.1322461 | 0.035605668 |
| SNP_A.K_992581_6_5659693 | 6 | 5659693 | 5.27E-05 | 0.49833 | 299 | 0.13224427 | 0.035605668 |
| SNP_A.K_992583_6_5659700 | 6 | 5659700 | 5.27E-05 | 0.49833 | 299 | 0.13224427 | 0.035605668 |
| SNP_A.K_1001161_6_6449818 | 6 | 6449818 | 5.27E-05 | 0.31104 | 299 | 0.132239193 | 0.035620949 |
| SNP_A.K_1830742_11_10577129 | 11 | 10577129 | 5.28E-05 | 0.39298 | 299 | 0.132235661 | 0.035620949 |
| SNP_A.K_198079_1_31695460 | 1 | 31695460 | 5.28E-05 | 0.301 | 299 | 0.132235489 | 0.035620949 |
| SNP_A.K_1834781_11_10971546 | 11 | 10971546 | 5.28E-05 | 0.25251 | 299 | 0.132231474 | 0.035632244 |
| SNP_A.K_326924_2_10070831 | 2 | 10070831 | 5.29E-05 | 0.07525 | 299 | 0.132221215 | 0.03567911 |
| SNP_A.K_1600712_9_18822800 | 9 | 18822800 | 5.29E-05 | 0.12375 | 299 | 0.132215872 | 0.035698003 |
| SNP_A.K_1001806_6_6522087 | 6 | 6522087 | 5.32E-05 | 0.29933 | 299 | 0.132181257 | 0.03588429 |
| SNP_A.K_1001803_6_6521650 | 6 | 6521650 | 5.34E-05 | 0.33779 | 299 | 0.132164107 | 0.035971074 |
| SNP_A.K_1001256_6_6459260 | 6 | 6459260 | 5.35E-05 | 0.28261 | 299 | 0.132149077 | 0.036045868 |
| SNP_A.K_1830685_11_10574298 | 11 | 10574298 | 5.35E-05 | 0.46823 | 299 | 0.132146883 | 0.036046834 |
| SNP_A.K_481527_3_2743339 | 3 | 2743339 | 5.36E-05 | 0.03679 | 299 | 0.132134009 | 0.036109359 |
| SNP_A.K_978100_6_1647185 | 6 | 1647185 | 5.38E-05 | 0.26254 | 299 | 0.132115625 | 0.036203851 |
| SNP_A.K_978345_6_1679921 | 6 | 1679921 | 5.43E-05 | 0.25251 | 299 | 0.132061827 | 0.036504539 |
| SNP_A.K_307281_2_7391116 | 2 | 7391116 | 5.44E-05 | 0.26254 | 299 | 0.132050764 | 0.03655032 |

| | | | | | | | |
|---|---|---|---|---|---|---|---|
| SNP_A.K_456471_2_33041697 | 2 | 33041697 | 5.44E-05 | 0.08194 | 299 | 0.132049937 | 0.03655032 |
| SNP_A.K_710216_4_8322463 | 4 | 8322463 | 5.44E-05 | 0.09197 | 299 | 0.132042425 | 0.036578525 |
| SNP_A.K_977880_6_1610216 | 6 | 1610216 | 5.44E-05 | 0.26087 | 299 | 0.132041067 | 0.036578525 |
| SNP_A.K_326495_2_10015069 | 2 | 10015069 | 5.45E-05 | 0.07191 | 299 | 0.132036927 | 0.036590914 |
| SNP_A.K_996072_6_5960700 | 6 | 5960700 | 5.46E-05 | 0.29431 | 299 | 0.132024932 | 0.036649281 |
| SNP_A.K_201312_1_32162596 | 1 | 32162596 | 5.46E-05 | 0.43478 | 299 | 0.132021662 | 0.036656609 |
| SNP_A.K_978949_6_1822540 | 6 | 1822540 | 5.47E-05 | 0.46823 | 299 | 0.132012648 | 0.036697607 |
| SNP_A.K_709885_4_8300285 | 4 | 8300285 | 5.47E-05 | 0.07525 | 299 | 0.132010012 | 0.036701235 |
| SNP_A.K_1001280_6_6461013 | 6 | 6461013 | 5.48E-05 | 0.37458 | 299 | 0.132002901 | 0.036721559 |
| SNP_A.K_1144895_6_27617770 | 6 | 27617770 | 5.48E-05 | 0.04013 | 299 | 0.132002514 | 0.036721559 |
| SNP_A.K_902240_5_16463207 | 5 | 16463207 | 5.49E-05 | 0.05351 | 299 | 0.131991725 | 0.036773079 |
| SNP_A.K_307269_2_7389562 | 2 | 7389562 | 5.49E-05 | 0.27592 | 299 | 0.131987124 | 0.036788279 |
| SNP_A.K_1019185_6_9019957 | 6 | 9019957 | 5.50E-05 | 0.11706 | 299 | 0.131981967 | 0.036806759 |
| SNP_A.K_710131_4_8317731 | 4 | 8317731 | 5.50E-05 | 0.07525 | 299 | 0.131979126 | 0.036811627 |
| SNP_A.K_1595495_9_17675180 | 9 | 17675180 | 5.50E-05 | 0.03679 | 299 | 0.131974459 | 0.036827244 |
| SNP_A.K_709582_4_8274798 | 4 | 8274798 | 5.51E-05 | 0.08528 | 299 | 0.131966769 | 0.036852088 |
| SNP_A.K_992006_6_5608032 | 6 | 5608032 | 5.51E-05 | 0.18729 | 299 | 0.131966215 | 0.036852088 |
| SNP_A.K_751795_4_17707948 | 4 | 17707948 | 5.52E-05 | 0.02341 | 299 | 0.131959106 | 0.036882121 |
| SNP_A.K_1925823_11_24765641 | 11 | 24765641 | 5.52E-05 | 0.02676 | 299 | 0.131954773 | 0.036895814 |
| SNP_A.K_1001484_6_6483625 | 6 | 6483625 | 5.52E-05 | 0.29599 | 299 | 0.131950248 | 0.036910652 |
| SNP_A.K_992582_6_5659697 | 6 | 5659697 | 5.53E-05 | 0.5 | 299 | 0.131945686 | 0.036925712 |
| SNP_A.K_1924721_11_24664444 | 11 | 24664444 | 5.53E-05 | 0.02007 | 299 | 0.131939855 | 0.03694828 |
| SNP_A.K_456677_2_33104538 | 2 | 33104538 | 5.53E-05 | 0.02843 | 299 | 0.131937443 | 0.036950667 |
| SNP_A.K_1001980_6_6533310 | 6 | 6533310 | 5.54E-05 | 0.42809 | 299 | 0.131933094 | 0.0369645 |
| SNP_A.K_990246_6_4702953 | 6 | 4702953 | 5.54E-05 | 0.2107 | 299 | 0.131927447 | 0.036986015 |
| SNP_A.K_306733_2_7346454 | 2 | 7346454 | 5.57E-05 | 0.29097 | 299 | 0.131903585 | 0.037115431 |
| SNP_A.K_709502_4_8270852 | 4 | 8270852 | 5.58E-05 | 0.10368 | 299 | 0.13188884 | 0.037191077 |
| SNP_A.K_995528_6_5940256 | 6 | 5940256 | 5.59E-05 | 0.48829 | 299 | 0.131876186 | 0.037241996 |
| SNP_A.K_991238_6_4882965 | 6 | 4882965 | 5.59E-05 | 0.04682 | 299 | 0.131874531 | 0.037241996 |
| SNP_A.K_1001716_6_6508659 | 6 | 6508659 | 5.59E-05 | 0.31271 | 299 | 0.131874269 | 0.037241996 |
| SNP_A.K_195042_1_31039638 | 1 | 31039638 | 5.60E-05 | 0.37793 | 299 | 0.131868973 | 0.03726161 |
| SNP_A.K_978327_6_1678577 | 6 | 1678577 | 5.60E-05 | 0.26756 | 299 | 0.131864911 | 0.037273881 |
| SNP_A.K_1001711_6_6507958 | 6 | 6507958 | 5.63E-05 | 0.29933 | 299 | 0.131835594 | 0.037436972 |
| SNP_A.K_1830671_11_10573467 | 11 | 10573467 | 5.63E-05 | 0.36957 | 299 | 0.131827566 | 0.037472143 |
| SNP_A.K_181485_1_29141129 | 1 | 29141129 | 5.63E-05 | 0.43813 | 299 | 0.13182572 | 0.037472143 |
| SNP_A.K_978064_6_1644682 | 6 | 1644682 | 5.64E-05 | 0.21405 | 299 | 0.131822817 | 0.037477558 |
| SNP_A.K_1001583_6_6492984 | 6 | 6492984 | 5.64E-05 | 0.31104 | 299 | 0.131819628 | 0.037484698 |
| SNP_A.K_992571_6_5659468 | 6 | 5659468 | 5.65E-05 | 0.49833 | 299 | 0.131804263 | 0.037562821 |
| SNP_A.K_522889_3_10225662 | 3 | 10225662 | 5.66E-05 | 0.2291 | 299 | 0.131800899 | 0.037562821 |
| SNP_A.K_901334_5_16287066 | 5 | 16287066 | 5.66E-05 | 0.04013 | 299 | 0.131799223 | 0.037562821 |
| SNP_A.K_194465_1_30875707 | 1 | 30875707 | 5.66E-05 | 0.40803 | 299 | 0.131798617 | 0.037562821 |
| SNP_A.K_307498_2_7413097 | 2 | 7413097 | 5.67E-05 | 0.24415 | 299 | 0.131786461 | 0.03762388 |
| SNP_A.K_1364880_8_6050393 | 8 | 6050393 | 5.67E-05 | 0.11873 | 299 | 0.131783356 | 0.037630564 |
| SNP_A.K_708723_4_8089101 | 4 | 8089101 | 5.68E-05 | 0.09866 | 299 | 0.131776518 | 0.037659718 |
| SNP_A.K_673070_4_314722 | 4 | 314722 | 5.68E-05 | 0.1505 | 299 | 0.131771252 | 0.037668449 |
| SNP_A.K_218884_1_35377052 | 1 | 35377052 | 5.68E-05 | 0.16221 | 299 | 0.131771083 | 0.037668449 |
| SNP_A.K_978433_6_1691449 | 6 | 1691449 | 5.70E-05 | 0.26589 | 299 | 0.131755805 | 0.03774854 |
| SNP_A.K_1608714_9_20820151 | 9 | 20820151 | 5.70E-05 | 0.24749 | 299 | 0.131753224 | 0.037752106 |
| SNP_A.K_978451_6_1694719 | 6 | 1694719 | 5.72E-05 | 0.27759 | 299 | 0.131733135 | 0.037861488 |
| SNP_A.K_710133_4_8317751 | 4 | 8317751 | 5.73E-05 | 0.07692 | 299 | 0.131726191 | 0.037891492 |

| | | | | | | | |
|---|---|---|---|---|---|---|---|
| SNP_A.K_1001282_6_6461312 | 6 | 6461312 | 5.74E-05 | 0.36789 | 299 | 0.131714531 | 0.03795012 |
| SNP_A.K_1002034_6_6537591 | 6 | 6537591 | 5.75E-05 | 0.32107 | 299 | 0.131697438 | 0.038041878 |
| SNP_A.K_991236_6_4882861 | 6 | 4882861 | 5.76E-05 | 0.04013 | 299 | 0.131690643 | 0.038071128 |
| SNP_A.K_1713933_10_15025418 | 10 | 15025418 | 5.76E-05 | 0.25418 | 299 | 0.131684995 | 0.038089143 |
| SNP_A.K_1803408_11_6911996 | 11 | 6911996 | 5.76E-05 | 0.02174 | 299 | 0.131683712 | 0.038089143 |
| SNP_A.K_1000808_6_6417296 | 6 | 6417296 | 5.78E-05 | 0.3311 | 299 | 0.131670651 | 0.038156649 |
| SNP_A.K_1002050_6_6538246 | 6 | 6538246 | 5.78E-05 | 0.32107 | 299 | 0.13166483 | 0.038180058 |
| SNP_A.K_1183402_7_4541134 | 7 | 4541134 | 5.79E-05 | 0.4097 | 299 | 0.131657214 | 0.038214452 |
| SNP_A.K_308292_2_7488561 | 2 | 7488561 | 5.80E-05 | 0.20903 | 299 | 0.131645608 | 0.038262216 |
| SNP_A.K_991137_6_4831548 | 6 | 4831548 | 5.80E-05 | 0.04013 | 299 | 0.131643961 | 0.038262216 |
| SNP_A.K_1001326_6_6465288 | 6 | 6465288 | 5.80E-05 | 0.30602 | 299 | 0.131643454 | 0.038262216 |
| SNP_A.K_1831326_11_10619447 | 11 | 10619447 | 5.81E-05 | 0.11873 | 299 | 0.131636886 | 0.038281373 |
| SNP_A.K_996370_6_5995006 | 6 | 5995006 | 5.81E-05 | 0.47659 | 299 | 0.131636361 | 0.038281373 |
| SNP_A.K_995734_6_5949212 | 6 | 5949212 | 5.81E-05 | 0.40134 | 299 | 0.131631784 | 0.038297273 |
| SNP_A.K_1083969_6_17403949 | 6 | 17403949 | 5.81E-05 | 0.21237 | 299 | 0.131629699 | 0.038297923 |
| SNP_A.K_181159_1_29077823 | 1 | 29077823 | 5.82E-05 | 0.42809 | 299 | 0.131623627 | 0.038308764 |
| SNP_A.K_312487_2_7965672 | 2 | 7965672 | 5.82E-05 | 0.19064 | 299 | 0.131623473 | 0.038308764 |
| SNP_A.K_1804320_11_7061281 | 11 | 7061281 | 5.82E-05 | 0.01171 | 299 | 0.131620017 | 0.038308764 |
| SNP_A.K_1803414_11_6912303 | 11 | 6912303 | 5.82E-05 | 0.01171 | 299 | 0.131620017 | 0.038308764 |
| SNP_A.K_218481_1_35296610 | 1 | 35296610 | 5.83E-05 | 0.1689 | 299 | 0.13161521 | 0.038326108 |
| SNP_A.K_194457_1_30874598 | 1 | 30874598 | 5.83E-05 | 0.36622 | 299 | 0.131611914 | 0.038334198 |
| SNP_A.K_999248_6_6270673 | 6 | 6270673 | 5.83E-05 | 0.16555 | 299 | 0.131608858 | 0.038340824 |
| SNP_A.K_709952_4_8302863 | 4 | 8302863 | 5.84E-05 | 0.09532 | 299 | 0.131606259 | 0.038344134 |
| SNP_A.K_20610_1_3099378 | 1 | 3099378 | 5.84E-05 | 0.00502 | 299 | 0.13160437 | 0.038344134 |
| SNP_A.K_195523_1_31222507 | 1 | 31222507 | 5.84E-05 | 0.49164 | 299 | 0.131599146 | 0.038356347 |
| SNP_A.K_709802_4_8298031 | 4 | 8298031 | 5.84E-05 | 0.07525 | 299 | 0.131598433 | 0.038356347 |
| SNP_A.K_1001212_6_6455205 | 6 | 6455205 | 5.85E-05 | 0.28763 | 299 | 0.131592145 | 0.03836468 |
| SNP_A.K_1001425_6_6481372 | 6 | 6481372 | 5.85E-05 | 0.30769 | 299 | 0.131591931 | 0.03836468 |
| SNP_A.K_998869_6_6229807 | 6 | 6229807 | 5.85E-05 | 0.27926 | 299 | 0.13158955 | 0.03836468 |
| SNP_A.K_201232_1_32153804 | 1 | 32153804 | 5.85E-05 | 0.42809 | 299 | 0.131587697 | 0.03836468 |
| SNP_A.K_1383135_8_9131670 | 8 | 9131670 | 5.85E-05 | 0.27258 | 299 | 0.131587222 | 0.03836468 |
| SNP_A.K_978129_6_1649024 | 6 | 1649024 | 5.88E-05 | 0.1806 | 299 | 0.131560591 | 0.038508652 |
| SNP_A.K_1001465_6_6482468 | 6 | 6482468 | 5.88E-05 | 0.30602 | 299 | 0.131559862 | 0.038508652 |
| SNP_A.K_978390_6_1685715 | 6 | 1685715 | 5.88E-05 | 0.26923 | 299 | 0.131555426 | 0.038523858 |
| SNP_A.K_194459_1_30874622 | 1 | 30874622 | 5.89E-05 | 0.36957 | 299 | 0.131552755 | 0.038527419 |
| SNP_A.K_1000638_6_6402851 | 6 | 6402851 | 5.89E-05 | 0.35284 | 299 | 0.131550915 | 0.038527419 |
| SNP_A.K_709969_4_8303779 | 4 | 8303779 | 5.89E-05 | 0.07358 | 299 | 0.131543333 | 0.038549927 |
| SNP_A.K_709876_4_8300143 | 4 | 8300143 | 5.89E-05 | 0.07358 | 299 | 0.131543333 | 0.038549927 |
| SNP_A.K_1002025_6_6537424 | 6 | 6537424 | 5.90E-05 | 0.31271 | 299 | 0.131538397 | 0.038562691 |
| SNP_A.K_308535_2_7524767 | 2 | 7524767 | 5.90E-05 | 0.42977 | 299 | 0.131537335 | 0.038562691 |
| SNP_A.K_708722_4_8089099 | 4 | 8089099 | 5.91E-05 | 0.09866 | 299 | 0.131532373 | 0.038581193 |
| SNP_A.K_997905_6_6156521 | 6 | 6156521 | 5.91E-05 | 0.17057 | 299 | 0.131527673 | 0.038586099 |
| SNP_A.K_2065394_12_24383567 | 12 | 24383567 | 5.91E-05 | 0.03512 | 299 | 0.131527653 | 0.038586099 |
| SNP_A.K_710036_4_8310467 | 4 | 8310467 | 5.91E-05 | 0.07692 | 299 | 0.13152422 | 0.038595186 |
| SNP_A.K_687687_4_3571074 | 4 | 3571074 | 5.92E-05 | 0.10702 | 299 | 0.1315175 | 0.038624573 |
| SNP_A.K_309936_2_7688972 | 2 | 7688972 | 5.93E-05 | 0.10702 | 299 | 0.131505876 | 0.038671562 |
| SNP_A.K_1803735_11_6970719 | 11 | 6970719 | 5.93E-05 | 0.03512 | 299 | 0.131505329 | 0.038671562 |
| SNP_A.K_995731_6_5949184 | 6 | 5949184 | 5.93E-05 | 0.49498 | 299 | 0.131504021 | 0.038671562 |
| SNP_A.K_1924668_11_24662752 | 11 | 24662752 | 5.94E-05 | 0.02007 | 299 | 0.131495525 | 0.038712021 |
| SNP_A.K_992004_6_5607984 | 6 | 5607984 | 5.95E-05 | 0.2709 | 299 | 0.131487684 | 0.038737557 |

| | | | | | | | |
|---|---|---|---|---|---|---|---|
| SNP_A.K_194979_1_31019147 | 1 | 31019147 | 5.95E-05 | 0.44147 | 299 | 0.131487488 | 0.038737557 |
| SNP_A.K_1184234_7_4689402 | 7 | 4689402 | 5.95E-05 | 0.05518 | 299 | 0.131484634 | 0.038741417 |
| SNP_A.K_1001773_6_6519750 | 6 | 6519750 | 5.95E-05 | 0.30268 | 299 | 0.131482013 | 0.038741417 |
| SNP_A.K_218495_1_35297108 | 1 | 35297108 | 5.95E-05 | 0.17391 | 299 | 0.131480999 | 0.038741417 |
| SNP_A.K_708721_4_8089093 | 4 | 8089093 | 5.96E-05 | 0.09866 | 299 | 0.131477869 | 0.038748702 |
| SNP_A.K_725571_4_11822284 | 4 | 11822284 | 5.96E-05 | 0.0903 | 299 | 0.131473706 | 0.038761884 |
| SNP_A.K_977901_6_1617816 | 6 | 1617816 | 5.96E-05 | 0.26087 | 299 | 0.131471836 | 0.038761884 |
| SNP_A.K_996226_6_5976079 | 6 | 5976079 | 5.96E-05 | 0.38629 | 299 | 0.131469866 | 0.038761997 |
| SNP_A.K_710039_4_8311360 | 4 | 8311360 | 5.97E-05 | 0.07358 | 299 | 0.131463785 | 0.038768057 |
| SNP_A.K_978420_6_1689366 | 6 | 1689366 | 5.97E-05 | 0.26923 | 299 | 0.131461022 | 0.038768057 |
| SNP_A.K_1001494_6_6484010 | 6 | 6484010 | 5.97E-05 | 0.29933 | 299 | 0.131460888 | 0.038768057 |
| SNP_A.K_977870_6_1605936 | 6 | 1605936 | 5.97E-05 | 0.18562 | 299 | 0.131460335 | 0.038768057 |
| SNP_A.K_710536_4_8426700 | 4 | 8426700 | 5.98E-05 | 0.0786 | 299 | 0.131458571 | 0.038768057 |
| SNP_A.K_1608486_9_20763215 | 9 | 20763215 | 5.98E-05 | 0.29599 | 299 | 0.131457188 | 0.038768057 |
| SNP_A.K_218485_1_35296802 | 1 | 35296802 | 5.98E-05 | 0.16722 | 299 | 0.13145187 | 0.038788954 |
| SNP_A.K_978453_6_1694761 | 6 | 1694761 | 5.99E-05 | 0.27425 | 299 | 0.131447481 | 0.038796449 |
| SNP_A.K_1726583_10_16600997 | 10 | 16600997 | 5.99E-05 | 0.08361 | 299 | 0.131446769 | 0.038796449 |
| SNP_A.K_218033_1_35217472 | 1 | 35217472 | 6.01E-05 | 0.02007 | 299 | 0.131426804 | 0.038908425 |
| SNP_A.K_1526503_9_5069568 | 9 | 5069568 | 6.01E-05 | 0.02174 | 299 | 0.131424651 | 0.03890972 |
| SNP_A.K_1389907_8_10135203 | 8 | 10135203 | 6.02E-05 | 0.05853 | 299 | 0.131416955 | 0.03894044 |
| SNP_A.K_306944_2_7370256 | 2 | 7370256 | 6.02E-05 | 0.4097 | 299 | 0.131415828 | 0.03894044 |
| SNP_A.K_1001109_6_6444885 | 6 | 6444885 | 6.02E-05 | 0.38629 | 299 | 0.13141311 | 0.038945265 |
| SNP_A.K_992588_6_5659786 | 6 | 5659786 | 6.03E-05 | 0.5 | 299 | 0.131402029 | 0.039002236 |
| SNP_A.K_253132_1_40864632 | 1 | 40864632 | 6.03E-05 | 0.16054 | 299 | 0.131397633 | 0.039017545 |
| SNP_A.K_710035_4_8310403 | 4 | 8310403 | 6.04E-05 | 0.07692 | 299 | 0.131394347 | 0.03902594 |
| SNP_A.K_216399_1_34966419 | 1 | 34966419 | 6.05E-05 | 0.07692 | 299 | 0.131378186 | 0.039114817 |
| SNP_A.K_993152_6_5710655 | 6 | 5710655 | 6.06E-05 | 0.18227 | 299 | 0.131374433 | 0.039126161 |
| SNP_A.K_978462_6_1696048 | 6 | 1696048 | 6.06E-05 | 0.24582 | 299 | 0.131371691 | 0.039126514 |
| SNP_A.K_1001227_6_6456028 | 6 | 6456028 | 6.06E-05 | 0.37291 | 299 | 0.131370497 | 0.039126514 |
| SNP_A.K_709878_4_8300175 | 4 | 8300175 | 6.08E-05 | 0.07525 | 299 | 0.131354617 | 0.03921388 |
| SNP_A.K_992593_6_5660116 | 6 | 5660116 | 6.08E-05 | 0.49666 | 299 | 0.13134752 | 0.039246259 |
| SNP_A.K_900435_5_16188756 | 5 | 16188756 | 6.09E-05 | 0.06689 | 299 | 0.131344644 | 0.039252153 |
| SNP_A.K_1805292_11_7146168 | 11 | 7146168 | 6.09E-05 | 0.10535 | 299 | 0.131339313 | 0.039273469 |
| SNP_A.K_1601992_9_19102345 | 9 | 19102345 | 6.10E-05 | 0.0602 | 299 | 0.131329358 | 0.039313202 |
| SNP_A.K_1001568_6_6492304 | 6 | 6492304 | 6.10E-05 | 0.30268 | 299 | 0.131329122 | 0.039313202 |
| SNP_A.K_843882_5_1006233 | 5 | 1006233 | 6.11E-05 | 0.04181 | 299 | 0.131318514 | 0.039362375 |
| SNP_A.K_999116_6_6257653 | 6 | 6257653 | 6.11E-05 | 0.16054 | 299 | 0.131317439 | 0.039362375 |
| SNP_A.K_1004363_6_6876587 | 6 | 6876587 | 6.14E-05 | 0.34615 | 299 | 0.131283894 | 0.039554106 |
| SNP_A.K_307491_2_7412879 | 2 | 7412879 | 6.15E-05 | 0.26087 | 299 | 0.131282464 | 0.039554106 |
| SNP_A.K_978303_6_1671898 | 6 | 1671898 | 6.15E-05 | 0.24749 | 299 | 0.131280264 | 0.039554106 |
| SNP_A.K_523141_3_10260089 | 3 | 10260089 | 6.15E-05 | 0.22074 | 299 | 0.131279339 | 0.039554106 |
| SNP_A.K_1001078_6_6439943 | 6 | 6439943 | 6.17E-05 | 0.39465 | 299 | 0.131259118 | 0.039670032 |
| SNP_A.K_1602093_9_19123657 | 9 | 19123657 | 6.17E-05 | 0.04181 | 299 | 0.131256392 | 0.039675083 |
| SNP_A.K_1962888_12_5098184 | 12 | 5098184 | 6.18E-05 | 0.04348 | 299 | 0.131252081 | 0.039690204 |
| SNP_A.K_709829_4_8299088 | 4 | 8299088 | 6.19E-05 | 0.07692 | 299 | 0.131242296 | 0.039740127 |
| SNP_A.K_2078883_12_27031072 | 12 | 27031072 | 6.20E-05 | 0.02007 | 299 | 0.131227282 | 0.039822219 |
| SNP_A.K_2081243_12_27449747 | 12 | 27449747 | 6.20E-05 | 0.0301 | 299 | 0.131225544 | 0.039822219 |
| SNP_A.K_1002120_6_6541964 | 6 | 6541964 | 6.21E-05 | 0.32776 | 299 | 0.131213203 | 0.039882308 |
| SNP_A.K_1103427_6_20632795 | 6 | 20632795 | 6.22E-05 | 0.03846 | 299 | 0.131212268 | 0.039882308 |
| SNP_A.K_218526_1_35302712 | 1 | 35302712 | 6.22E-05 | 0.15385 | 299 | 0.131208269 | 0.039886395 |

| | | | | | | | |
|---|---|---|---|---|---|---|---|
| SNP_A.K_992714_6_5675130 | 6 | 5675130 | 6.22E-05 | 0.47157 | 299 | 0.131207776 | 0.039886395 |
| SNP_A.K_1001531_6_6487586 | 6 | 6487586 | 6.22E-05 | 0.29933 | 299 | 0.13120468 | 0.039892185 |
| SNP_A.K_218781_1_35351224 | 1 | 35351224 | 6.22E-05 | 0.16221 | 299 | 0.13120302 | 0.039892185 |
| SNP_A.K_978738_6_1772768 | 6 | 1772768 | 6.23E-05 | 0.3612 | 299 | 0.13120076 | 0.039894328 |
| SNP_A.K_312486_2_7965659 | 2 | 7965659 | 6.24E-05 | 0.08194 | 299 | 0.13119008 | 0.039927266 |
| SNP_A.K_753058_4_18150504 | 4 | 18150504 | 6.24E-05 | 0.27425 | 299 | 0.131189995 | 0.039927266 |
| SNP_A.K_978854_6_1788336 | 6 | 1788336 | 6.24E-05 | 0.36288 | 299 | 0.131189002 | 0.039927266 |
| SNP_A.K_1924730_11_24664729 | 11 | 24664729 | 6.24E-05 | 0.02007 | 299 | 0.131187574 | 0.039927266 |
| SNP_A.K_1608602_9_20792561 | 9 | 20792561 | 6.24E-05 | 0.24749 | 299 | 0.131185994 | 0.039927266 |
| SNP_A.K_673449_4_362758 | 4 | 362758 | 6.25E-05 | 0.0602 | 299 | 0.131179727 | 0.039945204 |
| SNP_A.K_286420_2_4764123 | 2 | 4764123 | 6.25E-05 | 0.07358 | 299 | 0.131179348 | 0.039945204 |
| SNP_A.K_1608496_9_20763808 | 9 | 20763808 | 6.26E-05 | 0.301 | 299 | 0.131170853 | 0.039987265 |
| SNP_A.K_1364856_8_6049652 | 8 | 6049652 | 6.26E-05 | 0.09365 | 299 | 0.131167746 | 0.039994875 |
| SNP_A.K_710418_4_8360802 | 4 | 8360802 | 6.27E-05 | 0.12375 | 299 | 0.131162249 | 0.040017794 |
| SNP_A.K_1162247_6_30626209 | 6 | 30626209 | 6.27E-05 | 0.2893 | 299 | 0.131155535 | 0.040048527 |
| SNP_A.K_218480_1_35296510 | 1 | 35296510 | 6.28E-05 | 0.23077 | 299 | 0.131147454 | 0.040088061 |
| SNP_A.K_993630_6_5780447 | 6 | 5780447 | 6.29E-05 | 0.29264 | 299 | 0.131139474 | 0.040126985 |
| SNP_A.K_1503680_9_1049850 | 9 | 1049850 | 6.29E-05 | 0.0602 | 299 | 0.131135912 | 0.040137561 |
| SNP_A.K_978306_6_1672070 | 6 | 1672070 | 6.30E-05 | 0.18227 | 299 | 0.131129456 | 0.040157521 |
| SNP_A.K_978279_6_1666920 | 6 | 1666920 | 6.30E-05 | 0.17726 | 299 | 0.131128978 | 0.040157521 |
| SNP_A.K_1804317_11_7061234 | 11 | 7061234 | 6.31E-05 | 0.01171 | 299 | 0.131117489 | 0.040215371 |
| SNP_A.K_1001146_6_6449374 | 6 | 6449374 | 6.31E-05 | 0.2893 | 299 | 0.131115271 | 0.040215371 |
| SNP_A.K_708831_4_8115405 | 4 | 8115405 | 6.31E-05 | 0.09365 | 299 | 0.131114245 | 0.040215371 |
| SNP_A.K_978309_6_1672380 | 6 | 1672380 | 6.32E-05 | 0.17559 | 299 | 0.131108012 | 0.0402432 |
| SNP_A.K_1001148_6_6449411 | 6 | 6449411 | 6.33E-05 | 0.29766 | 299 | 0.131101932 | 0.040270061 |
| SNP_A.K_990607_6_4728216 | 6 | 4728216 | 6.33E-05 | 0.37793 | 299 | 0.131095029 | 0.040297269 |
| SNP_A.K_307041_2_7374680 | 2 | 7374680 | 6.33E-05 | 0.2107 | 299 | 0.131093893 | 0.040297269 |
| SNP_A.K_218967_1_35389720 | 1 | 35389720 | 6.34E-05 | 0.15719 | 299 | 0.131086787 | 0.040330802 |
| SNP_A.K_1831299_11_10617499 | 11 | 10617499 | 6.35E-05 | 0.13545 | 299 | 0.131082026 | 0.040349218 |
| SNP_A.K_210165_1_33958333 | 1 | 33958333 | 6.35E-05 | 0.02341 | 299 | 0.13107726 | 0.040367683 |
| SNP_A.K_678657_4_993420 | 4 | 993420 | 6.35E-05 | 0.07191 | 299 | 0.13107411 | 0.040375712 |
| SNP_A.K_978452_6_1694730 | 6 | 1694730 | 6.36E-05 | 0.27425 | 299 | 0.131069032 | 0.040396211 |
| SNP_A.K_1001469_6_6482736 | 6 | 6482736 | 6.36E-05 | 0.28428 | 299 | 0.131066354 | 0.040401206 |
| SNP_A.K_709813_4_8298307 | 4 | 8298307 | 6.37E-05 | 0.07358 | 299 | 0.131059057 | 0.040434305 |
| SNP_A.K_871271_5_5373815 | 5 | 5373815 | 6.37E-05 | 0.2107 | 299 | 0.131057428 | 0.040434305 |
| SNP_A.K_709697_4_8282154 | 4 | 8282154 | 6.37E-05 | 0.09197 | 299 | 0.131055105 | 0.040437013 |
| SNP_A.K_1004966_6_6958803 | 6 | 6958803 | 6.38E-05 | 0.30435 | 299 | 0.131046641 | 0.040479499 |
| SNP_A.K_1004283_6_6862737 | 6 | 6862737 | 6.39E-05 | 0.17391 | 299 | 0.131034553 | 0.040545551 |
| SNP_A.K_277566_2_2951780 | 2 | 2951780 | 6.40E-05 | 0.08696 | 299 | 0.131031171 | 0.040555154 |
| SNP_A.K_1382891_8_9121263 | 8 | 9121263 | 6.40E-05 | 0.27258 | 299 | 0.131027631 | 0.040565792 |
| SNP_A.K_1001333_6_6466095 | 6 | 6466095 | 6.41E-05 | 0.28595 | 299 | 0.131020212 | 0.040601637 |
| SNP_A.K_1600936_9_18862695 | 9 | 18862695 | 6.42E-05 | 0.12876 | 299 | 0.131008446 | 0.040665811 |
| SNP_A.K_978955_6_1828735 | 6 | 1828735 | 6.42E-05 | 0.44649 | 299 | 0.13100628 | 0.040667541 |
| SNP_A.K_1399766_8_11032536 | 8 | 11032536 | 6.43E-05 | 0.07358 | 299 | 0.131002684 | 0.040678588 |
| SNP_A.K_307035_2_7374410 | 2 | 7374410 | 6.44E-05 | 0.2291 | 299 | 0.13099146 | 0.040739361 |
| SNP_A.K_978724_6_1771541 | 6 | 1771541 | 6.44E-05 | 0.44314 | 299 | 0.130985868 | 0.040763462 |
| SNP_A.K_1001538_6_6487981 | 6 | 6487981 | 6.46E-05 | 0.301 | 299 | 0.130975951 | 0.040815828 |
| SNP_A.K_1001424_6_6481306 | 6 | 6481306 | 6.47E-05 | 0.32776 | 299 | 0.130965715 | 0.04086866 |
| SNP_A.K_2030037_12_17237414 | 12 | 17237414 | 6.47E-05 | 0.16722 | 299 | 0.130962706 | 0.04086866 |
| SNP_A.K_389606_2_19886611 | 2 | 19886611 | 6.47E-05 | 0.0602 | 299 | 0.130962182 | 0.04086866 |

| | | | | | | | | |
|---|---|---|---|---|---|---|---|---|
| SNP_A.K_1600922_9_18862177 | 9 | 18862177 | 6.47E-05 | 0.17224 | 299 | 0.130958884 | | 0.040868756 |
| SNP_A.K_307209_2_7384979 | 2 | 7384979 | 6.47E-05 | 0.44816 | 299 | 0.130958377 | | 0.040868756 |
| SNP_A.K_1383138_8_9131767 | 8 | 9131767 | 6.48E-05 | 0.27592 | 299 | 0.130954206 | | 0.040879381 |
| SNP_A.K_139103_1_22285115 | 1 | 22285115 | 6.48E-05 | 0.13211 | 299 | 0.130952966 | | 0.040879381 |
| SNP_A.K_310109_2_7698068 | 2 | 7698068 | 6.48E-05 | 0.35452 | 299 | 0.130949416 | | 0.040890232 |
| SNP_A.K_1421832_8_14601562 | 8 | 14601562 | 6.49E-05 | 0.01338 | 299 | 0.130945993 | | 0.040900253 |
| SNP_A.K_995768_6_5950267 | 6 | 5950267 | 6.49E-05 | 0.48662 | 299 | 0.130941207 | | 0.040919211 |
| SNP_A.K_671553_4_163298 | 4 | 163298 | 6.49E-05 | 0.05853 | 299 | 0.130939153 | | 0.040920274 |
| SNP_A.K_1001486_6_6483641 | 6 | 6483641 | 6.50E-05 | 0.29933 | 299 | 0.130937165 | | 0.040920911 |
| SNP_A.K_1383035_8_9128069 | 8 | 9128069 | 6.51E-05 | 0.15719 | 299 | 0.13092758 | | 0.040971362 |
| SNP_A.K_1002031_6_6537546 | 6 | 6537546 | 6.51E-05 | 0.32107 | 299 | 0.130923148 | | 0.040987611 |
| SNP_A.K_1960654_12_4797089 | 12 | 4797089 | 6.51E-05 | 0.03177 | 299 | 0.130920537 | | 0.040987611 |
| SNP_A.K_709933_4_8302431 | 4 | 8302431 | 6.51E-05 | 0.09699 | 299 | 0.130919437 | | 0.040987611 |
| SNP_A.K_996821_6_6029884 | 6 | 6029884 | 6.53E-05 | 0.3796 | 299 | 0.13090055 | | 0.041090615 |
| SNP_A.K_307010_2_7372969 | 2 | 7372969 | 6.53E-05 | 0.24582 | 299 | 0.130899987 | | 0.041090615 |
| SNP_A.K_672587_4_264369 | 4 | 264369 | 6.55E-05 | 0.14047 | 299 | 0.130881511 | | 0.041199911 |
| SNP_A.K_498995_3_5793736 | 3 | 5793736 | 6.58E-05 | 0.03846 | 299 | 0.130860898 | | 0.041312224 |
| SNP_A.K_978075_6_1645459 | 6 | 1645459 | 6.58E-05 | 0.25084 | 299 | 0.13086074 | | 0.041312224 |
| SNP_A.K_326843_2_10060386 | 2 | 10060386 | 6.59E-05 | 0.07358 | 299 | 0.130847687 | | 0.041386171 |
| SNP_A.K_995349_6_5927522 | 6 | 5927522 | 6.59E-05 | 0.23077 | 299 | 0.130845147 | | 0.041390524 |
| SNP_A.K_709246_4_8242605 | 4 | 8242605 | 6.61E-05 | 0.1087 | 299 | 0.130831731 | | 0.041463474 |
| SNP_A.K_29237_1_4218802 | 1 | 4218802 | 6.61E-05 | 0.08361 | 299 | 0.130830385 | | 0.041463474 |
| SNP_A.K_1001312_6_6464052 | 6 | 6464052 | 6.61E-05 | 0.28595 | 299 | 0.130828151 | | 0.041465803 |
| SNP_A.K_533405_3_12166294 | 3 | 12166294 | 6.63E-05 | 0.04682 | 299 | 0.1308111 | | 0.041566663 |
| SNP_A.K_1001507_6_6485400 | 6 | 6485400 | 6.64E-05 | 0.29933 | 299 | 0.130802976 | | 0.04160789 |
| SNP_A.K_978465_6_1696692 | 6 | 1696692 | 6.64E-05 | 0.33612 | 299 | 0.130801149 | | 0.04160789 |
| SNP_A.K_1390037_8_10146112 | 8 | 10146112 | 6.66E-05 | 0.05518 | 299 | 0.130777181 | | 0.04174488 |
| SNP_A.K_978870_6_1793665 | 6 | 1793665 | 6.66E-05 | 0.31104 | 299 | 0.130776866 | | 0.04174488 |
| SNP_A.K_978165_6_1656674 | 6 | 1656674 | 6.67E-05 | 0.17559 | 299 | 0.130767344 | | 0.041796016 |
| SNP_A.K_1005052_6_6966455 | 6 | 6966455 | 6.68E-05 | 0.07525 | 299 | 0.130763863 | | 0.041797733 |
| SNP_A.K_992587_6_5659762 | 6 | 5659762 | 6.68E-05 | 0.49833 | 299 | 0.130763332 | | 0.041797733 |
| SNP_A.K_710046_4_8312233 | 4 | 8312233 | 6.69E-05 | 0.07358 | 299 | 0.130755336 | | 0.041838719 |
| SNP_A.K_195041_1_31039631 | 1 | 31039631 | 6.70E-05 | 0.41639 | 299 | 0.130748032 | | 0.041875108 |
| SNP_A.K_308787_2_7553888 | 2 | 7553888 | 6.70E-05 | 0.41472 | 299 | 0.130741024 | | 0.041883532 |
| SNP_A.K_278117_2_3027954 | 2 | 3027954 | 6.70E-05 | 0.08027 | 299 | 0.130740676 | | 0.041883532 |
| SNP_A.K_993450_6_5752285 | 6 | 5752285 | 6.70E-05 | 0.15217 | 299 | 0.130739712 | | 0.041883532 |
| SNP_A.K_1004994_6_6960871 | 6 | 6960871 | 6.70E-05 | 0.30936 | 299 | 0.130739276 | | 0.041883532 |
| SNP_A.K_710490_4_8424788 | 4 | 8424788 | 6.73E-05 | 0.07358 | 299 | 0.130717925 | | 0.042012652 |
| SNP_A.K_2059179_12_23313148 | 12 | 23313148 | 6.73E-05 | 0.02508 | 299 | 0.130714796 | | 0.042012652 |
| SNP_A.K_1924639_11_24661532 | 11 | 24661532 | 6.73E-05 | 0.04348 | 299 | 0.130714441 | | 0.042012652 |
| SNP_A.K_978685_6_1761242 | 6 | 1761242 | 6.73E-05 | 0.38462 | 299 | 0.130711918 | | 0.042017034 |
| SNP_A.K_1002042_6_6537823 | 6 | 6537823 | 6.74E-05 | 0.30602 | 299 | 0.13070811 | | 0.042030072 |
| SNP_A.K_306797_2_7351618 | 2 | 7351618 | 6.74E-05 | 0.26589 | 299 | 0.130704872 | | 0.042039285 |
| SNP_A.K_1000684_6_6411598 | 6 | 6411598 | 6.75E-05 | 0.28428 | 299 | 0.130697848 | | 0.042074006 |
| SNP_A.K_999053_6_6250072 | 6 | 6250072 | 6.76E-05 | 0.13211 | 299 | 0.130690323 | | 0.042112137 |
| SNP_A.K_29324_1_4224401 | 1 | 4224401 | 6.76E-05 | 0.08696 | 299 | 0.130687109 | | 0.042121215 |
| SNP_A.K_993659_6_5781869 | 6 | 5781869 | 6.77E-05 | 0.28595 | 299 | 0.130680265 | | 0.0421548 |
| SNP_A.K_235244_1_38434142 | 1 | 38434142 | 6.78E-05 | 0.02676 | 299 | 0.130671194 | | 0.042203473 |
| SNP_A.K_307534_2_7416090 | 2 | 7416090 | 6.78E-05 | 0.27425 | 299 | 0.130667088 | | 0.0422109 |
| SNP_A.K_456674_2_33104137 | 2 | 33104137 | 6.78E-05 | 0.02341 | 299 | 0.130665202 | | 0.0422109 |

| | | | | | | | |
|---|---|---|---|---|---|---|---|
| SNP_A.K_710028_4_8309717 | 4 | 8309717 | 6.79E-05 | 0.07525 | 299 | 0.130664496 | 0.0422109 |
| SNP_A.K_1924938_11_24680768 | 11 | 24680768 | 6.79E-05 | 0.02174 | 299 | 0.130658188 | 0.042240958 |
| SNP_A.K_1000886_6_6422273 | 6 | 6422273 | 6.80E-05 | 0.28595 | 299 | 0.130646683 | 0.042302159 |
| SNP_A.K_1001758_6_6515000 | 6 | 6515000 | 6.81E-05 | 0.30602 | 299 | 0.130645421 | 0.042302159 |
| SNP_A.K_990319_6_4710290 | 6 | 4710290 | 6.81E-05 | 0.20234 | 299 | 0.130643266 | 0.042304138 |
| SNP_A.K_306984_2_7371889 | 2 | 7371889 | 6.81E-05 | 0.24916 | 299 | 0.130640845 | 0.042307917 |
| SNP_A.K_990375_6_4712689 | 6 | 4712689 | 6.82E-05 | 0.20067 | 299 | 0.130636483 | 0.042315281 |
| SNP_A.K_218580_1_35319113 | 1 | 35319113 | 6.82E-05 | 0.29097 | 299 | 0.130636034 | 0.042315281 |
| SNP_A.K_307473_2_7409900 | 2 | 7409900 | 6.82E-05 | 0.33278 | 299 | 0.130629486 | 0.042335046 |
| SNP_A.K_523324_3_10297916 | 3 | 10297916 | 6.82E-05 | 0.21405 | 299 | 0.130629398 | 0.042335046 |
| SNP_A.K_534804_3_12562190 | 3 | 12562190 | 6.83E-05 | 0.02007 | 299 | 0.130621373 | 0.042376875 |
| SNP_A.K_1804394_11_7066859 | 11 | 7066859 | 6.84E-05 | 0.01338 | 299 | 0.13061813 | 0.042386267 |
| SNP_A.K_978136_6_1650499 | 6 | 1650499 | 6.86E-05 | 0.1689 | 299 | 0.130592426 | 0.04254851 |
| SNP_A.K_1001262_6_6459770 | 6 | 6459770 | 6.88E-05 | 0.39967 | 299 | 0.130578732 | 0.04262927 |
| SNP_A.K_310991_2_7785307 | 2 | 7785307 | 6.90E-05 | 0.19231 | 299 | 0.130556716 | 0.042767186 |
| SNP_A.K_307030_2_7374032 | 2 | 7374032 | 6.91E-05 | 0.22742 | 299 | 0.13055262 | 0.042777536 |
| SNP_A.K_709831_4_8299168 | 4 | 8299168 | 6.91E-05 | 0.07525 | 299 | 0.130551492 | 0.042777536 |
| SNP_A.K_691046_4_4526203 | 4 | 4526203 | 6.92E-05 | 0.05853 | 299 | 0.130545241 | 0.042806204 |
| SNP_A.K_1001706_6_6507517 | 6 | 6507517 | 6.92E-05 | 0.30268 | 299 | 0.130543599 | 0.042806204 |
| SNP_A.K_306753_2_7347883 | 2 | 7347883 | 6.92E-05 | 0.35619 | 299 | 0.130539003 | 0.042825013 |
| SNP_A.K_1004983_6_6959674 | 6 | 6959674 | 6.93E-05 | 0.31438 | 299 | 0.130535217 | 0.042838271 |
| SNP_A.K_28837_1_4169679 | 1 | 4169679 | 6.93E-05 | 0.04849 | 299 | 0.130530445 | 0.042858302 |
| SNP_A.K_307666_2_7439471 | 2 | 7439471 | 6.94E-05 | 0.26756 | 299 | 0.130524541 | 0.042886131 |
| SNP_A.K_1383010_8_9127220 | 8 | 9127220 | 6.94E-05 | 0.26923 | 299 | 0.130519496 | 0.042908081 |
| SNP_A.K_710333_4_8336697 | 4 | 8336697 | 6.96E-05 | 0.07692 | 299 | 0.130506634 | 0.042983849 |
| SNP_A.K_1925185_11_24711727 | 11 | 24711727 | 6.97E-05 | 0.02007 | 299 | 0.130495841 | 0.043045479 |
| SNP_A.K_1383030_8_9127826 | 8 | 9127826 | 6.98E-05 | 0.30602 | 299 | 0.130486621 | 0.043060497 |
| SNP_A.K_1002011_6_6537068 | 6 | 6537068 | 6.98E-05 | 0.29933 | 299 | 0.130486575 | 0.043060497 |
| SNP_A.K_2078688_12_27009239 | 12 | 27009239 | 6.98E-05 | 0.02007 | 299 | 0.130484441 | 0.043060497 |
| SNP_A.K_709201_4_8238580 | 4 | 8238580 | 6.98E-05 | 0.09532 | 299 | 0.130484396 | 0.043060497 |
| SNP_A.K_2078570_12_26996490 | 12 | 26996490 | 6.98E-05 | 0.02007 | 299 | 0.130484341 | 0.043060497 |
| SNP_A.K_760869_4_19812951 | 4 | 19812951 | 6.99E-05 | 0.04348 | 299 | 0.130481429 | 0.043060497 |
| SNP_A.K_725432_4_11813682 | 4 | 11813682 | 6.99E-05 | 0.09699 | 299 | 0.130478501 | 0.043060497 |
| SNP_A.K_673338_4_349585 | 4 | 349585 | 6.99E-05 | 0.09532 | 299 | 0.130477959 | 0.043060497 |
| SNP_A.K_192889_1_30593972 | 1 | 30593972 | 6.99E-05 | 0.5 | 299 | 0.130477027 | 0.043060497 |
| SNP_A.K_1001483_6_6483425 | 6 | 6483425 | 7.00E-05 | 0.30268 | 299 | 0.130466676 | 0.043119231 |
| SNP_A.K_1001617_6_6498514 | 6 | 6498514 | 7.01E-05 | 0.40301 | 299 | 0.130462706 | 0.043133909 |
| SNP_A.K_1830600_11_10569524 | 11 | 10569524 | 7.01E-05 | 0.17726 | 299 | 0.130459761 | 0.043141521 |
| SNP_A.K_996619_6_6012312 | 6 | 6012312 | 7.02E-05 | 0.39967 | 299 | 0.130450356 | 0.043190309 |
| SNP_A.K_1005592_6_7028921 | 6 | 7028921 | 7.03E-05 | 0.32609 | 299 | 0.130447658 | 0.043190309 |
| SNP_A.K_1001543_6_6488610 | 6 | 6488610 | 7.03E-05 | 0.30769 | 299 | 0.130445592 | 0.043190309 |
| SNP_A.K_218899_1_35380756 | 1 | 35380756 | 7.03E-05 | 0.16388 | 299 | 0.130445335 | 0.043190309 |
| SNP_A.K_990256_6_4703429 | 6 | 4703429 | 7.03E-05 | 0.16388 | 299 | 0.130440934 | 0.043208029 |
| SNP_A.K_805664_4_29334216 | 4 | 29334216 | 7.04E-05 | 0.0301 | 299 | 0.130435669 | 0.043231741 |
| SNP_A.K_990317_6_4710279 | 6 | 4710279 | 7.04E-05 | 0.20401 | 299 | 0.130431037 | 0.043251083 |
| SNP_A.K_1025773_6_10085549 | 6 | 10085549 | 7.05E-05 | 0.03512 | 299 | 0.130426435 | 0.043270232 |
| SNP_A.K_307163_2_7382840 | 2 | 7382840 | 7.05E-05 | 0.26421 | 299 | 0.130423313 | 0.043279127 |
| SNP_A.K_1001329_6_6466014 | 6 | 6466014 | 7.06E-05 | 0.30602 | 299 | 0.130418362 | 0.043297893 |
| SNP_A.K_978326_6_1678478 | 6 | 1678478 | 7.06E-05 | 0.2709 | 299 | 0.13041693 | 0.043297893 |
| SNP_A.K_277247_2_2932074 | 2 | 2932074 | 7.07E-05 | 0.07525 | 299 | 0.13041227 | 0.043317482 |

| | | | | | | | |
|---|---|---|---|---|---|---|---|
| SNP_A.K_1000988_6_6434182 | 6 | 6434182 | 7.08E-05 | 0.33445 | 299 | 0.130400635 | 0.043375013 |
| SNP_A.K_1432903_8_16598101 | 8 | 16598101 | 7.08E-05 | 0.10368 | 299 | 0.130400315 | 0.043375013 |
| SNP_A.K_2078561_12_26996305 | 12 | 26996305 | 7.08E-05 | 0.09866 | 299 | 0.130398022 | 0.043378186 |
| SNP_A.K_709695_4_8281716 | 4 | 8281716 | 7.09E-05 | 0.08696 | 299 | 0.130386058 | 0.043442157 |
| SNP_A.K_993223_6_5722509 | 6 | 5722509 | 7.10E-05 | 0.24247 | 299 | 0.130383457 | 0.043442157 |
| SNP_A.K_1382988_8_9125963 | 8 | 9125963 | 7.10E-05 | 0.27258 | 299 | 0.130383322 | 0.043442157 |
| SNP_A.K_1364861_8_6049795 | 8 | 6049795 | 7.13E-05 | 0.1204 | 299 | 0.130356131 | 0.043619091 |
| SNP_A.K_1364932_8_6060163 | 8 | 6060163 | 7.13E-05 | 0.1204 | 299 | 0.13035322 | 0.043626621 |
| SNP_A.K_995660_6_5946669 | 6 | 5946669 | 7.14E-05 | 0.05184 | 299 | 0.130348001 | 0.043650307 |
| SNP_A.K_311098_2_7789692 | 2 | 7789692 | 7.14E-05 | 0.19398 | 299 | 0.130343547 | 0.043668655 |
| SNP_A.K_990393_6_4715003 | 6 | 4715003 | 7.15E-05 | 0.45485 | 299 | 0.130339935 | 0.043674202 |
| SNP_A.K_1365370_8_6107312 | 8 | 6107312 | 7.15E-05 | 0.09699 | 299 | 0.130339092 | 0.043674202 |
| SNP_A.K_307429_2_7404088 | 2 | 7404088 | 7.15E-05 | 0.25585 | 299 | 0.130333997 | 0.043697059 |
| SNP_A.K_523295_3_10290840 | 3 | 10290840 | 7.16E-05 | 0.21572 | 299 | 0.130331665 | 0.043700574 |
| SNP_A.K_709226_4_8241858 | 4 | 8241858 | 7.17E-05 | 0.0903 | 299 | 0.130317935 | 0.043784019 |
| SNP_A.K_312167_2_7918709 | 2 | 7918709 | 7.19E-05 | 0.33278 | 299 | 0.130305399 | 0.043859236 |
| SNP_A.K_793835_4_26300907 | 4 | 26300907 | 7.19E-05 | 0.03344 | 299 | 0.130303542 | 0.043859435 |
| SNP_A.K_995630_6_5944101 | 6 | 5944101 | 7.20E-05 | 0.45485 | 299 | 0.130296549 | 0.04388857 |
| SNP_A.K_1000683_6_6411592 | 6 | 6411592 | 7.20E-05 | 0.2893 | 299 | 0.130295745 | 0.04388857 |
| SNP_A.K_207451_1_33404904 | 1 | 33404904 | 7.20E-05 | 0.02174 | 299 | 0.130293436 | 0.043891957 |
| SNP_A.K_1001641_6_6499888 | 6 | 6499888 | 7.21E-05 | 0.46154 | 299 | 0.130284423 | 0.043942562 |
| SNP_A.K_992586_6_5659756 | 6 | 5659756 | 7.22E-05 | 0.5 | 299 | 0.130279603 | 0.043963659 |
| SNP_A.K_992579_6_5659615 | 6 | 5659615 | 7.24E-05 | 0.5 | 299 | 0.130262104 | 0.044074275 |
| SNP_A.K_77910_1_10906282 | 1 | 10906282 | 7.25E-05 | 0.29599 | 299 | 0.130255019 | 0.044111459 |
| SNP_A.K_1389916_8_10135422 | 8 | 10135422 | 7.25E-05 | 0.0602 | 299 | 0.130248291 | 0.044138356 |
| SNP_A.K_1001286_6_6461809 | 6 | 6461809 | 7.26E-05 | 0.28595 | 299 | 0.130245988 | 0.044138356 |
| SNP_A.K_201153_1_32138883 | 1 | 32138883 | 7.26E-05 | 0.42977 | 299 | 0.130245747 | 0.044138356 |
| SNP_A.K_728240_4_12294071 | 4 | 12294071 | 7.26E-05 | 0.21739 | 299 | 0.130243648 | 0.044140311 |
| SNP_A.K_29061_1_4201884 | 1 | 4201884 | 7.27E-05 | 0.14381 | 299 | 0.130233083 | 0.044202231 |
| SNP_A.K_709015_4_8212878 | 4 | 8212878 | 7.30E-05 | 0.09699 | 299 | 0.130211225 | 0.044344467 |
| SNP_A.K_978436_6_1693411 | 6 | 1693411 | 7.30E-05 | 0.25418 | 299 | 0.130205917 | 0.044368665 |
| SNP_A.K_1004291_6_6863017 | 6 | 6863017 | 7.30E-05 | 0.36789 | 299 | 0.130204182 | 0.044368665 |
| SNP_A.K_456510_2_33052726 | 2 | 33052726 | 7.32E-05 | 0.02341 | 299 | 0.130193286 | 0.044433027 |
| SNP_A.K_710041_4_8311863 | 4 | 8311863 | 7.32E-05 | 0.07358 | 299 | 0.130191503 | 0.044433027 |
| SNP_A.K_996823_6_6029917 | 6 | 6029917 | 7.32E-05 | 0.3796 | 299 | 0.130187237 | 0.044450463 |
| SNP_A.K_978286_6_1669137 | 6 | 1669137 | 7.33E-05 | 0.25585 | 299 | 0.130182023 | 0.044474677 |
| SNP_A.K_1002049_6_6538171 | 6 | 6538171 | 7.34E-05 | 0.30769 | 299 | 0.130177526 | 0.044493794 |
| SNP_A.K_277525_2_2949239 | 2 | 2949239 | 7.35E-05 | 0.07525 | 299 | 0.130165134 | 0.044569309 |
| SNP_A.K_757998_4_19215066 | 4 | 19215066 | 7.36E-05 | 0.00502 | 299 | 0.130159772 | 0.044594657 |
| SNP_A.K_978537_6_1719932 | 6 | 1719932 | 7.37E-05 | 0.26756 | 299 | 0.130145801 | 0.0446709 |
| SNP_A.K_278279_2_3049113 | 2 | 3049113 | 7.37E-05 | 0.07525 | 299 | 0.130145488 | 0.0446709 |
| SNP_A.K_995675_6_5947207 | 6 | 5947207 | 7.38E-05 | 0.37124 | 299 | 0.130141798 | 0.044680242 |
| SNP_A.K_1004280_6_6862136 | 6 | 6862136 | 7.38E-05 | 0.40134 | 299 | 0.130138985 | 0.044680242 |
| SNP_A.K_2079044_12_27050415 | 12 | 27050415 | 7.38E-05 | 0.05351 | 299 | 0.130138742 | 0.044680242 |
| SNP_A.K_524473_3_10507188 | 3 | 10507188 | 7.39E-05 | 0.00502 | 299 | 0.130132552 | 0.044711615 |
| SNP_A.K_990287_6_4704907 | 6 | 4704907 | 7.40E-05 | 0.16722 | 299 | 0.130124299 | 0.044737762 |
| SNP_A.K_1383185_8_9133665 | 8 | 9133665 | 7.40E-05 | 0.27926 | 299 | 0.130123471 | 0.044737762 |
| SNP_A.K_1383184_8_9133661 | 8 | 9133661 | 7.40E-05 | 0.27926 | 299 | 0.130123471 | 0.044737762 |
| SNP_A.K_978662_6_1751718 | 6 | 1751718 | 7.41E-05 | 0.29097 | 299 | 0.130119101 | 0.044756126 |
| SNP_A.K_673206_4_333457 | 4 | 333457 | 7.43E-05 | 0.15217 | 299 | 0.130100983 | 0.044854422 |

| | | | | | | | |
|---|---|---|---|---|---|---|---|
| SNP_A.K_375081_2_17933736 | 2 | 17933736 | 7.43E-05 | 0.04515 | 299 | 0.130100712 | 0.044854422 |
| SNP_A.K_196656_1_31473294 | 1 | 31473294 | 7.43E-05 | 0.31773 | 299 | 0.130099992 | 0.044854422 |
| SNP_A.K_2058944_12_23271722 | 12 | 23271722 | 7.44E-05 | 0.01839 | 299 | 0.130093671 | 0.044886895 |
| SNP_A.K_29303_1_4222831 | 1 | 4222831 | 7.45E-05 | 0.09699 | 299 | 0.130081594 | 0.044960886 |
| SNP_A.K_690127_4_4406786 | 4 | 4406786 | 7.46E-05 | 0.03679 | 299 | 0.130075756 | 0.044989959 |
| SNP_A.K_909063_5_17581001 | 5 | 17581001 | 7.48E-05 | 0.04013 | 299 | 0.130053457 | 0.045138063 |
| SNP_A.K_462837_2_34434580 | 2 | 34434580 | 7.49E-05 | 0.04849 | 299 | 0.130043969 | 0.045189625 |
| SNP_A.K_1383140_8_9131871 | 8 | 9131871 | 7.50E-05 | 0.27425 | 299 | 0.130042727 | 0.045189625 |
| SNP_A.K_978348_6_1680899 | 6 | 1680899 | 7.50E-05 | 0.35953 | 299 | 0.130037725 | 0.045203687 |
| SNP_A.K_218489_1_35296904 | 1 | 35296904 | 7.50E-05 | 0.16722 | 299 | 0.130037177 | 0.045203687 |
| SNP_A.K_710393_4_8339904 | 4 | 8339904 | 7.52E-05 | 0.07525 | 299 | 0.130026762 | 0.045266169 |
| SNP_A.K_307475_2_7409949 | 2 | 7409949 | 7.54E-05 | 0.25418 | 299 | 0.130003864 | 0.045419586 |
| SNP_A.K_307488_2_7412777 | 2 | 7412777 | 7.56E-05 | 0.25418 | 299 | 0.129986107 | 0.045535972 |
| SNP_A.K_990251_6_4703307 | 6 | 4703307 | 7.59E-05 | 0.17726 | 299 | 0.12996693 | 0.045663066 |
| SNP_A.K_1924859_11_24671201 | 11 | 24671201 | 7.59E-05 | 0.02007 | 299 | 0.129963707 | 0.045673477 |
| SNP_A.K_77577_1_10868646 | 1 | 10868646 | 7.61E-05 | 0.27759 | 299 | 0.129952603 | 0.045730933 |
| SNP_A.K_760918_4_19817420 | 4 | 19817420 | 7.61E-05 | 0.01672 | 299 | 0.129952268 | 0.045730933 |
| SNP_A.K_990347_6_4711819 | 6 | 4711819 | 7.61E-05 | 0.16221 | 299 | 0.129947189 | 0.045754997 |
| SNP_A.K_1602004_9_19102864 | 9 | 19102864 | 7.62E-05 | 0.12542 | 299 | 0.129941273 | 0.045785217 |
| SNP_A.K_752436_4_17906178 | 4 | 17906178 | 7.63E-05 | 0.06522 | 299 | 0.129932301 | 0.045822094 |
| SNP_A.K_709863_4_8299769 | 4 | 8299769 | 7.63E-05 | 0.07692 | 299 | 0.129931541 | 0.045822094 |
| SNP_A.K_709368_4_8259022 | 4 | 8259022 | 7.63E-05 | 0.09699 | 299 | 0.129930859 | 0.045822094 |
| SNP_A.K_709991_4_8304676 | 4 | 8304676 | 7.64E-05 | 0.09197 | 299 | 0.129923654 | 0.045861859 |
| SNP_A.K_885770_5_8585545 | 5 | 8585545 | 7.65E-05 | 0.03846 | 299 | 0.129919387 | 0.045880037 |
| SNP_A.K_978585_6_1733945 | 6 | 1733945 | 7.65E-05 | 0.27425 | 299 | 0.129914652 | 0.045901674 |
| SNP_A.K_690553_4_4450554 | 4 | 4450554 | 7.65E-05 | 0.04515 | 299 | 0.129912624 | 0.045903382 |
| SNP_A.K_673438_4_361780 | 4 | 361780 | 7.66E-05 | 0.1505 | 299 | 0.129905476 | 0.045920448 |
| SNP_A.K_201302_1_32161653 | 1 | 32161653 | 7.66E-05 | 0.43478 | 299 | 0.129904641 | 0.045920448 |
| SNP_A.K_1383155_8_9132243 | 8 | 9132243 | 7.67E-05 | 0.27425 | 299 | 0.129903412 | 0.045920448 |
| SNP_A.K_978810_6_1783455 | 6 | 1783455 | 7.67E-05 | 0.32274 | 299 | 0.129903128 | 0.045920448 |
| SNP_A.K_456210_2_32978883 | 2 | 32978883 | 7.68E-05 | 0.12542 | 299 | 0.12988964 | 0.046006691 |
| SNP_A.K_1534507_9_5994196 | 9 | 5994196 | 7.69E-05 | 0.09365 | 299 | 0.129883843 | 0.046036255 |
| SNP_A.K_2078523_12_26994309 | 12 | 26994309 | 7.70E-05 | 0.01839 | 299 | 0.129877093 | 0.046072889 |
| SNP_A.K_2079977_12_27187455 | 12 | 27187455 | 7.70E-05 | 0.02508 | 299 | 0.129873466 | 0.046079924 |
| SNP_A.K_710332_4_8336677 | 4 | 8336677 | 7.70E-05 | 0.07692 | 299 | 0.129872559 | 0.046079924 |
| SNP_A.K_1017829_6_8736472 | 6 | 8736472 | 7.71E-05 | 0.03177 | 299 | 0.129865746 | 0.04611707 |
| SNP_A.K_673135_4_322170 | 4 | 322170 | 7.72E-05 | 0.14883 | 299 | 0.129862763 | 0.046125896 |
| SNP_A.K_978272_6_1666603 | 6 | 1666603 | 7.72E-05 | 0.27926 | 299 | 0.129855807 | 0.04616415 |
| SNP_A.K_977889_6_1612775 | 6 | 1612775 | 7.73E-05 | 0.18562 | 299 | 0.129852927 | 0.046172235 |
| SNP_A.K_1534458_9_5987012 | 9 | 5987012 | 7.73E-05 | 0.13043 | 299 | 0.129848238 | 0.046193123 |
| SNP_A.K_978085_6_1646532 | 6 | 1646532 | 7.74E-05 | 0.1806 | 299 | 0.129846531 | 0.046193123 |
| SNP_A.K_498011_3_5645820 | 3 | 5645820 | 7.74E-05 | 0.03846 | 299 | 0.129842321 | 0.046197837 |
| SNP_A.K_498438_3_5706397 | 3 | 5706397 | 7.74E-05 | 0.03846 | 299 | 0.129842321 | 0.046197837 |
| SNP_A.K_1001478_6_6483168 | 6 | 6483168 | 7.75E-05 | 0.2893 | 299 | 0.129835881 | 0.046232342 |
| SNP_A.K_1001230_6_6456577 | 6 | 6456577 | 7.76E-05 | 0.37625 | 299 | 0.129825588 | 0.046295494 |
| SNP_A.K_710047_4_8312669 | 4 | 8312669 | 7.77E-05 | 0.07525 | 299 | 0.129821729 | 0.046301869 |
| SNP_A.K_996188_6_5973569 | 6 | 5973569 | 7.77E-05 | 0.47659 | 299 | 0.12982116 | 0.046301869 |
| SNP_A.K_1001455_6_6482052 | 6 | 6482052 | 7.78E-05 | 0.29264 | 299 | 0.129809836 | 0.046360021 |
| SNP_A.K_374899_2_17914649 | 2 | 17914649 | 7.78E-05 | 0.04181 | 299 | 0.12980977 | 0.046360021 |
| SNP_A.K_992578_6_5659599 | 6 | 5659599 | 7.79E-05 | 0.49498 | 299 | 0.129805493 | 0.046378576 |

| | | | | | | | |
|---|---|---|---|---|---|---|---|
| SNP_A.K_306939_2_7369641 | 2 | 7369641 | 7.79E-05 | 0.24247 | 299 | 0.129801915 | 0.046391944 |
| SNP_A.K_307423_2_7403730 | 2 | 7403730 | 7.80E-05 | 0.43478 | 299 | 0.129798966 | 0.04640063 |
| SNP_A.K_990281_6_4704450 | 6 | 4704450 | 7.80E-05 | 0.18729 | 299 | 0.129793242 | 0.046429994 |
| SNP_A.K_1830837_11_10584295 | 11 | 10584295 | 7.82E-05 | 0.40301 | 299 | 0.129778557 | 0.046516129 |
| SNP_A.K_978398_6_1686655 | 6 | 1686655 | 7.82E-05 | 0.25251 | 299 | 0.129777978 | 0.046516129 |
| SNP_A.K_995644_6_5946130 | 6 | 5946130 | 7.83E-05 | 0.38127 | 299 | 0.129775183 | 0.046516129 |
| SNP_A.K_1382903_8_9121485 | 8 | 9121485 | 7.83E-05 | 0.27759 | 299 | 0.129774113 | 0.046516129 |
| SNP_A.K_994839_6_5893953 | 6 | 5893953 | 7.83E-05 | 0.31271 | 299 | 0.129772791 | 0.046516129 |
| SNP_A.K_1005039_6_6964303 | 6 | 6964303 | 7.83E-05 | 0.41304 | 299 | 0.129769668 | 0.046526166 |
| SNP_A.K_709639_4_8278928 | 4 | 8278928 | 7.85E-05 | 0.09699 | 299 | 0.129757149 | 0.046606436 |
| SNP_A.K_607341_3_25613137 | 3 | 25613137 | 7.85E-05 | 0.08528 | 299 | 0.129754904 | 0.046609936 |
| SNP_A.K_1383008_8_9127083 | 8 | 9127083 | 7.85E-05 | 0.27926 | 299 | 0.129752382 | 0.046615508 |
| SNP_A.K_1382832_8_9118497 | 8 | 9118497 | 7.86E-05 | 0.27759 | 299 | 0.129750029 | 0.046619825 |
| SNP_A.K_374698_2_17874706 | 2 | 17874706 | 7.87E-05 | 0.04181 | 299 | 0.129737334 | 0.046701591 |
| SNP_A.K_194469_1_30876557 | 1 | 30876557 | 7.88E-05 | 0.38127 | 299 | 0.129729866 | 0.046744276 |
| SNP_A.K_710528_4_8426482 | 4 | 8426482 | 7.89E-05 | 0.0786 | 299 | 0.12972534 | 0.046764922 |
| SNP_A.K_710125_4_8317518 | 4 | 8317518 | 7.90E-05 | 0.08194 | 299 | 0.12971227 | 0.046836445 |
| SNP_A.K_307490_2_7412869 | 2 | 7412869 | 7.91E-05 | 0.25084 | 299 | 0.129711243 | 0.046836445 |
| SNP_A.K_1608481_9_20762678 | 9 | 20762678 | 7.91E-05 | 0.29766 | 299 | 0.129708929 | 0.046836445 |
| SNP_A.K_374680_2_17871556 | 2 | 17871556 | 7.91E-05 | 0.04348 | 299 | 0.129708694 | 0.046836445 |
| SNP_A.K_709583_4_8274808 | 4 | 8274808 | 7.91E-05 | 0.08696 | 299 | 0.129706951 | 0.046836445 |
| SNP_A.K_83728_1_11508046 | 1 | 11508046 | 7.92E-05 | 0.19064 | 299 | 0.129702614 | 0.046855738 |
| SNP_A.K_201192_1_32147880 | 1 | 32147880 | 7.92E-05 | 0.18395 | 299 | 0.129700001 | 0.046862069 |
| SNP_A.K_1005259_6_7000081 | 6 | 7000081 | 7.93E-05 | 0.29264 | 299 | 0.1296893 | 0.046922132 |
| SNP_A.K_991323_6_4929340 | 6 | 4929340 | 7.93E-05 | 0.03512 | 299 | 0.129688479 | 0.046922132 |
| SNP_A.K_992591_6_5660018 | 6 | 5660018 | 7.94E-05 | 0.49498 | 299 | 0.129683551 | 0.046932599 |
| SNP_A.K_992592_6_5660023 | 6 | 5660023 | 7.94E-05 | 0.49498 | 299 | 0.129683551 | 0.046932599 |
| SNP_A.K_306783_2_7350081 | 2 | 7350081 | 7.95E-05 | 0.2709 | 299 | 0.129678918 | 0.046954183 |
| SNP_A.K_900709_5_16208228 | 5 | 16208228 | 7.95E-05 | 0.04348 | 299 | 0.129675951 | 0.046963217 |
| SNP_A.K_1001341_6_6466246 | 6 | 6466246 | 7.96E-05 | 0.28595 | 299 | 0.129672422 | 0.046976499 |
| SNP_A.K_978475_6_1699335 | 6 | 1699335 | 7.96E-05 | 0.26421 | 299 | 0.129669375 | 0.046986152 |
| SNP_A.K_307247_2_7387909 | 2 | 7387909 | 7.97E-05 | 0.2893 | 299 | 0.129660423 | 0.047028037 |
| SNP_A.K_308425_2_7512172 | 2 | 7512172 | 7.97E-05 | 0.42977 | 299 | 0.129659406 | 0.047028037 |
| SNP_A.K_1001214_6_6455275 | 6 | 6455275 | 7.97E-05 | 0.29264 | 299 | 0.129658525 | 0.047028037 |
| SNP_A.K_77851_1_10901266 | 1 | 10901266 | 7.99E-05 | 0.2893 | 299 | 0.129647411 | 0.047098663 |
| SNP_A.K_996674_6_6016494 | 6 | 6016494 | 8.00E-05 | 0.48328 | 299 | 0.129639367 | 0.047146167 |
| SNP_A.K_1004140_6_6836824 | 6 | 6836824 | 8.00E-05 | 0.35786 | 299 | 0.129637025 | 0.047148495 |
| SNP_A.K_963595_5_28294113 | 5 | 28294113 | 8.01E-05 | 0.01338 | 299 | 0.129632809 | 0.047148495 |
| SNP_A.K_1001628_6_6499457 | 6 | 6499457 | 8.01E-05 | 0.32609 | 299 | 0.129631185 | 0.047148495 |
| SNP_A.K_1382854_8_9119512 | 8 | 9119512 | 8.01E-05 | 0.27258 | 299 | 0.129630514 | 0.047148495 |
| SNP_A.K_1382970_8_9125489 | 8 | 9125489 | 8.01E-05 | 0.14214 | 299 | 0.129630244 | 0.047148495 |
| SNP_A.K_1382695_8_9113466 | 8 | 9113466 | 8.02E-05 | 0.27926 | 299 | 0.129619927 | 0.047213297 |
| SNP_A.K_308452_2_7517022 | 2 | 7517022 | 8.04E-05 | 0.18896 | 299 | 0.129603344 | 0.047325769 |
| SNP_A.K_307470_2_7409427 | 2 | 7409427 | 8.05E-05 | 0.2709 | 299 | 0.129600486 | 0.047334111 |
| SNP_A.K_306917_2_7366603 | 2 | 7366603 | 8.05E-05 | 0.27258 | 299 | 0.12959809 | 0.047338942 |
| SNP_A.K_8623_1_1623800 | 1 | 1623800 | 8.05E-05 | 0.08194 | 299 | 0.129596162 | 0.047340223 |
| SNP_A.K_1293946_7_23121204 | 7 | 23121204 | 8.06E-05 | 0.03512 | 299 | 0.129592627 | 0.047353729 |
| SNP_A.K_978776_6_1780088 | 6 | 1780088 | 8.07E-05 | 0.30268 | 299 | 0.129580886 | 0.04742916 |
| SNP_A.K_708666_4_8072070 | 4 | 8072070 | 8.08E-05 | 0.09365 | 299 | 0.129579196 | 0.04742916 |
| SNP_A.K_1925579_11_24746935 | 11 | 24746935 | 8.08E-05 | 0.03512 | 299 | 0.129576243 | 0.04743096 |

| | | | | | | | |
|---|---|---|---|---|---|---|---|
| SNP_A.K_307151_2_7382007 | 2 | 7382007 | 8.08E-05 | 0.39967 | 299 | 0.129573532 | 0.04743096 |
| SNP_A.K_1000480_6_6388682 | 6 | 6388682 | 8.08E-05 | 0.25418 | 299 | 0.129572854 | 0.04743096 |
| SNP_A.K_312022_2_7899217 | 2 | 7899217 | 8.09E-05 | 0.26589 | 299 | 0.129569568 | 0.04743096 |
| SNP_A.K_1383163_8_9132629 | 8 | 9132629 | 8.09E-05 | 0.28261 | 299 | 0.129567423 | 0.04743096 |
| SNP_A.K_992573_6_5659517 | 6 | 5659517 | 8.09E-05 | 0.49331 | 299 | 0.129565347 | 0.04743096 |
| SNP_A.K_978006_6_1635119 | 6 | 1635119 | 8.10E-05 | 0.26756 | 299 | 0.129562682 | 0.04743096 |
| SNP_A.K_1382763_8_9116039 | 8 | 9116039 | 8.10E-05 | 0.27425 | 299 | 0.129560315 | 0.04743096 |
| SNP_A.K_710535_4_8426696 | 4 | 8426696 | 8.10E-05 | 0.08027 | 299 | 0.129558593 | 0.04743096 |
| SNP_A.K_1433100_8_16617641 | 8 | 16617641 | 8.10E-05 | 0.0903 | 299 | 0.129558131 | 0.04743096 |
| SNP_A.K_218895_1_35379698 | 1 | 35379698 | 8.11E-05 | 0.15385 | 299 | 0.129552819 | 0.04743096 |
| SNP_A.K_482888_3_3103485 | 3 | 3103485 | 8.11E-05 | 0.00502 | 299 | 0.129552122 | 0.04743096 |
| SNP_A.K_1389807_8_10127899 | 8 | 10127899 | 8.11E-05 | 0.04013 | 299 | 0.129551714 | 0.04743096 |
| SNP_A.K_1001203_6_6454957 | 6 | 6454957 | 8.11E-05 | 0.32107 | 299 | 0.129548689 | 0.04743096 |
| SNP_A.K_978558_6_1724409 | 6 | 1724409 | 8.12E-05 | 0.26087 | 299 | 0.129547463 | 0.04743096 |
| SNP_A.K_868704_5_4771673 | 5 | 4771673 | 8.12E-05 | 0.16722 | 299 | 0.129546712 | 0.04743096 |
| SNP_A.K_1830868_11_10585599 | 11 | 10585599 | 8.12E-05 | 0.21405 | 299 | 0.129546091 | 0.04743096 |
| SNP_A.K_307198_2_7384350 | 2 | 7384350 | 8.12E-05 | 0.40134 | 299 | 0.129544482 | 0.04743096 |
| SNP_A.K_1018131_6_8840415 | 6 | 8840415 | 8.12E-05 | 0.07692 | 299 | 0.1295443 | 0.04743096 |
| SNP_A.K_1389914_8_10135405 | 8 | 10135405 | 8.12E-05 | 0.06187 | 299 | 0.129543908 | 0.04743096 |
| SNP_A.K_1349626_8_3725597 | 8 | 3725597 | 8.12E-05 | 0.02174 | 299 | 0.129542145 | 0.047431084 |
| SNP_A.K_1001244_6_6457890 | 6 | 6457890 | 8.13E-05 | 0.29097 | 299 | 0.129537738 | 0.047451343 |
| SNP_A.K_1004408_6_6880002 | 6 | 6880002 | 8.14E-05 | 0.47492 | 299 | 0.129532536 | 0.047477681 |
| SNP_A.K_1383102_8_9130726 | 8 | 9130726 | 8.15E-05 | 0.27425 | 299 | 0.129522277 | 0.047542622 |
| SNP_A.K_421707_2_24819079 | 2 | 24819079 | 8.17E-05 | 0.02843 | 299 | 0.129505881 | 0.047654586 |
| SNP_A.K_710093_4_8315772 | 4 | 8315772 | 8.18E-05 | 0.11037 | 299 | 0.129500785 | 0.047680236 |
| SNP_A.K_863270_5_3670844 | 5 | 3670844 | 8.18E-05 | 0.06522 | 299 | 0.129498088 | 0.04768753 |
| SNP_A.K_1804588_11_7090062 | 11 | 7090062 | 8.21E-05 | 0.02508 | 299 | 0.12947793 | 0.047828749 |
| SNP_A.K_1001519_6_6486695 | 6 | 6486695 | 8.23E-05 | 0.301 | 299 | 0.129463023 | 0.047929957 |
| SNP_A.K_993250_6_5724272 | 6 | 5724272 | 8.27E-05 | 0.2291 | 299 | 0.129428338 | 0.048180573 |
| SNP_A.K_1391374_8_10301324 | 8 | 10301324 | 8.28E-05 | 0.05017 | 299 | 0.129423507 | 0.048180573 |
| SNP_A.K_978417_6_1689309 | 6 | 1689309 | 8.28E-05 | 0.26756 | 299 | 0.129421261 | 0.048180573 |
| SNP_A.K_1600889_9_18860007 | 9 | 18860007 | 8.28E-05 | 0.13712 | 299 | 0.129420052 | 0.048180573 |
| SNP_A.K_1261351_7_17339078 | 7 | 17339078 | 8.28E-05 | 0.01839 | 299 | 0.129419592 | 0.048180573 |
| SNP_A.K_1000761_6_6415497 | 6 | 6415497 | 8.29E-05 | 0.34783 | 299 | 0.12941874 | 0.048180573 |
| SNP_A.K_308603_2_7528963 | 2 | 7528963 | 8.29E-05 | 0.25084 | 299 | 0.129418359 | 0.048180573 |
| SNP_A.K_1726517_10_16594449 | 10 | 16594449 | 8.29E-05 | 0.06187 | 299 | 0.129414466 | 0.048194557 |
| SNP_A.K_710394_4_8339949 | 4 | 8339949 | 8.29E-05 | 0.07692 | 299 | 0.129413075 | 0.048194557 |
| SNP_A.K_308604_2_7528965 | 2 | 7528965 | 8.30E-05 | 0.22074 | 299 | 0.12940637 | 0.048233011 |
| SNP_A.K_1001540_6_6488536 | 6 | 6488536 | 8.31E-05 | 0.32274 | 299 | 0.129403802 | 0.048239447 |
| SNP_A.K_709867_4_8299910 | 4 | 8299910 | 8.31E-05 | 0.08194 | 299 | 0.129398207 | 0.048269344 |
| SNP_A.K_1001732_6_6511710 | 6 | 6511710 | 8.33E-05 | 0.31271 | 299 | 0.129384052 | 0.048365711 |
| SNP_A.K_995672_6_5946897 | 6 | 5946897 | 8.34E-05 | 0.47826 | 299 | 0.129382051 | 0.048367766 |
| SNP_A.K_992005_6_5607999 | 6 | 5607999 | 8.34E-05 | 0.27759 | 299 | 0.129377809 | 0.048387244 |
| SNP_A.K_20616_1_3100204 | 1 | 3100204 | 8.35E-05 | 0.00502 | 299 | 0.129372491 | 0.048415098 |
| SNP_A.K_1608494_9_20763771 | 9 | 20763771 | 8.35E-05 | 0.30769 | 299 | 0.129370487 | 0.048417188 |
| SNP_A.K_977989_6_1631020 | 6 | 1631020 | 8.35E-05 | 0.19565 | 299 | 0.129367668 | 0.048425631 |
| SNP_A.K_1001715_6_6508592 | 6 | 6508592 | 8.36E-05 | 0.31104 | 299 | 0.129361599 | 0.048459358 |
| SNP_A.K_1831031_11_10596750 | 11 | 10596750 | 8.38E-05 | 0.41973 | 299 | 0.129350795 | 0.048530017 |
| SNP_A.K_308024_2_7462905 | 2 | 7462905 | 8.41E-05 | 0.45652 | 299 | 0.129323452 | 0.048730058 |
| SNP_A.K_1001708_6_6507860 | 6 | 6507860 | 8.42E-05 | 0.30268 | 299 | 0.129317815 | 0.048752289 |

| | | | | | | | |
|---|---|---|---|---|---|---|---|
| SNP_A.K_1001747_6_6514498 | 6 | 6514498 | 8.42E-05 | 0.31104 | 299 | 0.129317149 | 0.048752289 |
| SNP_A.K_218819_1_35367739 | 1 | 35367739 | 8.43E-05 | 0.1689 | 299 | 0.12931257 | 0.048774596 |
| SNP_A.K_978342_6_1679898 | 6 | 1679898 | 8.43E-05 | 0.25585 | 299 | 0.129309895 | 0.048781998 |
| SNP_A.K_326689_2_10039219 | 2 | 10039219 | 8.45E-05 | 0.09197 | 299 | 0.129296294 | 0.048875105 |
| SNP_A.K_1004699_6_6919601 | 6 | 6919601 | 8.46E-05 | 0.28094 | 299 | 0.129291161 | 0.048901838 |
| SNP_A.K_996813_6_6027424 | 6 | 6027424 | 8.48E-05 | 0.46488 | 299 | 0.129278169 | 0.048990394 |
| SNP_A.K_992548_6_5655951 | 6 | 5655951 | 8.48E-05 | 0.19398 | 299 | 0.129274666 | 0.049004359 |
| SNP_A.K_1382754_8_9115710 | 8 | 9115710 | 8.49E-05 | 0.27258 | 299 | 0.129268153 | 0.04904204 |
| SNP_A.K_1364912_8_6055399 | 8 | 6055399 | 8.49E-05 | 0.05518 | 299 | 0.12926526 | 0.049045288 |
| SNP_A.K_998860_6_6229196 | 6 | 6229196 | 8.49E-05 | 0.26756 | 299 | 0.129264287 | 0.049045288 |
| SNP_A.K_1608336_9_20721637 | 9 | 20721637 | 8.51E-05 | 0.28261 | 299 | 0.129249772 | 0.049129272 |
| SNP_A.K_978344_6_1679907 | 6 | 1679907 | 8.52E-05 | 0.26087 | 299 | 0.129248948 | 0.049129272 |
| SNP_A.K_478916_3_2117334 | 3 | 2117334 | 8.52E-05 | 0.21237 | 299 | 0.129248459 | 0.049129272 |
| SNP_A.K_978928_6_1807960 | 6 | 1807960 | 8.52E-05 | 0.47157 | 299 | 0.129243636 | 0.049153727 |
| SNP_A.K_1383104_8_9130735 | 8 | 9130735 | 8.53E-05 | 0.27759 | 299 | 0.129238855 | 0.049156823 |
| SNP_A.K_1323468_7_29126048 | 7 | 29126048 | 8.53E-05 | 0.03846 | 299 | 0.129238447 | 0.049156823 |
| SNP_A.K_1004980_6_6959612 | 6 | 6959612 | 8.53E-05 | 0.33445 | 299 | 0.129238072 | 0.049156823 |
| SNP_A.K_1001941_6_6531338 | 6 | 6531338 | 8.53E-05 | 0.3194 | 299 | 0.129235091 | 0.049166755 |
| SNP_A.K_1382684_8_9113252 | 8 | 9113252 | 8.54E-05 | 0.28261 | 299 | 0.129231879 | 0.049177968 |
| SNP_A.K_709663_4_8279629 | 4 | 8279629 | 8.54E-05 | 0.08863 | 299 | 0.129230227 | 0.049177968 |
| SNP_A.K_1001571_6_6492428 | 6 | 6492428 | 8.55E-05 | 0.2893 | 299 | 0.129226907 | 0.049190602 |
| SNP_A.K_992594_6_5660145 | 6 | 5660145 | 8.56E-05 | 0.49164 | 299 | 0.129219433 | 0.049236088 |
| SNP_A.K_198704_1_31778131 | 1 | 31778131 | 8.57E-05 | 0.35619 | 299 | 0.129208413 | 0.049309704 |
| SNP_A.K_308668_2_7534932 | 2 | 7534932 | 8.57E-05 | 0.18896 | 299 | 0.129206247 | 0.049310257 |
| SNP_A.K_992021_6_5613140 | 6 | 5613140 | 8.58E-05 | 0.23913 | 299 | 0.129204903 | 0.049310257 |
| SNP_A.K_307901_2_7453477 | 2 | 7453477 | 8.58E-05 | 0.45987 | 299 | 0.129202993 | 0.049311774 |
| SNP_A.K_1001655_6_6501015 | 6 | 6501015 | 8.58E-05 | 0.44816 | 299 | 0.129199511 | 0.049318572 |
| SNP_A.K_993116_6_5708781 | 6 | 5708781 | 8.58E-05 | 0.11204 | 299 | 0.129198698 | 0.049318572 |
| SNP_A.K_580686_3_21128225 | 3 | 21128225 | 8.59E-05 | 0.48328 | 299 | 0.129191073 | 0.049342827 |
| SNP_A.K_513806_3_8342538 | 3 | 8342538 | 8.60E-05 | 0.02508 | 299 | 0.129188973 | 0.049342827 |
| SNP_A.K_218773_1_35348651 | 1 | 35348651 | 8.60E-05 | 0.15552 | 299 | 0.129186192 | 0.049342827 |
| SNP_A.K_278112_2_3027686 | 2 | 3027686 | 8.60E-05 | 0.0903 | 299 | 0.129185766 | 0.049342827 |
| SNP_A.K_995240_6_5921101 | 6 | 5921101 | 8.60E-05 | 0.2408 | 299 | 0.12918544 | 0.049342827 |
| SNP_A.K_758817_4_19402229 | 4 | 19402229 | 8.60E-05 | 0.00502 | 299 | 0.129185339 | 0.049342827 |
| SNP_A.K_1816215_11_9006025 | 11 | 9006025 | 8.61E-05 | 0.39967 | 299 | 0.12917965 | 0.049364271 |
| SNP_A.K_1924519_11_24649992 | 11 | 24649992 | 8.61E-05 | 0.02341 | 299 | 0.129179205 | 0.049364271 |
| SNP_A.K_1019147_6_9008417 | 6 | 9008417 | 8.62E-05 | 0.10201 | 299 | 0.129172361 | 0.049404983 |
| SNP_A.K_218471_1_35269885 | 1 | 35269885 | 8.63E-05 | 0.15886 | 299 | 0.129168786 | 0.049419758 |
| SNP_A.K_1001541_6_6488561 | 6 | 6488561 | 8.64E-05 | 0.31773 | 299 | 0.129158056 | 0.049491418 |
| SNP_A.K_1004998_6_6961047 | 6 | 6961047 | 8.65E-05 | 0.3194 | 299 | 0.129150103 | 0.049541065 |
| SNP_A.K_977972_6_1627112 | 6 | 1627112 | 8.66E-05 | 0.2709 | 299 | 0.129143514 | 0.049579908 |
| SNP_A.K_1608679_9_20815871 | 9 | 20815871 | 8.69E-05 | 0.30435 | 299 | 0.129125416 | 0.049703386 |
| SNP_A.K_1001360_6_6466803 | 6 | 6466803 | 8.69E-05 | 0.2893 | 299 | 0.129124612 | 0.049703386 |
| SNP_A.K_710543_4_8426850 | 4 | 8426850 | 8.70E-05 | 0.0786 | 299 | 0.129117971 | 0.049731111 |
| SNP_A.K_1726512_10_16593653 | 10 | 16593653 | 8.70E-05 | 0.0602 | 299 | 0.129117721 | 0.049731111 |
| SNP_A.K_1162924_6_30904814 | 6 | 30904814 | 8.70E-05 | 0.03512 | 299 | 0.129113633 | 0.049739833 |
| SNP_A.K_1001742_6_6512543 | 6 | 6512543 | 8.70E-05 | 0.31271 | 299 | 0.129113212 | 0.049739833 |
| SNP_A.K_995532_6_5940401 | 6 | 5940401 | 8.72E-05 | 0.46656 | 299 | 0.129099563 | 0.049835362 |
| SNP_A.K_1803420_11_6912569 | 11 | 6912569 | 8.73E-05 | 0.01171 | 299 | 0.129093116 | 0.049873333 |
| SNP_A.K_995726_6_5948984 | 6 | 5948984 | 8.74E-05 | 0.49498 | 299 | 0.129090217 | 0.049875624 |

| | | | | | | | |
|---|---|---|---|---|---|---|---|
| SNP_A.K_1019295_6_9032330 | 6 | 9032330 | 8.74E-05 | 0.03512 | 299 | 0.129086003 | 0.049875624 |
| SNP_A.K_1019353_6_9043307 | 6 | 9043307 | 8.74E-05 | 0.03512 | 299 | 0.129086003 | 0.049875624 |
| SNP_A.K_1021064_6_9317440 | 6 | 9317440 | 8.74E-05 | 0.03512 | 299 | 0.129086003 | 0.049875624 |
| SNP_A.K_1383168_8_9133136 | 8 | 9133136 | 8.75E-05 | 0.10702 | 299 | 0.129077102 | 0.049921295 |
| SNP_A.K_218582_1_35319214 | 1 | 35319214 | 8.75E-05 | 0.29933 | 299 | 0.129076897 | 0.049921295 |
| SNP_A.K_1433105_8_16617724 | 8 | 16617724 | 8.77E-05 | 0.10535 | 299 | 0.129068738 | 0.049973107 |

Grain_length

| SNP | Chrc | Position | P.value | maf | nobs | Rsquare.of.Model.w | FDR_Adjusted_P-values |
|---|---|---|---|---|---|---|---|
| SNP_A.K_553163_3_16733441 | 3 | 16733441 | 7.16E-10 | 0.24688 | 320 | 0.322481071 | 0.001491214 |
| SNP_A.K_553157_3_16692213 | 3 | 16692213 | 3.44E-08 | 0.04219 | 320 | 0.30180039 | 0.035789542 |
| SNP_A.K_553158_3_16696473 | 3 | 16696473 | 5.27E-08 | 0.04063 | 320 | 0.299555021 | 0.036583814 |

Grain_width

| SNP | Chrc | Position | P.value | maf | nobs | Rsquare.of.Model.w | FDR_Adjusted_P-values |
|---|---|---|---|---|---|---|---|
| SNP_A.K_871244_5_5371949 | 5 | 5371949 | 3.23E-10 | 0.46719 | 320 | 0.342419716 | 0.000288826 |
| SNP_A.K_871245_5_5371952 | 5 | 5371952 | 4.06E-10 | 0.47188 | 320 | 0.34120786 | 0.000288826 |
| SNP_A.K_871237_5_5371529 | 5 | 5371529 | 5.14E-10 | 0.46719 | 320 | 0.339950934 | 0.000288826 |
| SNP_A.K_871243_5_5371916 | 5 | 5371916 | 5.86E-10 | 0.46406 | 320 | 0.339257308 | 0.000288826 |
| SNP_A.K_871272_5_5373841 | 5 | 5373841 | 6.94E-10 | 0.4625 | 320 | 0.338365666 | 0.000288826 |
| SNP_A.K_871258_5_5373422 | 5 | 5373422 | 2.21E-09 | 0.49844 | 320 | 0.332248326 | 0.00063476 |
| SNP_A.K_871239_5_5371686 | 5 | 5371686 | 2.28E-09 | 0.47813 | 320 | 0.332096651 | 0.00063476 |
| SNP_A.K_871241_5_5371716 | 5 | 5371716 | 2.44E-09 | 0.47813 | 320 | 0.331737021 | 0.00063476 |
| SNP_A.K_871238_5_5371609 | 5 | 5371609 | 3.31E-09 | 0.47188 | 320 | 0.330138572 | 0.000765454 |
| SNP_A.K_871242_5_5371772 | 5 | 5371772 | 3.72E-09 | 0.48125 | 320 | 0.329532069 | 0.000773574 |
| SNP_A.K_871265_5_5373560 | 5 | 5373560 | 5.93E-09 | 0.48281 | 320 | 0.327091098 | 0.000985037 |
| SNP_A.K_871147_5_5361894 | 5 | 5361894 | 6.08E-09 | 0.37969 | 320 | 0.326962577 | 0.000985037 |
| SNP_A.K_871141_5_5361276 | 5 | 5361276 | 6.15E-09 | 0.38594 | 320 | 0.32690106 | 0.000985037 |
| SNP_A.K_871277_5_5374961 | 5 | 5374961 | 9.81E-09 | 0.44219 | 320 | 0.324471703 | 0.001459283 |
| SNP_A.K_871294_5_5376138 | 5 | 5376138 | 1.11E-08 | 0.43594 | 320 | 0.323810771 | 0.001538233 |
| SNP_A.K_871114_5_5359520 | 5 | 5359520 | 1.18E-08 | 0.36406 | 320 | 0.323505686 | 0.001538233 |
| SNP_A.K_871116_5_5359681 | 5 | 5359681 | 1.37E-08 | 0.36875 | 320 | 0.322732282 | 0.001596788 |
| SNP_A.K_871115_5_5359598 | 5 | 5359598 | 1.38E-08 | 0.36406 | 320 | 0.322701957 | 0.001596788 |
| SNP_A.K_871283_5_5375201 | 5 | 5375201 | 1.64E-08 | 0.44531 | 320 | 0.321803555 | 0.001799603 |
| SNP_A.K_871260_5_5373491 | 5 | 5373491 | 1.75E-08 | 0.475 | 320 | 0.321490408 | 0.001816393 |
| SNP_A.K_871103_5_5358972 | 5 | 5358972 | 2.33E-08 | 0.37188 | 320 | 0.319996347 | 0.002310564 |
| SNP_A.K_871289_5_5375786 | 5 | 5375786 | 2.79E-08 | 0.43906 | 320 | 0.319068785 | 0.002640564 |
| SNP_A.K_871290_5_5375793 | 5 | 5375793 | 3.69E-08 | 0.44375 | 320 | 0.317634758 | 0.003337973 |
| SNP_A.K_871102_5_5358870 | 5 | 5358870 | 4.32E-08 | 0.38125 | 320 | 0.316825473 | 0.003704623 |
| SNP_A.K_871280_5_5375138 | 5 | 5375138 | 4.69E-08 | 0.44844 | 320 | 0.316400072 | 0.003704623 |
| SNP_A.K_871108_5_5359246 | 5 | 5359246 | 4.70E-08 | 0.36406 | 320 | 0.316390994 | 0.003704623 |
| SNP_A.K_871288_5_5375764 | 5 | 5375764 | 4.81E-08 | 0.44063 | 320 | 0.316277037 | 0.003704623 |
| SNP_A.K_871311_5_5377198 | 5 | 5377198 | 5.30E-08 | 0.40781 | 320 | 0.315771451 | 0.003942734 |
| SNP_A.K_871234_5_5371042 | 5 | 5371042 | 6.01E-08 | 0.37344 | 320 | 0.315131249 | 0.004185944 |
| SNP_A.K_871255_5_5373133 | 5 | 5373133 | 6.03E-08 | 0.49375 | 320 | 0.315111761 | 0.004185944 |
| SNP_A.K_871282_5_5375167 | 5 | 5375167 | 6.97E-08 | 0.44844 | 320 | 0.314372658 | 0.004570922 |
| SNP_A.K_871281_5_5375144 | 5 | 5375144 | 7.03E-08 | 0.44844 | 320 | 0.314331773 | 0.004570922 |
| SNP_A.K_871322_5_5378228 | 5 | 5378228 | 7.26E-08 | 0.43438 | 320 | 0.314161628 | 0.004582475 |
| SNP_A.K_871270_5_5373750 | 5 | 5373750 | 8.94E-08 | 0.5 | 320 | 0.313102073 | 0.005473591 |
| SNP_A.K_871292_5_5375922 | 5 | 5375922 | 1.02E-07 | 0.45 | 320 | 0.312448851 | 0.006044074 |
| SNP_A.K_871297_5_5376161 | 5 | 5376161 | 1.09E-07 | 0.43125 | 320 | 0.31208433 | 0.006312096 |

| | | | | | | | |
|---|---|---|---|---|---|---|---|
| SNP_A.K_871171_5_5363611 | 5 | 5363611 | 1.37E-07 | 0.39531 | 320 | 0.31092526 | 0.007712874 |
| SNP_A.K_871098_5_5358629 | 5 | 5358629 | 1.69E-07 | 0.36719 | 320 | 0.309869016 | 0.00888159 |
| SNP_A.K_871216_5_5369802 | 5 | 5369802 | 1.72E-07 | 0.38281 | 320 | 0.309764431 | 0.00888159 |
| SNP_A.K_871268_5_5373724 | 5 | 5373724 | 1.72E-07 | 0.48438 | 320 | 0.309760641 | 0.00888159 |
| SNP_A.K_871210_5_5369555 | 5 | 5369555 | 1.77E-07 | 0.39688 | 320 | 0.309627763 | 0.00888159 |
| SNP_A.K_871257_5_5373379 | 5 | 5373379 | 1.79E-07 | 0.49688 | 320 | 0.309565388 | 0.00888159 |
| SNP_A.K_871274_5_5374196 | 5 | 5374196 | 2.02E-07 | 0.46875 | 320 | 0.308955855 | 0.009783517 |
| SNP_A.K_871227_5_5370747 | 5 | 5370747 | 2.08E-07 | 0.36406 | 320 | 0.308818888 | 0.009823202 |
| SNP_A.K_871195_5_5368150 | 5 | 5368150 | 2.14E-07 | 0.38594 | 320 | 0.308663786 | 0.009903608 |
| SNP_A.K_871286_5_5375574 | 5 | 5375574 | 2.66E-07 | 0.46406 | 320 | 0.307560011 | 0.012050304 |
| SNP_A.K_871181_5_5364561 | 5 | 5364561 | 2.75E-07 | 0.36563 | 320 | 0.307392322 | 0.012191805 |
| SNP_A.K_871313_5_5377607 | 5 | 5377607 | 2.91E-07 | 0.40469 | 320 | 0.307118884 | 0.012498799 |
| SNP_A.K_871326_5_5378446 | 5 | 5378446 | 2.94E-07 | 0.42344 | 320 | 0.307056173 | 0.012498799 |
| SNP_A.K_871273_5_5374168 | 5 | 5374168 | 3.03E-07 | 0.475 | 320 | 0.306912815 | 0.012601544 |
| SNP_A.K_871205_5_5369111 | 5 | 5369111 | 3.11E-07 | 0.39375 | 320 | 0.30676938 | 0.012710497 |
| SNP_A.K_871276_5_5374221 | 5 | 5374221 | 3.20E-07 | 0.46563 | 320 | 0.306637608 | 0.012795808 |
| SNP_A.K_871327_5_5378981 | 5 | 5378981 | 3.44E-07 | 0.41563 | 320 | 0.306261035 | 0.013527308 |
| SNP_A.K_1161223_6_30339552 | 6 | 30339552 | 3.68E-07 | 0.10938 | 320 | 0.305933031 | 0.013921663 |
| SNP_A.K_871232_5_5370906 | 5 | 5370906 | 3.68E-07 | 0.36563 | 320 | 0.305929381 | 0.013921663 |
| SNP_A.K_871318_5_5377857 | 5 | 5377857 | 4.36E-07 | 0.43438 | 320 | 0.305078084 | 0.016190688 |
| SNP_A.K_871247_5_5372050 | 5 | 5372050 | 4.57E-07 | 0.475 | 320 | 0.304835454 | 0.016692361 |
| SNP_A.K_871291_5_5375849 | 5 | 5375849 | 6.12E-07 | 0.37969 | 320 | 0.303367899 | 0.021968686 |
| SNP_A.K_871157_5_5362848 | 5 | 5362848 | 6.35E-07 | 0.3875 | 320 | 0.303185379 | 0.022396371 |
| SNP_A.K_871197_5_5368219 | 5 | 5368219 | 7.10E-07 | 0.3875 | 320 | 0.302621905 | 0.024642266 |
| SNP_A.K_871198_5_5368312 | 5 | 5368312 | 9.09E-07 | 0.38281 | 320 | 0.301385331 | 0.031029237 |
| SNP_A.K_871253_5_5372955 | 5 | 5372955 | 1.03E-06 | 0.48906 | 320 | 0.300780105 | 0.03445852 |
| SNP_A.K_871284_5_5375267 | 5 | 5375267 | 1.14E-06 | 0.46094 | 320 | 0.300275797 | 0.037515444 |
| SNP_A.K_871309_5_5377112 | 5 | 5377112 | 1.33E-06 | 0.4125 | 320 | 0.299485367 | 0.043271191 |

Panicle_axis

| SNP | Chr | Position | P.value | maf | nobs | Rsquare.of.Model.w | FDR_Adjusted_P-values |
|---|---|---|---|---|---|---|---|
| SNP_A.K_679358_4_1075655 | 4 | 1075655 | 3.17E-10 | 0.01302 | 192 | 0.240455921 | 0.000375146 |
| SNP_A.K_1151705_6_28676456 | 6 | 28676456 | 6.49E-10 | 0.00781 | 192 | 0.231306619 | 0.000375146 |
| SNP_A.K_1712473_10_14829875 | 10 | 14829875 | 6.66E-10 | 0.00781 | 192 | 0.23097996 | 0.000375146 |
| SNP_A.K_1681573_10_10214034 | 10 | 10214034 | 7.21E-10 | 0.00781 | 192 | 0.229984796 | 0.000375146 |
| SNP_A.K_288341_2_5011285 | 2 | 5011285 | 6.39E-09 | 0.01042 | 192 | 0.202709095 | 0.002393125 |
| SNP_A.K_1545254_9_7356966 | 9 | 7356966 | 7.71E-09 | 0.04427 | 192 | 0.200384457 | 0.002393125 |
| SNP_A.K_1543746_9_7148402 | 9 | 7148402 | 8.05E-09 | 0.01823 | 192 | 0.19986044 | 0.002393125 |
| SNP_A.K_1803408_11_6911996 | 11 | 6911996 | 3.24E-08 | 0.02604 | 192 | 0.182876074 | 0.008419603 |
| SNP_A.K_288314_2_5007783 | 2 | 5007783 | 4.60E-08 | 0.01302 | 192 | 0.178630124 | 0.009865679 |
| SNP_A.K_288469_2_5032381 | 2 | 5032381 | 8.93E-08 | 0.01302 | 192 | 0.170668312 | 0.009865679 |
| SNP_A.K_2076621_12_26474657 | 12 | 26474657 | 9.59E-08 | 0.03646 | 192 | 0.169820499 | 0.009865679 |
| SNP_A.K_1535441_9_6111975 | 9 | 6111975 | 1.01E-07 | 0.01042 | 192 | 0.169228264 | 0.009865679 |
| SNP_A.K_651649_3_32604953 | 3 | 32604953 | 1.03E-07 | 0.01823 | 192 | 0.168963853 | 0.009865679 |
| SNP_A.K_1192398_7_7094693 | 7 | 7094693 | 1.05E-07 | 0.01302 | 192 | 0.168746184 | 0.009865679 |
| SNP_A.K_288109_2_4988475 | 2 | 4988475 | 1.05E-07 | 0.01302 | 192 | 0.1687002 | 0.009865679 |
| SNP_A.K_288538_2_5037464 | 2 | 5037464 | 1.30E-07 | 0.01302 | 192 | 0.166199152 | 0.009865679 |
| SNP_A.K_1804778_11_7101390 | 11 | 7101390 | 1.36E-07 | 0.01042 | 192 | 0.16567253 | 0.009865679 |
| SNP_A.K_228078_1_37144984 | 1 | 37144984 | 1.66E-07 | 0.01042 | 192 | 0.163318027 | 0.009865679 |
| SNP_A.K_902536_5_16534350 | 5 | 16534350 | 1.79E-07 | 0.01042 | 192 | 0.162382738 | 0.009865679 |
| SNP_A.K_1557619_9_10009442 | 9 | 10009442 | 1.80E-07 | 0.00521 | 192 | 0.162300093 | 0.009865679 |

| | | | | | | | |
|---|---|---|---|---|---|---|---|
| SNP_A.K_1805532_11_7199542 | 11 | 7199542 | 1.80E-07 | 0.00521 | 192 | 0.162300093 | 0.009865679 |
| SNP_A.K_652308_3_32791155 | 3 | 32791155 | 1.80E-07 | 0.00521 | 192 | 0.162300093 | 0.009865679 |
| SNP_A.K_902551_5_16545655 | 5 | 16545655 | 1.80E-07 | 0.00521 | 192 | 0.162300093 | 0.009865679 |
| SNP_A.K_20343_1_3082541 | 1 | 3082541 | 1.90E-07 | 0.00781 | 192 | 0.161705289 | 0.009865679 |
| SNP_A.K_829651_4_33918753 | 4 | 33918753 | 1.98E-07 | 0.00521 | 192 | 0.16117166 | 0.009865679 |
| SNP_A.K_870468_5_5303956 | 5 | 5303956 | 1.98E-07 | 0.00521 | 192 | 0.16117166 | 0.009865679 |
| SNP_A.K_953607_5_25097135 | 5 | 25097135 | 1.98E-07 | 0.00521 | 192 | 0.16117166 | 0.009865679 |
| SNP_A.K_1606956_9_20241256 | 9 | 20241256 | 1.98E-07 | 0.00521 | 192 | 0.16117166 | 0.009865679 |
| SNP_A.K_384409_2_19153339 | 2 | 19153339 | 1.98E-07 | 0.00521 | 192 | 0.16117166 | 0.009865679 |
| SNP_A.K_526677_3_10964889 | 3 | 10964889 | 1.98E-07 | 0.00521 | 192 | 0.16117166 | 0.009865679 |
| SNP_A.K_1288155_7_22249791 | 7 | 22249791 | 1.98E-07 | 0.00521 | 192 | 0.16117166 | 0.009865679 |
| SNP_A.K_648754_3_31937585 | 3 | 31937585 | 2.07E-07 | 0.01042 | 192 | 0.160645541 | 0.009865679 |
| SNP_A.K_526222_3_10849783 | 3 | 10849783 | 2.30E-07 | 0.00521 | 192 | 0.15942989 | 0.009865679 |
| SNP_A.K_863296_5_3678971 | 5 | 3678971 | 2.30E-07 | 0.00521 | 192 | 0.15942989 | 0.009865679 |
| SNP_A.K_806606_4_29505446 | 4 | 29505446 | 2.30E-07 | 0.00521 | 192 | 0.15942989 | 0.009865679 |
| SNP_A.K_288218_2_4998843 | 2 | 4998843 | 2.32E-07 | 0.00521 | 192 | 0.159309678 | 0.009865679 |
| SNP_A.K_288228_2_4999616 | 2 | 4999616 | 2.32E-07 | 0.00521 | 192 | 0.159309678 | 0.009865679 |
| SNP_A.K_288229_2_4999648 | 2 | 4999648 | 2.32E-07 | 0.00521 | 192 | 0.159309678 | 0.009865679 |
| SNP_A.K_315911_2_8310508 | 2 | 8310508 | 2.32E-07 | 0.00521 | 192 | 0.159309678 | 0.009865679 |
| SNP_A.K_817407_4_31495768 | 4 | 31495768 | 2.32E-07 | 0.00521 | 192 | 0.159309678 | 0.009865679 |
| SNP_A.K_885802_5_8945161 | 5 | 8945161 | 2.32E-07 | 0.00521 | 192 | 0.159309678 | 0.009865679 |
| SNP_A.K_886511_5_13003572 | 5 | 13003572 | 2.32E-07 | 0.00521 | 192 | 0.159309678 | 0.009865679 |
| SNP_A.K_1371552_8_7852191 | 8 | 7852191 | 2.32E-07 | 0.00521 | 192 | 0.159309678 | 0.009865679 |
| SNP_A.K_363616_2_16455997 | 2 | 16455997 | 2.32E-07 | 0.00521 | 192 | 0.159309678 | 0.009865679 |
| SNP_A.K_528009_3_11220328 | 3 | 11220328 | 2.32E-07 | 0.00521 | 192 | 0.159309678 | 0.009865679 |
| SNP_A.K_1154455_6_29052368 | 6 | 29052368 | 2.32E-07 | 0.00521 | 192 | 0.159309678 | 0.009865679 |
| SNP_A.K_1371604_8_7858316 | 8 | 7858316 | 2.32E-07 | 0.00521 | 192 | 0.159309678 | 0.009865679 |
| SNP_A.K_1605756_9_19946683 | 9 | 19946683 | 2.32E-07 | 0.00521 | 192 | 0.159309678 | 0.009865679 |
| SNP_A.K_1752246_10_19901886 | 10 | 19901886 | 2.32E-07 | 0.00521 | 192 | 0.159309678 | 0.009865679 |
| SNP_A.K_889109_5_14206551 | 5 | 14206551 | 2.60E-07 | 0.00521 | 192 | 0.157999695 | 0.010622622 |
| SNP_A.K_366621_2_16762351 | 2 | 16762351 | 2.76E-07 | 0.00521 | 192 | 0.157291193 | 0.010622622 |
| SNP_A.K_1342595_8_2940419 | 8 | 2940419 | 2.76E-07 | 0.00521 | 192 | 0.157291193 | 0.010622622 |
| SNP_A.K_1647732_10_4648118 | 10 | 4648118 | 2.76E-07 | 0.00521 | 192 | 0.157291193 | 0.010622622 |
| SNP_A.K_1948607_12_2150360 | 12 | 2150360 | 2.76E-07 | 0.00521 | 192 | 0.157291193 | 0.010622622 |
| SNP_A.K_324792_2_9780295 | 2 | 9780295 | 2.99E-07 | 0.00521 | 192 | 0.156346969 | 0.010903451 |
| SNP_A.K_1371510_8_7850476 | 8 | 7850476 | 2.99E-07 | 0.00521 | 192 | 0.156346969 | 0.010903451 |
| SNP_A.K_20116_1_3038515 | 1 | 3038515 | 2.99E-07 | 0.00521 | 192 | 0.156346969 | 0.010903451 |
| SNP_A.K_1031878_6_11251082 | 6 | 11251082 | 3.19E-07 | 0.05729 | 192 | 0.155580251 | 0.01143683 |
| SNP_A.K_1739574_10_18161913 | 10 | 18161913 | 3.39E-07 | 0.01563 | 192 | 0.154867798 | 0.011945243 |
| SNP_A.K_288315_2_5007791 | 2 | 5007791 | 4.06E-07 | 0.01563 | 192 | 0.152731017 | 0.01387566 |
| SNP_A.K_1489858_8_27232205 | 8 | 27232205 | 4.07E-07 | 0.05469 | 192 | 0.152717642 | 0.01387566 |
| SNP_A.K_1794053_11_5377516 | 11 | 5377516 | 5.23E-07 | 0.04948 | 192 | 0.149765601 | 0.01756694 |
| SNP_A.K_1535465_9_6113565 | 9 | 6113565 | 6.51E-07 | 0.01563 | 192 | 0.147221977 | 0.02128225 |
| SNP_A.K_686965_4_2890383 | 4 | 2890383 | 6.54E-07 | 0.08333 | 192 | 0.147155718 | 0.02128225 |
| SNP_A.K_1685877_10_10754153 | 10 | 10754153 | 8.67E-07 | 0.01563 | 192 | 0.143887956 | 0.022991894 |
| SNP_A.K_864827_5_4097990 | 5 | 4097990 | 8.85E-07 | 0.02083 | 192 | 0.14363822 | 0.022991894 |
| SNP_A.K_1648152_10_4704356 | 10 | 4704356 | 9.04E-07 | 0.01563 | 192 | 0.143392915 | 0.022991894 |
| SNP_A.K_707999_4_7494052 | 4 | 7494052 | 1.00E-06 | 0.15625 | 192 | 0.142200898 | 0.022991894 |
| SNP_A.K_288103_2_4987974 | 2 | 4987974 | 1.15E-06 | 0.00781 | 192 | 0.140620569 | 0.022991894 |
| SNP_A.K_288114_2_4988672 | 2 | 4988672 | 1.15E-06 | 0.00781 | 192 | 0.140620569 | 0.022991894 |

| | | | | | | | |
|---|---|---|---|---|---|---|---|
| SNP_A.K_288116_2_4988805 | 2 | 4988805 | 1.15E-06 | 0.00781 | 192 | 0.140620569 | 0.022991894 |
| SNP_A.K_288122_2_4989029 | 2 | 4989029 | 1.15E-06 | 0.00781 | 192 | 0.140620569 | 0.022991894 |
| SNP_A.K_288123_2_4989036 | 2 | 4989036 | 1.15E-06 | 0.00781 | 192 | 0.140620569 | 0.022991894 |
| SNP_A.K_288132_2_4990502 | 2 | 4990502 | 1.15E-06 | 0.00781 | 192 | 0.140620569 | 0.022991894 |
| SNP_A.K_288136_2_4991126 | 2 | 4991126 | 1.15E-06 | 0.00781 | 192 | 0.140620569 | 0.022991894 |
| SNP_A.K_288144_2_4991792 | 2 | 4991792 | 1.15E-06 | 0.00781 | 192 | 0.140620569 | 0.022991894 |
| SNP_A.K_288145_2_4991816 | 2 | 4991816 | 1.15E-06 | 0.00781 | 192 | 0.140620569 | 0.022991894 |
| SNP_A.K_288150_2_4992218 | 2 | 4992218 | 1.15E-06 | 0.00781 | 192 | 0.140620569 | 0.022991894 |
| SNP_A.K_288157_2_4992816 | 2 | 4992816 | 1.15E-06 | 0.00781 | 192 | 0.140620569 | 0.022991894 |
| SNP_A.K_288158_2_4992821 | 2 | 4992821 | 1.15E-06 | 0.00781 | 192 | 0.140620569 | 0.022991894 |
| SNP_A.K_288163_2_4993339 | 2 | 4993339 | 1.15E-06 | 0.00781 | 192 | 0.140620569 | 0.022991894 |
| SNP_A.K_288164_2_4993351 | 2 | 4993351 | 1.15E-06 | 0.00781 | 192 | 0.140620569 | 0.022991894 |
| SNP_A.K_288178_2_4994046 | 2 | 4994046 | 1.15E-06 | 0.00781 | 192 | 0.140620569 | 0.022991894 |
| SNP_A.K_288189_2_4995195 | 2 | 4995195 | 1.15E-06 | 0.00781 | 192 | 0.140620569 | 0.022991894 |
| SNP_A.K_288197_2_4997655 | 2 | 4997655 | 1.15E-06 | 0.00781 | 192 | 0.140620569 | 0.022991894 |
| SNP_A.K_288279_2_5003424 | 2 | 5003424 | 1.15E-06 | 0.00781 | 192 | 0.140620569 | 0.022991894 |
| SNP_A.K_288290_2_5004458 | 2 | 5004458 | 1.15E-06 | 0.00781 | 192 | 0.140620569 | 0.022991894 |
| SNP_A.K_288493_2_5034560 | 2 | 5034560 | 1.15E-06 | 0.00781 | 192 | 0.140620569 | 0.022991894 |
| SNP_A.K_288563_2_5041476 | 2 | 5041476 | 1.15E-06 | 0.00781 | 192 | 0.140620569 | 0.022991894 |
| SNP_A.K_288565_2_5042437 | 2 | 5042437 | 1.15E-06 | 0.00781 | 192 | 0.140620569 | 0.022991894 |
| SNP_A.K_288092_2_4986793 | 2 | 4986793 | 1.15E-06 | 0.00781 | 192 | 0.140620569 | 0.022991894 |
| SNP_A.K_288113_2_4988667 | 2 | 4988667 | 1.15E-06 | 0.00781 | 192 | 0.140620569 | 0.022991894 |
| SNP_A.K_288120_2_4988990 | 2 | 4988990 | 1.15E-06 | 0.00781 | 192 | 0.140620569 | 0.022991894 |
| SNP_A.K_288125_2_4989212 | 2 | 4989212 | 1.15E-06 | 0.00781 | 192 | 0.140620569 | 0.022991894 |
| SNP_A.K_288146_2_4991828 | 2 | 4991828 | 1.15E-06 | 0.00781 | 192 | 0.140620569 | 0.022991894 |
| SNP_A.K_288156_2_4992518 | 2 | 4992518 | 1.15E-06 | 0.00781 | 192 | 0.140620569 | 0.022991894 |
| SNP_A.K_288184_2_4994991 | 2 | 4994991 | 1.15E-06 | 0.00781 | 192 | 0.140620569 | 0.022991894 |
| SNP_A.K_288194_2_4996097 | 2 | 4996097 | 1.15E-06 | 0.00781 | 192 | 0.140620569 | 0.022991894 |
| SNP_A.K_288199_2_4997717 | 2 | 4997717 | 1.15E-06 | 0.00781 | 192 | 0.140620569 | 0.022991894 |
| SNP_A.K_288285_2_5004065 | 2 | 5004065 | 1.15E-06 | 0.00781 | 192 | 0.140620569 | 0.022991894 |
| SNP_A.K_288287_2_5004370 | 2 | 5004370 | 1.15E-06 | 0.00781 | 192 | 0.140620569 | 0.022991894 |
| SNP_A.K_288399_2_5021434 | 2 | 5021434 | 1.15E-06 | 0.00781 | 192 | 0.140620569 | 0.022991894 |
| SNP_A.K_288492_2_5034545 | 2 | 5034545 | 1.15E-06 | 0.00781 | 192 | 0.140620569 | 0.022991894 |
| SNP_A.K_288500_2_5034670 | 2 | 5034670 | 1.15E-06 | 0.00781 | 192 | 0.140620569 | 0.022991894 |
| SNP_A.K_1319559_7_27994549 | 7 | 27994549 | 1.24E-06 | 0.1276 | 192 | 0.139745505 | 0.024563099 |
| SNP_A.K_1348256_8_3577908 | 8 | 3577908 | 1.53E-06 | 0.06771 | 192 | 0.137311694 | 0.030043156 |
| SNP_A.K_1895518_11_19527653 | 11 | 19527653 | 1.67E-06 | 0.01302 | 192 | 0.136327774 | 0.03241629 |
| SNP_A.K_1535491_9_6115356 | 9 | 6115356 | 1.73E-06 | 0.01823 | 192 | 0.135906264 | 0.033314097 |
| SNP_A.K_1535444_9_6112050 | 9 | 6112050 | 1.85E-06 | 0.01302 | 192 | 0.135124702 | 0.035329405 |
| SNP_A.K_635764_3_29600462 | 3 | 29600462 | 2.00E-06 | 0.01302 | 192 | 0.134252287 | 0.037769687 |
| SNP_A.K_1451681_8_19195204 | 8 | 19195204 | 2.09E-06 | 0.01302 | 192 | 0.133735238 | 0.038811359 |
| SNP_A.K_1681526_10_10209690 | 10 | 10209690 | 2.09E-06 | 0.02083 | 192 | 0.133733048 | 0.038811359 |
| SNP_A.K_1319500_7_27992765 | 7 | 27992765 | 2.18E-06 | 0.13802 | 192 | 0.133252989 | 0.040111403 |
| SNP_A.K_1319502_7_27992803 | 7 | 27992803 | 2.26E-06 | 0.13802 | 192 | 0.132810986 | 0.041321829 |
| SNP_A.K_833731_4_34877438 | 4 | 34877438 | 2.49E-06 | 0.16406 | 192 | 0.131717518 | 0.045064259 |
| SNP_A.K_732137_4_13283083 | 4 | 13283083 | 2.61E-06 | 0.01302 | 192 | 0.131198068 | 0.046749777 |
| SNP_A.K_1319571_7_27995114 | 7 | 27995114 | 2.67E-06 | 0.1224 | 192 | 0.130914771 | 0.047512162 |
| SNP_A.K_287596_2_4900668 | 2 | 4900668 | 2.70E-06 | 0.01823 | 192 | 0.130773894 | 0.047622921 |
| SNP_A.K_205192_1_32937176 | 1 | 32937176 | 2.72E-06 | 0.21875 | 192 | 0.130694263 | 0.047622921 |
| SNP_A.K_690585_4_4456947 | 4 | 4456947 | 2.79E-06 | 0.08333 | 192 | 0.130407954 | 0.048423449 |

| | | | | | | | |
|---|---|---|---|---|---|---|---|
| SNP_A.K_288630_2_5052237 | 2 | 5052237 | 2.90E-06 | 0.01823 | 192 | 0.129982885 | 0.049842802 |

Secondary branching

| SNP | Chr | Position | P.value | maf | nobs | Rsquare.of.Model.w | FDR_Adjusted_P-values |
|---|---|---|---|---|---|---|---|
| SNP_A.K_943938_5_23236471 | 5 | 23236471 | 2.84E-13 | 0.03646 | 192 | 0.336997249 | 5.90E-07 |
| SNP_A.K_288472_2_5032535 | 2 | 5032535 | 6.16E-13 | 0.01302 | 192 | 0.326360828 | 6.41E-07 |
| SNP_A.K_288170_2_4993555 | 2 | 4993555 | 1.47E-10 | 0.00521 | 192 | 0.253814777 | 6.54E-05 |
| SNP_A.K_288426_2_5026842 | 2 | 5026842 | 1.47E-10 | 0.0026 | 192 | 0.253814777 | 6.54E-05 |
| SNP_A.K_288394_2_5019623 | 2 | 5019623 | 1.57E-10 | 0.01823 | 192 | 0.252979335 | 6.54E-05 |
| SNP_A.K_1804778_11_7101390 | 11 | 7101390 | 2.12E-10 | 0.01042 | 192 | 0.249155177 | 7.35E-05 |
| SNP_A.K_685981_4_2521459 | 4 | 2521459 | 6.42E-10 | 0.05208 | 192 | 0.235063036 | 0.000168791 |
| SNP_A.K_729010_4_12427420 | 4 | 12427420 | 6.49E-10 | 0.20833 | 192 | 0.234932229 | 0.000168791 |
| SNP_A.K_730543_4_12698858 | 4 | 12698858 | 2.11E-09 | 0.21615 | 192 | 0.220132701 | 0.000316358 |
| SNP_A.K_157462_1_24620999 | 1 | 24620999 | 2.42E-09 | 0.05469 | 192 | 0.218459856 | 0.000316358 |
| SNP_A.K_730545_4_12698917 | 4 | 12698917 | 2.81E-09 | 0.21875 | 192 | 0.216585041 | 0.000316358 |
| SNP_A.K_730538_4_12698678 | 4 | 12698678 | 2.81E-09 | 0.21875 | 192 | 0.21657872 | 0.000316358 |
| SNP_A.K_729056_4_12506341 | 4 | 12506341 | 6.32E-09 | 0.21354 | 192 | 0.206587188 | 0.000316358 |
| SNP_A.K_1174598_7_1762446 | 7 | 1762446 | 6.74E-09 | 0.02865 | 192 | 0.205797728 | 0.000316358 |
| SNP_A.K_288103_2_4987974 | 2 | 4987974 | 7.60E-09 | 0.00781 | 192 | 0.204321736 | 0.000316358 |
| SNP_A.K_288114_2_4988672 | 2 | 4988672 | 7.60E-09 | 0.00781 | 192 | 0.204321736 | 0.000316358 |
| SNP_A.K_288116_2_4988805 | 2 | 4988805 | 7.60E-09 | 0.00781 | 192 | 0.204321736 | 0.000316358 |
| SNP_A.K_288122_2_4989029 | 2 | 4989029 | 7.60E-09 | 0.00781 | 192 | 0.204321736 | 0.000316358 |
| SNP_A.K_288123_2_4989036 | 2 | 4989036 | 7.60E-09 | 0.00781 | 192 | 0.204321736 | 0.000316358 |
| SNP_A.K_288132_2_4990502 | 2 | 4990502 | 7.60E-09 | 0.00781 | 192 | 0.204321736 | 0.000316358 |
| SNP_A.K_288136_2_4991126 | 2 | 4991126 | 7.60E-09 | 0.00781 | 192 | 0.204321736 | 0.000316358 |
| SNP_A.K_288144_2_4991792 | 2 | 4991792 | 7.60E-09 | 0.00781 | 192 | 0.204321736 | 0.000316358 |
| SNP_A.K_288145_2_4991816 | 2 | 4991816 | 7.60E-09 | 0.00781 | 192 | 0.204321736 | 0.000316358 |
| SNP_A.K_288150_2_4992218 | 2 | 4992218 | 7.60E-09 | 0.00781 | 192 | 0.204321736 | 0.000316358 |
| SNP_A.K_288157_2_4992816 | 2 | 4992816 | 7.60E-09 | 0.00781 | 192 | 0.204321736 | 0.000316358 |
| SNP_A.K_288158_2_4992821 | 2 | 4992821 | 7.60E-09 | 0.00781 | 192 | 0.204321736 | 0.000316358 |
| SNP_A.K_288163_2_4993339 | 2 | 4993339 | 7.60E-09 | 0.00781 | 192 | 0.204321736 | 0.000316358 |
| SNP_A.K_288164_2_4993351 | 2 | 4993351 | 7.60E-09 | 0.00781 | 192 | 0.204321736 | 0.000316358 |
| SNP_A.K_288178_2_4994046 | 2 | 4994046 | 7.60E-09 | 0.00781 | 192 | 0.204321736 | 0.000316358 |
| SNP_A.K_288189_2_4995195 | 2 | 4995195 | 7.60E-09 | 0.00781 | 192 | 0.204321736 | 0.000316358 |
| SNP_A.K_288197_2_4997655 | 2 | 4997655 | 7.60E-09 | 0.00781 | 192 | 0.204321736 | 0.000316358 |
| SNP_A.K_288279_2_5003424 | 2 | 5003424 | 7.60E-09 | 0.00781 | 192 | 0.204321736 | 0.000316358 |
| SNP_A.K_288290_2_5004458 | 2 | 5004458 | 7.60E-09 | 0.00781 | 192 | 0.204321736 | 0.000316358 |
| SNP_A.K_288493_2_5034560 | 2 | 5034560 | 7.60E-09 | 0.00781 | 192 | 0.204321736 | 0.000316358 |
| SNP_A.K_288563_2_5041476 | 2 | 5041476 | 7.60E-09 | 0.00781 | 192 | 0.204321736 | 0.000316358 |
| SNP_A.K_288565_2_5042437 | 2 | 5042437 | 7.60E-09 | 0.00781 | 192 | 0.204321736 | 0.000316358 |
| SNP_A.K_288092_2_4986793 | 2 | 4986793 | 7.60E-09 | 0.00781 | 192 | 0.204321736 | 0.000316358 |
| SNP_A.K_288113_2_4988667 | 2 | 4988667 | 7.60E-09 | 0.00781 | 192 | 0.204321736 | 0.000316358 |
| SNP_A.K_288120_2_4988990 | 2 | 4988990 | 7.60E-09 | 0.00781 | 192 | 0.204321736 | 0.000316358 |
| SNP_A.K_288125_2_4989212 | 2 | 4989212 | 7.60E-09 | 0.00781 | 192 | 0.204321736 | 0.000316358 |
| SNP_A.K_288146_2_4991828 | 2 | 4991828 | 7.60E-09 | 0.00781 | 192 | 0.204321736 | 0.000316358 |
| SNP_A.K_288156_2_4992518 | 2 | 4992518 | 7.60E-09 | 0.00781 | 192 | 0.204321736 | 0.000316358 |
| SNP_A.K_288184_2_4994991 | 2 | 4994991 | 7.60E-09 | 0.00781 | 192 | 0.204321736 | 0.000316358 |
| SNP_A.K_288194_2_4996097 | 2 | 4996097 | 7.60E-09 | 0.00781 | 192 | 0.204321736 | 0.000316358 |
| SNP_A.K_288199_2_4997717 | 2 | 4997717 | 7.60E-09 | 0.00781 | 192 | 0.204321736 | 0.000316358 |
| SNP_A.K_288285_2_5004065 | 2 | 5004065 | 7.60E-09 | 0.00781 | 192 | 0.204321736 | 0.000316358 |
| SNP_A.K_288287_2_5004370 | 2 | 5004370 | 7.60E-09 | 0.00781 | 192 | 0.204321736 | 0.000316358 |

| | | | | | | | |
|---|---|---|---|---|---|---|---|
| SNP_A.K_288399_2_5021434 | 2 | 5021434 | 7.60E-09 | 0.00781 | 192 | 0.204321736 | 0.000316358 |
| SNP_A.K_288492_2_5034545 | 2 | 5034545 | 7.60E-09 | 0.00781 | 192 | 0.204321736 | 0.000316358 |
| SNP_A.K_288500_2_5034670 | 2 | 5034670 | 7.60E-09 | 0.00781 | 192 | 0.204321736 | 0.000316358 |
| SNP_A.K_1924289_11_24625870 | 11 | 24625870 | 8.06E-09 | 0.00781 | 192 | 0.203592749 | 0.000329165 |
| SNP_A.K_1159169_6_29751394 | 6 | 29751394 | 9.57E-09 | 0.04427 | 192 | 0.201505066 | 0.000380769 |
| SNP_A.K_679358_4_1075655 | 4 | 1075655 | 9.70E-09 | 0.01302 | 192 | 0.201339957 | 0.000380769 |
| SNP_A.K_288425_2_5026783 | 2 | 5026783 | 9.89E-09 | 0.00781 | 192 | 0.201100453 | 0.000381113 |
| SNP_A.K_730546_4_12698925 | 4 | 12698925 | 1.76E-08 | 0.22396 | 192 | 0.194063514 | 0.000666865 |
| SNP_A.K_730533_4_12698403 | 4 | 12698403 | 2.22E-08 | 0.21875 | 192 | 0.191244483 | 0.000821583 |
| SNP_A.K_1175144_7_1827324 | 7 | 1827324 | 2.29E-08 | 0.03125 | 192 | 0.190891979 | 0.000821583 |
| SNP_A.K_1175146_7_1827355 | 7 | 1827355 | 2.29E-08 | 0.03125 | 192 | 0.190891979 | 0.000821583 |
| SNP_A.K_709666_4_8279704 | 4 | 8279704 | 2.85E-08 | 0.07552 | 192 | 0.188253272 | 0.001004809 |
| SNP_A.K_725892_4_11878318 | 4 | 11878318 | 3.30E-08 | 0.13281 | 192 | 0.186472334 | 0.001145376 |
| SNP_A.K_725933_4_11886168 | 4 | 11886168 | 3.38E-08 | 0.125 | 192 | 0.186198339 | 0.001152527 |
| SNP_A.K_706653_4_6930584 | 4 | 6930584 | 3.56E-08 | 0.14844 | 192 | 0.185578044 | 0.001193908 |
| SNP_A.K_709709_4_8282833 | 4 | 8282833 | 3.71E-08 | 0.07552 | 192 | 0.185057843 | 0.001226881 |
| SNP_A.K_708854_4_8117317 | 4 | 8117317 | 3.97E-08 | 0.07813 | 192 | 0.184260118 | 0.001279343 |
| SNP_A.K_709711_4_8282848 | 4 | 8282848 | 4.04E-08 | 0.07292 | 192 | 0.18403749 | 0.001279343 |
| SNP_A.K_708636_4_8069030 | 4 | 8069030 | 4.06E-08 | 0.07292 | 192 | 0.183995413 | 0.001279343 |
| SNP_A.K_708905_4_8123294 | 4 | 8123294 | 4.39E-08 | 0.07292 | 192 | 0.183047585 | 0.00136376 |
| SNP_A.K_709119_4_8229490 | 4 | 8229490 | 4.92E-08 | 0.08073 | 192 | 0.181672855 | 0.001506903 |
| SNP_A.K_1175140_7_1827092 | 7 | 1827092 | 5.33E-08 | 0.03385 | 192 | 0.18072936 | 0.001606762 |
| SNP_A.K_1028690_6_10678950 | 6 | 10678950 | 5.54E-08 | 0.04167 | 192 | 0.180248307 | 0.001648754 |
| SNP_A.K_708866_4_8119634 | 4 | 8119634 | 5.80E-08 | 0.07552 | 192 | 0.179704488 | 0.001682483 |
| SNP_A.K_1813672_11_8565228 | 11 | 8565228 | 5.82E-08 | 0.14323 | 192 | 0.179669008 | 0.001682483 |
| SNP_A.K_1552916_9_9224803 | 9 | 9224803 | 6.07E-08 | 0.23177 | 192 | 0.179169635 | 0.001730208 |
| SNP_A.K_708816_4_8115086 | 4 | 8115086 | 6.39E-08 | 0.05469 | 192 | 0.178546584 | 0.001798178 |
| SNP_A.K_708864_4_8119296 | 4 | 8119296 | 7.16E-08 | 0.07292 | 192 | 0.177189339 | 0.001944605 |
| SNP_A.K_709754_4_8284876 | 4 | 8284876 | 7.26E-08 | 0.07552 | 192 | 0.177024612 | 0.001944605 |
| SNP_A.K_709718_4_8283165 | 4 | 8283165 | 7.32E-08 | 0.07552 | 192 | 0.176926237 | 0.001944605 |
| SNP_A.K_708762_4_8111293 | 4 | 8111293 | 7.37E-08 | 0.07552 | 192 | 0.176845436 | 0.001944605 |
| SNP_A.K_288155_2_4992514 | 2 | 4992514 | 7.38E-08 | 0.01042 | 192 | 0.176831818 | 0.001944605 |
| SNP_A.K_709712_4_8282869 | 4 | 8282869 | 7.50E-08 | 0.07552 | 192 | 0.17663705 | 0.001947865 |
| SNP_A.K_288501_2_5034949 | 2 | 5034949 | 7.66E-08 | 0.01042 | 192 | 0.176387863 | 0.001947865 |
| SNP_A.K_708808_4_8114964 | 4 | 8114964 | 7.76E-08 | 0.08073 | 192 | 0.176230756 | 0.001947865 |
| SNP_A.K_288133_2_4990519 | 2 | 4990519 | 7.77E-08 | 0.01042 | 192 | 0.176223025 | 0.001947865 |
| SNP_A.K_724652_4_11653571 | 4 | 11653571 | 8.02E-08 | 0.21615 | 192 | 0.175846218 | 0.001963953 |
| SNP_A.K_709717_4_8283123 | 4 | 8283123 | 8.02E-08 | 0.08333 | 192 | 0.175841311 | 0.001963953 |
| SNP_A.K_1552868_9_9222748 | 9 | 9222748 | 8.31E-08 | 0.19271 | 192 | 0.175417889 | 0.001999632 |
| SNP_A.K_709278_4_8245620 | 4 | 8245620 | 8.46E-08 | 0.08073 | 192 | 0.175211783 | 0.001999632 |
| SNP_A.K_708852_4_8117281 | 4 | 8117281 | 8.46E-08 | 0.07813 | 192 | 0.175201379 | 0.001999632 |
| SNP_A.K_1552915_9_9224800 | 9 | 9224800 | 8.55E-08 | 0.2474 | 192 | 0.175079327 | 0.001999632 |
| SNP_A.K_288499_2_5034621 | 2 | 5034621 | 8.72E-08 | 0.01042 | 192 | 0.174839166 | 0.00201772 |
| SNP_A.K_1552942_9_9225590 | 9 | 9225590 | 8.89E-08 | 0.1901 | 192 | 0.174611672 | 0.002034067 |
| SNP_A.K_709714_4_8282955 | 4 | 8282955 | 9.45E-08 | 0.08073 | 192 | 0.173887109 | 0.002041189 |
| SNP_A.K_288384_2_5019012 | 2 | 5019012 | 9.57E-08 | 0.01042 | 192 | 0.173742773 | 0.002041189 |
| SNP_A.K_288112_2_4988622 | 2 | 4988622 | 9.57E-08 | 0.01042 | 192 | 0.173739233 | 0.002041189 |
| SNP_A.K_288322_2_5007971 | 2 | 5007971 | 9.57E-08 | 0.01042 | 192 | 0.173739233 | 0.002041189 |
| SNP_A.K_288550_2_5038140 | 2 | 5038140 | 9.57E-08 | 0.01042 | 192 | 0.173739233 | 0.002041189 |
| SNP_A.K_1939603_11_28480520 | 11 | 28480520 | 9.92E-08 | 0.03646 | 192 | 0.173308585 | 0.002041189 |

| | | | | | | | |
|---|---|---|---|---|---|---|---|
| SNP_A.K_1552943_9_9225600 | 9 | 9225600 | 1.01E-07 | 0.1875 | 192 | 0.173115523 | 0.002041189 |
| SNP_A.K_708809_4_8114977 | 4 | 8114977 | 1.02E-07 | 0.08073 | 192 | 0.173017302 | 0.002041189 |
| SNP_A.K_708645_4_8070065 | 4 | 8070065 | 1.02E-07 | 0.08333 | 192 | 0.172944751 | 0.002041189 |
| SNP_A.K_1552874_9_9223041 | 9 | 9223041 | 1.08E-07 | 0.20052 | 192 | 0.172312673 | 0.002041189 |
| SNP_A.K_730540_4_12698747 | 4 | 12698747 | 1.08E-07 | 0.22917 | 192 | 0.172255916 | 0.002041189 |
| SNP_A.K_725879_4_11877029 | 4 | 11877029 | 1.09E-07 | 0.13281 | 192 | 0.172196628 | 0.002041189 |
| SNP_A.K_288497_2_5034592 | 2 | 5034592 | 1.10E-07 | 0.01042 | 192 | 0.172034238 | 0.002041189 |
| SNP_A.K_288139_2_4991452 | 2 | 4991452 | 1.14E-07 | 0.01042 | 192 | 0.171711709 | 0.002041189 |
| SNP_A.K_288398_2_5021121 | 2 | 5021121 | 1.14E-07 | 0.01042 | 192 | 0.17164415 | 0.002041189 |
| SNP_A.K_708810_4_8114983 | 4 | 8114983 | 1.14E-07 | 0.08073 | 192 | 0.171631196 | 0.002041189 |
| SNP_A.K_708811_4_8114995 | 4 | 8114995 | 1.19E-07 | 0.07813 | 192 | 0.171130021 | 0.002041189 |
| SNP_A.K_288376_2_5017884 | 2 | 5017884 | 1.19E-07 | 0.01042 | 192 | 0.171125272 | 0.002041189 |
| SNP_A.K_724877_4_11690622 | 4 | 11690622 | 1.20E-07 | 0.21615 | 192 | 0.17102922 | 0.002041189 |
| SNP_A.K_708655_4_8071479 | 4 | 8071479 | 1.21E-07 | 0.07292 | 192 | 0.170962382 | 0.002041189 |
| SNP_A.K_288449_2_5028561 | 2 | 5028561 | 1.21E-07 | 0.01042 | 192 | 0.170914639 | 0.002041189 |
| SNP_A.K_709546_4_8273241 | 4 | 8273241 | 1.22E-07 | 0.07813 | 192 | 0.170899031 | 0.002041189 |
| SNP_A.K_1953646_12_3610642 | 12 | 3610642 | 1.23E-07 | 0.07552 | 192 | 0.170714354 | 0.002041189 |
| SNP_A.K_1027527_6_10421615 | 6 | 10421615 | 1.26E-07 | 0.03125 | 192 | 0.170491664 | 0.002041189 |
| SNP_A.K_1552866_9_9222708 | 9 | 9222708 | 1.26E-07 | 0.20052 | 192 | 0.170457063 | 0.002041189 |
| SNP_A.K_708807_4_8114955 | 4 | 8114955 | 1.27E-07 | 0.08333 | 192 | 0.170379353 | 0.002041189 |
| SNP_A.K_725248_4_11767129 | 4 | 11767129 | 1.28E-07 | 0.21354 | 192 | 0.170320189 | 0.002041189 |
| SNP_A.K_709741_4_8284570 | 4 | 8284570 | 1.29E-07 | 0.07552 | 192 | 0.170167379 | 0.002041189 |
| SNP_A.K_708859_4_8118449 | 4 | 8118449 | 1.30E-07 | 0.07813 | 192 | 0.170086088 | 0.002041189 |
| SNP_A.K_709729_4_8284091 | 4 | 8284091 | 1.36E-07 | 0.07552 | 192 | 0.169586071 | 0.002041189 |
| SNP_A.K_288159_2_4992884 | 2 | 4992884 | 1.37E-07 | 0.01042 | 192 | 0.169464472 | 0.002041189 |
| SNP_A.K_723720_4_11498296 | 4 | 11498296 | 1.38E-07 | 0.21354 | 192 | 0.169417047 | 0.002041189 |
| SNP_A.K_288173_2_4993935 | 2 | 4993935 | 1.39E-07 | 0.01042 | 192 | 0.169329644 | 0.002041189 |
| SNP_A.K_288177_2_4994000 | 2 | 4994000 | 1.39E-07 | 0.01042 | 192 | 0.169329644 | 0.002041189 |
| SNP_A.K_1487843_8_26986462 | 8 | 26986462 | 1.41E-07 | 0.03906 | 192 | 0.169121119 | 0.002041189 |
| SNP_A.K_709279_4_8245639 | 4 | 8245639 | 1.43E-07 | 0.08333 | 192 | 0.168987881 | 0.002041189 |
| SNP_A.K_709273_4_8245255 | 4 | 8245255 | 1.43E-07 | 0.07813 | 192 | 0.168961616 | 0.002041189 |
| SNP_A.K_1487950_8_26990220 | 8 | 26990220 | 1.45E-07 | 0.03385 | 192 | 0.168792833 | 0.002041189 |
| SNP_A.K_725895_4_11878735 | 4 | 11878735 | 1.46E-07 | 0.13021 | 192 | 0.168725207 | 0.002041189 |
| SNP_A.K_1552848_9_9222236 | 9 | 9222236 | 1.47E-07 | 0.1849 | 192 | 0.168673885 | 0.002041189 |
| SNP_A.K_709696_4_8281788 | 4 | 8281788 | 1.48E-07 | 0.08073 | 192 | 0.168592023 | 0.002041189 |
| SNP_A.K_288160_2_4993128 | 2 | 4993128 | 1.48E-07 | 0.01042 | 192 | 0.168574117 | 0.002041189 |
| SNP_A.K_288119_2_4988952 | 2 | 4988952 | 1.52E-07 | 0.01042 | 192 | 0.16822123 | 0.002041189 |
| SNP_A.K_724618_4_11646202 | 4 | 11646202 | 1.53E-07 | 0.17188 | 192 | 0.168177821 | 0.002041189 |
| SNP_A.K_288105_2_4988403 | 2 | 4988403 | 1.53E-07 | 0.01042 | 192 | 0.168155529 | 0.002041189 |
| SNP_A.K_288110_2_4988498 | 2 | 4988498 | 1.53E-07 | 0.01042 | 192 | 0.168155529 | 0.002041189 |
| SNP_A.K_288111_2_4988565 | 2 | 4988565 | 1.53E-07 | 0.01042 | 192 | 0.168155529 | 0.002041189 |
| SNP_A.K_288118_2_4988884 | 2 | 4988884 | 1.53E-07 | 0.01042 | 192 | 0.168155529 | 0.002041189 |
| SNP_A.K_288131_2_4990339 | 2 | 4990339 | 1.53E-07 | 0.01042 | 192 | 0.168155529 | 0.002041189 |
| SNP_A.K_288152_2_4992317 | 2 | 4992317 | 1.53E-07 | 0.01042 | 192 | 0.168155529 | 0.002041189 |
| SNP_A.K_288196_2_4997387 | 2 | 4997387 | 1.53E-07 | 0.01042 | 192 | 0.168155529 | 0.002041189 |
| SNP_A.K_288106_2_4988412 | 2 | 4988412 | 1.53E-07 | 0.01042 | 192 | 0.168155529 | 0.002041189 |
| SNP_A.K_288153_2_4992344 | 2 | 4992344 | 1.53E-07 | 0.01042 | 192 | 0.168155529 | 0.002041189 |
| SNP_A.K_288180_2_4994173 | 2 | 4994173 | 1.53E-07 | 0.01042 | 192 | 0.168155529 | 0.002041189 |
| SNP_A.K_288214_2_4998711 | 2 | 4998711 | 1.53E-07 | 0.01042 | 192 | 0.168155529 | 0.002041189 |
| SNP_A.K_726928_4_12057294 | 4 | 12057294 | 1.53E-07 | 0.22396 | 192 | 0.16815213 | 0.002041189 |

| | | | | | | | |
|---|---|---|---|---|---|---|---|
| SNP_A.K_288108_2_4988457 | 2 | 4988457 | 1.54E-07 | 0.01042 | 192 | 0.168130261 | 0.002041189 |
| SNP_A.K_288511_2_5035693 | 2 | 5035693 | 1.56E-07 | 0.01042 | 192 | 0.167926711 | 0.002041189 |
| SNP_A.K_1552865_9_9222655 | 9 | 9222655 | 1.57E-07 | 0.20573 | 192 | 0.167885856 | 0.002041189 |
| SNP_A.K_288583_2_5045331 | 2 | 5045331 | 1.57E-07 | 0.01042 | 192 | 0.167860935 | 0.002041189 |
| SNP_A.K_288713_2_5061585 | 2 | 5061585 | 1.57E-07 | 0.01042 | 192 | 0.167860935 | 0.002041189 |
| SNP_A.K_709177_4_8236729 | 4 | 8236729 | 1.57E-07 | 0.02865 | 192 | 0.167853945 | 0.002041189 |
| SNP_A.K_288121_2_4988997 | 2 | 4988997 | 1.57E-07 | 0.01042 | 192 | 0.167840867 | 0.002041189 |
| SNP_A.K_288166_2_4993376 | 2 | 4993376 | 1.58E-07 | 0.01042 | 192 | 0.167783942 | 0.002041189 |
| SNP_A.K_288186_2_4995030 | 2 | 4995030 | 1.60E-07 | 0.01042 | 192 | 0.167637558 | 0.002041189 |
| SNP_A.K_288187_2_4995034 | 2 | 4995034 | 1.60E-07 | 0.01042 | 192 | 0.167637558 | 0.002041189 |
| SNP_A.K_288138_2_4991416 | 2 | 4991416 | 1.61E-07 | 0.01042 | 192 | 0.167594887 | 0.002041189 |
| SNP_A.K_708911_4_8124310 | 4 | 8124310 | 1.62E-07 | 0.07813 | 192 | 0.167538547 | 0.002041189 |
| SNP_A.K_288307_2_5006319 | 2 | 5006319 | 1.62E-07 | 0.01042 | 192 | 0.167467315 | 0.002041189 |
| SNP_A.K_288212_2_4998626 | 2 | 4998626 | 1.64E-07 | 0.01042 | 192 | 0.167379437 | 0.002041189 |
| SNP_A.K_708869_4_8120145 | 4 | 8120145 | 1.64E-07 | 0.07552 | 192 | 0.167331857 | 0.002041189 |
| SNP_A.K_709740_4_8284561 | 4 | 8284561 | 1.65E-07 | 0.07552 | 192 | 0.167311345 | 0.002041189 |
| SNP_A.K_288140_2_4991514 | 2 | 4991514 | 1.66E-07 | 0.01042 | 192 | 0.167218687 | 0.002041189 |
| SNP_A.K_288185_2_4995012 | 2 | 4995012 | 1.66E-07 | 0.01042 | 192 | 0.167197817 | 0.002041189 |
| SNP_A.K_288341_2_5011285 | 2 | 5011285 | 1.66E-07 | 0.01042 | 192 | 0.167182423 | 0.002041189 |
| SNP_A.K_709069_4_8216219 | 4 | 8216219 | 1.67E-07 | 0.07552 | 192 | 0.167120799 | 0.002041189 |
| SNP_A.K_709114_4_8227467 | 4 | 8227467 | 1.67E-07 | 0.08333 | 192 | 0.167111604 | 0.002041189 |
| SNP_A.K_708815_4_8115069 | 4 | 8115069 | 1.68E-07 | 0.08333 | 192 | 0.167039277 | 0.002041189 |
| SNP_A.K_708574_4_8062246 | 4 | 8062246 | 1.70E-07 | 0.08333 | 192 | 0.166938331 | 0.002041189 |
| SNP_A.K_1552921_9_9224901 | 9 | 9224901 | 1.70E-07 | 0.21615 | 192 | 0.166901486 | 0.002041189 |
| SNP_A.K_709171_4_8236402 | 4 | 8236402 | 1.71E-07 | 0.08333 | 192 | 0.166892615 | 0.002041189 |
| SNP_A.K_730537_4_12698650 | 4 | 12698650 | 1.71E-07 | 0.20573 | 192 | 0.166849437 | 0.002041189 |
| SNP_A.K_709627_4_8278535 | 4 | 8278535 | 1.71E-07 | 0.07552 | 192 | 0.166840211 | 0.002041189 |
| SNP_A.K_709698_4_8282162 | 4 | 8282162 | 1.72E-07 | 0.08333 | 192 | 0.166821933 | 0.002041189 |
| SNP_A.K_724792_4_11681871 | 4 | 11681871 | 1.74E-07 | 0.22135 | 192 | 0.166669976 | 0.002054887 |
| SNP_A.K_708580_4_8062495 | 4 | 8062495 | 1.76E-07 | 0.07552 | 192 | 0.166550973 | 0.002054887 |
| SNP_A.K_1552922_9_9224908 | 9 | 9224908 | 1.76E-07 | 0.21615 | 192 | 0.166542475 | 0.002054887 |
| SNP_A.K_723464_4_11458037 | 4 | 11458037 | 1.78E-07 | 0.21875 | 192 | 0.166423226 | 0.002058082 |
| SNP_A.K_726053_4_11897104 | 4 | 11897104 | 1.78E-07 | 0.22135 | 192 | 0.166392351 | 0.002058082 |
| SNP_A.K_1813026_11_8487695 | 11 | 8487695 | 1.83E-07 | 0.13542 | 192 | 0.166095055 | 0.002091963 |
| SNP_A.K_1552867_9_9222718 | 9 | 9222718 | 1.83E-07 | 0.20052 | 192 | 0.166069487 | 0.002091963 |
| SNP_A.K_730495_4_12696561 | 4 | 12696561 | 1.84E-07 | 0.22135 | 192 | 0.165973231 | 0.002097586 |
| SNP_A.K_1028765_6_10682951 | 6 | 10682951 | 1.88E-07 | 0.02083 | 192 | 0.165730205 | 0.002129642 |
| SNP_A.K_709239_4_8242287 | 4 | 8242287 | 1.92E-07 | 0.07813 | 192 | 0.165519078 | 0.00215592 |
| SNP_A.K_709227_4_8241958 | 4 | 8241958 | 1.93E-07 | 0.07552 | 192 | 0.16545828 | 0.00215592 |
| SNP_A.K_723779_4_11515937 | 4 | 11515937 | 2.11E-07 | 0.22396 | 192 | 0.164405144 | 0.002344994 |
| SNP_A.K_723257_4_11423648 | 4 | 11423648 | 2.18E-07 | 0.25781 | 192 | 0.164007295 | 0.002412731 |
| SNP_A.K_727195_4_12092846 | 4 | 12092846 | 2.19E-07 | 0.21354 | 192 | 0.163938447 | 0.002414052 |
| SNP_A.K_709719_4_8283243 | 4 | 8283243 | 2.22E-07 | 0.08073 | 192 | 0.163774767 | 0.002423456 |
| SNP_A.K_725554_4_11821281 | 4 | 11821281 | 2.22E-07 | 0.17448 | 192 | 0.16376639 | 0.002423456 |
| SNP_A.K_728188_4_12288051 | 4 | 12288051 | 2.24E-07 | 0.22135 | 192 | 0.163702381 | 0.002423456 |
| SNP_A.K_730552_4_12699438 | 4 | 12699438 | 2.25E-07 | 0.19792 | 192 | 0.163646382 | 0.002423456 |
| SNP_A.K_723345_4_11434442 | 4 | 11434442 | 2.27E-07 | 0.22135 | 192 | 0.163546458 | 0.002431538 |
| SNP_A.K_708853_4_8117289 | 4 | 8117289 | 2.32E-07 | 0.08073 | 192 | 0.163294595 | 0.002471448 |
| SNP_A.K_709382_4_8263107 | 4 | 8263107 | 2.36E-07 | 0.07813 | 192 | 0.163090263 | 0.002501955 |
| SNP_A.K_708764_4_8111318 | 4 | 8111318 | 2.38E-07 | 0.08073 | 192 | 0.162960482 | 0.002516896 |

| | | | | | | | |
|---|---|---|---|---|---|---|---|
| SNP_A.K_725139_4_11740931 | 4 | 11740931 | 2.44E-07 | 0.26302 | 192 | 0.162675312 | 0.002565706 |
| SNP_A.K_709534_4_8272471 | 4 | 8272471 | 2.48E-07 | 0.07813 | 192 | 0.162497344 | 0.002591783 |
| SNP_A.K_253295_1_40881735 | 1 | 40881735 | 2.51E-07 | 0.14844 | 192 | 0.162359188 | 0.002609339 |
| SNP_A.K_1552927_9_9225039 | 9 | 9225039 | 2.57E-07 | 0.20573 | 192 | 0.162057897 | 0.00266384 |
| SNP_A.K_708766_4_8111373 | 4 | 8111373 | 2.61E-07 | 0.08594 | 192 | 0.161871862 | 0.002692994 |
| SNP_A.K_708736_4_8089953 | 4 | 8089953 | 2.63E-07 | 0.07813 | 192 | 0.161777085 | 0.002701455 |
| SNP_A.K_709111_4_8226407 | 4 | 8226407 | 2.72E-07 | 0.08333 | 192 | 0.161395276 | 0.002777121 |
| SNP_A.K_708763_4_8111298 | 4 | 8111298 | 2.74E-07 | 0.07813 | 192 | 0.16130778 | 0.002784263 |
| SNP_A.K_723460_4_11457805 | 4 | 11457805 | 2.77E-07 | 0.21094 | 192 | 0.161181122 | 0.002800828 |
| SNP_A.K_709426_4_8265823 | 4 | 8265823 | 2.79E-07 | 0.11458 | 192 | 0.16108772 | 0.002809584 |
| SNP_A.K_726174_4_11926870 | 4 | 11926870 | 2.87E-07 | 0.10677 | 192 | 0.160785916 | 0.002868977 |
| SNP_A.K_709200_4_8238535 | 4 | 8238535 | 2.89E-07 | 0.0625 | 192 | 0.160711343 | 0.00287347 |
| SNP_A.K_1552876_9_9223555 | 9 | 9223555 | 2.96E-07 | 0.20313 | 192 | 0.160425267 | 0.002930444 |
| SNP_A.K_1815425_11_8851660 | 11 | 8851660 | 2.98E-07 | 0.14323 | 192 | 0.160326649 | 0.002941203 |
| SNP_A.K_725249_4_11767161 | 4 | 11767161 | 3.05E-07 | 0.21875 | 192 | 0.160047912 | 0.002997807 |
| SNP_A.K_709665_4_8279691 | 4 | 8279691 | 3.17E-07 | 0.22135 | 192 | 0.159608701 | 0.003097753 |
| SNP_A.K_724241_4_11583139 | 4 | 11583139 | 3.24E-07 | 0.11198 | 192 | 0.159357484 | 0.003150154 |
| SNP_A.K_730550_4_12699355 | 4 | 12699355 | 3.29E-07 | 0.20313 | 192 | 0.159166856 | 0.003184067 |
| SNP_A.K_1953652_12_3611028 | 12 | 3611028 | 3.30E-07 | 0.08333 | 192 | 0.15912328 | 0.003184067 |
| SNP_A.K_709737_4_8284482 | 4 | 8284482 | 3.41E-07 | 0.08073 | 192 | 0.158751375 | 0.003271753 |
| SNP_A.K_1815428_11_8851722 | 11 | 8851722 | 3.45E-07 | 0.15104 | 192 | 0.158632163 | 0.003290106 |
| SNP_A.K_708902_4_8122881 | 4 | 8122881 | 3.48E-07 | 0.08073 | 192 | 0.158527658 | 0.003304477 |
| SNP_A.K_709738_4_8284489 | 4 | 8284489 | 3.55E-07 | 0.08073 | 192 | 0.158271901 | 0.003362196 |
| SNP_A.K_1552894_9_9224143 | 9 | 9224143 | 3.62E-07 | 0.21354 | 192 | 0.158047692 | 0.003411793 |
| SNP_A.K_1813663_11_8564877 | 11 | 8564877 | 3.68E-07 | 0.14063 | 192 | 0.15785987 | 0.003444294 |
| SNP_A.K_1552953_9_9225791 | 9 | 9225791 | 3.69E-07 | 0.20833 | 192 | 0.157823402 | 0.003444294 |
| SNP_A.K_710508_4_8425491 | 4 | 8425491 | 3.71E-07 | 0.08594 | 192 | 0.1577793 | 0.003444294 |
| SNP_A.K_1552878_9_9223677 | 9 | 9223677 | 3.76E-07 | 0.20833 | 192 | 0.157613642 | 0.003477947 |
| SNP_A.K_1552935_9_9225150 | 9 | 9225150 | 3.79E-07 | 0.21094 | 192 | 0.157504879 | 0.003484237 |
| SNP_A.K_730535_4_12698463 | 4 | 12698463 | 3.80E-07 | 0.20833 | 192 | 0.157489147 | 0.003484237 |
| SNP_A.K_1552920_9_9224844 | 9 | 9224844 | 3.97E-07 | 0.21354 | 192 | 0.156980299 | 0.003611979 |
| SNP_A.K_1815426_11_8851679 | 11 | 8851679 | 3.97E-07 | 0.15365 | 192 | 0.1569662 | 0.003611979 |
| SNP_A.K_708834_4_8115542 | 4 | 8115542 | 3.99E-07 | 0.08594 | 192 | 0.156904646 | 0.003615285 |
| SNP_A.K_709282_4_8245664 | 4 | 8245664 | 4.08E-07 | 0.08594 | 192 | 0.156646602 | 0.003680097 |
| SNP_A.K_708733_4_8089817 | 4 | 8089817 | 4.26E-07 | 0.08073 | 192 | 0.156154365 | 0.003803738 |
| SNP_A.K_708765_4_8111323 | 4 | 8111323 | 4.27E-07 | 0.08073 | 192 | 0.156132996 | 0.003803738 |
| SNP_A.K_709228_4_8241962 | 4 | 8241962 | 4.28E-07 | 0.08073 | 192 | 0.156110492 | 0.003803738 |
| SNP_A.K_708830_4_8115387 | 4 | 8115387 | 4.35E-07 | 0.08073 | 192 | 0.155903938 | 0.003851781 |
| SNP_A.K_1552923_9_9224924 | 9 | 9224924 | 4.39E-07 | 0.20573 | 192 | 0.155810635 | 0.003851781 |
| SNP_A.K_724233_4_11582596 | 4 | 11582596 | 4.42E-07 | 0.13021 | 192 | 0.155721576 | 0.003851781 |
| SNP_A.K_709556_4_8273703 | 4 | 8273703 | 4.42E-07 | 0.08594 | 192 | 0.15571795 | 0.003851781 |
| SNP_A.K_1813673_11_8565241 | 11 | 8565241 | 4.42E-07 | 0.15625 | 192 | 0.155717567 | 0.003851781 |
| SNP_A.K_1487409_8_26904720 | 8 | 26904720 | 4.46E-07 | 0.01823 | 192 | 0.155609772 | 0.003871362 |
| SNP_A.K_709046_4_8214906 | 4 | 8214906 | 4.51E-07 | 0.08073 | 192 | 0.155486637 | 0.003884706 |
| SNP_A.K_1175143_7_1827317 | 7 | 1827317 | 4.53E-07 | 0.04167 | 192 | 0.155435542 | 0.003884706 |
| SNP_A.K_1552871_9_9222892 | 9 | 9222892 | 4.54E-07 | 0.20833 | 192 | 0.155424858 | 0.003884706 |
| SNP_A.K_709027_4_8213731 | 4 | 8213731 | 4.60E-07 | 0.08073 | 192 | 0.155248922 | 0.003922582 |
| SNP_A.K_708973_4_8211776 | 4 | 8211776 | 4.62E-07 | 0.08594 | 192 | 0.155216255 | 0.003922582 |
| SNP_A.K_2023179_12_15535035 | 12 | 15535035 | 4.65E-07 | 0.09635 | 192 | 0.155141153 | 0.003931898 |
| SNP_A.K_709693_4_8281558 | 4 | 8281558 | 4.71E-07 | 0.08854 | 192 | 0.154988463 | 0.003967638 |

| | | | | | | | |
|---|---|---|---|---|---|---|---|
| SNP_A.K_709557_4_8273707 | 4 | 8273707 | 4.74E-07 | 0.08594 | 192 | 0.154922056 | 0.00397423 |
| SNP_A.K_725782_4_11865314 | 4 | 11865314 | 4.84E-07 | 0.18229 | 192 | 0.154671435 | 0.004044365 |
| SNP_A.K_1551668_9_9119953 | 9 | 9119953 | 4.91E-07 | 0.33594 | 192 | 0.154495708 | 0.004089437 |
| SNP_A.K_710534_4_8426668 | 4 | 8426668 | 4.93E-07 | 0.08333 | 192 | 0.154444535 | 0.004091086 |
| SNP_A.K_709124_4_8231016 | 4 | 8231016 | 5.13E-07 | 0.08333 | 192 | 0.15399812 | 0.004234167 |
| SNP_A.K_940970_5_22811089 | 5 | 22811089 | 5.18E-07 | 0.02604 | 192 | 0.153876746 | 0.004261652 |
| SNP_A.K_1552392_9_9174341 | 9 | 9174341 | 5.24E-07 | 0.15625 | 192 | 0.15373881 | 0.0042955 |
| SNP_A.K_724243_4_11583164 | 4 | 11583164 | 5.30E-07 | 0.11979 | 192 | 0.153612046 | 0.004319163 |
| SNP_A.K_1028761_6_10682850 | 6 | 10682850 | 5.31E-07 | 0.02344 | 192 | 0.153582283 | 0.004319163 |
| SNP_A.K_1552847_9_9222219 | 9 | 9222219 | 5.35E-07 | 0.21354 | 192 | 0.153505553 | 0.004319163 |
| SNP_A.K_708863_4_8118805 | 4 | 8118805 | 5.35E-07 | 0.09115 | 192 | 0.153493153 | 0.004319163 |
| SNP_A.K_709092_4_8222010 | 4 | 8222010 | 5.45E-07 | 0.08594 | 192 | 0.153282072 | 0.004381294 |
| SNP_A.K_724334_4_11588870 | 4 | 11588870 | 5.70E-07 | 0.1276 | 192 | 0.152763051 | 0.004555473 |
| SNP_A.K_288401_2_5021522 | 2 | 5021522 | 5.72E-07 | 0.01302 | 192 | 0.152723179 | 0.004555473 |
| SNP_A.K_709017_4_8212961 | 4 | 8212961 | 5.75E-07 | 0.08594 | 192 | 0.15266348 | 0.004555473 |
| SNP_A.K_1028692_6_10679118 | 6 | 10679118 | 5.76E-07 | 0.02344 | 192 | 0.152650777 | 0.004555473 |
| SNP_A.K_1028707_6_10680163 | 6 | 10680163 | 5.80E-07 | 0.02344 | 192 | 0.152562927 | 0.004572661 |
| SNP_A.K_709105_4_8224626 | 4 | 8224626 | 5.85E-07 | 0.08333 | 192 | 0.152468023 | 0.004585208 |
| SNP_A.K_1551737_9_9124097 | 9 | 9124097 | 5.86E-07 | 0.28385 | 192 | 0.152443411 | 0.004585208 |
| SNP_A.K_1551753_9_9124517 | 9 | 9124517 | 5.93E-07 | 0.28385 | 192 | 0.152313221 | 0.00461952 |
| SNP_A.K_1552946_9_9225741 | 9 | 9225741 | 6.00E-07 | 0.20833 | 192 | 0.152176045 | 0.004656963 |
| SNP_A.K_729015_4_12427666 | 4 | 12427666 | 6.14E-07 | 0.24479 | 192 | 0.151901047 | 0.004750842 |
| SNP_A.K_709409_4_8265347 | 4 | 8265347 | 6.22E-07 | 0.08333 | 192 | 0.151757706 | 0.004784878 |
| SNP_A.K_708972_4_8211753 | 4 | 8211753 | 6.23E-07 | 0.09115 | 192 | 0.151732186 | 0.004784878 |
| SNP_A.K_1552957_9_9226039 | 9 | 9226039 | 6.36E-07 | 0.24479 | 192 | 0.151495948 | 0.004832091 |
| SNP_A.K_708821_4_8115237 | 4 | 8115237 | 6.37E-07 | 0.08333 | 192 | 0.15147354 | 0.004832091 |
| SNP_A.K_1551752_9_9124501 | 9 | 9124501 | 6.41E-07 | 0.28906 | 192 | 0.151396045 | 0.004832091 |
| SNP_A.K_726909_4_12056420 | 4 | 12056420 | 6.41E-07 | 0.22917 | 192 | 0.151394474 | 0.004832091 |
| SNP_A.K_708587_4_8062950 | 4 | 8062950 | 6.42E-07 | 0.08594 | 192 | 0.151387077 | 0.004832091 |
| SNP_A.K_708833_4_8115517 | 4 | 8115517 | 6.43E-07 | 0.08333 | 192 | 0.151364096 | 0.004832091 |
| SNP_A.K_730534_4_12698429 | 4 | 12698429 | 6.53E-07 | 0.21094 | 192 | 0.151193773 | 0.004885914 |
| SNP_A.K_709115_4_8227722 | 4 | 8227722 | 6.57E-07 | 0.09115 | 192 | 0.151108988 | 0.004904115 |
| SNP_A.K_686026_4_2522606 | 4 | 2522606 | 6.76E-07 | 0.08073 | 192 | 0.150779962 | 0.005027221 |
| SNP_A.K_709554_4_8273677 | 4 | 8273677 | 7.00E-07 | 0.08854 | 192 | 0.150380465 | 0.005174339 |
| SNP_A.K_729300_4_12590909 | 4 | 12590909 | 7.01E-07 | 0.22135 | 192 | 0.150363071 | 0.005174339 |
| SNP_A.K_709116_4_8227885 | 4 | 8227885 | 7.04E-07 | 0.09115 | 192 | 0.150318661 | 0.00517585 |
| SNP_A.K_709281_4_8245647 | 4 | 8245647 | 7.14E-07 | 0.1224 | 192 | 0.1501572 | 0.00522999 |
| SNP_A.K_709463_4_8267868 | 4 | 8267868 | 7.17E-07 | 0.09115 | 192 | 0.150106525 | 0.005234483 |
| SNP_A.K_708912_4_8124548 | 4 | 8124548 | 7.33E-07 | 0.08854 | 192 | 0.149846085 | 0.005334773 |
| SNP_A.K_1551904_9_9131739 | 9 | 9131739 | 7.48E-07 | 0.28125 | 192 | 0.149606301 | 0.005427398 |
| SNP_A.K_724300_4_11587032 | 4 | 11587032 | 7.57E-07 | 0.11979 | 192 | 0.14947625 | 0.005463958 |
| SNP_A.K_709036_4_8214282 | 4 | 8214282 | 7.59E-07 | 0.08594 | 192 | 0.149448154 | 0.005463958 |
| SNP_A.K_708813_4_8115020 | 4 | 8115020 | 7.69E-07 | 0.08594 | 192 | 0.149284813 | 0.005522487 |
| SNP_A.K_1552913_9_9224737 | 9 | 9224737 | 7.74E-07 | 0.20833 | 192 | 0.14921981 | 0.005534502 |
| SNP_A.K_1552940_9_9225513 | 9 | 9225513 | 7.81E-07 | 0.21354 | 192 | 0.149110857 | 0.005567713 |
| SNP_A.K_708554_4_8061283 | 4 | 8061283 | 7.94E-07 | 0.08594 | 192 | 0.148923438 | 0.005619941 |
| SNP_A.K_708814_4_8115034 | 4 | 8115034 | 7.95E-07 | 0.09375 | 192 | 0.14890824 | 0.005619941 |
| SNP_A.K_708822_4_8115248 | 4 | 8115248 | 7.96E-07 | 0.08594 | 192 | 0.148884525 | 0.005619941 |
| SNP_A.K_709739_4_8284495 | 4 | 8284495 | 8.01E-07 | 0.08854 | 192 | 0.148824625 | 0.005630027 |
| SNP_A.K_709084_4_8219294 | 4 | 8219294 | 8.05E-07 | 0.09635 | 192 | 0.148757145 | 0.005643893 |

| | | | | | | | |
|---|---|---|---|---|---|---|---|
| SNP_A.K_1551709_9_9122364 | 9 | 9122364 | 8.16E-07 | 0.17448 | 192 | 0.148604107 | 0.005686099 |
| SNP_A.K_708753_4_8111008 | 4 | 8111008 | 8.21E-07 | 0.09375 | 192 | 0.148540711 | 0.005686099 |
| SNP_A.K_1487790_8_26983558 | 8 | 26983558 | 8.21E-07 | 0.03646 | 192 | 0.148529379 | 0.005686099 |
| SNP_A.K_709566_4_8273973 | 4 | 8273973 | 8.22E-07 | 0.08854 | 192 | 0.148516213 | 0.005686099 |
| SNP_A.K_1813029_11_8488078 | 11 | 8488078 | 8.30E-07 | 0.14323 | 192 | 0.148403312 | 0.005722867 |
| SNP_A.K_708594_4_8063331 | 4 | 8063331 | 8.36E-07 | 0.08854 | 192 | 0.148319068 | 0.005745686 |
| SNP_A.K_725605_4_11823655 | 4 | 11823655 | 8.50E-07 | 0.17969 | 192 | 0.148131975 | 0.005820208 |
| SNP_A.K_1552892_9_9224033 | 9 | 9224033 | 8.67E-07 | 0.21615 | 192 | 0.147902143 | 0.00591523 |
| SNP_A.K_288383_2_5018846 | 2 | 5018846 | 8.72E-07 | 0.01302 | 192 | 0.14784059 | 0.00591523 |
| SNP_A.K_1813733_11_8574694 | 11 | 8574694 | 8.74E-07 | 0.14323 | 192 | 0.147807447 | 0.00591523 |
| SNP_A.K_709290_4_8246220 | 4 | 8246220 | 8.75E-07 | 0.19531 | 192 | 0.147793696 | 0.00591523 |
| SNP_A.K_288137_2_4991246 | 2 | 4991246 | 8.81E-07 | 0.01302 | 192 | 0.147713739 | 0.005937024 |
| SNP_A.K_1552929_9_9225079 | 9 | 9225079 | 8.98E-07 | 0.21354 | 192 | 0.147496196 | 0.005995159 |
| SNP_A.K_709346_4_8250325 | 4 | 8250325 | 9.00E-07 | 0.09115 | 192 | 0.147471184 | 0.005995159 |
| SNP_A.K_288314_2_5007783 | 2 | 5007783 | 9.04E-07 | 0.01302 | 192 | 0.147423859 | 0.005995159 |
| SNP_A.K_724994_4_11719940 | 4 | 11719940 | 9.04E-07 | 0.22656 | 192 | 0.147420272 | 0.005995159 |
| SNP_A.K_708642_4_8069500 | 4 | 8069500 | 9.07E-07 | 0.20573 | 192 | 0.147386919 | 0.005995159 |
| SNP_A.K_686028_4_2522613 | 4 | 2522613 | 9.07E-07 | 0.08333 | 192 | 0.147378979 | 0.005995159 |
| SNP_A.K_288215_2_4998717 | 2 | 4998717 | 9.10E-07 | 0.01302 | 192 | 0.147341111 | 0.005995812 |
| SNP_A.K_1814804_11_8776880 | 11 | 8776880 | 9.18E-07 | 0.13281 | 192 | 0.14724836 | 0.006025088 |
| SNP_A.K_709280_4_8245645 | 4 | 8245645 | 9.40E-07 | 0.125 | 192 | 0.146974663 | 0.006150199 |
| SNP_A.K_693760_4_5059626 | 4 | 5059626 | 9.52E-07 | 0.08333 | 192 | 0.146818104 | 0.006214628 |
| SNP_A.K_693646_4_5045827 | 4 | 5045827 | 9.76E-07 | 0.04688 | 192 | 0.146538381 | 0.006347187 |
| SNP_A.K_288115_2_4988802 | 2 | 4988802 | 1.00E-06 | 0.01302 | 192 | 0.146251399 | 0.006486766 |
| SNP_A.K_288117_2_4988837 | 2 | 4988837 | 1.01E-06 | 0.01302 | 192 | 0.146176586 | 0.006508696 |
| SNP_A.K_724527_4_11613406 | 4 | 11613406 | 1.02E-06 | 0.08073 | 192 | 0.146058387 | 0.006555378 |
| SNP_A.K_728967_4_12421929 | 4 | 12421929 | 1.03E-06 | 0.22396 | 192 | 0.14595941 | 0.006591472 |
| SNP_A.K_1487792_8_26983656 | 8 | 26983656 | 1.03E-06 | 0.04427 | 192 | 0.145861174 | 0.006609523 |
| SNP_A.K_1815427_11_8851705 | 11 | 8851705 | 1.04E-06 | 0.15885 | 192 | 0.14585691 | 0.006609523 |
| SNP_A.K_288188_2_4995116 | 2 | 4995116 | 1.04E-06 | 0.01302 | 192 | 0.145812749 | 0.006614594 |
| SNP_A.K_730551_4_12699428 | 4 | 12699428 | 1.05E-06 | 0.20052 | 192 | 0.145688221 | 0.006650712 |
| SNP_A.K_731705_4_13199434 | 4 | 13199434 | 1.05E-06 | 0.01823 | 192 | 0.145679663 | 0.006650712 |
| SNP_A.K_1815418_11_8851377 | 11 | 8851377 | 1.08E-06 | 0.15365 | 192 | 0.145415472 | 0.006784282 |
| SNP_A.K_708974_4_8211787 | 4 | 8211787 | 1.11E-06 | 0.09115 | 192 | 0.1450924 | 0.006956106 |
| SNP_A.K_708548_4_8061131 | 4 | 8061131 | 1.13E-06 | 0.20052 | 192 | 0.144827025 | 0.007096778 |
| SNP_A.K_724319_4_11588460 | 4 | 11588460 | 1.15E-06 | 0.1276 | 192 | 0.14463423 | 0.007156991 |
| SNP_A.K_288179_2_4994059 | 2 | 4994059 | 1.15E-06 | 0.01302 | 192 | 0.144608548 | 0.007156991 |
| SNP_A.K_698302_4_5662955 | 4 | 5662955 | 1.15E-06 | 0.02865 | 192 | 0.144608327 | 0.007156991 |
| SNP_A.K_1028724_6_10681142 | 6 | 10681142 | 1.16E-06 | 0.02604 | 192 | 0.144584819 | 0.007156991 |
| SNP_A.K_1028188_6_10584217 | 6 | 10584217 | 1.16E-06 | 0.02344 | 192 | 0.144557595 | 0.007156991 |
| SNP_A.K_1552896_9_9224257 | 9 | 9224257 | 1.17E-06 | 0.21094 | 192 | 0.144493722 | 0.007167105 |
| SNP_A.K_709049_4_8215210 | 4 | 8215210 | 1.17E-06 | 0.09635 | 192 | 0.14447321 | 0.007167105 |
| SNP_A.K_709425_4_8265815 | 4 | 8265815 | 1.18E-06 | 0.09375 | 192 | 0.144369631 | 0.007210612 |
| SNP_A.K_288134_2_4990664 | 2 | 4990664 | 1.18E-06 | 0.01302 | 192 | 0.144321365 | 0.007219675 |
| SNP_A.K_288469_2_5032381 | 2 | 5032381 | 1.20E-06 | 0.01302 | 192 | 0.14415514 | 0.007303283 |
| SNP_A.K_729016_4_12427674 | 4 | 12427674 | 1.21E-06 | 0.25 | 192 | 0.144034787 | 0.007358548 |
| SNP_A.K_288175_2_4993980 | 2 | 4993980 | 1.24E-06 | 0.01302 | 192 | 0.143790362 | 0.007472974 |
| SNP_A.K_288174_2_4993961 | 2 | 4993961 | 1.24E-06 | 0.01302 | 192 | 0.143790362 | 0.007472974 |
| SNP_A.K_278117_2_3027954 | 2 | 3027954 | 1.25E-06 | 0.0651 | 192 | 0.14369332 | 0.007509867 |
| SNP_A.K_1552907_9_9224464 | 9 | 9224464 | 1.25E-06 | 0.21615 | 192 | 0.143667222 | 0.007509867 |

| | | | | | | | |
|---|---|---|---|---|---|---|---|
| SNP_A.K_709112_4_8226414 | 4 | 8226414 | 1.28E-06 | 0.09375 | 192 | 0.143390722 | 0.007670509 |
| SNP_A.K_726215_4_11929224 | 4 | 11929224 | 1.29E-06 | 0.22396 | 192 | 0.14333708 | 0.007680404 |
| SNP_A.K_725217_4_11763173 | 4 | 11763173 | 1.29E-06 | 0.25 | 192 | 0.143310008 | 0.007680404 |
| SNP_A.K_1813515_11_8543843 | 11 | 8543843 | 1.30E-06 | 0.14583 | 192 | 0.143207755 | 0.007698152 |
| SNP_A.K_708599_4_8063460 | 4 | 8063460 | 1.30E-06 | 0.08854 | 192 | 0.143198016 | 0.007698152 |
| SNP_A.K_710525_4_8426312 | 4 | 8426312 | 1.31E-06 | 0.09115 | 192 | 0.143158267 | 0.007698152 |
| SNP_A.K_729013_4_12427498 | 4 | 12427498 | 1.31E-06 | 0.2526 | 192 | 0.143152831 | 0.007698152 |
| SNP_A.K_708598_4_8063409 | 4 | 8063409 | 1.32E-06 | 0.09375 | 192 | 0.143053697 | 0.007703612 |
| SNP_A.K_1815448_11_8852393 | 11 | 8852393 | 1.32E-06 | 0.14583 | 192 | 0.143048936 | 0.007703612 |
| SNP_A.K_709555_4_8273679 | 4 | 8273679 | 1.33E-06 | 0.08854 | 192 | 0.142992417 | 0.007703612 |
| SNP_A.K_709294_4_8246409 | 4 | 8246409 | 1.33E-06 | 0.13802 | 192 | 0.142970423 | 0.007703612 |
| SNP_A.K_363696_2_16460925 | 2 | 16460925 | 1.33E-06 | 0.10938 | 192 | 0.142953394 | 0.007703612 |
| SNP_A.K_730516_4_12697714 | 4 | 12697714 | 1.33E-06 | 0.21094 | 192 | 0.142951501 | 0.007703612 |
| SNP_A.K_729055_4_12506050 | 4 | 12506050 | 1.35E-06 | 0.20833 | 192 | 0.142832186 | 0.007747072 |
| SNP_A.K_1552937_9_9225382 | 9 | 9225382 | 1.35E-06 | 0.22135 | 192 | 0.142766708 | 0.007747072 |
| SNP_A.K_709289_4_8245934 | 4 | 8245934 | 1.35E-06 | 0.10677 | 192 | 0.142761083 | 0.007747072 |
| SNP_A.K_288109_2_4988475 | 2 | 4988475 | 1.36E-06 | 0.01302 | 192 | 0.142730925 | 0.007747072 |
| SNP_A.K_1487782_8_26983036 | 8 | 26983036 | 1.36E-06 | 0.03906 | 192 | 0.14269692 | 0.007747072 |
| SNP_A.K_1487789_8_26983445 | 8 | 26983445 | 1.36E-06 | 0.03906 | 192 | 0.14269692 | 0.007747072 |
| SNP_A.K_288151_2_4992282 | 2 | 4992282 | 1.39E-06 | 0.01302 | 192 | 0.142473968 | 0.007877371 |
| SNP_A.K_1028760_6_10682840 | 6 | 10682840 | 1.40E-06 | 0.02604 | 192 | 0.142379868 | 0.007920591 |
| SNP_A.K_288714_2_5061588 | 2 | 5061588 | 1.41E-06 | 0.01302 | 192 | 0.142315412 | 0.00793863 |
| SNP_A.K_708595_4_8063346 | 4 | 8063346 | 1.41E-06 | 0.09375 | 192 | 0.142291496 | 0.00793863 |
| SNP_A.K_288209_2_4998353 | 2 | 4998353 | 1.42E-06 | 0.01302 | 192 | 0.142199275 | 0.00798107 |
| SNP_A.K_1698635_10_12921082 | 10 | 12921082 | 1.44E-06 | 0.17708 | 192 | 0.142079268 | 0.008043245 |
| SNP_A.K_288107_2_4988418 | 2 | 4988418 | 1.44E-06 | 0.01302 | 192 | 0.142032058 | 0.008054735 |
| SNP_A.K_1487785_8_26983139 | 8 | 26983139 | 1.46E-06 | 0.03906 | 192 | 0.1419311 | 0.008104157 |
| SNP_A.K_708832_4_8115505 | 4 | 8115505 | 1.46E-06 | 0.08854 | 192 | 0.141876071 | 0.008119178 |
| SNP_A.K_288462_2_5031896 | 2 | 5031896 | 1.47E-06 | 0.01302 | 192 | 0.141832144 | 0.008119178 |
| SNP_A.K_708596_4_8063402 | 4 | 8063402 | 1.47E-06 | 0.10156 | 192 | 0.1418144 | 0.008119178 |
| SNP_A.K_708004_4_7494976 | 4 | 7494976 | 1.47E-06 | 0.22135 | 192 | 0.141787736 | 0.008119178 |
| SNP_A.K_1028764_6_10682948 | 6 | 10682948 | 1.49E-06 | 0.02604 | 192 | 0.141632358 | 0.008204026 |
| SNP_A.K_1552269_9_9164344 | 9 | 9164344 | 1.50E-06 | 0.21615 | 192 | 0.141607871 | 0.008204026 |
| SNP_A.K_288538_2_5037464 | 2 | 5037464 | 1.52E-06 | 0.01302 | 192 | 0.141459606 | 0.008288914 |
| SNP_A.K_1552933_9_9225119 | 9 | 9225119 | 1.54E-06 | 0.21615 | 192 | 0.141311309 | 0.008374779 |
| SNP_A.K_1487784_8_26983106 | 8 | 26983106 | 1.56E-06 | 0.04427 | 192 | 0.14113221 | 0.008484367 |
| SNP_A.K_288304_2_5005747 | 2 | 5005747 | 1.59E-06 | 0.01563 | 192 | 0.140909248 | 0.008628421 |
| SNP_A.K_709267_4_8244898 | 4 | 8244898 | 1.61E-06 | 0.21094 | 192 | 0.140770432 | 0.008710847 |
| SNP_A.K_1763047_10_21616432 | 10 | 21616432 | 1.62E-06 | 0.20833 | 192 | 0.140691774 | 0.008726412 |
| SNP_A.K_253412_1_40888392 | 1 | 40888392 | 1.62E-06 | 0.13281 | 192 | 0.140690573 | 0.008726412 |
| SNP_A.K_1027131_6_10364196 | 6 | 10364196 | 1.66E-06 | 0.01302 | 192 | 0.140442345 | 0.008894491 |
| SNP_A.K_725790_4_11865823 | 4 | 11865823 | 1.67E-06 | 0.17708 | 192 | 0.140390137 | 0.008912139 |
| SNP_A.K_1552887_9_9223868 | 9 | 9223868 | 1.68E-06 | 0.22396 | 192 | 0.140313374 | 0.008949041 |
| SNP_A.K_1552910_9_9224624 | 9 | 9224624 | 1.69E-06 | 0.22135 | 192 | 0.140211462 | 0.009004423 |
| SNP_A.K_709596_4_8277494 | 4 | 8277494 | 1.70E-06 | 0.23177 | 192 | 0.140158 | 0.009004423 |
| SNP_A.K_1552879_9_9223690 | 9 | 9223690 | 1.70E-06 | 0.22396 | 192 | 0.140154903 | 0.009004423 |
| SNP_A.K_1552863_9_9222619 | 9 | 9222619 | 1.73E-06 | 0.23438 | 192 | 0.139961928 | 0.009134168 |
| SNP_A.K_730553_4_12699446 | 4 | 12699446 | 1.74E-06 | 0.20313 | 192 | 0.139859391 | 0.009192986 |
| SNP_A.K_724242_4_11583159 | 4 | 11583159 | 1.79E-06 | 0.1276 | 192 | 0.13954017 | 0.009429032 |
| SNP_A.K_1552864_9_9222626 | 9 | 9222626 | 1.82E-06 | 0.23177 | 192 | 0.139367981 | 0.009547839 |

| | | | | | | | |
|---|---|---|---|---|---|---|---|
| SNP_A.K_709288_4_8245927 | 4 | 8245927 | 1.88E-06 | 0.10417 | 192 | 0.139015125 | 0.009814931 |
| SNP_A.K_1552944_9_9225672 | 9 | 9225672 | 1.88E-06 | 0.25781 | 192 | 0.138994784 | 0.009814931 |
| SNP_A.K_709399_4_8265050 | 4 | 8265050 | 1.90E-06 | 0.09896 | 192 | 0.138864347 | 0.009902671 |
| SNP_A.K_708940_4_8150849 | 4 | 8150849 | 1.93E-06 | 0.16667 | 192 | 0.138712338 | 0.010010138 |
| SNP_A.K_724762_4_11676549 | 4 | 11676549 | 1.93E-06 | 0.23698 | 192 | 0.13867698 | 0.010016158 |
| SNP_A.K_728941_4_12410616 | 4 | 12410616 | 1.95E-06 | 0.22135 | 192 | 0.138583375 | 0.010073426 |
| SNP_A.K_1813989_11_8594000 | 11 | 8594000 | 1.98E-06 | 0.15365 | 192 | 0.13839159 | 0.010217991 |
| SNP_A.K_708597_4_8063404 | 4 | 8063404 | 1.99E-06 | 0.09635 | 192 | 0.138363867 | 0.010217991 |
| SNP_A.K_709283_4_8245671 | 4 | 8245671 | 2.01E-06 | 0.08854 | 192 | 0.138261222 | 0.010284764 |
| SNP_A.K_1815447_11_8852386 | 11 | 8852386 | 2.01E-06 | 0.14844 | 192 | 0.138213628 | 0.010302304 |
| SNP_A.K_1953659_12_3611869 | 12 | 3611869 | 2.03E-06 | 0.09375 | 192 | 0.138121982 | 0.010335602 |
| SNP_A.K_1815436_11_8851973 | 11 | 8851973 | 2.03E-06 | 0.14844 | 192 | 0.138120724 | 0.010335602 |
| SNP_A.K_1815446_11_8852379 | 11 | 8852379 | 2.04E-06 | 0.14844 | 192 | 0.138088272 | 0.010339714 |
| SNP_A.K_1815453_11_8852501 | 11 | 8852501 | 2.06E-06 | 0.14323 | 192 | 0.137970213 | 0.010421674 |
| SNP_A.K_1552095_9_9145397 | 9 | 9145397 | 2.07E-06 | 0.33333 | 192 | 0.137878495 | 0.010480171 |
| SNP_A.K_724200_4_11579766 | 4 | 11579766 | 2.09E-06 | 0.13021 | 192 | 0.13777797 | 0.010547196 |
| SNP_A.K_724463_4_11601086 | 4 | 11601086 | 2.10E-06 | 0.1875 | 192 | 0.137718009 | 0.010565996 |
| SNP_A.K_709731_4_8284154 | 4 | 8284154 | 2.11E-06 | 0.09635 | 192 | 0.137702442 | 0.010565996 |
| SNP_A.K_709626_4_8278533 | 4 | 8278533 | 2.14E-06 | 0.23177 | 192 | 0.137541927 | 0.01065285 |
| SNP_A.K_1552951_9_9225776 | 9 | 9225776 | 2.14E-06 | 0.25521 | 192 | 0.137538035 | 0.01065285 |
| SNP_A.K_134461_1_19249290 | 1 | 19249290 | 2.14E-06 | 0.02865 | 192 | 0.137499986 | 0.01065285 |
| SNP_A.K_1552928_9_9225076 | 9 | 9225076 | 2.15E-06 | 0.2474 | 192 | 0.13748724 | 0.01065285 |
| SNP_A.K_1815417_11_8851360 | 11 | 8851360 | 2.15E-06 | 0.16146 | 192 | 0.137472163 | 0.01065285 |
| SNP_A.K_730465_4_12694910 | 4 | 12694910 | 2.24E-06 | 0.20313 | 192 | 0.137013187 | 0.011063511 |
| SNP_A.K_1028783_6_10683434 | 6 | 10683434 | 2.25E-06 | 0.02865 | 192 | 0.136966865 | 0.011082183 |
| SNP_A.K_685401_4_2467389 | 4 | 2467389 | 2.27E-06 | 0.03385 | 192 | 0.136871107 | 0.011149144 |
| SNP_A.K_1028712_6_10680306 | 6 | 10680306 | 2.32E-06 | 0.02865 | 192 | 0.136606129 | 0.011384191 |
| SNP_A.K_1551787_9_9126244 | 9 | 9126244 | 2.33E-06 | 0.32031 | 192 | 0.136565231 | 0.011398196 |
| SNP_A.K_709730_4_8284150 | 4 | 8284150 | 2.34E-06 | 0.09896 | 192 | 0.136513694 | 0.011422932 |
| SNP_A.K_709238_4_8242278 | 4 | 8242278 | 2.39E-06 | 0.20573 | 192 | 0.136245769 | 0.011667073 |
| SNP_A.K_372682_2_17460818 | 2 | 17460818 | 2.41E-06 | 0.01563 | 192 | 0.136176695 | 0.011710539 |
| SNP_A.K_1815132_11_8806510 | 11 | 8806510 | 2.42E-06 | 0.13021 | 192 | 0.136116452 | 0.011745136 |
| SNP_A.K_418958_2_24508877 | 2 | 24508877 | 2.44E-06 | 0.04427 | 192 | 0.136012253 | 0.011825411 |
| SNP_A.K_334565_2_11211324 | 2 | 11211324 | 2.46E-06 | 0.01563 | 192 | 0.135950848 | 0.011854356 |
| SNP_A.K_1552952_9_9225783 | 9 | 9225783 | 2.46E-06 | 0.25781 | 192 | 0.135931488 | 0.011854356 |
| SNP_A.K_728197_4_12288860 | 4 | 12288860 | 2.52E-06 | 0.22656 | 192 | 0.135667371 | 0.012104264 |
| SNP_A.K_1552918_9_9224829 | 9 | 9224829 | 2.53E-06 | 0.22656 | 192 | 0.135615513 | 0.012131458 |
| SNP_A.K_1552899_9_9224344 | 9 | 9224344 | 2.55E-06 | 0.23698 | 192 | 0.135538722 | 0.01218542 |
| SNP_A.K_1815486_11_8853479 | 11 | 8853479 | 2.61E-06 | 0.14844 | 192 | 0.135279585 | 0.012437198 |
| SNP_A.K_710409_4_8348757 | 4 | 8348757 | 2.64E-06 | 0.10417 | 192 | 0.135109218 | 0.012595758 |
| SNP_A.K_710510_4_8425607 | 4 | 8425607 | 2.66E-06 | 0.10156 | 192 | 0.135044544 | 0.012638579 |
| SNP_A.K_1815620_11_8857077 | 11 | 8857077 | 2.71E-06 | 0.14844 | 192 | 0.134838637 | 0.012828626 |
| SNP_A.K_1815438_11_8852015 | 11 | 8852015 | 2.71E-06 | 0.14844 | 192 | 0.134822749 | 0.012828626 |
| SNP_A.K_727016_4_12069358 | 4 | 12069358 | 2.73E-06 | 0.05729 | 192 | 0.134760821 | 0.012855739 |
| SNP_A.K_730529_4_12698210 | 4 | 12698210 | 2.73E-06 | 0.21615 | 192 | 0.134747092 | 0.012855739 |
| SNP_A.K_1393181_8_10466636 | 8 | 10466636 | 2.77E-06 | 0.0625 | 192 | 0.13457387 | 0.01301953 |
| SNP_A.K_1552948_9_9225759 | 9 | 9225759 | 2.78E-06 | 0.22917 | 192 | 0.134551618 | 0.01301953 |
| SNP_A.K_724567_4_11634757 | 4 | 11634757 | 2.81E-06 | 0.18229 | 192 | 0.134433204 | 0.013126157 |
| SNP_A.K_1815334_11_8836292 | 11 | 8836292 | 2.81E-06 | 0.17708 | 192 | 0.134397574 | 0.013137803 |
| SNP_A.K_687579_4_3565945 | 4 | 3565945 | 2.84E-06 | 0.13281 | 192 | 0.134294109 | 0.013228163 |

| | | | | | | | |
|---|---|---|---|---|---|---|---|
| SNP_A.K_725306_4_11791599 | 4 | 11791599 | 2.85E-06 | 0.23958 | 192 | 0.134238568 | 0.013263233 |
| SNP_A.K_706654_4_6930631 | 4 | 6930631 | 2.89E-06 | 0.18229 | 192 | 0.134109953 | 0.013384174 |
| SNP_A.K_1551786_9_9126240 | 9 | 9126240 | 2.96E-06 | 0.29427 | 192 | 0.13382338 | 0.013695235 |
| SNP_A.K_1814803_11_8776861 | 11 | 8776861 | 2.97E-06 | 0.14583 | 192 | 0.133784933 | 0.013711155 |
| SNP_A.K_1813046_11_8488787 | 11 | 8488787 | 2.99E-06 | 0.15104 | 192 | 0.133706586 | 0.013775424 |
| SNP_A.K_727022_4_12069883 | 4 | 12069883 | 3.04E-06 | 0.05729 | 192 | 0.133535148 | 0.013942371 |
| SNP_A.K_1196918_7_7667699 | 7 | 7667699 | 3.05E-06 | 0.02083 | 192 | 0.133472885 | 0.013942371 |
| SNP_A.K_1815585_11_8855936 | 11 | 8855936 | 3.06E-06 | 0.14583 | 192 | 0.133441939 | 0.013942371 |
| SNP_A.K_1815564_11_8855479 | 11 | 8855479 | 3.06E-06 | 0.14583 | 192 | 0.133438907 | 0.013942371 |
| SNP_A.K_708060_4_7522296 | 4 | 7522296 | 3.07E-06 | 0.20833 | 192 | 0.133420485 | 0.013942371 |
| SNP_A.K_1813301_11_8528262 | 11 | 8528262 | 3.07E-06 | 0.15104 | 192 | 0.133419768 | 0.013942371 |
| SNP_A.K_708749_4_8110906 | 4 | 8110906 | 3.08E-06 | 0.09896 | 192 | 0.133377149 | 0.013964267 |
| SNP_A.K_1552897_9_9224260 | 9 | 9224260 | 3.11E-06 | 0.23177 | 192 | 0.133251552 | 0.014087697 |
| SNP_A.K_1552909_9_9224603 | 9 | 9224603 | 3.12E-06 | 0.22917 | 192 | 0.133222868 | 0.014087697 |
| SNP_A.K_724630_4_11648139 | 4 | 11648139 | 3.13E-06 | 0.20313 | 192 | 0.133203129 | 0.014087697 |
| SNP_A.K_624824_3_27997787 | 3 | 27997787 | 3.18E-06 | 0.25781 | 192 | 0.133005874 | 0.014299039 |
| SNP_A.K_1487783_8_26983084 | 8 | 26983084 | 3.20E-06 | 0.04427 | 192 | 0.13295255 | 0.014299039 |
| SNP_A.K_1551961_9_9135127 | 9 | 9135127 | 3.20E-06 | 0.3151 | 192 | 0.132946617 | 0.014299039 |
| SNP_A.K_624512_3_27966375 | 3 | 27966375 | 3.20E-06 | 0.1901 | 192 | 0.132936044 | 0.014299039 |
| SNP_A.K_709467_4_8267957 | 4 | 8267957 | 3.23E-06 | 0.10417 | 192 | 0.132845443 | 0.014340613 |
| SNP_A.K_725561_4_11821553 | 4 | 11821553 | 3.24E-06 | 0.17969 | 192 | 0.132798703 | 0.014340613 |
| SNP_A.K_1813732_11_8574690 | 11 | 8574690 | 3.24E-06 | 0.16406 | 192 | 0.132791126 | 0.014340613 |
| SNP_A.K_1488914_8_27113919 | 8 | 27113919 | 3.24E-06 | 0.11198 | 192 | 0.132786873 | 0.014340613 |
| SNP_A.K_708774_4_8112498 | 4 | 8112498 | 3.24E-06 | 0.10156 | 192 | 0.132781865 | 0.014340613 |
| SNP_A.K_1551968_9_9135579 | 9 | 9135579 | 3.30E-06 | 0.2526 | 192 | 0.132576641 | 0.014485246 |
| SNP_A.K_1211151_7_9887065 | 7 | 9887065 | 3.31E-06 | 0.07031 | 192 | 0.132554234 | 0.014485246 |
| SNP_A.K_1813027_11_8487897 | 11 | 8487897 | 3.32E-06 | 0.15365 | 192 | 0.132514248 | 0.014485246 |
| SNP_A.K_730524_4_12698045 | 4 | 12698045 | 3.32E-06 | 0.25521 | 192 | 0.132505605 | 0.014485246 |
| SNP_A.K_1815431_11_8851771 | 11 | 8851771 | 3.33E-06 | 0.14063 | 192 | 0.132493875 | 0.014485246 |
| SNP_A.K_1814805_11_8776897 | 11 | 8776897 | 3.34E-06 | 0.13542 | 192 | 0.132470066 | 0.014485246 |
| SNP_A.K_1487878_8_26988976 | 8 | 26988976 | 3.34E-06 | 0.03906 | 192 | 0.132453027 | 0.014485246 |
| SNP_A.K_1487879_8_26988980 | 8 | 26988980 | 3.34E-06 | 0.03906 | 192 | 0.132453027 | 0.014485246 |
| SNP_A.K_1487899_8_26989186 | 8 | 26989186 | 3.34E-06 | 0.03906 | 192 | 0.132453027 | 0.014485246 |
| SNP_A.K_1813140_11_8518282 | 11 | 8518282 | 3.36E-06 | 0.15885 | 192 | 0.132395346 | 0.014528774 |
| SNP_A.K_1813261_11_8526065 | 11 | 8526065 | 3.38E-06 | 0.15365 | 192 | 0.132324333 | 0.014560163 |
| SNP_A.K_1815485_11_8853476 | 11 | 8853476 | 3.38E-06 | 0.14844 | 192 | 0.132323754 | 0.014560163 |
| SNP_A.K_730528_4_12698185 | 4 | 12698185 | 3.39E-06 | 0.21094 | 192 | 0.132274255 | 0.01459359 |
| SNP_A.K_708797_4_8114726 | 4 | 8114726 | 3.41E-06 | 0.07292 | 192 | 0.132210685 | 0.014645308 |
| SNP_A.K_709474_4_8268290 | 4 | 8268290 | 3.44E-06 | 0.09896 | 192 | 0.132112502 | 0.014742179 |
| SNP_A.K_1815555_11_8855282 | 11 | 8855282 | 3.46E-06 | 0.14844 | 192 | 0.132039207 | 0.01480725 |
| SNP_A.K_1815566_11_8855519 | 11 | 8855519 | 3.48E-06 | 0.14844 | 192 | 0.131978517 | 0.014856166 |
| SNP_A.K_1552891_9_9223996 | 9 | 9223996 | 3.55E-06 | 0.22917 | 192 | 0.131756208 | 0.015119207 |
| SNP_A.K_709599_4_8277661 | 4 | 8277661 | 3.59E-06 | 0.09896 | 192 | 0.131633657 | 0.015222422 |
| SNP_A.K_1551806_9_9126815 | 9 | 9126815 | 3.63E-06 | 0.36979 | 192 | 0.131498018 | 0.015222422 |
| SNP_A.K_1551705_9_9122135 | 9 | 9122135 | 3.64E-06 | 0.26302 | 192 | 0.131491177 | 0.015222422 |
| SNP_A.K_707033_4_6997430 | 4 | 6997430 | 3.64E-06 | 0.21094 | 192 | 0.131482528 | 0.015222422 |
| SNP_A.K_1551941_9_9134015 | 9 | 9134015 | 3.65E-06 | 0.26042 | 192 | 0.131455691 | 0.015222422 |
| SNP_A.K_708945_4_8151290 | 4 | 8151290 | 3.66E-06 | 0.1901 | 192 | 0.131425456 | 0.015222422 |
| SNP_A.K_730786_4_12954074 | 4 | 12954074 | 3.66E-06 | 0.19792 | 192 | 0.131418544 | 0.015222422 |
| SNP_A.K_709308_4_8248006 | 4 | 8248006 | 3.66E-06 | 0.11198 | 192 | 0.131417328 | 0.015222422 |

| | | | | | | | |
|---|---|---|---|---|---|---|---|
| SNP_A.K_1810489_11_8088821 | 11 | 8088821 | 3.67E-06 | 0.15625 | 192 | 0.131377114 | 0.015222422 |
| SNP_A.K_288218_2_4998843 | 2 | 4998843 | 3.74E-06 | 0.00521 | 192 | 0.131157961 | 0.015222422 |
| SNP_A.K_288228_2_4999616 | 2 | 4999616 | 3.74E-06 | 0.00521 | 192 | 0.131157961 | 0.015222422 |
| SNP_A.K_288229_2_4999648 | 2 | 4999648 | 3.74E-06 | 0.00521 | 192 | 0.131157961 | 0.015222422 |
| SNP_A.K_315911_2_8310508 | 2 | 8310508 | 3.74E-06 | 0.00521 | 192 | 0.131157961 | 0.015222422 |
| SNP_A.K_817407_4_31495768 | 4 | 31495768 | 3.74E-06 | 0.00521 | 192 | 0.131157961 | 0.015222422 |
| SNP_A.K_885802_5_8945161 | 5 | 8945161 | 3.74E-06 | 0.00521 | 192 | 0.131157961 | 0.015222422 |
| SNP_A.K_886511_5_13003572 | 5 | 13003572 | 3.74E-06 | 0.00521 | 192 | 0.131157961 | 0.015222422 |
| SNP_A.K_1371552_8_7852191 | 8 | 7852191 | 3.74E-06 | 0.00521 | 192 | 0.131157961 | 0.015222422 |
| SNP_A.K_363616_2_16455997 | 2 | 16455997 | 3.74E-06 | 0.00521 | 192 | 0.131157961 | 0.015222422 |
| SNP_A.K_528009_3_11220328 | 3 | 11220328 | 3.74E-06 | 0.00521 | 192 | 0.131157961 | 0.015222422 |
| SNP_A.K_1154455_6_29052368 | 6 | 29052368 | 3.74E-06 | 0.00521 | 192 | 0.131157961 | 0.015222422 |
| SNP_A.K_1371604_8_7858316 | 8 | 7858316 | 3.74E-06 | 0.00521 | 192 | 0.131157961 | 0.015222422 |
| SNP_A.K_1605756_9_19946683 | 9 | 19946683 | 3.74E-06 | 0.00521 | 192 | 0.131157961 | 0.015222422 |
| SNP_A.K_1752246_10_19901886 | 10 | 19901886 | 3.74E-06 | 0.00521 | 192 | 0.131157961 | 0.015222422 |
| SNP_A.K_725216_4_11763133 | 4 | 11763133 | 3.76E-06 | 0.14323 | 192 | 0.131106777 | 0.015261518 |
| SNP_A.K_1552881_9_9223720 | 9 | 9223720 | 3.80E-06 | 0.24479 | 192 | 0.130979824 | 0.015403426 |
| SNP_A.K_253133_1_40864747 | 1 | 40864747 | 3.82E-06 | 0.14323 | 192 | 0.130933486 | 0.015436515 |
| SNP_A.K_708922_4_8125665 | 4 | 8125665 | 3.87E-06 | 0.0651 | 192 | 0.1307915 | 0.015600894 |
| SNP_A.K_685874_4_2518130 | 4 | 2518130 | 3.90E-06 | 0.04948 | 192 | 0.13071047 | 0.015682494 |
| SNP_A.K_424818_2_25205727 | 2 | 25205727 | 3.90E-06 | 0.01563 | 192 | 0.130683019 | 0.015690196 |
| SNP_A.K_1815552_11_8855237 | 11 | 8855237 | 3.92E-06 | 0.14583 | 192 | 0.130629981 | 0.01570767 |
| SNP_A.K_1815567_11_8855642 | 11 | 8855642 | 3.92E-06 | 0.14583 | 192 | 0.130625569 | 0.01570767 |
| SNP_A.K_886241_5_11405244 | 5 | 11405244 | 3.93E-06 | 0.00521 | 192 | 0.130605003 | 0.01570767 |
| SNP_A.K_709446_4_8266337 | 4 | 8266337 | 3.96E-06 | 0.10156 | 192 | 0.130519626 | 0.015796211 |
| SNP_A.K_1815459_11_8852717 | 11 | 8852717 | 3.98E-06 | 0.15104 | 192 | 0.130479432 | 0.015822064 |
| SNP_A.K_440181_2_28272190 | 2 | 28272190 | 3.98E-06 | 0.01563 | 192 | 0.130454916 | 0.015826094 |
| SNP_A.K_1551855_9_9129525 | 9 | 9129525 | 4.05E-06 | 0.25521 | 192 | 0.13027985 | 0.015961858 |
| SNP_A.K_1815461_11_8852743 | 11 | 8852743 | 4.05E-06 | 0.15104 | 192 | 0.13026603 | 0.015961858 |
| SNP_A.K_730519_4_12697846 | 4 | 12697846 | 4.06E-06 | 0.22396 | 192 | 0.130253129 | 0.015961858 |
| SNP_A.K_374611_2_17863769 | 2 | 17863769 | 4.06E-06 | 0.10938 | 192 | 0.130251637 | 0.015961858 |
| SNP_A.K_806606_4_29505446 | 4 | 29505446 | 4.08E-06 | 0.00521 | 192 | 0.130191342 | 0.015961858 |
| SNP_A.K_526222_3_10849783 | 3 | 10849783 | 4.08E-06 | 0.00521 | 192 | 0.130191342 | 0.015961858 |
| SNP_A.K_863296_5_3678971 | 5 | 3678971 | 4.08E-06 | 0.00521 | 192 | 0.130191342 | 0.015961858 |
| SNP_A.K_1488988_8_27119845 | 8 | 27119845 | 4.08E-06 | 0.11719 | 192 | 0.130186651 | 0.015961858 |
| SNP_A.K_1551738_9_9124100 | 9 | 9124100 | 4.11E-06 | 0.35677 | 192 | 0.13011058 | 0.016039336 |
| SNP_A.K_1551684_9_9120567 | 9 | 9120567 | 4.13E-06 | 0.15104 | 192 | 0.1300463 | 0.016063366 |
| SNP_A.K_1552930_9_9225086 | 9 | 9225086 | 4.13E-06 | 0.2474 | 192 | 0.130035913 | 0.016063366 |
| SNP_A.K_708975_4_8211883 | 4 | 8211883 | 4.14E-06 | 0.10156 | 192 | 0.130030106 | 0.016063366 |
| SNP_A.K_708681_4_8085071 | 4 | 8085071 | 4.15E-06 | 0.21354 | 192 | 0.129993551 | 0.016085321 |
| SNP_A.K_710506_4_8425311 | 4 | 8425311 | 4.20E-06 | 0.07552 | 192 | 0.129855942 | 0.016251846 |
| SNP_A.K_1552954_9_9225976 | 9 | 9225976 | 4.22E-06 | 0.22656 | 192 | 0.129810542 | 0.016286908 |
| SNP_A.K_709762_4_8285211 | 4 | 8285211 | 4.23E-06 | 0.08073 | 192 | 0.129775421 | 0.016307286 |
| SNP_A.K_1552430_9_9176508 | 9 | 9176508 | 4.29E-06 | 0.24479 | 192 | 0.129625419 | 0.016494391 |
| SNP_A.K_709246_4_8242605 | 4 | 8242605 | 4.29E-06 | 0.09635 | 192 | 0.129604385 | 0.016494599 |
| SNP_A.K_1552959_9_9226408 | 9 | 9226408 | 4.30E-06 | 0.1901 | 192 | 0.129582844 | 0.016495604 |
| SNP_A.K_1552889_9_9223960 | 9 | 9223960 | 4.32E-06 | 0.22917 | 192 | 0.129537119 | 0.016531974 |
| SNP_A.K_709132_4_8232773 | 4 | 8232773 | 4.34E-06 | 0.0625 | 192 | 0.129478321 | 0.016560586 |
| SNP_A.K_1552936_9_9225269 | 9 | 9225269 | 4.34E-06 | 0.26302 | 192 | 0.129476051 | 0.016560586 |
| SNP_A.K_1552532_9_9192773 | 9 | 9192773 | 4.38E-06 | 0.15365 | 192 | 0.129381404 | 0.016669224 |

| | | | | | | | |
|---|---|---|---|---|---|---|---|
| SNP_A.K_1551683_9_9120529 | 9 | 9120529 | 4.42E-06 | 0.15625 | 192 | 0.129287872 | 0.016776994 |
| SNP_A.K_1815551_11_8855223 | 11 | 8855223 | 4.44E-06 | 0.14844 | 192 | 0.129238822 | 0.016813374 |
| SNP_A.K_1815601_11_8856277 | 11 | 8856277 | 4.46E-06 | 0.14844 | 192 | 0.129188215 | 0.016813374 |
| SNP_A.K_1815580_11_8855793 | 11 | 8855793 | 4.46E-06 | 0.14844 | 192 | 0.12918257 | 0.016813374 |
| SNP_A.K_1815450_11_8852407 | 11 | 8852407 | 4.46E-06 | 0.15365 | 192 | 0.129166792 | 0.016813374 |
| SNP_A.K_1552886_9_9223863 | 9 | 9223863 | 4.47E-06 | 0.23438 | 192 | 0.129160681 | 0.016813374 |
| SNP_A.K_1552634_9_9203413 | 9 | 9203413 | 4.48E-06 | 0.16146 | 192 | 0.129134208 | 0.016816606 |
| SNP_A.K_1813624_11_8561348 | 11 | 8561348 | 4.48E-06 | 0.17188 | 192 | 0.129117693 | 0.016816606 |
| SNP_A.K_708889_4_8121990 | 4 | 8121990 | 4.62E-06 | 0.20833 | 192 | 0.128774448 | 0.017290212 |
| SNP_A.K_1551724_9_9123486 | 9 | 9123486 | 4.63E-06 | 0.26302 | 192 | 0.128753786 | 0.017290212 |
| SNP_A.K_1487908_8_26989399 | 8 | 26989399 | 4.64E-06 | 0.04167 | 192 | 0.128742832 | 0.017290212 |
| SNP_A.K_1487914_8_26989490 | 8 | 26989490 | 4.67E-06 | 0.04167 | 192 | 0.128654233 | 0.017395139 |
| SNP_A.K_708515_4_8059492 | 4 | 8059492 | 4.73E-06 | 0.10417 | 192 | 0.128524018 | 0.017565359 |
| SNP_A.K_1543942_9_7179036 | 9 | 7179036 | 4.77E-06 | 0.03906 | 192 | 0.128409963 | 0.017711975 |
| SNP_A.K_288315_2_5007791 | 2 | 5007791 | 4.80E-06 | 0.01563 | 192 | 0.128351951 | 0.017771498 |
| SNP_A.K_708381_4_8041630 | 4 | 8041630 | 4.82E-06 | 0.07031 | 192 | 0.128310709 | 0.017804826 |
| SNP_A.K_1815441_11_8852154 | 11 | 8852154 | 4.84E-06 | 0.15104 | 192 | 0.128253582 | 0.017863386 |
| SNP_A.K_1813478_11_8540416 | 11 | 8540416 | 4.91E-06 | 0.15625 | 192 | 0.128088719 | 0.018025144 |
| SNP_A.K_1815617_11_8856875 | 11 | 8856875 | 4.92E-06 | 0.16146 | 192 | 0.128065982 | 0.018025144 |
| SNP_A.K_636157_3_29648616 | 3 | 29648616 | 4.93E-06 | 0.04427 | 192 | 0.12805086 | 0.018025144 |
| SNP_A.K_1813259_11_8525961 | 11 | 8525961 | 4.93E-06 | 0.15104 | 192 | 0.128041179 | 0.018025144 |
| SNP_A.K_709028_4_8213733 | 4 | 8213733 | 4.93E-06 | 0.21094 | 192 | 0.128037689 | 0.018025144 |
| SNP_A.K_1782845_11_3505689 | 11 | 3505689 | 4.94E-06 | 0.03125 | 192 | 0.128024632 | 0.018025144 |
| SNP_A.K_1815458_11_8852702 | 11 | 8852702 | 4.96E-06 | 0.15104 | 192 | 0.127981676 | 0.018025144 |
| SNP_A.K_1552924_9_9224936 | 9 | 9224936 | 4.97E-06 | 0.25781 | 192 | 0.127946102 | 0.018025144 |
| SNP_A.K_724597_4_11643216 | 4 | 11643216 | 4.98E-06 | 0.1875 | 192 | 0.127936882 | 0.018025144 |
| SNP_A.K_613471_3_26710142 | 3 | 26710142 | 4.99E-06 | 0.15885 | 192 | 0.127914371 | 0.018025144 |
| SNP_A.K_730797_4_12954926 | 4 | 12954926 | 5.00E-06 | 0.20052 | 192 | 0.127892179 | 0.018025144 |
| SNP_A.K_1815470_11_8853087 | 11 | 8853087 | 5.02E-06 | 0.15104 | 192 | 0.12785046 | 0.018025144 |
| SNP_A.K_1815579_11_8855791 | 11 | 8855791 | 5.02E-06 | 0.15104 | 192 | 0.12784712 | 0.018025144 |
| SNP_A.K_1815537_11_8854907 | 11 | 8854907 | 5.03E-06 | 0.15104 | 192 | 0.127825275 | 0.018025144 |
| SNP_A.K_1815531_11_8854811 | 11 | 8854811 | 5.03E-06 | 0.15104 | 192 | 0.127825275 | 0.018025144 |
| SNP_A.K_1815536_11_8854878 | 11 | 8854878 | 5.03E-06 | 0.15104 | 192 | 0.127825275 | 0.018025144 |
| SNP_A.K_1815716_11_8883590 | 11 | 8883590 | 5.03E-06 | 0.15104 | 192 | 0.12781651 | 0.018025144 |
| SNP_A.K_1815444_11_8852306 | 11 | 8852306 | 5.05E-06 | 0.15104 | 192 | 0.127782454 | 0.018048547 |
| SNP_A.K_1174425_7_1714065 | 7 | 1714065 | 5.12E-06 | 0.0625 | 192 | 0.127618443 | 0.018281336 |
| SNP_A.K_709761_4_8285205 | 4 | 8285205 | 5.15E-06 | 0.08594 | 192 | 0.127550857 | 0.018345375 |
| SNP_A.K_727458_4_12136897 | 4 | 12136897 | 5.16E-06 | 0.04427 | 192 | 0.127540339 | 0.018345375 |
| SNP_A.K_709525_4_8271593 | 4 | 8271593 | 5.19E-06 | 0.07292 | 192 | 0.127466293 | 0.018434649 |
| SNP_A.K_724644_4_11650814 | 4 | 11650814 | 5.24E-06 | 0.20313 | 192 | 0.127359482 | 0.018578302 |
| SNP_A.K_708767_4_8111399 | 4 | 8111399 | 5.25E-06 | 0.10417 | 192 | 0.127333742 | 0.018589073 |
| SNP_A.K_709527_4_8271604 | 4 | 8271604 | 5.26E-06 | 0.07552 | 192 | 0.127309944 | 0.018596704 |
| SNP_A.K_1552030_9_9139882 | 9 | 9139882 | 5.31E-06 | 0.25781 | 192 | 0.127198256 | 0.018728799 |
| SNP_A.K_730549_4_12699272 | 4 | 12699272 | 5.32E-06 | 0.21354 | 192 | 0.127191869 | 0.018728799 |
| SNP_A.K_1777345_11_2489144 | 11 | 2489144 | 5.39E-06 | 0.13281 | 192 | 0.127049824 | 0.018908068 |
| SNP_A.K_1132994_6_25647636 | 6 | 25647636 | 5.39E-06 | 0.01823 | 192 | 0.127046331 | 0.018908068 |
| SNP_A.K_730504_4_12697046 | 4 | 12697046 | 5.42E-06 | 0.21875 | 192 | 0.126982882 | 0.018946659 |
| SNP_A.K_727008_4_12068565 | 4 | 12068565 | 5.42E-06 | 0.0599 | 192 | 0.126973231 | 0.018946659 |
| SNP_A.K_1487905_8_26989374 | 8 | 26989374 | 5.42E-06 | 0.04167 | 192 | 0.126966437 | 0.018946659 |
| SNP_A.K_1028669_6_10676525 | 6 | 10676525 | 5.44E-06 | 0.03125 | 192 | 0.126925759 | 0.018983284 |

| | | | | | | | |
|---|---|---|---|---|---|---|---|
| SNP_A.K_1812688_11_8462724 | 11 | 8462724 | 5.47E-06 | 0.16667 | 192 | 0.126876906 | 0.019033832 |
| SNP_A.K_726156_4_11923161 | 4 | 11923161 | 5.48E-06 | 0.03385 | 192 | 0.126843057 | 0.019059192 |
| SNP_A.K_685240_4_2447337 | 4 | 2447337 | 5.50E-06 | 0.01823 | 192 | 0.126805183 | 0.019091456 |
| SNP_A.K_1815525_11_8854768 | 11 | 8854768 | 5.59E-06 | 0.15365 | 192 | 0.126632903 | 0.019348516 |
| SNP_A.K_388507_2_19730767 | 2 | 19730767 | 5.61E-06 | 0.07031 | 192 | 0.126597012 | 0.019348516 |
| SNP_A.K_1279730_7_20912600 | 7 | 20912600 | 5.62E-06 | 0.20313 | 192 | 0.126565943 | 0.019348516 |
| SNP_A.K_1815578_11_8855779 | 11 | 8855779 | 5.63E-06 | 0.15365 | 192 | 0.126547327 | 0.019348516 |
| SNP_A.K_1028408_6_10600462 | 6 | 10600462 | 5.64E-06 | 0.0599 | 192 | 0.126535234 | 0.019348516 |
| SNP_A.K_1814893_11_8784439 | 11 | 8784439 | 5.64E-06 | 0.15885 | 192 | 0.126526112 | 0.019348516 |
| SNP_A.K_708735_4_8089934 | 4 | 8089934 | 5.65E-06 | 0.10156 | 192 | 0.126514177 | 0.019348516 |
| SNP_A.K_1815499_11_8854115 | 11 | 8854115 | 5.65E-06 | 0.15365 | 192 | 0.126505137 | 0.019348516 |
| SNP_A.K_1813006_11_8486470 | 11 | 8486470 | 5.68E-06 | 0.1276 | 192 | 0.126439488 | 0.019372031 |
| SNP_A.K_1815514_11_8854504 | 11 | 8854504 | 5.70E-06 | 0.15365 | 192 | 0.126417557 | 0.019372031 |
| SNP_A.K_1815498_11_8854102 | 11 | 8854102 | 5.70E-06 | 0.15365 | 192 | 0.126417557 | 0.019372031 |
| SNP_A.K_1815515_11_8854508 | 11 | 8854508 | 5.70E-06 | 0.15365 | 192 | 0.126417557 | 0.019372031 |
| SNP_A.K_623825_3_27887046 | 3 | 27887046 | 5.74E-06 | 0.1875 | 192 | 0.126324598 | 0.019500651 |
| SNP_A.K_1552912_9_9224710 | 9 | 9224710 | 5.75E-06 | 0.26823 | 192 | 0.12630159 | 0.019508689 |
| SNP_A.K_1813020_11_8487380 | 11 | 8487380 | 5.77E-06 | 0.11719 | 192 | 0.126265223 | 0.01953994 |
| SNP_A.K_1812993_11_8485723 | 11 | 8485723 | 5.82E-06 | 0.16406 | 192 | 0.126170154 | 0.019673547 |
| SNP_A.K_288264_2_5001299 | 2 | 5001299 | 5.88E-06 | 0.01563 | 192 | 0.126068251 | 0.019820162 |
| SNP_A.K_288590_2_5045817 | 2 | 5045817 | 5.98E-06 | 0.01042 | 192 | 0.125878042 | 0.020125143 |
| SNP_A.K_1815481_11_8853309 | 11 | 8853309 | 6.02E-06 | 0.15625 | 192 | 0.125790238 | 0.020201933 |
| SNP_A.K_288233_2_5000218 | 2 | 5000218 | 6.02E-06 | 0.01563 | 192 | 0.12578839 | 0.020201933 |
| SNP_A.K_1815440_11_8852122 | 11 | 8852122 | 6.03E-06 | 0.16146 | 192 | 0.125780632 | 0.020201933 |
| SNP_A.K_710019_4_8308488 | 4 | 8308488 | 6.12E-06 | 0.08073 | 192 | 0.125606634 | 0.020483604 |
| SNP_A.K_727011_4_12068704 | 4 | 12068704 | 6.21E-06 | 0.0599 | 192 | 0.125449926 | 0.020501251 |
| SNP_A.K_685900_4_2519168 | 4 | 2519168 | 6.22E-06 | 0.07813 | 192 | 0.125422284 | 0.020501251 |
| SNP_A.K_1813135_11_8517675 | 11 | 8517675 | 6.23E-06 | 0.15625 | 192 | 0.125400171 | 0.020501251 |
| SNP_A.K_1815571_11_8855679 | 11 | 8855679 | 6.24E-06 | 0.15625 | 192 | 0.125397065 | 0.020501251 |
| SNP_A.K_729053_4_12505814 | 4 | 12505814 | 6.25E-06 | 0.21615 | 192 | 0.125364499 | 0.020501251 |
| SNP_A.K_1814806_11_8776900 | 11 | 8776900 | 6.31E-06 | 0.14063 | 192 | 0.125266488 | 0.020501251 |
| SNP_A.K_709390_4_8264164 | 4 | 8264164 | 6.31E-06 | 0.06771 | 192 | 0.125265042 | 0.020501251 |
| SNP_A.K_1552856_9_9222461 | 9 | 9222461 | 6.34E-06 | 0.35938 | 192 | 0.125203118 | 0.020501251 |
| SNP_A.K_1815465_11_8852885 | 11 | 8852885 | 6.36E-06 | 0.15104 | 192 | 0.125176182 | 0.020501251 |
| SNP_A.K_1810466_11_8086737 | 11 | 8086737 | 6.36E-06 | 0.15104 | 192 | 0.125166877 | 0.020501251 |
| SNP_A.K_1815509_11_8854358 | 11 | 8854358 | 6.38E-06 | 0.15104 | 192 | 0.125134819 | 0.020501251 |
| SNP_A.K_1815563_11_8855398 | 11 | 8855398 | 6.38E-06 | 0.15104 | 192 | 0.125134819 | 0.020501251 |
| SNP_A.K_1815503_11_8854182 | 11 | 8854182 | 6.39E-06 | 0.15104 | 192 | 0.125127242 | 0.020501251 |
| SNP_A.K_1815504_11_8854202 | 11 | 8854202 | 6.39E-06 | 0.15104 | 192 | 0.125127242 | 0.020501251 |
| SNP_A.K_1815505_11_8854228 | 11 | 8854228 | 6.39E-06 | 0.15104 | 192 | 0.125127242 | 0.020501251 |
| SNP_A.K_726145_4_11920509 | 4 | 11920509 | 6.40E-06 | 0.06771 | 192 | 0.125108482 | 0.020501251 |
| SNP_A.K_1812692_11_8462782 | 11 | 8462782 | 6.43E-06 | 0.16406 | 192 | 0.125057686 | 0.020501251 |
| SNP_A.K_708572_4_8061987 | 4 | 8061987 | 6.44E-06 | 0.06771 | 192 | 0.125032424 | 0.020501251 |
| SNP_A.K_708696_4_8086923 | 4 | 8086923 | 6.44E-06 | 0.06771 | 192 | 0.125032424 | 0.020501251 |
| SNP_A.K_708983_4_8212173 | 4 | 8212173 | 6.44E-06 | 0.06771 | 192 | 0.125032424 | 0.020501251 |
| SNP_A.K_709363_4_8258876 | 4 | 8258876 | 6.44E-06 | 0.06771 | 192 | 0.125032424 | 0.020501251 |
| SNP_A.K_709372_4_8259282 | 4 | 8259282 | 6.44E-06 | 0.06771 | 192 | 0.125032424 | 0.020501251 |
| SNP_A.K_709383_4_8263151 | 4 | 8263151 | 6.44E-06 | 0.06771 | 192 | 0.125032424 | 0.020501251 |
| SNP_A.K_709580_4_8274751 | 4 | 8274751 | 6.44E-06 | 0.06771 | 192 | 0.125032424 | 0.020501251 |
| SNP_A.K_709681_4_8280871 | 4 | 8280871 | 6.44E-06 | 0.06771 | 192 | 0.125032424 | 0.020501251 |

| | | | | | | | |
|---|---|---|---|---|---|---|---|
| SNP_A.K_708650_4_8070918 | 4 | 8070918 | 6.44E-06 | 0.06771 | 192 | 0.125032424 | 0.020501251 |
| SNP_A.K_708697_4_8086929 | 4 | 8086929 | 6.44E-06 | 0.06771 | 192 | 0.125032424 | 0.020501251 |
| SNP_A.K_708724_4_8089128 | 4 | 8089128 | 6.44E-06 | 0.06771 | 192 | 0.125032424 | 0.020501251 |
| SNP_A.K_709532_4_8271878 | 4 | 8271878 | 6.44E-06 | 0.06771 | 192 | 0.125032424 | 0.020501251 |
| SNP_A.K_709544_4_8272941 | 4 | 8272941 | 6.44E-06 | 0.06771 | 192 | 0.125032424 | 0.020501251 |
| SNP_A.K_709545_4_8272956 | 4 | 8272956 | 6.44E-06 | 0.06771 | 192 | 0.125032424 | 0.020501251 |
| SNP_A.K_709581_4_8274764 | 4 | 8274764 | 6.44E-06 | 0.06771 | 192 | 0.125032424 | 0.020501251 |
| SNP_A.K_708896_4_8122472 | 4 | 8122472 | 6.45E-06 | 0.06771 | 192 | 0.125012331 | 0.020502961 |
| SNP_A.K_253411_1_40888350 | 1 | 40888350 | 6.46E-06 | 0.14063 | 192 | 0.12499298 | 0.020502961 |
| SNP_A.K_1814372_11_8622044 | 11 | 8622044 | 6.47E-06 | 0.15885 | 192 | 0.124980005 | 0.020502961 |
| SNP_A.K_1552007_9_9138358 | 9 | 9138358 | 6.54E-06 | 0.26823 | 192 | 0.124862416 | 0.020686956 |
| SNP_A.K_709123_4_8230845 | 4 | 8230845 | 6.57E-06 | 0.06771 | 192 | 0.12481355 | 0.020714195 |
| SNP_A.K_1681526_10_10209690 | 10 | 10209690 | 6.57E-06 | 0.02083 | 192 | 0.124813488 | 0.020714195 |
| SNP_A.K_1552872_9_9223007 | 9 | 9223007 | 6.62E-06 | 0.3099 | 192 | 0.12472466 | 0.020821187 |
| SNP_A.K_1813689_11_8566185 | 11 | 8566185 | 6.63E-06 | 0.17708 | 192 | 0.12471192 | 0.020821187 |
| SNP_A.K_613662_3_26726882 | 3 | 26726882 | 6.63E-06 | 0.16146 | 192 | 0.124704558 | 0.020821187 |
| SNP_A.K_1552902_9_9224373 | 9 | 9224373 | 6.65E-06 | 0.24219 | 192 | 0.124679208 | 0.020836754 |
| SNP_A.K_971666_6_276357 | 6 | 276357 | 6.78E-06 | 0.20313 | 192 | 0.124456639 | 0.021221424 |
| SNP_A.K_29885_1_4292522 | 1 | 4292522 | 6.87E-06 | 0.04948 | 192 | 0.124309067 | 0.021398293 |
| SNP_A.K_726738_4_12039893 | 4 | 12039893 | 6.87E-06 | 0.04167 | 192 | 0.124306277 | 0.021398293 |
| SNP_A.K_708751_4_8110947 | 4 | 8110947 | 6.87E-06 | 0.10677 | 192 | 0.124300163 | 0.021398293 |
| SNP_A.K_727017_4_12069543 | 4 | 12069543 | 6.88E-06 | 0.0625 | 192 | 0.124295948 | 0.021398293 |
| SNP_A.K_1813170_11_8520979 | 11 | 8520979 | 6.90E-06 | 0.15885 | 192 | 0.124265644 | 0.021424047 |
| SNP_A.K_1552895_9_9224249 | 9 | 9224249 | 6.91E-06 | 0.24479 | 192 | 0.124243746 | 0.021433844 |
| SNP_A.K_708775_4_8112515 | 4 | 8112515 | 6.94E-06 | 0.10156 | 192 | 0.124188294 | 0.021507821 |
| SNP_A.K_727015_4_12069326 | 4 | 12069326 | 6.98E-06 | 0.0599 | 192 | 0.124127658 | 0.021571495 |
| SNP_A.K_1812846_11_8473363 | 11 | 8473363 | 7.01E-06 | 0.15885 | 192 | 0.124076736 | 0.021571495 |
| SNP_A.K_616205_3_26976558 | 3 | 26976558 | 7.06E-06 | 0.20573 | 192 | 0.124000804 | 0.021571495 |
| SNP_A.K_1812415_11_8416067 | 11 | 8416067 | 7.06E-06 | 0.15885 | 192 | 0.1239975 | 0.021571495 |
| SNP_A.K_1552956_9_9226016 | 9 | 9226016 | 7.11E-06 | 0.30729 | 192 | 0.123926448 | 0.021571495 |
| SNP_A.K_1812901_11_8478682 | 11 | 8478682 | 7.13E-06 | 0.15365 | 192 | 0.123882922 | 0.021571495 |
| SNP_A.K_723566_4_11474881 | 4 | 11474881 | 7.14E-06 | 0.20573 | 192 | 0.123871536 | 0.021571495 |
| SNP_A.K_1815492_11_8853812 | 11 | 8853812 | 7.16E-06 | 0.15365 | 192 | 0.123849464 | 0.021571495 |
| SNP_A.K_1815518_11_8854583 | 11 | 8854583 | 7.16E-06 | 0.15365 | 192 | 0.123849464 | 0.021571495 |
| SNP_A.K_1815526_11_8854775 | 11 | 8854775 | 7.16E-06 | 0.15365 | 192 | 0.123849464 | 0.021571495 |
| SNP_A.K_1815528_11_8854790 | 11 | 8854790 | 7.16E-06 | 0.15365 | 192 | 0.123849464 | 0.021571495 |
| SNP_A.K_1815529_11_8854805 | 11 | 8854805 | 7.16E-06 | 0.15365 | 192 | 0.123849464 | 0.021571495 |
| SNP_A.K_1815540_11_8854983 | 11 | 8854983 | 7.16E-06 | 0.15365 | 192 | 0.123849464 | 0.021571495 |
| SNP_A.K_1815541_11_8854985 | 11 | 8854985 | 7.16E-06 | 0.15365 | 192 | 0.123849464 | 0.021571495 |
| SNP_A.K_1815651_11_8859767 | 11 | 8859767 | 7.16E-06 | 0.15365 | 192 | 0.123849464 | 0.021571495 |
| SNP_A.K_1815466_11_8852919 | 11 | 8852919 | 7.16E-06 | 0.15365 | 192 | 0.123849464 | 0.021571495 |
| SNP_A.K_1815519_11_8854609 | 11 | 8854609 | 7.16E-06 | 0.15365 | 192 | 0.123849464 | 0.021571495 |
| SNP_A.K_1815720_11_8884036 | 11 | 8884036 | 7.16E-06 | 0.15365 | 192 | 0.123849464 | 0.021571495 |
| SNP_A.K_1813302_11_8528273 | 11 | 8528273 | 7.16E-06 | 0.15885 | 192 | 0.12384185 | 0.021571495 |
| SNP_A.K_1815522_11_8854673 | 11 | 8854673 | 7.18E-06 | 0.15365 | 192 | 0.123812618 | 0.021596458 |
| SNP_A.K_1815451_11_8852451 | 11 | 8852451 | 7.21E-06 | 0.15885 | 192 | 0.123768977 | 0.021632899 |
| SNP_A.K_1815506_11_8854242 | 11 | 8854242 | 7.21E-06 | 0.15365 | 192 | 0.123761269 | 0.021632899 |
| SNP_A.K_1487932_8_26989815 | 8 | 26989815 | 7.27E-06 | 0.04427 | 192 | 0.123670276 | 0.021638064 |
| SNP_A.K_1487974_8_26990831 | 8 | 26990831 | 7.27E-06 | 0.04427 | 192 | 0.123670276 | 0.021638064 |
| SNP_A.K_1487995_8_26991351 | 8 | 26991351 | 7.27E-06 | 0.04427 | 192 | 0.123670276 | 0.021638064 |

| | | | | | | | |
|---|---|---|---|---|---|---|---|
| SNP_A.K_1487942_8_26990147 | 8 | 26990147 | 7.27E-06 | 0.04427 | 192 | 0.123670276 | 0.021638064 |
| SNP_A.K_1487994_8_26991338 | 8 | 26991338 | 7.27E-06 | 0.04427 | 192 | 0.123670276 | 0.021638064 |
| SNP_A.K_707143_4_7162084 | 4 | 7162084 | 7.28E-06 | 0.21615 | 192 | 0.123661915 | 0.021638064 |
| SNP_A.K_708524_4_8060184 | 4 | 8060184 | 7.30E-06 | 0.07552 | 192 | 0.123623653 | 0.021680945 |
| SNP_A.K_709190_4_8237874 | 4 | 8237874 | 7.33E-06 | 0.10417 | 192 | 0.123580886 | 0.021732677 |
| SNP_A.K_508366_3_7553566 | 3 | 7553566 | 7.38E-06 | 0.10677 | 192 | 0.123509771 | 0.02183144 |
| SNP_A.K_724645_4_11650816 | 4 | 11650816 | 7.38E-06 | 0.20573 | 192 | 0.123498035 | 0.02183144 |
| SNP_A.K_725926_4_11885014 | 4 | 11885014 | 7.44E-06 | 0.15885 | 192 | 0.123412682 | 0.021966851 |
| SNP_A.K_623833_3_27887401 | 3 | 27887401 | 7.51E-06 | 0.17969 | 192 | 0.123309747 | 0.022113984 |
| SNP_A.K_1487861_8_26987202 | 8 | 26987202 | 7.52E-06 | 0.04427 | 192 | 0.123298268 | 0.022113984 |
| SNP_A.K_1551750_9_9124402 | 9 | 9124402 | 7.52E-06 | 0.27083 | 192 | 0.123290082 | 0.022113984 |
| SNP_A.K_1813669_11_8565183 | 11 | 8565183 | 7.55E-06 | 0.44792 | 192 | 0.123253896 | 0.022154103 |
| SNP_A.K_1763468_10_21740953 | 10 | 21740953 | 7.60E-06 | 0.17708 | 192 | 0.123180536 | 0.022267979 |
| SNP_A.K_709319_4_8248576 | 4 | 8248576 | 7.64E-06 | 0.21875 | 192 | 0.123121671 | 0.022317097 |
| SNP_A.K_1699032_10_12967177 | 10 | 12967177 | 7.64E-06 | 0.19792 | 192 | 0.123108704 | 0.022317097 |
| SNP_A.K_1815424_11_8851608 | 11 | 8851608 | 7.64E-06 | 0.13802 | 192 | 0.123108492 | 0.022317097 |
| SNP_A.K_1815464_11_8852876 | 11 | 8852876 | 7.69E-06 | 0.15625 | 192 | 0.123042289 | 0.02241654 |
| SNP_A.K_1815491_11_8853768 | 11 | 8853768 | 7.70E-06 | 0.15625 | 192 | 0.123024508 | 0.02241654 |
| SNP_A.K_1815524_11_8854764 | 11 | 8854764 | 7.72E-06 | 0.15625 | 192 | 0.12300265 | 0.02241654 |
| SNP_A.K_708514_4_8059398 | 4 | 8059398 | 7.72E-06 | 0.10417 | 192 | 0.12299583 | 0.02241654 |
| SNP_A.K_724167_4_11577615 | 4 | 11577615 | 7.74E-06 | 0.2526 | 192 | 0.122975771 | 0.022425375 |
| SNP_A.K_1815527_11_8854788 | 11 | 8854788 | 7.76E-06 | 0.15625 | 192 | 0.122940212 | 0.022441455 |
| SNP_A.K_614827_3_26816165 | 3 | 26816165 | 7.78E-06 | 0.16406 | 192 | 0.122909112 | 0.022441455 |
| SNP_A.K_2005748_12_13661416 | 12 | 13661416 | 7.79E-06 | 0.03646 | 192 | 0.122900623 | 0.022441455 |
| SNP_A.K_1815568_11_8855663 | 11 | 8855663 | 7.79E-06 | 0.15625 | 192 | 0.122897417 | 0.022441455 |
| SNP_A.K_709081_4_8218481 | 4 | 8218481 | 7.79E-06 | 0.22135 | 192 | 0.122889873 | 0.022441455 |
| SNP_A.K_1812958_11_8483284 | 11 | 8483284 | 7.88E-06 | 0.15625 | 192 | 0.122768183 | 0.022628308 |
| SNP_A.K_727014_4_12068983 | 4 | 12068983 | 7.88E-06 | 0.0599 | 192 | 0.12276587 | 0.022628308 |
| SNP_A.K_288697_2_5058170 | 2 | 5058170 | 7.91E-06 | 0.01563 | 192 | 0.122720991 | 0.022639919 |
| SNP_A.K_708750_4_8110911 | 4 | 8110911 | 7.92E-06 | 0.10938 | 192 | 0.12271388 | 0.022639919 |
| SNP_A.K_709721_4_8283492 | 4 | 8283492 | 7.92E-06 | 0.11458 | 192 | 0.122713799 | 0.022639919 |
| SNP_A.K_622454_3_27743329 | 3 | 27743329 | 7.97E-06 | 0.17708 | 192 | 0.12264749 | 0.022742927 |
| SNP_A.K_1815715_11_8883252 | 11 | 8883252 | 7.98E-06 | 0.15625 | 192 | 0.122624054 | 0.022759283 |
| SNP_A.K_1487881_8_26989003 | 8 | 26989003 | 8.01E-06 | 0.04427 | 192 | 0.12258164 | 0.022807523 |
| SNP_A.K_613526_3_26712409 | 3 | 26712409 | 8.04E-06 | 0.15885 | 192 | 0.122541623 | 0.022807523 |
| SNP_A.K_1814039_11_8596495 | 11 | 8596495 | 8.04E-06 | 0.15625 | 192 | 0.122535965 | 0.022807523 |
| SNP_A.K_1815516_11_8854522 | 11 | 8854522 | 8.05E-06 | 0.15625 | 192 | 0.122524397 | 0.022807523 |
| SNP_A.K_1815493_11_8853831 | 11 | 8853831 | 8.05E-06 | 0.15625 | 192 | 0.122523763 | 0.022807523 |
| SNP_A.K_1815473_11_8853133 | 11 | 8853133 | 8.10E-06 | 0.15625 | 192 | 0.122459294 | 0.022907866 |
| SNP_A.K_729367_4_12595036 | 4 | 12595036 | 8.16E-06 | 0.20573 | 192 | 0.122378664 | 0.023041894 |
| SNP_A.K_253046_1_40848308 | 1 | 40848308 | 8.19E-06 | 0.14583 | 192 | 0.122334539 | 0.023101417 |
| SNP_A.K_1551925_9_9132746 | 9 | 9132746 | 8.21E-06 | 0.26823 | 192 | 0.122309469 | 0.023121806 |
| SNP_A.K_709503_4_8270885 | 4 | 8270885 | 8.35E-06 | 0.10417 | 192 | 0.122112118 | 0.023500705 |
| SNP_A.K_710546_4_8426906 | 4 | 8426906 | 8.41E-06 | 0.10156 | 192 | 0.122036365 | 0.023628164 |
| SNP_A.K_1777240_11_2482491 | 11 | 2482491 | 8.44E-06 | 0.16927 | 192 | 0.122004569 | 0.023663367 |
| SNP_A.K_1487830_8_26985948 | 8 | 26985948 | 8.49E-06 | 0.04427 | 192 | 0.121937969 | 0.023772389 |
| SNP_A.K_708541_4_8060888 | 4 | 8060888 | 8.50E-06 | 0.10677 | 192 | 0.121922364 | 0.023773524 |
| SNP_A.K_1028139_6_10567125 | 6 | 10567125 | 8.60E-06 | 0.0599 | 192 | 0.121784869 | 0.024034767 |
| SNP_A.K_716331_4_9712846 | 4 | 9712846 | 8.64E-06 | 0.08854 | 192 | 0.121733709 | 0.02408694 |
| SNP_A.K_1815666_11_8860273 | 11 | 8860273 | 8.68E-06 | 0.15885 | 192 | 0.121682109 | 0.02408694 |

| | | | | | | | |
|---|---|---|---|---|---|---|---|
| SNP_A.K_726235_4_11930224 | 4 | 11930224 | 8.69E-06 | 0.2474 | 192 | 0.121666816 | 0.02408694 |
| SNP_A.K_1812418_11_8416286 | 11 | 8416286 | 8.71E-06 | 0.17448 | 192 | 0.12164061 | 0.02408694 |
| SNP_A.K_1813527_11_8544616 | 11 | 8544616 | 8.73E-06 | 0.15885 | 192 | 0.121614258 | 0.02408694 |
| SNP_A.K_1813538_11_8553553 | 11 | 8553553 | 8.73E-06 | 0.15885 | 192 | 0.121614258 | 0.02408694 |
| SNP_A.K_1552875_9_9223466 | 9 | 9223466 | 8.73E-06 | 0.32292 | 192 | 0.121613691 | 0.02408694 |
| SNP_A.K_708571_4_8061962 | 4 | 8061962 | 8.75E-06 | 0.06771 | 192 | 0.12159843 | 0.02408694 |
| SNP_A.K_1813700_11_8567885 | 11 | 8567885 | 8.75E-06 | 0.15885 | 192 | 0.121593949 | 0.02408694 |
| SNP_A.K_1814140_11_8603156 | 11 | 8603156 | 8.75E-06 | 0.15885 | 192 | 0.121593949 | 0.02408694 |
| SNP_A.K_1812883_11_8476548 | 11 | 8476548 | 8.76E-06 | 0.15885 | 192 | 0.121581327 | 0.02408694 |
| SNP_A.K_1813692_11_8566360 | 11 | 8566360 | 8.77E-06 | 0.15885 | 192 | 0.121562374 | 0.02408694 |
| SNP_A.K_1812944_11_8482325 | 11 | 8482325 | 8.78E-06 | 0.15885 | 192 | 0.121560162 | 0.02408694 |
| SNP_A.K_1813735_11_8574808 | 11 | 8574808 | 8.80E-06 | 0.15885 | 192 | 0.121535291 | 0.02408694 |
| SNP_A.K_1815187_11_8821972 | 11 | 8821972 | 8.81E-06 | 0.15885 | 192 | 0.121514201 | 0.02408694 |
| SNP_A.K_1814120_11_8602300 | 11 | 8602300 | 8.81E-06 | 0.15885 | 192 | 0.121513096 | 0.02408694 |
| SNP_A.K_1814209_11_8607690 | 11 | 8607690 | 8.82E-06 | 0.15885 | 192 | 0.121507862 | 0.02408694 |
| SNP_A.K_253453_1_40892441 | 1 | 40892441 | 8.87E-06 | 0.3099 | 192 | 0.121446689 | 0.024160771 |
| SNP_A.K_1814251_11_8610219 | 11 | 8610219 | 8.87E-06 | 0.15885 | 192 | 0.121444235 | 0.024160771 |
| SNP_A.K_623361_3_27826478 | 3 | 27826478 | 8.92E-06 | 0.1901 | 192 | 0.121376756 | 0.024275034 |
| SNP_A.K_728998_4_12425251 | 4 | 12425251 | 8.94E-06 | 0.19271 | 192 | 0.121354535 | 0.024291505 |
| SNP_A.K_1815711_11_8882193 | 11 | 8882193 | 8.97E-06 | 0.15885 | 192 | 0.121321903 | 0.024330642 |
| SNP_A.K_1815649_11_8859743 | 11 | 8859743 | 8.99E-06 | 0.15885 | 192 | 0.121296317 | 0.024354554 |
| SNP_A.K_709683_4_8280900 | 4 | 8280900 | 9.02E-06 | 0.07292 | 192 | 0.121253002 | 0.024417167 |
| SNP_A.K_709276_4_8245560 | 4 | 8245560 | 9.10E-06 | 0.07292 | 192 | 0.12115704 | 0.024595392 |
| SNP_A.K_687591_4_3566516 | 4 | 3566516 | 9.23E-06 | 0.14583 | 192 | 0.121002167 | 0.024905732 |
| SNP_A.K_624391_3_27958906 | 3 | 27958906 | 9.24E-06 | 0.1875 | 192 | 0.120981379 | 0.02490606 |
| SNP_A.K_1551890_9_9131125 | 9 | 9131125 | 9.27E-06 | 0.27865 | 192 | 0.120950611 | 0.02490606 |
| SNP_A.K_709518_4_8271479 | 4 | 8271479 | 9.27E-06 | 0.07031 | 192 | 0.120946628 | 0.02490606 |
| SNP_A.K_726167_4_11925273 | 4 | 11925273 | 9.27E-06 | 0.23958 | 192 | 0.120944108 | 0.02490606 |
| SNP_A.K_1478146_8_24957861 | 8 | 24957861 | 9.29E-06 | 0.17708 | 192 | 0.120926228 | 0.024913739 |
| SNP_A.K_253016_1_40844876 | 1 | 40844876 | 9.37E-06 | 0.15365 | 192 | 0.120833212 | 0.025089363 |
| SNP_A.K_89994_1_12300538 | 1 | 12300538 | 9.44E-06 | 0.14323 | 192 | 0.120744698 | 0.025256127 |
| SNP_A.K_724252_4_11583763 | 4 | 11583763 | 9.46E-06 | 0.25 | 192 | 0.120718351 | 0.025283177 |
| SNP_A.K_1814081_11_8599597 | 11 | 8599597 | 9.51E-06 | 0.17188 | 192 | 0.120664096 | 0.025288739 |
| SNP_A.K_725920_4_11884523 | 4 | 11884523 | 9.51E-06 | 0.1224 | 192 | 0.120657858 | 0.025288739 |
| SNP_A.K_1813749_11_8575447 | 11 | 8575447 | 9.52E-06 | 0.17188 | 192 | 0.12065577 | 0.025288739 |
| SNP_A.K_727013_4_12068877 | 4 | 12068877 | 9.52E-06 | 0.0651 | 192 | 0.120647025 | 0.025288739 |
| SNP_A.K_315436_2_8285183 | 2 | 8285183 | 9.52E-06 | 0.01823 | 192 | 0.120644315 | 0.025288739 |
| SNP_A.K_1813543_11_8554160 | 11 | 8554160 | 9.58E-06 | 0.16146 | 192 | 0.12057825 | 0.025383914 |
| SNP_A.K_706659_4_6930917 | 4 | 6930917 | 9.59E-06 | 0.19271 | 192 | 0.120573795 | 0.025383914 |
| SNP_A.K_1813955_11_8590141 | 11 | 8590141 | 9.60E-06 | 0.16146 | 192 | 0.120558405 | 0.025386564 |
| SNP_A.K_723478_4_11461663 | 4 | 11461663 | 9.63E-06 | 0.21094 | 192 | 0.120520117 | 0.025423037 |
| SNP_A.K_724354_4_11590517 | 4 | 11590517 | 9.71E-06 | 0.23958 | 192 | 0.120433634 | 0.025423037 |
| SNP_A.K_1814780_11_8775095 | 11 | 8775095 | 9.73E-06 | 0.16146 | 192 | 0.1204011 | 0.025423037 |
| SNP_A.K_1813231_11_8524957 | 11 | 8524957 | 9.77E-06 | 0.16146 | 192 | 0.120363646 | 0.025423037 |
| SNP_A.K_709528_4_8271656 | 4 | 8271656 | 9.77E-06 | 0.07031 | 192 | 0.120362906 | 0.025423037 |
| SNP_A.K_709753_4_8284869 | 4 | 8284869 | 9.77E-06 | 0.07031 | 192 | 0.120362906 | 0.025423037 |
| SNP_A.K_373792_2_17594939 | 2 | 17594939 | 9.77E-06 | 0.0651 | 192 | 0.120356043 | 0.025423037 |
| SNP_A.K_1813572_11_8556911 | 11 | 8556911 | 9.79E-06 | 0.16146 | 192 | 0.120341477 | 0.025423037 |
| SNP_A.K_1812021_11_8371174 | 11 | 8371174 | 9.79E-06 | 0.16667 | 192 | 0.120339692 | 0.025423037 |
| SNP_A.K_1813150_11_8518739 | 11 | 8518739 | 9.79E-06 | 0.16146 | 192 | 0.120338209 | 0.025423037 |

| | | | | | | | |
|---|---|---|---|---|---|---|---|
| SNP_A.K_709118_4_8228979 | 4 | 8228979 | 9.81E-06 | 0.07552 | 192 | 0.120315583 | 0.025423037 |
| SNP_A.K_1814109_11_8601687 | 11 | 8601687 | 9.82E-06 | 0.16146 | 192 | 0.120303738 | 0.025423037 |
| SNP_A.K_723428_4_11453962 | 4 | 11453962 | 9.82E-06 | 0.24479 | 192 | 0.120302962 | 0.025423037 |
| SNP_A.K_1815314_11_8834222 | 11 | 8834222 | 9.82E-06 | 0.16146 | 192 | 0.120298947 | 0.025423037 |
| SNP_A.K_1815610_11_8856811 | 11 | 8856811 | 9.83E-06 | 0.16146 | 192 | 0.120295243 | 0.025423037 |
| SNP_A.K_1173613_7_1567280 | 7 | 1567280 | 9.83E-06 | 0.17188 | 192 | 0.120290661 | 0.025423037 |
| SNP_A.K_1814037_11_8596463 | 11 | 8596463 | 9.85E-06 | 0.16146 | 192 | 0.120270493 | 0.025423037 |
| SNP_A.K_1815611_11_8856819 | 11 | 8856819 | 9.85E-06 | 0.16146 | 192 | 0.120268384 | 0.025423037 |
| SNP_A.K_1812744_11_8465140 | 11 | 8465140 | 9.86E-06 | 0.16146 | 192 | 0.120256992 | 0.025423037 |
| SNP_A.K_1813654_11_8564508 | 11 | 8564508 | 9.87E-06 | 0.16146 | 192 | 0.120251314 | 0.025423037 |
| SNP_A.K_1815507_11_8854246 | 11 | 8854246 | 9.87E-06 | 0.15625 | 192 | 0.120247853 | 0.025423037 |
| SNP_A.K_1814858_11_8783237 | 11 | 8783237 | 9.88E-06 | 0.16667 | 192 | 0.120230546 | 0.02542737 |
| SNP_A.K_709128_4_8231909 | 4 | 8231909 | 9.91E-06 | 0.07031 | 192 | 0.120204512 | 0.02542737 |
| SNP_A.K_709130_4_8232406 | 4 | 8232406 | 9.91E-06 | 0.07031 | 192 | 0.120204512 | 0.02542737 |
| SNP_A.K_1812849_11_8473410 | 11 | 8473410 | 9.94E-06 | 0.16146 | 192 | 0.120163498 | 0.025489391 |
| SNP_A.K_1813015_11_8487058 | 11 | 8487058 | 9.98E-06 | 0.16406 | 192 | 0.120124119 | 0.02554787 |
| SNP_A.K_727215_4_12095252 | 4 | 12095252 | 9.99E-06 | 0.06771 | 192 | 0.120109282 | 0.025550375 |
| SNP_A.K_725925_4_11885007 | 4 | 11885007 | 1.00E-05 | 0.16146 | 192 | 0.120064295 | 0.025611476 |
| SNP_A.K_621205_3_27593787 | 3 | 27593787 | 1.00E-05 | 0.14583 | 192 | 0.120055149 | 0.025611476 |
| SNP_A.K_710410_4_8348771 | 4 | 8348771 | 1.01E-05 | 0.10156 | 192 | 0.120010985 | 0.025650213 |
| SNP_A.K_1551976_9_9135874 | 9 | 9135874 | 1.01E-05 | 0.27344 | 192 | 0.120010892 | 0.025650213 |
| SNP_A.K_709221_4_8241644 | 4 | 8241644 | 1.01E-05 | 0.07031 | 192 | 0.119988131 | 0.025671116 |
| SNP_A.K_708758_4_8111191 | 4 | 8111191 | 1.02E-05 | 0.07031 | 192 | 0.119930491 | 0.025674455 |
| SNP_A.K_709529_4_8271720 | 4 | 8271720 | 1.02E-05 | 0.07031 | 192 | 0.11991865 | 0.025674455 |
| SNP_A.K_709531_4_8271851 | 4 | 8271851 | 1.02E-05 | 0.07031 | 192 | 0.11991865 | 0.025674455 |
| SNP_A.K_709688_4_8281178 | 4 | 8281178 | 1.02E-05 | 0.07031 | 192 | 0.11991865 | 0.025674455 |
| SNP_A.K_708817_4_8115098 | 4 | 8115098 | 1.02E-05 | 0.07031 | 192 | 0.11991865 | 0.025674455 |
| SNP_A.K_1551680_9_9120360 | 9 | 9120360 | 1.02E-05 | 0.15625 | 192 | 0.119900062 | 0.025686019 |
| SNP_A.K_708851_4_8117273 | 4 | 8117273 | 1.02E-05 | 0.07292 | 192 | 0.11988176 | 0.02569697 |
| SNP_A.K_709504_4_8270934 | 4 | 8270934 | 1.02E-05 | 0.11458 | 192 | 0.119836259 | 0.025753843 |
| SNP_A.K_709689_4_8281236 | 4 | 8281236 | 1.02E-05 | 0.07031 | 192 | 0.119830028 | 0.025753843 |
| SNP_A.K_708725_4_8089158 | 4 | 8089158 | 1.03E-05 | 0.07031 | 192 | 0.119789125 | 0.025817102 |
| SNP_A.K_708692_4_8086415 | 4 | 8086415 | 1.03E-05 | 0.07031 | 192 | 0.119754856 | 0.025865199 |
| SNP_A.K_935612_5_21781134 | 5 | 21781134 | 1.04E-05 | 0.02604 | 192 | 0.11962806 | 0.026128918 |
| SNP_A.K_709018_4_8213055 | 4 | 8213055 | 1.05E-05 | 0.06771 | 192 | 0.119596299 | 0.026171812 |
| SNP_A.K_709542_4_8272829 | 4 | 8272829 | 1.05E-05 | 0.11198 | 192 | 0.119558964 | 0.026227901 |
| SNP_A.K_1744581_10_18871372 | 10 | 18871372 | 1.05E-05 | 0.08333 | 192 | 0.119527651 | 0.026229696 |
| SNP_A.K_708981_4_8212139 | 4 | 8212139 | 1.05E-05 | 0.07031 | 192 | 0.119518041 | 0.026229696 |
| SNP_A.K_708982_4_8212141 | 4 | 8212141 | 1.05E-05 | 0.07031 | 192 | 0.119518041 | 0.026229696 |
| SNP_A.K_1551676_9_9120167 | 9 | 9120167 | 1.06E-05 | 0.17448 | 192 | 0.119493707 | 0.026255492 |
| SNP_A.K_253251_1_40878818 | 1 | 40878818 | 1.06E-05 | 0.42969 | 192 | 0.119462475 | 0.026297587 |
| SNP_A.K_710492_4_8424885 | 4 | 8424885 | 1.06E-05 | 0.06771 | 192 | 0.119447285 | 0.026301988 |
| SNP_A.K_729659_4_12624469 | 4 | 12624469 | 1.06E-05 | 0.19531 | 192 | 0.119418183 | 0.026339214 |
| SNP_A.K_727124_4_12083242 | 4 | 12083242 | 1.07E-05 | 0.21354 | 192 | 0.119390741 | 0.026372614 |
| SNP_A.K_1814188_11_8605851 | 11 | 8605851 | 1.07E-05 | 0.16406 | 192 | 0.11936748 | 0.026388058 |
| SNP_A.K_1551954_9_9134635 | 9 | 9134635 | 1.07E-05 | 0.26042 | 192 | 0.119347441 | 0.026388058 |
| SNP_A.K_1744591_10_18873229 | 10 | 18873229 | 1.07E-05 | 0.08073 | 192 | 0.119323509 | 0.026388058 |
| SNP_A.K_1815413_11_8851132 | 11 | 8851132 | 1.07E-05 | 0.11458 | 192 | 0.119309349 | 0.026388058 |
| SNP_A.K_1140708_6_26884056 | 6 | 26884056 | 1.07E-05 | 0.36198 | 192 | 0.119299468 | 0.026388058 |
| SNP_A.K_970650_6_123745 | 6 | 123745 | 1.07E-05 | 0.22135 | 192 | 0.119297838 | 0.026388058 |

| | | | | | | | |
|---|---|---|---|---|---|---|---|
| SNP_A.K_467565_2_35155221 | 2 | 35155221 | 1.08E-05 | 0.05469 | 192 | 0.119237417 | 0.026388058 |
| SNP_A.K_1813839_11_8583008 | 11 | 8583008 | 1.08E-05 | 0.16406 | 192 | 0.11922663 | 0.026388058 |
| SNP_A.K_253292_1_40881667 | 1 | 40881667 | 1.08E-05 | 0.40885 | 192 | 0.11920265 | 0.026388058 |
| SNP_A.K_1814248_11_8610136 | 11 | 8610136 | 1.09E-05 | 0.16406 | 192 | 0.119179614 | 0.026388058 |
| SNP_A.K_253270_1_40880221 | 1 | 40880221 | 1.09E-05 | 0.41927 | 192 | 0.119171202 | 0.026388058 |
| SNP_A.K_1814316_11_8613129 | 11 | 8613129 | 1.09E-05 | 0.16406 | 192 | 0.119156004 | 0.026388058 |
| SNP_A.K_725901_4_11879947 | 4 | 11879947 | 1.09E-05 | 0.125 | 192 | 0.119152648 | 0.026388058 |
| SNP_A.K_1279731_7_20912634 | 7 | 20912634 | 1.09E-05 | 0.25 | 192 | 0.119147512 | 0.026388058 |
| SNP_A.K_725204_4_11762492 | 4 | 11762492 | 1.09E-05 | 0.11198 | 192 | 0.119144871 | 0.026388058 |
| SNP_A.K_253227_1_40876471 | 1 | 40876471 | 1.09E-05 | 0.42708 | 192 | 0.119136902 | 0.026388058 |
| SNP_A.K_253253_1_40878855 | 1 | 40878855 | 1.09E-05 | 0.42188 | 192 | 0.119128037 | 0.026388058 |
| SNP_A.K_1813990_11_8594033 | 11 | 8594033 | 1.09E-05 | 0.16406 | 192 | 0.119120506 | 0.026388058 |
| SNP_A.K_1813149_11_8518735 | 11 | 8518735 | 1.09E-05 | 0.16406 | 192 | 0.119116108 | 0.026388058 |
| SNP_A.K_1813283_11_8527538 | 11 | 8527538 | 1.09E-05 | 0.15885 | 192 | 0.119108792 | 0.026388058 |
| SNP_A.K_1813284_11_8527568 | 11 | 8527568 | 1.09E-05 | 0.15885 | 192 | 0.119108792 | 0.026388058 |
| SNP_A.K_1812796_11_8468766 | 11 | 8468766 | 1.10E-05 | 0.16406 | 192 | 0.119085399 | 0.026412765 |
| SNP_A.K_708868_4_8119732 | 4 | 8119732 | 1.10E-05 | 0.07552 | 192 | 0.11904993 | 0.026445648 |
| SNP_A.K_1552858_9_9222496 | 9 | 9222496 | 1.10E-05 | 0.2474 | 192 | 0.119038388 | 0.026445648 |
| SNP_A.K_613555_3_26715552 | 3 | 26715552 | 1.10E-05 | 0.15365 | 192 | 0.119032768 | 0.026445648 |
| SNP_A.K_693278_4_4944727 | 4 | 4944727 | 1.10E-05 | 0.04948 | 192 | 0.118998808 | 0.026495624 |
| SNP_A.K_1763561_10_21773985 | 10 | 21773985 | 1.11E-05 | 0.32552 | 192 | 0.118931669 | 0.026498716 |
| SNP_A.K_723072_4_11368341 | 4 | 11368341 | 1.11E-05 | 0.20833 | 192 | 0.118919881 | 0.026498716 |
| SNP_A.K_1812615_11_8455677 | 11 | 8455677 | 1.11E-05 | 0.16927 | 192 | 0.118910459 | 0.026498716 |
| SNP_A.K_1814147_11_8603570 | 11 | 8603570 | 1.11E-05 | 0.15885 | 192 | 0.118907959 | 0.026498716 |
| SNP_A.K_1813863_11_8583786 | 11 | 8583786 | 1.11E-05 | 0.15885 | 192 | 0.118907959 | 0.026498716 |
| SNP_A.K_1814148_11_8603574 | 11 | 8603574 | 1.11E-05 | 0.15885 | 192 | 0.118907959 | 0.026498716 |
| SNP_A.K_253223_1_40875847 | 1 | 40875847 | 1.11E-05 | 0.40885 | 192 | 0.118907744 | 0.026498716 |
| SNP_A.K_724187_4_11578865 | 4 | 11578865 | 1.11E-05 | 0.25781 | 192 | 0.118893401 | 0.026502466 |
| SNP_A.K_253458_1_40893321 | 1 | 40893321 | 1.12E-05 | 0.36458 | 192 | 0.118852283 | 0.026569915 |
| SNP_A.K_1953667_12_3612128 | 12 | 3612128 | 1.12E-05 | 0.07292 | 192 | 0.118838436 | 0.026572566 |
| SNP_A.K_726056_4_11897126 | 4 | 11897126 | 1.12E-05 | 0.2526 | 192 | 0.11881273 | 0.026603507 |
| SNP_A.K_1487989_8_26991145 | 8 | 26991145 | 1.13E-05 | 0.04948 | 192 | 0.118730155 | 0.026759481 |
| SNP_A.K_1487924_8_26989708 | 8 | 26989708 | 1.13E-05 | 0.04688 | 192 | 0.1187221 | 0.026759481 |
| SNP_A.K_253219_1_40875041 | 1 | 40875041 | 1.13E-05 | 0.41146 | 192 | 0.118694124 | 0.026760841 |
| SNP_A.K_624515_3_27966450 | 3 | 27966450 | 1.14E-05 | 0.19271 | 192 | 0.118683709 | 0.026760841 |
| SNP_A.K_1551989_9_9136691 | 9 | 9136691 | 1.14E-05 | 0.13281 | 192 | 0.118683556 | 0.026760841 |
| SNP_A.K_1487990_8_26991268 | 8 | 26991268 | 1.15E-05 | 0.04688 | 192 | 0.118511395 | 0.027146228 |
| SNP_A.K_709845_4_8299480 | 4 | 8299480 | 1.16E-05 | 0.05208 | 192 | 0.118462032 | 0.027235805 |
| SNP_A.K_1744838_10_18891229 | 10 | 18891229 | 1.16E-05 | 0.0651 | 192 | 0.118446337 | 0.027243367 |
| SNP_A.K_725849_4_11869726 | 4 | 11869726 | 1.16E-05 | 0.125 | 192 | 0.118425373 | 0.027263843 |
| SNP_A.K_708123_4_7543268 | 4 | 7543268 | 1.16E-05 | 0.21615 | 192 | 0.118401874 | 0.027290572 |
| SNP_A.K_614245_3_26780445 | 3 | 26780445 | 1.17E-05 | 0.17448 | 192 | 0.118379153 | 0.027315457 |
| SNP_A.K_725284_4_11781085 | 4 | 11781085 | 1.17E-05 | 0.11458 | 192 | 0.118345501 | 0.027367209 |
| SNP_A.K_1316099_7_27551126 | 7 | 27551126 | 1.17E-05 | 0.08594 | 192 | 0.118326606 | 0.027382844 |
| SNP_A.K_1488948_8_27117165 | 8 | 27117165 | 1.18E-05 | 0.13281 | 192 | 0.118258191 | 0.027520448 |
| SNP_A.K_253205_1_40874424 | 1 | 40874424 | 1.18E-05 | 0.41927 | 192 | 0.118225853 | 0.027548752 |
| SNP_A.K_615255_3_26852039 | 3 | 26852039 | 1.18E-05 | 0.17188 | 192 | 0.118213219 | 0.027548752 |
| SNP_A.K_1804565_11_7086542 | 11 | 7086542 | 1.18E-05 | 0.09115 | 192 | 0.118209283 | 0.027548752 |
| SNP_A.K_708504_4_8055739 | 4 | 8055739 | 1.19E-05 | 0.07813 | 192 | 0.118173251 | 0.027607064 |
| SNP_A.K_1813336_11_8530834 | 11 | 8530834 | 1.20E-05 | 0.16146 | 192 | 0.118108688 | 0.027736415 |

| | | | | | | | |
|---|---|---|---|---|---|---|---|
| SNP_A.K_693811_4_5064993 | 4 | 5064993 | 1.20E-05 | 0.03906 | 192 | 0.118088667 | 0.027755321 |
| SNP_A.K_1487817_8_26985421 | 8 | 26985421 | 1.20E-05 | 0.04688 | 192 | 0.118038094 | 0.027755986 |
| SNP_A.K_253206_1_40874430 | 1 | 40874430 | 1.20E-05 | 0.3099 | 192 | 0.118029693 | 0.027755986 |
| SNP_A.K_1813711_11_8568938 | 11 | 8568938 | 1.21E-05 | 0.16146 | 192 | 0.117960987 | 0.027755986 |
| SNP_A.K_1814722_11_8771460 | 11 | 8771460 | 1.21E-05 | 0.17188 | 192 | 0.117954196 | 0.027755986 |
| SNP_A.K_1777352_11_2490224 | 11 | 2490224 | 1.21E-05 | 0.17188 | 192 | 0.117943948 | 0.027755986 |
| SNP_A.K_1812962_11_8483743 | 11 | 8483743 | 1.21E-05 | 0.16146 | 192 | 0.117939846 | 0.027755986 |
| SNP_A.K_1813017_11_8487219 | 11 | 8487219 | 1.21E-05 | 0.16146 | 192 | 0.117939846 | 0.027755986 |
| SNP_A.K_1813131_11_8517245 | 11 | 8517245 | 1.21E-05 | 0.16146 | 192 | 0.117939846 | 0.027755986 |
| SNP_A.K_1815497_11_8854083 | 11 | 8854083 | 1.22E-05 | 0.16667 | 192 | 0.11791422 | 0.027755986 |
| SNP_A.K_1813834_11_8582468 | 11 | 8582468 | 1.22E-05 | 0.16146 | 192 | 0.117907268 | 0.027755986 |
| SNP_A.K_1813864_11_8583790 | 11 | 8583790 | 1.22E-05 | 0.16146 | 192 | 0.117907268 | 0.027755986 |
| SNP_A.K_1552882_9_9223791 | 9 | 9223791 | 1.22E-05 | 0.20573 | 192 | 0.117869833 | 0.027755986 |
| SNP_A.K_1812957_11_8483224 | 11 | 8483224 | 1.22E-05 | 0.16146 | 192 | 0.117868147 | 0.027755986 |
| SNP_A.K_1812086_11_8377491 | 11 | 8377491 | 1.22E-05 | 0.16146 | 192 | 0.117865734 | 0.027755986 |
| SNP_A.K_1812827_11_8472145 | 11 | 8472145 | 1.22E-05 | 0.16146 | 192 | 0.117865734 | 0.027755986 |
| SNP_A.K_1812355_11_8410465 | 11 | 8410465 | 1.22E-05 | 0.16146 | 192 | 0.117853764 | 0.027755986 |
| SNP_A.K_1814627_11_8765891 | 11 | 8765891 | 1.22E-05 | 0.17188 | 192 | 0.117847369 | 0.027755986 |
| SNP_A.K_1815183_11_8821911 | 11 | 8821911 | 1.23E-05 | 0.16667 | 192 | 0.117826978 | 0.027755986 |
| SNP_A.K_710339_4_8336880 | 4 | 8336880 | 1.23E-05 | 0.05208 | 192 | 0.117824109 | 0.027755986 |
| SNP_A.K_614549_3_26798087 | 3 | 26798087 | 1.23E-05 | 0.16927 | 192 | 0.117823109 | 0.027755986 |
| SNP_A.K_709222_4_8241651 | 4 | 8241651 | 1.23E-05 | 0.07292 | 192 | 0.117819786 | 0.027755986 |
| SNP_A.K_709562_4_8273825 | 4 | 8273825 | 1.23E-05 | 0.22135 | 192 | 0.117806528 | 0.027755986 |
| SNP_A.K_709760_4_8285202 | 4 | 8285202 | 1.23E-05 | 0.09115 | 192 | 0.117806469 | 0.027755986 |
| SNP_A.K_1812025_11_8371435 | 11 | 8371435 | 1.23E-05 | 0.16667 | 192 | 0.11778651 | 0.027775576 |
| SNP_A.K_813337_4_31054643 | 4 | 31054643 | 1.23E-05 | 0.07552 | 192 | 0.117765317 | 0.027798291 |
| SNP_A.K_708570_4_8061896 | 4 | 8061896 | 1.23E-05 | 0.07813 | 192 | 0.117750553 | 0.027805013 |
| SNP_A.K_253457_1_40893176 | 1 | 40893176 | 1.25E-05 | 0.36458 | 192 | 0.117570632 | 0.02822703 |
| SNP_A.K_725196_4_11762123 | 4 | 11762123 | 1.26E-05 | 0.11198 | 192 | 0.117510971 | 0.028347934 |
| SNP_A.K_709014_4_8212856 | 4 | 8212856 | 1.27E-05 | 0.11198 | 192 | 0.117464516 | 0.028435672 |
| SNP_A.K_1815484_11_8853460 | 11 | 8853460 | 1.27E-05 | 0.16406 | 192 | 0.117438801 | 0.028470671 |
| SNP_A.K_253491_1_40897248 | 1 | 40897248 | 1.27E-05 | 0.16927 | 192 | 0.117394908 | 0.028552299 |
| SNP_A.K_730514_4_12697659 | 4 | 12697659 | 1.28E-05 | 0.22135 | 192 | 0.117381003 | 0.028557221 |
| SNP_A.K_1551732_9_9123876 | 9 | 9123876 | 1.28E-05 | 0.26823 | 192 | 0.117329988 | 0.028657491 |
| SNP_A.K_970963_6_162183 | 6 | 162183 | 1.29E-05 | 0.26042 | 192 | 0.117278178 | 0.028760205 |
| SNP_A.K_709416_4_8265516 | 4 | 8265516 | 1.29E-05 | 0.07552 | 192 | 0.11721668 | 0.028866339 |
| SNP_A.K_725270_4_11777022 | 4 | 11777022 | 1.30E-05 | 0.21354 | 192 | 0.117180809 | 0.028866339 |
| SNP_A.K_709206_4_8238891 | 4 | 8238891 | 1.30E-05 | 0.08073 | 192 | 0.117154092 | 0.028866339 |
| SNP_A.K_1180612_7_3237598 | 7 | 3237598 | 1.30E-05 | 0.09115 | 192 | 0.117130269 | 0.028866339 |
| SNP_A.K_709691_4_8281401 | 4 | 8281401 | 1.31E-05 | 0.07552 | 192 | 0.117096658 | 0.028866339 |
| SNP_A.K_1814996_11_8794482 | 11 | 8794482 | 1.31E-05 | 0.16927 | 192 | 0.117095407 | 0.028866339 |
| SNP_A.K_1815280_11_8830682 | 11 | 8830682 | 1.31E-05 | 0.16406 | 192 | 0.117087888 | 0.028866339 |
| SNP_A.K_709286_4_8245822 | 4 | 8245822 | 1.31E-05 | 0.07552 | 192 | 0.117086358 | 0.028866339 |
| SNP_A.K_1812553_11_8450365 | 11 | 8450365 | 1.31E-05 | 0.16927 | 192 | 0.11707758 | 0.028866339 |
| SNP_A.K_709207_4_8238911 | 4 | 8238911 | 1.32E-05 | 0.07552 | 192 | 0.117034587 | 0.028866339 |
| SNP_A.K_1812458_11_8437193 | 11 | 8437193 | 1.32E-05 | 0.16406 | 192 | 0.117032288 | 0.028866339 |
| SNP_A.K_727018_4_12069629 | 4 | 12069629 | 1.32E-05 | 0.06771 | 192 | 0.117031144 | 0.028866339 |
| SNP_A.K_253271_1_40880525 | 1 | 40880525 | 1.32E-05 | 0.42188 | 192 | 0.117029527 | 0.028866339 |
| SNP_A.K_1027060_6_10359412 | 6 | 10359412 | 1.32E-05 | 0.03125 | 192 | 0.117005383 | 0.028866339 |
| SNP_A.K_726924_4_12057240 | 4 | 12057240 | 1.32E-05 | 0.20833 | 192 | 0.116991394 | 0.028866339 |

| | | | | | | | |
|---|---|---|---|---|---|---|---|
| SNP_A.K_817414_4_31496720 | 4 | 31496720 | 1.32E-05 | 0.02083 | 192 | 0.116989238 | 0.028866339 |
| SNP_A.K_708818_4_8115105 | 4 | 8115105 | 1.33E-05 | 0.07292 | 192 | 0.116922505 | 0.028866339 |
| SNP_A.K_1141392_6_26999034 | 6 | 26999034 | 1.33E-05 | 0.26823 | 192 | 0.116902085 | 0.028866339 |
| SNP_A.K_1812973_11_8484308 | 11 | 8484308 | 1.34E-05 | 0.16406 | 192 | 0.1168232 | 0.028866339 |
| SNP_A.K_1813869_11_8583987 | 11 | 8583987 | 1.35E-05 | 0.16406 | 192 | 0.116768997 | 0.028866339 |
| SNP_A.K_1815144_11_8807874 | 11 | 8807874 | 1.35E-05 | 0.16406 | 192 | 0.1167583 | 0.028866339 |
| SNP_A.K_1813805_11_8580036 | 11 | 8580036 | 1.35E-05 | 0.16406 | 192 | 0.1167583 | 0.028866339 |
| SNP_A.K_1814153_11_8603831 | 11 | 8603831 | 1.35E-05 | 0.16406 | 192 | 0.116757792 | 0.028866339 |
| SNP_A.K_1814318_11_8613190 | 11 | 8613190 | 1.35E-05 | 0.16406 | 192 | 0.116757792 | 0.028866339 |
| SNP_A.K_1815098_11_8803478 | 11 | 8803478 | 1.35E-05 | 0.16406 | 192 | 0.116757792 | 0.028866339 |
| SNP_A.K_1813868_11_8583861 | 11 | 8583861 | 1.35E-05 | 0.16406 | 192 | 0.116757792 | 0.028866339 |
| SNP_A.K_1814101_11_8600921 | 11 | 8600921 | 1.35E-05 | 0.16406 | 192 | 0.116757792 | 0.028866339 |
| SNP_A.K_1814321_11_8613369 | 11 | 8613369 | 1.35E-05 | 0.16406 | 192 | 0.116757792 | 0.028866339 |
| SNP_A.K_1813659_11_8564710 | 11 | 8564710 | 1.35E-05 | 0.16406 | 192 | 0.11675337 | 0.028866339 |
| SNP_A.K_1812673_11_8461889 | 11 | 8461889 | 1.35E-05 | 0.16406 | 192 | 0.116739604 | 0.028866339 |
| SNP_A.K_1812528_11_8447848 | 11 | 8447848 | 1.35E-05 | 0.16406 | 192 | 0.116738081 | 0.028866339 |
| SNP_A.K_253235_1_40877112 | 1 | 40877112 | 1.35E-05 | 0.41667 | 192 | 0.116734781 | 0.028866339 |
| SNP_A.K_708872_4_8120638 | 4 | 8120638 | 1.35E-05 | 0.07292 | 192 | 0.116728095 | 0.028866339 |
| SNP_A.K_708679_4_8084885 | 4 | 8084885 | 1.35E-05 | 0.07292 | 192 | 0.116724717 | 0.028866339 |
| SNP_A.K_1812423_11_8416830 | 11 | 8416830 | 1.35E-05 | 0.16406 | 192 | 0.116719682 | 0.028866339 |
| SNP_A.K_1812535_11_8448437 | 11 | 8448437 | 1.35E-05 | 0.16406 | 192 | 0.116719682 | 0.028866339 |
| SNP_A.K_1812551_11_8450228 | 11 | 8450228 | 1.35E-05 | 0.16406 | 192 | 0.116719682 | 0.028866339 |
| SNP_A.K_1812534_11_8448330 | 11 | 8448330 | 1.35E-05 | 0.16406 | 192 | 0.116719682 | 0.028866339 |
| SNP_A.K_1812831_11_8472271 | 11 | 8472271 | 1.35E-05 | 0.16406 | 192 | 0.116719682 | 0.028866339 |
| SNP_A.K_709635_4_8278828 | 4 | 8278828 | 1.35E-05 | 0.07031 | 192 | 0.116715077 | 0.028866339 |
| SNP_A.K_1812003_11_8370003 | 11 | 8370003 | 1.35E-05 | 0.16927 | 192 | 0.116713762 | 0.028866339 |
| SNP_A.K_1813723_11_8571004 | 11 | 8571004 | 1.35E-05 | 0.16406 | 192 | 0.116713598 | 0.028866339 |
| SNP_A.K_1814118_11_8602101 | 11 | 8602101 | 1.36E-05 | 0.16927 | 192 | 0.116708446 | 0.028866339 |
| SNP_A.K_1813726_11_8571662 | 11 | 8571662 | 1.36E-05 | 0.16406 | 192 | 0.116699087 | 0.028866339 |
| SNP_A.K_1814149_11_8603612 | 11 | 8603612 | 1.36E-05 | 0.16406 | 192 | 0.11669641 | 0.028866339 |
| SNP_A.K_1812597_11_8454973 | 11 | 8454973 | 1.36E-05 | 0.16406 | 192 | 0.116684379 | 0.028866339 |
| SNP_A.K_419091_2_24516471 | 2 | 24516471 | 1.36E-05 | 0.04948 | 192 | 0.116679248 | 0.028866339 |
| SNP_A.K_725245_4_11766665 | 4 | 11766665 | 1.36E-05 | 0.11198 | 192 | 0.116658685 | 0.028866339 |
| SNP_A.K_1812894_11_8477443 | 11 | 8477443 | 1.36E-05 | 0.16406 | 192 | 0.116651115 | 0.028866339 |
| SNP_A.K_253222_1_40875720 | 1 | 40875720 | 1.36E-05 | 0.42448 | 192 | 0.116647702 | 0.028866339 |
| SNP_A.K_709370_4_8259079 | 4 | 8259079 | 1.36E-05 | 0.07292 | 192 | 0.116644005 | 0.028866339 |
| SNP_A.K_1814750_11_8772404 | 11 | 8772404 | 1.37E-05 | 0.16406 | 192 | 0.116628159 | 0.028878093 |
| SNP_A.K_614553_3_26798280 | 3 | 26798280 | 1.37E-05 | 0.16927 | 192 | 0.116588876 | 0.028950788 |
| SNP_A.K_708648_4_8070775 | 4 | 8070775 | 1.37E-05 | 0.07552 | 192 | 0.116553724 | 0.028985426 |
| SNP_A.K_706763_4_6934952 | 4 | 6934952 | 1.38E-05 | 0.20313 | 192 | 0.116544322 | 0.028985426 |
| SNP_A.K_708947_4_8151434 | 4 | 8151434 | 1.38E-05 | 0.20313 | 192 | 0.116536376 | 0.028985426 |
| SNP_A.K_1744583_10_18871643 | 10 | 18871643 | 1.38E-05 | 0.08073 | 192 | 0.116530473 | 0.028985426 |
| SNP_A.K_1310463_7_26336641 | 7 | 26336641 | 1.38E-05 | 0.1849 | 192 | 0.116515809 | 0.028994332 |
| SNP_A.K_1812614_11_8455655 | 11 | 8455655 | 1.39E-05 | 0.16406 | 192 | 0.116454503 | 0.029096007 |
| SNP_A.K_1551808_9_9126913 | 9 | 9126913 | 1.39E-05 | 0.27083 | 192 | 0.116454403 | 0.029096007 |
| SNP_A.K_723325_4_11432828 | 4 | 11432828 | 1.39E-05 | 0.20573 | 192 | 0.116414596 | 0.029123614 |
| SNP_A.K_1812126_11_8381841 | 11 | 8381841 | 1.39E-05 | 0.16406 | 192 | 0.116398442 | 0.029123614 |
| SNP_A.K_725208_4_11762688 | 4 | 11762688 | 1.39E-05 | 0.11198 | 192 | 0.116390522 | 0.029123614 |
| SNP_A.K_272449_2_1512770 | 2 | 1512770 | 1.40E-05 | 0.04427 | 192 | 0.116385099 | 0.029123614 |
| SNP_A.K_728226_4_12292000 | 4 | 12292000 | 1.40E-05 | 0.21094 | 192 | 0.11637726 | 0.029123614 |

| | | | | | | | |
|---|---|---|---|---|---|---|---|
| SNP_A.K_709087_4_8221014 | 4 | 8221014 | 1.40E-05 | 0.07552 | 192 | 0.116376761 | 0.029123614 |
| SNP_A.K_253201_1_40873876 | 1 | 40873876 | 1.40E-05 | 0.42448 | 192 | 0.11635584 | 0.029149228 |
| SNP_A.K_689530_4_4260652 | 4 | 4260652 | 1.40E-05 | 0.14063 | 192 | 0.116311387 | 0.029232879 |
| SNP_A.K_253252_1_40878834 | 1 | 40878834 | 1.41E-05 | 0.42708 | 192 | 0.116301712 | 0.029232879 |
| SNP_A.K_709522_4_8271574 | 4 | 8271574 | 1.41E-05 | 0.07552 | 192 | 0.116266126 | 0.029297278 |
| SNP_A.K_760861_4_19811966 | 4 | 19811966 | 1.42E-05 | 0.03125 | 192 | 0.116188217 | 0.029423545 |
| SNP_A.K_708706_4_8087942 | 4 | 8087942 | 1.42E-05 | 0.07292 | 192 | 0.116184402 | 0.029423545 |
| SNP_A.K_613533_3_26712753 | 3 | 26712753 | 1.42E-05 | 0.16146 | 192 | 0.116171346 | 0.029423545 |
| SNP_A.K_710467_4_8423720 | 4 | 8423720 | 1.42E-05 | 0.1276 | 192 | 0.116165063 | 0.029423545 |
| SNP_A.K_1744586_10_18872160 | 10 | 18872160 | 1.42E-05 | 0.08594 | 192 | 0.116162926 | 0.029423545 |
| SNP_A.K_970628_6_120967 | 6 | 120967 | 1.43E-05 | 0.21094 | 192 | 0.116121589 | 0.02950381 |
| SNP_A.K_709001_4_8212587 | 4 | 8212587 | 1.43E-05 | 0.07031 | 192 | 0.116094644 | 0.029510641 |
| SNP_A.K_1551733_9_9123916 | 9 | 9123916 | 1.43E-05 | 0.27344 | 192 | 0.116093609 | 0.029510641 |
| SNP_A.K_729028_4_12430012 | 4 | 12430012 | 1.43E-05 | 0.23698 | 192 | 0.116085965 | 0.029510641 |
| SNP_A.K_728236_4_12293902 | 4 | 12293902 | 1.44E-05 | 0.21094 | 192 | 0.116050967 | 0.029574406 |
| SNP_A.K_613976_3_26750990 | 3 | 26750990 | 1.44E-05 | 0.17448 | 192 | 0.116014227 | 0.029642986 |
| SNP_A.K_970730_6_128233 | 6 | 128233 | 1.45E-05 | 0.25521 | 192 | 0.115993251 | 0.029669669 |
| SNP_A.K_730518_4_12697769 | 4 | 12697769 | 1.45E-05 | 0.21354 | 192 | 0.115963534 | 0.029719759 |
| SNP_A.K_1814068_11_8599167 | 11 | 8599167 | 1.46E-05 | 0.17188 | 192 | 0.115904396 | 0.029796773 |
| SNP_A.K_1814151_11_8603709 | 11 | 8603709 | 1.46E-05 | 0.16667 | 192 | 0.115898644 | 0.029796773 |
| SNP_A.K_725259_4_11772124 | 4 | 11772124 | 1.46E-05 | 0.10938 | 192 | 0.115882632 | 0.029796773 |
| SNP_A.K_1812629_11_8457301 | 11 | 8457301 | 1.46E-05 | 0.16667 | 192 | 0.115874679 | 0.029796773 |
| SNP_A.K_708900_4_8122790 | 4 | 8122790 | 1.46E-05 | 0.07292 | 192 | 0.115872107 | 0.029796773 |
| SNP_A.K_708380_4_8041596 | 4 | 8041596 | 1.46E-05 | 0.07292 | 192 | 0.115858663 | 0.029796773 |
| SNP_A.K_709310_4_8248073 | 4 | 8248073 | 1.46E-05 | 0.11719 | 192 | 0.115855798 | 0.029796773 |
| SNP_A.K_724865_4_11690165 | 4 | 11690165 | 1.47E-05 | 0.20573 | 192 | 0.115837542 | 0.029796773 |
| SNP_A.K_1814049_11_8597252 | 11 | 8597252 | 1.47E-05 | 0.16667 | 192 | 0.11583448 | 0.029796773 |
| SNP_A.K_708127_4_7544397 | 4 | 7544397 | 1.47E-05 | 0.22917 | 192 | 0.115825757 | 0.029796773 |
| SNP_A.K_709646_4_8279119 | 4 | 8279119 | 1.47E-05 | 0.07292 | 192 | 0.115790895 | 0.029861228 |
| SNP_A.K_1815124_11_8806254 | 11 | 8806254 | 1.48E-05 | 0.16667 | 192 | 0.115706038 | 0.029970517 |
| SNP_A.K_724627_4_11647596 | 4 | 11647596 | 1.48E-05 | 0.21094 | 192 | 0.115696948 | 0.029970517 |
| SNP_A.K_708635_4_8068976 | 4 | 8068976 | 1.49E-05 | 0.07292 | 192 | 0.115652097 | 0.029970517 |
| SNP_A.K_1813647_11_8564287 | 11 | 8564287 | 1.49E-05 | 0.16667 | 192 | 0.115646881 | 0.029970517 |
| SNP_A.K_709396_4_8264967 | 4 | 8264967 | 1.49E-05 | 0.11198 | 192 | 0.115635948 | 0.029970517 |
| SNP_A.K_707120_4_7157830 | 4 | 7157830 | 1.49E-05 | 0.22135 | 192 | 0.115628242 | 0.029970517 |
| SNP_A.K_1813747_11_8575436 | 11 | 8575436 | 1.49E-05 | 0.17188 | 192 | 0.115625906 | 0.029970517 |
| SNP_A.K_1763063_10_21617798 | 10 | 21617798 | 1.50E-05 | 0.26042 | 192 | 0.115612522 | 0.029970517 |
| SNP_A.K_709585_4_8274829 | 4 | 8274829 | 1.50E-05 | 0.07292 | 192 | 0.115594795 | 0.029970517 |
| SNP_A.K_709519_4_8271517 | 4 | 8271517 | 1.50E-05 | 0.07292 | 192 | 0.115593844 | 0.029970517 |
| SNP_A.K_1813639_11_8563318 | 11 | 8563318 | 1.50E-05 | 0.16667 | 192 | 0.115587599 | 0.029970517 |
| SNP_A.K_1813904_11_8586993 | 11 | 8586993 | 1.50E-05 | 0.16667 | 192 | 0.115586387 | 0.029970517 |
| SNP_A.K_1813698_11_8566910 | 11 | 8566910 | 1.50E-05 | 0.16667 | 192 | 0.115580553 | 0.029970517 |
| SNP_A.K_709434_4_8265939 | 4 | 8265939 | 1.50E-05 | 0.07292 | 192 | 0.11557616 | 0.029970517 |
| SNP_A.K_1812753_11_8465735 | 11 | 8465735 | 1.50E-05 | 0.16667 | 192 | 0.115574267 | 0.029970517 |
| SNP_A.K_1813870_11_8584215 | 11 | 8584215 | 1.51E-05 | 0.16667 | 192 | 0.115542911 | 0.029970517 |
| SNP_A.K_709652_4_8279416 | 4 | 8279416 | 1.51E-05 | 0.07292 | 192 | 0.115537477 | 0.029970517 |
| SNP_A.K_1814880_11_8784070 | 11 | 8784070 | 1.51E-05 | 0.17188 | 192 | 0.115527544 | 0.029970517 |
| SNP_A.K_1552238_9_9162362 | 9 | 9162362 | 1.51E-05 | 0.29948 | 192 | 0.115526656 | 0.029970517 |
| SNP_A.K_708721_4_8089093 | 4 | 8089093 | 1.51E-05 | 0.08073 | 192 | 0.115519439 | 0.029970517 |
| SNP_A.K_708723_4_8089101 | 4 | 8089101 | 1.51E-05 | 0.08073 | 192 | 0.115519439 | 0.029970517 |

| | | | | | | | |
|---|---|---|---|---|---|---|---|
| SNP_A.K_705198_4_6538131 | 4 | 6538131 | 1.51E-05 | 0.17969 | 192 | 0.115514484 | 0.029970517 |
| SNP_A.K_707034_4_6997494 | 4 | 6997494 | 1.51E-05 | 0.23177 | 192 | 0.115473726 | 0.030051971 |
| SNP_A.K_708909_4_8124192 | 4 | 8124192 | 1.52E-05 | 0.07292 | 192 | 0.115459219 | 0.030062571 |
| SNP_A.K_709644_4_8279097 | 4 | 8279097 | 1.52E-05 | 0.07292 | 192 | 0.115405216 | 0.030180292 |
| SNP_A.K_1815299_11_8832849 | 11 | 8832849 | 1.53E-05 | 0.16146 | 192 | 0.115393196 | 0.03018424 |
| SNP_A.K_1814615_11_8765159 | 11 | 8765159 | 1.53E-05 | 0.17188 | 192 | 0.115371207 | 0.030197989 |
| SNP_A.K_725187_4_11748161 | 4 | 11748161 | 1.53E-05 | 0.11979 | 192 | 0.115367033 | 0.030197989 |
| SNP_A.K_1170399_7_1023301 | 7 | 1023301 | 1.53E-05 | 0.10938 | 192 | 0.115330832 | 0.030267828 |
| SNP_A.K_1552038_9_9140319 | 9 | 9140319 | 1.54E-05 | 0.32813 | 192 | 0.115303347 | 0.030314068 |
| SNP_A.K_730512_4_12697611 | 4 | 12697611 | 1.54E-05 | 0.27604 | 192 | 0.115277483 | 0.030355978 |
| SNP_A.K_1813886_11_8585843 | 11 | 8585843 | 1.55E-05 | 0.17708 | 192 | 0.115232743 | 0.030429254 |
| SNP_A.K_723448_4_11457373 | 4 | 11457373 | 1.55E-05 | 0.07031 | 192 | 0.1152297 | 0.030429254 |
| SNP_A.K_1813175_11_8521293 | 11 | 8521293 | 1.55E-05 | 0.17708 | 192 | 0.115188898 | 0.030444295 |
| SNP_A.K_708899_4_8122743 | 4 | 8122743 | 1.56E-05 | 0.07292 | 192 | 0.115178812 | 0.030444295 |
| SNP_A.K_288265_2_5001429 | 2 | 5001429 | 1.56E-05 | 0.01823 | 192 | 0.115169862 | 0.030444295 |
| SNP_A.K_708652_4_8071149 | 4 | 8071149 | 1.56E-05 | 0.07292 | 192 | 0.115161454 | 0.030444295 |
| SNP_A.K_709578_4_8274664 | 4 | 8274664 | 1.56E-05 | 0.07292 | 192 | 0.115161454 | 0.030444295 |
| SNP_A.K_709579_4_8274667 | 4 | 8274667 | 1.56E-05 | 0.07292 | 192 | 0.115161454 | 0.030444295 |
| SNP_A.K_624814_3_27997593 | 3 | 27997593 | 1.56E-05 | 0.26563 | 192 | 0.115126824 | 0.030477789 |
| SNP_A.K_391733_2_20195362 | 2 | 20195362 | 1.56E-05 | 0.03906 | 192 | 0.115122292 | 0.030477789 |
| SNP_A.K_253224_1_40875874 | 1 | 40875874 | 1.56E-05 | 0.40885 | 192 | 0.115117997 | 0.030477789 |
| SNP_A.K_723453_4_11457638 | 4 | 11457638 | 1.57E-05 | 0.21094 | 192 | 0.115082926 | 0.030545579 |
| SNP_A.K_1744831_10_18890945 | 10 | 18890945 | 1.58E-05 | 0.08333 | 192 | 0.114988479 | 0.030777668 |
| SNP_A.K_809301_4_30033826 | 4 | 30033826 | 1.58E-05 | 0.01302 | 192 | 0.114970948 | 0.030797511 |
| SNP_A.K_726963_4_12061610 | 4 | 12061610 | 1.59E-05 | 0.21615 | 192 | 0.114934335 | 0.030865012 |
| SNP_A.K_708895_4_8122443 | 4 | 8122443 | 1.59E-05 | 0.07292 | 192 | 0.114924047 | 0.030865012 |
| SNP_A.K_547271_3_15448670 | 3 | 15448670 | 1.59E-05 | 0.1276 | 192 | 0.114912146 | 0.030865012 |
| SNP_A.K_687582_4_3566163 | 4 | 3566163 | 1.59E-05 | 0.08594 | 192 | 0.114905242 | 0.030865012 |
| SNP_A.K_1552883_9_9223800 | 9 | 9223800 | 1.60E-05 | 0.20833 | 192 | 0.114830575 | 0.031037573 |
| SNP_A.K_709003_4_8212623 | 4 | 8212623 | 1.61E-05 | 0.07031 | 192 | 0.114822701 | 0.031037573 |
| SNP_A.K_253213_1_40874598 | 1 | 40874598 | 1.61E-05 | 0.41406 | 192 | 0.114790472 | 0.03109893 |
| SNP_A.K_253159_1_40869276 | 1 | 40869276 | 1.61E-05 | 0.42188 | 192 | 0.114767955 | 0.03111104 |
| SNP_A.K_724240_4_11583094 | 4 | 11583094 | 1.62E-05 | 0.13802 | 192 | 0.114754139 | 0.03111104 |
| SNP_A.K_1552903_9_9224383 | 9 | 9224383 | 1.62E-05 | 0.25781 | 192 | 0.114742061 | 0.03111104 |
| SNP_A.K_253268_1_40879996 | 1 | 40879996 | 1.62E-05 | 0.42188 | 192 | 0.114737793 | 0.03111104 |
| SNP_A.K_1812565_11_8452162 | 11 | 8452162 | 1.62E-05 | 0.16927 | 192 | 0.114734777 | 0.03111104 |
| SNP_A.K_1812803_11_8470002 | 11 | 8470002 | 1.63E-05 | 0.16927 | 192 | 0.114672327 | 0.031207586 |
| SNP_A.K_1813982_11_8592755 | 11 | 8592755 | 1.63E-05 | 0.16927 | 192 | 0.114671784 | 0.031207586 |
| SNP_A.K_1316100_7_27551212 | 7 | 27551212 | 1.63E-05 | 0.08594 | 192 | 0.114659774 | 0.031207586 |
| SNP_A.K_710381_4_8339403 | 4 | 8339403 | 1.63E-05 | 0.36458 | 192 | 0.114633981 | 0.031207586 |
| SNP_A.K_1814063_11_8597610 | 11 | 8597610 | 1.64E-05 | 0.16406 | 192 | 0.114619359 | 0.031207586 |
| SNP_A.K_727818_4_12215577 | 4 | 12215577 | 1.64E-05 | 0.20313 | 192 | 0.114586127 | 0.031207586 |
| SNP_A.K_1310431_7_26333039 | 7 | 26333039 | 1.64E-05 | 0.11198 | 192 | 0.114569242 | 0.031207586 |
| SNP_A.K_1813731_11_8574588 | 11 | 8574588 | 1.64E-05 | 0.16927 | 192 | 0.114561358 | 0.031207586 |
| SNP_A.K_1813943_11_8589370 | 11 | 8589370 | 1.65E-05 | 0.16927 | 192 | 0.114551508 | 0.031207586 |
| SNP_A.K_710088_4_8315732 | 4 | 8315732 | 1.65E-05 | 0.38021 | 192 | 0.114488987 | 0.031207586 |
| SNP_A.K_1814297_11_8612075 | 11 | 8612075 | 1.66E-05 | 0.16927 | 192 | 0.114484055 | 0.031207586 |
| SNP_A.K_1812420_11_8416383 | 11 | 8416383 | 1.66E-05 | 0.1875 | 192 | 0.114475124 | 0.031207586 |
| SNP_A.K_709052_4_8215598 | 4 | 8215598 | 1.66E-05 | 0.11458 | 192 | 0.114474997 | 0.031207586 |
| SNP_A.K_1551913_9_9132049 | 9 | 9132049 | 1.66E-05 | 0.27083 | 192 | 0.114467878 | 0.031207586 |

| | | | | | | | |
|---|---|---|---|---|---|---|---|
| SNP_A.K_725263_4_11773621 | 4 | 11773621 | 1.66E-05 | 0.11198 | 192 | 0.114462633 | 0.031207586 |
| SNP_A.K_1814263_11_8610879 | 11 | 8610879 | 1.66E-05 | 0.16406 | 192 | 0.114462584 | 0.031207586 |
| SNP_A.K_1552885_9_9223861 | 9 | 9223861 | 1.66E-05 | 0.26042 | 192 | 0.11446145 | 0.031207586 |
| SNP_A.K_1813655_11_8564618 | 11 | 8564618 | 1.66E-05 | 0.16406 | 192 | 0.114458892 | 0.031207586 |
| SNP_A.K_1813710_11_8568799 | 11 | 8568799 | 1.66E-05 | 0.16406 | 192 | 0.114450245 | 0.031207586 |
| SNP_A.K_1814157_11_8604017 | 11 | 8604017 | 1.66E-05 | 0.16406 | 192 | 0.114450245 | 0.031207586 |
| SNP_A.K_1814171_11_8604636 | 11 | 8604636 | 1.66E-05 | 0.16406 | 192 | 0.114450245 | 0.031207586 |
| SNP_A.K_1814215_11_8607924 | 11 | 8607924 | 1.66E-05 | 0.16406 | 192 | 0.114450245 | 0.031207586 |
| SNP_A.K_1813907_11_8587148 | 11 | 8587148 | 1.66E-05 | 0.16406 | 192 | 0.114450245 | 0.031207586 |
| SNP_A.K_1815263_11_8828450 | 11 | 8828450 | 1.66E-05 | 0.16406 | 192 | 0.114450245 | 0.031207586 |
| SNP_A.K_1812136_11_8382229 | 11 | 8382229 | 1.66E-05 | 0.16406 | 192 | 0.114423913 | 0.031207586 |
| SNP_A.K_708929_4_8150353 | 4 | 8150353 | 1.66E-05 | 0.22135 | 192 | 0.114422326 | 0.031207586 |
| SNP_A.K_726055_4_11897120 | 4 | 11897120 | 1.67E-05 | 0.25781 | 192 | 0.114414498 | 0.031207586 |
| SNP_A.K_1552838_9_9221010 | 9 | 9221010 | 1.67E-05 | 0.1276 | 192 | 0.114414199 | 0.031207586 |
| SNP_A.K_1782951_11_3523512 | 11 | 3523512 | 1.67E-05 | 0.03646 | 192 | 0.114407098 | 0.031207586 |
| SNP_A.K_725285_4_11781186 | 4 | 11781186 | 1.67E-05 | 0.11458 | 192 | 0.11437895 | 0.03125874 |
| SNP_A.K_1813379_11_8535300 | 11 | 8535300 | 1.68E-05 | 0.14323 | 192 | 0.114306313 | 0.031427956 |
| SNP_A.K_1552846_9_9222184 | 9 | 9222184 | 1.68E-05 | 0.20052 | 192 | 0.114299128 | 0.031427956 |
| SNP_A.K_672649_4_268874 | 4 | 268874 | 1.69E-05 | 0.07813 | 192 | 0.114286997 | 0.031434147 |
| SNP_A.K_1813797_11_8579062 | 11 | 8579062 | 1.69E-05 | 0.17448 | 192 | 0.114248716 | 0.031514558 |
| SNP_A.K_709292_4_8246367 | 4 | 8246367 | 1.70E-05 | 0.13021 | 192 | 0.114222466 | 0.031560959 |
| SNP_A.K_709026_4_8213700 | 4 | 8213700 | 1.70E-05 | 0.07292 | 192 | 0.114177005 | 0.03166224 |
| SNP_A.K_725291_4_11782974 | 4 | 11782974 | 1.71E-05 | 0.11458 | 192 | 0.114152813 | 0.031703036 |
| SNP_A.K_708992_4_8212378 | 4 | 8212378 | 1.72E-05 | 0.07292 | 192 | 0.114093391 | 0.031844895 |
| SNP_A.K_705469_4_6566546 | 4 | 6566546 | 1.72E-05 | 0.17448 | 192 | 0.114067991 | 0.031889456 |
| SNP_A.K_693757_4_5059137 | 4 | 5059137 | 1.74E-05 | 0.17448 | 192 | 0.113946028 | 0.032213368 |
| SNP_A.K_253334_1_40883725 | 1 | 40883725 | 1.75E-05 | 0.3151 | 192 | 0.113883954 | 0.032365375 |
| SNP_A.K_725867_4_11875373 | 4 | 11875373 | 1.75E-05 | 0.13281 | 192 | 0.113846271 | 0.03242422 |
| SNP_A.K_1922418_11_24286933 | 11 | 24286933 | 1.75E-05 | 0.02604 | 192 | 0.113844099 | 0.03242422 |
| SNP_A.K_730548_4_12699131 | 4 | 12699131 | 1.76E-05 | 0.21615 | 192 | 0.11382539 | 0.032450157 |
| SNP_A.K_727005_4_12068428 | 4 | 12068428 | 1.76E-05 | 0.06771 | 192 | 0.113781135 | 0.032551054 |
| SNP_A.K_253460_1_40893476 | 1 | 40893476 | 1.77E-05 | 0.3724 | 192 | 0.113749347 | 0.032615601 |
| SNP_A.K_730470_4_12695231 | 4 | 12695231 | 1.77E-05 | 0.21094 | 192 | 0.113726003 | 0.032653741 |
| SNP_A.K_709755_4_8284891 | 4 | 8284891 | 1.77E-05 | 0.07552 | 192 | 0.113716768 | 0.032653741 |
| SNP_A.K_1551942_9_9134079 | 9 | 9134079 | 1.78E-05 | 0.16667 | 192 | 0.113674942 | 0.032681314 |
| SNP_A.K_724287_4_11586157 | 4 | 11586157 | 1.78E-05 | 0.21094 | 192 | 0.113662603 | 0.032681314 |
| SNP_A.K_1812701_11_8463412 | 11 | 8463412 | 1.79E-05 | 0.17188 | 192 | 0.113643615 | 0.032681314 |
| SNP_A.K_1815119_11_8805475 | 11 | 8805475 | 1.79E-05 | 0.16667 | 192 | 0.113616278 | 0.032681314 |
| SNP_A.K_726927_4_12057266 | 4 | 12057266 | 1.79E-05 | 0.21615 | 192 | 0.113605807 | 0.032681314 |
| SNP_A.K_1813274_11_8526818 | 11 | 8526818 | 1.80E-05 | 0.16667 | 192 | 0.113574327 | 0.032681314 |
| SNP_A.K_1551917_9_9132096 | 9 | 9132096 | 1.80E-05 | 0.26823 | 192 | 0.113527433 | 0.032681314 |
| SNP_A.K_708562_4_8061486 | 4 | 8061486 | 1.81E-05 | 0.07552 | 192 | 0.113521923 | 0.032681314 |
| SNP_A.K_709706_4_8282569 | 4 | 8282569 | 1.81E-05 | 0.07813 | 192 | 0.113496496 | 0.032681314 |
| SNP_A.K_1812816_11_8471166 | 11 | 8471166 | 1.81E-05 | 0.17188 | 192 | 0.113496239 | 0.032681314 |
| SNP_A.K_1551884_9_9131043 | 9 | 9131043 | 1.81E-05 | 0.27865 | 192 | 0.113483137 | 0.032681314 |
| SNP_A.K_613984_3_26751295 | 3 | 26751295 | 1.82E-05 | 0.16406 | 192 | 0.113426218 | 0.032681314 |
| SNP_A.K_1814689_11_8769690 | 11 | 8769690 | 1.82E-05 | 0.17188 | 192 | 0.113417904 | 0.032681314 |
| SNP_A.K_1814691_11_8769712 | 11 | 8769712 | 1.82E-05 | 0.17188 | 192 | 0.113416433 | 0.032681314 |
| SNP_A.K_1812430_11_8418154 | 11 | 8418154 | 1.83E-05 | 0.17188 | 192 | 0.113399294 | 0.032681314 |
| SNP_A.K_724244_4_11583169 | 4 | 11583169 | 1.83E-05 | 0.25521 | 192 | 0.113398602 | 0.032681314 |

| | | | | | | | |
|---|---|---|---|---|---|---|---|
| SNP_A.K_1813859_11_8583551 | 11 | 8583551 | 1.83E-05 | 0.17188 | 192 | 0.113373553 | 0.032681314 |
| SNP_A.K_1028785_6_10683483 | 6 | 10683483 | 1.83E-05 | 0.03646 | 192 | 0.113369594 | 0.032681314 |
| SNP_A.K_709550_4_8273583 | 4 | 8273583 | 1.83E-05 | 0.17188 | 192 | 0.113365824 | 0.032681314 |
| SNP_A.K_1813997_11_8594883 | 11 | 8594883 | 1.83E-05 | 0.16667 | 192 | 0.113356991 | 0.032681314 |
| SNP_A.K_1814150_11_8603677 | 11 | 8603677 | 1.83E-05 | 0.16667 | 192 | 0.113356991 | 0.032681314 |
| SNP_A.K_1814271_11_8611080 | 11 | 8611080 | 1.83E-05 | 0.16667 | 192 | 0.113356991 | 0.032681314 |
| SNP_A.K_1813898_11_8586669 | 11 | 8586669 | 1.83E-05 | 0.16667 | 192 | 0.113356991 | 0.032681314 |
| SNP_A.K_1814512_11_8759802 | 11 | 8759802 | 1.83E-05 | 0.16667 | 192 | 0.113356991 | 0.032681314 |
| SNP_A.K_1812466_11_8439678 | 11 | 8439678 | 1.83E-05 | 0.16667 | 192 | 0.113354287 | 0.032681314 |
| SNP_A.K_726002_4_11894055 | 4 | 11894055 | 1.83E-05 | 0.1276 | 192 | 0.113353732 | 0.032681314 |
| SNP_A.K_1814208_11_8607487 | 11 | 8607487 | 1.84E-05 | 0.16667 | 192 | 0.113338007 | 0.032681314 |
| SNP_A.K_1814601_11_8763765 | 11 | 8763765 | 1.84E-05 | 0.16667 | 192 | 0.113331378 | 0.032681314 |
| SNP_A.K_1814686_11_8769295 | 11 | 8769295 | 1.84E-05 | 0.16667 | 192 | 0.113325832 | 0.032681314 |
| SNP_A.K_1812830_11_8472220 | 11 | 8472220 | 1.84E-05 | 0.16667 | 192 | 0.113322298 | 0.032681314 |
| SNP_A.K_253228_1_40876601 | 1 | 40876601 | 1.85E-05 | 0.42708 | 192 | 0.113263018 | 0.032681314 |
| SNP_A.K_708695_4_8086917 | 4 | 8086917 | 1.85E-05 | 0.07031 | 192 | 0.113246762 | 0.032681314 |
| SNP_A.K_708699_4_8087095 | 4 | 8087095 | 1.85E-05 | 0.07031 | 192 | 0.113246762 | 0.032681314 |
| SNP_A.K_708915_4_8124997 | 4 | 8124997 | 1.85E-05 | 0.07031 | 192 | 0.113246762 | 0.032681314 |
| SNP_A.K_708921_4_8125451 | 4 | 8125451 | 1.85E-05 | 0.07031 | 192 | 0.113246762 | 0.032681314 |
| SNP_A.K_709209_4_8239090 | 4 | 8239090 | 1.85E-05 | 0.07031 | 192 | 0.113246762 | 0.032681314 |
| SNP_A.K_709746_4_8284640 | 4 | 8284640 | 1.85E-05 | 0.07031 | 192 | 0.113246762 | 0.032681314 |
| SNP_A.K_708612_4_8064067 | 4 | 8064067 | 1.85E-05 | 0.07031 | 192 | 0.113246762 | 0.032681314 |
| SNP_A.K_708613_4_8064081 | 4 | 8064081 | 1.85E-05 | 0.07031 | 192 | 0.113246762 | 0.032681314 |
| SNP_A.K_708700_4_8087148 | 4 | 8087148 | 1.85E-05 | 0.07031 | 192 | 0.113246762 | 0.032681314 |
| SNP_A.K_708701_4_8087151 | 4 | 8087151 | 1.85E-05 | 0.07031 | 192 | 0.113246762 | 0.032681314 |
| SNP_A.K_708702_4_8087181 | 4 | 8087181 | 1.85E-05 | 0.07031 | 192 | 0.113246762 | 0.032681314 |
| SNP_A.K_708703_4_8087285 | 4 | 8087285 | 1.85E-05 | 0.07031 | 192 | 0.113246762 | 0.032681314 |
| SNP_A.K_708914_4_8124990 | 4 | 8124990 | 1.85E-05 | 0.07031 | 192 | 0.113246762 | 0.032681314 |
| SNP_A.K_709145_4_8234002 | 4 | 8234002 | 1.85E-05 | 0.07031 | 192 | 0.113246762 | 0.032681314 |
| SNP_A.K_709385_4_8263322 | 4 | 8263322 | 1.85E-05 | 0.07031 | 192 | 0.113246762 | 0.032681314 |
| SNP_A.K_709386_4_8263378 | 4 | 8263378 | 1.85E-05 | 0.07031 | 192 | 0.113246762 | 0.032681314 |
| SNP_A.K_709685_4_8281044 | 4 | 8281044 | 1.85E-05 | 0.07031 | 192 | 0.113246762 | 0.032681314 |
| SNP_A.K_1814857_11_8783220 | 11 | 8783220 | 1.85E-05 | 0.17188 | 192 | 0.113232238 | 0.032687535 |
| SNP_A.K_364186_2_16502487 | 2 | 16502487 | 1.85E-05 | 0.08073 | 192 | 0.113225875 | 0.032687535 |
| SNP_A.K_1814881_11_8784076 | 11 | 8784076 | 1.86E-05 | 0.17188 | 192 | 0.113201345 | 0.032712761 |
| SNP_A.K_1812365_11_8411055 | 11 | 8411055 | 1.86E-05 | 0.18229 | 192 | 0.113190009 | 0.032712761 |
| SNP_A.K_253471_1_40894488 | 1 | 40894488 | 1.86E-05 | 0.43229 | 192 | 0.113189222 | 0.032712761 |
| SNP_A.K_708584_4_8062818 | 4 | 8062818 | 1.86E-05 | 0.07552 | 192 | 0.113174825 | 0.032713545 |
| SNP_A.K_1813778_11_8576759 | 11 | 8576759 | 1.86E-05 | 0.16667 | 192 | 0.113170258 | 0.032713545 |
| SNP_A.K_1551899_9_9131434 | 9 | 9131434 | 1.87E-05 | 0.27865 | 192 | 0.113160505 | 0.032714776 |
| SNP_A.K_277541_2_2949744 | 2 | 2949744 | 1.88E-05 | 0.04688 | 192 | 0.11308179 | 0.032870253 |
| SNP_A.K_725255_4_11769797 | 4 | 11769797 | 1.89E-05 | 0.11458 | 192 | 0.113033236 | 0.032870253 |
| SNP_A.K_708999_4_8212549 | 4 | 8212549 | 1.89E-05 | 0.06771 | 192 | 0.113015432 | 0.032870253 |
| SNP_A.K_693921_4_5074737 | 4 | 5074737 | 1.89E-05 | 0.2474 | 192 | 0.113011493 | 0.032870253 |
| SNP_A.K_710172_4_8319193 | 4 | 8319193 | 1.90E-05 | 0.08333 | 192 | 0.112955715 | 0.032870253 |
| SNP_A.K_723324_4_11432817 | 4 | 11432817 | 1.90E-05 | 0.20573 | 192 | 0.112935536 | 0.032870253 |
| SNP_A.K_1812035_11_8372022 | 11 | 8372022 | 1.90E-05 | 0.17448 | 192 | 0.112929566 | 0.032870253 |
| SNP_A.K_253161_1_40869751 | 1 | 40869751 | 1.91E-05 | 0.42708 | 192 | 0.112883911 | 0.032870253 |
| SNP_A.K_726003_4_11894144 | 4 | 11894144 | 1.92E-05 | 0.11979 | 192 | 0.112870889 | 0.032870253 |
| SNP_A.K_685281_4_2452547 | 4 | 2452547 | 1.92E-05 | 0.0625 | 192 | 0.112855566 | 0.032870253 |

| | | | | | | | |
|---|---|---|---|---|---|---|---|
| SNP_A.K_253166_1_40870839 | 1 | 40870839 | 1.92E-05 | 0.42188 | 192 | 0.11285321 | 0.032870253 |
| SNP_A.K_1813772_11_8576545 | 11 | 8576545 | 1.92E-05 | 0.16927 | 192 | 0.112847019 | 0.032870253 |
| SNP_A.K_708968_4_8210955 | 4 | 8210955 | 1.93E-05 | 0.1224 | 192 | 0.112783934 | 0.032870253 |
| SNP_A.K_726647_4_12028645 | 4 | 12028645 | 1.93E-05 | 0.07031 | 192 | 0.112773958 | 0.032870253 |
| SNP_A.K_724210_4_11580714 | 4 | 11580714 | 1.94E-05 | 0.11198 | 192 | 0.112738825 | 0.032870253 |
| SNP_A.K_730746_4_12726533 | 4 | 12726533 | 1.94E-05 | 0.22656 | 192 | 0.112738185 | 0.032870253 |
| SNP_A.K_253195_1_40873317 | 1 | 40873317 | 1.94E-05 | 0.43229 | 192 | 0.112733485 | 0.032870253 |
| SNP_A.K_1744592_10_18873245 | 10 | 18873245 | 1.94E-05 | 0.07813 | 192 | 0.112703301 | 0.032870253 |
| SNP_A.K_707035_4_6997500 | 4 | 6997500 | 1.95E-05 | 0.23177 | 192 | 0.112696029 | 0.032870253 |
| SNP_A.K_709574_4_8274366 | 4 | 8274366 | 1.95E-05 | 0.11458 | 192 | 0.112643672 | 0.032870253 |
| SNP_A.K_727119_4_12082758 | 4 | 12082758 | 1.96E-05 | 0.21094 | 192 | 0.112623004 | 0.032870253 |
| SNP_A.K_1552850_9_9222268 | 9 | 9222268 | 1.97E-05 | 0.21094 | 192 | 0.112560925 | 0.032870253 |
| SNP_A.K_709006_4_8212655 | 4 | 8212655 | 1.97E-05 | 0.07292 | 192 | 0.112556218 | 0.032870253 |
| SNP_A.K_709007_4_8212657 | 4 | 8212657 | 1.97E-05 | 0.07292 | 192 | 0.112556218 | 0.032870253 |
| SNP_A.K_1813965_11_8590897 | 11 | 8590897 | 1.97E-05 | 0.17969 | 192 | 0.112535036 | 0.032870253 |
| SNP_A.K_615449_3_26867985 | 3 | 26867985 | 1.98E-05 | 0.17969 | 192 | 0.112486933 | 0.032870253 |
| SNP_A.K_1812714_11_8464100 | 11 | 8464100 | 1.98E-05 | 0.16927 | 192 | 0.112485133 | 0.032870253 |
| SNP_A.K_708891_4_8122090 | 4 | 8122090 | 1.99E-05 | 0.07552 | 192 | 0.112470226 | 0.032870253 |
| SNP_A.K_288569_2_5043799 | 2 | 5043799 | 1.99E-05 | 0.01302 | 192 | 0.112469338 | 0.032870253 |
| SNP_A.K_288586_2_5045547 | 2 | 5045547 | 1.99E-05 | 0.01302 | 192 | 0.112469338 | 0.032870253 |
| SNP_A.K_288620_2_5052058 | 2 | 5052058 | 1.99E-05 | 0.01302 | 192 | 0.112469338 | 0.032870253 |
| SNP_A.K_288621_2_5052061 | 2 | 5052061 | 1.99E-05 | 0.01302 | 192 | 0.112469338 | 0.032870253 |
| SNP_A.K_288631_2_5052285 | 2 | 5052285 | 1.99E-05 | 0.01302 | 192 | 0.112469338 | 0.032870253 |
| SNP_A.K_288666_2_5054896 | 2 | 5054896 | 1.99E-05 | 0.01302 | 192 | 0.112469338 | 0.032870253 |
| SNP_A.K_288668_2_5054968 | 2 | 5054968 | 1.99E-05 | 0.01302 | 192 | 0.112469338 | 0.032870253 |
| SNP_A.K_288670_2_5055022 | 2 | 5055022 | 1.99E-05 | 0.01302 | 192 | 0.112469338 | 0.032870253 |
| SNP_A.K_288671_2_5055034 | 2 | 5055034 | 1.99E-05 | 0.01302 | 192 | 0.112469338 | 0.032870253 |
| SNP_A.K_288676_2_5055206 | 2 | 5055206 | 1.99E-05 | 0.01302 | 192 | 0.112469338 | 0.032870253 |
| SNP_A.K_288680_2_5055487 | 2 | 5055487 | 1.99E-05 | 0.01302 | 192 | 0.112469338 | 0.032870253 |
| SNP_A.K_288689_2_5057033 | 2 | 5057033 | 1.99E-05 | 0.01302 | 192 | 0.112469338 | 0.032870253 |
| SNP_A.K_288690_2_5057099 | 2 | 5057099 | 1.99E-05 | 0.01302 | 192 | 0.112469338 | 0.032870253 |
| SNP_A.K_288707_2_5060504 | 2 | 5060504 | 1.99E-05 | 0.01302 | 192 | 0.112469338 | 0.032870253 |
| SNP_A.K_288711_2_5061280 | 2 | 5061280 | 1.99E-05 | 0.01302 | 192 | 0.112469338 | 0.032870253 |
| SNP_A.K_288716_2_5061728 | 2 | 5061728 | 1.99E-05 | 0.01302 | 192 | 0.112469338 | 0.032870253 |
| SNP_A.K_288576_2_5044298 | 2 | 5044298 | 1.99E-05 | 0.01302 | 192 | 0.112469338 | 0.032870253 |
| SNP_A.K_288587_2_5045562 | 2 | 5045562 | 1.99E-05 | 0.01302 | 192 | 0.112469338 | 0.032870253 |
| SNP_A.K_288588_2_5045773 | 2 | 5045773 | 1.99E-05 | 0.01302 | 192 | 0.112469338 | 0.032870253 |
| SNP_A.K_288592_2_5045945 | 2 | 5045945 | 1.99E-05 | 0.01302 | 192 | 0.112469338 | 0.032870253 |
| SNP_A.K_288601_2_5046597 | 2 | 5046597 | 1.99E-05 | 0.01302 | 192 | 0.112469338 | 0.032870253 |
| SNP_A.K_288609_2_5051563 | 2 | 5051563 | 1.99E-05 | 0.01302 | 192 | 0.112469338 | 0.032870253 |
| SNP_A.K_288610_2_5051601 | 2 | 5051601 | 1.99E-05 | 0.01302 | 192 | 0.112469338 | 0.032870253 |
| SNP_A.K_288612_2_5051623 | 2 | 5051623 | 1.99E-05 | 0.01302 | 192 | 0.112469338 | 0.032870253 |
| SNP_A.K_288614_2_5051672 | 2 | 5051672 | 1.99E-05 | 0.01302 | 192 | 0.112469338 | 0.032870253 |
| SNP_A.K_288624_2_5052122 | 2 | 5052122 | 1.99E-05 | 0.01302 | 192 | 0.112469338 | 0.032870253 |
| SNP_A.K_288625_2_5052136 | 2 | 5052136 | 1.99E-05 | 0.01302 | 192 | 0.112469338 | 0.032870253 |
| SNP_A.K_288626_2_5052163 | 2 | 5052163 | 1.99E-05 | 0.01302 | 192 | 0.112469338 | 0.032870253 |
| SNP_A.K_288627_2_5052180 | 2 | 5052180 | 1.99E-05 | 0.01302 | 192 | 0.112469338 | 0.032870253 |
| SNP_A.K_288634_2_5052472 | 2 | 5052472 | 1.99E-05 | 0.01302 | 192 | 0.112469338 | 0.032870253 |
| SNP_A.K_288635_2_5052511 | 2 | 5052511 | 1.99E-05 | 0.01302 | 192 | 0.112469338 | 0.032870253 |
| SNP_A.K_288637_2_5052644 | 2 | 5052644 | 1.99E-05 | 0.01302 | 192 | 0.112469338 | 0.032870253 |

| | | | | | | | |
|---|---|---|---|---|---|---|---|
| SNP_A.K_288662_2_5054600 | 2 | 5054600 | 1.99E-05 | 0.01302 | 192 | 0.112469338 | 0.032870253 |
| SNP_A.K_288665_2_5054890 | 2 | 5054890 | 1.99E-05 | 0.01302 | 192 | 0.112469338 | 0.032870253 |
| SNP_A.K_288667_2_5054899 | 2 | 5054899 | 1.99E-05 | 0.01302 | 192 | 0.112469338 | 0.032870253 |
| SNP_A.K_288672_2_5055049 | 2 | 5055049 | 1.99E-05 | 0.01302 | 192 | 0.112469338 | 0.032870253 |
| SNP_A.K_288677_2_5055357 | 2 | 5055357 | 1.99E-05 | 0.01302 | 192 | 0.112469338 | 0.032870253 |
| SNP_A.K_288678_2_5055402 | 2 | 5055402 | 1.99E-05 | 0.01302 | 192 | 0.112469338 | 0.032870253 |
| SNP_A.K_288679_2_5055415 | 2 | 5055415 | 1.99E-05 | 0.01302 | 192 | 0.112469338 | 0.032870253 |
| SNP_A.K_288684_2_5056206 | 2 | 5056206 | 1.99E-05 | 0.01302 | 192 | 0.112469338 | 0.032870253 |
| SNP_A.K_288685_2_5056259 | 2 | 5056259 | 1.99E-05 | 0.01302 | 192 | 0.112469338 | 0.032870253 |
| SNP_A.K_1814840_11_8781444 | 11 | 8781444 | 1.99E-05 | 0.16927 | 192 | 0.112424779 | 0.032870253 |
| SNP_A.K_724312_4_11587850 | 4 | 11587850 | 1.99E-05 | 0.1224 | 192 | 0.112420474 | 0.032870253 |
| SNP_A.K_1813944_11_8589401 | 11 | 8589401 | 2.01E-05 | 0.17448 | 192 | 0.112361443 | 0.032870253 |
| SNP_A.K_709204_4_8238794 | 4 | 8238794 | 2.01E-05 | 0.07552 | 192 | 0.112345825 | 0.032870253 |
| SNP_A.K_973805_6_619845 | 6 | 619845 | 2.02E-05 | 0.13542 | 192 | 0.112307009 | 0.032870253 |
| SNP_A.K_1814287_11_8611726 | 11 | 8611726 | 2.02E-05 | 0.16927 | 192 | 0.11229985 | 0.032870253 |
| SNP_A.K_1812807_11_8470329 | 11 | 8470329 | 2.02E-05 | 0.16927 | 192 | 0.112297251 | 0.032870253 |
| SNP_A.K_1814266_11_8611016 | 11 | 8611016 | 2.02E-05 | 0.16927 | 192 | 0.112272073 | 0.032870253 |
| SNP_A.K_709392_4_8264359 | 4 | 8264359 | 2.03E-05 | 0.07552 | 192 | 0.112248912 | 0.032870253 |
| SNP_A.K_709075_4_8217405 | 4 | 8217405 | 2.03E-05 | 0.07292 | 192 | 0.112246371 | 0.032870253 |
| SNP_A.K_1812050_11_8373086 | 11 | 8373086 | 2.03E-05 | 0.16927 | 192 | 0.112241043 | 0.032870253 |
| SNP_A.K_1814319_11_8613260 | 11 | 8613260 | 2.03E-05 | 0.16927 | 192 | 0.112237551 | 0.032870253 |
| SNP_A.K_708378_4_8041577 | 4 | 8041577 | 2.03E-05 | 0.07813 | 192 | 0.112221852 | 0.032870253 |
| SNP_A.K_708950_4_8208164 | 4 | 8208164 | 2.03E-05 | 0.11458 | 192 | 0.112220971 | 0.032870253 |
| SNP_A.K_1812738_11_8464976 | 11 | 8464976 | 2.03E-05 | 0.16927 | 192 | 0.112217565 | 0.032870253 |
| SNP_A.K_1812536_11_8448556 | 11 | 8448556 | 2.03E-05 | 0.16927 | 192 | 0.112215701 | 0.032870253 |
| SNP_A.K_1813684_11_8566024 | 11 | 8566024 | 2.03E-05 | 0.16927 | 192 | 0.112214443 | 0.032870253 |
| SNP_A.K_730515_4_12697672 | 4 | 12697672 | 2.03E-05 | 0.22656 | 192 | 0.1122003 | 0.032870253 |
| SNP_A.K_1815094_11_8803263 | 11 | 8803263 | 2.04E-05 | 0.16927 | 192 | 0.112197206 | 0.032870253 |
| SNP_A.K_1813640_11_8563933 | 11 | 8563933 | 2.04E-05 | 0.16927 | 192 | 0.112171343 | 0.032870253 |
| SNP_A.K_1378502_8_8710992 | 8 | 8710992 | 2.04E-05 | 0.2526 | 192 | 0.112163705 | 0.032870253 |
| SNP_A.K_253515_1_40899218 | 1 | 40899218 | 2.04E-05 | 0.15885 | 192 | 0.11216152 | 0.032870253 |
| SNP_A.K_709361_4_8258822 | 4 | 8258822 | 2.04E-05 | 0.07552 | 192 | 0.11216103 | 0.032870253 |
| SNP_A.K_726199_4_11928183 | 4 | 11928183 | 2.04E-05 | 0.25 | 192 | 0.112144937 | 0.032870253 |
| SNP_A.K_287317_2_4877873 | 2 | 4877873 | 2.06E-05 | 0.01302 | 192 | 0.112069958 | 0.032870253 |
| SNP_A.K_287329_2_4879115 | 2 | 4879115 | 2.06E-05 | 0.01302 | 192 | 0.112069958 | 0.032870253 |
| SNP_A.K_287793_2_4941146 | 2 | 4941146 | 2.06E-05 | 0.01302 | 192 | 0.112069958 | 0.032870253 |
| SNP_A.K_287821_2_4945921 | 2 | 4945921 | 2.06E-05 | 0.01302 | 192 | 0.112069958 | 0.032870253 |
| SNP_A.K_287841_2_4947679 | 2 | 4947679 | 2.06E-05 | 0.01302 | 192 | 0.112069958 | 0.032870253 |
| SNP_A.K_287881_2_4951805 | 2 | 4951805 | 2.06E-05 | 0.01302 | 192 | 0.112069958 | 0.032870253 |
| SNP_A.K_287883_2_4952868 | 2 | 4952868 | 2.06E-05 | 0.01302 | 192 | 0.112069958 | 0.032870253 |
| SNP_A.K_287885_2_4953550 | 2 | 4953550 | 2.06E-05 | 0.01302 | 192 | 0.112069958 | 0.032870253 |
| SNP_A.K_287918_2_4961246 | 2 | 4961246 | 2.06E-05 | 0.01302 | 192 | 0.112069958 | 0.032870253 |
| SNP_A.K_287919_2_4961262 | 2 | 4961262 | 2.06E-05 | 0.01302 | 192 | 0.112069958 | 0.032870253 |
| SNP_A.K_288057_2_4979972 | 2 | 4979972 | 2.06E-05 | 0.01302 | 192 | 0.112069958 | 0.032870253 |
| SNP_A.K_288058_2_4980170 | 2 | 4980170 | 2.06E-05 | 0.01302 | 192 | 0.112069958 | 0.032870253 |
| SNP_A.K_288065_2_4982311 | 2 | 4982311 | 2.06E-05 | 0.01302 | 192 | 0.112069958 | 0.032870253 |
| SNP_A.K_288066_2_4982752 | 2 | 4982752 | 2.06E-05 | 0.01302 | 192 | 0.112069958 | 0.032870253 |
| SNP_A.K_288072_2_4983228 | 2 | 4983228 | 2.06E-05 | 0.01302 | 192 | 0.112069958 | 0.032870253 |
| SNP_A.K_288082_2_4985548 | 2 | 4985548 | 2.06E-05 | 0.01302 | 192 | 0.112069958 | 0.032870253 |
| SNP_A.K_287468_2_4893030 | 2 | 4893030 | 2.06E-05 | 0.01302 | 192 | 0.112069958 | 0.032870253 |

| | | | | | | | |
|---|---|---|---|---|---|---|---|
| SNP_A.K_287784_2_4938963 | 2 | 4938963 | 2.06E-05 | 0.01302 | 192 | 0.112069958 | 0.032870253 |
| SNP_A.K_287794_2_4941196 | 2 | 4941196 | 2.06E-05 | 0.01302 | 192 | 0.112069958 | 0.032870253 |
| SNP_A.K_287825_2_4946275 | 2 | 4946275 | 2.06E-05 | 0.01302 | 192 | 0.112069958 | 0.032870253 |
| SNP_A.K_287878_2_4951756 | 2 | 4951756 | 2.06E-05 | 0.01302 | 192 | 0.112069958 | 0.032870253 |
| SNP_A.K_288049_2_4977951 | 2 | 4977951 | 2.06E-05 | 0.01302 | 192 | 0.112069958 | 0.032870253 |
| SNP_A.K_288074_2_4984256 | 2 | 4984256 | 2.06E-05 | 0.01302 | 192 | 0.112069958 | 0.032870253 |
| SNP_A.K_724654_4_11653634 | 4 | 11653634 | 2.06E-05 | 0.11719 | 192 | 0.112067356 | 0.032870253 |
| SNP_A.K_721632_4_11010253 | 4 | 11010253 | 2.07E-05 | 0.22656 | 192 | 0.112029389 | 0.0329373 |
| SNP_A.K_730456_4_12694654 | 4 | 12694654 | 2.07E-05 | 0.03125 | 192 | 0.112023687 | 0.0329373 |
| SNP_A.K_708720_4_8089085 | 4 | 8089085 | 2.07E-05 | 0.08333 | 192 | 0.112013273 | 0.0329373 |
| SNP_A.K_253452_1_40892280 | 1 | 40892280 | 2.07E-05 | 0.42969 | 192 | 0.112003465 | 0.0329373 |
| SNP_A.K_708707_4_8087954 | 4 | 8087954 | 2.07E-05 | 0.07552 | 192 | 0.112002484 | 0.0329373 |
| SNP_A.K_1812366_11_8411150 | 11 | 8411150 | 2.08E-05 | 0.17969 | 192 | 0.111978392 | 0.032979172 |
| SNP_A.K_708964_4_8209986 | 4 | 8209986 | 2.08E-05 | 0.13021 | 192 | 0.111970602 | 0.032979172 |
| SNP_A.K_685845_4_2517504 | 4 | 2517504 | 2.08E-05 | 0.05729 | 192 | 0.111945832 | 0.032979172 |
| SNP_A.K_685846_4_2517506 | 4 | 2517506 | 2.08E-05 | 0.05729 | 192 | 0.111945832 | 0.032979172 |
| SNP_A.K_708759_4_8111255 | 4 | 8111255 | 2.08E-05 | 0.07552 | 192 | 0.111938493 | 0.032979172 |
| SNP_A.K_253160_1_40869735 | 1 | 40869735 | 2.08E-05 | 0.42448 | 192 | 0.111937851 | 0.032979172 |
| SNP_A.K_253498_1_40898211 | 1 | 40898211 | 2.10E-05 | 0.15885 | 192 | 0.111871091 | 0.033153639 |
| SNP_A.K_708640_4_8069362 | 4 | 8069362 | 2.10E-05 | 0.07552 | 192 | 0.111844244 | 0.033208986 |
| SNP_A.K_724800_4_11682810 | 4 | 11682810 | 2.11E-05 | 0.21354 | 192 | 0.111778886 | 0.033376577 |
| SNP_A.K_1550890_9_9016684 | 9 | 9016684 | 2.11E-05 | 0.26042 | 192 | 0.111771799 | 0.033376577 |
| SNP_A.K_708794_4_8114574 | 4 | 8114574 | 2.13E-05 | 0.07552 | 192 | 0.111709363 | 0.03353796 |
| SNP_A.K_708549_4_8061152 | 4 | 8061152 | 2.13E-05 | 0.07552 | 192 | 0.111701712 | 0.03353796 |
| SNP_A.K_708653_4_8071288 | 4 | 8071288 | 2.14E-05 | 0.07552 | 192 | 0.111656043 | 0.03362587 |
| SNP_A.K_708656_4_8071665 | 4 | 8071665 | 2.14E-05 | 0.07552 | 192 | 0.111656043 | 0.03362587 |
| SNP_A.K_708910_4_8124228 | 4 | 8124228 | 2.14E-05 | 0.07552 | 192 | 0.111628034 | 0.033685706 |
| SNP_A.K_1813549_11_8555046 | 11 | 8555046 | 2.15E-05 | 0.17188 | 192 | 0.111603473 | 0.033687992 |
| SNP_A.K_687432_4_3530754 | 4 | 3530754 | 2.15E-05 | 0.125 | 192 | 0.11160323 | 0.033687992 |
| SNP_A.K_708122_4_7543134 | 4 | 7543134 | 2.15E-05 | 0.21615 | 192 | 0.111602263 | 0.033687992 |
| SNP_A.K_253439_1_40890909 | 1 | 40890909 | 2.15E-05 | 0.23958 | 192 | 0.111585497 | 0.033713714 |
| SNP_A.K_1813718_11_8569792 | 11 | 8569792 | 2.15E-05 | 0.17188 | 192 | 0.111575054 | 0.033720187 |
| SNP_A.K_1812897_11_8477811 | 11 | 8477811 | 2.16E-05 | 0.17188 | 192 | 0.111516681 | 0.033873234 |
| SNP_A.K_709558_4_8273713 | 4 | 8273713 | 2.17E-05 | 0.08073 | 192 | 0.111498865 | 0.033902383 |
| SNP_A.K_725333_4_11794539 | 4 | 11794539 | 2.17E-05 | 0.10938 | 192 | 0.111474009 | 0.033953198 |
| SNP_A.K_1813598_11_8559126 | 11 | 8559126 | 2.18E-05 | 0.17188 | 192 | 0.111458626 | 0.03397498 |
| SNP_A.K_709070_4_8216354 | 4 | 8216354 | 2.19E-05 | 0.07292 | 192 | 0.11140972 | 0.034100066 |
| SNP_A.K_310430_2_7728334 | 2 | 7728334 | 2.19E-05 | 0.375 | 192 | 0.111366893 | 0.034206821 |
| SNP_A.K_1813740_11_8574948 | 11 | 8574948 | 2.20E-05 | 0.17188 | 192 | 0.111348331 | 0.034238677 |
| SNP_A.K_708583_4_8062652 | 4 | 8062652 | 2.20E-05 | 0.07813 | 192 | 0.111321872 | 0.034295082 |
| SNP_A.K_709513_4_8271340 | 4 | 8271340 | 2.20E-05 | 0.07552 | 192 | 0.11131194 | 0.034300262 |
| SNP_A.K_725314_4_11792263 | 4 | 11792263 | 2.21E-05 | 0.11719 | 192 | 0.111277444 | 0.034346102 |
| SNP_A.K_709628_4_8278618 | 4 | 8278618 | 2.21E-05 | 0.22135 | 192 | 0.111275987 | 0.034346102 |
| SNP_A.K_253200_1_40873698 | 1 | 40873698 | 2.21E-05 | 0.43229 | 192 | 0.111272433 | 0.034346102 |
| SNP_A.K_709645_4_8279109 | 4 | 8279109 | 2.23E-05 | 0.07552 | 192 | 0.111207638 | 0.034522355 |
| SNP_A.K_708379_4_8041586 | 4 | 8041586 | 2.23E-05 | 0.07552 | 192 | 0.111170522 | 0.034598498 |
| SNP_A.K_1814636_11_8766498 | 11 | 8766498 | 2.23E-05 | 0.17188 | 192 | 0.111162892 | 0.034598498 |
| SNP_A.K_708722_4_8089099 | 4 | 8089099 | 2.24E-05 | 0.08333 | 192 | 0.111158624 | 0.034598498 |
| SNP_A.K_1814136_11_8602854 | 11 | 8602854 | 2.24E-05 | 0.17188 | 192 | 0.111133141 | 0.034627053 |
| SNP_A.K_1953653_12_3611534 | 12 | 3611534 | 2.24E-05 | 0.08073 | 192 | 0.111128873 | 0.034627053 |

| | | | | | | | |
|---|---|---|---|---|---|---|---|
| SNP_A.K_1814751_11_8772409 | 11 | 8772409 | 2.24E-05 | 0.17188 | 192 | 0.111124894 | 0.034627053 |
| SNP_A.K_1814875_11_8783817 | 11 | 8783817 | 2.24E-05 | 0.17188 | 192 | 0.111115372 | 0.034630977 |
| SNP_A.K_724192_4_11579308 | 4 | 11579308 | 2.25E-05 | 0.11458 | 192 | 0.111082387 | 0.034630977 |
| SNP_A.K_1028960_6_10713119 | 6 | 10713119 | 2.25E-05 | 0.01302 | 192 | 0.111080482 | 0.034630977 |
| SNP_A.K_1028961_6_10714264 | 6 | 10714264 | 2.25E-05 | 0.01302 | 192 | 0.111080482 | 0.034630977 |
| SNP_A.K_1777242_11_2482507 | 11 | 2482507 | 2.25E-05 | 0.1901 | 192 | 0.111075309 | 0.034630977 |
| SNP_A.K_1812020_11_8371130 | 11 | 8371130 | 2.25E-05 | 0.17188 | 192 | 0.111069855 | 0.034630977 |
| SNP_A.K_709752_4_8284859 | 4 | 8284859 | 2.25E-05 | 0.07552 | 192 | 0.111066423 | 0.034630977 |
| SNP_A.K_1812151_11_8383055 | 11 | 8383055 | 2.26E-05 | 0.17448 | 192 | 0.111050128 | 0.034633833 |
| SNP_A.K_709136_4_8233299 | 4 | 8233299 | 2.26E-05 | 0.07552 | 192 | 0.111028546 | 0.034633833 |
| SNP_A.K_1814872_11_8783768 | 11 | 8783768 | 2.26E-05 | 0.17188 | 192 | 0.111026203 | 0.034633833 |
| SNP_A.K_705321_4_6545625 | 4 | 6545625 | 2.26E-05 | 0.17969 | 192 | 0.111025823 | 0.034633833 |
| SNP_A.K_1813804_11_8579976 | 11 | 8579976 | 2.26E-05 | 0.17448 | 192 | 0.111021684 | 0.034633833 |
| SNP_A.K_1814855_11_8783162 | 11 | 8783162 | 2.26E-05 | 0.17188 | 192 | 0.111016704 | 0.034633833 |
| SNP_A.K_726902_4_12055821 | 4 | 12055821 | 2.28E-05 | 0.26042 | 192 | 0.11096522 | 0.034770088 |
| SNP_A.K_1812396_11_8414208 | 11 | 8414208 | 2.29E-05 | 0.18229 | 192 | 0.110910041 | 0.034912428 |
| SNP_A.K_709835_4_8299235 | 4 | 8299235 | 2.29E-05 | 0.05469 | 192 | 0.11090389 | 0.034912428 |
| SNP_A.K_709284_4_8245685 | 4 | 8245685 | 2.30E-05 | 0.22656 | 192 | 0.11085723 | 0.035034554 |
| SNP_A.K_277435_2_2943367 | 2 | 2943367 | 2.30E-05 | 0.08333 | 192 | 0.110841805 | 0.035057838 |
| SNP_A.K_709751_4_8284749 | 4 | 8284749 | 2.30E-05 | 0.07813 | 192 | 0.11083174 | 0.035064133 |
| SNP_A.K_1310472_7_26337252 | 7 | 26337252 | 2.30E-05 | 0.14063 | 192 | 0.110822395 | 0.035068165 |
| SNP_A.K_253124_1_40862588 | 1 | 40862588 | 2.32E-05 | 0.32292 | 192 | 0.110745247 | 0.035288223 |
| SNP_A.K_1551936_9_9133820 | 9 | 9133820 | 2.32E-05 | 0.34896 | 192 | 0.110731255 | 0.035307174 |
| SNP_A.K_307048_2_7375415 | 2 | 7375415 | 2.33E-05 | 0.20052 | 192 | 0.110710578 | 0.035334508 |
| SNP_A.K_613493_3_26711137 | 3 | 26711137 | 2.33E-05 | 0.15885 | 192 | 0.110705173 | 0.035334508 |
| SNP_A.K_723729_4_11498524 | 4 | 11498524 | 2.33E-05 | 0.21875 | 192 | 0.110698558 | 0.035334508 |
| SNP_A.K_1777239_11_2482485 | 11 | 2482485 | 2.33E-05 | 0.1849 | 192 | 0.110678887 | 0.035371751 |
| SNP_A.K_1812559_11_8451697 | 11 | 8451697 | 2.34E-05 | 0.17448 | 192 | 0.110652734 | 0.035429849 |
| SNP_A.K_709692_4_8281511 | 4 | 8281511 | 2.35E-05 | 0.07813 | 192 | 0.110620782 | 0.035506712 |
| SNP_A.K_868729_5_4774933 | 5 | 4774933 | 2.36E-05 | 0.02604 | 192 | 0.110563449 | 0.035618016 |
| SNP_A.K_709736_4_8284382 | 4 | 8284382 | 2.36E-05 | 0.07813 | 192 | 0.110557551 | 0.035618016 |
| SNP_A.K_709398_4_8265000 | 4 | 8265000 | 2.36E-05 | 0.11719 | 192 | 0.110556346 | 0.035618016 |
| SNP_A.K_708547_4_8061100 | 4 | 8061100 | 2.36E-05 | 0.07552 | 192 | 0.110554183 | 0.035618016 |
| SNP_A.K_1813743_11_8575019 | 11 | 8575019 | 2.36E-05 | 0.17969 | 192 | 0.110544405 | 0.035623762 |
| SNP_A.K_708561_4_8061468 | 4 | 8061468 | 2.37E-05 | 0.07552 | 192 | 0.110501462 | 0.035696225 |
| SNP_A.K_708991_4_8212370 | 4 | 8212370 | 2.37E-05 | 0.07813 | 192 | 0.110501173 | 0.035696225 |
| SNP_A.K_708980_4_8212115 | 4 | 8212115 | 2.37E-05 | 0.07552 | 192 | 0.110498019 | 0.035696225 |
| SNP_A.K_157019_1_24522971 | 1 | 24522971 | 2.39E-05 | 0.10677 | 192 | 0.110433177 | 0.035880613 |
| SNP_A.K_708792_4_8114534 | 4 | 8114534 | 2.39E-05 | 0.08073 | 192 | 0.11040425 | 0.035938423 |
| SNP_A.K_709664_4_8279660 | 4 | 8279660 | 2.39E-05 | 0.07031 | 192 | 0.110399481 | 0.035938423 |
| SNP_A.K_708783_4_8113709 | 4 | 8113709 | 2.41E-05 | 0.11458 | 192 | 0.110327942 | 0.036146072 |
| SNP_A.K_708666_4_8072070 | 4 | 8072070 | 2.41E-05 | 0.07813 | 192 | 0.110313057 | 0.036151301 |
| SNP_A.K_1812851_11_8473503 | 11 | 8473503 | 2.42E-05 | 0.17448 | 192 | 0.110300906 | 0.036151301 |
| SNP_A.K_708927_4_8126092 | 4 | 8126092 | 2.42E-05 | 0.08073 | 192 | 0.110297914 | 0.036151301 |
| SNP_A.K_725329_4_11794183 | 4 | 11794183 | 2.42E-05 | 0.11979 | 192 | 0.110294588 | 0.036151301 |
| SNP_A.K_1813270_11_8526576 | 11 | 8526576 | 2.42E-05 | 0.16927 | 192 | 0.110276613 | 0.036184241 |
| SNP_A.K_708939_4_8150815 | 4 | 8150815 | 2.43E-05 | 0.20573 | 192 | 0.110258239 | 0.036218538 |
| SNP_A.K_1552435_9_9176659 | 9 | 9176659 | 2.43E-05 | 0.22917 | 192 | 0.110248333 | 0.03622508 |
| SNP_A.K_724251_4_11583707 | 4 | 11583707 | 2.43E-05 | 0.28385 | 192 | 0.110219219 | 0.036279851 |
| SNP_A.K_708793_4_8114538 | 4 | 8114538 | 2.43E-05 | 0.07813 | 192 | 0.110215853 | 0.036279851 |

| | | | | | | | |
|---|---|---|---|---|---|---|---|
| SNP_A.K_1744588_10_18872954 | 10 | 18872954 | 2.44E-05 | 0.08594 | 192 | 0.110198189 | 0.036291333 |
| SNP_A.K_613333_3_26690871 | 3 | 26690871 | 2.44E-05 | 0.13021 | 192 | 0.110187621 | 0.036291333 |
| SNP_A.K_1697693_10_12770016 | 10 | 12770016 | 2.44E-05 | 0.20052 | 192 | 0.110182978 | 0.036291333 |
| SNP_A.K_709344_4_8250123 | 4 | 8250123 | 2.44E-05 | 0.21875 | 192 | 0.110180813 | 0.036291333 |
| SNP_A.K_1552843_9_9221520 | 9 | 9221520 | 2.45E-05 | 0.29688 | 192 | 0.110160838 | 0.036295289 |
| SNP_A.K_1552067_9_9142806 | 9 | 9142806 | 2.45E-05 | 0.30729 | 192 | 0.110157103 | 0.036295289 |
| SNP_A.K_708628_4_8066752 | 4 | 8066752 | 2.45E-05 | 0.07813 | 192 | 0.110150714 | 0.036295289 |
| SNP_A.K_1552120_9_9148626 | 9 | 9148626 | 2.45E-05 | 0.18229 | 192 | 0.110148153 | 0.036295289 |
| SNP_A.K_1813675_11_8565315 | 11 | 8565315 | 2.45E-05 | 0.15885 | 192 | 0.110137219 | 0.036295404 |
| SNP_A.K_1814634_11_8766328 | 11 | 8766328 | 2.45E-05 | 0.17448 | 192 | 0.110125321 | 0.036295404 |
| SNP_A.K_1813916_11_8587458 | 11 | 8587458 | 2.46E-05 | 0.16927 | 192 | 0.110118989 | 0.036295404 |
| SNP_A.K_1813777_11_8576720 | 11 | 8576720 | 2.46E-05 | 0.17969 | 192 | 0.110116752 | 0.036295404 |
| SNP_A.K_709629_4_8278643 | 4 | 8278643 | 2.46E-05 | 0.07813 | 192 | 0.110095154 | 0.036340736 |
| SNP_A.K_1814685_11_8769206 | 11 | 8769206 | 2.46E-05 | 0.16927 | 192 | 0.110084931 | 0.036348647 |
| SNP_A.K_1812053_11_8373667 | 11 | 8373667 | 2.47E-05 | 0.16927 | 192 | 0.110038491 | 0.03646886 |
| SNP_A.K_253216_1_40874633 | 1 | 40874633 | 2.48E-05 | 0.43229 | 192 | 0.110025586 | 0.03646886 |
| SNP_A.K_1813860_11_8583632 | 11 | 8583632 | 2.48E-05 | 0.18229 | 192 | 0.110025075 | 0.03646886 |
| SNP_A.K_708806_4_8114939 | 4 | 8114939 | 2.48E-05 | 0.08333 | 192 | 0.11000997 | 0.036493023 |
| SNP_A.K_1814344_11_8614167 | 11 | 8614167 | 2.48E-05 | 0.14063 | 192 | 0.109991237 | 0.036508317 |
| SNP_A.K_709433_4_8265906 | 4 | 8265906 | 2.49E-05 | 0.07552 | 192 | 0.109987973 | 0.036508317 |
| SNP_A.K_253211_1_40874522 | 1 | 40874522 | 2.49E-05 | 0.4349 | 192 | 0.109981489 | 0.036508317 |
| SNP_A.K_1814859_11_8783256 | 11 | 8783256 | 2.49E-05 | 0.17448 | 192 | 0.109971127 | 0.036508317 |
| SNP_A.K_253508_1_40898965 | 1 | 40898965 | 2.49E-05 | 0.17448 | 192 | 0.109961676 | 0.036508317 |
| SNP_A.K_727219_4_12095623 | 4 | 12095623 | 2.49E-05 | 0.21354 | 192 | 0.109953346 | 0.036508317 |
| SNP_A.K_730507_4_12697448 | 4 | 12697448 | 2.49E-05 | 0.23438 | 192 | 0.109950913 | 0.036508317 |
| SNP_A.K_1814900_11_8784687 | 11 | 8784687 | 2.50E-05 | 0.17448 | 192 | 0.109911802 | 0.036601778 |
| SNP_A.K_709376_4_8259816 | 4 | 8259816 | 2.50E-05 | 0.07813 | 192 | 0.10990721 | 0.036601778 |
| SNP_A.K_709381_4_8263095 | 4 | 8263095 | 2.51E-05 | 0.07813 | 192 | 0.109881147 | 0.036662625 |
| SNP_A.K_708719_4_8089008 | 4 | 8089008 | 2.52E-05 | 0.1224 | 192 | 0.10984436 | 0.036742047 |
| SNP_A.K_708610_4_8064025 | 4 | 8064025 | 2.52E-05 | 0.07552 | 192 | 0.109841812 | 0.036742047 |
| SNP_A.K_709088_4_8221435 | 4 | 8221435 | 2.52E-05 | 0.07552 | 192 | 0.109828294 | 0.03676135 |
| SNP_A.K_708942_4_8150941 | 4 | 8150941 | 2.55E-05 | 0.21354 | 192 | 0.109722586 | 0.037089454 |
| SNP_A.K_1028147_6_10567378 | 6 | 10567378 | 2.55E-05 | 0.03906 | 192 | 0.109708174 | 0.03711199 |
| SNP_A.K_730525_4_12698095 | 4 | 12698095 | 2.56E-05 | 0.22917 | 192 | 0.109678878 | 0.03718472 |
| SNP_A.K_709387_4_8263723 | 4 | 8263723 | 2.56E-05 | 0.07292 | 192 | 0.109654611 | 0.037240623 |
| SNP_A.K_685893_4_2518826 | 4 | 2518826 | 2.59E-05 | 0.05469 | 192 | 0.109529548 | 0.037573968 |
| SNP_A.K_1815435_11_8851897 | 11 | 8851897 | 2.59E-05 | 0.14583 | 192 | 0.109522105 | 0.037573968 |
| SNP_A.K_708659_4_8071880 | 4 | 8071880 | 2.60E-05 | 0.07292 | 192 | 0.109498138 | 0.037573968 |
| SNP_A.K_709040_4_8214465 | 4 | 8214465 | 2.60E-05 | 0.07292 | 192 | 0.109498138 | 0.037573968 |
| SNP_A.K_709743_4_8284612 | 4 | 8284612 | 2.60E-05 | 0.07292 | 192 | 0.109498138 | 0.037573968 |
| SNP_A.K_709744_4_8284627 | 4 | 8284627 | 2.60E-05 | 0.07292 | 192 | 0.109498138 | 0.037573968 |
| SNP_A.K_709745_4_8284636 | 4 | 8284636 | 2.60E-05 | 0.07292 | 192 | 0.109498138 | 0.037573968 |
| SNP_A.K_1552032_9_9140140 | 9 | 9140140 | 2.60E-05 | 0.32813 | 192 | 0.109494948 | 0.037573968 |
| SNP_A.K_842261_5_810097 | 5 | 810097 | 2.61E-05 | 0.11198 | 192 | 0.109457746 | 0.037674824 |
| SNP_A.K_1814317_11_8613137 | 11 | 8613137 | 2.61E-05 | 0.17708 | 192 | 0.109440674 | 0.037707054 |
| SNP_A.K_709818_4_8298376 | 4 | 8298376 | 2.62E-05 | 0.07292 | 192 | 0.109427465 | 0.037726104 |
| SNP_A.K_1812054_11_8373713 | 11 | 8373713 | 2.62E-05 | 0.17708 | 192 | 0.109389047 | 0.037818687 |
| SNP_A.K_725288_4_11781692 | 4 | 11781692 | 2.63E-05 | 0.21875 | 192 | 0.109384311 | 0.037818687 |
| SNP_A.K_1812393_11_8414058 | 11 | 8414058 | 2.63E-05 | 0.1901 | 192 | 0.109377558 | 0.037818687 |
| SNP_A.K_1175357_7_1866178 | 7 | 1866178 | 2.64E-05 | 0.1224 | 192 | 0.109311206 | 0.037994507 |

| | | | | | | | |
|---|---|---|---|---|---|---|---|
| SNP_A.K_1175358_7_1866184 | 7 | 1866184 | 2.64E-05 | 0.1224 | 192 | 0.109311206 | 0.037994507 |
| SNP_A.K_710517_4_8425693 | 4 | 8425693 | 2.65E-05 | 0.11719 | 192 | 0.109271516 | 0.038105274 |
| SNP_A.K_374318_2_17805532 | 2 | 17805532 | 2.66E-05 | 0.06771 | 192 | 0.109257919 | 0.038105985 |
| SNP_A.K_709343_4_8250121 | 4 | 8250121 | 2.66E-05 | 0.21875 | 192 | 0.10925611 | 0.038105985 |
| SNP_A.K_1552126_9_9149371 | 9 | 9149371 | 2.66E-05 | 0.29948 | 192 | 0.109239134 | 0.038138449 |
| SNP_A.K_708920_4_8125431 | 4 | 8125431 | 2.67E-05 | 0.07292 | 192 | 0.109209949 | 0.038172025 |
| SNP_A.K_708919_4_8125425 | 4 | 8125425 | 2.67E-05 | 0.07292 | 192 | 0.109209949 | 0.038172025 |
| SNP_A.K_1552101_9_9147266 | 9 | 9147266 | 2.67E-05 | 0.17969 | 192 | 0.109199431 | 0.038172025 |
| SNP_A.K_1813888_11_8585907 | 11 | 8585907 | 2.67E-05 | 0.18229 | 192 | 0.109199122 | 0.038172025 |
| SNP_A.K_1552906_9_9224418 | 9 | 9224418 | 2.67E-05 | 0.25521 | 192 | 0.109182171 | 0.038204555 |
| SNP_A.K_708985_4_8212224 | 4 | 8212224 | 2.68E-05 | 0.07292 | 192 | 0.109151154 | 0.038233831 |
| SNP_A.K_708614_4_8064118 | 4 | 8064118 | 2.69E-05 | 0.07292 | 192 | 0.10910844 | 0.038233831 |
| SNP_A.K_1814016_11_8595437 | 11 | 8595437 | 2.69E-05 | 0.17188 | 192 | 0.10910228 | 0.038233831 |
| SNP_A.K_962070_5_28029286 | 5 | 28029286 | 2.70E-05 | 0.14323 | 192 | 0.1090927 | 0.038233831 |
| SNP_A.K_1813954_11_8590096 | 11 | 8590096 | 2.70E-05 | 0.17708 | 192 | 0.109083509 | 0.038233831 |
| SNP_A.K_725299_4_11790658 | 4 | 11790658 | 2.70E-05 | 0.21615 | 192 | 0.109073446 | 0.038233831 |
| SNP_A.K_1814735_11_8771913 | 11 | 8771913 | 2.70E-05 | 0.17188 | 192 | 0.109064506 | 0.038233831 |
| SNP_A.K_1814842_11_8782027 | 11 | 8782027 | 2.70E-05 | 0.17188 | 192 | 0.109058324 | 0.038233831 |
| SNP_A.K_1814844_11_8782049 | 11 | 8782049 | 2.70E-05 | 0.17188 | 192 | 0.109058324 | 0.038233831 |
| SNP_A.K_1814905_11_8784922 | 11 | 8784922 | 2.70E-05 | 0.17188 | 192 | 0.109058324 | 0.038233831 |
| SNP_A.K_1814682_11_8768983 | 11 | 8768983 | 2.71E-05 | 0.17708 | 192 | 0.109021921 | 0.038233831 |
| SNP_A.K_725335_4_11794567 | 4 | 11794567 | 2.71E-05 | 0.21615 | 192 | 0.109015621 | 0.038233831 |
| SNP_A.K_1551924_9_9132727 | 9 | 9132727 | 2.72E-05 | 0.33854 | 192 | 0.109011813 | 0.038233831 |
| SNP_A.K_708986_4_8212252 | 4 | 8212252 | 2.72E-05 | 0.07292 | 192 | 0.109010349 | 0.038233831 |
| SNP_A.K_709146_4_8234031 | 4 | 8234031 | 2.72E-05 | 0.07292 | 192 | 0.109010349 | 0.038233831 |
| SNP_A.K_709621_4_8278416 | 4 | 8278416 | 2.72E-05 | 0.07292 | 192 | 0.109010349 | 0.038233831 |
| SNP_A.K_709670_4_8280243 | 4 | 8280243 | 2.72E-05 | 0.07292 | 192 | 0.109010349 | 0.038233831 |
| SNP_A.K_709147_4_8234036 | 4 | 8234036 | 2.72E-05 | 0.07292 | 192 | 0.109010349 | 0.038233831 |
| SNP_A.K_709151_4_8234170 | 4 | 8234170 | 2.72E-05 | 0.07292 | 192 | 0.109010349 | 0.038233831 |
| SNP_A.K_709747_4_8284684 | 4 | 8284684 | 2.72E-05 | 0.07292 | 192 | 0.109010349 | 0.038233831 |
| SNP_A.K_1813428_11_8538044 | 11 | 8538044 | 2.72E-05 | 0.17708 | 192 | 0.109006597 | 0.038233831 |
| SNP_A.K_363021_2_16391839 | 2 | 16391839 | 2.72E-05 | 0.11198 | 192 | 0.109003826 | 0.038233831 |
| SNP_A.K_709684_4_8281014 | 4 | 8281014 | 2.72E-05 | 0.07292 | 192 | 0.108999341 | 0.038233831 |
| SNP_A.K_1813835_11_8582501 | 11 | 8582501 | 2.72E-05 | 0.17188 | 192 | 0.108993729 | 0.038233831 |
| SNP_A.K_725356_4_11797372 | 4 | 11797372 | 2.73E-05 | 0.11198 | 192 | 0.108972952 | 0.038280166 |
| SNP_A.K_708988_4_8212323 | 4 | 8212323 | 2.73E-05 | 0.07292 | 192 | 0.108953781 | 0.038295164 |
| SNP_A.K_709211_4_8239305 | 4 | 8239305 | 2.73E-05 | 0.07292 | 192 | 0.108953781 | 0.038295164 |
| SNP_A.K_708609_4_8063995 | 4 | 8063995 | 2.73E-05 | 0.07292 | 192 | 0.108934574 | 0.038336166 |
| SNP_A.K_708585_4_8062823 | 4 | 8062823 | 2.74E-05 | 0.07813 | 192 | 0.108919493 | 0.038348138 |
| SNP_A.K_253393_1_40887243 | 1 | 40887243 | 2.74E-05 | 0.1224 | 192 | 0.108916309 | 0.038348138 |
| SNP_A.K_687588_4_3566285 | 4 | 3566285 | 2.76E-05 | 0.15104 | 192 | 0.108845865 | 0.038542509 |
| SNP_A.K_1310467_7_26336854 | 7 | 26336854 | 2.76E-05 | 0.13542 | 192 | 0.10884582 | 0.038542509 |
| SNP_A.K_709354_4_8258458 | 4 | 8258458 | 2.77E-05 | 0.07292 | 192 | 0.108783365 | 0.038735683 |
| SNP_A.K_708997_4_8212485 | 4 | 8212485 | 2.79E-05 | 0.07031 | 192 | 0.108733062 | 0.038886928 |
| SNP_A.K_1813110_11_8515527 | 11 | 8515527 | 2.79E-05 | 0.17708 | 192 | 0.108724538 | 0.038890957 |
| SNP_A.K_708901_4_8122854 | 4 | 8122854 | 2.79E-05 | 0.07292 | 192 | 0.108715235 | 0.038897756 |
| SNP_A.K_1815430_11_8851744 | 11 | 8851744 | 2.80E-05 | 0.15885 | 192 | 0.10869366 | 0.038947958 |
| SNP_A.K_708831_4_8115405 | 4 | 8115405 | 2.80E-05 | 0.07813 | 192 | 0.10865817 | 0.039047555 |
| SNP_A.K_709019_4_8213062 | 4 | 8213062 | 2.81E-05 | 0.07031 | 192 | 0.108641981 | 0.039055514 |
| SNP_A.K_1552854_9_9222428 | 9 | 9222428 | 2.81E-05 | 0.21875 | 192 | 0.108641206 | 0.039055514 |

| | | | | | | | |
|---|---|---|---|---|---|---|---|
| SNP_A.K_708615_4_8064167 | 4 | 8064167 | 2.82E-05 | 0.07292 | 192 | 0.108613967 | 0.039126116 |
| SNP_A.K_708622_4_8065092 | 4 | 8065092 | 2.82E-05 | 0.07031 | 192 | 0.108594153 | 0.039170443 |
| SNP_A.K_709650_4_8279251 | 4 | 8279251 | 2.82E-05 | 0.07813 | 192 | 0.108578649 | 0.039179011 |
| SNP_A.K_1814250_11_8610181 | 11 | 8610181 | 2.83E-05 | 0.17708 | 192 | 0.108577066 | 0.039179011 |
| SNP_A.K_443758_2_29266078 | 2 | 29266078 | 2.83E-05 | 0.3125 | 192 | 0.108553182 | 0.039237955 |
| SNP_A.K_1812474_11_8441962 | 11 | 8441962 | 2.85E-05 | 0.17448 | 192 | 0.108475423 | 0.039489799 |
| SNP_A.K_709362_4_8258837 | 4 | 8258837 | 2.87E-05 | 0.07813 | 192 | 0.108420056 | 0.039662537 |
| SNP_A.K_709617_4_8278341 | 4 | 8278341 | 2.89E-05 | 0.07552 | 192 | 0.108324262 | 0.03996907 |
| SNP_A.K_723287_4_11429138 | 4 | 11429138 | 2.89E-05 | 0.20313 | 192 | 0.108320692 | 0.03996907 |
| SNP_A.K_1777314_11_2486044 | 11 | 2486044 | 2.92E-05 | 0.17448 | 192 | 0.108231324 | 0.040268223 |
| SNP_A.K_1552851_9_9222334 | 9 | 9222334 | 2.92E-05 | 0.26563 | 192 | 0.108218253 | 0.040289349 |
| SNP_A.K_1813800_11_8579307 | 11 | 8579307 | 2.93E-05 | 0.17448 | 192 | 0.108192087 | 0.040358503 |
| SNP_A.K_708552_4_8061203 | 4 | 8061203 | 2.94E-05 | 0.07813 | 192 | 0.108155191 | 0.040467237 |
| SNP_A.K_1815421_11_8851526 | 11 | 8851526 | 2.94E-05 | 0.15885 | 192 | 0.10812427 | 0.040554262 |
| SNP_A.K_1814756_11_8772488 | 11 | 8772488 | 2.95E-05 | 0.17969 | 192 | 0.108096588 | 0.04061577 |
| SNP_A.K_1814707_11_8770764 | 11 | 8770764 | 2.95E-05 | 0.17969 | 192 | 0.108093042 | 0.04061577 |
| SNP_A.K_1813842_11_8583219 | 11 | 8583219 | 2.96E-05 | 0.17448 | 192 | 0.108074394 | 0.040640484 |
| SNP_A.K_708760_4_8111267 | 4 | 8111267 | 2.96E-05 | 0.07813 | 192 | 0.108069264 | 0.040640484 |
| SNP_A.K_708998_4_8212500 | 4 | 8212500 | 2.96E-05 | 0.07292 | 192 | 0.108064555 | 0.040640484 |
| SNP_A.K_1814734_11_8771851 | 11 | 8771851 | 2.96E-05 | 0.17448 | 192 | 0.108045645 | 0.040659983 |
| SNP_A.K_708855_4_8117540 | 4 | 8117540 | 2.97E-05 | 0.07813 | 192 | 0.108039614 | 0.040659983 |
| SNP_A.K_1814647_11_8767206 | 11 | 8767206 | 2.97E-05 | 0.17969 | 192 | 0.108037528 | 0.040659983 |
| SNP_A.K_1814649_11_8767299 | 11 | 8767299 | 2.97E-05 | 0.17969 | 192 | 0.108014168 | 0.040694567 |
| SNP_A.K_708726_4_8089178 | 4 | 8089178 | 2.97E-05 | 0.07813 | 192 | 0.108010027 | 0.040694567 |
| SNP_A.K_1813951_11_8589683 | 11 | 8589683 | 2.98E-05 | 0.17969 | 192 | 0.108006469 | 0.040694567 |
| SNP_A.K_1812397_11_8414220 | 11 | 8414220 | 2.99E-05 | 0.1849 | 192 | 0.107971221 | 0.040798363 |
| SNP_A.K_711566_4_8637122 | 4 | 8637122 | 2.99E-05 | 0.20573 | 192 | 0.107942549 | 0.040856559 |
| SNP_A.K_1813441_11_8538578 | 11 | 8538578 | 2.99E-05 | 0.17969 | 192 | 0.107937018 | 0.040856559 |
| SNP_A.K_253039_1_40848183 | 1 | 40848183 | 3.00E-05 | 0.2526 | 192 | 0.107928449 | 0.040856559 |
| SNP_A.K_708865_4_8119500 | 4 | 8119500 | 3.00E-05 | 0.07813 | 192 | 0.107911664 | 0.040856559 |
| SNP_A.K_1814887_11_8784151 | 11 | 8784151 | 3.01E-05 | 0.17448 | 192 | 0.10789386 | 0.040856559 |
| SNP_A.K_1814888_11_8784160 | 11 | 8784160 | 3.01E-05 | 0.17448 | 192 | 0.10789386 | 0.040856559 |
| SNP_A.K_1814861_11_8783356 | 11 | 8783356 | 3.01E-05 | 0.17448 | 192 | 0.10789386 | 0.040856559 |
| SNP_A.K_970962_6_162154 | 6 | 162154 | 3.01E-05 | 0.27083 | 192 | 0.107891542 | 0.040856559 |
| SNP_A.K_709715_4_8283044 | 4 | 8283044 | 3.01E-05 | 0.07552 | 192 | 0.107884927 | 0.040856559 |
| SNP_A.K_1814849_11_8782798 | 11 | 8782798 | 3.01E-05 | 0.17448 | 192 | 0.107883552 | 0.040856559 |
| SNP_A.K_1814837_11_8780957 | 11 | 8780957 | 3.01E-05 | 0.17448 | 192 | 0.107868577 | 0.040885553 |
| SNP_A.K_739791_4_15302574 | 4 | 15302574 | 3.02E-05 | 0.04948 | 192 | 0.107835808 | 0.040980834 |
| SNP_A.K_724245_4_11583196 | 4 | 11583196 | 3.03E-05 | 0.26042 | 192 | 0.107823036 | 0.041001739 |
| SNP_A.K_1552855_9_9222436 | 9 | 9222436 | 3.03E-05 | 0.21354 | 192 | 0.107798918 | 0.041013253 |
| SNP_A.K_1362713_8_5699051 | 8 | 5699051 | 3.04E-05 | 0.30208 | 192 | 0.107787264 | 0.041013253 |
| SNP_A.K_1814045_11_8597154 | 11 | 8597154 | 3.04E-05 | 0.17708 | 192 | 0.107780316 | 0.041013253 |
| SNP_A.K_708660_4_8071891 | 4 | 8071891 | 3.04E-05 | 0.07552 | 192 | 0.107776355 | 0.041013253 |
| SNP_A.K_613492_3_26711110 | 3 | 26711110 | 3.04E-05 | 0.16406 | 192 | 0.107774998 | 0.041013253 |
| SNP_A.K_1744711_10_18881890 | 10 | 18881890 | 3.04E-05 | 0.08594 | 192 | 0.107770086 | 0.041013253 |
| SNP_A.K_1551761_9_9124839 | 9 | 9124839 | 3.04E-05 | 0.32813 | 192 | 0.107769942 | 0.041013253 |
| SNP_A.K_709252_4_8242969 | 4 | 8242969 | 3.05E-05 | 0.22135 | 192 | 0.107750714 | 0.041028641 |
| SNP_A.K_119163_1_17040583 | 1 | 17040583 | 3.05E-05 | 0.02344 | 192 | 0.107750045 | 0.041028641 |
| SNP_A.K_1814885_11_8784112 | 11 | 8784112 | 3.05E-05 | 0.17448 | 192 | 0.107744455 | 0.041028641 |
| SNP_A.K_709089_4_8221477 | 4 | 8221477 | 3.06E-05 | 0.07552 | 192 | 0.107706222 | 0.041143405 |

| | | | | | | | |
|---|---|---|---|---|---|---|---|
| SNP_A.K_1552967_9_9227581 | 9 | 9227581 | 3.06E-05 | 0.23698 | 192 | 0.107699522 | 0.041143405 |
| SNP_A.K_1813185_11_8521615 | 11 | 8521615 | 3.07E-05 | 0.17969 | 192 | 0.107673766 | 0.041213262 |
| SNP_A.K_288389_2_5019477 | 2 | 5019477 | 3.07E-05 | 0.01823 | 192 | 0.107654099 | 0.041260403 |
| SNP_A.K_364196_2_16502679 | 2 | 16502679 | 3.08E-05 | 0.02083 | 192 | 0.107639342 | 0.041289178 |
| SNP_A.K_253121_1_40862453 | 1 | 40862453 | 3.10E-05 | 0.32031 | 192 | 0.107549205 | 0.041602244 |
| SNP_A.K_1550883_9_9016610 | 9 | 9016610 | 3.10E-05 | 0.1224 | 192 | 0.107541525 | 0.041604506 |
| SNP_A.K_740602_4_15920108 | 4 | 15920108 | 3.13E-05 | 0.03385 | 192 | 0.107457175 | 0.041871657 |
| SNP_A.K_708675_4_8084755 | 4 | 8084755 | 3.13E-05 | 0.07813 | 192 | 0.107457012 | 0.041871657 |
| SNP_A.K_709526_4_8271597 | 4 | 8271597 | 3.13E-05 | 0.20573 | 192 | 0.10743769 | 0.041918363 |
| SNP_A.K_1552778_9_9217929 | 9 | 9217929 | 3.15E-05 | 0.13021 | 192 | 0.107390911 | 0.042043096 |
| SNP_A.K_1552552_9_9194726 | 9 | 9194726 | 3.15E-05 | 0.13021 | 192 | 0.107390911 | 0.042043096 |
| SNP_A.K_708678_4_8084838 | 4 | 8084838 | 3.15E-05 | 0.07813 | 192 | 0.107382607 | 0.042047882 |
| SNP_A.K_708649_4_8070826 | 4 | 8070826 | 3.16E-05 | 0.07813 | 192 | 0.107345125 | 0.042164471 |
| SNP_A.K_709223_4_8241661 | 4 | 8241661 | 3.17E-05 | 0.07552 | 192 | 0.107302263 | 0.042283779 |
| SNP_A.K_725936_4_11886583 | 4 | 11886583 | 3.17E-05 | 0.13021 | 192 | 0.107299988 | 0.042283779 |
| SNP_A.K_726982_4_12065488 | 4 | 12065488 | 3.18E-05 | 0.06771 | 192 | 0.107287731 | 0.042303882 |
| SNP_A.K_288698_2_5058182 | 2 | 5058182 | 3.19E-05 | 0.01823 | 192 | 0.107236045 | 0.042451872 |
| SNP_A.K_710107_4_8316101 | 4 | 8316101 | 3.19E-05 | 0.19792 | 192 | 0.107235299 | 0.042451872 |
| SNP_A.K_1812587_11_8453121 | 11 | 8453121 | 3.20E-05 | 0.17708 | 192 | 0.10722066 | 0.04248132 |
| SNP_A.K_1813967_11_8591339 | 11 | 8591339 | 3.22E-05 | 0.17708 | 192 | 0.107124003 | 0.042816419 |
| SNP_A.K_724390_4_11592274 | 4 | 11592274 | 3.23E-05 | 0.26042 | 192 | 0.107120297 | 0.042816419 |
| SNP_A.K_693670_4_5046897 | 4 | 5046897 | 3.23E-05 | 0.13281 | 192 | 0.107108117 | 0.042836592 |
| SNP_A.K_1815414_11_8851157 | 11 | 8851157 | 3.23E-05 | 0.11979 | 192 | 0.107098686 | 0.042846068 |
| SNP_A.K_642926_3_30888585 | 3 | 30888585 | 3.24E-05 | 0.30208 | 192 | 0.107081456 | 0.042885997 |
| SNP_A.K_1814901_11_8784709 | 11 | 8784709 | 3.24E-05 | 0.1875 | 192 | 0.107070221 | 0.042902563 |
| SNP_A.K_1814668_11_8768087 | 11 | 8768087 | 3.25E-05 | 0.18229 | 192 | 0.107051303 | 0.042949178 |
| SNP_A.K_708617_4_8064290 | 4 | 8064290 | 3.25E-05 | 0.07552 | 192 | 0.107041915 | 0.042958584 |
| SNP_A.K_1951052_12_2939047 | 12 | 2939047 | 3.26E-05 | 0.03125 | 192 | 0.107011834 | 0.04302798 |
| SNP_A.K_709654_4_8279460 | 4 | 8279460 | 3.26E-05 | 0.07552 | 192 | 0.10698808 | 0.04302798 |
| SNP_A.K_374577_2_17851726 | 2 | 17851726 | 3.27E-05 | 0.08854 | 192 | 0.106983479 | 0.04302798 |
| SNP_A.K_374578_2_17851732 | 2 | 17851732 | 3.27E-05 | 0.08854 | 192 | 0.106983479 | 0.04302798 |
| SNP_A.K_1812994_11_8485770 | 11 | 8485770 | 3.27E-05 | 0.17708 | 192 | 0.106983333 | 0.04302798 |
| SNP_A.K_710173_4_8319366 | 4 | 8319366 | 3.27E-05 | 0.09115 | 192 | 0.106982274 | 0.04302798 |
| SNP_A.K_724784_4_11680029 | 4 | 11680029 | 3.27E-05 | 0.21615 | 192 | 0.106975434 | 0.04302798 |
| SNP_A.K_709896_4_8300648 | 4 | 8300648 | 3.27E-05 | 0.0651 | 192 | 0.106961804 | 0.043054166 |
| SNP_A.K_1813685_11_8566031 | 11 | 8566031 | 3.28E-05 | 0.1849 | 192 | 0.106949017 | 0.043055277 |
| SNP_A.K_729033_4_12440522 | 4 | 12440522 | 3.28E-05 | 0.27083 | 192 | 0.106941666 | 0.043055277 |
| SNP_A.K_1813722_11_8570967 | 11 | 8570967 | 3.28E-05 | 0.18229 | 192 | 0.106933965 | 0.043055277 |
| SNP_A.K_1814637_11_8766551 | 11 | 8766551 | 3.28E-05 | 0.17708 | 192 | 0.106931321 | 0.043055277 |
| SNP_A.K_1813449_11_8538938 | 11 | 8538938 | 3.28E-05 | 0.17708 | 192 | 0.106920201 | 0.043055277 |
| SNP_A.K_709022_4_8213401 | 4 | 8213401 | 3.29E-05 | 0.07813 | 192 | 0.106910884 | 0.043055277 |
| SNP_A.K_1814656_11_8767580 | 11 | 8767580 | 3.29E-05 | 0.17708 | 192 | 0.106909716 | 0.043055277 |
| SNP_A.K_1953593_12_3604966 | 12 | 3604966 | 3.29E-05 | 0.10938 | 192 | 0.106906112 | 0.043055277 |
| SNP_A.K_1813164_11_8520536 | 11 | 8520536 | 3.31E-05 | 0.18229 | 192 | 0.10683071 | 0.043308707 |
| SNP_A.K_374431_2_17835716 | 2 | 17835716 | 3.31E-05 | 0.07813 | 192 | 0.106827843 | 0.043308707 |
| SNP_A.K_709563_4_8273832 | 4 | 8273832 | 3.32E-05 | 0.36458 | 192 | 0.106817366 | 0.043322824 |
| SNP_A.K_1814883_11_8784097 | 11 | 8784097 | 3.33E-05 | 0.17708 | 192 | 0.106777496 | 0.043442329 |
| SNP_A.K_724815_4_11684980 | 4 | 11684980 | 3.33E-05 | 0.1224 | 192 | 0.106773333 | 0.043442329 |
| SNP_A.K_708898_4_8122627 | 4 | 8122627 | 3.33E-05 | 0.07813 | 192 | 0.106755107 | 0.04348722 |
| SNP_A.K_706801_4_6936589 | 4 | 6936589 | 3.34E-05 | 0.21875 | 192 | 0.106730374 | 0.043507508 |

| | | | | | | | |
|---|---|---|---|---|---|---|---|
| SNP_A.K_709117_4_8228308 | 4 | 8228308 | 3.34E-05 | 0.08073 | 192 | 0.106729148 | 0.043507508 |
| SNP_A.K_687549_4_3564120 | 4 | 3564120 | 3.34E-05 | 0.15104 | 192 | 0.106723277 | 0.043507508 |
| SNP_A.K_624595_3_27970881 | 3 | 27970881 | 3.34E-05 | 0.29167 | 192 | 0.106722496 | 0.043507508 |
| SNP_A.K_687546_4_3563933 | 4 | 3563933 | 3.36E-05 | 0.14844 | 192 | 0.106673146 | 0.043664621 |
| SNP_A.K_726200_4_11928202 | 4 | 11928202 | 3.36E-05 | 0.26042 | 192 | 0.106669195 | 0.043664621 |
| SNP_A.K_725962_4_11889084 | 4 | 11889084 | 3.38E-05 | 0.17188 | 192 | 0.106598611 | 0.043918799 |
| SNP_A.K_624209_3_27934904 | 3 | 27934904 | 3.38E-05 | 0.21615 | 192 | 0.106590744 | 0.043922879 |
| SNP_A.K_613868_3_26743395 | 3 | 26743395 | 3.40E-05 | 0.16667 | 192 | 0.106550683 | 0.044036943 |
| SNP_A.K_708925_4_8125940 | 4 | 8125940 | 3.40E-05 | 0.07292 | 192 | 0.106548588 | 0.044036943 |
| SNP_A.K_709608_4_8277958 | 4 | 8277958 | 3.40E-05 | 0.07813 | 192 | 0.106529735 | 0.044066146 |
| SNP_A.K_709355_4_8258567 | 4 | 8258567 | 3.40E-05 | 0.07813 | 192 | 0.106527646 | 0.044066146 |
| SNP_A.K_710051_4_8312772 | 4 | 8312772 | 3.41E-05 | 0.05729 | 192 | 0.106505101 | 0.044129291 |
| SNP_A.K_1362119_8_5625112 | 8 | 5625112 | 3.42E-05 | 0.25781 | 192 | 0.106466266 | 0.044258171 |
| SNP_A.K_815664_4_31300328 | 4 | 31300328 | 3.43E-05 | 0.09896 | 192 | 0.106439545 | 0.044338496 |
| SNP_A.K_709182_4_8237215 | 4 | 8237215 | 3.45E-05 | 0.07552 | 192 | 0.106386212 | 0.044526806 |
| SNP_A.K_1777289_11_2484957 | 11 | 2484957 | 3.48E-05 | 0.21094 | 192 | 0.106284306 | 0.044914289 |
| SNP_A.K_723920_4_11545967 | 4 | 11545967 | 3.48E-05 | 0.21875 | 192 | 0.106272541 | 0.044934608 |
| SNP_A.K_708631_4_8068448 | 4 | 8068448 | 3.49E-05 | 0.07552 | 192 | 0.106252414 | 0.044980286 |
| SNP_A.K_709614_4_8278208 | 4 | 8278208 | 3.49E-05 | 0.07552 | 192 | 0.106242636 | 0.044980286 |
| SNP_A.K_709641_4_8278999 | 4 | 8278999 | 3.50E-05 | 0.07292 | 192 | 0.106236979 | 0.044980286 |
| SNP_A.K_253178_1_40872462 | 1 | 40872462 | 3.50E-05 | 0.4401 | 192 | 0.106228169 | 0.044980286 |
| SNP_A.K_651397_3_32507406 | 3 | 32507406 | 3.50E-05 | 0.19792 | 192 | 0.106227446 | 0.044980286 |
| SNP_A.K_708239_4_7871281 | 4 | 7871281 | 3.51E-05 | 0.24479 | 192 | 0.106204458 | 0.045021498 |
| SNP_A.K_710184_4_8320602 | 4 | 8320602 | 3.51E-05 | 0.09896 | 192 | 0.106200436 | 0.045021498 |
| SNP_A.K_1813021_11_8487443 | 11 | 8487443 | 3.51E-05 | 0.15885 | 192 | 0.106196818 | 0.045021498 |
| SNP_A.K_1813889_11_8585929 | 11 | 8585929 | 3.51E-05 | 0.1849 | 192 | 0.106190317 | 0.045021498 |
| SNP_A.K_1028784_6_10683469 | 6 | 10683469 | 3.52E-05 | 0.03906 | 192 | 0.106161432 | 0.045075891 |
| SNP_A.K_709475_4_8268650 | 4 | 8268650 | 3.52E-05 | 0.08333 | 192 | 0.106150989 | 0.045075891 |
| SNP_A.K_685847_4_2517527 | 4 | 2517527 | 3.52E-05 | 0.05729 | 192 | 0.10614924 | 0.045075891 |
| SNP_A.K_709360_4_8258818 | 4 | 8258818 | 3.52E-05 | 0.07552 | 192 | 0.106145244 | 0.045075891 |
| SNP_A.K_1777351_11_2490187 | 11 | 2490187 | 3.53E-05 | 0.17969 | 192 | 0.106126685 | 0.045075891 |
| SNP_A.K_709620_4_8278402 | 4 | 8278402 | 3.53E-05 | 0.07552 | 192 | 0.106125535 | 0.045075891 |
| SNP_A.K_722570_4_11224191 | 4 | 11224191 | 3.53E-05 | 0.21875 | 192 | 0.106119724 | 0.045075891 |
| SNP_A.K_724599_4_11643380 | 4 | 11643380 | 3.54E-05 | 0.16146 | 192 | 0.10611281 | 0.045075891 |
| SNP_A.K_1812107_11_8379190 | 11 | 8379190 | 3.54E-05 | 0.19792 | 192 | 0.106105598 | 0.045075891 |
| SNP_A.K_709697_4_8282154 | 4 | 8282154 | 3.54E-05 | 0.08333 | 192 | 0.106104297 | 0.045075891 |
| SNP_A.K_709013_4_8212782 | 4 | 8212782 | 3.54E-05 | 0.07552 | 192 | 0.106102951 | 0.045075891 |
| SNP_A.K_1551802_9_9126780 | 9 | 9126780 | 3.55E-05 | 0.29688 | 192 | 0.106069103 | 0.045178329 |
| SNP_A.K_725233_4_11764122 | 4 | 11764122 | 3.55E-05 | 0.1224 | 192 | 0.106064624 | 0.045178329 |
| SNP_A.K_1777330_11_2487773 | 11 | 2487773 | 3.55E-05 | 0.17969 | 192 | 0.106056337 | 0.045184847 |
| SNP_A.K_708592_4_8063241 | 4 | 8063241 | 3.56E-05 | 0.09115 | 192 | 0.106028179 | 0.045271459 |
| SNP_A.K_708995_4_8212398 | 4 | 8212398 | 3.56E-05 | 0.07813 | 192 | 0.106021932 | 0.045271459 |
| SNP_A.K_709612_4_8278177 | 4 | 8278177 | 3.57E-05 | 0.07552 | 192 | 0.10601353 | 0.045278517 |
| SNP_A.K_1814651_11_8767369 | 11 | 8767369 | 3.57E-05 | 0.17969 | 192 | 0.105999363 | 0.045309399 |
| SNP_A.K_709742_4_8284588 | 4 | 8284588 | 3.57E-05 | 0.07813 | 192 | 0.105992147 | 0.0453116 |
| SNP_A.K_716913_4_9934647 | 4 | 9934647 | 3.58E-05 | 0.15885 | 192 | 0.105972051 | 0.045367054 |
| SNP_A.K_693667_4_5046796 | 4 | 5046796 | 3.58E-05 | 0.14063 | 192 | 0.105964527 | 0.045370565 |
| SNP_A.K_1814889_11_8784206 | 11 | 8784206 | 3.59E-05 | 0.17969 | 192 | 0.10594894 | 0.045407457 |
| SNP_A.K_1813998_11_8594911 | 11 | 8594911 | 3.59E-05 | 0.17448 | 192 | 0.10594118 | 0.045411983 |
| SNP_A.K_708903_4_8123035 | 4 | 8123035 | 3.60E-05 | 0.07813 | 192 | 0.105921631 | 0.045465365 |

| | | | | | | | |
|---|---|---|---|---|---|---|---|
| SNP_A.K_1812936_11_8481733 | 11 | 8481733 | 3.60E-05 | 0.1849 | 192 | 0.105902564 | 0.045516828 |
| SNP_A.K_709033_4_8214040 | 4 | 8214040 | 3.61E-05 | 0.07552 | 192 | 0.105887538 | 0.045551583 |
| SNP_A.K_1027593_6_10431515 | 6 | 10431515 | 3.62E-05 | 0.08594 | 192 | 0.105867129 | 0.045587609 |
| SNP_A.K_1362543_8_5674448 | 8 | 5674448 | 3.62E-05 | 0.25 | 192 | 0.105859992 | 0.045587609 |
| SNP_A.K_1814573_11_8762758 | 11 | 8762758 | 3.62E-05 | 0.14844 | 192 | 0.105845034 | 0.045587609 |
| SNP_A.K_418327_2_24389648 | 2 | 24389648 | 3.62E-05 | 0.13542 | 192 | 0.10584467 | 0.045587609 |
| SNP_A.K_623709_3_27872385 | 3 | 27872385 | 3.62E-05 | 0.20052 | 192 | 0.105843729 | 0.045587609 |
| SNP_A.K_654685_3_33856238 | 3 | 33856238 | 3.63E-05 | 0.28125 | 192 | 0.105837793 | 0.045587609 |
| SNP_A.K_1814221_11_8608175 | 11 | 8608175 | 3.63E-05 | 0.22656 | 192 | 0.105832404 | 0.045587609 |
| SNP_A.K_708683_4_8085306 | 4 | 8085306 | 3.63E-05 | 0.07552 | 192 | 0.105813353 | 0.045638819 |
| SNP_A.K_1488056_8_26996305 | 8 | 26996305 | 3.64E-05 | 0.0625 | 192 | 0.105806853 | 0.045638819 |
| SNP_A.K_708709_4_8088140 | 4 | 8088140 | 3.64E-05 | 0.08073 | 192 | 0.105779703 | 0.045724362 |
| SNP_A.K_1274046_7_20012345 | 7 | 20012345 | 3.65E-05 | 0.07031 | 192 | 0.105767505 | 0.045747673 |
| SNP_A.K_709394_4_8264814 | 4 | 8264814 | 3.65E-05 | 0.07813 | 192 | 0.105757901 | 0.045760186 |
| SNP_A.K_727030_4_12070996 | 4 | 12070996 | 3.66E-05 | 0.22135 | 192 | 0.105747197 | 0.045777312 |
| SNP_A.K_723644_4_11485007 | 4 | 11485007 | 3.67E-05 | 0.21354 | 192 | 0.105695163 | 0.04596738 |
| SNP_A.K_1814335_11_8613970 | 11 | 8613970 | 3.68E-05 | 0.14323 | 192 | 0.105680708 | 0.045981895 |
| SNP_A.K_1813738_11_8574897 | 11 | 8574897 | 3.68E-05 | 0.1849 | 192 | 0.105678521 | 0.045981895 |
| SNP_A.K_730498_4_12696618 | 4 | 12696618 | 3.68E-05 | 0.21354 | 192 | 0.105658571 | 0.046037978 |
| SNP_A.K_709240_4_8242332 | 4 | 8242332 | 3.69E-05 | 0.07552 | 192 | 0.105638612 | 0.046094189 |
| SNP_A.K_709630_4_8278656 | 4 | 8278656 | 3.70E-05 | 0.08073 | 192 | 0.105613903 | 0.046170479 |
| SNP_A.K_710058_4_8313170 | 4 | 8313170 | 3.72E-05 | 0.05729 | 192 | 0.105565553 | 0.046346748 |
| SNP_A.K_708989_4_8212358 | 4 | 8212358 | 3.72E-05 | 0.07813 | 192 | 0.105549012 | 0.04636117 |
| SNP_A.K_708990_4_8212360 | 4 | 8212360 | 3.72E-05 | 0.07813 | 192 | 0.105549012 | 0.04636117 |
| SNP_A.K_1813072_11_8490320 | 11 | 8490320 | 3.73E-05 | 0.17969 | 192 | 0.105510801 | 0.046377726 |
| SNP_A.K_1362679_8_5697697 | 8 | 5697697 | 3.74E-05 | 0.27344 | 192 | 0.105504446 | 0.046377726 |
| SNP_A.K_1488053_8_26996242 | 8 | 26996242 | 3.74E-05 | 0.0625 | 192 | 0.105498361 | 0.046377726 |
| SNP_A.K_725218_4_11763287 | 4 | 11763287 | 3.74E-05 | 0.11719 | 192 | 0.105494558 | 0.046377726 |
| SNP_A.K_1552845_9_9222177 | 9 | 9222177 | 3.74E-05 | 0.22396 | 192 | 0.10549437 | 0.046377726 |
| SNP_A.K_1552372_9_9170885 | 9 | 9170885 | 3.74E-05 | 0.15104 | 192 | 0.105481759 | 0.046377726 |
| SNP_A.K_709012_4_8212726 | 4 | 8212726 | 3.75E-05 | 0.07552 | 192 | 0.105470059 | 0.046377726 |
| SNP_A.K_708862_4_8118733 | 4 | 8118733 | 3.75E-05 | 0.08073 | 192 | 0.105468127 | 0.046377726 |
| SNP_A.K_709748_4_8284701 | 4 | 8284701 | 3.75E-05 | 0.07552 | 192 | 0.105465827 | 0.046377726 |
| SNP_A.K_709671_4_8280257 | 4 | 8280257 | 3.75E-05 | 0.07552 | 192 | 0.105465827 | 0.046377726 |
| SNP_A.K_2044323_12_19674278 | 12 | 19674278 | 3.75E-05 | 0.21094 | 192 | 0.105460529 | 0.046377726 |
| SNP_A.K_726962_4_12061583 | 4 | 12061583 | 3.75E-05 | 0.22656 | 192 | 0.105460142 | 0.046377726 |
| SNP_A.K_1812893_11_8477436 | 11 | 8477436 | 3.75E-05 | 0.17969 | 192 | 0.105460139 | 0.046377726 |
| SNP_A.K_624105_3_27923525 | 3 | 27923525 | 3.76E-05 | 0.20833 | 192 | 0.105450911 | 0.046389234 |
| SNP_A.K_709700_4_8282371 | 4 | 8282371 | 3.77E-05 | 0.07813 | 192 | 0.105420082 | 0.046483701 |
| SNP_A.K_708876_4_8120724 | 4 | 8120724 | 3.77E-05 | 0.07813 | 192 | 0.105409481 | 0.046483701 |
| SNP_A.K_1552693_9_9210864 | 9 | 9210864 | 3.77E-05 | 0.14063 | 192 | 0.105409113 | 0.046483701 |
| SNP_A.K_709582_4_8274798 | 4 | 8274798 | 3.80E-05 | 0.07552 | 192 | 0.105320898 | 0.046799623 |
| SNP_A.K_709583_4_8274808 | 4 | 8274808 | 3.80E-05 | 0.07552 | 192 | 0.105320898 | 0.046799623 |
| SNP_A.K_1815533_11_8854827 | 11 | 8854827 | 3.80E-05 | 0.18229 | 192 | 0.105314145 | 0.046799623 |
| SNP_A.K_708619_4_8064651 | 4 | 8064651 | 3.80E-05 | 0.07552 | 192 | 0.105308929 | 0.046799623 |
| SNP_A.K_709374_4_8259506 | 4 | 8259506 | 3.81E-05 | 0.07552 | 192 | 0.105280709 | 0.046884937 |
| SNP_A.K_709133_4_8233148 | 4 | 8233148 | 3.82E-05 | 0.08073 | 192 | 0.105276022 | 0.046884937 |
| SNP_A.K_709351_4_8258358 | 4 | 8258358 | 3.82E-05 | 0.08073 | 192 | 0.105255344 | 0.046932466 |
| SNP_A.K_709662_4_8279620 | 4 | 8279620 | 3.82E-05 | 0.07552 | 192 | 0.105251986 | 0.046932466 |
| SNP_A.K_725803_4_11866461 | 4 | 11866461 | 3.83E-05 | 0.17969 | 192 | 0.105221156 | 0.047036954 |

| | | | | | | | |
|---|---|---|---|---|---|---|---|
| SNP_A.K_708823_4_8115278 | 4 | 8115278 | 3.84E-05 | 0.07552 | 192 | 0.105206417 | 0.047040378 |
| SNP_A.K_709690_4_8281328 | 4 | 8281328 | 3.84E-05 | 0.07552 | 192 | 0.105193195 | 0.047040378 |
| SNP_A.K_709226_4_8241858 | 4 | 8241858 | 3.84E-05 | 0.07552 | 192 | 0.105192816 | 0.047040378 |
| SNP_A.K_1813429_11_8538050 | 11 | 8538050 | 3.85E-05 | 0.18229 | 192 | 0.105191133 | 0.047040378 |
| SNP_A.K_709576_4_8274572 | 4 | 8274572 | 3.85E-05 | 0.08073 | 192 | 0.105179321 | 0.047040378 |
| SNP_A.K_1813276_11_8526906 | 11 | 8526906 | 3.85E-05 | 0.18229 | 192 | 0.105175463 | 0.047040378 |
| SNP_A.K_709611_4_8278146 | 4 | 8278146 | 3.85E-05 | 0.07552 | 192 | 0.105175249 | 0.047040378 |
| SNP_A.K_1744595_10_18874750 | 10 | 18874750 | 3.86E-05 | 0.08594 | 192 | 0.105160506 | 0.047076089 |
| SNP_A.K_723328_4_11432881 | 4 | 11432881 | 3.87E-05 | 0.21354 | 192 | 0.105112692 | 0.0472438 |
| SNP_A.K_689582_4_4270640 | 4 | 4270640 | 3.87E-05 | 0.07552 | 192 | 0.105108694 | 0.0472438 |
| SNP_A.K_709675_4_8280356 | 4 | 8280356 | 3.88E-05 | 0.07552 | 192 | 0.105093954 | 0.0472797 |
| SNP_A.K_709553_4_8273667 | 4 | 8273667 | 3.89E-05 | 0.08073 | 192 | 0.105074589 | 0.047307662 |
| SNP_A.K_932655_5_20903915 | 5 | 20903915 | 3.89E-05 | 0.22917 | 192 | 0.105069232 | 0.047307662 |
| SNP_A.K_1814658_11_8767714 | 11 | 8767714 | 3.89E-05 | 0.18229 | 192 | 0.105064655 | 0.047307662 |
| SNP_A.K_685800_4_2515817 | 4 | 2515817 | 3.90E-05 | 0.0599 | 192 | 0.105047926 | 0.047307662 |
| SNP_A.K_724878_4_11690680 | 4 | 11690680 | 3.90E-05 | 0.11458 | 192 | 0.105044372 | 0.047307662 |
| SNP_A.K_708560_4_8061430 | 4 | 8061430 | 3.90E-05 | 0.07552 | 192 | 0.10502871 | 0.047307662 |
| SNP_A.K_708693_4_8086575 | 4 | 8086575 | 3.90E-05 | 0.07552 | 192 | 0.105026865 | 0.047307662 |
| SNP_A.K_1777272_11_2484065 | 11 | 2484065 | 3.91E-05 | 0.1849 | 192 | 0.105019454 | 0.047307662 |
| SNP_A.K_932653_5_20903302 | 5 | 20903302 | 3.91E-05 | 0.23177 | 192 | 0.10501228 | 0.047307662 |
| SNP_A.K_709389_4_8264076 | 4 | 8264076 | 3.91E-05 | 0.07552 | 192 | 0.105004536 | 0.047307662 |
| SNP_A.K_710348_4_8337372 | 4 | 8337372 | 3.91E-05 | 0.09115 | 192 | 0.105003121 | 0.047307662 |
| SNP_A.K_1813097_11_8514861 | 11 | 8514861 | 3.92E-05 | 0.1901 | 192 | 0.104990717 | 0.047307662 |
| SNP_A.K_708657_4_8071788 | 4 | 8071788 | 3.92E-05 | 0.07552 | 192 | 0.104980285 | 0.047307662 |
| SNP_A.K_708987_4_8212272 | 4 | 8212272 | 3.92E-05 | 0.07552 | 192 | 0.104980285 | 0.047307662 |
| SNP_A.K_709432_4_8265884 | 4 | 8265884 | 3.92E-05 | 0.07552 | 192 | 0.104980285 | 0.047307662 |
| SNP_A.K_709615_4_8278276 | 4 | 8278276 | 3.92E-05 | 0.07552 | 192 | 0.104980285 | 0.047307662 |
| SNP_A.K_253138_1_40864925 | 1 | 40864925 | 3.92E-05 | 0.43229 | 192 | 0.104978249 | 0.047307662 |
| SNP_A.K_1813395_11_8536289 | 11 | 8536289 | 3.92E-05 | 0.18229 | 192 | 0.104968846 | 0.047307662 |
| SNP_A.K_1777336_11_2488556 | 11 | 2488556 | 3.93E-05 | 0.17708 | 192 | 0.104966279 | 0.047307662 |
| SNP_A.K_1953581_12_3603155 | 12 | 3603155 | 3.93E-05 | 0.11198 | 192 | 0.104954735 | 0.047330134 |
| SNP_A.K_1814860_11_8783303 | 11 | 8783303 | 3.94E-05 | 0.17708 | 192 | 0.104915903 | 0.047388521 |
| SNP_A.K_1814833_11_8780383 | 11 | 8780383 | 3.94E-05 | 0.17708 | 192 | 0.104915903 | 0.047388521 |
| SNP_A.K_1814853_11_8783155 | 11 | 8783155 | 3.94E-05 | 0.17708 | 192 | 0.104915903 | 0.047388521 |
| SNP_A.K_1814897_11_8784559 | 11 | 8784559 | 3.94E-05 | 0.17708 | 192 | 0.104915903 | 0.047388521 |
| SNP_A.K_709616_4_8278295 | 4 | 8278295 | 3.95E-05 | 0.07552 | 192 | 0.104893565 | 0.047457857 |
| SNP_A.K_729005_4_12425668 | 4 | 12425668 | 3.96E-05 | 0.1901 | 192 | 0.10486797 | 0.047541455 |
| SNP_A.K_708732_4_8089798 | 4 | 8089798 | 3.97E-05 | 0.07552 | 192 | 0.104853345 | 0.047577539 |
| SNP_A.K_1552011_9_9138555 | 9 | 9138555 | 3.98E-05 | 0.26563 | 192 | 0.104812487 | 0.047704924 |
| SNP_A.K_1794053_11_5377516 | 11 | 5377516 | 3.98E-05 | 0.04948 | 192 | 0.104803543 | 0.047704924 |
| SNP_A.K_709735_4_8284322 | 4 | 8284322 | 3.99E-05 | 0.07552 | 192 | 0.104799968 | 0.047704924 |
| SNP_A.K_726267_4_11932218 | 4 | 11932218 | 3.99E-05 | 0.21354 | 192 | 0.104798838 | 0.047704924 |
| SNP_A.K_708691_4_8086292 | 4 | 8086292 | 3.99E-05 | 0.07552 | 192 | 0.104780743 | 0.047707943 |
| SNP_A.K_709366_4_8258985 | 4 | 8258985 | 3.99E-05 | 0.07552 | 192 | 0.104779268 | 0.047707943 |
| SNP_A.K_709367_4_8259002 | 4 | 8259002 | 3.99E-05 | 0.07552 | 192 | 0.104779268 | 0.047707943 |
| SNP_A.K_708011_4_7496094 | 4 | 7496094 | 4.00E-05 | 0.23438 | 192 | 0.104760394 | 0.047734434 |
| SNP_A.K_1815433_11_8851795 | 11 | 8851795 | 4.00E-05 | 0.15625 | 192 | 0.104756597 | 0.047734434 |
| SNP_A.K_709680_4_8280834 | 4 | 8280834 | 4.00E-05 | 0.07552 | 192 | 0.104754344 | 0.047734434 |
| SNP_A.K_723682_4_11487985 | 4 | 11487985 | 4.02E-05 | 0.1224 | 192 | 0.104703564 | 0.047928832 |
| SNP_A.K_1813770_11_8576517 | 11 | 8576517 | 4.03E-05 | 0.18229 | 192 | 0.104681625 | 0.047997461 |

| | | | | | | | |
|---|---|---|---|---|---|---|---|
| SNP_A.K_2078742_12_27020700 | 12 | 27020700 | 4.04E-05 | 0.02083 | 192 | 0.104653571 | 0.048093061 |
| SNP_A.K_709618_4_8278356 | 4 | 8278356 | 4.04E-05 | 0.07813 | 192 | 0.104636723 | 0.048139579 |
| SNP_A.K_709643_4_8279064 | 4 | 8279064 | 4.05E-05 | 0.07813 | 192 | 0.104629963 | 0.048141782 |
| SNP_A.K_814121_4_31132534 | 4 | 31132534 | 4.05E-05 | 0.05208 | 192 | 0.104615419 | 0.048169324 |
| SNP_A.K_1812977_11_8484543 | 11 | 8484543 | 4.05E-05 | 0.14583 | 192 | 0.104611195 | 0.048169324 |
| SNP_A.K_709004_4_8212637 | 4 | 8212637 | 4.07E-05 | 0.07292 | 192 | 0.104582389 | 0.048268667 |
| SNP_A.K_709375_4_8259632 | 4 | 8259632 | 4.07E-05 | 0.07552 | 192 | 0.104573489 | 0.048280379 |
| SNP_A.K_1813584_11_8557400 | 11 | 8557400 | 4.08E-05 | 0.17969 | 192 | 0.104530909 | 0.048440901 |
| SNP_A.K_708003_4_7494623 | 4 | 7494623 | 4.09E-05 | 0.23698 | 192 | 0.104513788 | 0.048489088 |
| SNP_A.K_724859_4_11689480 | 4 | 11689480 | 4.10E-05 | 0.19271 | 192 | 0.104480561 | 0.048608804 |
| SNP_A.K_724479_4_11607080 | 4 | 11607080 | 4.11E-05 | 0.29427 | 192 | 0.104460133 | 0.048671893 |
| SNP_A.K_1552266_9_9164047 | 9 | 9164047 | 4.12E-05 | 0.15365 | 192 | 0.104436477 | 0.048730039 |
| SNP_A.K_709098_4_8223160 | 4 | 8223160 | 4.12E-05 | 0.08073 | 192 | 0.10442951 | 0.048730039 |
| SNP_A.K_708801_4_8114788 | 4 | 8114788 | 4.12E-05 | 0.07813 | 192 | 0.1044284 | 0.048730039 |
| SNP_A.K_725252_4_11768581 | 4 | 11768581 | 4.13E-05 | 0.21875 | 192 | 0.104419643 | 0.04874136 |
| SNP_A.K_1763159_10_21646632 | 10 | 21646632 | 4.15E-05 | 0.1875 | 192 | 0.104367877 | 0.048944641 |
| SNP_A.K_708316_4_7902431 | 4 | 7902431 | 4.15E-05 | 0.23177 | 192 | 0.104348654 | 0.048981332 |
| SNP_A.K_710497_4_8424991 | 4 | 8424991 | 4.15E-05 | 0.07813 | 192 | 0.104347265 | 0.048981332 |
| SNP_A.K_290404_2_5328871 | 2 | 5328871 | 4.16E-05 | 0.08594 | 192 | 0.104340848 | 0.048982306 |
| SNP_A.K_1552033_9_9140182 | 9 | 9140182 | 4.18E-05 | 0.33854 | 192 | 0.104288524 | 0.049182201 |
| SNP_A.K_709074_4_8217101 | 4 | 8217101 | 4.18E-05 | 0.08073 | 192 | 0.104283888 | 0.049182201 |
| SNP_A.K_1813626_11_8561415 | 11 | 8561415 | 4.18E-05 | 0.16927 | 192 | 0.104277039 | 0.04918517 |
| SNP_A.K_728153_4_12282417 | 4 | 12282417 | 4.19E-05 | 0.25781 | 192 | 0.104249252 | 0.049282355 |
| SNP_A.K_1814869_11_8783655 | 11 | 8783655 | 4.20E-05 | 0.1849 | 192 | 0.104228932 | 0.049338364 |
| SNP_A.K_725305_4_11791553 | 4 | 11791553 | 4.20E-05 | 0.1224 | 192 | 0.104224463 | 0.049338364 |
| SNP_A.K_253068_1_40850409 | 1 | 40850409 | 4.21E-05 | 0.16406 | 192 | 0.10420404 | 0.049402654 |
| SNP_A.K_709208_4_8239037 | 4 | 8239037 | 4.22E-05 | 0.07813 | 192 | 0.104185856 | 0.049456921 |
| SNP_A.K_1777337_11_2488638 | 11 | 2488638 | 4.22E-05 | 0.17969 | 192 | 0.104171163 | 0.049475335 |
| SNP_A.K_1813180_11_8521517 | 11 | 8521517 | 4.22E-05 | 0.1849 | 192 | 0.104169453 | 0.049475335 |
| SNP_A.K_650908_3_32280631 | 3 | 32280631 | 4.23E-05 | 0.21615 | 192 | 0.104139441 | 0.049583306 |
| SNP_A.K_1814644_11_8767095 | 11 | 8767095 | 4.24E-05 | 0.1849 | 192 | 0.104123067 | 0.049629635 |
| SNP_A.K_1648670_10_4748578 | 10 | 4748578 | 4.24E-05 | 0.05469 | 192 | 0.104114848 | 0.049639009 |

Seed_coat_color

| SNP | Chrc | Position | P.value | maf | nobs | Rsquare.of.Model.w | FDR_Adjusted_P-values |
|---|---|---|---|---|---|---|---|
| SNP_A.K_1188748_7_6124457 | 7 | 6124457 | 2.17E-14 | 0.45652 | 299 | 0.616237178 | 4.52E-08 |
| SNP_A.K_1188775_7_6195128 | 7 | 6195128 | 2.10E-12 | 0.23746 | 299 | 0.598211775 | 1.60E-06 |
| SNP_A.K_1189244_7_6660825 | 7 | 6660825 | 2.31E-12 | 0.45485 | 299 | 0.597852159 | 1.60E-06 |
| SNP_A.K_1189255_7_6663350 | 7 | 6663350 | 6.55E-12 | 0.46154 | 299 | 0.593832819 | 3.18E-06 |
| SNP_A.K_1188749_7_6128379 | 7 | 6128379 | 7.63E-12 | 0.199 | 299 | 0.593244316 | 3.18E-06 |
| SNP_A.K_1189276_7_6665511 | 7 | 6665511 | 1.30E-11 | 0.45318 | 299 | 0.591212513 | 4.50E-06 |
| SNP_A.K_1189246_7_6661169 | 7 | 6661169 | 1.85E-11 | 0.46154 | 299 | 0.589861025 | 4.82E-06 |
| SNP_A.K_1188751_7_6133394 | 7 | 6133394 | 1.85E-11 | 0.3495 | 299 | 0.589853064 | 4.82E-06 |
| SNP_A.K_1189214_7_6658617 | 7 | 6658617 | 4.64E-11 | 0.47492 | 299 | 0.586363754 | 1.07E-05 |
| SNP_A.K_1188921_7_6434432 | 7 | 6434432 | 6.70E-11 | 0.42642 | 299 | 0.584976528 | 1.26E-05 |
| SNP_A.K_1189309_7_6670284 | 7 | 6670284 | 6.72E-11 | 0.46488 | 299 | 0.58496537 | 1.26E-05 |
| SNP_A.K_1189258_7_6663434 | 7 | 6663434 | 7.28E-11 | 0.46154 | 299 | 0.584666378 | 1.26E-05 |
| SNP_A.K_1189226_7_6659524 | 7 | 6659524 | 8.62E-11 | 0.38294 | 299 | 0.584026021 | 1.38E-05 |
| SNP_A.K_1189282_7_6666905 | 7 | 6666905 | 1.41E-10 | 0.45485 | 299 | 0.582181927 | 2.04E-05 |
| SNP_A.K_1188838_7_6361858 | 7 | 6361858 | 1.47E-10 | 0.49331 | 299 | 0.582017513 | 2.04E-05 |
| SNP_A.K_1189339_7_6685920 | 7 | 6685920 | 1.79E-10 | 0.43645 | 299 | 0.581281759 | 2.28E-05 |

| | | | | | | | |
|---|---|---|---|---|---|---|---|
| SNP_A.K_1189250_7_6662451 | 7 | 6662451 | 1.99E-10 | 0.45987 | 299 | 0.580881558 | 2.28E-05 |
| SNP_A.K_1188817_7_6342940 | 7 | 6342940 | 2.05E-10 | 0.49498 | 299 | 0.580781978 | 2.28E-05 |
| SNP_A.K_1189294_7_6668920 | 7 | 6668920 | 2.08E-10 | 0.45987 | 299 | 0.580724401 | 2.28E-05 |
| SNP_A.K_1188773_7_6192768 | 7 | 6192768 | 3.15E-10 | 0.28094 | 299 | 0.579172248 | 3.28E-05 |
| SNP_A.K_1189249_7_6661790 | 7 | 6661790 | 3.42E-10 | 0.45819 | 299 | 0.578860056 | 3.36E-05 |
| SNP_A.K_1189215_7_6658783 | 7 | 6658783 | 3.55E-10 | 0.44482 | 299 | 0.578727326 | 3.36E-05 |
| SNP_A.K_1189285_7_6667624 | 7 | 6667624 | 4.65E-10 | 0.44314 | 299 | 0.577719996 | 4.21E-05 |
| SNP_A.K_1189336_7_6685769 | 7 | 6685769 | 7.00E-10 | 0.45819 | 299 | 0.576201084 | 6.07E-05 |
| SNP_A.K_1189245_7_6660873 | 7 | 6660873 | 7.52E-10 | 0.44649 | 299 | 0.575936089 | 6.26E-05 |
| SNP_A.K_1188742_7_6062744 | 7 | 6062744 | 7.95E-10 | 0.4398 | 299 | 0.575730482 | 6.36E-05 |
| SNP_A.K_1189239_7_6660353 | 7 | 6660353 | 8.50E-10 | 0.45485 | 299 | 0.575482821 | 6.55E-05 |
| SNP_A.K_1188769_7_6185067 | 7 | 6185067 | 9.10E-10 | 0.40635 | 299 | 0.575227013 | 6.77E-05 |
| SNP_A.K_1189280_7_6666646 | 7 | 6666646 | 1.03E-09 | 0.45485 | 299 | 0.574761275 | 7.41E-05 |
| SNP_A.K_1188764_7_6170573 | 7 | 6170573 | 1.15E-09 | 0.41973 | 299 | 0.574357637 | 7.99E-05 |
| SNP_A.K_1189216_7_6658844 | 7 | 6658844 | 1.24E-09 | 0.4398 | 299 | 0.574097256 | 8.30E-05 |
| SNP_A.K_1188816_7_6341838 | 7 | 6341838 | 1.57E-09 | 0.48662 | 299 | 0.573218726 | 9.77E-05 |
| SNP_A.K_1189213_7_6658535 | 7 | 6658535 | 1.60E-09 | 0.45151 | 299 | 0.573148832 | 9.77E-05 |
| SNP_A.K_1189287_7_6667699 | 7 | 6667699 | 1.61E-09 | 0.44983 | 299 | 0.573120591 | 9.77E-05 |
| SNP_A.K_1188866_7_6416752 | 7 | 6416752 | 1.67E-09 | 0.43144 | 299 | 0.572982216 | 9.77E-05 |
| SNP_A.K_1189234_7_6659826 | 7 | 6659826 | 1.69E-09 | 0.46154 | 299 | 0.572945073 | 9.77E-05 |
| SNP_A.K_1189305_7_6670198 | 7 | 6670198 | 1.94E-09 | 0.44816 | 299 | 0.572433882 | 0.000109214 |
| SNP_A.K_1188778_7_6204740 | 7 | 6204740 | 2.10E-09 | 0.2893 | 299 | 0.572146249 | 0.000114995 |
| SNP_A.K_1188867_7_6417000 | 7 | 6417000 | 3.06E-09 | 0.41304 | 299 | 0.570764127 | 0.0001633 |
| SNP_A.K_1189286_7_6667646 | 7 | 6667646 | 3.59E-09 | 0.49666 | 299 | 0.570182066 | 0.000186652 |
| SNP_A.K_1189201_7_6655953 | 7 | 6655953 | 5.03E-09 | 0.46321 | 299 | 0.568943035 | 0.000255605 |
| SNP_A.K_1189281_7_6666734 | 7 | 6666734 | 5.36E-09 | 0.44816 | 299 | 0.568717635 | 0.000265422 |
| SNP_A.K_1188797_7_6299925 | 7 | 6299925 | 6.51E-09 | 0.49666 | 299 | 0.568008606 | 0.0003113 |
| SNP_A.K_1188695_7_6049723 | 7 | 6049723 | 6.58E-09 | 0.41806 | 299 | 0.567967089 | 0.0003113 |
| SNP_A.K_1188795_7_6292339 | 7 | 6292339 | 9.82E-09 | 0.31605 | 299 | 0.566511298 | 0.000448477 |
| SNP_A.K_1188837_7_6360143 | 7 | 6360143 | 9.91E-09 | 0.49164 | 299 | 0.566478333 | 0.000448477 |
| SNP_A.K_1189292_7_6668810 | 7 | 6668810 | 1.04E-08 | 0.46488 | 299 | 0.566287823 | 0.00046254 |
| SNP_A.K_1189217_7_6658871 | 7 | 6658871 | 1.07E-08 | 0.46823 | 299 | 0.566211911 | 0.00046254 |
| SNP_A.K_1188818_7_6356862 | 7 | 6356862 | 1.10E-08 | 0.49666 | 299 | 0.566116582 | 0.000465169 |
| SNP_A.K_1188801_7_6303351 | 7 | 6303351 | 1.28E-08 | 0.42809 | 299 | 0.565559276 | 0.000531653 |
| SNP_A.K_1189224_7_6659256 | 7 | 6659256 | 1.34E-08 | 0.40803 | 299 | 0.565394775 | 0.000545456 |
| SNP_A.K_1189203_7_6656052 | 7 | 6656052 | 1.63E-08 | 0.42308 | 299 | 0.564675863 | 0.000645922 |
| SNP_A.K_1188785_7_6220391 | 7 | 6220391 | 1.64E-08 | 0.33612 | 299 | 0.564644164 | 0.000645922 |
| SNP_A.K_1188662_7_6040708 | 7 | 6040708 | 2.64E-08 | 0.43478 | 299 | 0.56293053 | 0.001019536 |
| SNP_A.K_1188822_7_6357186 | 7 | 6357186 | 2.85E-08 | 0.48161 | 299 | 0.562664807 | 0.001065557 |
| SNP_A.K_1189221_7_6659045 | 7 | 6659045 | 2.87E-08 | 0.42642 | 299 | 0.562640795 | 0.001065557 |
| SNP_A.K_1189222_7_6659158 | 7 | 6659158 | 3.36E-08 | 0.37291 | 299 | 0.562067864 | 0.001224508 |
| SNP_A.K_1188495_7_6021907 | 7 | 6021907 | 3.41E-08 | 0.3612 | 299 | 0.562014862 | 0.001224508 |
| SNP_A.K_1188850_7_6378583 | 7 | 6378583 | 3.88E-08 | 0.47993 | 299 | 0.561557885 | 0.00136716 |
| SNP_A.K_1188806_7_6309365 | 7 | 6309365 | 3.94E-08 | 0.43813 | 299 | 0.561496625 | 0.001367525 |
| SNP_A.K_1188871_7_6419242 | 7 | 6419242 | 4.35E-08 | 0.25585 | 299 | 0.561141965 | 0.001484938 |
| SNP_A.K_1189231_7_6659699 | 7 | 6659699 | 4.67E-08 | 0.4214 | 299 | 0.560892147 | 0.001566471 |
| SNP_A.K_1189227_7_6659547 | 7 | 6659547 | 5.10E-08 | 0.39465 | 299 | 0.560574554 | 0.001675705 |
| SNP_A.K_1189304_7_6670188 | 7 | 6670188 | 5.15E-08 | 0.42642 | 299 | 0.560537071 | 0.001675705 |
| SNP_A.K_1189302_7_6670108 | 7 | 6670108 | 5.28E-08 | 0.40301 | 299 | 0.560448295 | 0.001691362 |
| SNP_A.K_1189297_7_6668991 | 7 | 6668991 | 6.18E-08 | 0.47659 | 299 | 0.559886855 | 0.001948882 |

| | | | | | | | |
|---|---|---|---|---|---|---|---|
| SNP_A.K_1189273_7_6664854 | 7 | 6664854 | 6.59E-08 | 0.47659 | 299 | 0.559656691 | 0.002047537 |
| SNP_A.K_1188842_7_6368381 | 7 | 6368381 | 7.52E-08 | 0.47324 | 299 | 0.559185778 | 0.002301899 |
| SNP_A.K_1188926_7_6434733 | 7 | 6434733 | 8.21E-08 | 0.34281 | 299 | 0.55887317 | 0.00247635 |
| SNP_A.K_1189284_7_6667552 | 7 | 6667552 | 9.45E-08 | 0.47492 | 299 | 0.558369823 | 0.002811355 |
| SNP_A.K_1188723_7_6057246 | 7 | 6057246 | 1.08E-07 | 0.3913 | 299 | 0.557882603 | 0.003178438 |
| SNP_A.K_1189194_7_6655801 | 7 | 6655801 | 1.15E-07 | 0.46321 | 299 | 0.557683328 | 0.003314962 |
| SNP_A.K_1189240_7_6660546 | 7 | 6660546 | 1.17E-07 | 0.4214 | 299 | 0.557610343 | 0.003337378 |
| SNP_A.K_1188698_7_6049928 | 7 | 6049928 | 1.23E-07 | 0.42308 | 299 | 0.55744269 | 0.003451328 |
| SNP_A.K_1189197_7_6655874 | 7 | 6655874 | 1.33E-07 | 0.44314 | 299 | 0.557146915 | 0.003701053 |
| SNP_A.K_1188874_7_6420399 | 7 | 6420399 | 1.52E-07 | 0.26087 | 299 | 0.556672925 | 0.004174354 |
| SNP_A.K_1189223_7_6659177 | 7 | 6659177 | 1.64E-07 | 0.40635 | 299 | 0.556422354 | 0.004421907 |
| SNP_A.K_1188704_7_6050283 | 7 | 6050283 | 1.74E-07 | 0.41973 | 299 | 0.556213312 | 0.004630521 |
| SNP_A.K_1188405_7_6015206 | 7 | 6015206 | 1.81E-07 | 0.41137 | 299 | 0.556057598 | 0.00471938 |
| SNP_A.K_1189315_7_6671037 | 7 | 6671037 | 1.81E-07 | 0.40635 | 299 | 0.556056338 | 0.00471938 |
| SNP_A.K_1188706_7_6050559 | 7 | 6050559 | 1.92E-07 | 0.42809 | 299 | 0.555854521 | 0.004934588 |
| SNP_A.K_1188626_7_6033981 | 7 | 6033981 | 2.00E-07 | 0.42308 | 299 | 0.555702075 | 0.005089016 |
| SNP_A.K_1188389_7_6012422 | 7 | 6012422 | 2.04E-07 | 0.46823 | 299 | 0.555646995 | 0.005106604 |
| SNP_A.K_1188702_7_6050170 | 7 | 6050170 | 2.08E-07 | 0.42977 | 299 | 0.555569332 | 0.005136578 |
| SNP_A.K_1188758_7_6151209 | 7 | 6151209 | 2.10E-07 | 0.24415 | 299 | 0.555542088 | 0.005136578 |
| SNP_A.K_1843481_11_12777382 | 11 | 12777382 | 2.15E-07 | 0.43645 | 299 | 0.555448265 | 0.005213374 |
| SNP_A.K_1188629_7_6034130 | 7 | 6034130 | 2.18E-07 | 0.42977 | 299 | 0.55539937 | 0.005219636 |
| SNP_A.K_1189006_7_6440865 | 7 | 6440865 | 2.21E-07 | 0.25418 | 299 | 0.555362755 | 0.005219636 |
| SNP_A.K_1188705_7_6050288 | 7 | 6050288 | 2.40E-07 | 0.42809 | 299 | 0.555071991 | 0.005603671 |
| SNP_A.K_1189259_7_6663452 | 7 | 6663452 | 2.54E-07 | 0.41137 | 299 | 0.554869897 | 0.005867814 |
| SNP_A.K_1188620_7_6033330 | 7 | 6033330 | 2.60E-07 | 0.43311 | 299 | 0.554787117 | 0.005941039 |
| SNP_A.K_1189209_7_6657226 | 7 | 6657226 | 2.68E-07 | 0.40635 | 299 | 0.554678264 | 0.006060552 |
| SNP_A.K_1189303_7_6670110 | 7 | 6670110 | 2.84E-07 | 0.38127 | 299 | 0.554476423 | 0.006305549 |
| SNP_A.K_1188429_7_6018267 | 7 | 6018267 | 2.85E-07 | 0.41806 | 299 | 0.554462611 | 0.006305549 |
| SNP_A.K_1188685_7_6046792 | 7 | 6046792 | 3.08E-07 | 0.4214 | 299 | 0.554181424 | 0.006757217 |
| SNP_A.K_1188590_7_6031361 | 7 | 6031361 | 3.38E-07 | 0.41806 | 299 | 0.55386246 | 0.007320574 |
| SNP_A.K_1188656_7_6038256 | 7 | 6038256 | 3.48E-07 | 0.42475 | 299 | 0.553754116 | 0.007471529 |
| SNP_A.K_1189141_7_6482231 | 7 | 6482231 | 3.68E-07 | 0.21906 | 299 | 0.553561486 | 0.007740499 |
| SNP_A.K_1188611_7_6032549 | 7 | 6032549 | 3.68E-07 | 0.41304 | 299 | 0.553557828 | 0.007740499 |
| SNP_A.K_1188625_7_6033966 | 7 | 6033966 | 3.78E-07 | 0.4214 | 299 | 0.553468949 | 0.007859162 |
| SNP_A.K_1188703_7_6050264 | 7 | 6050264 | 3.83E-07 | 0.42308 | 299 | 0.553416828 | 0.007897514 |
| SNP_A.K_1188722_7_6056506 | 7 | 6056506 | 3.92E-07 | 0.41806 | 299 | 0.553337281 | 0.007998987 |
| SNP_A.K_1188700_7_6050091 | 7 | 6050091 | 4.02E-07 | 0.42642 | 299 | 0.553251065 | 0.008117967 |
| SNP_A.K_1189196_7_6655864 | 7 | 6655864 | 4.12E-07 | 0.41304 | 299 | 0.553160726 | 0.008249208 |
| SNP_A.K_1188719_7_6053553 | 7 | 6053553 | 4.17E-07 | 0.43144 | 299 | 0.553119973 | 0.008265938 |
| SNP_A.K_1188678_7_6046423 | 7 | 6046423 | 4.61E-07 | 0.42642 | 299 | 0.55276528 | 0.009057883 |
| SNP_A.K_1188621_7_6033401 | 7 | 6033401 | 4.73E-07 | 0.42977 | 299 | 0.552674042 | 0.009209481 |
| SNP_A.K_1188718_7_6053077 | 7 | 6053077 | 4.80E-07 | 0.41137 | 299 | 0.552623464 | 0.009256633 |
| SNP_A.K_1188658_7_6040526 | 7 | 6040526 | 4.86E-07 | 0.42308 | 299 | 0.552584714 | 0.009273531 |
| SNP_A.K_1188708_7_6050841 | 7 | 6050841 | 4.99E-07 | 0.41973 | 299 | 0.552491653 | 0.00943618 |
| SNP_A.K_1188667_7_6042535 | 7 | 6042535 | 5.20E-07 | 0.41806 | 299 | 0.552347323 | 0.009743923 |
| SNP_A.K_1188666_7_6042531 | 7 | 6042531 | 5.30E-07 | 0.41639 | 299 | 0.552280023 | 0.009844026 |
| SNP_A.K_1188717_7_6052907 | 7 | 6052907 | 5.44E-07 | 0.41639 | 299 | 0.552184637 | 0.010025979 |
| SNP_A.K_1188707_7_6050767 | 7 | 6050767 | 5.52E-07 | 0.42475 | 299 | 0.552135884 | 0.010077207 |
| SNP_A.K_1189059_7_6452713 | 7 | 6452713 | 5.73E-07 | 0.26756 | 299 | 0.552004629 | 0.010370773 |
| SNP_A.K_1188896_7_6423302 | 7 | 6423302 | 5.82E-07 | 0.2592 | 299 | 0.551950643 | 0.010441001 |

| | | | | | | | |
|---|---|---|---|---|---|---|---|
| SNP_A.K_1188686_7_6046821 | 7 | 6046821 | 6.02E-07 | 0.42642 | 299 | 0.551833798 | 0.010702851 |
| SNP_A.K_1188691_7_6048797 | 7 | 6048797 | 7.20E-07 | 0.41806 | 299 | 0.551206826 | 0.012601011 |
| SNP_A.K_1188653_7_6035758 | 7 | 6035758 | 7.20E-07 | 0.41973 | 299 | 0.551203078 | 0.012601011 |
| SNP_A.K_1188612_7_6032562 | 7 | 6032562 | 7.41E-07 | 0.41806 | 299 | 0.551105254 | 0.012850558 |
| SNP_A.K_1188628_7_6034007 | 7 | 6034007 | 7.53E-07 | 0.42475 | 299 | 0.551050224 | 0.012946565 |
| SNP_A.K_1188847_7_6377569 | 7 | 6377569 | 7.88E-07 | 0.49331 | 299 | 0.550887757 | 0.013451483 |
| SNP_A.K_1189208_7_6657209 | 7 | 6657209 | 8.14E-07 | 0.40468 | 299 | 0.55077569 | 0.013777065 |
| SNP_A.K_1188589_7_6031314 | 7 | 6031314 | 8.24E-07 | 0.4214 | 299 | 0.550735051 | 0.013825894 |
| SNP_A.K_1188643_7_6035099 | 7 | 6035099 | 8.65E-07 | 0.41639 | 299 | 0.550563327 | 0.014274002 |
| SNP_A.K_1188526_7_6025243 | 7 | 6025243 | 8.70E-07 | 0.40301 | 299 | 0.550543977 | 0.014274002 |
| SNP_A.K_1189172_7_6492992 | 7 | 6492992 | 8.71E-07 | 0.25084 | 299 | 0.550540211 | 0.014274002 |
| SNP_A.K_1189232_7_6659714 | 7 | 6659714 | 8.93E-07 | 0.41304 | 299 | 0.550452169 | 0.014360698 |
| SNP_A.K_1188655_7_6036907 | 7 | 6036907 | 8.94E-07 | 0.42308 | 299 | 0.550448264 | 0.014360698 |
| SNP_A.K_1188776_7_6195235 | 7 | 6195235 | 8.97E-07 | 0.29097 | 299 | 0.550437588 | 0.014360698 |
| SNP_A.K_1188421_7_6017745 | 7 | 6017745 | 9.22E-07 | 0.41639 | 299 | 0.550341714 | 0.014648074 |
| SNP_A.K_1189140_7_6482223 | 7 | 6482223 | 9.41E-07 | 0.26421 | 299 | 0.550269148 | 0.014765607 |
| SNP_A.K_1188475_7_6020813 | 7 | 6020813 | 9.43E-07 | 0.41973 | 299 | 0.550260977 | 0.014765607 |
| SNP_A.K_1189160_7_6487025 | 7 | 6487025 | 9.52E-07 | 0.25753 | 299 | 0.550231287 | 0.014780697 |
| SNP_A.K_1978698_12_8876084 | 12 | 8876084 | 9.71E-07 | 0.08194 | 299 | 0.550160487 | 0.014972097 |
| SNP_A.K_1188805_7_6309120 | 7 | 6309120 | 9.83E-07 | 0.21237 | 299 | 0.55011877 | 0.015040879 |
| SNP_A.K_1188407_7_6015296 | 7 | 6015296 | 9.90E-07 | 0.41973 | 299 | 0.550093102 | 0.015041414 |
| SNP_A.K_1188385_7_6009785 | 7 | 6009785 | 1.00E-06 | 0.42475 | 299 | 0.550057292 | 0.015086585 |
| SNP_A.K_1188416_7_6017492 | 7 | 6017492 | 1.02E-06 | 0.41304 | 299 | 0.549974625 | 0.015337504 |
| SNP_A.K_1188420_7_6017734 | 7 | 6017734 | 1.10E-06 | 0.41806 | 299 | 0.549725954 | 0.016354486 |
| SNP_A.K_1188506_7_6023020 | 7 | 6023020 | 1.12E-06 | 0.39632 | 299 | 0.549672074 | 0.016491658 |
| SNP_A.K_1188482_7_6021197 | 7 | 6021197 | 1.14E-06 | 0.40301 | 299 | 0.549614719 | 0.016564457 |
| SNP_A.K_1188639_7_6034903 | 7 | 6034903 | 1.14E-06 | 0.41806 | 299 | 0.549603433 | 0.016564457 |
| SNP_A.K_1188406_7_6015289 | 7 | 6015289 | 1.15E-06 | 0.41639 | 299 | 0.54956284 | 0.016564457 |
| SNP_A.K_1189133_7_6481768 | 7 | 6481768 | 1.16E-06 | 0.2592 | 299 | 0.549554644 | 0.016564457 |
| SNP_A.K_1188721_7_6055990 | 7 | 6055990 | 1.16E-06 | 0.41304 | 299 | 0.549535403 | 0.016564457 |
| SNP_A.K_1188424_7_6017919 | 7 | 6017919 | 1.19E-06 | 0.41639 | 299 | 0.549448321 | 0.016868545 |
| SNP_A.K_1188780_7_6209682 | 7 | 6209682 | 1.20E-06 | 0.37793 | 299 | 0.549420871 | 0.016887235 |
| SNP_A.K_1188714_7_6052833 | 7 | 6052833 | 1.22E-06 | 0.42308 | 299 | 0.549362681 | 0.016925704 |
| SNP_A.K_1188399_7_6014534 | 7 | 6014534 | 1.23E-06 | 0.41137 | 299 | 0.549348291 | 0.016925704 |
| SNP_A.K_1188672_7_6044448 | 7 | 6044448 | 1.23E-06 | 0.41137 | 299 | 0.549343121 | 0.016925704 |
| SNP_A.K_1188422_7_6017905 | 7 | 6017905 | 1.25E-06 | 0.41472 | 299 | 0.549282808 | 0.017108374 |
| SNP_A.K_1188730_7_6061164 | 7 | 6061164 | 1.27E-06 | 0.42308 | 299 | 0.549212991 | 0.01734113 |
| SNP_A.K_1188644_7_6035140 | 7 | 6035140 | 1.30E-06 | 0.41304 | 299 | 0.549131282 | 0.017401551 |
| SNP_A.K_1189113_7_6472957 | 7 | 6472957 | 1.31E-06 | 0.26254 | 299 | 0.549112582 | 0.017401551 |
| SNP_A.K_1188720_7_6055841 | 7 | 6055841 | 1.32E-06 | 0.42809 | 299 | 0.549102098 | 0.017401551 |
| SNP_A.K_1188398_7_6014305 | 7 | 6014305 | 1.32E-06 | 0.41304 | 299 | 0.549092086 | 0.017401551 |
| SNP_A.K_1188595_7_6031556 | 7 | 6031556 | 1.32E-06 | 0.41472 | 299 | 0.54908818 | 0.017401551 |
| SNP_A.K_1188740_7_6062503 | 7 | 6062503 | 1.33E-06 | 0.39632 | 299 | 0.549059179 | 0.017401551 |
| SNP_A.K_1188423_7_6017907 | 7 | 6017907 | 1.34E-06 | 0.41806 | 299 | 0.549045328 | 0.017401551 |
| SNP_A.K_1188701_7_6050144 | 7 | 6050144 | 1.35E-06 | 0.41973 | 299 | 0.549020997 | 0.01741492 |
| SNP_A.K_1188592_7_6031410 | 7 | 6031410 | 1.39E-06 | 0.33445 | 299 | 0.548913604 | 0.017850501 |
| SNP_A.K_1188552_7_6026786 | 7 | 6026786 | 1.41E-06 | 0.4097 | 299 | 0.548865312 | 0.017989246 |
| SNP_A.K_1189242_7_6660594 | 7 | 6660594 | 1.45E-06 | 0.44816 | 299 | 0.548770791 | 0.018277611 |
| SNP_A.K_1188382_7_6009617 | 7 | 6009617 | 1.46E-06 | 0.41472 | 299 | 0.54875189 | 0.018277611 |
| SNP_A.K_1188741_7_6062705 | 7 | 6062705 | 1.46E-06 | 0.4214 | 299 | 0.548746695 | 0.018277611 |

| | | | | | | | |
|---|---|---|---|---|---|---|---|
| SNP_A.K_1188683_7_6046692 | 7 | 6046692 | 1.50E-06 | 0.42642 | 299 | 0.548640039 | 0.018734714 |
| SNP_A.K_1188597_7_6031761 | 7 | 6031761 | 1.55E-06 | 0.40301 | 299 | 0.548537256 | 0.019182697 |
| SNP_A.K_1188411_7_6016282 | 7 | 6016282 | 1.57E-06 | 0.41639 | 299 | 0.548489567 | 0.019332946 |
| SNP_A.K_1978596_12_8865436 | 12 | 8865436 | 1.60E-06 | 0.08696 | 299 | 0.548428179 | 0.019472549 |
| SNP_A.K_1188955_7_6436503 | 7 | 6436503 | 1.60E-06 | 0.25753 | 299 | 0.548423756 | 0.019472549 |
| SNP_A.K_1188415_7_6017211 | 7 | 6017211 | 1.62E-06 | 0.41472 | 299 | 0.548383585 | 0.019565596 |
| SNP_A.K_1980465_12_9078697 | 12 | 9078697 | 1.63E-06 | 0.12542 | 299 | 0.548353855 | 0.019565596 |
| SNP_A.K_1188954_7_6436498 | 7 | 6436498 | 1.65E-06 | 0.25418 | 299 | 0.548314384 | 0.019565596 |
| SNP_A.K_1978716_12_8877270 | 12 | 8877270 | 1.66E-06 | 0.08194 | 299 | 0.548297996 | 0.019565596 |
| SNP_A.K_1188419_7_6017702 | 7 | 6017702 | 1.67E-06 | 0.41472 | 299 | 0.548281418 | 0.019565596 |
| SNP_A.K_1188430_7_6018298 | 7 | 6018298 | 1.67E-06 | 0.41304 | 299 | 0.548281394 | 0.019565596 |
| SNP_A.K_1188671_7_6044440 | 7 | 6044440 | 1.67E-06 | 0.41137 | 299 | 0.548267999 | 0.019565596 |
| SNP_A.K_1189124_7_6474124 | 7 | 6474124 | 1.69E-06 | 0.26756 | 299 | 0.548236946 | 0.019631229 |
| SNP_A.K_1188674_7_6044536 | 7 | 6044536 | 1.74E-06 | 0.40134 | 299 | 0.548138585 | 0.020007921 |
| SNP_A.K_1188642_7_6035030 | 7 | 6035030 | 1.74E-06 | 0.4097 | 299 | 0.548132484 | 0.020007921 |
| SNP_A.K_1188649_7_6035275 | 7 | 6035275 | 1.78E-06 | 0.42308 | 299 | 0.548046789 | 0.020331619 |
| SNP_A.K_1189104_7_6470836 | 7 | 6470836 | 1.79E-06 | 0.25585 | 299 | 0.548038729 | 0.020331619 |
| SNP_A.K_1188953_7_6436491 | 7 | 6436491 | 1.80E-06 | 0.24582 | 299 | 0.548010697 | 0.020385285 |
| SNP_A.K_1188956_7_6436513 | 7 | 6436513 | 1.84E-06 | 0.25753 | 299 | 0.5479477 | 0.020646928 |
| SNP_A.K_1188693_7_6049406 | 7 | 6049406 | 1.90E-06 | 0.43645 | 299 | 0.547832269 | 0.021125943 |
| SNP_A.K_1188431_7_6018616 | 7 | 6018616 | 1.90E-06 | 0.37625 | 299 | 0.54782106 | 0.021125943 |
| SNP_A.K_1188651_7_6035509 | 7 | 6035509 | 1.91E-06 | 0.43144 | 299 | 0.54781247 | 0.021125943 |
| SNP_A.K_1189116_7_6473186 | 7 | 6473186 | 1.96E-06 | 0.24582 | 299 | 0.547717172 | 0.021489987 |
| SNP_A.K_1188465_7_6020475 | 7 | 6020475 | 1.98E-06 | 0.41304 | 299 | 0.547688839 | 0.021489987 |
| SNP_A.K_1188396_7_6013857 | 7 | 6013857 | 1.99E-06 | 0.40468 | 299 | 0.547669453 | 0.021489987 |
| SNP_A.K_1189138_7_6482154 | 7 | 6482154 | 1.99E-06 | 0.25753 | 299 | 0.54766498 | 0.021489987 |
| SNP_A.K_1189182_7_6493618 | 7 | 6493618 | 1.99E-06 | 0.24749 | 299 | 0.547662357 | 0.021489987 |
| SNP_A.K_1189018_7_6441680 | 7 | 6441680 | 2.03E-06 | 0.26254 | 299 | 0.547597959 | 0.021780489 |
| SNP_A.K_1188634_7_6034804 | 7 | 6034804 | 2.04E-06 | 0.39799 | 299 | 0.547579024 | 0.021787608 |
| SNP_A.K_1188669_7_6044135 | 7 | 6044135 | 2.06E-06 | 0.42475 | 299 | 0.54754456 | 0.021893285 |
| SNP_A.K_1188560_7_6027879 | 7 | 6027879 | 2.08E-06 | 0.37793 | 299 | 0.547511221 | 0.021992918 |
| SNP_A.K_1188729_7_6060779 | 7 | 6060779 | 2.10E-06 | 0.42308 | 299 | 0.547484552 | 0.022051067 |
| SNP_A.K_196290_1_31384954 | 1 | 31384954 | 2.12E-06 | 0.199 | 299 | 0.547445939 | 0.022186373 |
| SNP_A.K_1188673_7_6044479 | 7 | 6044479 | 2.17E-06 | 0.38462 | 299 | 0.547368191 | 0.022576958 |
| SNP_A.K_1188893_7_6422590 | 7 | 6422590 | 2.20E-06 | 0.24749 | 299 | 0.547323002 | 0.022709528 |
| SNP_A.K_1921982_11_24257745 | 11 | 24257745 | 2.20E-06 | 0.1689 | 299 | 0.547313501 | 0.022709528 |
| SNP_A.K_1188631_7_6034489 | 7 | 6034489 | 2.23E-06 | 0.42642 | 299 | 0.54727031 | 0.022881529 |
| SNP_A.K_1188725_7_6058314 | 7 | 6058314 | 2.29E-06 | 0.43144 | 299 | 0.547180386 | 0.023369032 |
| SNP_A.K_1188583_7_6030616 | 7 | 6030616 | 2.39E-06 | 0.42642 | 299 | 0.547026298 | 0.02431471 |
| SNP_A.K_1188414_7_6017158 | 7 | 6017158 | 2.41E-06 | 0.4214 | 299 | 0.546997731 | 0.024397452 |
| SNP_A.K_1188486_7_6021294 | 7 | 6021294 | 2.43E-06 | 0.38127 | 299 | 0.546970631 | 0.024470676 |
| SNP_A.K_1188594_7_6031437 | 7 | 6031437 | 2.45E-06 | 0.42475 | 299 | 0.546952183 | 0.024483352 |
| SNP_A.K_1189233_7_6659795 | 7 | 6659795 | 2.53E-06 | 0.41973 | 299 | 0.546841181 | 0.025161527 |
| SNP_A.K_1188650_7_6035315 | 7 | 6035315 | 2.56E-06 | 0.41973 | 299 | 0.546795912 | 0.025371956 |
| SNP_A.K_1189054_7_6450095 | 7 | 6450095 | 2.71E-06 | 0.2709 | 299 | 0.546602636 | 0.02661312 |
| SNP_A.K_1188713_7_6051639 | 7 | 6051639 | 2.71E-06 | 0.4097 | 299 | 0.546598208 | 0.02661312 |
| SNP_A.K_1188559_7_6027876 | 7 | 6027876 | 2.73E-06 | 0.37625 | 299 | 0.546576213 | 0.026657446 |
| SNP_A.K_1188390_7_6012820 | 7 | 6012820 | 2.77E-06 | 0.38294 | 299 | 0.54652187 | 0.026869571 |
| SNP_A.K_1189241_7_6660558 | 7 | 6660558 | 2.79E-06 | 0.38294 | 299 | 0.546500579 | 0.026869571 |
| SNP_A.K_1189158_7_6486632 | 7 | 6486632 | 2.81E-06 | 0.26087 | 299 | 0.546473744 | 0.026869571 |

| | | | | | | | |
|---|---|---|---|---|---|---|---|
| SNP_A.K_1189019_7_6441806 | 7 | 6441806 | 2.81E-06 | 0.26254 | 299 | 0.54647082 | 0.026869571 |
| SNP_A.K_1188438_7_6019232 | 7 | 6019232 | 2.83E-06 | 0.41639 | 299 | 0.546453817 | 0.026869571 |
| SNP_A.K_1188426_7_6018103 | 7 | 6018103 | 2.83E-06 | 0.41472 | 299 | 0.546452949 | 0.026869571 |
| SNP_A.K_1188381_7_6009561 | 7 | 6009561 | 2.92E-06 | 0.41639 | 299 | 0.546337869 | 0.027654401 |
| SNP_A.K_1188882_7_6421132 | 7 | 6421132 | 3.02E-06 | 0.25753 | 299 | 0.546228702 | 0.028414305 |
| SNP_A.K_1188400_7_6014616 | 7 | 6014616 | 3.07E-06 | 0.42308 | 299 | 0.546171088 | 0.028762775 |
| SNP_A.K_1188474_7_6020808 | 7 | 6020808 | 3.09E-06 | 0.41639 | 299 | 0.546151034 | 0.02880078 |
| SNP_A.K_1188929_7_6434967 | 7 | 6434967 | 3.13E-06 | 0.2592 | 299 | 0.546099132 | 0.029107018 |
| SNP_A.K_1188410_7_6016255 | 7 | 6016255 | 3.18E-06 | 0.42308 | 299 | 0.546048105 | 0.0294097 |
| SNP_A.K_1922047_11_24262683 | 11 | 24262683 | 3.20E-06 | 0.2107 | 299 | 0.546021338 | 0.029476246 |
| SNP_A.K_1188575_7_6030070 | 7 | 6030070 | 3.21E-06 | 0.41304 | 299 | 0.546009804 | 0.029476246 |
| SNP_A.K_1189060_7_6452776 | 7 | 6452776 | 3.24E-06 | 0.26756 | 299 | 0.54598267 | 0.029578884 |
| SNP_A.K_1189058_7_6452614 | 7 | 6452614 | 3.28E-06 | 0.39298 | 299 | 0.545939004 | 0.029680888 |
| SNP_A.K_1188945_7_6436025 | 7 | 6436025 | 3.30E-06 | 0.25251 | 299 | 0.545921743 | 0.029680888 |
| SNP_A.K_1188957_7_6436531 | 7 | 6436531 | 3.31E-06 | 0.25585 | 299 | 0.545909694 | 0.029680888 |
| SNP_A.K_1188413_7_6016671 | 7 | 6016671 | 3.32E-06 | 0.41472 | 299 | 0.545897355 | 0.029680888 |
| SNP_A.K_1188958_7_6436543 | 7 | 6436543 | 3.33E-06 | 0.26087 | 299 | 0.545887767 | 0.029680888 |
| SNP_A.K_1189143_7_6482578 | 7 | 6482578 | 3.35E-06 | 0.25585 | 299 | 0.54587053 | 0.029680888 |
| SNP_A.K_1188525_7_6025219 | 7 | 6025219 | 3.35E-06 | 0.40635 | 299 | 0.545866586 | 0.029680888 |
| SNP_A.K_1188488_7_6021310 | 7 | 6021310 | 3.40E-06 | 0.32943 | 299 | 0.545815332 | 0.029885843 |
| SNP_A.K_1976796_12_8034717 | 12 | 8034717 | 3.40E-06 | 0.11706 | 299 | 0.545813675 | 0.029885843 |
| SNP_A.K_1189005_7_6440863 | 7 | 6440863 | 3.53E-06 | 0.2592 | 299 | 0.545684462 | 0.030897815 |
| SNP_A.K_1188549_7_6026617 | 7 | 6026617 | 3.56E-06 | 0.40635 | 299 | 0.545657964 | 0.031006186 |
| SNP_A.K_1188670_7_6044425 | 7 | 6044425 | 3.59E-06 | 0.41137 | 299 | 0.545629309 | 0.031134999 |
| SNP_A.K_1188807_7_6310485 | 7 | 6310485 | 3.62E-06 | 0.38127 | 299 | 0.54560299 | 0.031243707 |
| SNP_A.K_1188527_7_6025253 | 7 | 6025253 | 3.71E-06 | 0.39799 | 299 | 0.545516552 | 0.031853009 |
| SNP_A.K_1188952_7_6436489 | 7 | 6436489 | 3.72E-06 | 0.24749 | 299 | 0.545508045 | 0.031853009 |
| SNP_A.K_1188613_7_6032582 | 7 | 6032582 | 3.82E-06 | 0.4214 | 299 | 0.545415138 | 0.032590437 |
| SNP_A.K_1188473_7_6020754 | 7 | 6020754 | 3.92E-06 | 0.41639 | 299 | 0.545322877 | 0.033339474 |
| SNP_A.K_1188534_7_6025564 | 7 | 6025564 | 3.96E-06 | 0.40468 | 299 | 0.545287798 | 0.033544248 |
| SNP_A.K_1189110_7_6472614 | 7 | 6472614 | 4.05E-06 | 0.2592 | 299 | 0.545214314 | 0.034129836 |
| SNP_A.K_1188648_7_6035256 | 7 | 6035256 | 4.17E-06 | 0.4214 | 299 | 0.545115398 | 0.034984177 |
| SNP_A.K_1188427_7_6018134 | 7 | 6018134 | 4.39E-06 | 0.41639 | 299 | 0.544938786 | 0.036592271 |
| SNP_A.K_1188681_7_6046537 | 7 | 6046537 | 4.39E-06 | 0.44816 | 299 | 0.544933307 | 0.036592271 |
| SNP_A.K_1188533_7_6025559 | 7 | 6025559 | 4.42E-06 | 0.40468 | 299 | 0.544912913 | 0.036661869 |
| SNP_A.K_1980397_12_9074823 | 12 | 9074823 | 4.44E-06 | 0.09365 | 299 | 0.544899396 | 0.036661869 |
| SNP_A.K_1976362_12_7992926 | 12 | 7992926 | 4.49E-06 | 0.06689 | 299 | 0.544856659 | 0.036974002 |
| SNP_A.K_1188409_7_6015931 | 7 | 6015931 | 4.60E-06 | 0.41639 | 299 | 0.544775969 | 0.037703707 |
| SNP_A.K_1188432_7_6018663 | 7 | 6018663 | 4.84E-06 | 0.4214 | 299 | 0.544603448 | 0.039490786 |
| SNP_A.K_1188601_7_6031939 | 7 | 6031939 | 4.89E-06 | 0.41639 | 299 | 0.544569492 | 0.039727541 |
| SNP_A.K_1188664_7_6041779 | 7 | 6041779 | 4.94E-06 | 0.42308 | 299 | 0.54452834 | 0.040050229 |
| SNP_A.K_1188627_7_6034001 | 7 | 6034001 | 4.98E-06 | 0.42977 | 299 | 0.54450591 | 0.040156567 |
| SNP_A.K_1188927_7_6434767 | 7 | 6434767 | 5.09E-06 | 0.27425 | 299 | 0.544429907 | 0.040870909 |
| SNP_A.K_1188457_7_6020338 | 7 | 6020338 | 5.12E-06 | 0.40134 | 299 | 0.544411762 | 0.040870909 |
| SNP_A.K_1189184_7_6493946 | 7 | 6493946 | 5.12E-06 | 0.2592 | 299 | 0.544405728 | 0.040870909 |
| SNP_A.K_1188819_7_6356872 | 7 | 6356872 | 5.16E-06 | 0.36957 | 299 | 0.544379697 | 0.040941341 |
| SNP_A.K_1188970_7_6437983 | 7 | 6437983 | 5.17E-06 | 0.26421 | 299 | 0.544373629 | 0.040941341 |
| SNP_A.K_1978694_12_8875824 | 12 | 8875824 | 5.29E-06 | 0.08361 | 299 | 0.544299935 | 0.041444788 |
| SNP_A.K_1188428_7_6018250 | 7 | 6018250 | 5.29E-06 | 0.41806 | 299 | 0.544297449 | 0.041444788 |
| SNP_A.K_1976632_12_8020430 | 12 | 8020430 | 5.32E-06 | 0.0903 | 299 | 0.544279995 | 0.041444788 |

| | | | | | | | |
|---|---|---|---|---|---|---|---|
| SNP_A.K_1188731_7_6061934 | 7 | 6061934 | 5.32E-06 | 0.4214 | 299 | 0.544275698 | 0.041444788 |
| SNP_A.K_1188966_7_6436877 | 7 | 6436877 | 5.34E-06 | 0.25418 | 299 | 0.544267102 | 0.041444788 |
| SNP_A.K_1980333_12_9070375 | 12 | 9070375 | 5.36E-06 | 0.08696 | 299 | 0.544252245 | 0.04146997 |
| SNP_A.K_1188694_7_6049467 | 7 | 6049467 | 5.46E-06 | 0.4398 | 299 | 0.544191082 | 0.042000912 |
| SNP_A.K_1189179_7_6493365 | 7 | 6493365 | 5.50E-06 | 0.25084 | 299 | 0.54416564 | 0.042000912 |
| SNP_A.K_1188462_7_6020459 | 7 | 6020459 | 5.50E-06 | 0.41137 | 299 | 0.544164867 | 0.042000912 |
| SNP_A.K_1188397_7_6014141 | 7 | 6014141 | 5.52E-06 | 0.40635 | 299 | 0.544151699 | 0.042000912 |
| SNP_A.K_1188402_7_6014760 | 7 | 6014760 | 5.53E-06 | 0.41806 | 299 | 0.544145469 | 0.042000912 |
| SNP_A.K_1188637_7_6034881 | 7 | 6034881 | 5.59E-06 | 0.4097 | 299 | 0.544108156 | 0.042306127 |
| SNP_A.K_1980436_12_9077045 | 12 | 9077045 | 5.65E-06 | 0.08027 | 299 | 0.544070703 | 0.042615921 |
| SNP_A.K_1188412_7_6016537 | 7 | 6016537 | 5.67E-06 | 0.41639 | 299 | 0.54405732 | 0.042628189 |
| SNP_A.K_1188524_7_6025199 | 7 | 6025199 | 5.73E-06 | 0.41304 | 299 | 0.544023405 | 0.042897223 |
| SNP_A.K_1188379_7_6009473 | 7 | 6009473 | 5.76E-06 | 0.41973 | 299 | 0.544005539 | 0.042966878 |
| SNP_A.K_1188596_7_6031719 | 7 | 6031719 | 5.84E-06 | 0.42308 | 299 | 0.543957757 | 0.043414573 |
| SNP_A.K_1189129_7_6479712 | 7 | 6479712 | 5.87E-06 | 0.20903 | 299 | 0.543940733 | 0.043475537 |
| SNP_A.K_1188600_7_6031936 | 7 | 6031936 | 5.94E-06 | 0.41304 | 299 | 0.543898446 | 0.043859359 |
| SNP_A.K_1188500_7_6022211 | 7 | 6022211 | 6.14E-06 | 0.38127 | 299 | 0.543787827 | 0.04509114 |
| SNP_A.K_1188680_7_6046495 | 7 | 6046495 | 6.15E-06 | 0.4214 | 299 | 0.543779363 | 0.04509114 |
| SNP_A.K_1188481_7_6021119 | 7 | 6021119 | 6.19E-06 | 0.41137 | 299 | 0.543756547 | 0.045142055 |
| SNP_A.K_254578_1_41048790 | 1 | 41048790 | 6.20E-06 | 0.41472 | 299 | 0.543751466 | 0.045142055 |
| SNP_A.K_1980241_12_9067475 | 12 | 9067475 | 6.24E-06 | 0.0903 | 299 | 0.543733421 | 0.045222432 |
| SNP_A.K_1188453_7_6020005 | 7 | 6020005 | 6.39E-06 | 0.4097 | 299 | 0.543649489 | 0.046183787 |
| SNP_A.K_1980196_12_9065692 | 12 | 9065692 | 6.56E-06 | 0.09532 | 299 | 0.543560862 | 0.047008616 |
| SNP_A.K_925899_5_19753862 | 5 | 19753862 | 6.57E-06 | 0.44314 | 299 | 0.543556719 | 0.047008616 |
| SNP_A.K_1188417_7_6017576 | 7 | 6017576 | 6.58E-06 | 0.41806 | 299 | 0.543546768 | 0.047008616 |
| SNP_A.K_1188542_7_6025907 | 7 | 6025907 | 6.59E-06 | 0.40635 | 299 | 0.543541683 | 0.047008616 |
| SNP_A.K_1979774_12_8963385 | 12 | 8963385 | 6.63E-06 | 0.09197 | 299 | 0.543523849 | 0.047092957 |
| SNP_A.K_1188946_7_6436307 | 7 | 6436307 | 6.75E-06 | 0.26923 | 299 | 0.543459605 | 0.047718853 |
| SNP_A.K_1188458_7_6020348 | 7 | 6020348 | 6.76E-06 | 0.40635 | 299 | 0.543455393 | 0.047718853 |
| SNP_A.K_1188630_7_6034470 | 7 | 6034470 | 6.81E-06 | 0.41973 | 299 | 0.543432297 | 0.047879769 |
| SNP_A.K_1922066_11_24263921 | 11 | 24263921 | 6.88E-06 | 0.20569 | 299 | 0.543395011 | 0.048127595 |
| SNP_A.K_1189038_7_6443328 | 7 | 6443328 | 6.89E-06 | 0.2709 | 299 | 0.543391598 | 0.048127595 |
| SNP_A.K_1189123_7_6474113 | 7 | 6474113 | 6.92E-06 | 0.26923 | 299 | 0.543374843 | 0.048202165 |
| SNP_A.K_1189094_7_6467839 | 7 | 6467839 | 6.97E-06 | 0.25585 | 299 | 0.54335123 | 0.048374283 |
| SNP_A.K_1188632_7_6034699 | 7 | 6034699 | 7.06E-06 | 0.41137 | 299 | 0.543307262 | 0.04883739 |
| SNP_A.K_1188991_7_6439209 | 7 | 6439209 | 7.18E-06 | 0.26254 | 299 | 0.543249809 | 0.049363331 |
| SNP_A.K_1189115_7_6473151 | 7 | 6473151 | 7.19E-06 | 0.25753 | 299 | 0.543247985 | 0.049363331 |
| SNP_A.K_1188418_7_6017581 | 7 | 6017581 | 7.21E-06 | 0.42475 | 299 | 0.543235091 | 0.049386844 |
| SNP_A.K_1188606_7_6032141 | 7 | 6032141 | 7.26E-06 | 0.41973 | 299 | 0.543212195 | 0.049442333 |
| SNP_A.K_1189142_7_6482564 | 7 | 6482564 | 7.27E-06 | 0.2592 | 299 | 0.543208832 | 0.049442333 |
